# Population genomics of the island thrush elucidates one of earth’s great archipelagic radiations

**DOI:** 10.1101/2022.04.21.488757

**Authors:** Andrew Hart Reeve, Graham Gower, José Martín Pujolar, Brian Tilston Smith, Bent Petersen, Urban Olsson, Tri Haryoko, Bonny Koane, Gibson Maiah, Mozes P. K. Blom, Per G. P. Ericson, Martin Irestedt, Fernando Racimo, Knud Andreas Jønsson

**Affiliations:** Natural History Museum of Denmark, University of Copenhagen, DK-2100 Copenhagen Ø, Denmark; Lundbeck GeoGenetics Centre, The GLOBE Institute, University of Copenhagen, DK-1350 Copenhagen K, Denmark; Centre for Ocean Life, DTU Aqua, Kemitorvet, Building 202, DK-2800 Kgs. Lyngby, Denmark; Department of Ornithology, American Museum of Natural History, Central Park West at 79^th^ Street, New York, NY 10024, United States; Center for Evolutionary Hologenomics, The GLOBE Institute, University of Copenhagen, DK-1353 Copenhagen K, Denmark; Centre of Excellence for Omics-Driven Computational Biodiscovery, Faculty of Applied Sciences, AIMST University, Kedah, Malaysia; Gothenburg Global Biodiversity Centre, Box 461, SE-405 30 Gothenburg, Sweden; Department of Biological and Environmental Sciences, University of Gothenburg, Box 463, SE-405 30 Gothenburg, Sweden; Museum Zoologicum Bogoriense Research Centre for Biology, National Research and Innovation Agency (BRIN), Jl. Raya Jakarta-Bogor Km 46, Cibinong 16911, Indonesia; New Guinea Binatang Research Centre, Madang, Papua New Guinea; Museum für Naturkunde Berlin, Leibniz Institut für Evolutions- und Biodiversitätsforschung, 10115 Berlin, Germany; Department of Bioinformatics and Genetics, Swedish Museum of Natural History, P.O. Box 50007, SE-104 05 Stockholm, Sweden

## Abstract

Tropical islands are renowned as natural laboratories for evolutionary study. Lineage radiations across tropical archipelagos are ideal systems for investigating how colonization, speciation, and extinction processes shape biodiversity patterns. The expansion of the island thrush across the Indo-Pacific represents one of the largest yet most perplexing island radiations of any songbird species. The island thrush exhibits a complex mosaic of pronounced plumage variation across its range, and is arguably the world’s most polytypic bird. It is a sedentary species largely restricted to mountain forests, yet it has colonized a vast island region spanning a quarter of the globe. We conducted comprehensive sampling of island thrush populations and obtained genome-wide SNP data, which we used to reconstruct its phylogeny, population structure, gene flow, and demographic history. The island thrush evolved from migratory Palearctic ancestors and radiated explosively across the Indo-Pacific during the Pleistocene, with numerous instances of gene flow between populations. Its bewildering plumage variation masks a biogeographically intuitive stepping stone colonization path from the Philippines through the Greater Sundas, Wallacea and New Guinea to Polynesia. The island thrush’s success in colonizing Indo-Pacific mountains can be understood in light of its ancestral mobility and adaptation to cool climates; however, shifts in elevational range, degree of plumage variation and apparent dispersal rates in the eastern part of its range raise further intriguing questions about its biology.

## INTRODUCTION

Tropical archipelagos are natural laboratories that have shaped scientific understanding of evolution and biogeography (Darwin 1859; Wallace 1869; Mayr 1942; MacArthur and Wilson 1967; Ricklefs and Cox 1978; Mayr and Diamond 2001). The processes of colonization, speciation, and extinction are manifested in the modern distribution of their biotas, from evolutionary relics stranded on single islands, to ultra-mobile colonizers ubiquitous across entire archipelagos. At the intersection of these extremes are the ‘great speciators’ (Diamond et al. 1976). These species (or lineages) are sufficiently dispersive to broadly colonize island systems, but paradoxically show distinct differentiation between island populations, indicating incipient speciation (and limited dispersal ability). This dynamic makes great speciators an alluring model for investigating how lineage expansion and diversification shape global biodiversity patterns (Moyle et al. 2009; Jønsson et al. 2014; Pepke et al. 2019). Molecular phylogenetic studies (Moyle et al. 2009; Irestedt et al. 2013; Jønsson et al. 2014; Andersen et al. 2013, 2014, 2015; Pedersen et al. 2018; Kearns et al. 2020) have confirmed that great speciators represent rapid and geographically complex lineage radiations. However, those same attributes, combined with limited genetic sampling, have impeded precise evolutionary reconstruction of these radiations (though see Gwee et al. [2020] and Manthey et al. [2020]).

Another similar group overlaps with the great speciators: the montane species and lineages that have undergone expansive radiations across archipelagic highlands. This group represents a striking component of island species diversity in the Indo-Pacific that has held longstanding interest for researchers studying the formation of montane biodiversity (Rensch et al. 1930; Stresemann 1939; Mayr 1944; Mayr and Diamond 1976, 2001). The rapid mountain colonizations inferred for these species seem doubly improbable because dispersers must overcome both terrestrial lowland and water barriers. The group is therefore central to the question of how past climatic oscillations contributed to modern species distribution patterns via land bridge formation and elevational habitat shifts (Rensch et al. 1930; Stresemann 1939; Mayr 1944; Mayr and Diamond 1976, 2001). Despite this, the great montane island radiations have never been subjected to detailed molecular study.

The island thrush (*Turdus poliocephalus*) is both an archetypal great speciator (Mayr and Diamond 2001) and one of the most prolific avian colonizers of island mountains (Clement and Hathaway 2000; Collar 2005). It is a sedentary species restricted to high montane forest across much of its range, yet it has radiated across islands spanning a 10,000 km distance from Sumatra to Samoa (Clement and Hathaway 2000; Collar 2005). Extraordinary differentiation between individual populations belies this evident propensity for inter-island dispersal. With some 50 recognized subspecies, the island thrush is arguably the world’s most polytypic bird (Clements et al. 2019; Gill et al. 2020), and certainly one of the most variably plumaged. Plumage color and pattern variation is both extreme and geographically incoherent, with similar color patterns often shared by widely separated populations (Peterson 2007). This variation — in addition to variation in sexual dimorphism, body size, and elevational distribution — have confounded interpretation of the island thrush’s evolution. Preliminary molecular work (Voelker et al. 2007; Jones and Kennedy 2008; Nylander et al. 2008; Batista et al. 2020) has left open the question of whether the island thrush is even monophyletic, or an artificial assemblage of unrelated forms.

For this study, we conducted a comprehensive sampling of island thrush populations, both living and historically extinct, and additionally sampled its hypothesized sister clade (Voelker et al. 2007; Nylander et al. 2008) from East Asia. We obtained genome-wide shotgun sequencing data and used single nucleotide polymorphisms (SNPs) to reconstruct the island thrush’s phylogeny, population structure, gene flow, and demographic history. This approach allows us to reveal, in unprecedented detail, the evolution of a great speciator.

## RESULTS

### Phylogenetic analyses

#### Phylogenetic analysis of SNP data

Both genome-wide phylogenetic trees built using SNP data recover the island thrush as monophyletic (Figs. 1, S1). The topologies recovered by the pairwise distance (Fig. 1) and pairwise F_ST_ (Fig. S1) analyses differ in some details. The pairwise distance tree shows a sequential branching pattern that indicates an origin in the Philippines, an expansion through the Greater Sundas and Wallacea, and further eastward colonization of the Pacific via New Guinea. The pairwise F_ST_ tree is broadly similar, but suggests a more general western origin not necessarily centered in the Philippines. F_ST_ trees reflect differentiation due to genetic drift along different lineages, which is a function of time, lineage-specific variation in population sizes, and patterns of isolation between lineages. Our discussion below is mostly focused on the pairwise distance tree (Fig. 1), as it provides a more direct estimate of phylogenetic distances, i.e., without confounding by genetic drift. Additionally, cross-population heterozygosity levels (Results: Population structure and heterozygosity levels) and demographic reconstructions (Results: Demographic history inference using PSMC) both support the hypothesis that the island thrush expanded out of the Philippines. The pairwise distance tree is highly resolved, and the few nodes with < 100% bootstrap support appear to represent recent divergences between geographically proximate populations.

**Figs. 1a, 1b.**
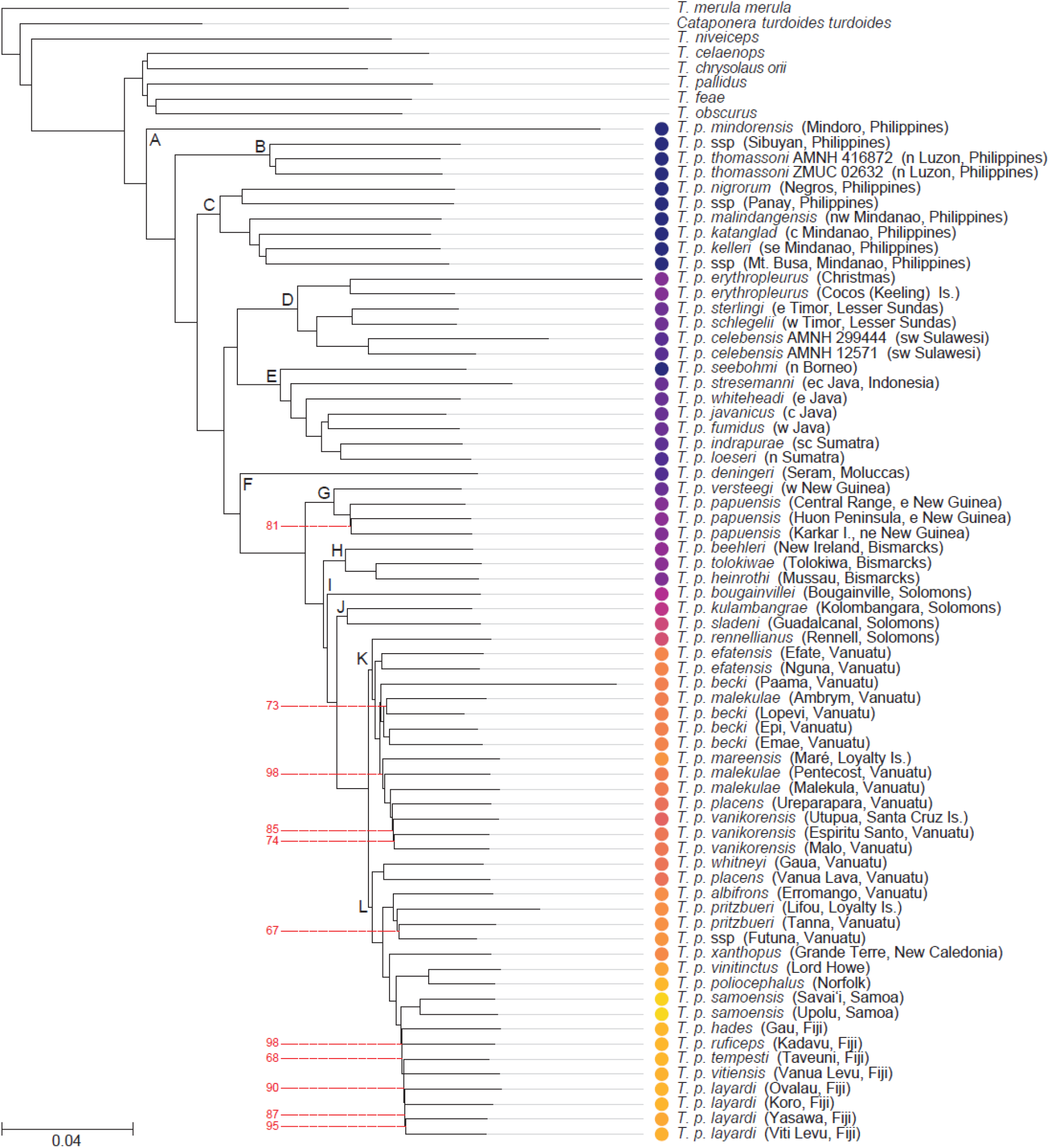
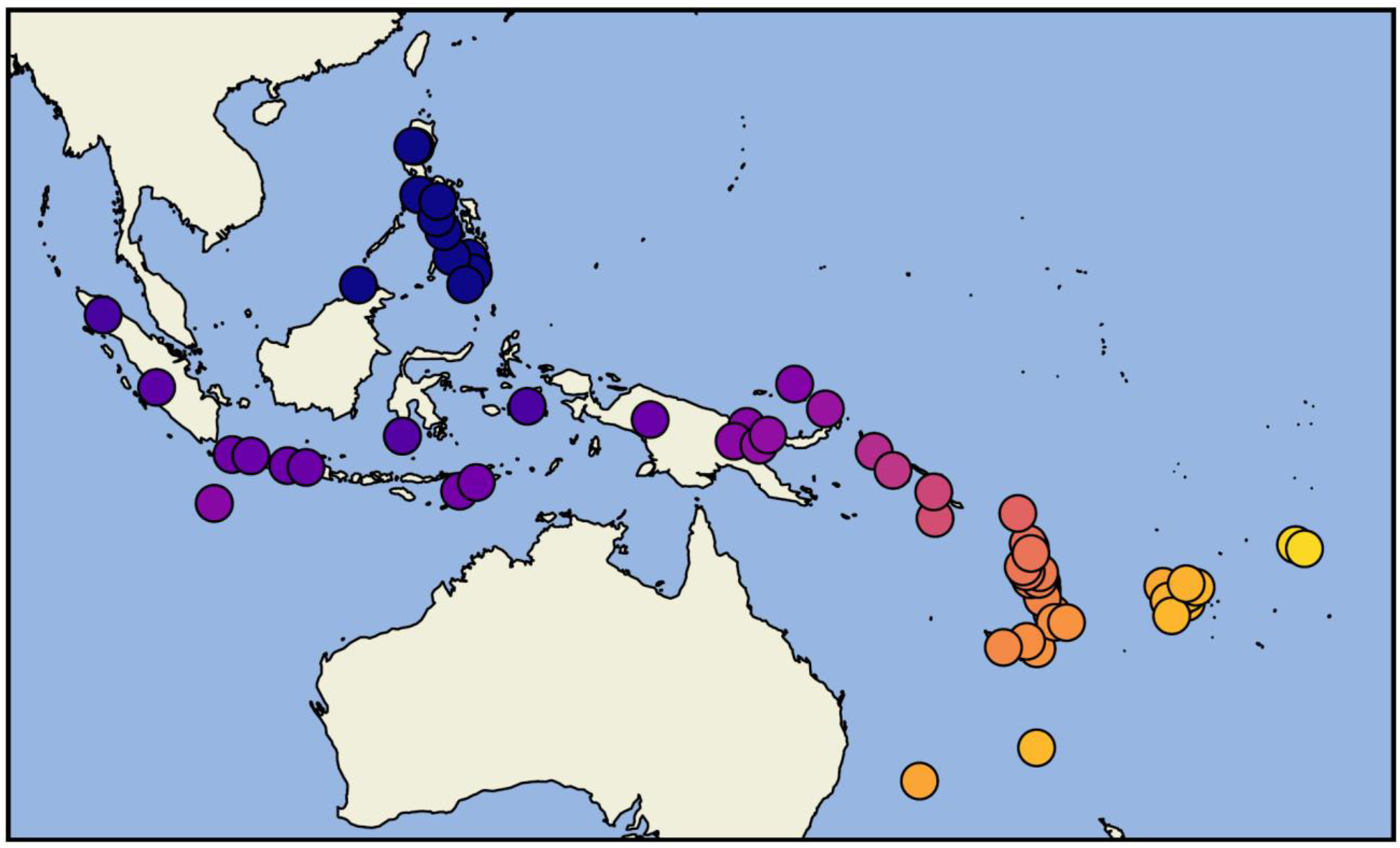
Phylogeny estimated from pairwise distances using neighbor-joining followed by subtree pruning and regrafting (Fig. 1a). Red lines indicate nodes with bootstrap support < 100% (from 100 non-parametric bootstrap replicates). Letters at nodes indicate clades referred to in the text. Leaf node colors on the tree match those used on the map (Fig. 1b). Island thrush individuals are colored by distance to a reference point at 30**°** N, 120**°** E, reflecting a hypothetical distribution of the species’ mainland ancestor.

Five East Asian species constitute the sister clade of the island thrush. This in turn contains two subclades. The first contains *T. chrysolaus* and *T. celaenops*, which together have a breeding range encompassing Japan, Sakhalin, and the Kuril Islands. The second contains *T. pallidus, T. feae*, and *T. obscurus*, which breed mostly on mainland East Asia.

Detailed maps are provided in Figs. S2a–c showing the geographic distribution of island thrush populations and their phylogenetic relationships as recovered by the pairwise distance analysis. Island thrush Clades A, B and C (Fig. 1) are composed of populations from the Philippines. Clade A represents the Mindoro population, *T. p. mindorensis*. Clade B contains *T. p. thomassoni* from northern Luzon and an undescribed population from Sibuyan. Clade C contains 1) a subclade from the central Philippines islands of Negros (*T. p. nigrorum*) and Panay (undescribed); and 2) four populations from disjunct mountain ranges across Mindanao, including an undescribed population from Mt. Busa in the island’s far south.

In Clade D, *T. p. erythropleurus* of Christmas Island in the Indian ocean is sister to a Wallacean group including *T. p. celebensis* from Sulawesi and two sister taxa on Timor. Clade E spans the Greater Sundas islands of Borneo, Java, and Sumatra. The Bornean population (*T. p. seebohmi*) is sister to the rest of the subclade, and Sumatran populations are embedded among Javan populations. The overall pattern indicates a southward spread from Borneo into eastern Java, followed by westward colonization across Java and into Sumatra.

Further eastward colonization into New Guinea appears to have proceeded via Seram in the Moluccas, represented by *T. p. deningeri* (Clade F). The relationships of *T. p. deningeri* with populations further west suggest that the island thrush may have crossed Wallace’s Line twice — either two eastward colonizations, or an eastward colonization followed by a westward back-colonization.

More recently, four clades diverged representing populations from New Guinea, the Bismarck Archipelago, and the Solomon Islands. Clade G contains New Guinea populations; *T. p. versteegi* from the west of the island is sister to more easterly populations, including the small offshore island of Karkar. Clade H contains populations from the Bismarcks. The uncollected population on the relatively large island of New Britain is not included; and we recover an unexpected sister relationship between the widely separated Tolokiwa and Mussau populations. Clade I comprises the Bougainville population, and Clade J contains populations further southeast on Kolombangara and Guadalcanal.

Two large sister clades (Clades K and L) represent broad expansions into the Pacific. Clade K is distributed across southern Melanesia, while Clade L represents an even broader radiation across southern Melanesia, remote Tasman Sea islands, and Samoa. Sister to the rest of Clade K is the population from Rennell in the southern Solomon Islands. The rest of the clade is mostly distributed across Vanuatu, but two populations lie outside Vanuatu’s central islands. The phylogenetic position of the extinct *T. p. mareensis* of Maré in New Caledonia’s Loyalty Islands is unexpected, as other populations from New Caledonia and the southernmost islands of Vanuatu belong to Clade L. The clade also reaches Temotu, north of Vanuatu (*T. p. vanikorensis*). Plumage variation within Clade K is subtle, and three subspecies are not recovered as monophyletic: *T. p. becki, T. p. malekulae*, and *T. p. placens* (the latter including populations both from Clades K and L).

Sister to the rest of Clade L are populations from Gaua and Vanua Lava in the Banks Islands of northern Vanuatu. These populations are oddly interspersed between populations of *T. p. vanikorensis* (Clade K) spanning northern Vanuatu and Temotu. The next clade to diverge represents a distributional leap, encompassing populations in some of the southernmost islands of Vanuatu (Erromango, Tanna, Futuna), as well the extinct population on Lifou in the Loyalty Islands. Our molecular results suggest that the undescribed Futuna population should be assigned to subspecies *pritzbueri*, which otherwise occurs on Tanna (and previously on Lifou). The next branch of the tree represents the extinct Grande Terre (New Caledonia) population of *T. p. xanthopus*. The remaining branches of Clade L represent the most extreme long-distance colonizations that can be inferred for the island thrush. The first branch represents a colonization of the distant Tasman Sea islands of Norfolk (the nominate subspecies) and Lord Howe (*T. p. vinitinctus*). Both taxa are now extinct. The second branch represents colonization of Fiji, where five subspecies form a clade, and the final branch represents colonization of Samoa (subspecies *samoensis*), which marks the eastern limit of the island thrush’s radiation across the Pacific.

### Phylogenetic analysis of mitochondrial genome data

The phylogenetic analysis of mitochondrial genome data (Fig. S3) also recovers the island thrush as monophyletic. The BEAST date estimate for divergence of the the island thrush from its five-species sister clade is 2.4 Mya (95% HPD 2.0–2.8 Mya), and population divergence within the island thrush itself is estimated to have begun 1.3 Mya (95% HPD 1.1–1.5 Mya). The tree shows a sequential branching pattern that is roughly similar to the trees built with nuclear SNP data, again indicating a west-to-east stepping stone colonization pattern. However, while most nodes are strongly supported (posterior probability ≥ 0.99), the topology of the mitogenome tree differs in many details from the nuclear trees, reflecting specific patterns of mitochondrial inheritance that are not recovered from the average autosomal tree. We stress that discordance between the mitrochondrial tree and the nuclear trees is not unexpected, as the nuclear data encompass many distinct gene trees. The mitogenome tree does recover the same Greater Sundas/Wallacea/Christmas Island clade as the nuclear trees, as well as the same clade containing populations from New Guinea and all points east. These clades are both dated to about 0.8 Mya, indicating a very rapid radiation out of the Philippines that quickly reached Melanesia.

### Population structure and heterozygosity levels

The PCAngsd MAP test suggests that six principal dimensions explain the population structure in the dataset, corresponding to 18.63% of the total variance (Fig. S4). Genetic correlation between pairs of individuals, which controls for individual variation in heterozygosity, is visualized in the heatmap in Fig. S5. Ancestry proportions for k=2 to k=8 putative ancestry components are illustrated in Fig. S6. These analyses suggest a genetic structure for the island thrush with strong differentiation between a western clade (Greater Sundas, Philippines, Wallacea) and an eastern clade (New Guinea and islands to the east). The eastern Clades K + L (Fig. 1) are heavily oversampled compared to the group’s many smaller, early branching clades. This has the effect of overrepresenting variation within the eastern clade, while underrepresenting variation between the other clades. To test this interpretation, we reran the latent mixed-membership model analyses on a dataset that included the outgroup taxa. This resulting plot (Fig. S7) shows the outgroup taxa to be homogenous, with a single common ancestry component for k=2 to k=6 ancestors, despite their deep divergences. The ingroup analysis (Fig. S6) suggests mixed ancestry in a number of populations, notably *T. p. mindorensis* (Mindoro, Philippines), *T. p. deningeri* (Seram, Moluccas), *T. p. seebohmi* (Borneo), *T. p. stressemanni* (east-central Java), and many populations in the islands east of New Guinea. While ascertainment bias due to uneven sampling is potentially problematic (Puechmaille 2016; Lawson et al. 2018), explicit tests for gene flow using D-statistics (Results: Gene flow) support multiple gene flow events. Heterozygosity levels of individuals are visualized in Fig. 2; there is a broad pattern of west-to-east decline, and levels tend to be higher in populations from larger islands.

**Fig. 2.**
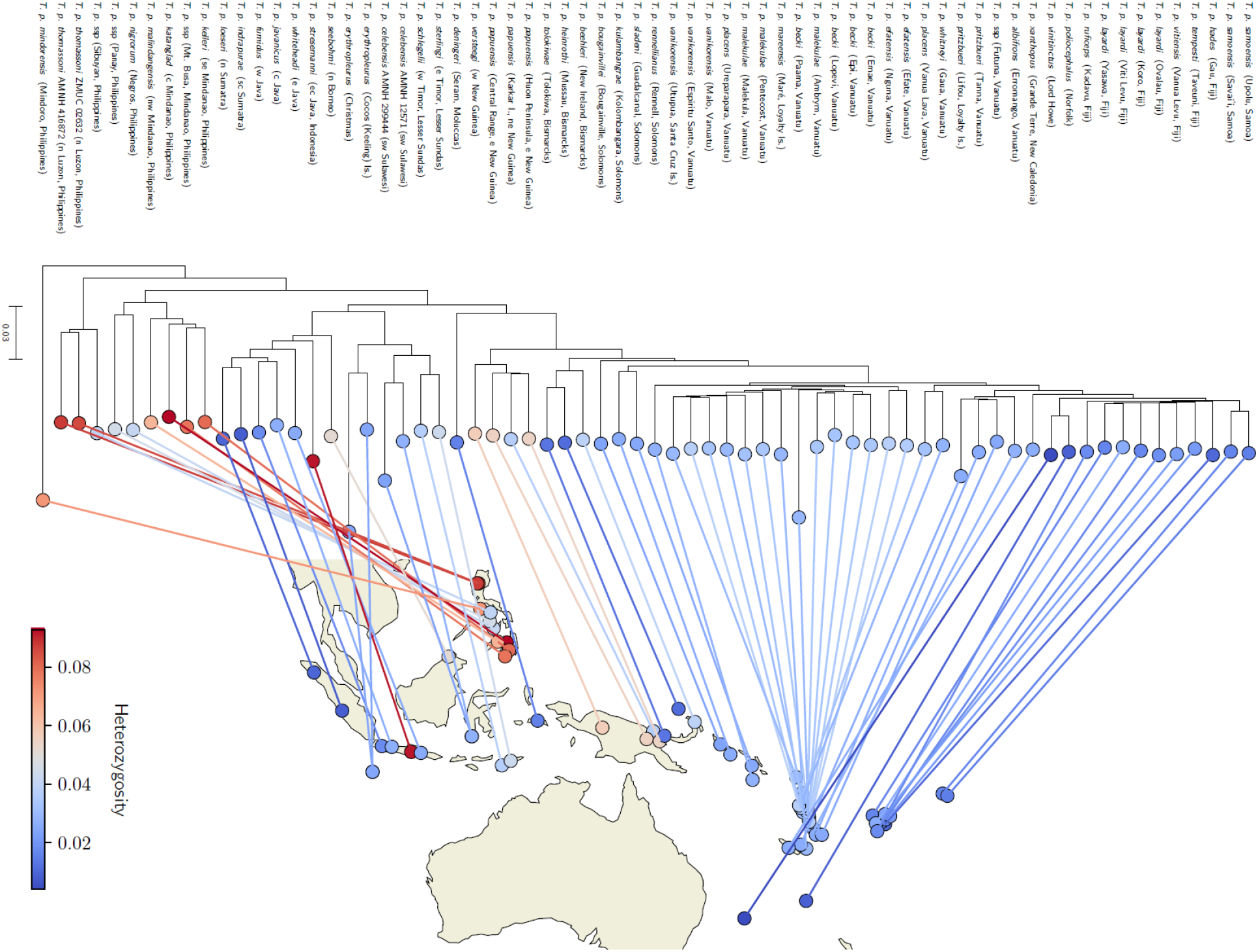
Heterozygosity levels of individuals, overlaid upon the pairwise distance tree. Lines connect leaf nodes to the geographic origin of the individual; crossing of lines has been reduced by sorting the left and right branches of internal nodes by the mean longitude of their respective leaf nodes. The overall pattern suggests a serial founder effect from a radiation that proceeded from the Philippines.

### Gene flow

Of the 67,525 calculated *D*_*b*_*(C)* statistics, 15,055 are significant at FWER < 0.05 (Fig. 3). The results indicate widespread ancient and recent gene flow within the island thrush and its five-species sister clade. Gene flow across early branches of the tree has in many cases left a visual pattern (Fig. 3) of long rows of similarly shaded cells, with the genetic signature of early admixture being inherited by descendent populations. Ancient gene flow is inferred within the island thrush’s sister clade, and also between members of this sister clade and the ancestral island thrush lineage. This is particularly evident in e.g. *T. pallidus*, and in the last common ancestor (LCA) of *T. celaenops* and *T. chrysolaus*. Ancient gene flow is also inferred within the island thrush itself. We detected substantial admixture between the ancestral lineages that gave rise to populations in 1) the Greater Sundas, 2) Wallacea, and 3) islands from the Moluccas east to Polynesia. Admixture is also widespread among the deeper ancestral nodes of the clades representing populations east of New Guinea (Clades H–L in Fig. 1). The results further indicate many instances of more recent gene flow between island thrush populations. Recent gene flow is much more prevalent among populations east of New Guinea, with gene flow inferred between several populations in the Bismarcks and Solomons (e.g., *T. p. heinrothi* and *T. p. beehleri*), and on many occasions between Clades K and L in the far east and south of the island thrush’s range. The few cases of inferred recent gene flow among western populations include those between e.g. *T. p. katanglad* (central Mindanao) and *T. p. malindangensis* (northwest Mindanao); and between *T. p. stresemanni* (east-central Java) and the LCA of *T. p. javanicus* and *T. p. fumidus* (west and central Java).

**Fig. 3.**
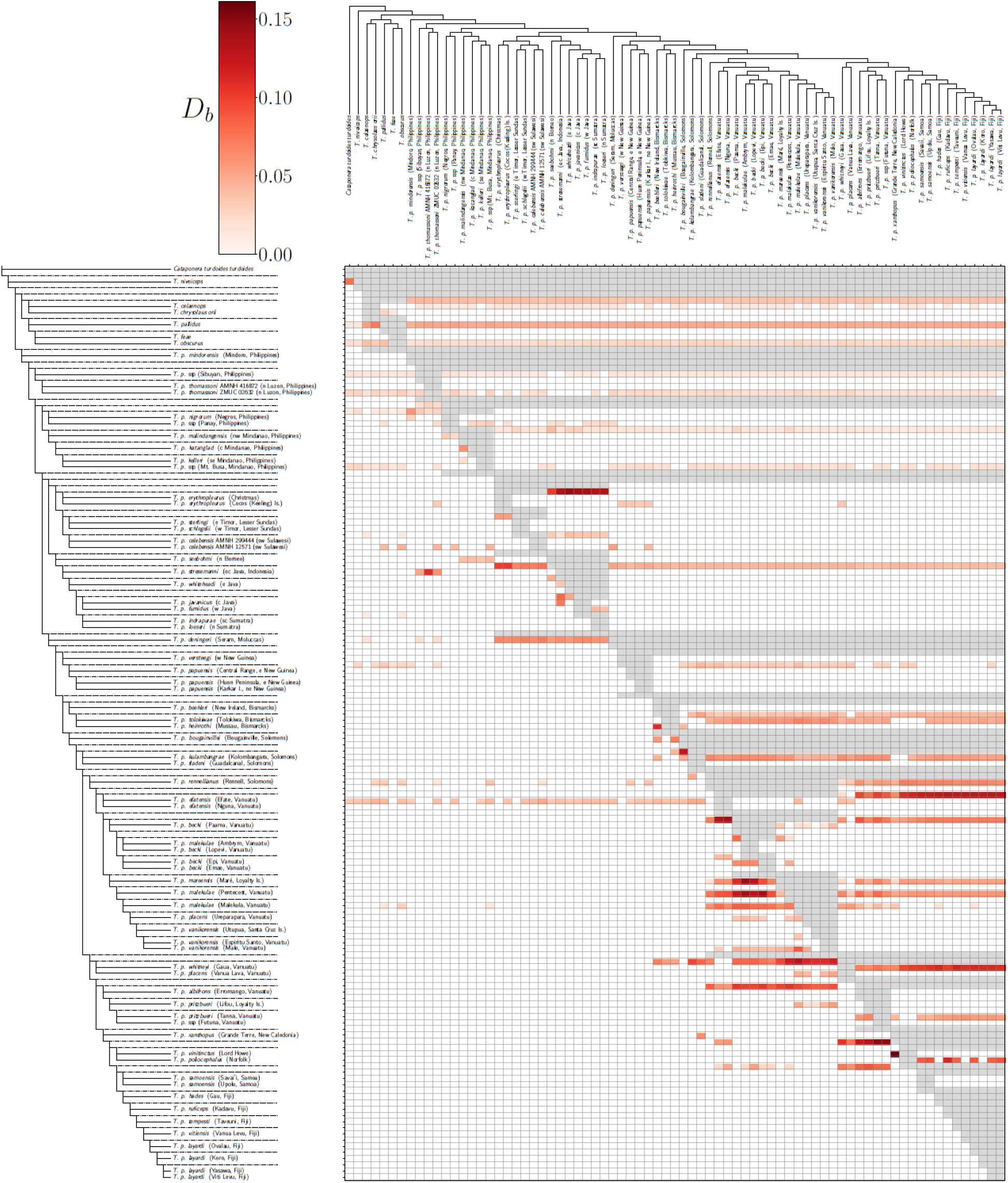
Tree violating branches and potential gene flow. The *D*_*b*_*(C)* statistic, analogous to the *f*_*b*_*(C)* statistic from Malinsky et al. (2018), summarizes the results of all D-statistic tests D(A, B, C, *T. merula*) that are consistent with the phylogenetic tree (Fig. 1). *D*_*b*_*(C)* measures excess allele sharing between individual (or ancestral node) C on the horizontal axis, and the branch of the tree *b* on the vertical axis (compared with *b*’s sister clade *a*). Each grid cell indicates one *D*_*b*_*(C)* statistic, where red cells correspond to significant values (more intense red indicates larger *D*_*b*_*(C)* values), white cells are non-significant, and gray corresponds to cells for which no statistic is consistent with the phylogenetic tree.

Patterns for certain populations suggest unlikely gene flow events. For example, the pattern for *T. p. efatensis* (Nguna, Vanuatu) implies numerous individual admixtures with most eastern island thrush populations, as well as the island thrush’s sister clade. This often occurs when two individuals from the same population (or two very closely-related populations) were sampled. In pairs belonging to *T. p. thomassoni, T. p. erythropleurus, T. p. celebensis*, and the aforementioned *T. p. efatensis*, one member of the pair shows an unrealistic pattern of gene flow, while the other does not. The individuals that constitute these pairs differ from one another quite markedly in data quality (Supplementary File 5), and these disparities likely caused the unrealistic patterns in the analysis results.

### Demographic history inference using PSMC

We used PSMC to infer the demographic histories of 60 individuals, representing 38 island thrush subspecies and six outgroup taxa (including *T*. [*poliocephalus*] *niveiceps*). We excluded individuals with sequencing depth < 10 (n = 16). The generated plots are presented in Fig. 4 and Supplementary File 4. The analyses cover the time period from c. 10 Mya to 10 Kya; the dates and effective population sizes (N_e_) reported here should be considered approximate, and reflective of temporal patterns of coalescent intensity (reflecting population structure), rather than census population sizes. However, the points at which PSMC curves diverge for different species correspond closely with the divergence time estimates from our mitochondrial genome tree (Fig. S3), and this guides our interpretation of the PSMC plots.

**Figs. 4a, 4b, 4c, 4d.**
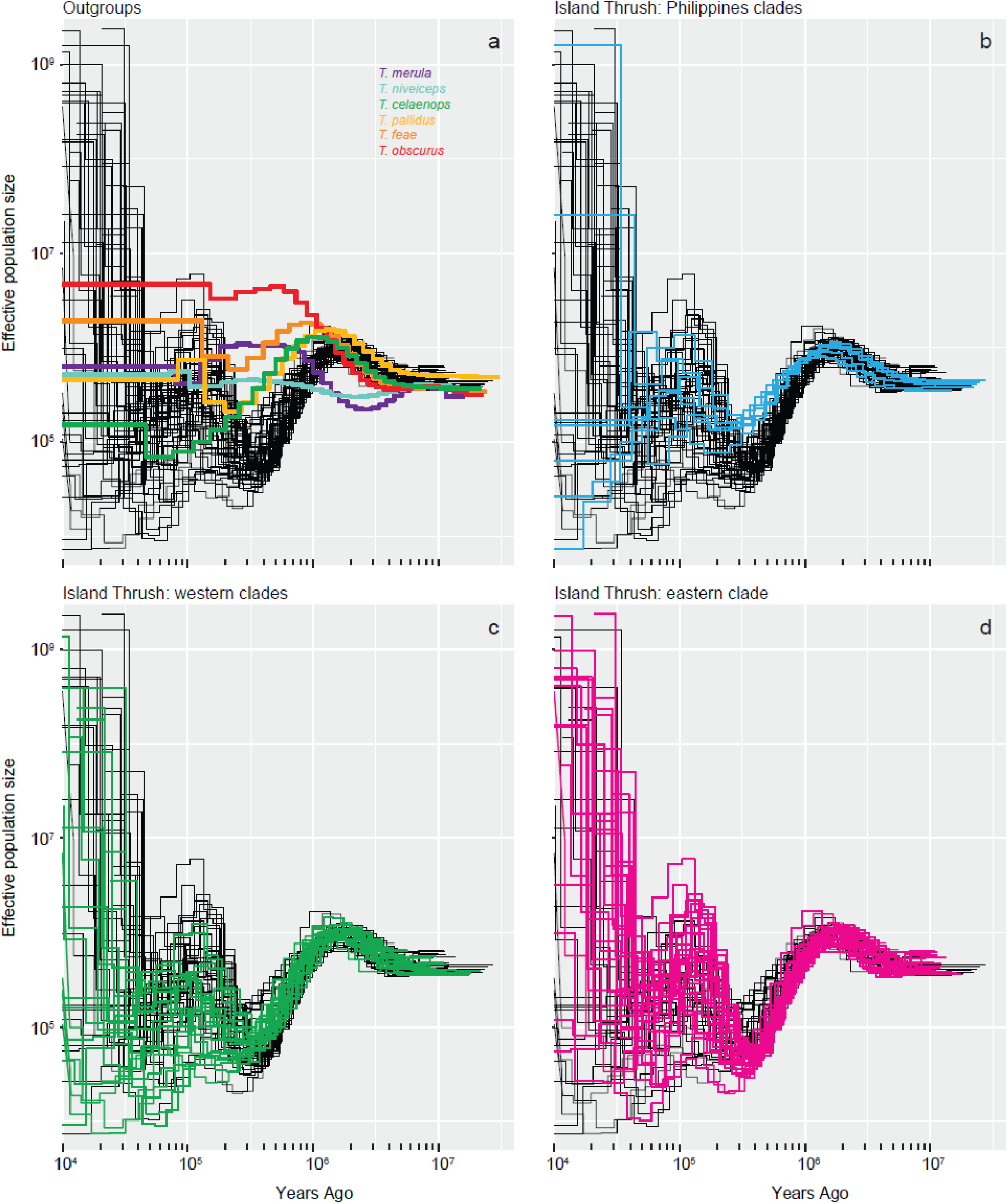
Pairwise sequentially Markovian coalescent (PSMC) plots illustrating demographic changes (effective population size; N_e_) over time for the island thrush and related *Turdus* thrushes. Generation time is set at two years. The four panels show curves from all island thrush individuals analyzed (black lines, with groups of interest highlighted in color). **4a** highlights outgroups; *T. celaenops, T. pallidus, T. feae*, and *T. obscurus* belong to the island thrush’s East Asian sister clade. **4b** highlights Philippines populations, inferred to be those earliest established. **4c** highlights all western clades outside the Philippines (Clades D–J in Fig. 1), as far east as the Solomons; and **4d** highlights the large eastern clade spanning southern Melanesia, Tasman Sea islands, and Polynesia (corresponding to sister clades K and L in Fig. 1). Individual N_e_ trajectories show little consistency more recently than c. 300 Kya, and within this recent timespan there is no clear west vs. east regional pattern, or montane (**4c**) vs. lowland (**4d**) pattern.

The common ancestral lineage had an estimated N_e_ of 600,000 individuals in the late Miocene. The outgroup taxon *T. merula* diverged from this lineage at 5–6 Mya, and maintained a fluctuating N_e_ of 200,000 to 1,000,000 until 10 Kya. The outgroup taxon *T. niveiceps* then diverged from the the island thrush ancestral lineage at c. 5 Mya, and maintained a fairly stable effective population size of 300,000–700,000 until 10 Kya. The ancestral island thrush lineage increased steadily from that point. The island thrush diverged from its sister clade at 2 Mya, which coincides with a N_e_ peak at 1,000,000. Species within the sister clade experienced a continued rise in N_e_ before diverging from one another slightly before 1 Mya. The island thrush’s N_e_ dropped steeply from its peak at 2 Mya. Philippines populations’ N_e_ curves started to subtly diverge at 1 Mya, and declined at a lower rate than the remainder of the island thrush lineage. Non-Philippines island thrushes declined steeply until reaching a low N_e_ of 50,000– 60,000 at 300 Kya; western clades began to stabilize slightly earlier than eastern clades. There is a poorly defined second hump of effective population growth and decline, which peaks, very roughly, at 100 Kya.

The curves present a very consistent overall pattern from 10 Mya to c. 300 Kya. The loss of concordance between 300 and 10 Kya likely reflects 1) that many populations were following variable individual trajectories at this point; and 2) that PSMC unable to adequately resolve very recent coalescent events (Li & Durbin, 2011).

### Geographic distance vs. genetic distance

We found a significant positive relationship between geographic distance and genetic distance (Fig. S8; r^2^ = 0.47, p < .001), indicating isolation by distance (Slatkin 1987, 1993) and supporting a stepping stone mode of colonization (Cibois et al. 2011; Irestedt et al. 2013).

### Colonization in light of Pleistocene land bridge formation

Populations with inferred Pleistocene land bridge connections share close phylogenetic relationships, indicating that Pleistocene cooling aided inter-island colonization. Results are presented in Fig. S9.

### Sexual dichromatism

Sexually dichromatic populations are scattered across the island thrush tree (Fig. S10), indicating that sexual dichromatism was gained and lost on numerous occasions.

## DISCUSSION

The island thrush represents a monophyletic island radiation that rapidly acquired its expansive geographic distribution within an estimated 1.3–2.4 million years. Its extreme plumage variation has obscured a biogeographically intuitive west-to-east stepping stone pattern of colonization from the Philippines through the Greater Sundas, Wallacea and New Guinea to Polynesia. With an aim to better understand the nature of archipelagic radiations and how they generate biodiversity, we here discuss the island thrush’s evolutionary origins, spatiotemporal radiation, population admixture, demographic history, and ecological and morphological variability.

### Evolutionary and geographic origins

The island thrush evolved from a clade of migratory *Turdus* thrushes with Palearctic/Sino-Himalayan breeding distributions (Nylander et al. 2008; Batista et al. 2020). Its sister clade comprises five East Asian species (Fig. 1) that range from short-distance partial migrants to long-distance migrants (Collar 2005). Four of five of these species are wholly or partly restricted to mountains within their breeding ranges (Collar 2005). Given this evolutionary background, the island thrush’s preference for cool (montane) habitat, and its evident ability to move across long distances, can be considered ancestral traits. The island thrush diverged from its continental sister clade c. 2.4 Mya, and began diversifying across the Indo-Pacific archipelagos c. 1.3 Mya (Fig. S3). Our results indicate that the first extant populations to be established were those from the Philippines, which correspond to the deepest splits in the tree (Fig. 1). How the ancestral island thrush reached the Philippines is unclear. Given that its ancestors were likely Palearctic migrants, and that two species from its sister clade winter in the Philippines, it is possible that colonization occurred via settling down of wintering birds (Rolland et al. 2014). However, the island thrush population on Mindoro is sister to the rest of the complex (Fig. 1) and appears to be one of the first established. Mindoro is one of the main arrival points for colonizers of the Philippines from Borneo (Diamond and Gilpin 1983; Jones and Kennedy 2008), which might suggest that a now-extinct population from the Greater Sundas colonized the Philippines.

### Spatiotemporal dynamics of the radiation

Diversification of the island thrush occurred during the second half of the Pleistocene, starting c. 1.3 Mya (Fig. S3). This is in line with dating estimates for other great speciators, which also radiated explosively during the Pleistocene (Moyle et al. 2009; Irestedt et al. 2013; Jønsson et al. 2014; Andersen et al. 2013, 2014, 2015; Pedersen et al. 2018; Kearns et al. 2020). The sequential branching pattern of the island thrush tree (Fig. 1) suggests that it expanded across most of its range following a stepping stone colonization path. Starting in the Philippines, it expanded into the Greater Sundas and Wallacea, colonized New Guinea and islands of Northern Melanesia, and then underwent overlapping radiations in southern Melanesia. In one of these radiations (Fig. 1, Clade L), the pattern of incremental advances gives way to long-distance oversea dispersals to reach far-away outposts in the Tasman Sea, Fiji, and Samoa. The overall stepping stone colonization process is supported by our regression analysis showing a positive relationship between geographic and genetic distance (Fig. S8). A similar pattern has been found for some other Indo-Pacific bird radiations (Cibois et al. 2011; Irestedt et al. 2013; Pedersen et al. 2018), but not all (Ericson et al. 2019).

Many of the islands that the island thrush inhabits have never shared subaerial connections (Fig. S9), so it is clear that repeated oversea colonizations have driven its current distribution. Nevertheless, water barriers have impeded its dispersal, and land bridge formation via Pleistocene cooling facilitated colonization. This is evident from the findings that populations are usually most closely related to other populations from the same island, and that populations connected by land bridges during Pleistocene glacial periods are in all cases closely related (Fig. S9). We find that the downslope expansion of montane forest habitat during the Pleistocene (Hewitt 2000, Garg et al. 2020) did not connect different populations on the same island (e.g. Mindanao, Greater Sundas, New Guinea) sufficiently to erase relatively deep genetic structure among them (Fig. S3).

### Gene flow between populations

The island thrush’s impressive dispersal capacity allowed it to repeatedly colonize new islands, but has also led to extensive admixture between established populations (Fig. 3). In western populations (Clades A–G in Fig. 1), most gene flow events appear to date back to the early phases of the radiation, when there was also admixture with the species’ East Asian sister clade. Gene flow between island populations to the east and south of New Guinea (Clades H–L in Fig. 1) is more recent and widespread. The many instances of gene flow between different branches of the island thrush phylogeny can help explain why topological inconsistencies exist across the phylogenetic trees (Figs. 1, S1, and S2).

The gene flow patterns provide new insight into the paradox of the great speciators (Diamond et al. 1976). A prominent hypothesis is that great speciators possess a uniformly moderate capacity for dispersal that is sufficient for colonization of new islands, but not sufficient for genetic and phenotypic homogenization across established populations (Diamond et al. 1976; Mayr and Diamond 2001). This is not the case in the island thrush: in recent times, population admixture and colonizations across deep-water barriers are much more frequent in eastern populations than in western populations. Another hypothesis is that the dispersal capacity of great speciators changes over time, usually imagined as an initial burst of rapid colonization followed by a sedentary phase of differentiation (Diamond et al. 1976). This model does not fit the island thrush as a whole, again because of the different dispersal patterns of eastern and western populations. The island thrush radiation might be better characterized as a rapidly advancing colonization front that leaves more sedentary populations in its wake. This is a dynamic seen (at much shorter timescales) in the spread of the invasive cane toad (*Rhinella marina*) across Australia (Phillips et al. 2007), where high dispersiveness is selected for at the edge of the expansion (Phillips et al. 2010). A similar mechanism could operate in the island thrush, assuming that island populations are founded by exceptionally dispersive individuals, but that dispersiveness is selected against in established populations because oversea dispersers leave those populations.

### Demographic history and genetic variation

The demographic history of the island thrush, and the modern genetic variation shown by its constituent populations, further elucidate its stepping stone colonization across the Indo-Pacific. The PSMC analyses (Fig. 4) imply that the lineage that spawned the island thrush and its sister clade experienced continuous growth in effective population size (N_e_) during the Pliocene and early Pleistocene, until beginning to diverge c. 2 Mya. This build-up was likely accompanied by range expansion across East Asia, as reflected by the clade’s current broad distribution across the region. As part of this expansion, the island thrush entered the Indo-Pacific archipelagos and experienced a steep N_e_ decline as gene flow with the continental lineage was lost. A similar rise-and-fall N_e_ dynamic is seen in *T. celaenops*, which colonized isolated Japanese islands (Fig. 4a), and in snowy-browed flycatcher (*Ficedula hyperythra*) (Pujolar et al. 2022), another passerine supercolonizer of island mountains with likely origins on the Asian mainland (Moyle et al. 2015). This pattern suggests that mainland population build-up can trigger archipelagic radiations. The oldest island thrush populations in the Philippines declined less steeply than other groups (Fig. 4b), and remaining western populations slowed their decline slightly earlier than eastern populations (Figs. 4c, 4d). This is likely because populations that were established earlier could regenerate genetic diversity earlier. The substantial population increases estimated for most populations starting c. 300 Kya are technical artifacts reflecting the limitations of the PSMC method at recent timescales (Li & Durbin, 2011; Nadachowska-Brzyska et al., 2015, 2016); most modern populations actually have low heterozygosity levels.

Heterozygosity levels of modern island thrush populations (Fig. 2) can be understood in light of the serial founder effect (Ramachandran et al. 2005). The oldest populations in the Philippines show the highest heterozygosity, probably retained in large part from the ancestral continental lineage. Repeated colonizations (and founder events) resulted in a general west-to-east pattern of decreasing heterozygosity. Most deviations from this pattern can be explained by island size (e.g., regionally low heterozygosity in small central Philippines island populations and regionally high heterozygosity in the large island of New Guinea). The most conspicuous exception to the overall pattern is the remarkably high heterozygosity of *T. p. stresemanni* of east-central Java, contrasted against the exceptionally low heterozygosity of the other six Javan and Sumatran subspecies. The analyses of population structure (Fig. S6) and gene flow (Fig. 3) both suggest that this is due to admixture of *T. p. stressemanni* with the island thrush lineage now inhabiting the northern Philippines.

### Elevational distribution

The island thrush is mostly restricted to mountains in the western part of its range from Sumatra to the Solomons. Here it rarely occurs below 1000 masl, but it reaches sea level on Christmas Island in the Indian Ocean, Mussau in the Bismarck Archipelago, and Rennell in the southern Solomon Islands (MacKinnon and Philipps 1993; Coates and Bishop 1997; Kennedy et al. 2000; Dutson 2011; Beehler and Pratt 2016). By contrast, it occurs down to sea level, or nearly so, on the generally small and low islands in the Tasman Sea, southern Melanesia, and Polynesia (Pratt et al. 1987; Collar 2005; Dutson 2011). Preference for cool (montane) climates can be regarded as an ancestral trait, based on the ecology of the island thrush’s close relatives (see above), and because montane populations correspond to the deepest splits in the island thrush phylogeny (Fig. 1). Following this interpretation, its variable elevational distribution is the result of three individual shifts (or expansions) into the lowlands (on Christmas Island, Mussau, and island south and east of the core Solomons). However, disentangling the mechanisms driving the island thrush’s elevational distribution is not straightforward. Diamond (1975) argues that populations’ elevational ranges are governed by diffuse competition, being pushed into the mountains on islands with high bird species richness in the lowlands. However, those isolated islands with impoverished bird faunas (e.g. southern Melanesia, Tasman Sea Islands, Polynesia) also lack key nest predators such as *Rattus* rats and *Boiga* snakes, or did at least prior to anthropogenic introductions. An alternative explanation is therefore that nest predation pressure restricts some populations to mountains (Skutch 1985; Boyle 2008; Jankowski et al. 2013), which could explain the susceptibility of lowland island thrush populations to introduced rats (Collar 2005, Villard et al. 2019). Either mechanism could explain the island thrush’s striking absence from Australia, which is highly biodiverse but mostly lacks tall mountains.

### Plumage color

Phylogenetic relationships alone do not explain the complex color variation in the island thrush, as populations that are not closely related often show convergent plumage types (e.g. *T. p. deningeri* of the Moluccas and *T. p. albifrons* / *T. p. pritzbueri* of southern Melanesia). Nevertheless, there is phylogenetic signal in its range-wide (male) plumage variation (Fig. 5). The three major clades from the Philippines represent some of the oldest populations (Fig. 1), and encompass a few distinct plumage types. Populations from the Greater Sundas, Christmas Island, and Wallacea west of Seram (Clades D and E in Fig. 1) are dark brown birds that vary primarily in the extent of reddish coloration of the underparts. The Seram population marks a shift to blackish body plumage, but has a white head. Overall blackish plumage — with varying degrees of white in the vent and undertail coverts — is common to most of the rest of the complex, from New Guinea to southern Melanesia. The striking exception from this pattern is the eastern Clade L (Fig. 1), which encompasses massive plumage variation, even among geographically and genetically similar Fiji populations. Based on this clade’s broad distribution in the Pacific, it is exceptionally dispersive, but it is not clear if or how this might be related to the pronounced variation in plumage. Overall, plumage variation is a poor proxy for phylogenetic distance in the island thrush.

**Fig. 5.**
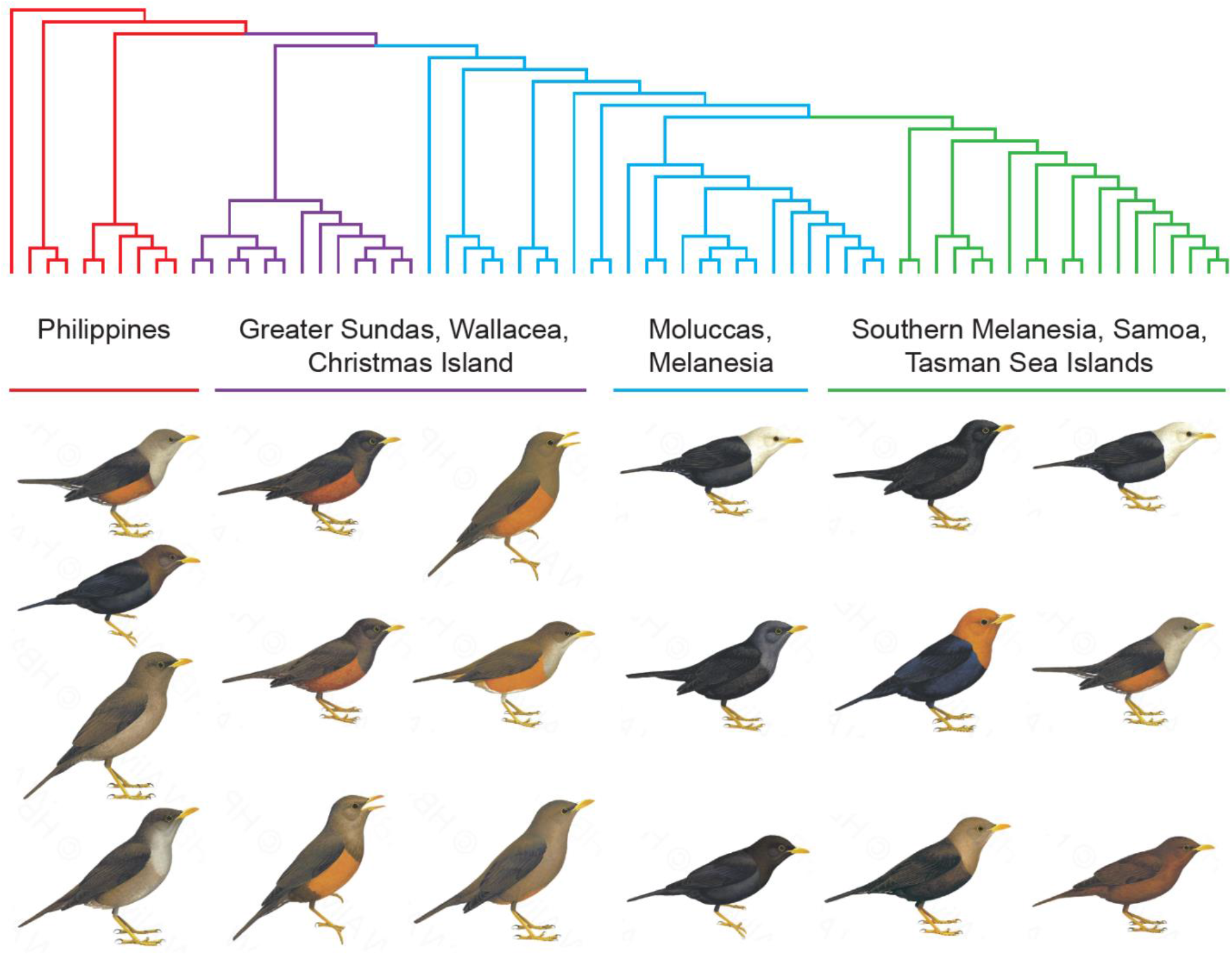
Plumage variation in light of phylogeny and geography. Tree topology matches that in Fig. 1. A selection of subspecies encompassing the range of male plumage variation found within each of four geographic groups are shown (illustrations: Lynx Edicions). The tree is colored to indicate the phylogenetic positions of those groups. The illustration of *T. p. albifrons* was used to represent the similar-looking *T. p. deningeri*, and the illustration of *T. p. mindorensis* was used to represent the similar-looking *T. p. layardi*.

The phylogenetic relationships of island thrush populations reveal numerous gains and losses of sexual dichromatism (Fig. S10). These transitions appear to occur haphazardly across the tree, having no obvious association with geographic region, island size, or elevational range. All members of the island thrush’s sister clade are sexually dichromatic (Collar 2005), as are most members of the broader Palearctic *Turdus* clade that it belongs to (Batista et al. 2020). Together with the fact that sexual dichromatism in the island thrush is mostly weak (Peterson 2007), this is consistent with a general pattern that island populations are less sexually dimorphic than their congeneric mainland populations (Omland 1997; Badyaev and Hill 2003). The seemingly random appearance and disappearance of sexual dichromatism may be attributable to repeated founder effects (Omland 1997; Kearns et al. 2020).

### Conclusion

Our study provides a detailed phylogenetic reconstruction of a great speciator. We demonstrate that the island thrush represents one of the most simultaneously explosive, expansive, and phenotypically diverse radiations among birds. The island thrush evolved from Palearctic ancestors and rapidly island hopped across a quarter of the globe, a journey that was facilitated by Pleistocene land bridge formation, but driven by repeated oversea dispersals. This stepping stone colonization left a clear signature of declining genetic variation from west to east. While representing extreme aspects of an archipelagic radiation, the island thrush provides a useful model for understanding Pleistocene interchange between Eurasian and Australo-Papuan faunas, and the role of mountains as pathways for temperate lineages to enter the tropics.

## MATERIALS AND METHODS

### Taxon sampling

Modern taxonomic treatments (Dickinson and Christidis 2014; Clements et al. 2019; Gill and Rasmussen 2020) recognize 50 island thrush subspecies, with slight variations in delimitation of a few forms. Aiming for comprehensive geographic and taxonomic coverage, we sampled 71 individuals representing 48 subspecies *sensu* IOC v10.2 (Gill and Rasmussen 2020) (Supplementary File 1). We sampled five recently extinct populations from the Pacific, as well as four undescribed populations from the Philippines and Vanuatu. Missing are *T. p. mayonensis* (s Luzon, Philippines) and the newly described *T. p. sukahujan* from Taliabu in Indonesia (Rheindt et al. 2020). We also sampled outside the island thrush complex to test its monophyly and elucidate its evolutionary background. We included the five members of its hypothesized east-Asian sister clade (Voelker et al. 2007; Nylander et al. 2008; Batista et al. 2020); the Sulawesi thrush (*Turdus* [*Cataponera*] *turdoides*), the island thrush’s Wallacean congener (Reeve et al. 2022); and the Taiwan thrush (*T*. [*poliocephalus*] *niveiceps*), removed from the island thrush complex on the basis of Nylander et al. (2008), but with uncertain phylogenetic placement. The common blackbird (*T. merula*), which diverged from the island thrush c. 5 Mya (Batista et al. 2020), was sampled to generate a *de novo* assembled reference genome (see Materials and methods: Bioinformatics: *de novo* reference genome assembly).

### Library preparation and sequencing

We extracted genomic DNA from toepad samples (n = 59) and from fresh blood and tissue samples (n = 19). Protocol for DNA extraction from toepad samples followed Irestedt et al. (2006). We followed the protocol of Meyer and Kircher (2010) to create sequencing libraries suitable for Illumina sequencing of toepad DNA extracts. Library preparation included blunt-end repair, adapter ligation, and adapter fill-in, followed by four independent index PCRs. The libraries were run on half a lane on Illumina HiSeq X, pooled at equal ratio with other museum samples. Genomic DNA was extracted from fresh samples with KingFisher Duo magnetic particle processor (ThermoFisher Scientific) using the KingFisher Cell and Tissue DNA Kit. Library preparation, using Illumina TruSeq DNA Library Preparation Kit, and sequencing on Illumina HiSeqX (2×151 bp) was performed by SciLifeLab. All raw reads generated for this study have been deposited at the NCBI Sequence Read Archive (SRA), accession number [pending].

### Bioinformatics

#### *de novo* reference genome assembly

A *Turdus merula* reference genome for mapping was generated by SciLifeLab using Neutronstar (https://github.com/nf-core/neutronstar), a NextFlow pipeline for the *de novo* assembly of 10X Chromium linked reads. Neutronstar employs Supernova (Weisenfeld et al. 2017) for *de novo* assembly and uses BUSCO (Simão et al. 2015) and QUAST (Gurevich et al. 2013) to evaluate assembly quality. Assembly statistics are summarized in Table S1.

#### Read cleaning and mapping

Illumina sequencing reads were processed using a custom-designed workflow to remove adapter contamination, low-quality bases, and low-complexity reads (available at https://github.com/mozesblom). Overlapping read pairs were merged using PEAR (v.0.9.10; Zhang et al. 2014), and Super Deduper (v.1.4; Petersen et al. 2015) was used to remove PCR duplicates. Trimming and adapter removal was performed using TRIMMOMATIC (v.0.32; Bolger et al. 2014; default settings). The overall quality and length distribution of sequence reads was inspected using FASTQC (v.0.11.5; Andrews 2010), before and after the cleaning.

Cleaned reads were mapped to the reference using BWA-MEM (Li 2013), PCR duplicates were removed with Picard MarkDuplicates (Broad Institute 2019), and GATK’s RealignerTargetCreator and IndelRealigner were used to realign reads around indels (McKenna et al. 2010) (mapping pipeline and scripts for subsequent analysis of the nuclear data are available from https://github.com/grahamgower/island_thrush_scripts). To avoid low-complexity reads and potential contamination, only reads with MAPQ ≥ 30 were retained, which resulted in a mean depth of coverage of 11.77 ± 3.69. The same pipeline was used to reconstruct mitochondrial genomes (mean depth of coverage 396.44 ± 314.27). Reads were mapped to a reference mitochondrial genome sequence from *Turdus mandarinus* (Genbank accession no. NC_028188). We then created consensus mitochondrial sequences for each individual using htsbox (https://github.com/lh3/htsbox).

#### Post-mortem damage

Many of our genetic samples derive from study skins collected in the early 20^th^ century, and we therefore assessed post-mortem DNA damage patterns. This was done using condamage (https://github.com/grahamgower/condamage), which revealed considerable cytosine deamination in many individuals (Supplementary File 3). Condamage also implements an approach by Meyer et al. (2016) to inspect per-read patterns of deamination on one read end, conditional on there also being damage at the other end. In the case of a contaminated specimen, the conditional and unconditional deamination rates can differ, due to the unconditional proportion being calculated from both endogenous and exogenous reads, while the conditional proportion is calculated from (mostly) endogenous reads. Similarly, the DNA fragment length distributions for damaged versus undamaged reads can differ for contaminated specimens, as exogenous reads are typically introduced at the time of sample processing and sequencing, and are thus longer than the degraded endogenous reads. However, no contamination was identified from this investigation.

#### Exclusion of sex-linked contigs

Sex chromosomes can have vastly different mutation rates, recombination rates, and effective population sizes compared with autosomes. We therefore wanted to restrict genomic analyses to autosomal data. One approach to identify sex-linked contigs is to map a reference assembly to a chromosome-level assembly of a closely related species. However, no such assembly was available for this purpose.

Under the assumption that reads are sampled from any given chromosome in proportion to the chromosome’s length and copy number, we then expect that the proportion of reads that map to any given sex-linked contig will differ between the two sexes. Given that both males and females are represented in our dataset, the contigs may be stratified into Z-linked or autosomal, depending on the proportion of reads mapping to the contigs in the respective male and female cohorts. In principle, we could use read dosage to identify W-linked contigs as well, but the individual used to construct our reference was male, and thus the assembly does not contain any W-linked contigs.

We stratified the contigs with length > 100 kbp, using principal components analysis (PCA) applied to **M**, an *m* × *n* matrix of *m* contigs and *n* individuals, which has entries

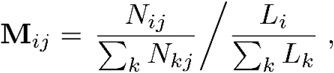

where *N*_*ij*_ is the number of reads mapped to contig *i* in individual *j*, and *L*_*i*_ is the length of contig *i*. The contig length normalization follows from our earlier assumption that the number of reads mapping to a given contig will be proportional to its length. PCA was performed on the mean-centered covariance matrix of **M**, which produced a clear separation of contigs into two clusters along PC2 (Fig. S11).

A well-known contributor to non-uniform sequencing coverage is an association between local GC% and sequencing depth (Benjamini and Speed 2012). By regressing PC1 against GC%, we determined that GC% is a contributor to the primary source of read dosage variation in our sample (Pearson *R*^2^=0.30, p=4.9e-66). To correct for this, we separately regressed each individual column in **M** against the contig GC proportions, then performed an additional PCA on the matrix of residuals. The GC-corrected PCA separated the contigs into two major groups along PC1 (Fig. S12), indicating that a systematic difference between two groups of contigs drives the remaining variation. We designated the smaller cluster of contigs, comprised of fewer nucleotides, as Z-linked, and the larger cluster as autosomal. The number of contigs for each group, and their total nucleotide count, are provided in Table S2.

To confirm that the two contig groups were indeed separating based on Z-chromosome copy number, we genetically sexed each individual using the autosomal and Z-linked contigs assigned from the GC-corrected PCA (Gower et al. 2019). Of the 65 individuals for which phenotypic sex information was available, 64 of the genetic sexes matched. We note that very similar results were obtained when using the non-GC-corrected PCA, but we restricted all further analyses to data mapped to the set of contigs identified as autosomal from the GC-corrected PCA. We also note that a similar method was recently developed by Nursyifa et al. (2021) to address the same problem we tackle here.

#### SNP ascertainment and genotype likelihoods

We obtained a set of candidate SNP sites by calling major and minor alleles using ANGSD (Korneliussen et al. 2014) with a SNP p-value cutoff of 10^−6^. We used only the individuals with average depth of sequencing greater than 10 (n=61). To reduce the impact of post-mortem damage in our ancient museum specimens, the set of sites was further filtered to exclude all transitions (C/T and G/A SNPs). Specifically, we used the following ANGSD parameters: - minMapQ 30 -minQ 20 -baq 2 -C 50 -uniqueOnly 1 -noTrans 1 -minInd 30 -GL 1 -doMaf 1 - doGlf 2 -doMajorMinor 1 -doSaf 1 -SNP_pval 1e-6. This yielded 8,476,511 SNPs across the autosomal contigs, at a density of approximately one SNP per 64 base pairs. Genotype likelihoods at the ascertained sites were then determined for all individuals (including those with low sequencing depth; n=77) using ANGSD with the same filtering options as before. In addition to the genotype likelihoods for the full set of individuals, we also used a genotype likelihood dataset comprising only *Turdus poliocephalus* individuals. We excluded *T. p. canescens* from further analysis, as it had low sequencing depth (= 0.74) and low coverage (0.47); preliminary investigations indicated a high error rate for this individual (i.e., negative pairwise F_ST_ values, and large pairwise distances, to all other individuals). We did not call SNPs for *T. p. hygroscopus*, which was a last-minute addition to the dataset. Both *T. p. canescens* and *T. p. hygroscopus* were included in the phylogenetic analysis of mitochondrial genome data (see Materials and methods: Phylogenetic analyses: Phylogenetic analysis of mitochondrial genome data).

### Phylogenetic analyses

#### Phylogenetic analysis of SNP data

We constructed genome-wide trees both from pairwise distances and from pairwise F_ST_, using *T. merula* as an outgroup to root the trees. Pairwise distances were calculated from genotype likelihoods using ngsDist (Vieira et al. 2016). Pairwise F_ST_ was calculated from genotype likelihoods using the folded joint (2D) allele frequency spectrum (Nielsen et al. 2012), as implemented in ANGSD (Korneliussen et al. 2014), configured to use the default Reynolds et al. (1983) F_ST_ estimator.

Trees were estimated from distance matrices using neighbor-joining (Saitou and Nei 1987), followed by subtree pruning and regrafting, as implemented in FastME (Lefort et al. 2015). In addition to constructing a whole-data pairwise distance tree, we obtained 100 bootstrap replicates of the pairwise distance matrix from ngsDist, configured to use blocks of 1500 SNPs (for an expected block size of 96 kbp), and constructed neighbor-joining trees for each replicate. Bootstrap support values were then assigned to internal nodes of the whole-data tree using RAxML (Stamatakis 2014). We note that bootstrap support values only indicate how consistent the data are across the genome, and do not reflect uncertainty of relationships due to e.g. gene flow or incomplete lineage sorting.

#### Phylogenetic analysis of mitochondrial genome data

We performed a Bayesian phylogenetic analysis of the newly generated mitochondrial genome data, supplemented with Genbank data from 12 additional outgroup taxa from the family Turdidae (see Supplementary File 1). Two island thrush taxa are included here that were excluded from the phylogenetic analyses described above due to poor quality SNP data (*T. p. canescens*) or lacking SNP data (*T. p. hygroscopus*). We excluded two mitochondrial genomes with suspected pseudogene contamination (*T. merula* and *T. poliocephalus papuensis* ZMUC 192303).

We built individual alignments for cytochrome *b* (cyt-*b*), NADH dehydrogenase 2 (ND2), and the remainder of the mitochondrial genome using MAFFT (Katoh et al. 2002) as implemented in SEAVIEW (Gouy et al. 2010). Subsequently, we analyzed the concatenated datasets, with cyt-*b* and ND2 partitioned, in BEAST v1.8.4 (Drummond et al. 2012) using the GTR nucleotide substitution model. We used a relaxed uncorrelated lognormal distribution for the molecular clock model, and assumed a birth-death speciation process as a tree prior. *Myadestes myadestinus* was set as the outgroup. The Markov chain Monte Carlo (MCMC) algorithm was run three times for 200 million iterations, with trees sampled every 10,000th generation. Convergence of individual runs was assessed using Tracer 1.6 (Rambaut et al. 2014), ensuring all ESS > 200, and graphically estimating an appropriate burn-in (55 million generations). TreeAnnotator 1.8.2 (Rambaut and Drummond 2015) was used to summarize a single maximum clade credibility (MCC) tree using mean node heights. To obtain absolute dates, we followed substitution rate estimates from Lerner et al. (2011), which derive from analysis of a Passerides songbird radiation across Pacific islands. We applied a rate of 0.0145 substitutions per site per lineage (2.9%) per Myr to our ND2 data; and a rate of 0.007 substitutions per site per lineage (1.4%) per Myr to our cyt-*b* data.

#### Population structure and heterozygosity levels

Population structure was analyzed for the island thrush, excluding outgroups, using PCAngsd (Meisner and Albrechtsen 2018). We performed a principal component analysis to explore the genetic differentiation represented in the data. A covariance matrix was estimated from genotype likelihoods, configured to exclude sites with minor allele frequency <0.05 and sites not in Hardy-Weinberg equilibrium. The PCAngsd minimum average partial (MAP) test was used to calculate the number of dimensions required to explain the population structure in this dataset. To account for individual variation in heterozygosity, we normalized the covariance matrix to obtain a correlation matrix, and then visualized the result as a heatmap. We used a latent mixed-membership model implemented in PCAngsd to estimate ancestry proportions for k=2 to k=8 ancestral components. Heterozygosity was calculated from the site allele frequency spectrum for each individual, as estimated by ANGSD.

#### Gene flow

We used D-statistics (Green et al. 2010, Patterson et al. 2012) to test for differential gene flow between populations. The *D*_*b*_*(C)* statistic is the D-statistic analogue of Malinsky et al.’s (2018) *f*_*b*_*(C)* statistic,

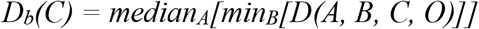

which minimizes over all clades B that are descendants of *b*, and takes the median over all clades A that are descendants of *b*’s sister branch *a*. To obtain this statistic, we first computed D-statistics of the form *D(A, B, C, T. merula)* for all sets of three individuals A, B, C that were consistent with the tree presented in Fig. 1, with the order of A and B chosen so that each statistic was positive. Computing D-statistics from genotype likelihoods using ANGSD was not computationally feasible. Instead, we first made hard genotype calls on the autosomal scaffolds (those with length > 100 kbp) using BCFtools call -m (Li 2011). Pseudohaploid genotypes were then obtained for each individual by taking the majority read at sites called as biallelic, excluding transversions (C-T or G-A substitutions) and sites within 10 bp of an indel call (https://github.com/grahamgower/eig-utils). Significance of the D-statistics were assessed via bootstrapping, with each bootstrap replicate obtained by sampling scaffolds with replacement to match the length of the autosome (544 mbp; see Table S2). To account for multiple testing, p-values were Holm-Bonferroni adjusted to achieve a family-wise error rate (FWER) of 0.05. We used Dsuite (Malinsky et al. 2021) to calculate the *D*_*b*_*(C)* statistics from our pairwise-distance tree and precalculated D-statistics, then plotted the results using Dsuite’s dtools.py script.

#### Demographic history inference using PSMC

We used the pairwise sequentially Markovian coalescent method (PSMC; Li and Durbin 2011) to infer the demographic histories of the island thrush and its relatives. PSMC uses the distribution of heterozygous sites across the genome of a single individual to infer the demographic history of an entire population or species. The method estimates the distribution of the time since the most recent common ancestor (TMRCA) of each allele pair at all loci, and uses this to estimate effective population size changes over time.

We performed individual PSMC analyses for all ingroup and outgroup taxa, including the *T. merula de novo* assembly. Non-autosomal regions were excluded, and variants were called with the mpileup and call -c subcommands in BCFtools (independently of the genotype calling described in Materials and methods: Gene flow). We filtered out sites where read depth was less than 10 or more than 100, sites with Phred quality scores below 20, and sites near indels. We then used the ‘consensus’ command in BCFtools to incorporate all variants into a single sequence using IUPAC codes. Next, the consensus sequence was divided into non-overlapping 100 bp bins, which were scored either as heterozygous (if there was at least one heterozygote nucleotide position in the bin), or homozygous. When running PSMC, the total number of expectation-maximization iterations was set to 25; T max (-t) was set to 15; the initial mutation/recombination ratio (-r) was set to 5; and the atomic time interval pattern (-p) was set to “4+25*2+4+6”. Results were scaled using a generation time of two years and a mutation rate of 3×10^−9^ per nucleotide per generation, based on the rates reported for passerine birds by Nadachowska-Brzyska et al. (2015).

#### Geographic distance vs. genetic distance

A positive relationship between pairwise geographic distance and genetic distance indicates isolation by distance (Slatkin 1987, 1993), a pattern that can result from stepping stone colonization (Cibois et al. 2011; Irestedt et al. 2013). To test this for the island thrush, we performed a Mantel test (10,000 permutations) using the ape package (Paradis and Schliep 2019) in R v3.5.2 (R Core Team 2018).

#### Colonization in light of Pleistocene land bridge formation

Pleistocene glacial cycles caused repeated drops in global sea levels, sometimes by as much as 120 m. This resulted in periodic land bridge connections between many Indo-Pacific islands (Voris 2000). To evaluate whether these connections facilitated inter-island colonization by the island thrush, we inferred which populations shared subaerial connections during the Pleistocene. Populations separated by water barriers deeper than 120 m were considered not to have been connected. These data were plotted across the tips of the pairwise distance tree.

#### Sexual dichromatism

In addition to high variation in plumage coloration and patterning, the island thrush represents a mosaic of sexually dichromatic and monochromatic populations. We used data from a comprehensive morphological study of the species by Peterson (2007) to determine how often sexually dichromatism arose by convergence. Peterson measured dichromatism as light, strong, or absent, but since only two taxa showed ‘strong’ dimorphism, including *T*. (*poliocephalus*) *niveiceps*, which is not an island thrush (Nylander et al. 2008), we scored dichromatism only as present or absent. Dichromatism scores were plotted across the tips of the pairwise distance tree; we were able to apply scores to 55 of 68 populations.

## SUPPLEMENTARY MATERIAL

**Supplementary File 1**. Specimen and sequence data accession information.

**Supplementary File 2**. Figures S1–12 and Tables S1–2.

**Supplementary File 3**. Post-mortem damage.

**Supplementary File 4**. PSMC plots.

**Supplementary File 5**. Data quality information and metadata.

## ACKNOWLEDGMENTS

Generous sample loans from several museums provided the basis for this study. We thank the American Museum of Natural History, New York, NY (Paul Sweet, Tom Trombone and Peter Capainolo); the British Museum of Natural History, Tring, UK (Robert Prys-Jones, Hein van Grouw and Mark Adams); the Burke Museum, Seattle, WA (Sharon Birks); the Cincinnati Museum Center, OH (Emily Imhoff); the Field Museum of Natural History, Chicago, IL (Ben Marks); the Natural History Museum of Denmark (Jan Bolding Kristensen); the Queensland Museum, South Brisbane, Australia (Heather Janetzki, Paul Oliver); Rijksmuseum van Natuurlijke Histoire, Leiden, the Netherlands (Steven van der Mije and Pepijn Kamminga); the Smithsonian Institution National Museum of Natural History, Washington, D.C. (Christopher Milensky); the Swedish Museum of Natural History, Stockholm (Ulf Johansson); and the Yale Peabody Museum of Natural History, New Haven, CT (Kristof Zyskowski). Leo Joseph, Frederick Sheldon, and Trevor Price provided valuable comments on the draft manuscript. We thank Thorfinn Korneliussen for adding folded 2D-SFS support into ANGSD. This work was supported by the Carlsberg Foundation (grant numbers CF15-0078 and CF15-0079 to K.A.J.); and the Villum Foundation (Young Investigator Programme grant number 15560 to K.A.J., and grant number 00025300 to F.R.). The authors acknowledge support from the National Genomics Infrastructure in Stockholm, funded by Science for Life Laboratory, the Knut and Alice Wallenberg Foundation and the Swedish Research Council. We also thank SNIC/Uppsala Multidisciplinary Center for Advanced Computational Science for assistance with massively parallel sequencing, and access to the UPPMAX computational infrastructure.

## AUTHOR CONTRIBUTIONS

A.H.R. and K.A.J. conceived the study. All authors contributed to build the dataset. A.H.R., G.G., F.R., and K.A.J. developed the analytical framework. G.G., M.P.K.B., B.P., and F.R. performed bioinformatics. G.G. and J.M.P. performed the phylogenomic analyses with input from A.H.R., F.R., and K.A.J. A.H.R. led the writing, and all authors contributed to the discussion of the results and the writing of the manuscript.

## DATA AVAILABILITY

Raw Illumina sequences and the *Turdus merula* genome assembly are deposited in the Sequence Reads Archive, National Center for Biotechnology Information, SRA accession [pending]. Some mitochondrial genome sequence data was downloaded from Genbank; accession numbers are provided in Supplementary File 1.

## REFERENCES

Andersen MJ, Nyári ÁS, Mason I, Joseph L, Dumbacher JP, Filardi CE, Moyle, RG. 2014. Molecular systematics of the world’s most polytypic bird: the Pachycephala pectoralis/melanura (Aves: Pachycephalidae) species complex. J Linn Soc Lond Zool. 170:566–588.

Andersen MJ, Oliveros CH, Filardi CE, Moyle RG. 2013. Phylogeography of the Variable Dwarf-Kingfisher Ceyx lepidus (Aves: Alcedinidae) inferred from mitochondrial and nuclear DNA sequences. Auk. 130:118–131.

Andersen MJ, Shult HT, Cibois A, Thibault JC, Filardi CE, Moyle, RG. 2015. Rapid diversification and secondary sympatry in Australo-Pacific kingfishers (Aves: Alcedinidae: Todiramphus). R Soc Open Sci. 2:140375.

Andrews S. 2010. FastQC: a quality control tool for high throughput sequence data. Available from: http://www.bioinformatics.babraham.ac.uk/projects/fastqc

Badyaev AV, Hill GE. 2003. Avian sexual dichromatism in relation to phylogeny and ecology. Annu Rev Ecol Evol Syst. 34:27–49.

Batista R, Olsson U, Andermann T, Aleixo A, Ribas CC, Antonelli A. 2020. Phylogenomics and biogeography of the world’s thrushes (Aves, Turdus): new evidence for a more parsimonious evolutionary history. Proc R Soc Lond B Biol Sci. 287:20192400.

Beehler BM, Pratt TK. 2016. Birds of New Guinea: distribution, taxonomy, and systematics. Princeton (NJ): Princeton University Press.

Benjamini Y, Speed TP. 2012. Summarizing and correcting the GC content bias in high-throughput sequencing. Nucleic Acids Res. 40:e72.

Bolger AM, Lohse M, Usadel B. 2014. Trimmomatic: a flexible trimmer for Illumina sequence data. Bioinformatics. 30:2114–2120.

Boyle WA. 2008. Can variation in risk of nest predation explain altitudinal migration in tropical birds?. Oecologia. 155:397–403.

Broad Institute 2019. Picard toolkit. Available from: http://broadinstitute.github.io/picard/

Cibois A, Beadell JS, Graves GR, Pasquet E, Slikas B, Sonsthagen SA, Thibault J-C, Fleischer RC. 2011. Charting the course of reed-warblers across the Pacific islands. J Biogeogr. 38:1963–1975.

Clement P, Hathway R. 2000. Thrushes. London (GB): Christopher Helm.

Clements JF, Schulenberg TS, Iliff MJ, Billerman SM, Fredericks TA, Sullivan BL, Wood CL. 2019. The eBird/Clements Checklist of Birds of the World: v2019. Available from: https://www.birds.cornell.edu/clementschecklist/download/

Coates BJ, Bishop KD. 1997. A guide to the birds of Wallacea. Alderley (QLD): Dove Publications.

Collar NJ. 2005. Family Turdidae (Thrushes). In: del Hoyo J, Elliott A, Christie D, editors. Handbook of the Birds of the World vol. 10. Barcelona (CAT): Lynx Edicions. p. 514–807.

Darwin C. 1859. On the origin of species by means of natural selection, or, the preservation of favoured races in the struggle for life. London (GB): J. Murray.

Diamond JM. 1975. Assembly of Species Communities. In: Cody M.L, Diamond J, editors. Ecology and Evolution of Species Communities. Cambridge (MA): Harvard University Press. p. 342–444.

Diamond JM, Gilpin ME. 1983. Biogeographic umbilici and the origin of the Philippine avifauna. Oikos. 41:307–321.

Diamond JM, Gilpin ME, Mayr E. 1976. Species-distance relation for birds of the Solomon Archipelago, and the paradox of the great speciators. Proc Natl Acad Sci U S A. 73:2160–2164.

Dickinson EC, Christidis L, editors. 2014. The Howard and Moore complete checklist of the birds of the world. 4th ed. vol. 2. Passerines. Eastbourne (GB): Aves Press.

Drummond AJ, Suchard MA, Xie D, Rambaut A. 2012. Bayesian phylogenetics with BEAUti and the BEAST 1.7. Mol Biol Evol. 29:1969–1973.

Dutson G. 2011. Birds of Melanesia: Bismarcks, Solomons, Vanuatu and New Caledonia. London (GB): Christopher Helm.

Ericson PG, Qu Y, Rasmussen PC, Blom MP, Rheindt FE, Irestedt M. 2019. Genomic differentiation tracks earth-historic isolation in an Indo-Australasian archipelagic pitta (Pittidae; Aves) complex. BMC Evol Biol. 19:1–13.

Garg KM, Chattopadhyay B, Koane B, Sam K, Rheindt FE. 2020. Last Glacial Maximum led to community-wide population expansion in a montane songbird radiation in highland Papua New Guinea. BMC Evol Biol. 20:1–10.

Gill F, Donsker D, Rasmussen P, editors. 2020. IOC World Bird List (v10.2). doi: 10.14344/IOC.ML.10.2.

Gouy M, Guindon S, Gascuel O. 2010. SeaView version 4: a multiplatform graphical user interface for sequence alignment and phylogenetic tree building. Mol Biol Evol. 27:221–224.

Gower G, Fenderson LE, Salis AT, Helgen KM, van Loenen AL, Heiniger H, Hofman-Kamińska E, Kowalczyk R, Mitchell KJ, Llamas B, et al. 2019. Widespread male sex bias in mammal fossil and museum collections. Proc Natl Acad Sci U S A. 116:19019–19024.

Green RE, Krause J, Briggs AW, Maricic T, Stenzel U, Kircher M, Patterson N, Li H, Zhai W, Fritz MH, et al. 2010. A draft sequence of the Neandertal genome. Science. 328:710–722.

Gurevich A, Saveliev V, Vyahhi N, Tesler G. 2013. QUAST: quality assessment tool for genome assemblies. Bioinformatics (Oxf). 29:1072–1075.

Gwee, CY, Garg, KM, Chattopadhyay, B, Sadanandan, KR, Prawiradilaga, DM, Irestedt, M, Lei F, Bloch LM, Lee JGH, Irham M, et al. 2020. Phylogenomics of white-eyes, a ‘great speciator’, reveals Indonesian archipelago as the center of lineage diversity. eLife. 9:e62765.

Hewitt G. 2000. The genetic legacy of the Quaternary ice ages. Nature. 405:907–913.

Irestedt M, Fabre PH, Batalha-Filho H, Jønsson KA, Roselaar CS, Sangster G, Ericson PG. 2013. The spatio-temporal colonization and diversification across the Indo-Pacific by a ‘great speciator’(Aves, Erythropitta erythrogaster). Proc R Soc Biol Sci Ser B. 280:20130309.

Jankowski JE, Londoño GA, Robinson SK, Chappell MA. 2013. Exploring the role of physiology and biotic interactions in determining elevational ranges of tropical animals. Ecography. 36:1–12.

Jones AW, Kennedy RS. 2008. Plumage convergence and evolutionary history of the Island Thrush in the Philippines. Condor. 110:35–44.

Jønsson KA, Irestedt M, Christidis L, Clegg SM, Holt BG, Fjeldså J. 2014. Evidence of taxon cycles in an Indo-Pacific passerine bird radiation (Aves: Pachycephala). Proc R Soc Biol Sci Ser B. 281:20131727.

Katoh K, Misawa K, Kuma KI, Miyata T. 2002. MAFFT: a novel method for rapid multiple sequence alignment based on fast Fourier transform. Nucleic Acids Res. 30:3059–3066.

Kearns AM, Joseph L, Austin JJ, Driskell AC, Omland KE. 2020. Complex mosaic of sexual dichromatism and monochromatism in Pacific robins results from both gains and losses of elaborate coloration. J Avian Biol. 51:1–19.

Kennedy R, Gonzales PC, Dickinson E, Miranda Jr HC, Fisher TH. 2000. A guide to the birds of the Philippines. New York (NY): Oxford University Press.

Korneliussen TS, Albrechtsen A, Nielsen R. 2014. ANGSD: analysis of next generation sequencing data. BMC Bioinformatics. 15:1–13.

Lawson DJ, Van Dorp L, Falush D. 2018. A tutorial on how not to over-interpret STRUCTURE and ADMIXTURE bar plots. Nat. Commun. 9:1–11.

Lefort V, Desper R, Gascuel O. 2015. FastME 2.0: a comprehensive, accurate, and fast distance-based phylogeny inference program. Mol Biol Evol. 32:2798–2800.

Lerner HR, Meyer M, James HF, Hofreiter M, Fleischer RC. 2011. Multilocus resolution of phylogeny and timescale in the extant adaptive radiation of Hawaiian honeycreepers. Curr Biol. 21:1838–1844.

Li H. 2011. A statistical framework for SNP calling, mutation discovery, association mapping and population genetical parameter estimation from sequencing data. Bioinformatics (Oxf). 27:2987–2993.

Li H. 2013. Aligning sequence reads, clone sequences and assembly contigs with BWA-MEM. arXiv. 1303.3997.

Li H, Durbin R. 2011. Inference of human population history from individual whole-genome sequences. Nature. 475:493–496.

Li H, Handsaker B, Wysoker A, Fennell T, Ruan J, Homer N, Marth G, Abecasis G, Durbin R. 2009. The sequence alignment/map format and SAMtools. Bioinformatics (Oxf.). 25:2078–2079.

MacArthur RH, Wilson EO. 1967. The theory of island biogeography. Princeton (NJ): Princeton University Press.

MacKinnon J, Phillipps K. 1993. A field guide to the birds of Borneo, Sumatra, Java, and Bali. New York (NY): Oxford University Press.

Malinsky M, Matschiner M, Svardal H. 2021. Dsuite-Fast D-statistics and related admixture evidence from VCF files. Mol Ecol Resour. 21:584–595.

Malinsky M, Svardal H, Tyers AM, Miska EA, Genner MJ, Turner GF, Durbin R. 2018. Whole-genome sequences of Malawi cichlids reveal multiple radiations interconnected by gene flow. Nat Ecol Evol. 2:1940–1955.

Manthey JD, Oliveros CH, Andersen MJ, Filardi CE, Moyle RG. 2020. Gene flow and rapid differentiation characterize a rapid insular radiation in the southwest Pacific (Aves: Zosterops). Evolution. 74:1788–1803.

Mayr E. 1942. Systematics and the origin of species, from the viewpoint of a zoologist. Cambridge (MA): Harvard University Press.

Mayr E. 1944. The birds of Timor and Sumba. Bull. Am Mus Nat Hist. 83:123–194.

Mayr E, Diamond JM. 1976. Birds on islands in the sky: origin of the montane avifauna of northern Melanesia. Proc Natl Acad Sci U S A. 73:1765–1769.

Mayr E, Diamond JM. 2001. The Birds of Northern Melanesia: Speciation, Ecology and Biogeography. New York (NY): Oxford University Press.

McKenna A, Hanna M, Banks E, Sivachenko A, Cibulskis K, Kernytsky A, Garimella K, Altshuler D, Gabriel S, Daly M, et al. 2010. The Genome Analysis Toolkit: a MapReduce framework for analyzing next-generation DNA sequencing data. Genome Res. 20:1297–1303.

Meisner J, Albrechtsen A. 2018. Inferring population structure and admixture proportions in low-depth NGS data. Genetics. 210:719–731.

Meyer M, Arsuaga JL, de Filippo C, Nagel S, Aximu-Petri A, Nickel B, Martínez I, Gracia A, de Castro JM, Carbonell E, et al. 2016. Nuclear DNA sequences from the Middle Pleistocene Sima de los Huesos hominins. Nature. 531:504–507.

Meyer M, Kircher M. 2010. Illumina sequencing library preparation for highly multiplexed target capture and sequencing. Cold Spring Harbor Protocols. doi:10.1101/pdb.prot5448.

Moyle RG, Filardi CE, Smith CE, Diamond J. 2009. Explosive Pleistocene diversification and hemispheric expansion of a “great speciator”. Proc Natl Acad Sci U S A. 106:1863–1868.

Moyle RG, Hosner PA, Jones AW, Outlaw DC. 2015. Phylogeny and biogeography of Ficedula flycatchers (Aves: Muscicapidae): novel results from fresh source material. Mol Phylogenet Evol. 82:87–94.

Nadachowska-Brzyska K, Li C, Smeds L, Zhang G, Ellegren H. 2015. Temporal dynamics of avian populations during Pleistocene revealed by whole-genome sequences. Curr Biol. 25:1375–1380.

Nadachowska-Brzyska K, Burri R, Smeds L, Ellegren H. 2016. PSMC analysis of effective population sizes in molecular ecology and its application to black-and-white Ficedula flycatchers. Mol Ecol. 25:1058–72.

Nielsen R, Korneliussen T, Albrechtsen A, Li Y, Wang J. 2012. SNP calling, genotype calling, and sample allele frequency estimation from new-generation sequencing data. PloS One. 7:e37558.

Nursyifa C, Bruniche-Olsen A, Garcia-Erill G, Heller R, Albrechtsen A. 2021. Joint identification of sex and sex-linked scaffolds in non-model organisms using low depth sequencing data. bioRxiv. 433779.

Nylander JA, Olsson U, Alström P, Sanmartín I. 2008. Accounting for phylogenetic uncertainty in biogeography: a Bayesian approach to dispersal-vicariance analysis of the thrushes (Aves: Turdus). Syst Biol. 57:257–268.

Omland KE. 1997. Examining two standard assumptions of ancestral reconstructions: repeated loss of dichromatism in dabbling ducks (Anatini). Evolution. 51:1636–1646.

Paradis E, Schliep K. 2019. ape 5.0: an environment for modern phylogenetics and evolutionary analyses in R. Bioinformatics (Oxf). 35:526–528.

Patterson N, Moorjani P, Luo Y, Mallick S, Rohland N, Zhan Y, Genschoreck T, Webster T, Reich D. 2012. Ancient admixture in human history. Genetics. 192:1065–1093.

Pedersen MP, Irestedt M, Joseph L, Rahbek C, Jønsson KA. 2018. Phylogeography of a ‘great speciator’(Aves: Edolisoma tenuirostre) reveals complex dispersal and diversification dynamics across the Indo-Pacific. J Biogeogr. 45:826–837.

Pepke ML, Irestedt M, Fjeldså J, Rahbek C, Jønsson KA. 2019. Reconciling supertramps, great speciators and relict species with the taxon cycle stages of a large island radiation (Aves: Campephagidae). J Biogeogr. 46:1214–1225.

Peterson AT. 2007. Geographic variation in size and coloration in the Turdus poliocephalus complex: a first review of species limits. Sci Pap Nat Hist Mus Univ Kans. 40:1–17.

Petersen KR, Streett DA, Gerritsen AT, Hunter SS, Settles ML. 2015, September. Super deduper, fast PCR duplicate detection in fastq files. In: Proceedings of the 6th ACM Conference on Bioinformatics, Computational Biology and Health Informatics; 2015 Sep 9–12; Atlanta. New York (NY): Association for Computing Machinery. p. 491–492.

Pratt HD, Bruner PL, Berrett DG. 1987. A field guide to the birds of Hawaii and the tropical Pacific. Princeton (NJ): Princeton University Press.

Pujolar JM, Blom MP, Reeve AH, Kennedy JD, Marki PZ, Korneliussen TS, Freeman BG, Sam K, Linck E, Haryoko T, et al. 2022. The formation of avian montane diversity across barriers and along elevational gradients. Nat Commun. 13:1–13.

R Core Team. 2018. R: A language and environment for statistical computing. Vienna: R Foundation for Statistical Computing. Available from: https://www.R-project.org.

Ramachandran S, Deshpande O, Roseman CC, Rosenberg NA, Feldman MW, Cavalli-Sforza LL. 2005. Support from the relationship of genetic and geographic distance in human populations for a serial founder effect originating in Africa. Proc Natl Acad Sci U S A. 102:15942–15947.

Rambaut A, Drummond AJ. 2015. TreeAnnotator v1.8.2: MCMC Output analysis. Available from: http://beast.bio.ed.ac.uk

Rambaut A, Suchard MA, Xie D, Drummond AJ. 2014. Tracer v1.6. Available from: http://beast.bio.ed.ac.uk

Reeve AH, Blom MPK, Marki PZ, Batista R, Olsson U, Edmark VN, Irestedt M, Jønsson KA. 2022. The Sulawesi Thrush (Cataponera turdoides; Aves: Passeriformes) belongs to the genus Turdus. Zool Scr. 51:32–40.

Rensch B, Heberer G, Lehmann W. 1930. Eine biologische reise nach den Kleinen Sunda-Inseln. Berlin: Gebrüder Borntraeger.

Reynolds J, Weir BS, Cockerham CC. 1983. Estimation of the coancestry coefficient: basis for a short-term genetic distance. Genetics. 105:767–779.

Rheindt FE, Prawiradilaga DM, Ashari H, Gwee CY, Lee GW, Wu MY, Ng, NS. 2020. A lost world in Wallacea: Description of a montane archipelagic avifauna. Science. 367:167–170.

Ricklefs RE, Cox GW. 1978. Stage of taxon cycle, habitat distribution, and population density in the avifauna of the West Indies. Am Nat. 112:875–895.

Rolland J, Jiguet F, Jønsson KA, Condamine, FL, and Morlon, H. 2014. Settling down of seasonal migrants promotes bird diversification. Proc R Soc Lond B Biol Sci. 281:20140473.

Saitou N, Nei M. 1987. The neighbor-joining method: a new method for reconstructing phylogenetic trees. Mol Biol Evol. 4:406–425.

Simão FA, Waterhouse RM, Ioannidis P, Kriventseva EV, Zdobnov EM. 2015. BUSCO: assessing genome assembly and annotation completeness with single-copy orthologs. Bioinformatics (Oxf). 31:3210–3212.

Skutch AF. 1985. Clutch size, nesting success, and predation on nests of Neotropical birds, reviewed. Ornithol Monogr. 36:575–594.

Slatkin M. 1987. Gene flow and the geographic structure of natural populations. Science. 236:787–792.

Slatkin M. 1993. Isolation by distance in equilibrium and non-equilibrium populations. Evolution. 47:264–279.

Stamatakis A. 2014. RAxML version 8: a tool for phylogenetic analysis and post-analysis of large phylogenies. Bioinformatics (Oxf). 30:1312–1313.

Stresemann E. 1939. Die Vögel von Celebes. J Ornithol. 87:299–425.

Vieira FG, Lassalle F, Korneliussen TS, Fumagalli M. 2016. Improving the estimation of genetic distances from Next-Generation Sequencing data. Biol J Linn Soc. 117:139–149.

Villard P, Duval T, Papineau C, Cassan JJ, Fuchs J. 2019. Notes on the biology of the threatened Island Thrush Turdus poliocephalus xanthopus in New Caledonia. Bird Conserv Int. 29:616–626.

Voelker G, Rohwer S, Bowie RC, Outlaw DC. 2007. Molecular systematics of a speciose, cosmopolitan songbird genus: defining the limits of, and relationships among, the Turdus thrushes. Mol Phylogenet Evol. 42:422–434.

Voris HK. 2000. Maps of Pleistocene sea levels in Southeast Asia: shorelines, river systems and time durations. J Biogeogr. 27:1153–1167.

Wallace AR. 1869. The Malay Archipelago: the land of the orang-utan and the bird of paradise; a narrative of travel, with studies of man and nature. London (GB): Macmillan.

Weisenfeld NI, Kumar V, Shah P, Church DM, Jaffe DB. 2017. Direct determination of diploid genome sequences. Genome Res. 27:757–767.

Zhang J, Kobert K, Flouri T, Stamatakis A. 2014. PEAR: a fast and accurate Illumina Paired-End reAd mergeR. Bioinformatics (Oxf). 30:614–620.

