## Supplementary File 2 for "Population genomics of the island thrush elucidates one of earth’s great archipelagic radiations"

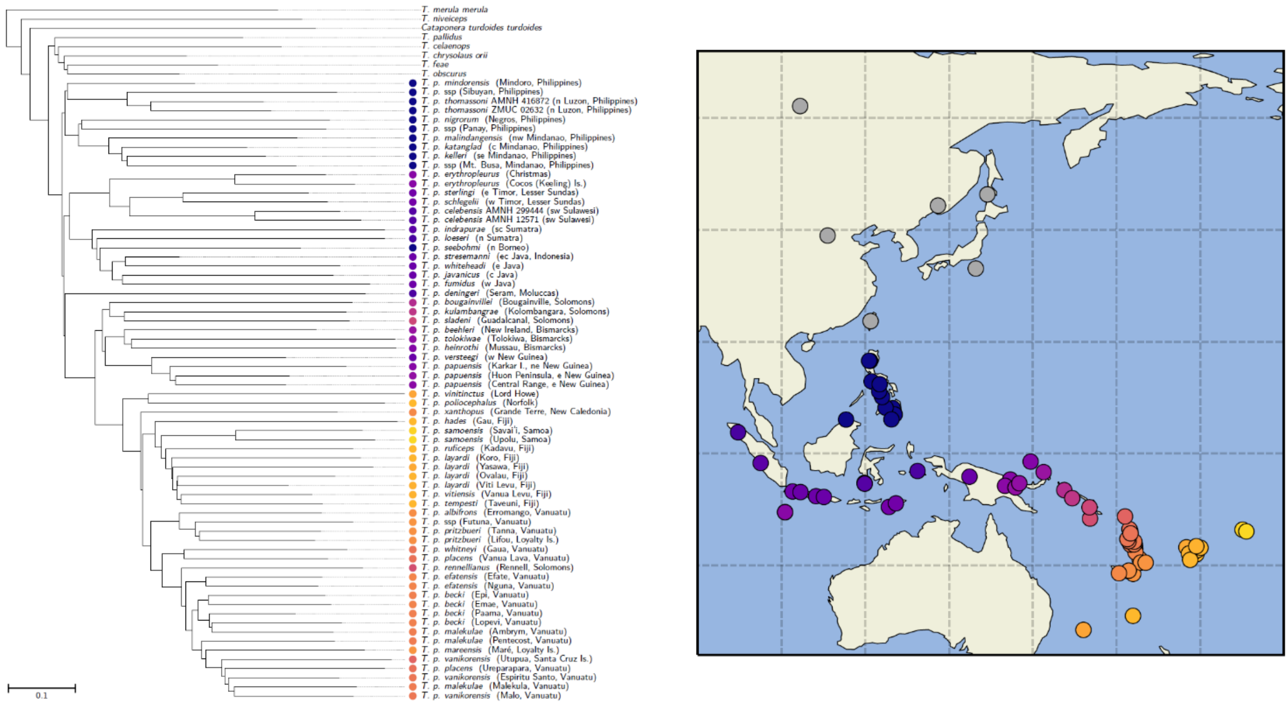


**Fig. S1.** Phylogeny estimated from pairwise $F_{ST}$ using neighbor-joining followed by subtree pruning and regrafting. Leaf node colors on the tree match those used on the map. Island thrush individuals are colored by distance to a reference point (30° N, 120° E,), while outgroups are shown in gray. The *Turdus merula* individual (not shown) is from Sweden.

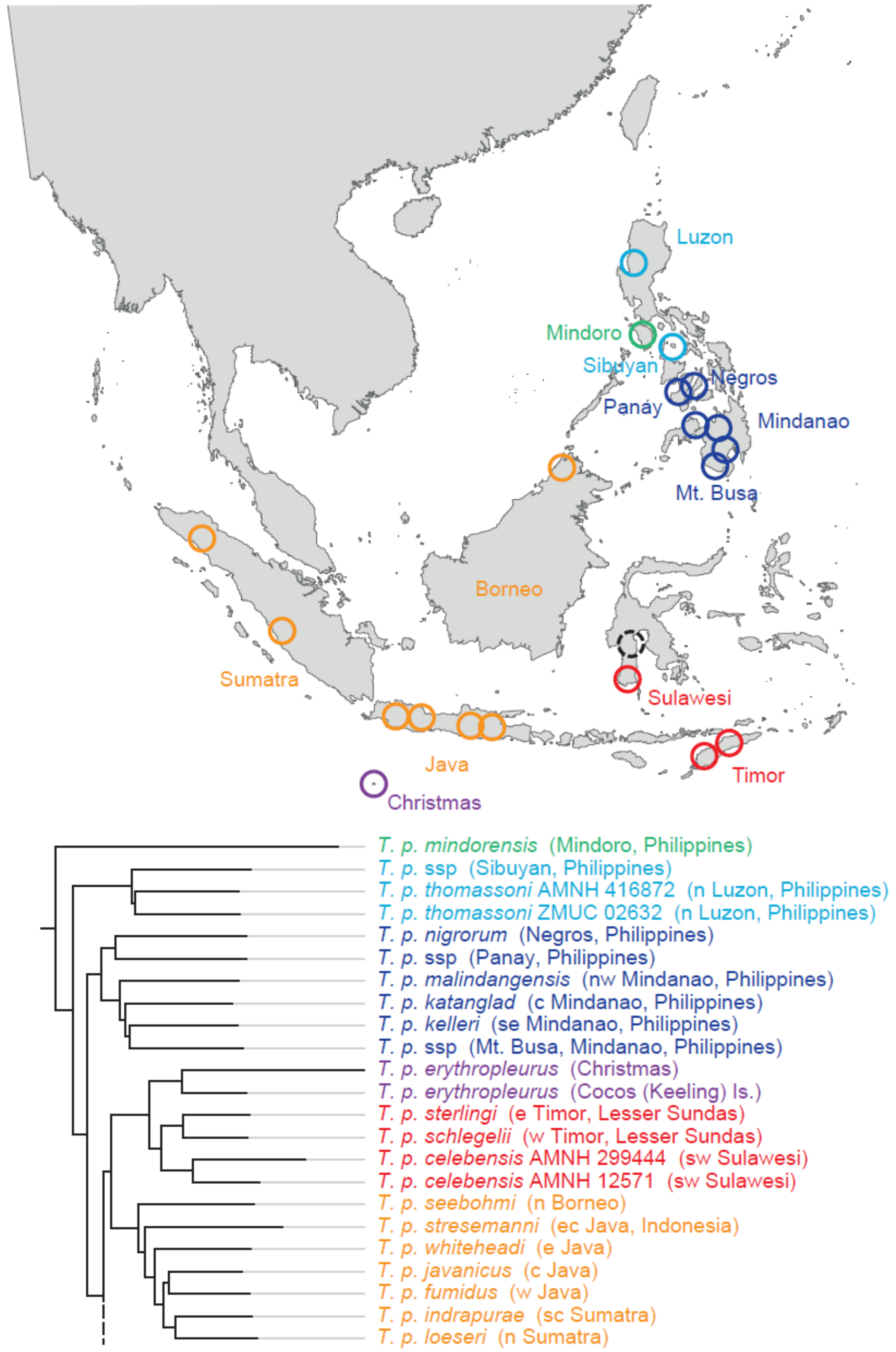


**Fig. S2a.** Geographic distribution and phylogenetic relationships of western island thrush populations. The tree is from Fig.1. Colored circles indicate sampling localities. Clades are color coded to aid interpretation. The extinct Cocos (Keeling) Is. population, introduced from Christmas I., is not shown. The black dashed circle indicates the sampling locality of *T. p. hygroscopus* (south Sulawesi), which was included in the mitogenome tree (Fig. S3), but not the pairwise distance tree (Fig. 1).

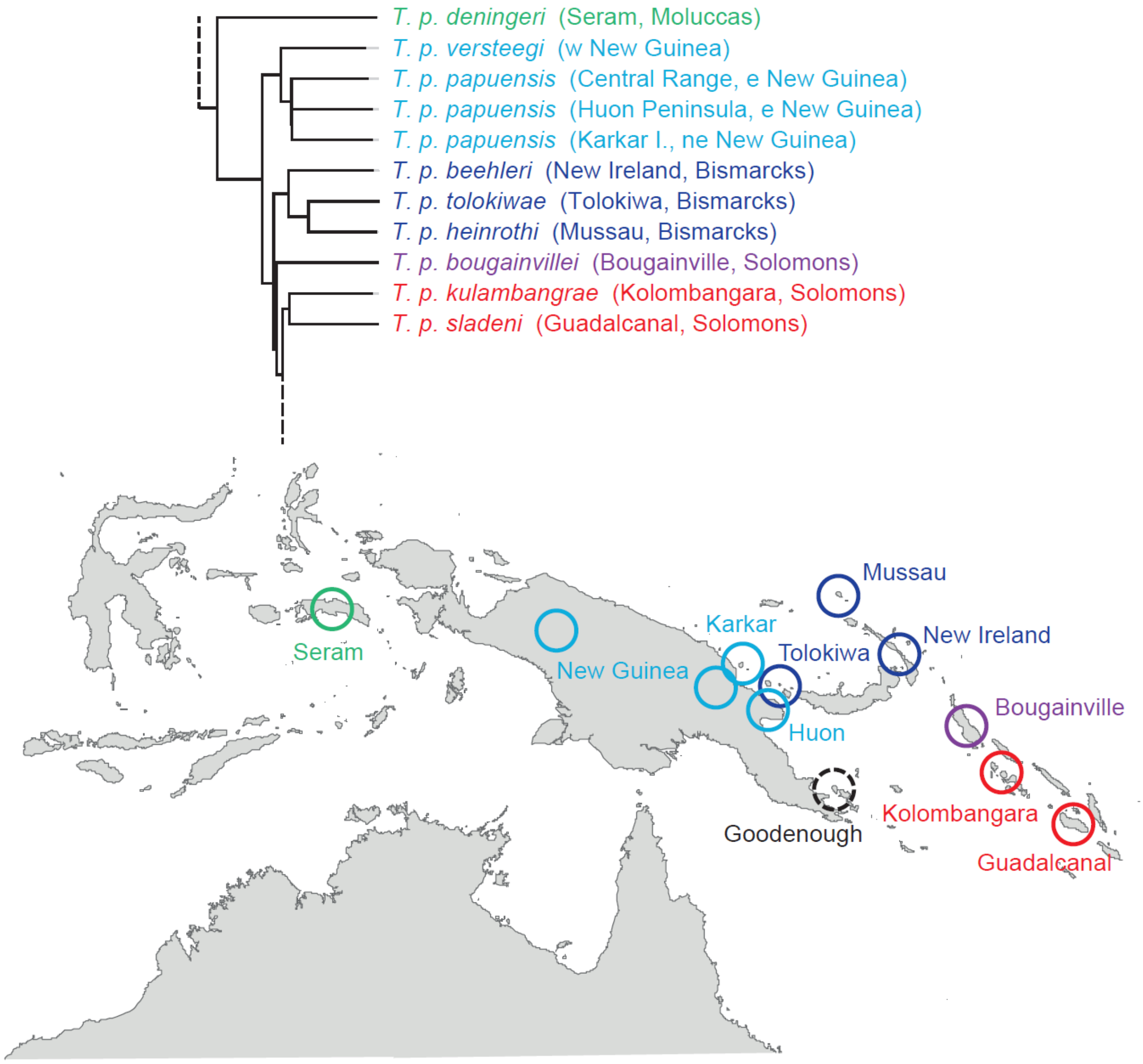


**Fig. S2b.** Geographic distribution and phylogenetic relationships of island thrush populations on New Guinea and nearby islands. The tree is from Fig.1. Colored circles indicate sampling localities. Clades are color coded to aid interpretation. The black dashed circle indicates the sampling locality of *T. p. canescens* (Goodenough, D'Entrecasteaux Is.), which was included in the mitogenome tree (Fig. S5), but not the pairwise distance tree (Fig. 1).

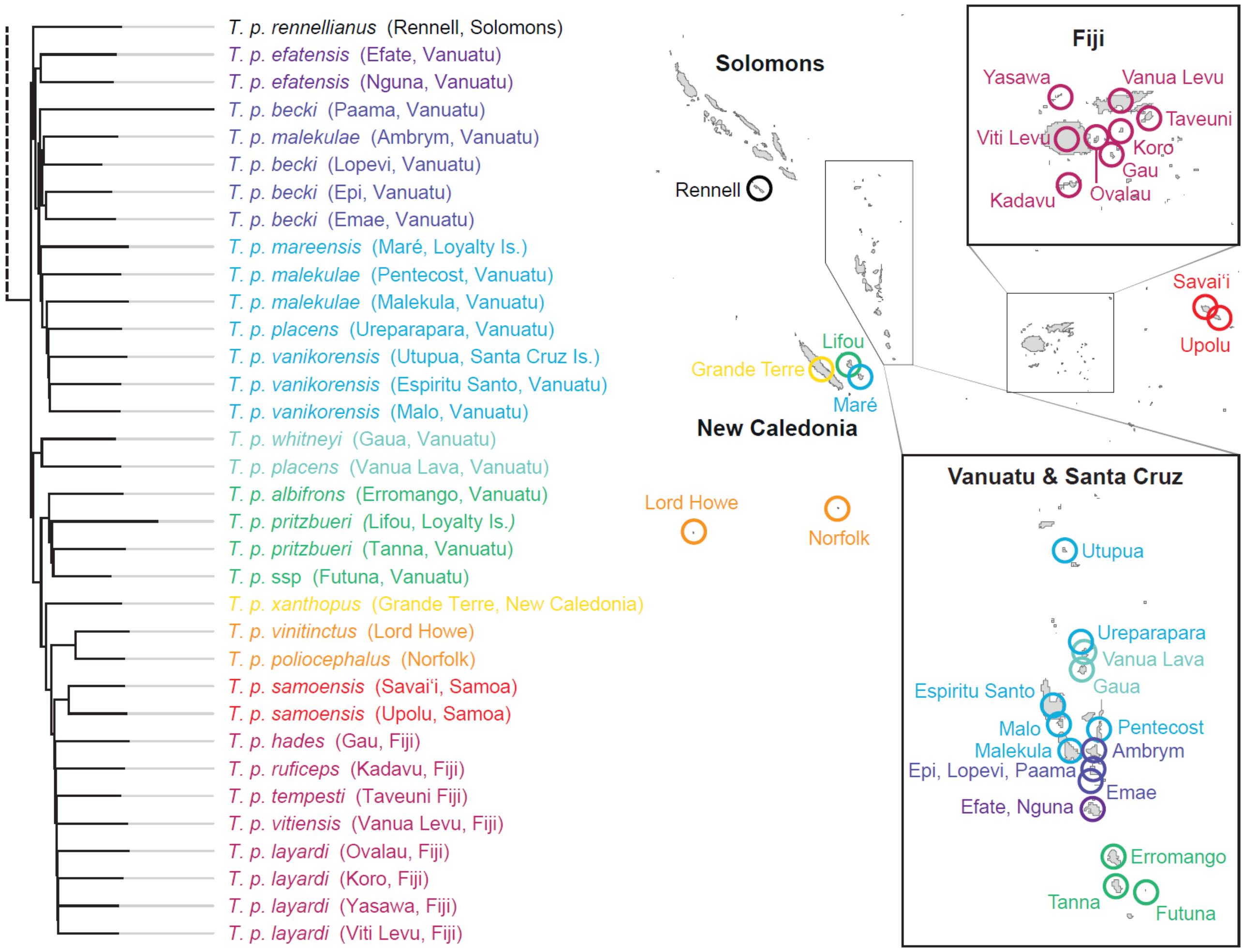


**Fig. S2c.** Geographic distribution and phylogenetic relationships of eastern island thrush populations. The tree is from Fig.1. Colored circles indicate sampling localities. Clades are color coded to aid interpretation.

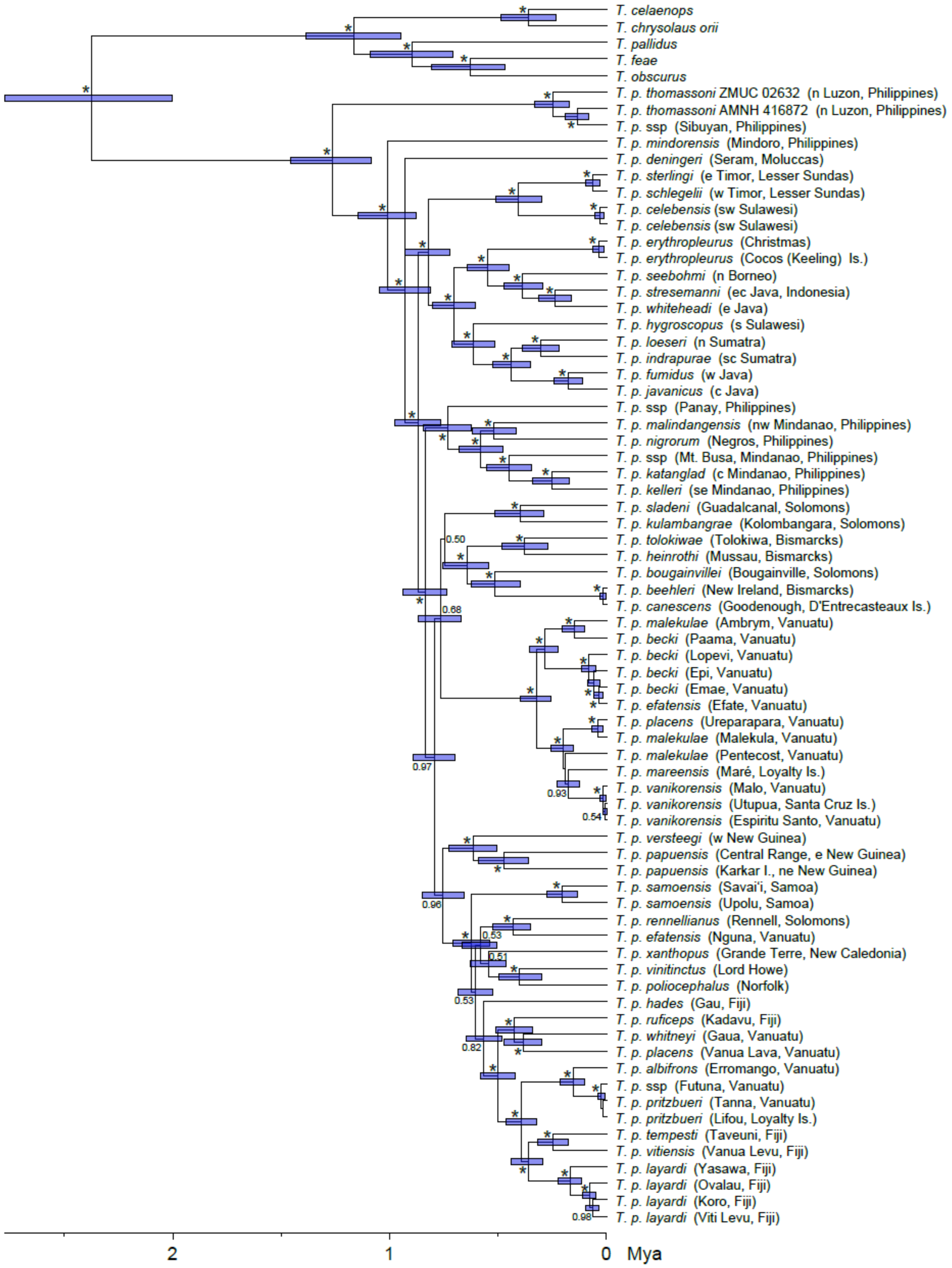


**Fig. S3.** Time-calibrated Bayesian consensus tree from analysis of mitochondrial genomes from the island thrush and its sister clade. Posterior probabilities ≥ 0.50 are indicated at the nodes; asterisks denote nodes with PP ≥ 0.99. Error bars indicate 95% highest posterior density (HPD) intervals. Outgroup taxa are not shown.

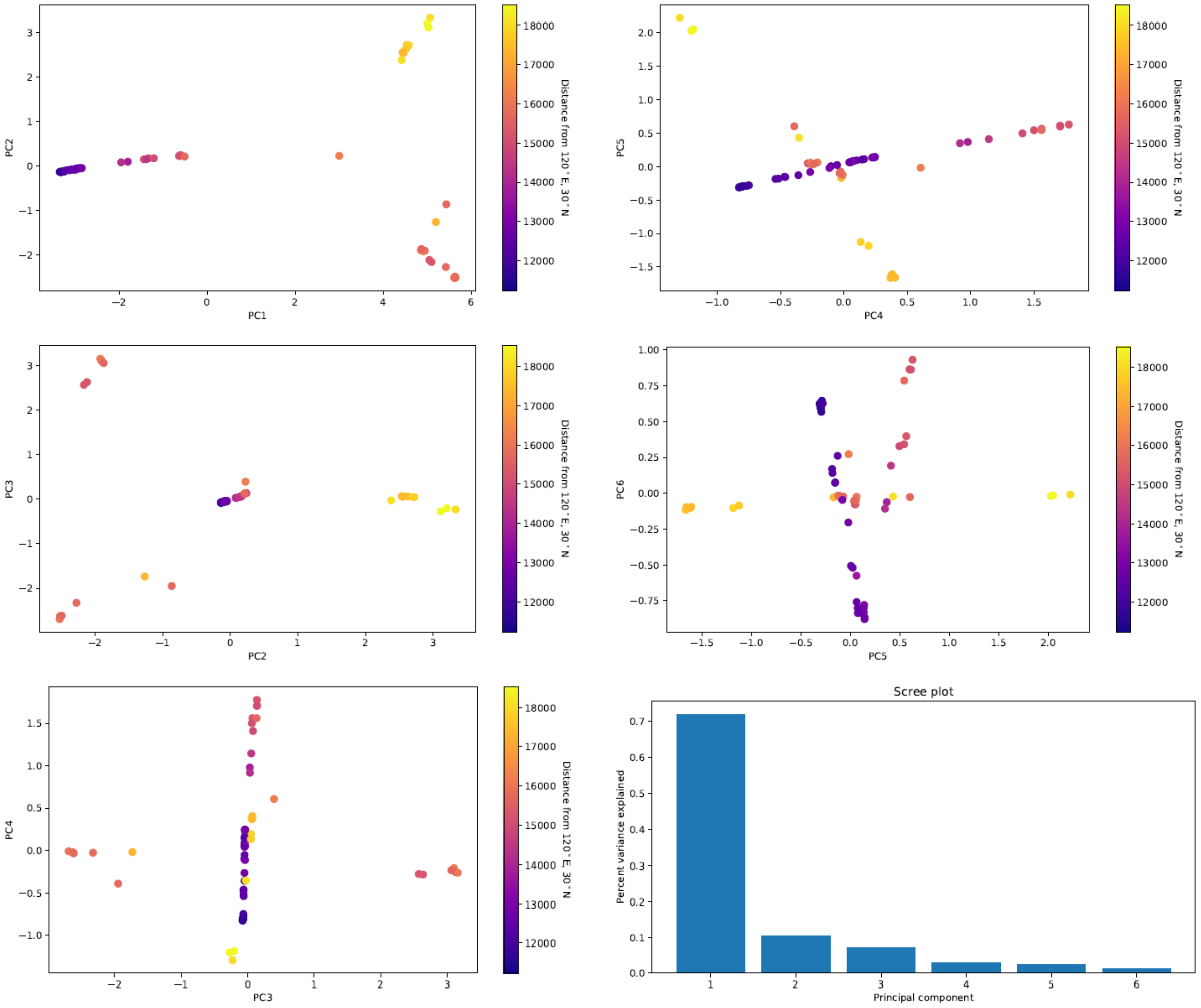


**Fig. S4.** PCA plots of island thrush individuals, with covariance matrix estimated from genotype likelihoods using PCAngsd. Colors indicate the geographic distance of an individual to a reference point (120˚E, 30˚N).

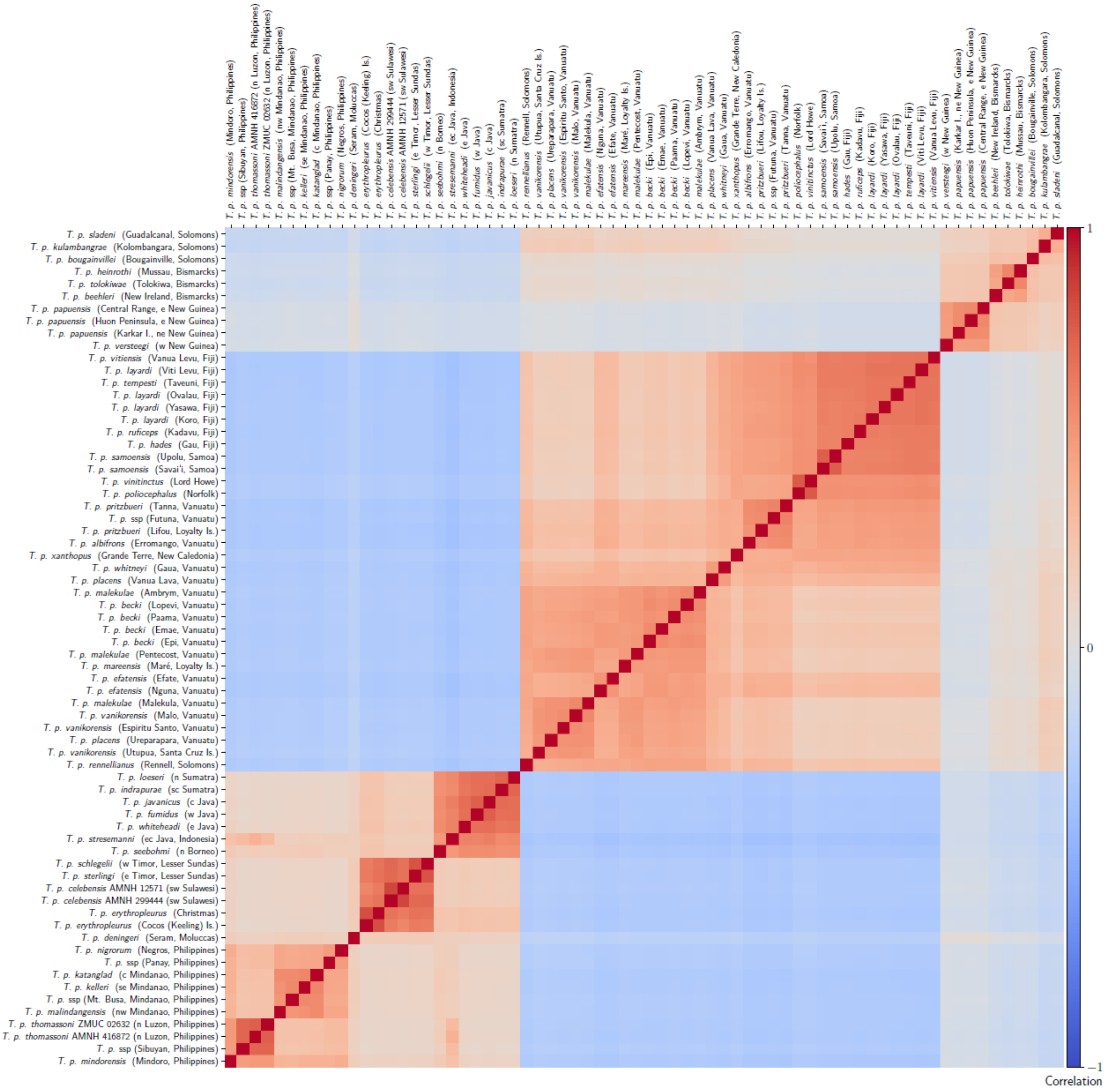


**Fig. S5.** Heatmap of the correlation matrix calculated from the PCAngsd covariance matrix.

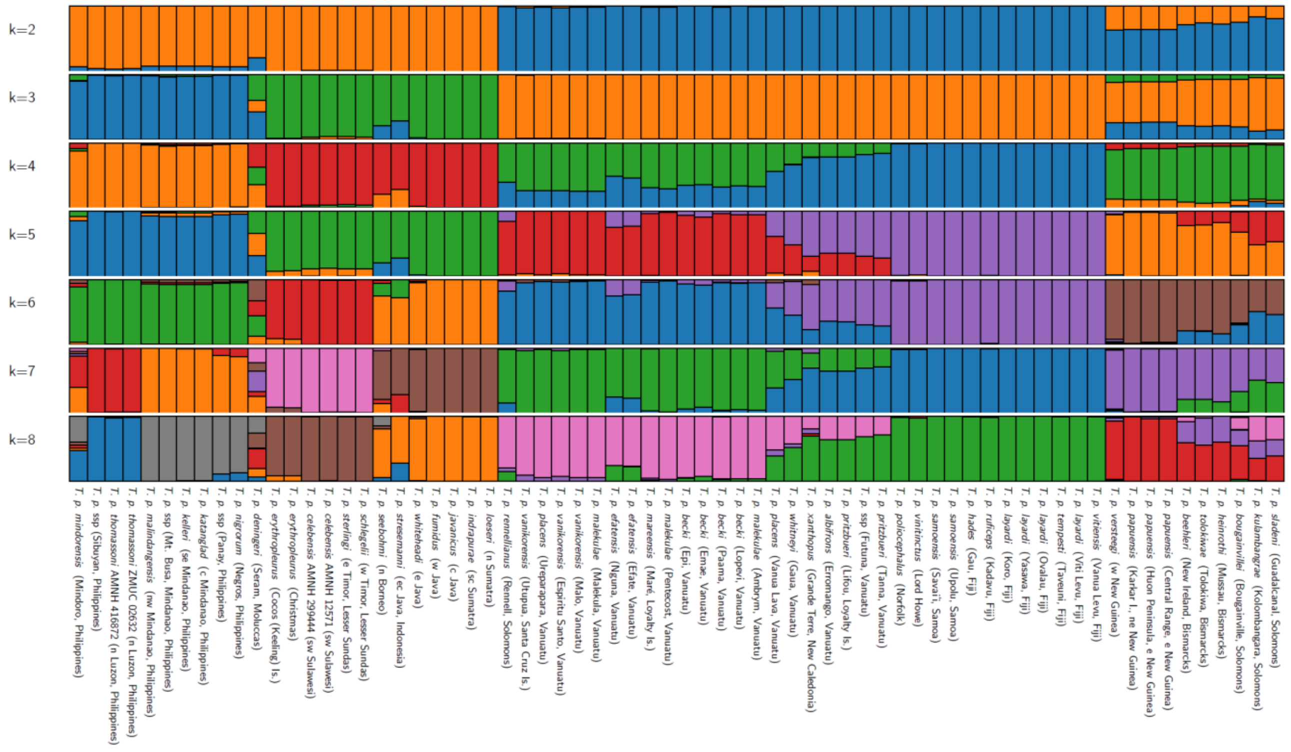


**Fig. S6.** Admixture output from PCAngsd -admix, for k=2 to k=8 clusters within the island thrush.

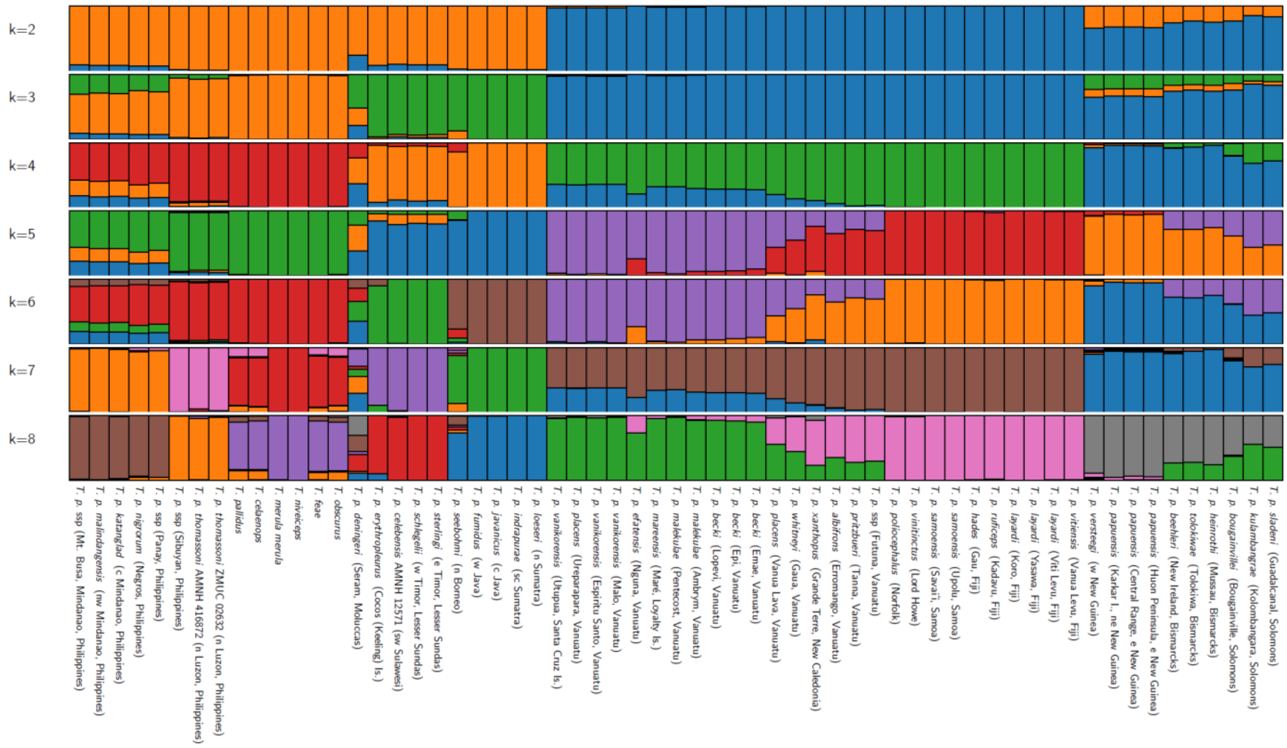


**Fig. S7**. Admixture output from PCAngsd -admix, for k=2 to k=8 clusters. Outgroups are included to illustrate the impact of ascertainment bias (see Fig. S6). Due to sampling imbalance, the analyses are oversensitive to differentiation within the eastern clades, and undersensitive to differentiation among the more basally-spitting branches of the tree (representing western groups and outgroups).

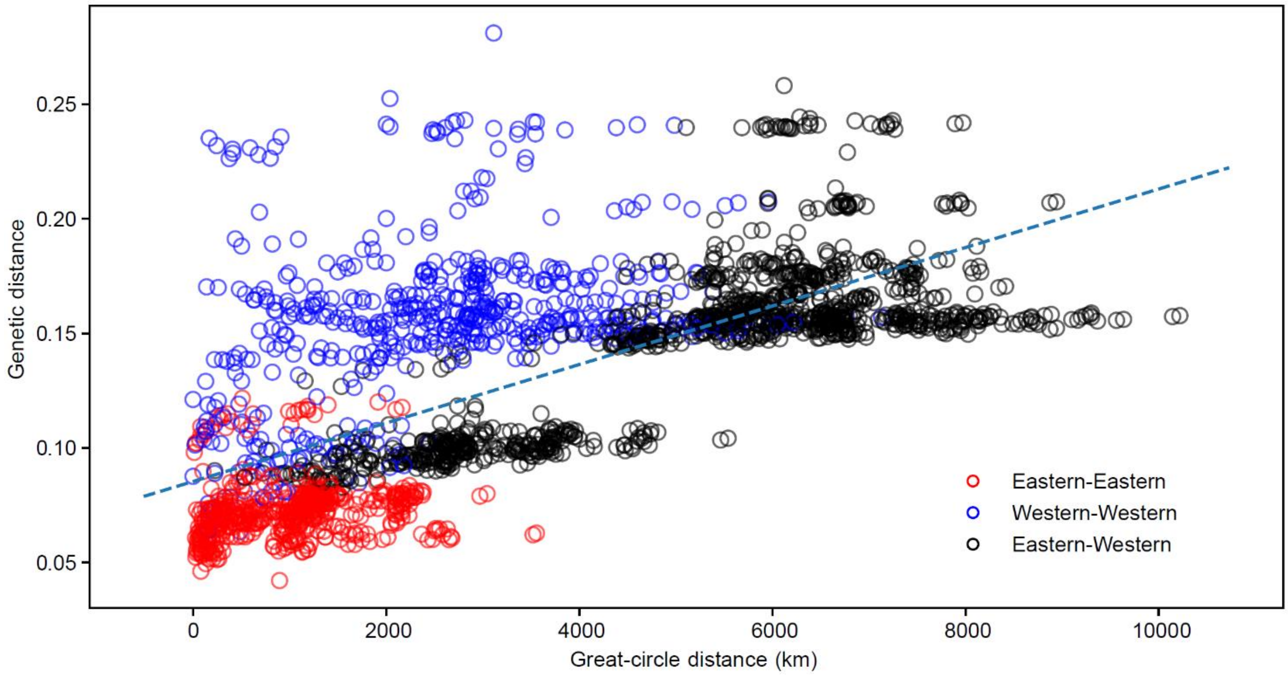


**Fig. S8.** Relationship between geographic distance (great-circle distance in km) and genetic distance (pairwise distance) in the island thrush. The recovered positive relationship (Mantel test: $r^2 = 0.47$, $p < .001$) supports a stepping stone colonization model. Blue points indicate comparisons between western populations (Clades A-J in Fig. 1); red points indicate comparisons between eastern populations (Clades K and L in Fig. 1). Some western populations show deep genetic divergence despite close proximity; eastern populations are genetically similar, but widespread.

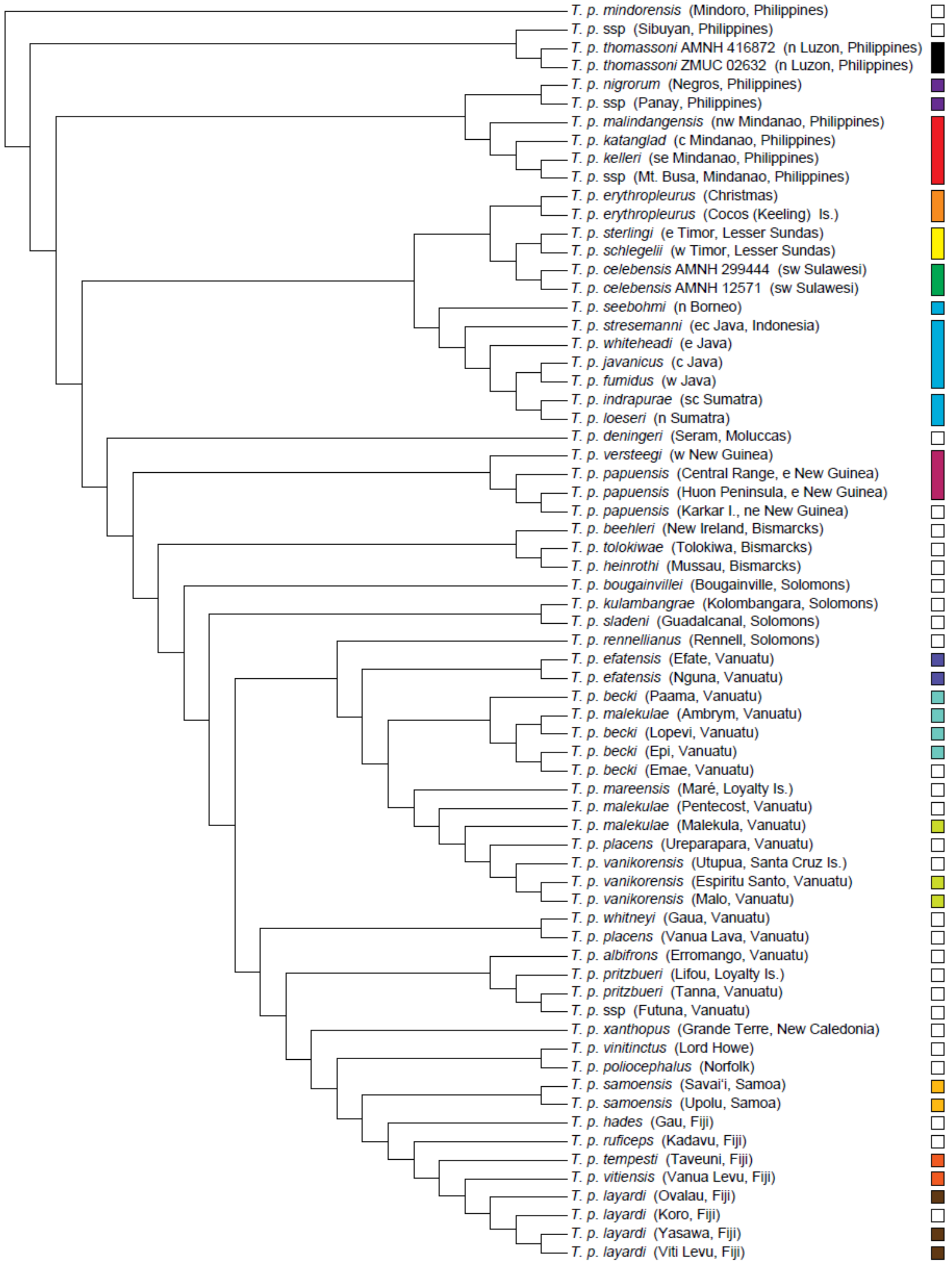


**Fig. S9.** Island thrush population connectivity. Tree topology matches that in Fig. 1. Similarly colored cells indicate populations inferred to have shared land connections during Pleistocene glacial periods. Merged cells indicate populations occuring on the same island at modern sea levels. White cells indicate populations never connected by land to any other sampled island thrush population. Close phylogenetic relationships are shared both by populations on the same island, and populations connected by Pleistocene land bridges. Land bridges assisted inter-island dispersal and colonization, but were not essential for the island thrush's spread across the Indo-Pacific.

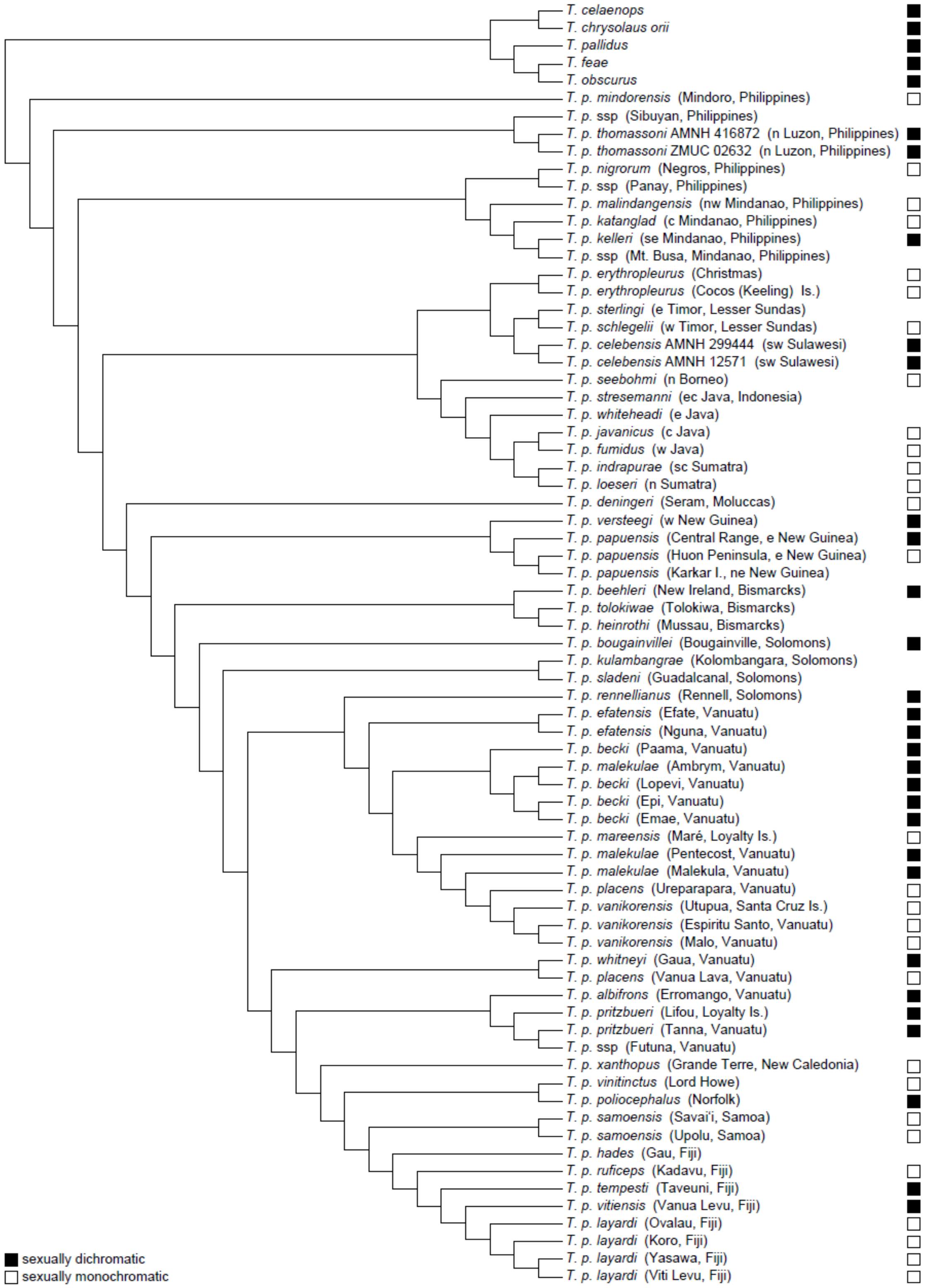


**Fig. S10.** Sexual dichromatism in the island thrush. Tree topology matches that in Fig. 1. Data is from Peterson (2007), but we additionally scored species from the five-species sister clade. Sexually dichromatic subspecies are indicated with black cells, and sexually monochromatic subspecies with white cells. Tips without cells represent subspecies not scored in the Peterson study. Sexual dichromatism was evidently lost and gained frequently across the radiation.

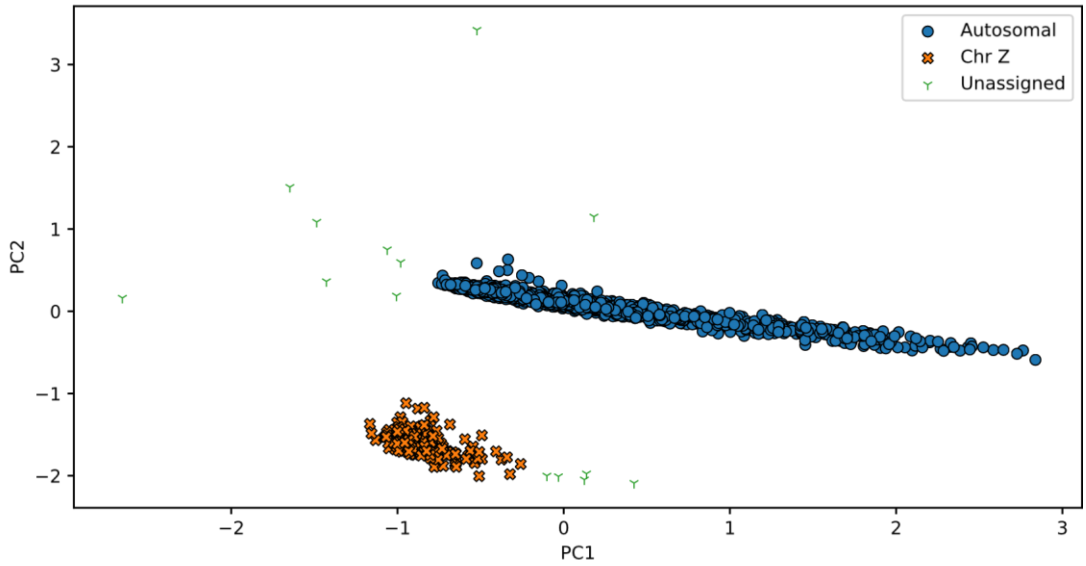


**Fig. S11.** PCA of contig read dosage, with clusters identified using DBSCAN.

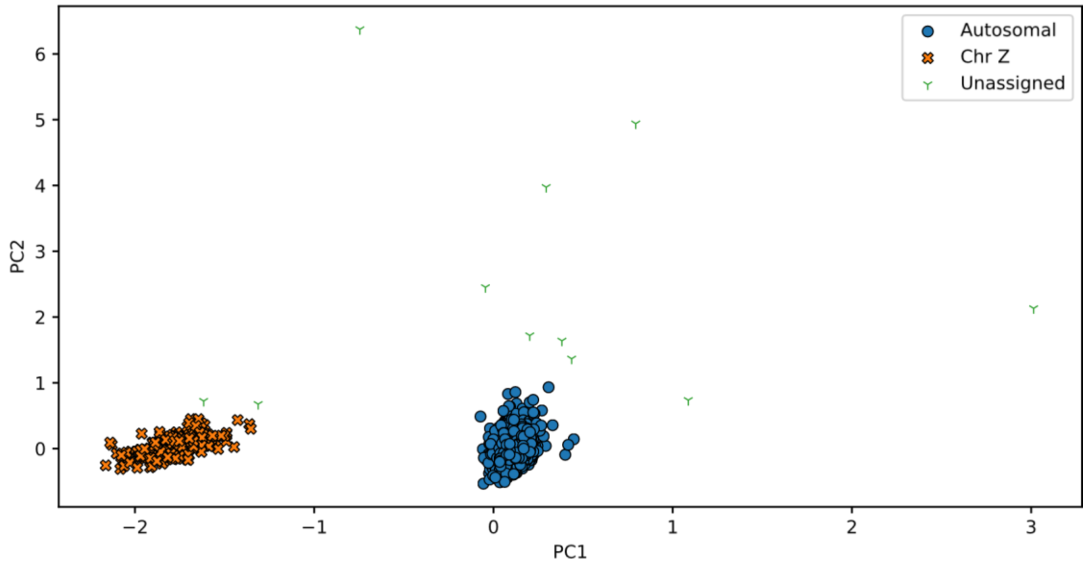


**Fig. S12.** GC-corrected PCA of contig read dosage, with clusters identified using DBSCAN.

**Table S1.** Assembly statistics for the *de novo Turdus merula* genome assembly, as calculated with Supernova, QUAST, and BUSCO.

| **SUPERNOVA** | | | | |
|---|---|---|---|---|
| Assembly length (Mbp) | %Missing >10Kb | Scaffold N50 (Kbp) | N Reads (M) | Coverage |
| 905.5 | 17.8 | 123.7 | 317.1 | 22.4x |
| **QUAST** | | | | |
| Assembly length (Mbp) | N50(Kbp) | Largest contig (Kpb) | | |
| 1067.2 | 111.1 | 2101.0 | | |
| **BUSCO** | | | | |
| %Single-copy | %Duplicated | %Fragmented | %Missing | |
| 68.0 | 3.3 | 17.5 | 11.2 | |

**Table S2.** Composition of Z-linked and autosomal contigs, as identified from the GC-corrected PCA. Only contigs with length > 100 kbp were considered for categorisation as Z-linked or autosomal.

| **Category** | **Number of contigs** | **Nucleotides** |
|---|---:|---:|
| Reference | 53,570 | 1,067,231,262 |
| length > 100 kbp | 3,110 | 576,091,170 |
| Autosomal | 2,905 | 543,947,081 |
| Z-linked | 194 | 30,354,447 |
| Unassigned | 11 | 1,789,642 |

### REFERENCES

Peterson A.T. 2007. Geographic variation in size and coloration in the *Turdus poliocephalus* complex: a first review of species limits. *Sci Pap Nat Hist Mus Univ Kans*. 40:1–17.
