## Supplementary File 3 for "Population genomics of the island thrush elucidates one of earth’s great archipelagic radiations"

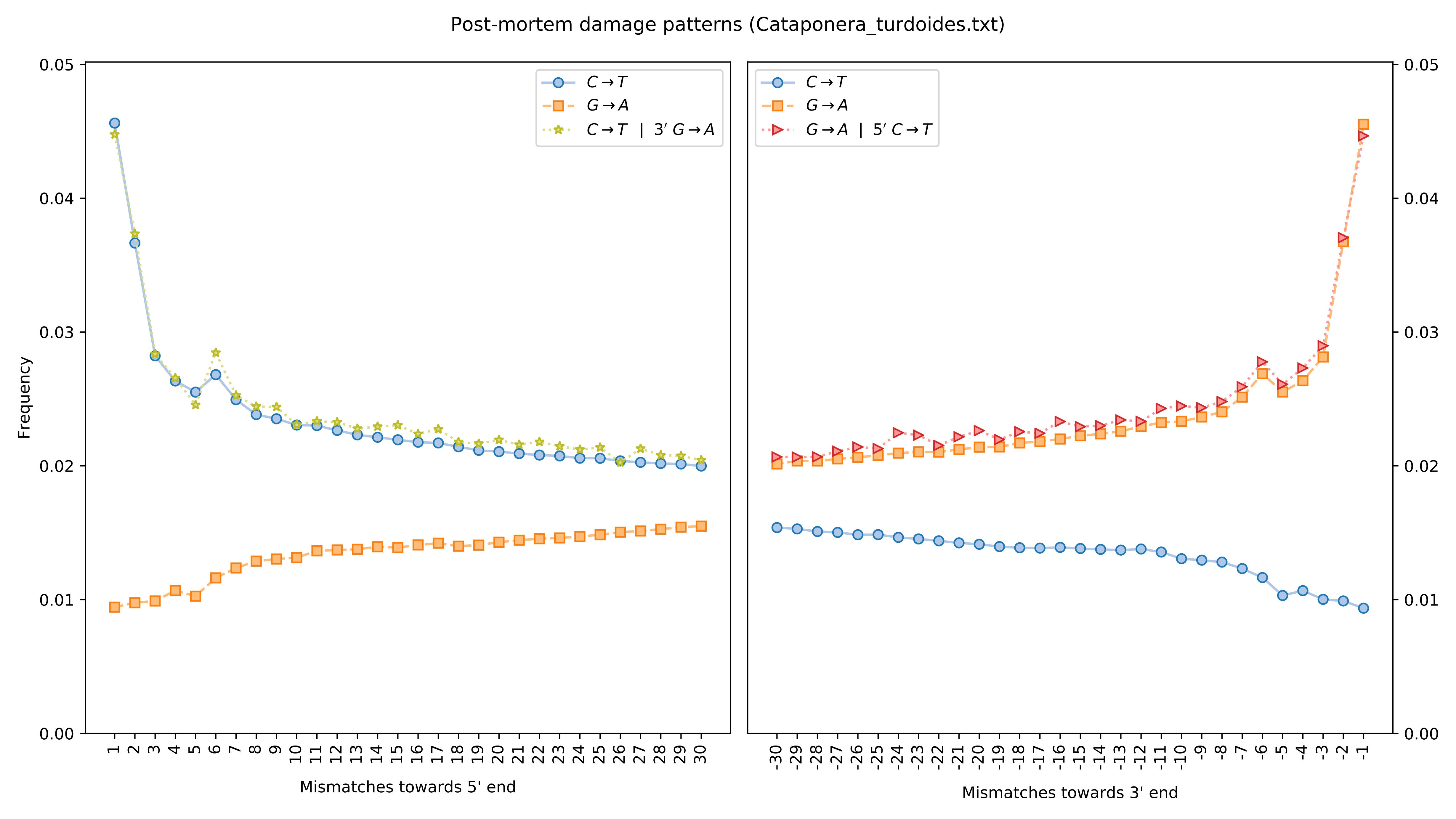

Post-mortem damage patterns (Cataponera_turdoides.txt)
Frequency
Mismatches towards 5' end
Mismatches towards 3' end
C→T
G→A
C→T | 3′ G→A
G→A | 5′ C→T

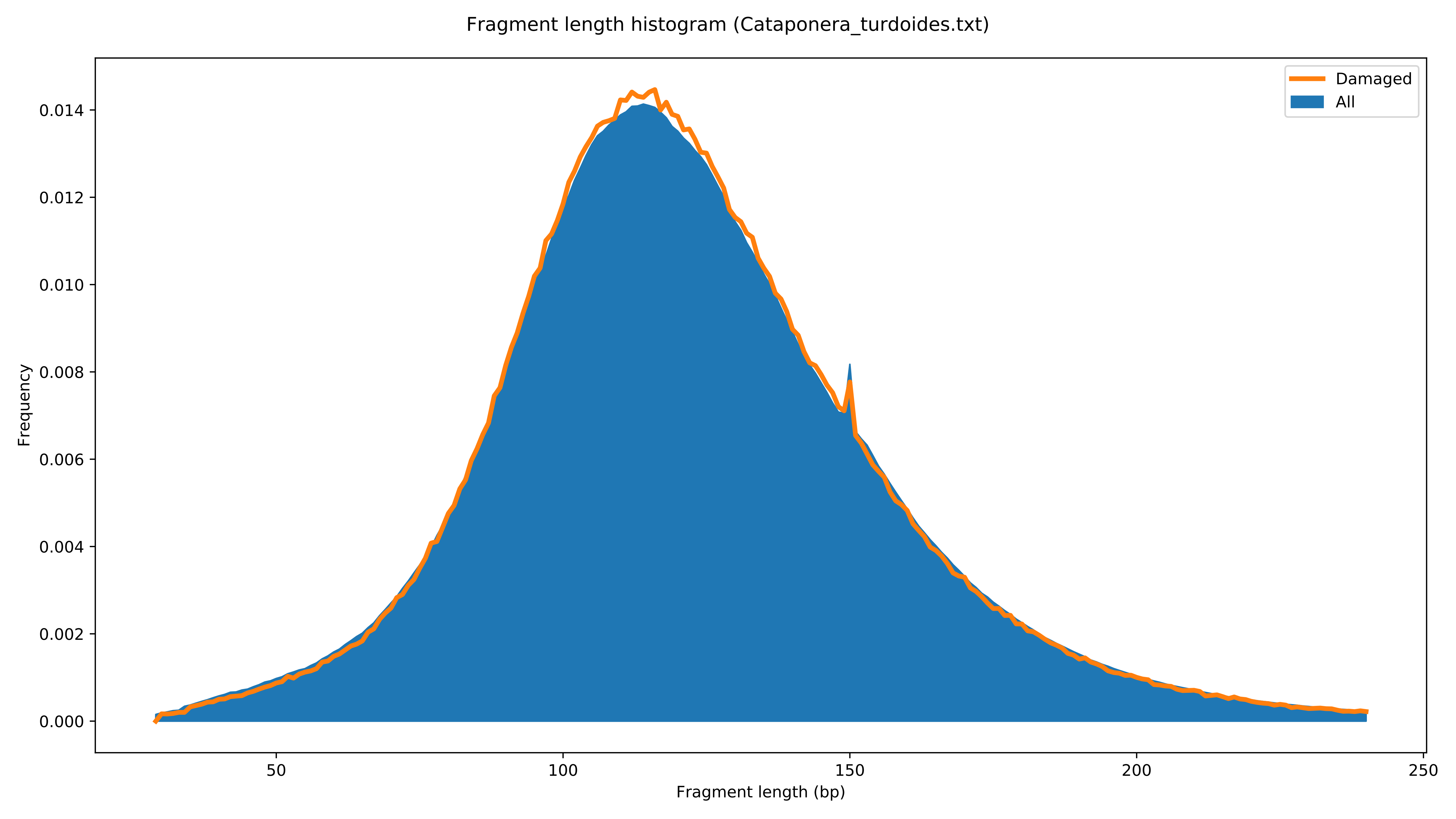

Fragment length histogram (Cataponera_turdoides.txt)
Damaged
All
Frequency
0.000
0.002
0.004
0.006
0.008
0.010
0.012
0.014
50
100
150
200
250
Fragment length (bp)

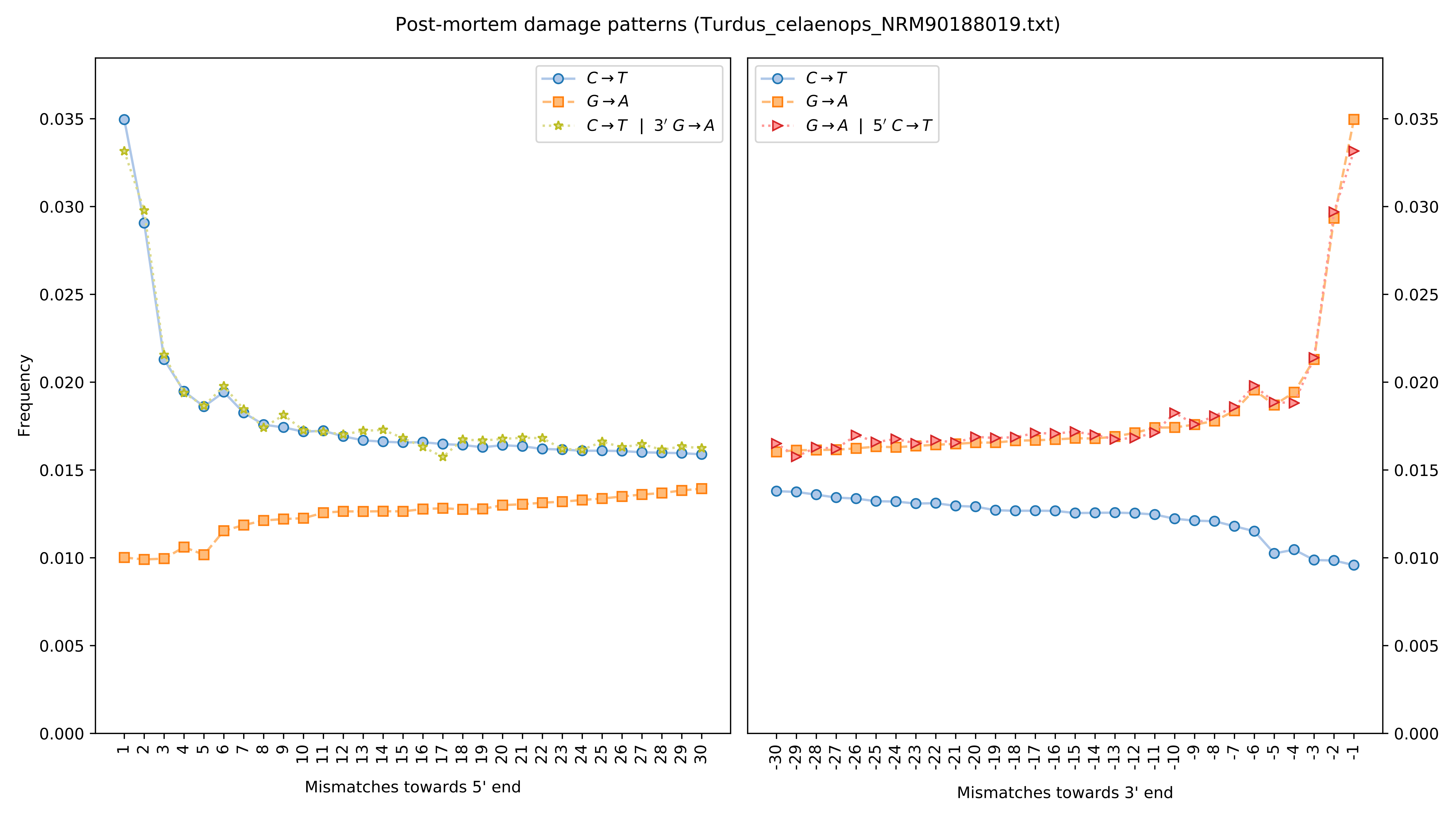

Post-mortem damage patterns (Turdus_celaenops_NRM90188019.txt)
C→T
G→A
C→T | 3′ G→A
G→A | 5′ C→T
Frequency
Mismatches towards 5' end
Mismatches towards 3' end

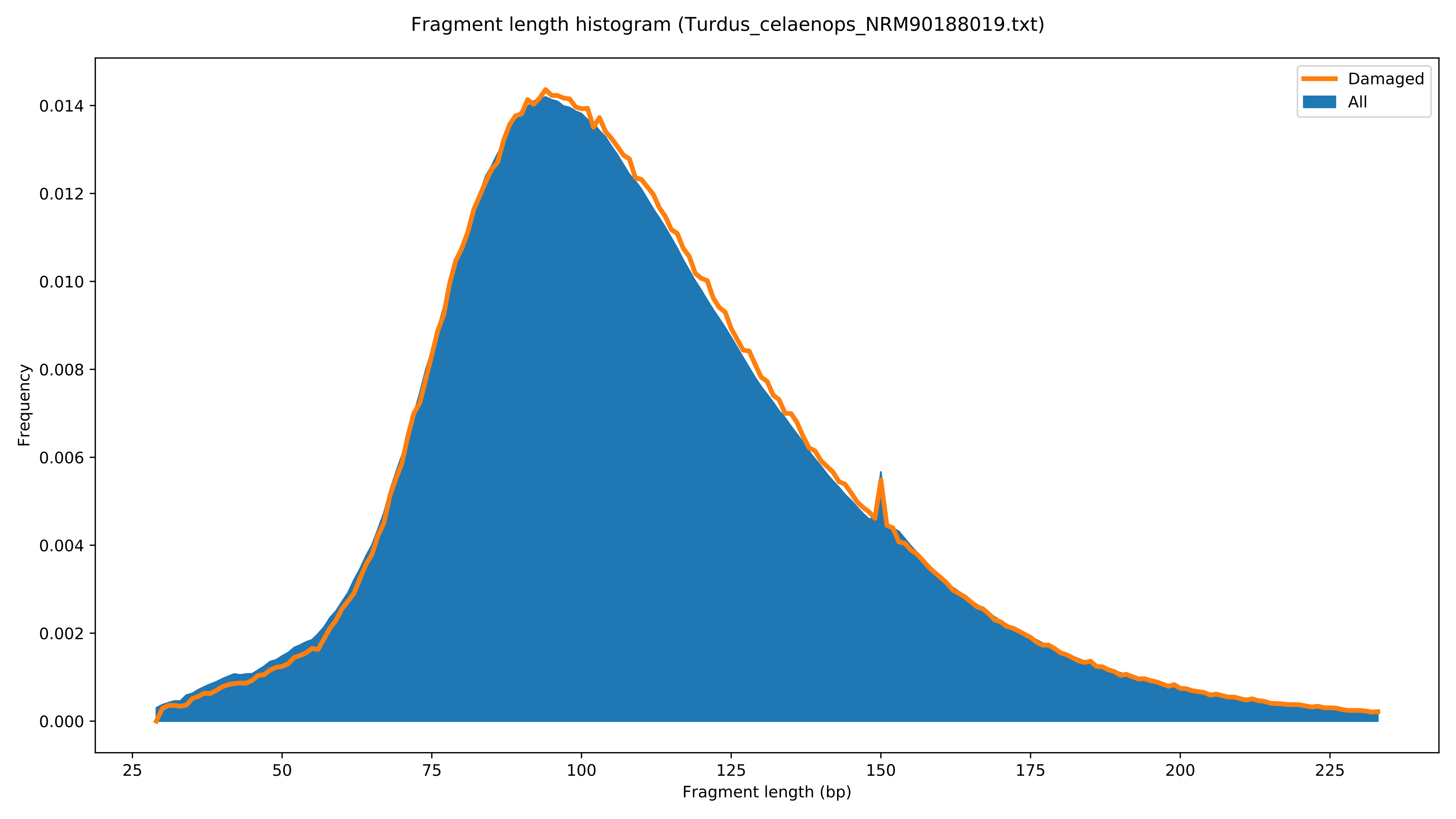
Fragment length histogram (Turdus_celaenops_NRM90188019.txt)
Damaged
All
Frequency
0.000
0.002
0.004
0.006
0.008
0.010
0.012
0.014
25
50
75
100
125
150
175
200
225
Fragment length (bp)

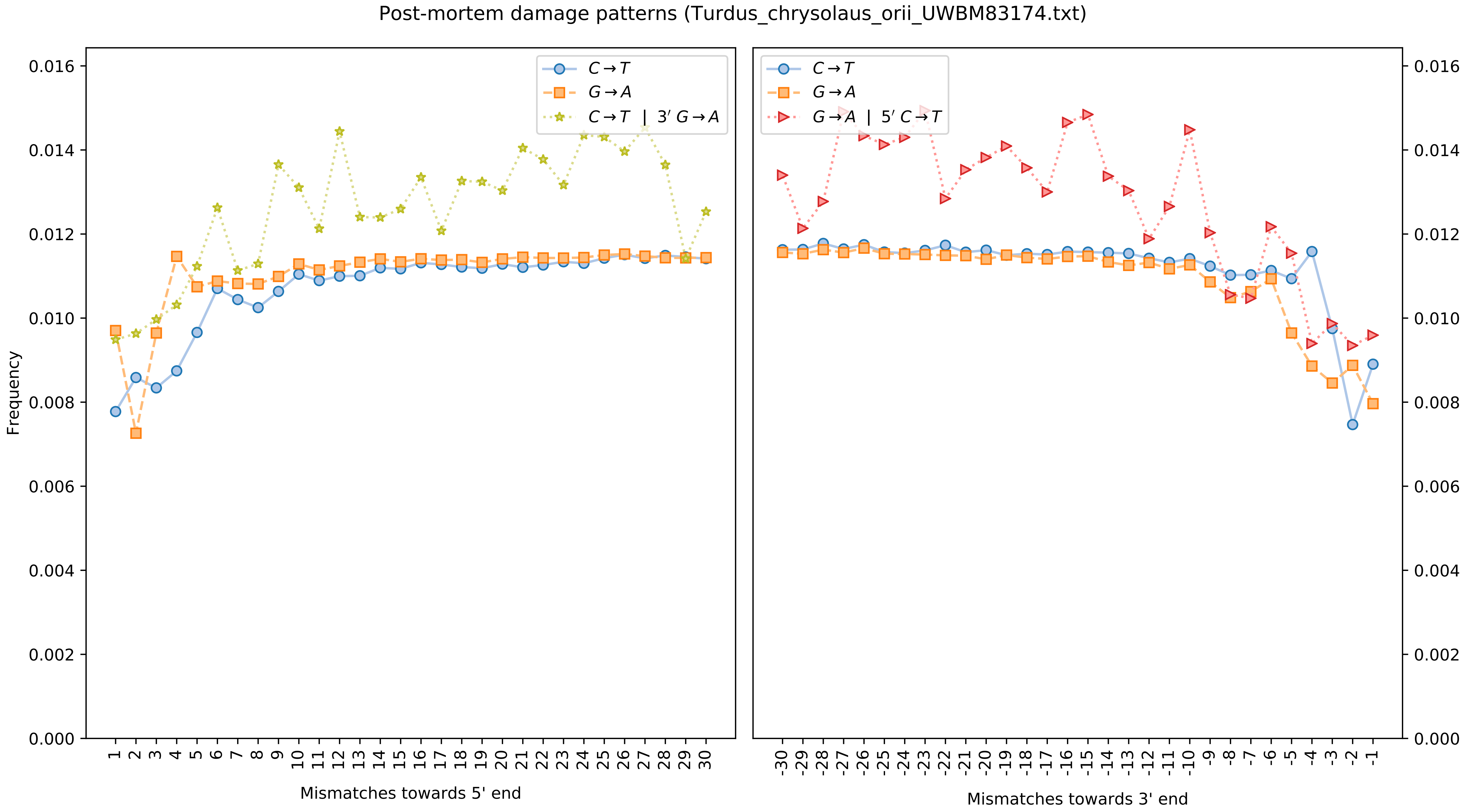
Post-mortem damage patterns (Turdus_chrysolaus_orii_UWBM83174.txt)
Frequency
Mismatches towards 5' end
Mismatches towards 3' end
C→T
G→A
C→T | 3′ G→A
G→A | 5′ C→T
0.000
0.002
0.004
0.006
0.008
0.010
0.012
0.014
0.016

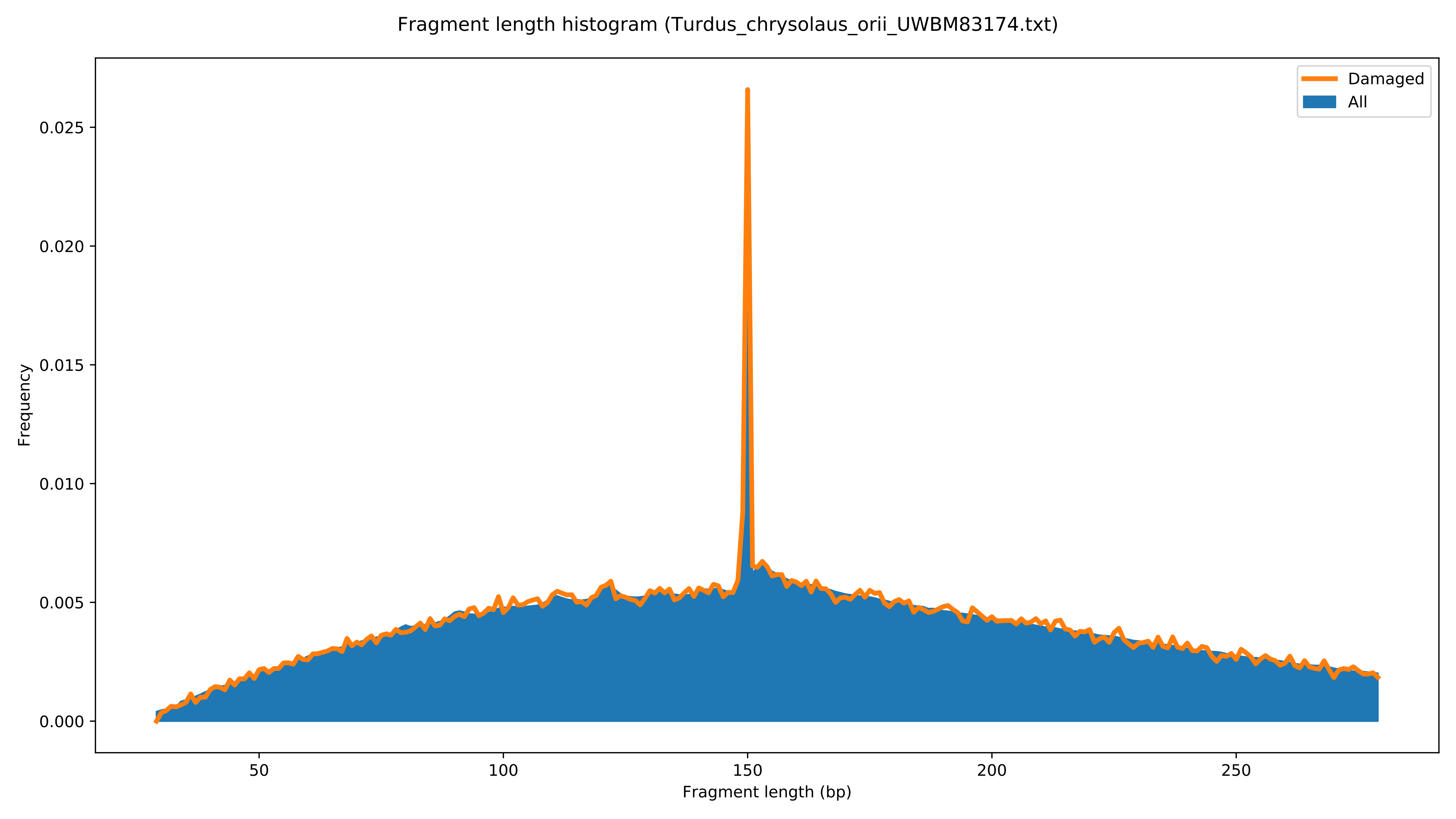

Fragment length histogram (Turdus_chrysolaus_orii_UWBM83174.txt)
Damaged
All
Frequency
0.000
0.005
0.010
0.015
0.020
0.025
50
100
150
200
250
Fragment length (bp)

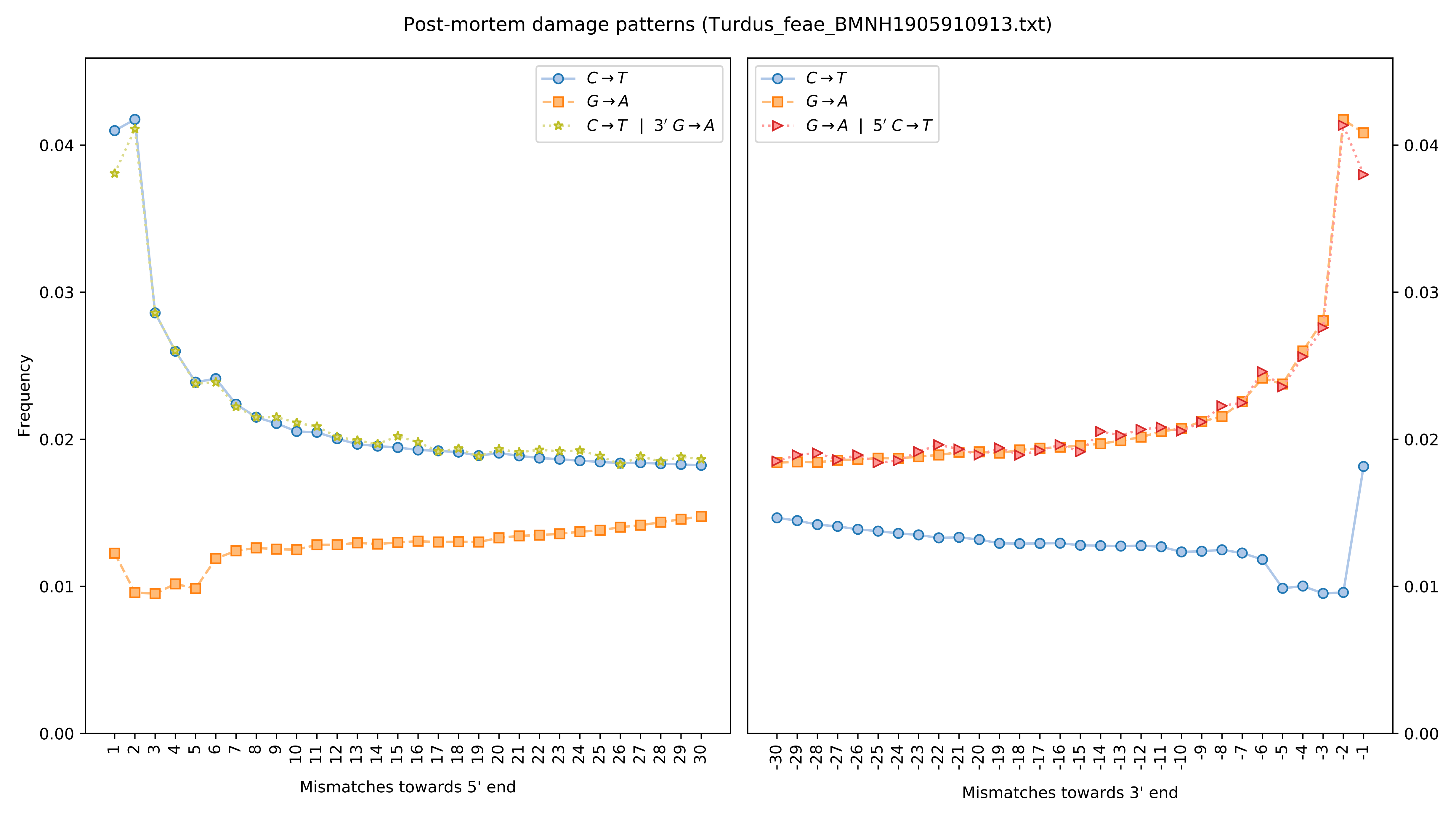
Post-mortem damage patterns (Turdus_feae_BMNH1905910913.txt)
Frequency
Mismatches towards 5' end
Mismatches towards 3' end
$C \rightarrow T$
$G \rightarrow A$
$C \rightarrow T$ | $3'$ $G \rightarrow A$
$G \rightarrow A$ | $5'$ $C \rightarrow T$
0.00
0.01
0.02
0.03
0.04

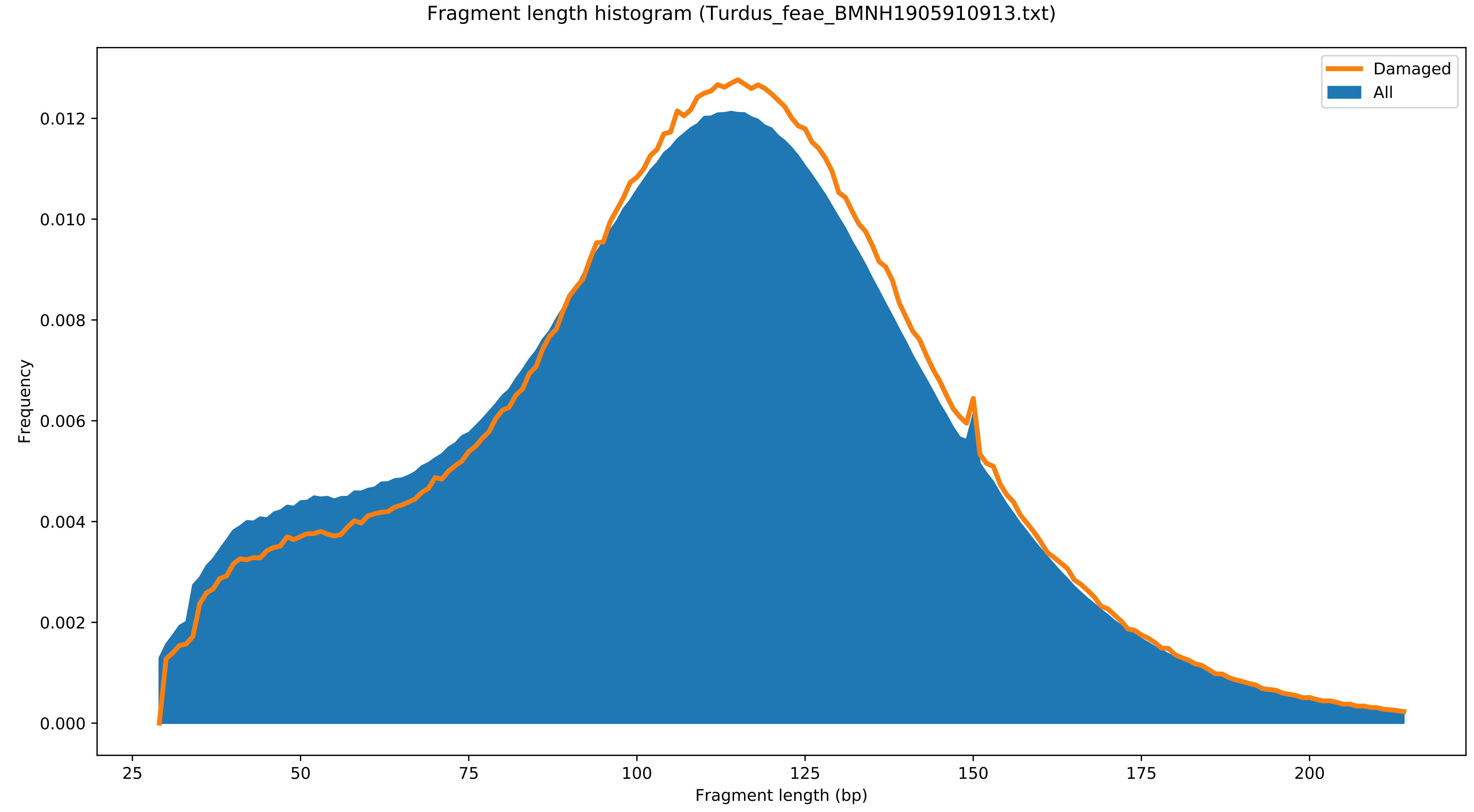

Fragment length histogram (Turdus_feae_BMNH1905910913.txt)
Damaged
All
Frequency
0.000
0.002
0.004
0.006
0.008
0.010
0.012
25
50
75
100
125
150
175
200
Fragment length (bp)

Post-mortem damage patterns (Turdus_merula.txt)

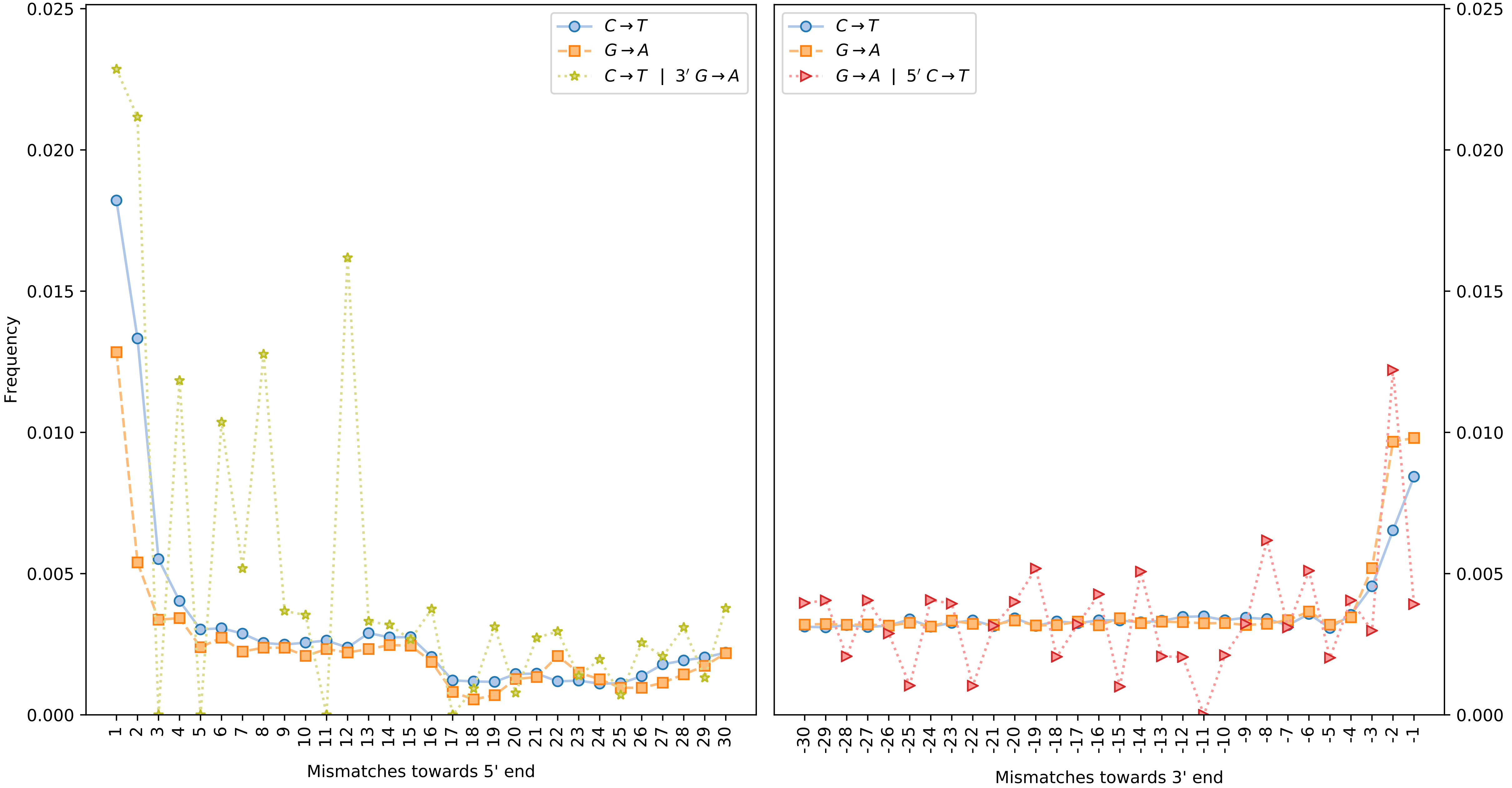

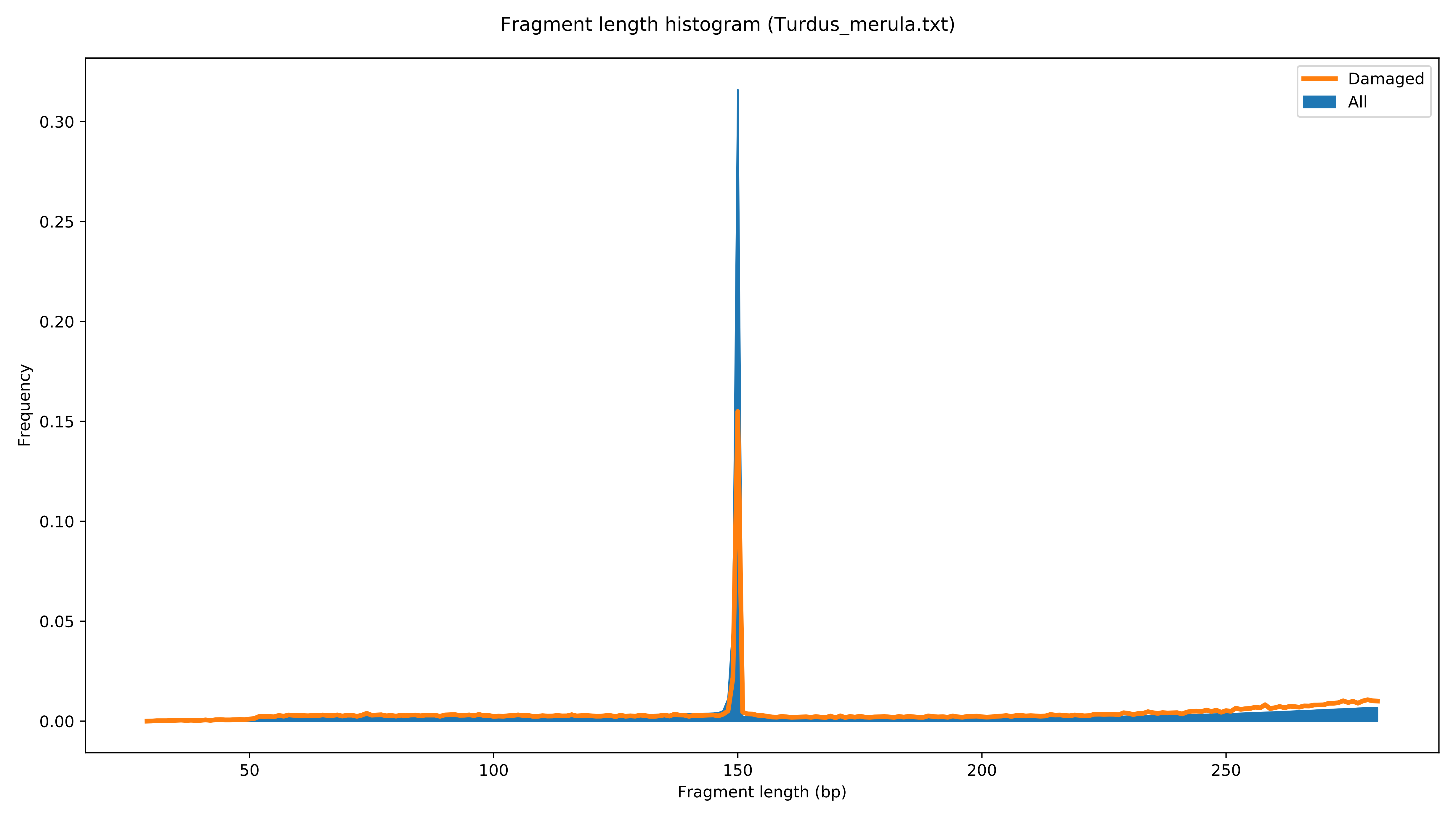
Fragment length histogram (Turdus_merula.txt)
Damaged
All
Frequency
0.00
0.05
0.10
0.15
0.20
0.25
0.30
50
100
150
200
250
Fragment length (bp)

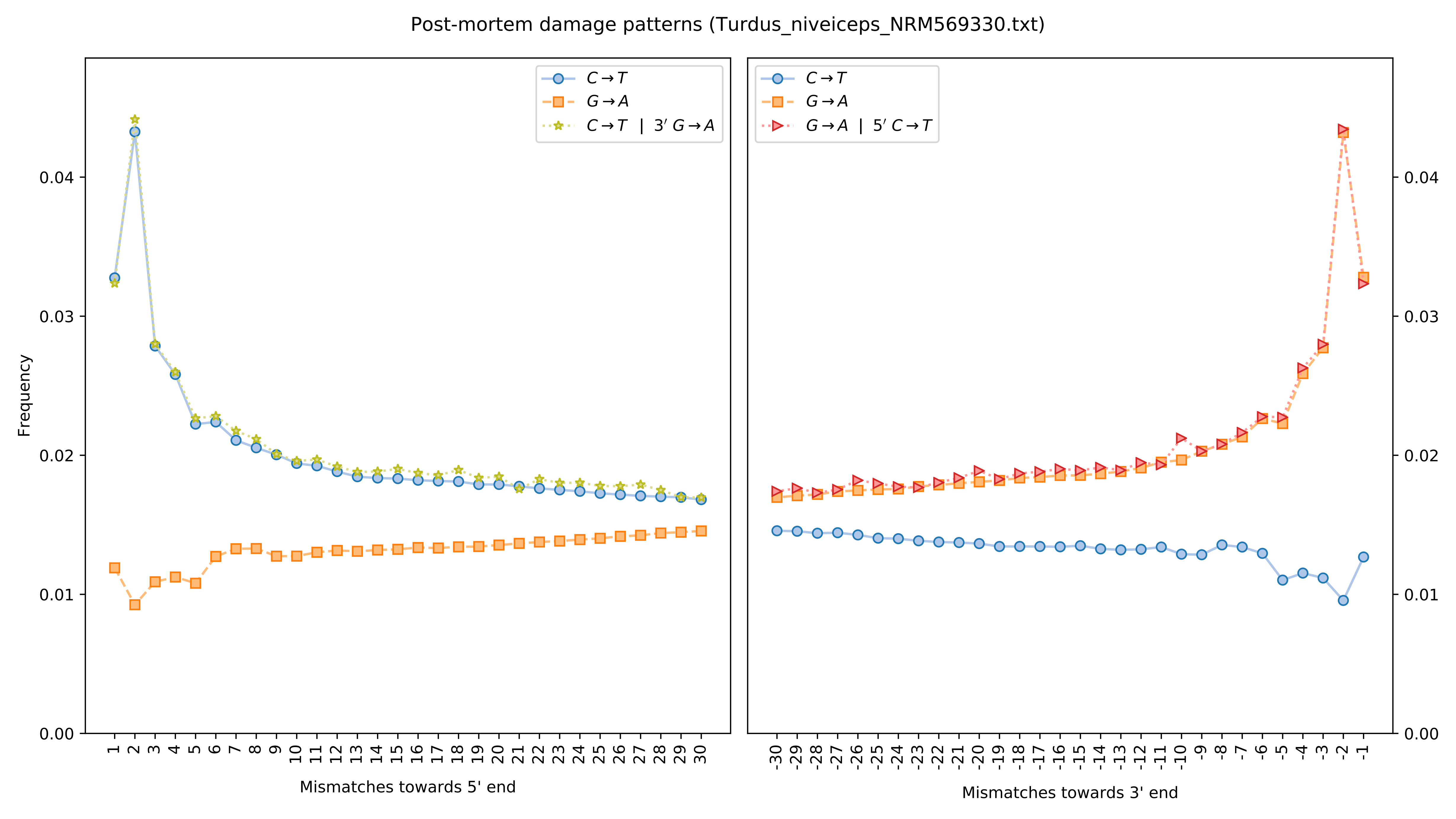

Post-mortem damage patterns (Turdus_niveiceps_NRM569330.txt)
C→T
G→A
C→T | 3′ G→A
G→A | 5′ C→T
Frequency
0.00
0.01
0.02
0.03
0.04
Mismatches towards 5' end
Mismatches towards 3' end

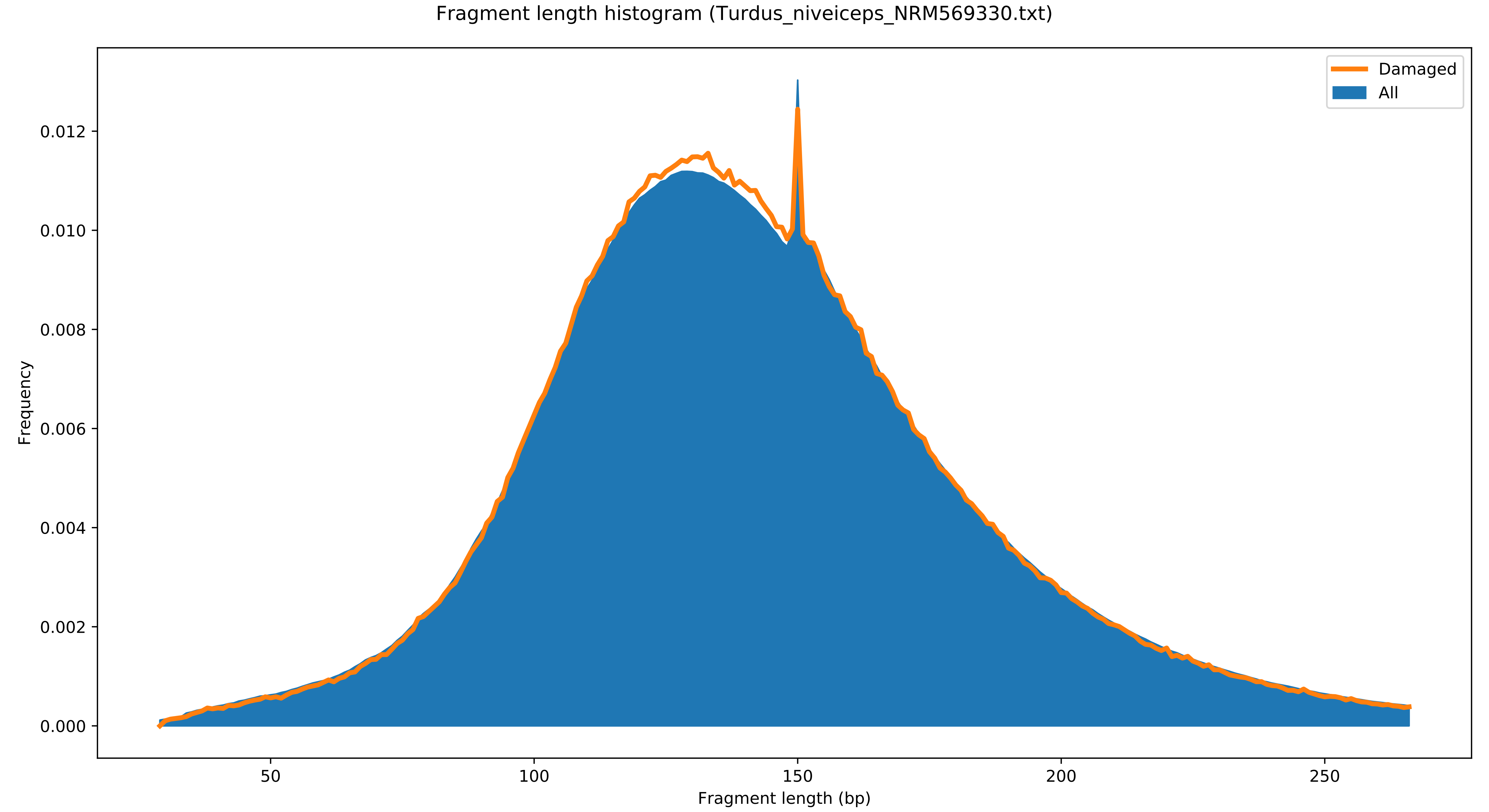

Fragment length histogram (Turdus_niveiceps_NRM569330.txt)
Damaged
All
Frequency
0.000
0.002
0.004
0.006
0.008
0.010
0.012
50
100
150
200
250
Fragment length (bp)

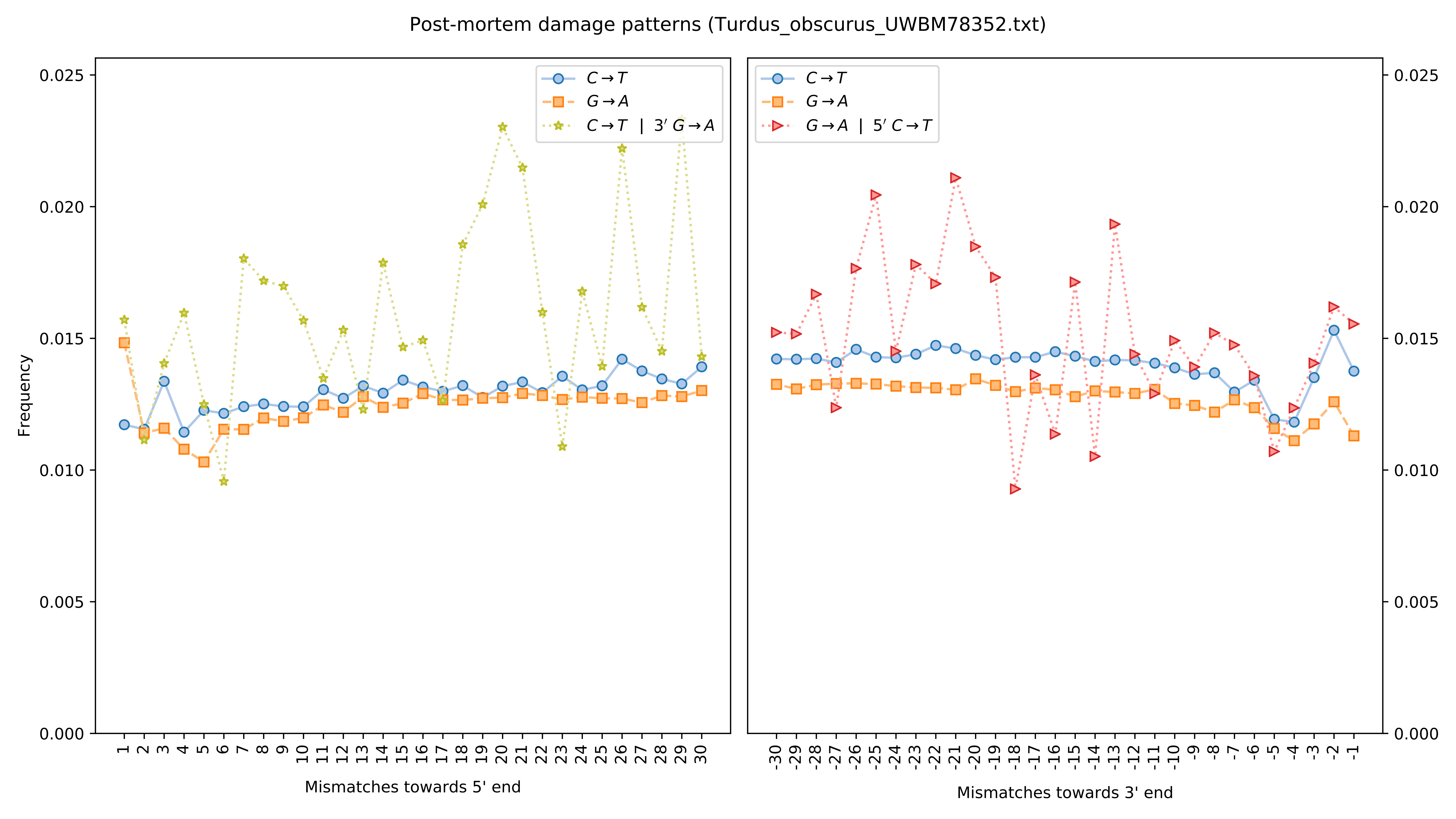
Post-mortem damage patterns (Turdus_obscurus_UWBM78352.txt)
Frequency
Mismatches towards 5' end
Mismatches towards 3' end
C→T
G→A
C→T | 3′ G→A
G→A | 5′ C→T
0.000
0.005
0.010
0.015
0.020
0.025

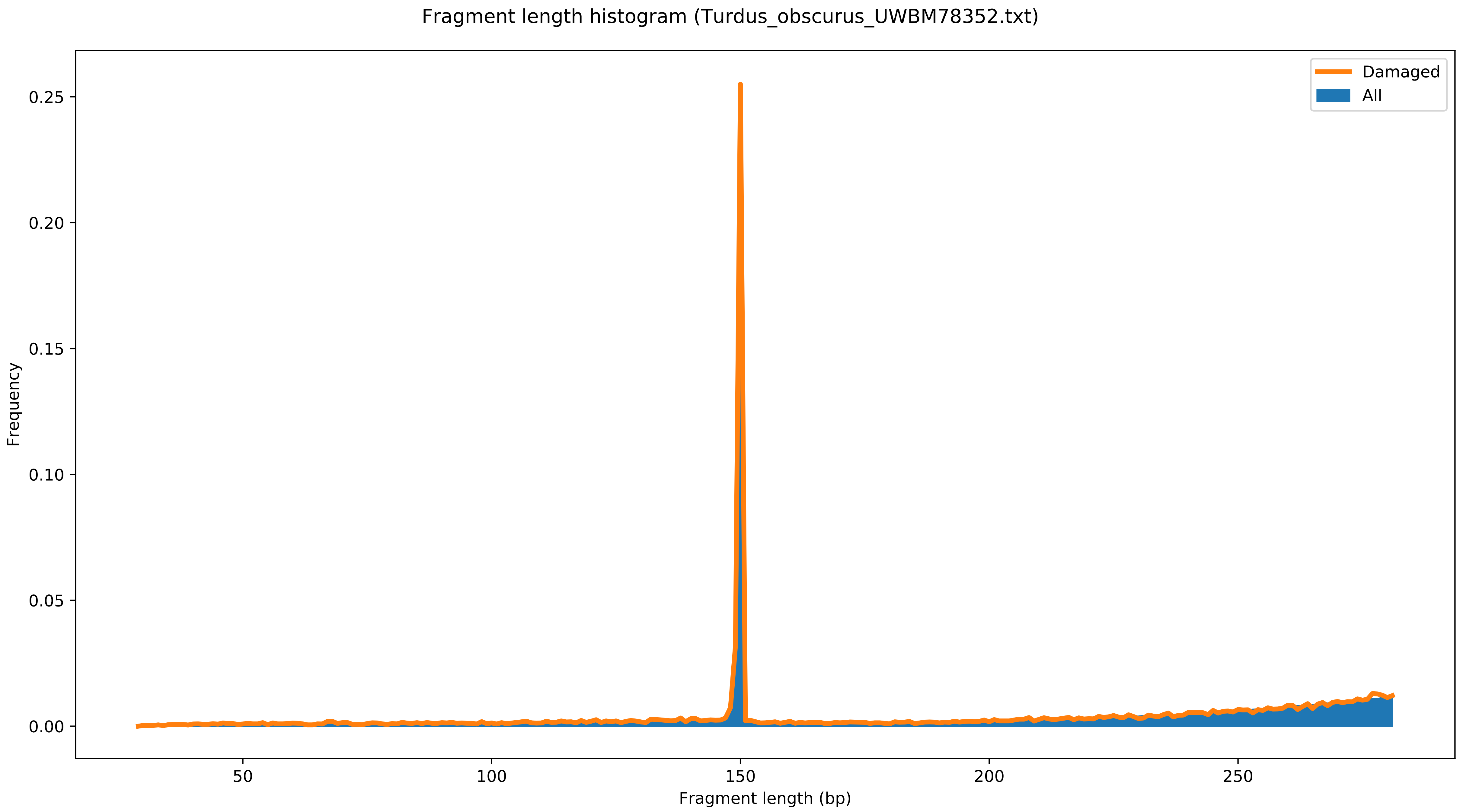
Fragment length histogram (Turdus_obscurus_UWBM78352.txt)
Damaged
All
Frequency
0.25
0.20
0.15
0.10
0.05
0.00
50
100
150
200
250
Fragment length (bp)

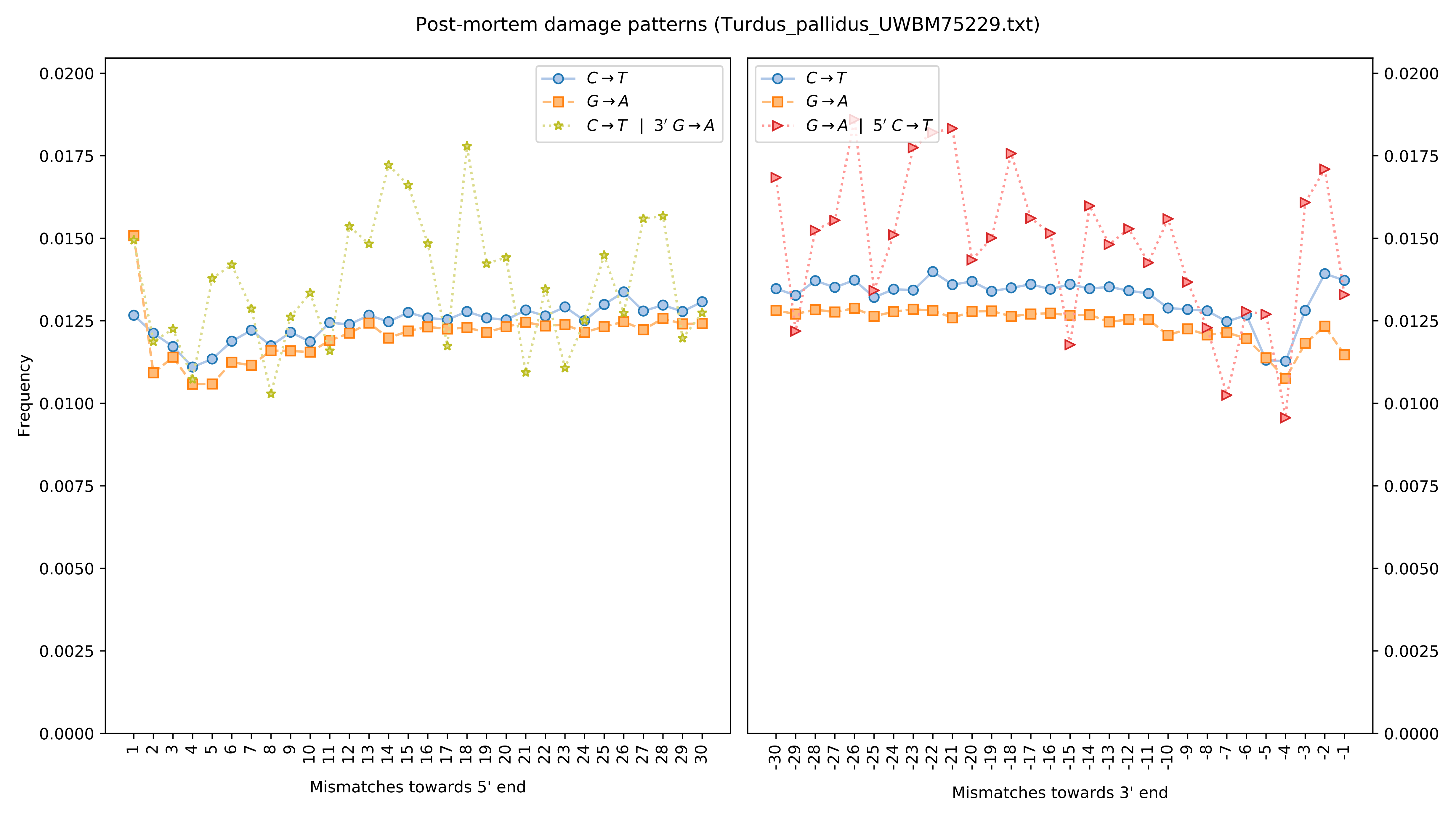

Post-mortem damage patterns (Turdus_pallidus_UWBM75229.txt)
C→T
G→A
C→T | 3′ G→A
G→A | 5′ C→T
Frequency
Mismatches towards 5' end
Mismatches towards 3' end

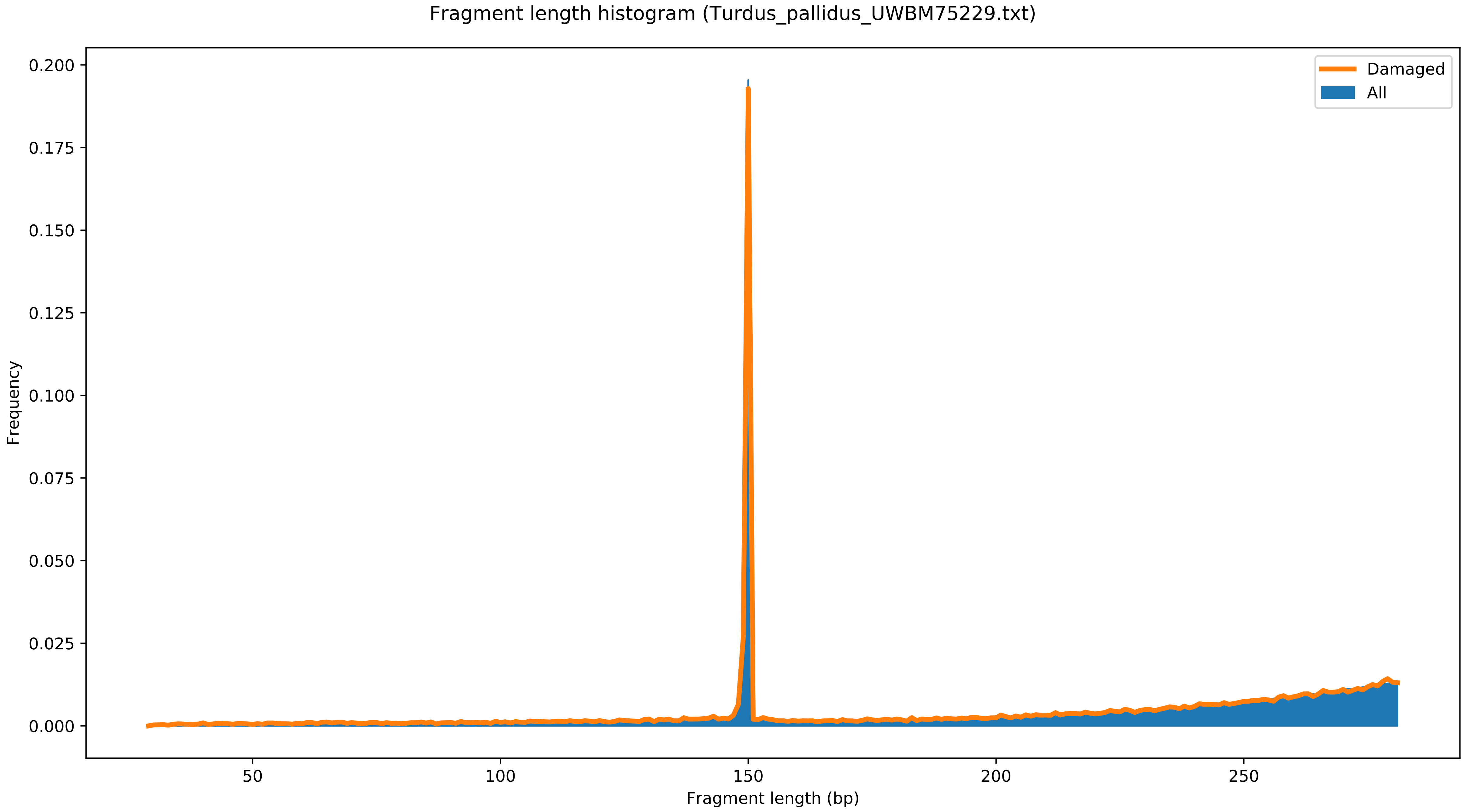

Fragment length histogram (Turdus_pallidus_UWBM75229.txt)
Damaged
All
Frequency
0.200
0.175
0.150
0.125
0.100
0.075
0.050
0.025
0.000
50
100
150
200
250
Fragment length (bp)

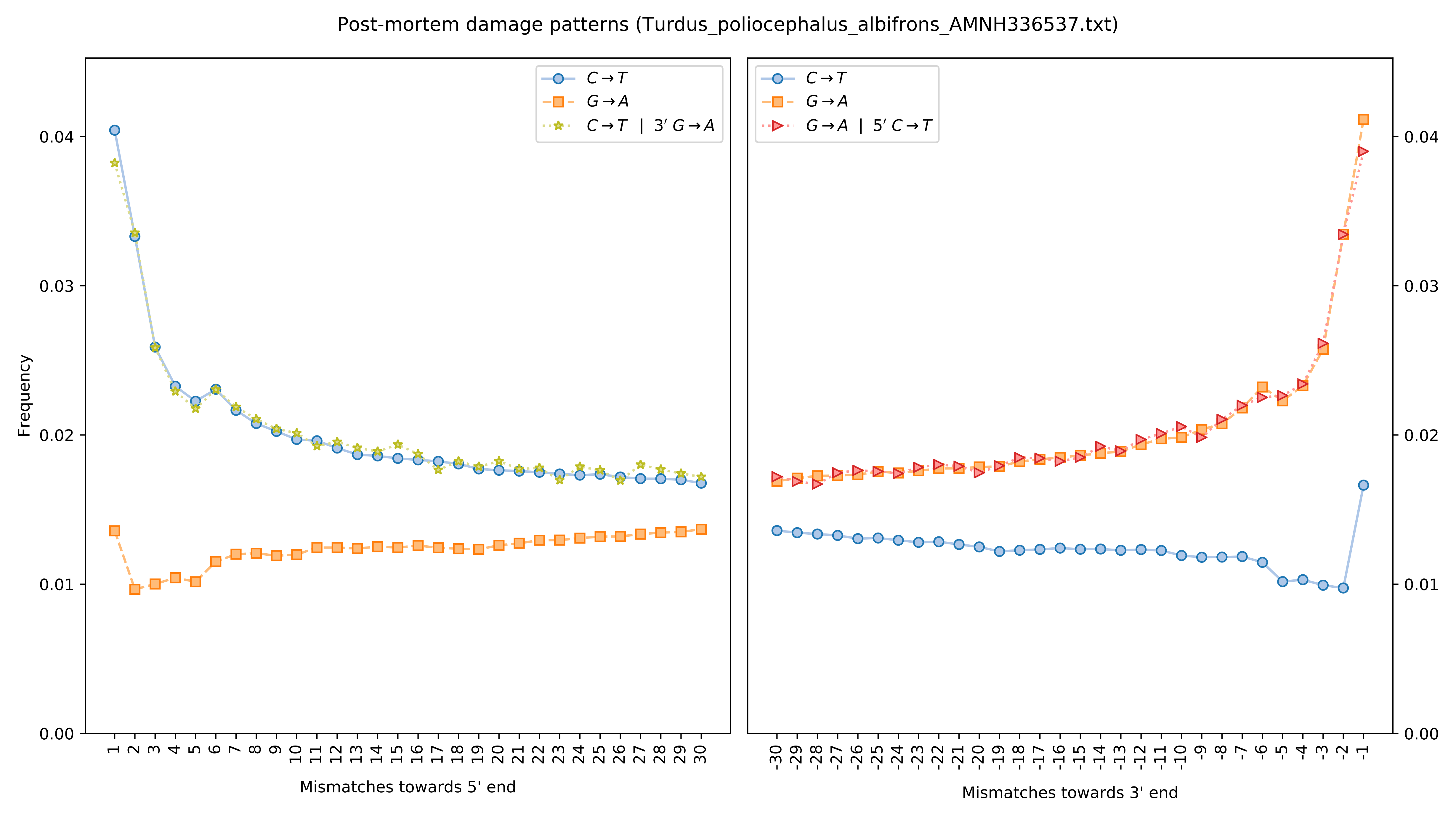

Post-mortem damage patterns (Turdus_poliocephalus_albifrons_AMNH336537.txt)
Frequency
Mismatches towards 5' end
Mismatches towards 3' end
C→T
G→A
C→T | 3′ G→A
G→A | 5′ C→T
0.00
0.01
0.02
0.03
0.04

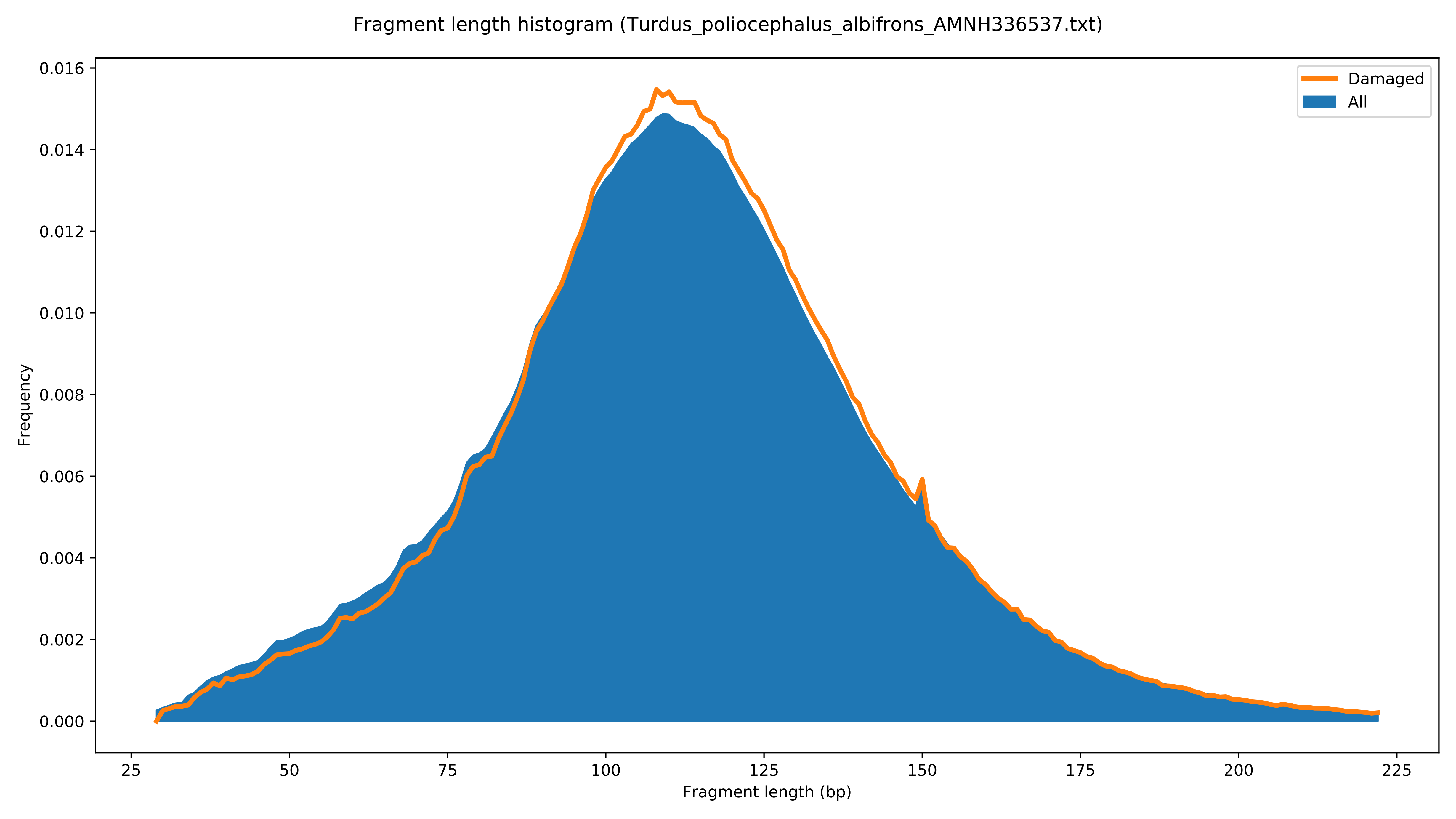

Fragment length histogram (Turdus_poliocephalus_albifrons_AMNH336537.txt)
Damaged
All
Frequency
0.000
0.002
0.004
0.006
0.008
0.010
0.012
0.014
0.016
25
50
75
100
125
150
175
200
225
Fragment length (bp)

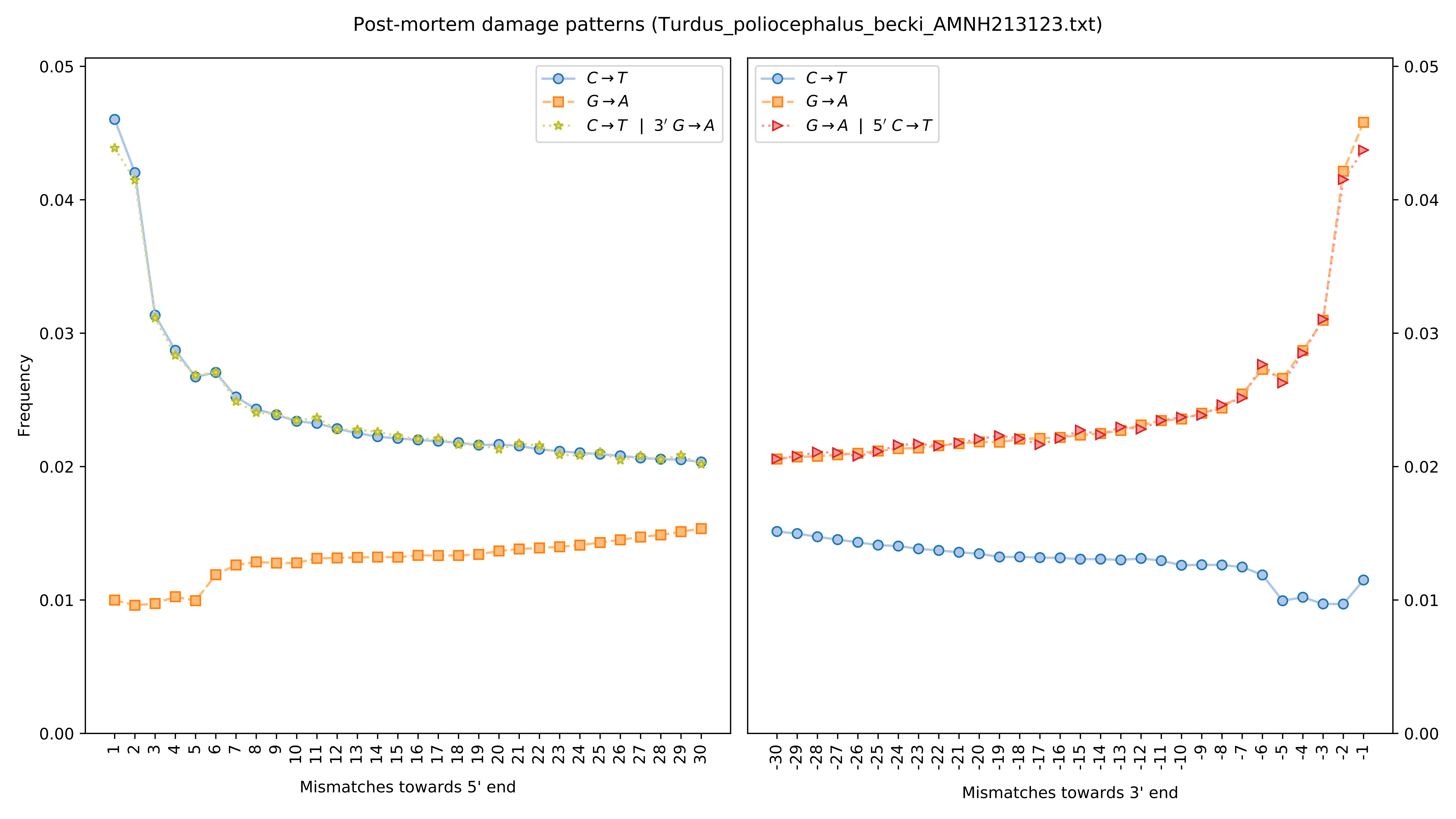
Post-mortem damage patterns (Turdus_poliocephalus_becki_AMNH213123.txt)
Frequency
Mismatches towards 5' end
Mismatches towards 3' end
C→T
G→A
C→T | 3′ G→A
G→A | 5′ C→T
0.00
0.01
0.02
0.03
0.04
0.05

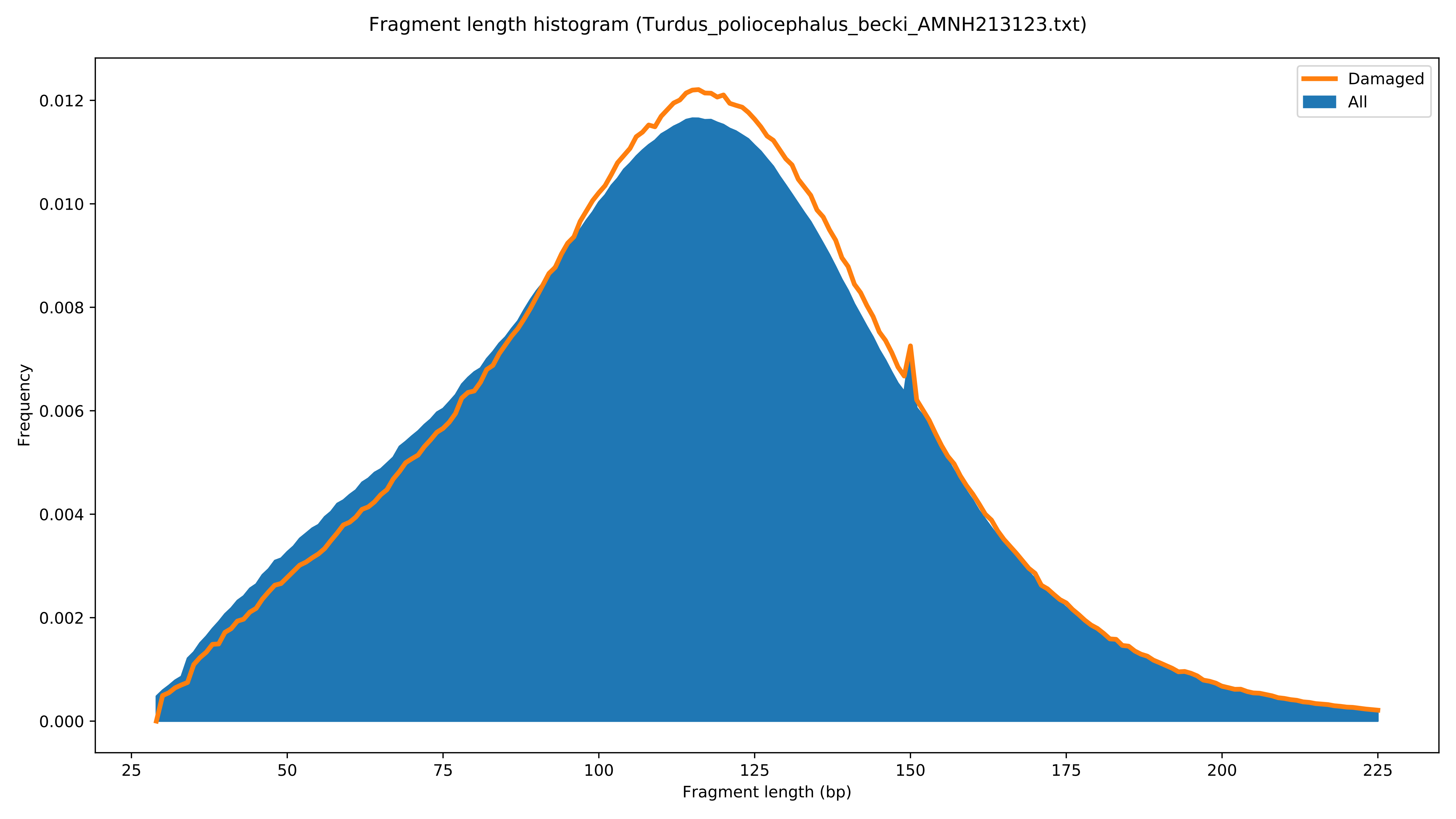

Fragment length histogram (Turdus_poliocephalus_becki_AMNH213123.txt)
Damaged
All
Frequency
0.000
0.002
0.004
0.006
0.008
0.010
0.012
25
50
75
100
125
150
175
200
225
Fragment length (bp)

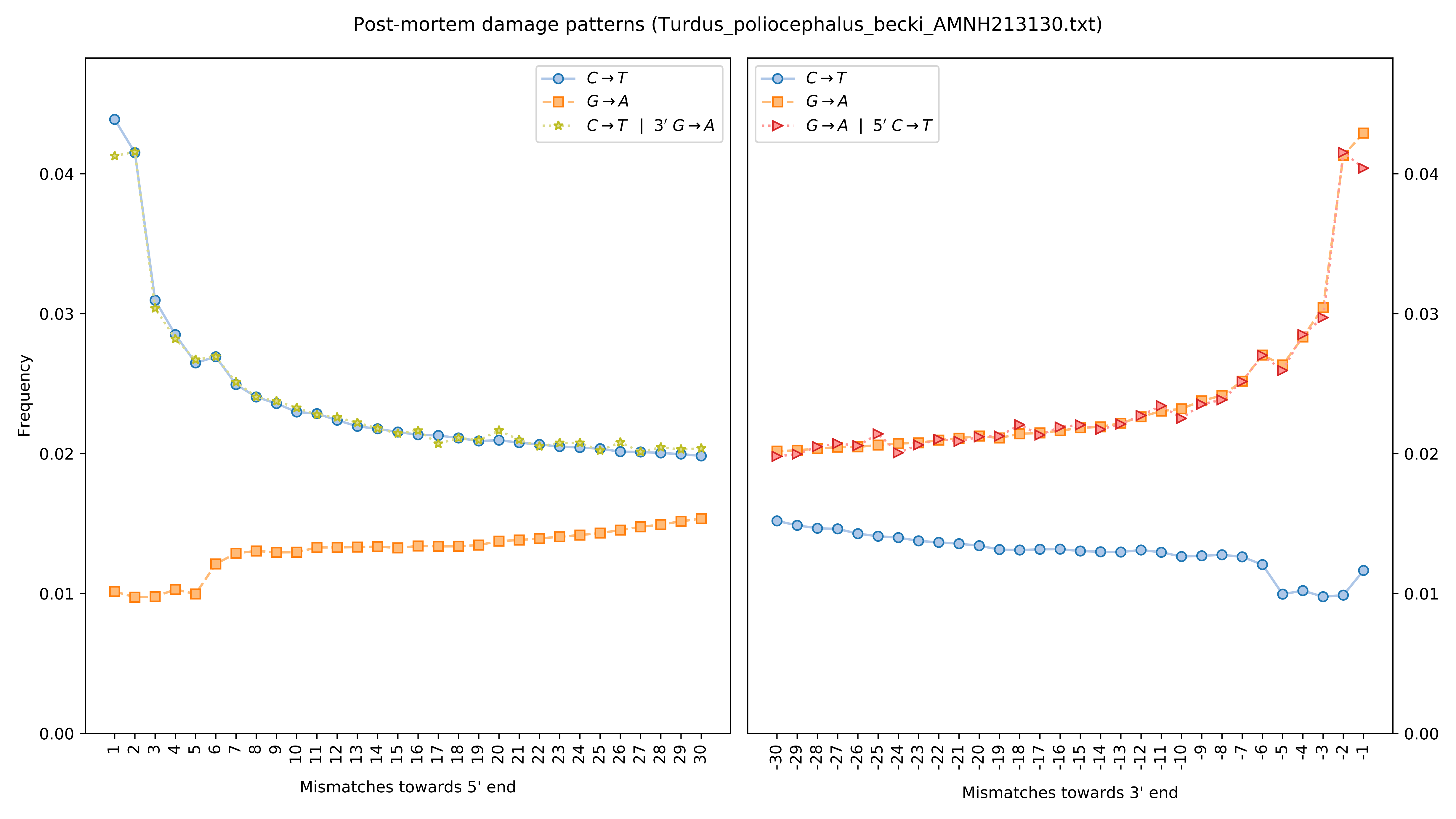

Post-mortem damage patterns (Turdus_poliocephalus_becki_AMNH213130.txt)
Frequency
Mismatches towards 5' end
Mismatches towards 3' end
C→T
G→A
C→T | 3′ G→A
G→A | 5′ C→T

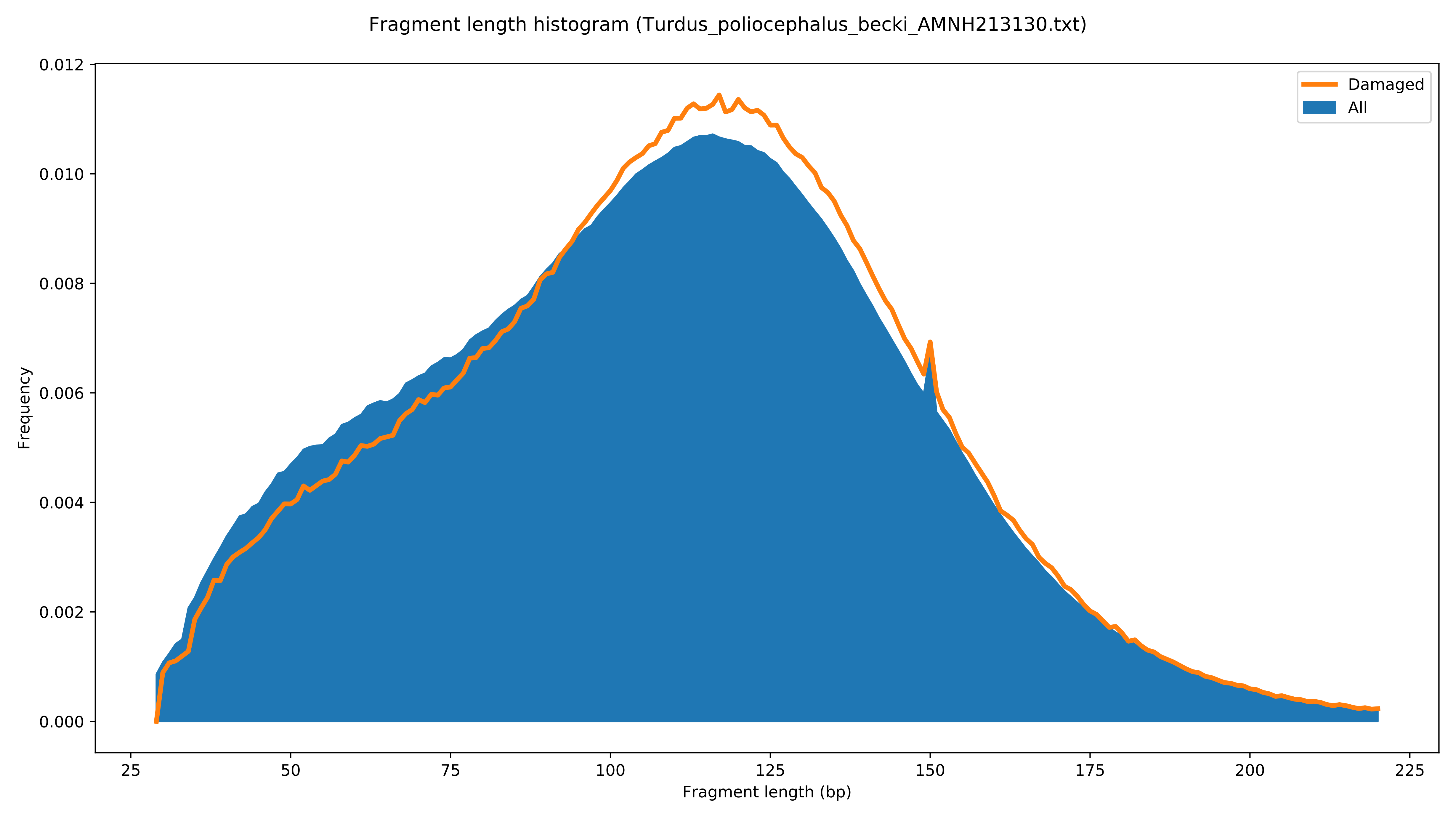

Fragment length histogram (Turdus_poliocephalus_becki_AMNH213130.txt)
Damaged
All
Frequency
0.000
0.002
0.004
0.006
0.008
0.010
0.012
25
50
75
100
125
150
175
200
225
Fragment length (bp)

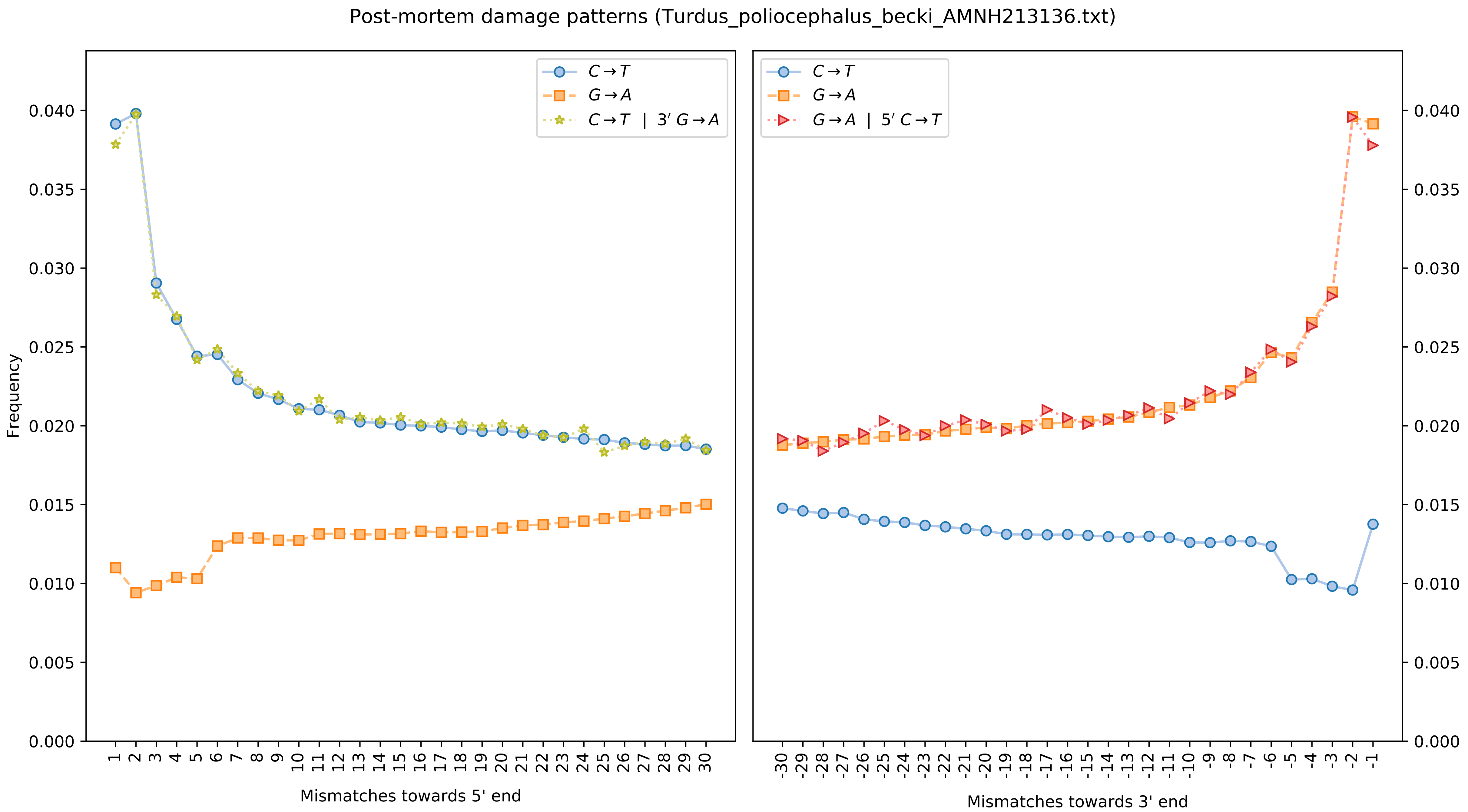

Post-mortem damage patterns (Turdus_poliocephalus_becki_AMNH213136.txt)
Frequency
$C \rightarrow T$
$G \rightarrow A$
$C \rightarrow T$ | $3' G \rightarrow A$
$G \rightarrow A$ | $5' C \rightarrow T$
Mismatches towards 5' end
Mismatches towards 3' end

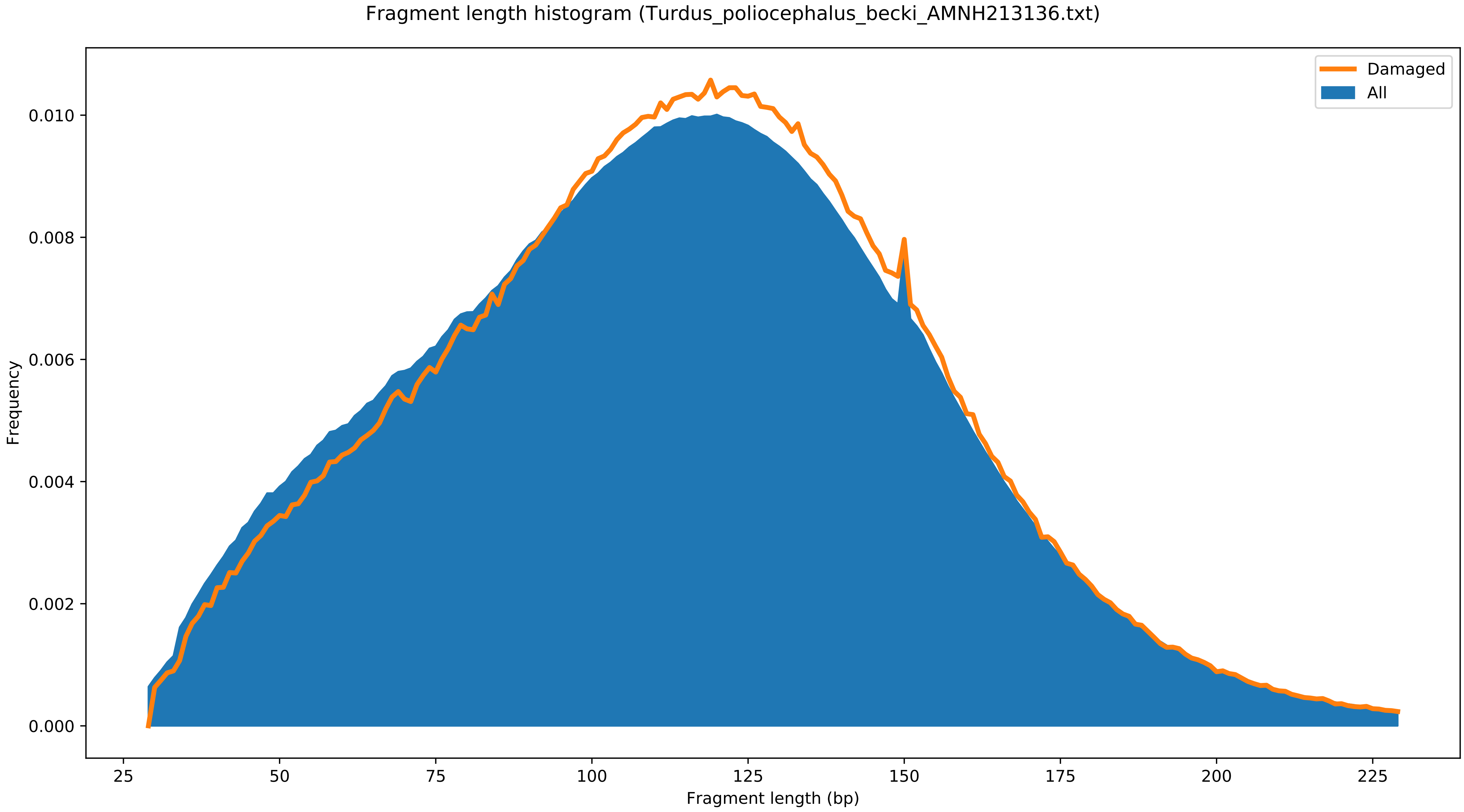
Fragment length histogram (Turdus_poliocephalus_becki_AMNH213136.txt)
Damaged
All
Frequency
0.000
0.002
0.004
0.006
0.008
0.010
25
50
75
100
125
150
175
200
225
Fragment length (bp)

Post-mortem damage patterns (Turdus_poliocephalus_becki_AMNH213163.txt)

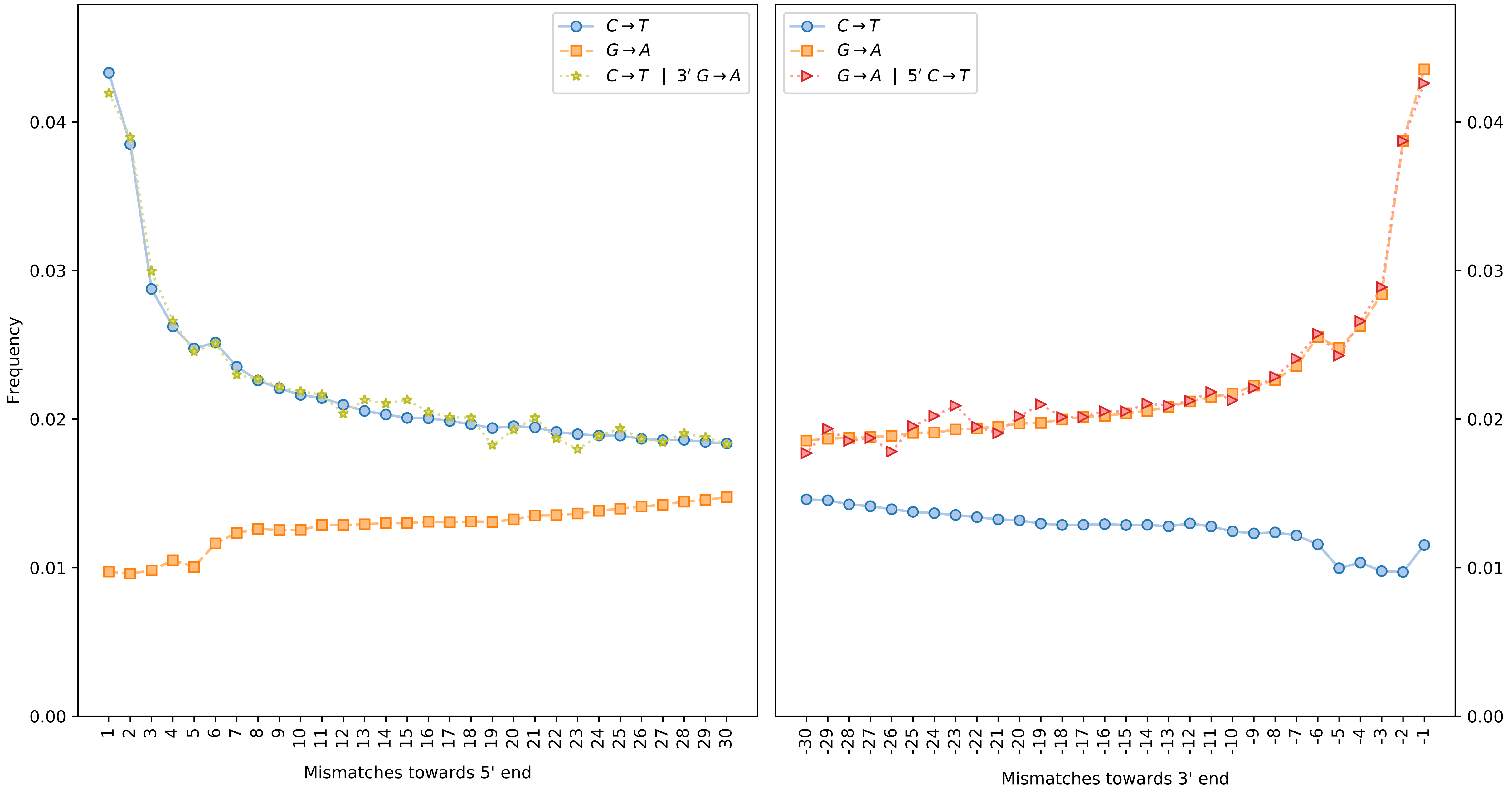

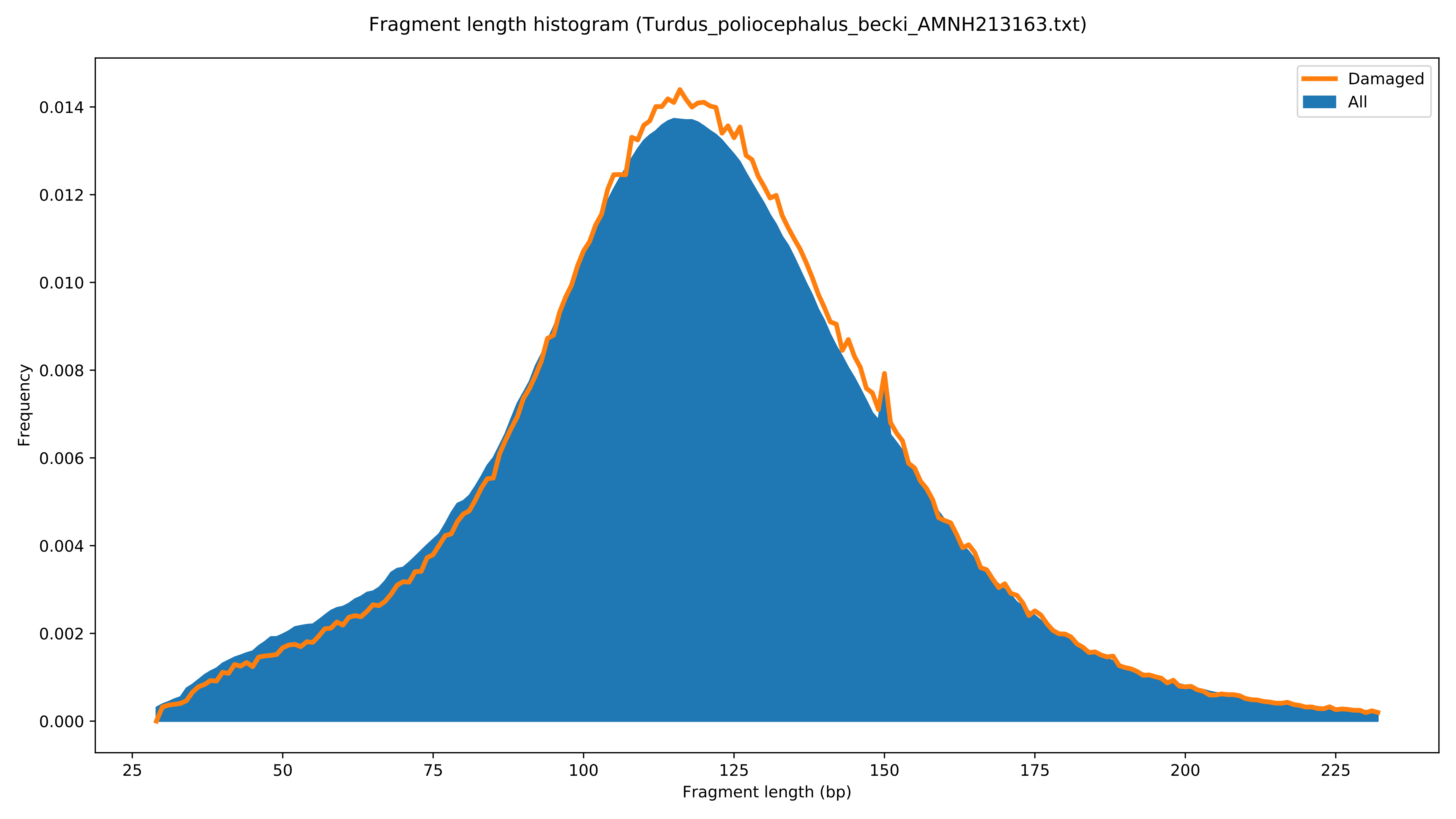

Fragment length histogram (Turdus_poliocephalus_becki_AMNH213163.txt)
Damaged
All
Frequency
0.000
0.002
0.004
0.006
0.008
0.010
0.012
0.014
25
50
75
100
125
150
175
200
225
Fragment length (bp)

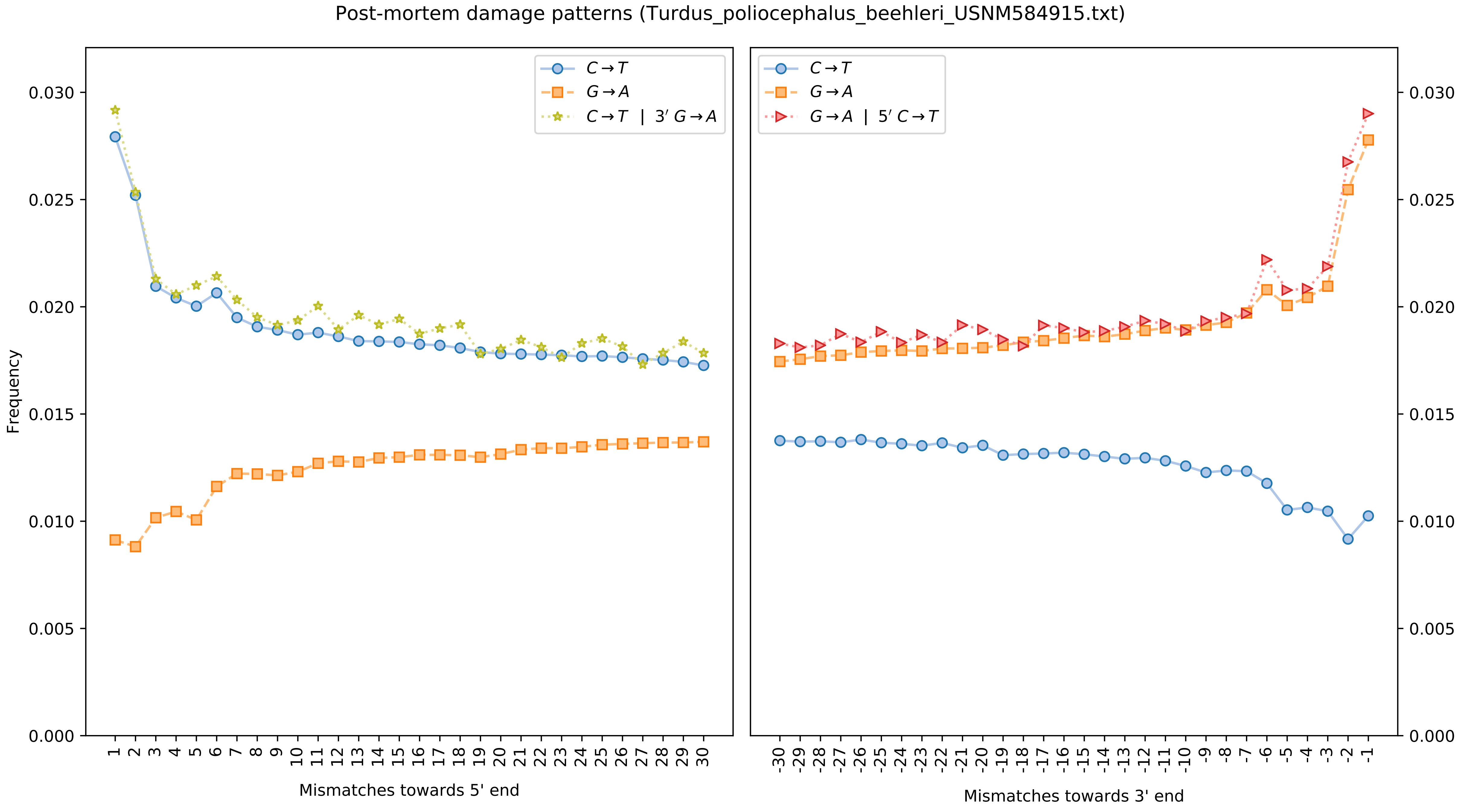
Post-mortem damage patterns (Turdus_poliocephalus_beehleri_USNM584915.txt)
C→T
G→A
C→T | 3′ G→A
G→A | 5′ C→T
Frequency
Mismatches towards 5' end
Mismatches towards 3' end

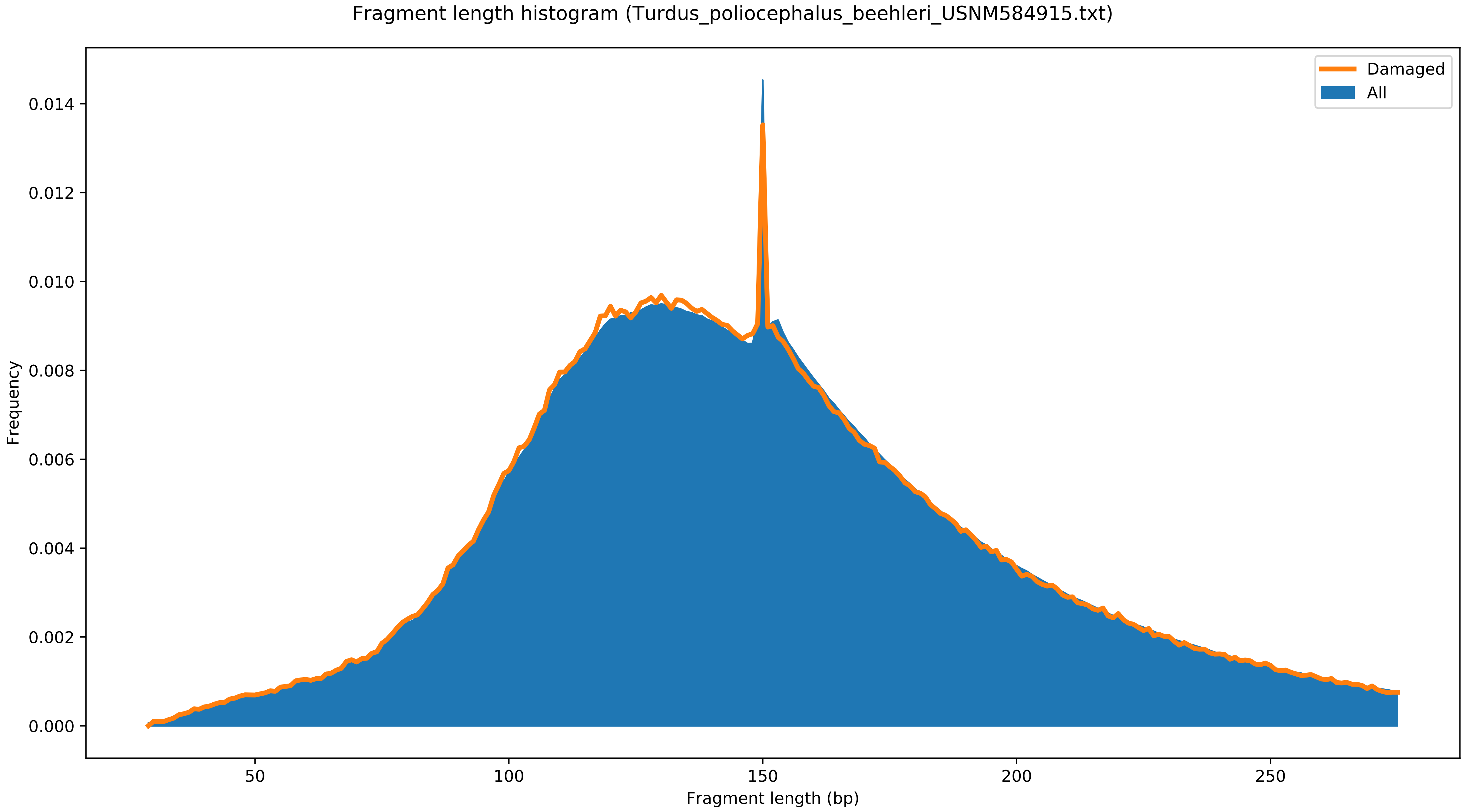
Fragment length histogram (Turdus_poliocephalus_beehleri_USNM584915.txt)
Damaged
All
Frequency
0.000
0.002
0.004
0.006
0.008
0.010
0.012
0.014
50
100
150
200
250
Fragment length (bp)

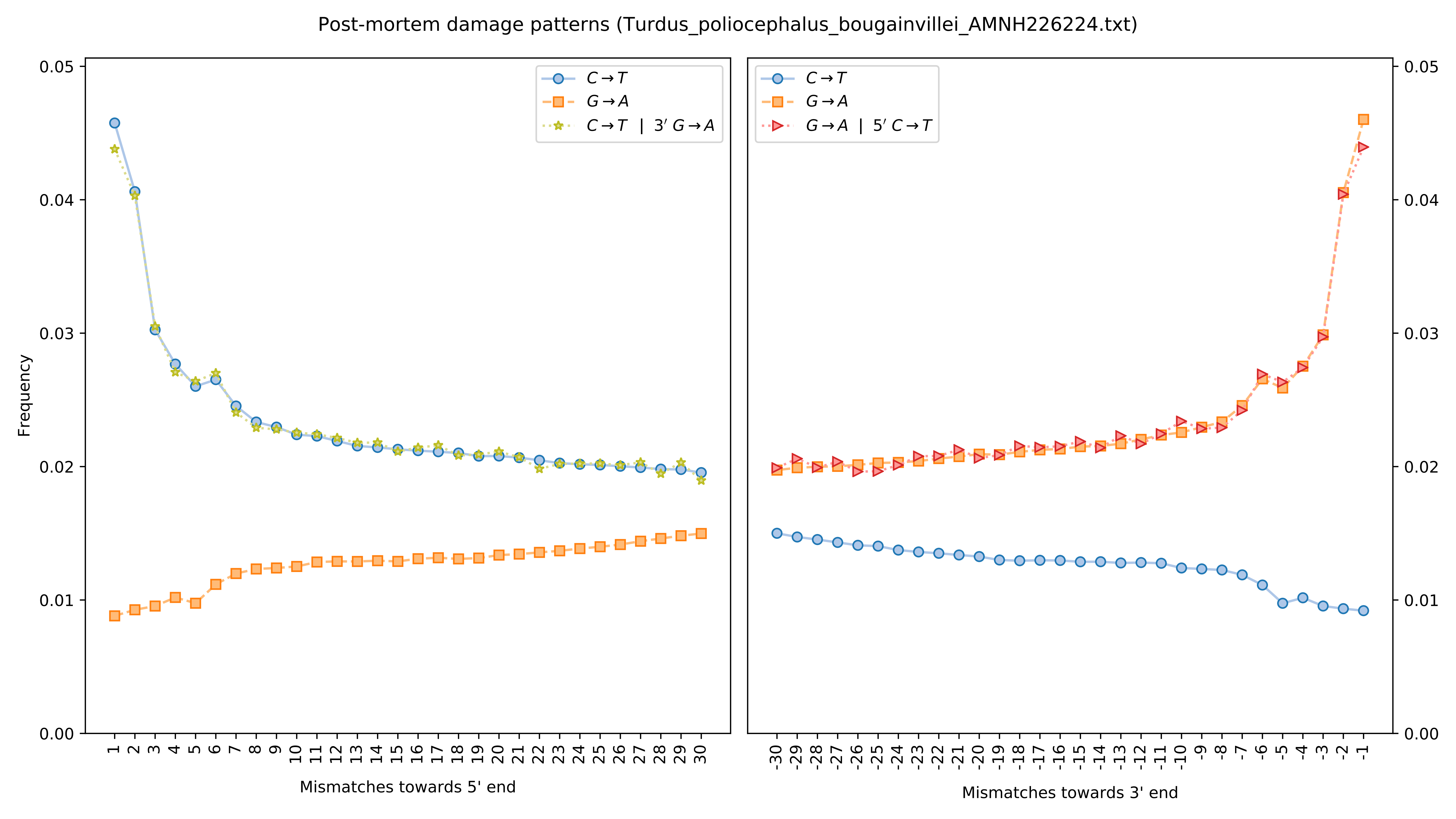
Post-mortem damage patterns (Turdus_poliocephalus_bougainvillei_AMNH226224.txt)
Frequency
C→T
G→A
C→T | 3′ G→A
G→A | 5′ C→T
Mismatches towards 5' end
Mismatches towards 3' end
0.00
0.01
0.02
0.03
0.04
0.05

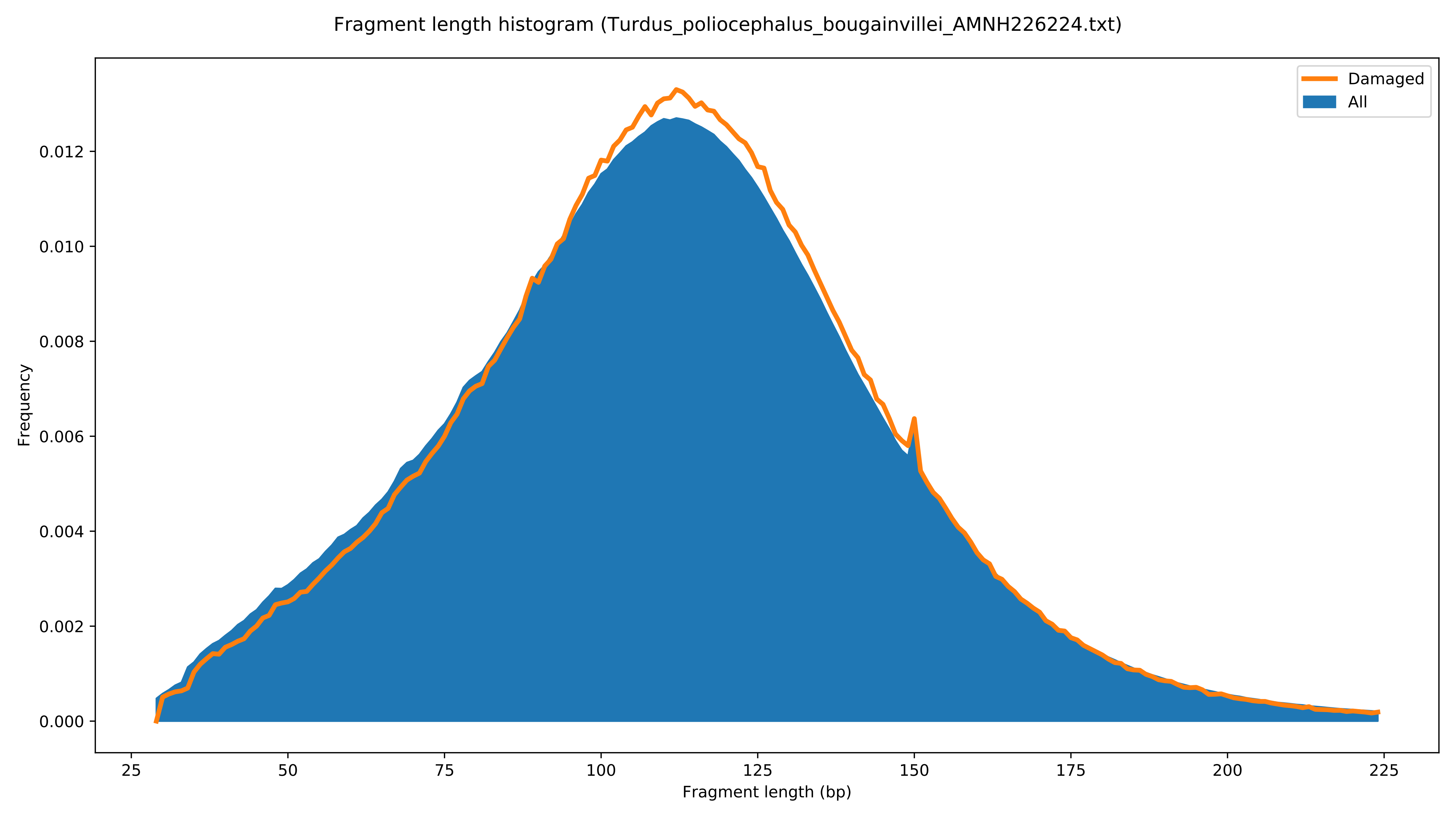

Fragment length histogram (Turdus_poliocephalus_bougainvillei_AMNH226224.txt)
Damaged
All
Frequency
Fragment length (bp)
0.000
0.002
0.004
0.006
0.008
0.010
0.012
25
50
75
100
125
150
175
200
225

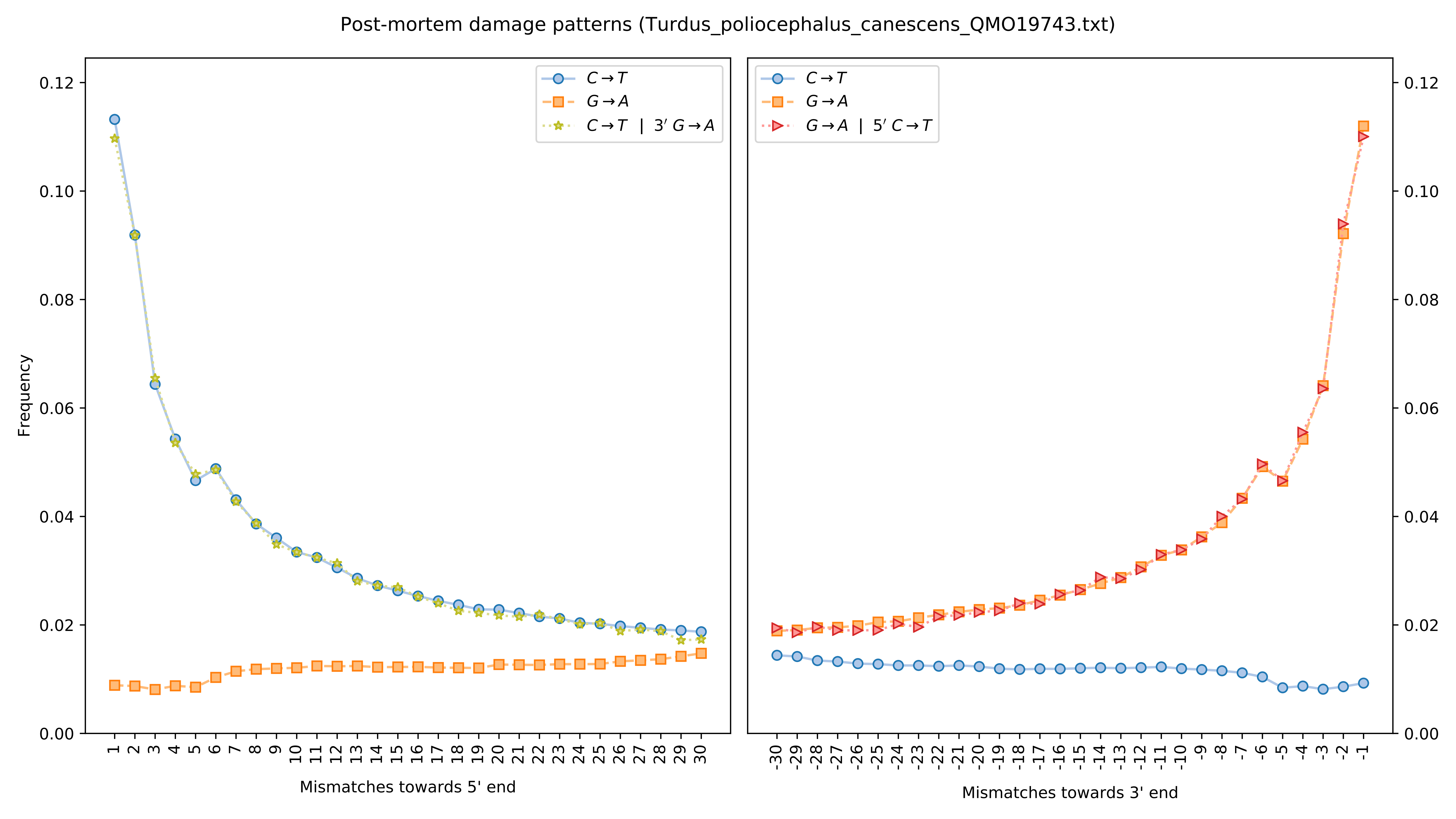

Post-mortem damage patterns (Turdus_poliocephalus_canescens_QMO19743.txt)
Frequency
Mismatches towards 5' end
Mismatches towards 3' end
C→T
G→A
C→T | 3′ G→A
G→A | 5′ C→T

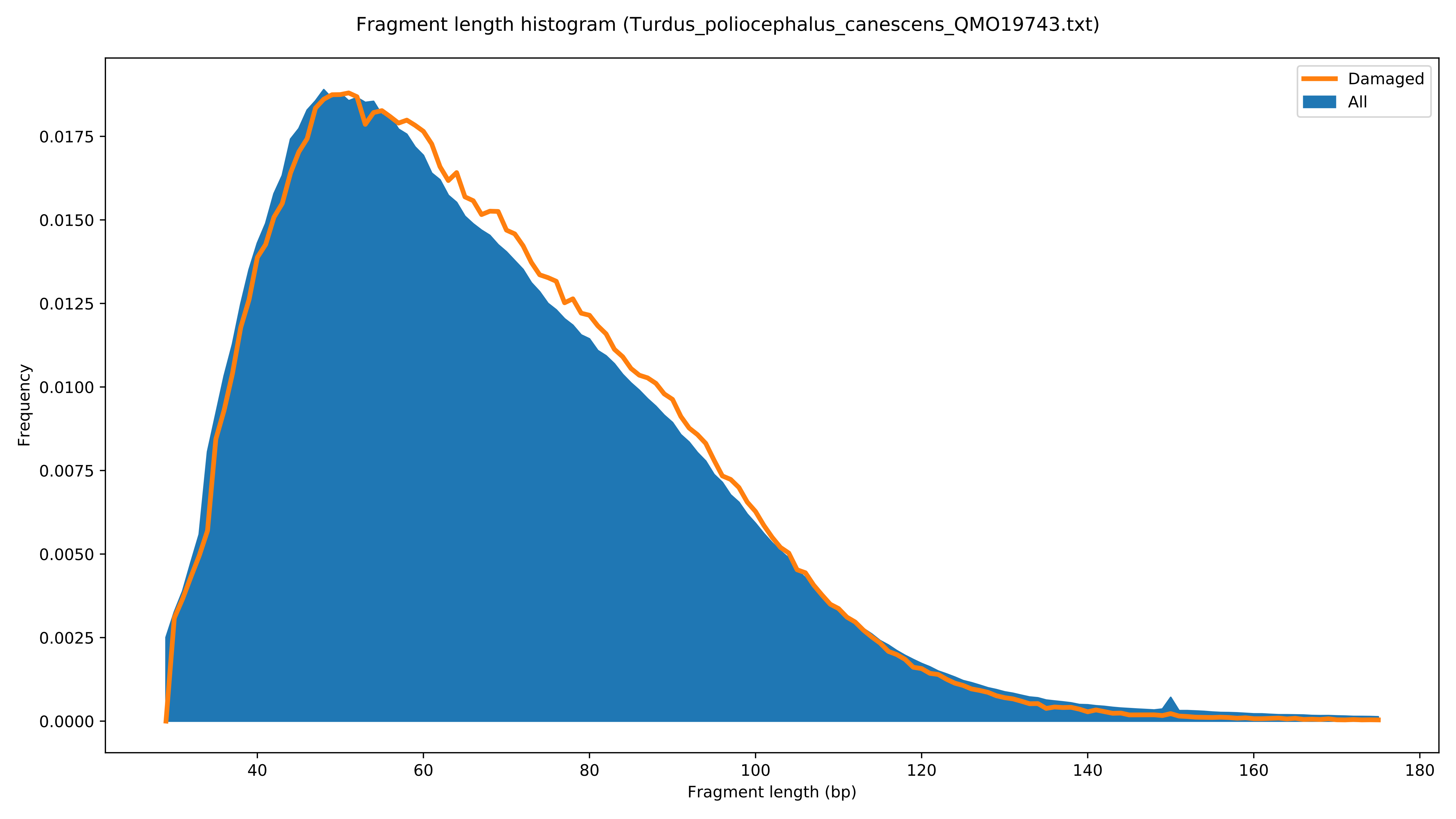

Fragment length histogram (Turdus_poliocephalus_canescens_QMO19743.txt)
Damaged
All
Frequency
0.0000
0.0025
0.0050
0.0075
0.0100
0.0125
0.0150
0.0175
40
60
80
100
120
140
160
180
Fragment length (bp)

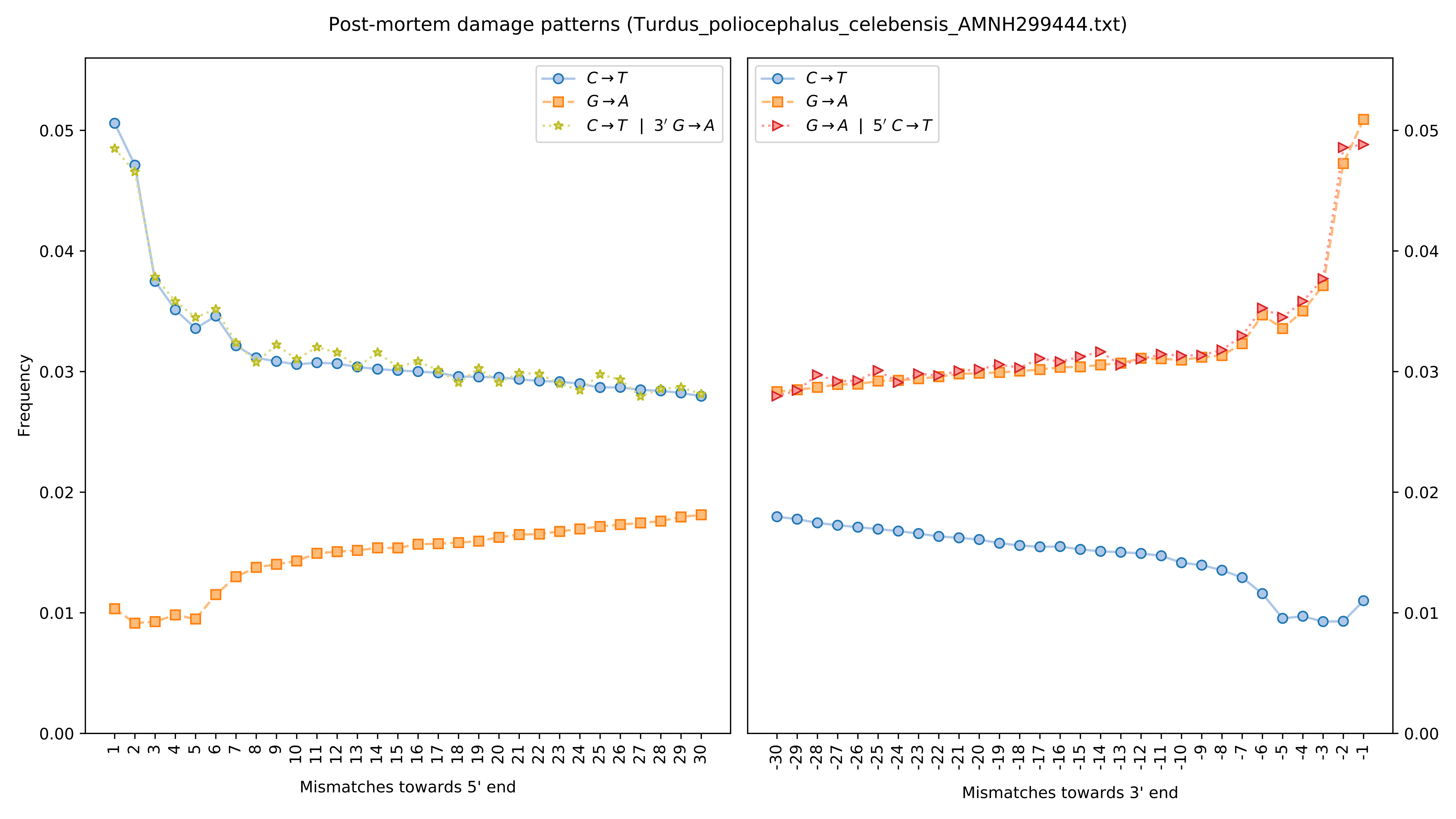
Post-mortem damage patterns (Turdus_poliocephalus_celebensis_AMNH299444.txt)
Frequency
Mismatches towards 5' end
Mismatches towards 3' end
C→T
G→A
C→T | 3′ G→A
G→A | 5′ C→T

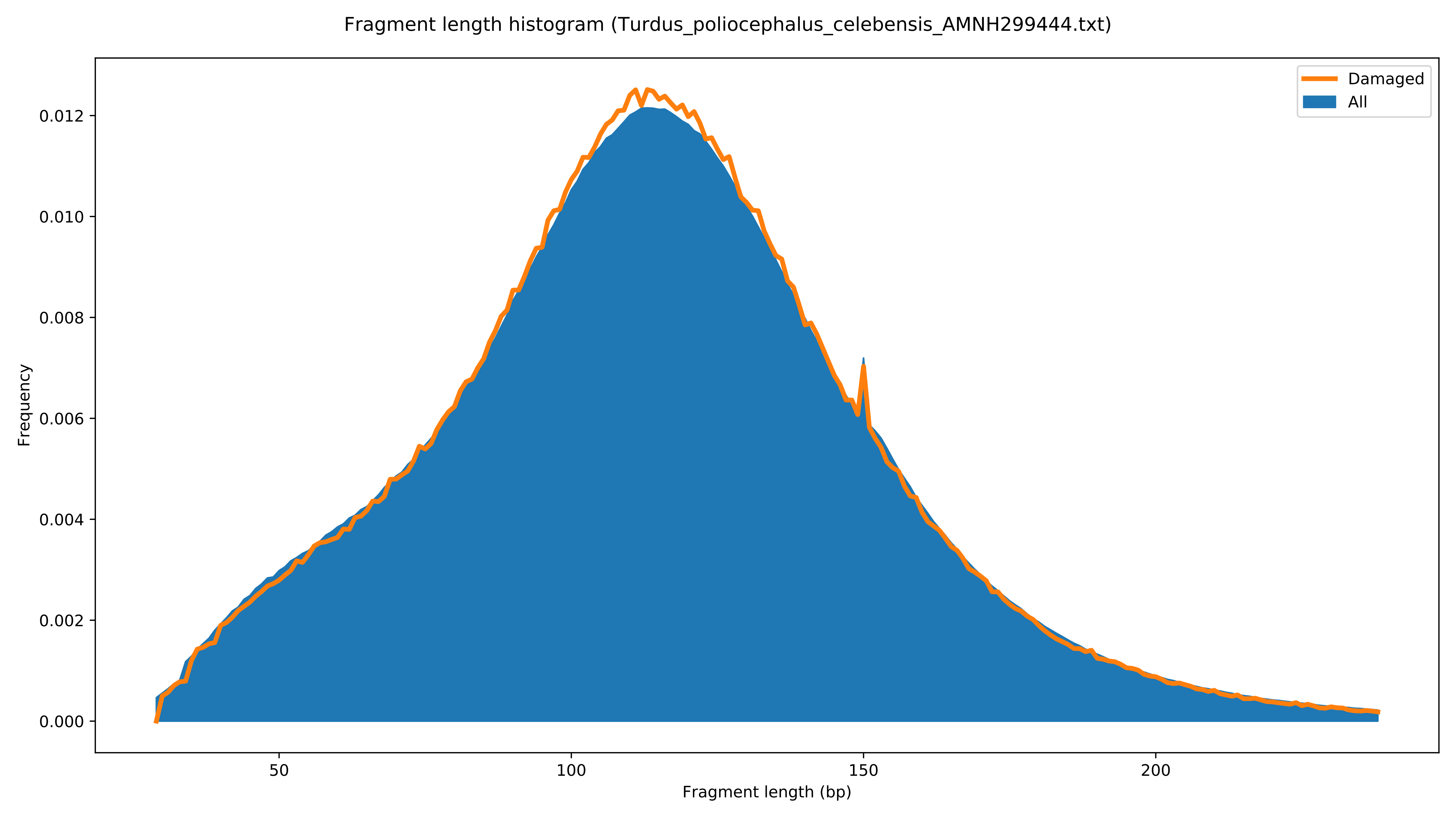
Fragment length histogram (Turdus_poliocephalus_celebensis_AMNH299444.txt)
Damaged
All
0.000
0.002
0.004
0.006
0.008
0.010
0.012
Frequency
50
100
150
200
Fragment length (bp)

Post-mortem damage patterns (Turdus_poliocephalus_deningeri_ZMUC148358.txt)

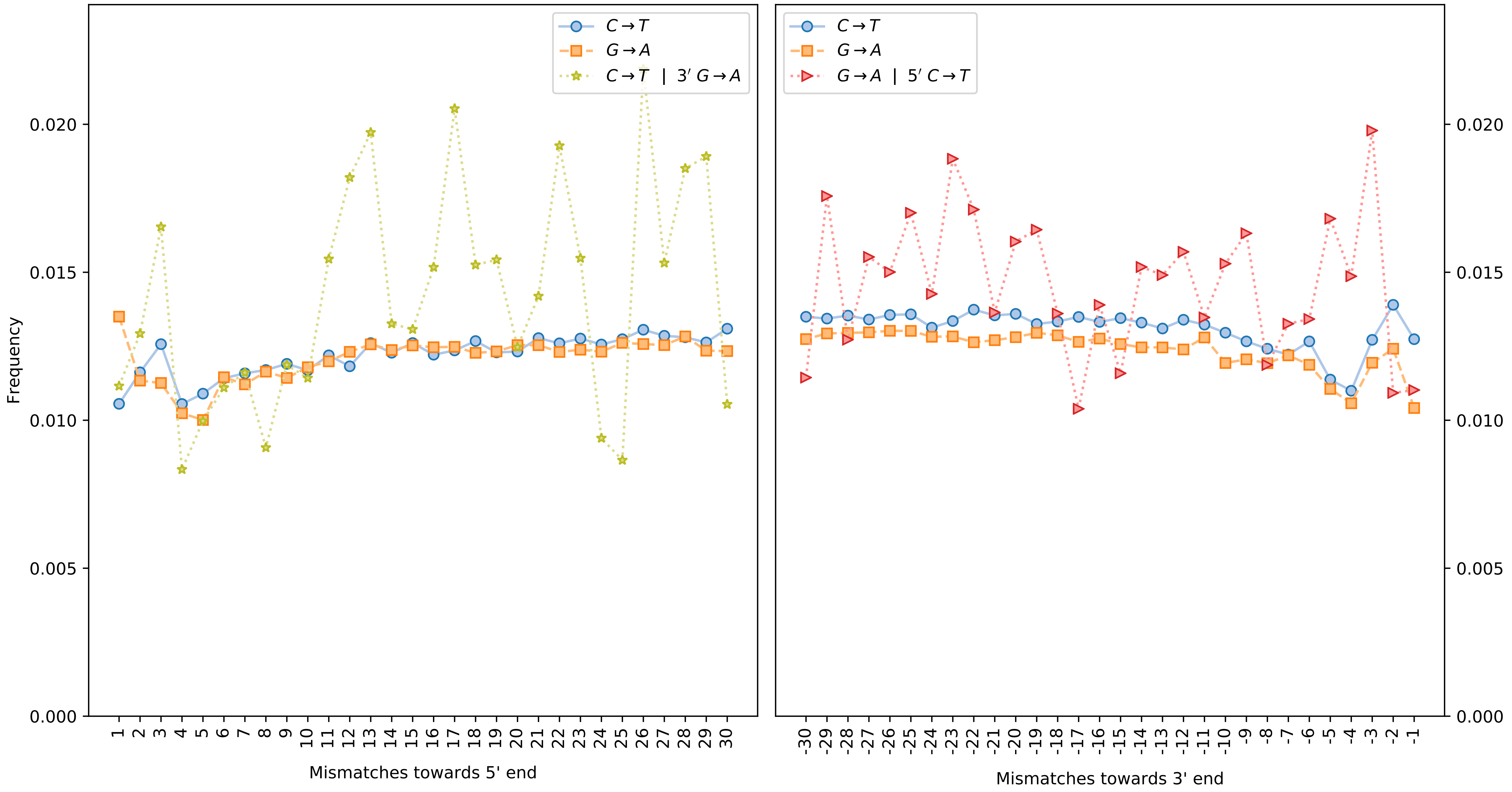

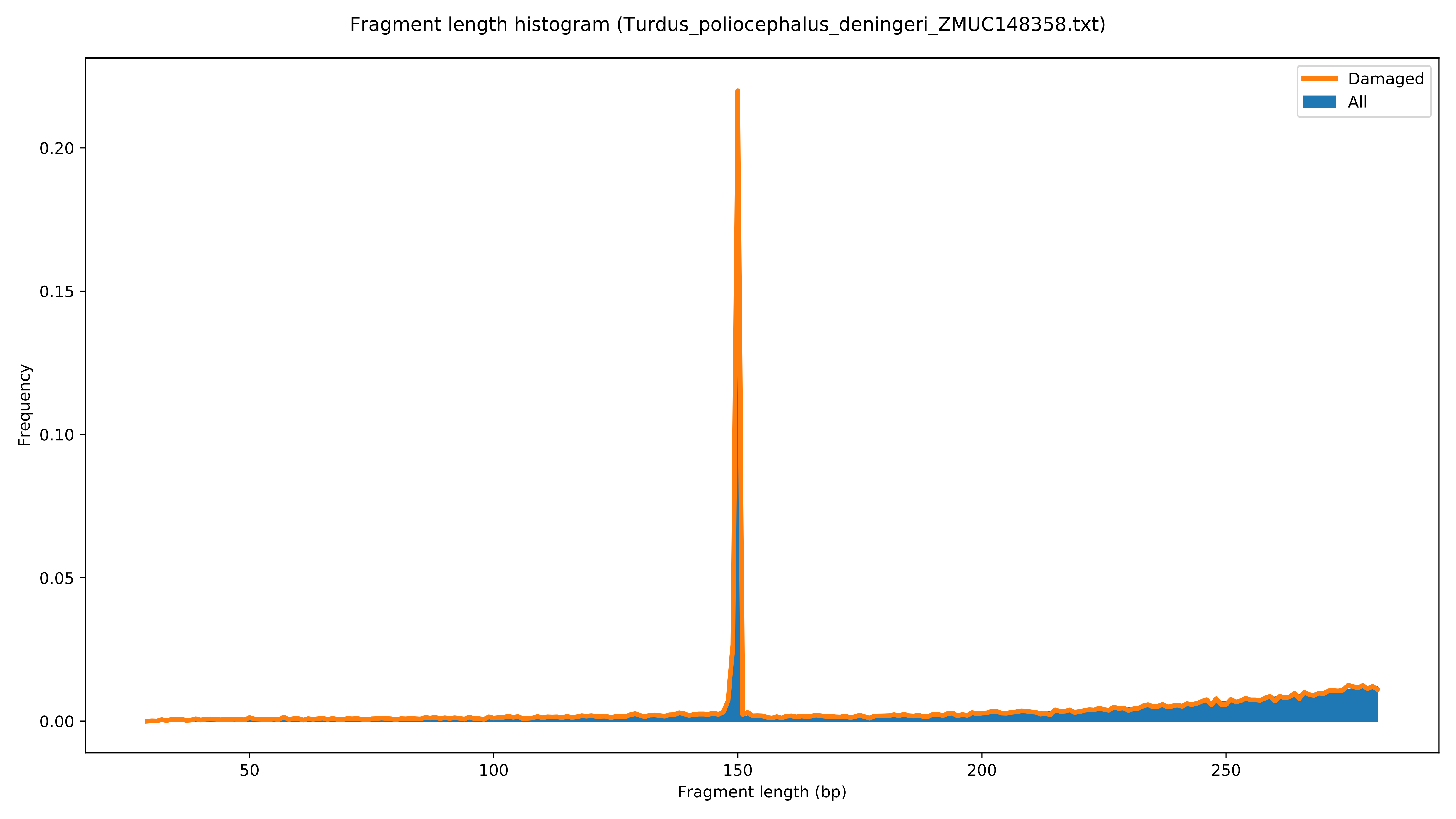
Fragment length histogram (Turdus_poliocephalus_deningeri_ZMUC148358.txt)
Damaged
All
Frequency
0.00
0.05
0.10
0.15
0.20
50
100
150
200
250
Fragment length (bp)

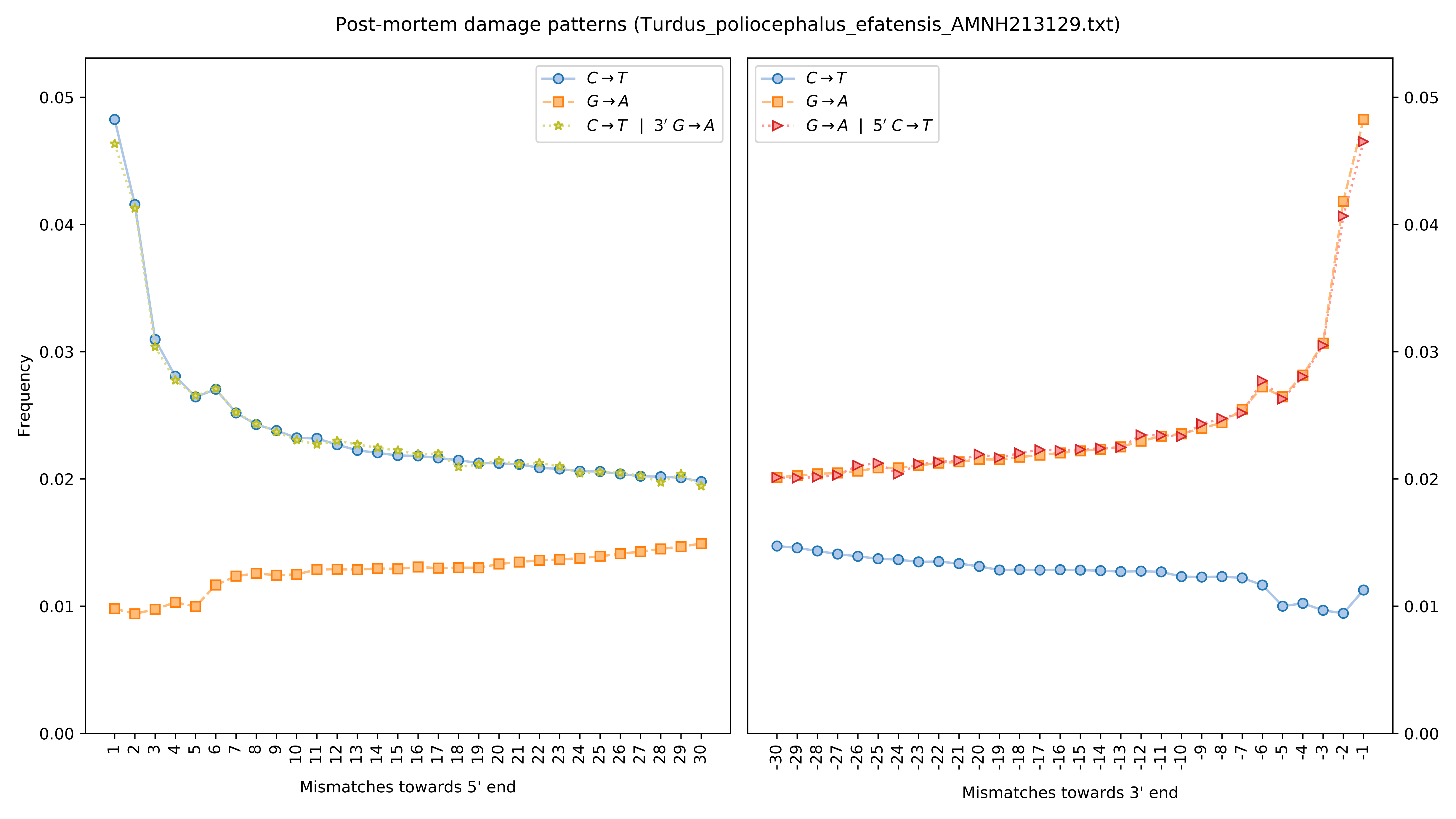

Post-mortem damage patterns (Turdus_poliocephalus_efatensis_AMNH213129.txt)
Frequency
Mismatches towards 5' end
Mismatches towards 3' end
C→T
G→A
C→T | 3′ G→A
G→A | 5′ C→T

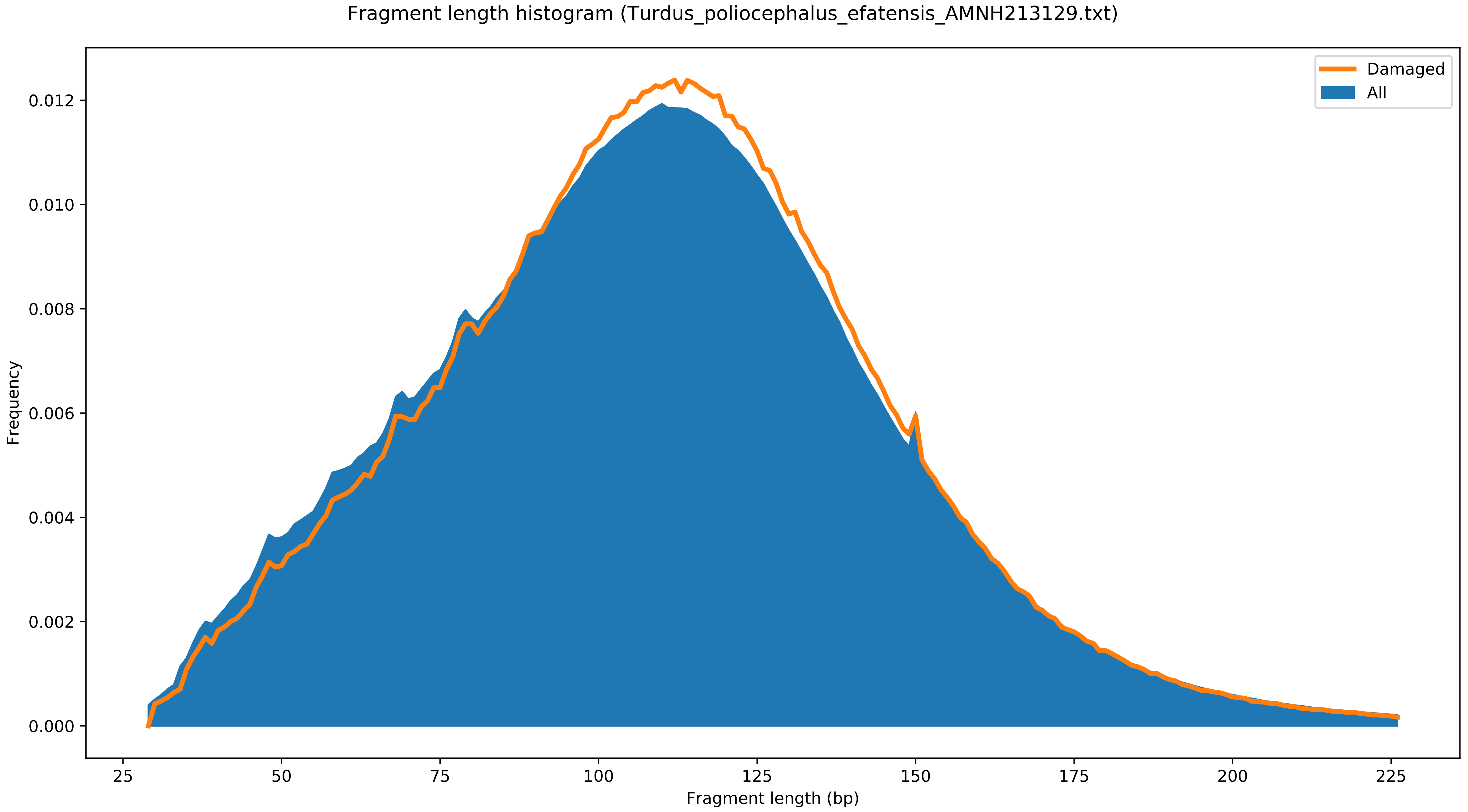

Fragment length histogram (Turdus_poliocephalus_efatensis_AMNH213129.txt)
Damaged
All
Frequency
0.000
0.002
0.004
0.006
0.008
0.010
0.012
25
50
75
100
125
150
175
200
225
Fragment length (bp)

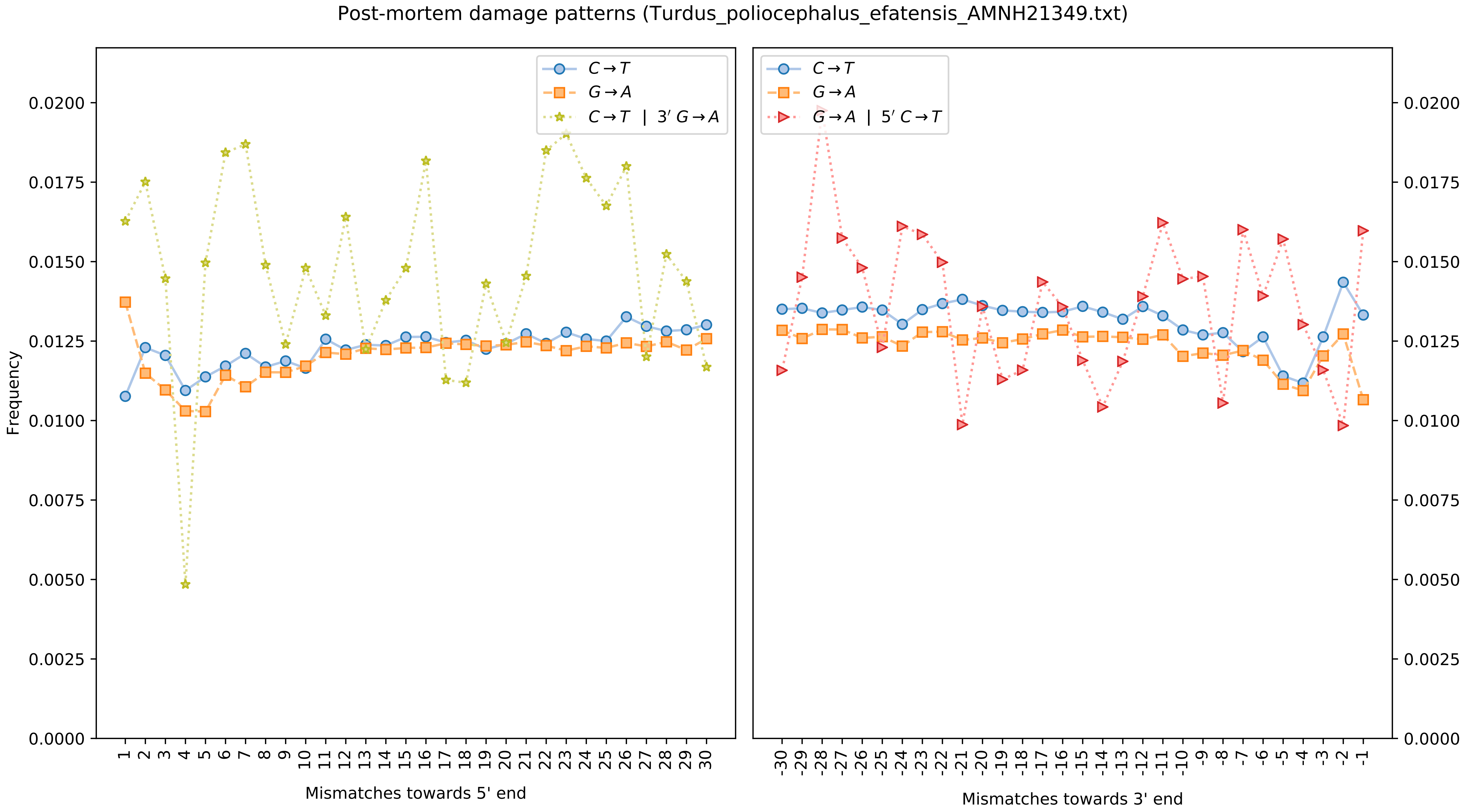

Post-mortem damage patterns (Turdus_poliocephalus_efatensis_AMNH21349.txt)
Frequency
Mismatches towards 5' end
Mismatches towards 3' end
C→T
G→A
C→T | 3′ G→A
G→A | 5′ C→T
0.0000
0.0025
0.0050
0.0075
0.0100
0.0125
0.0150
0.0175
0.0200

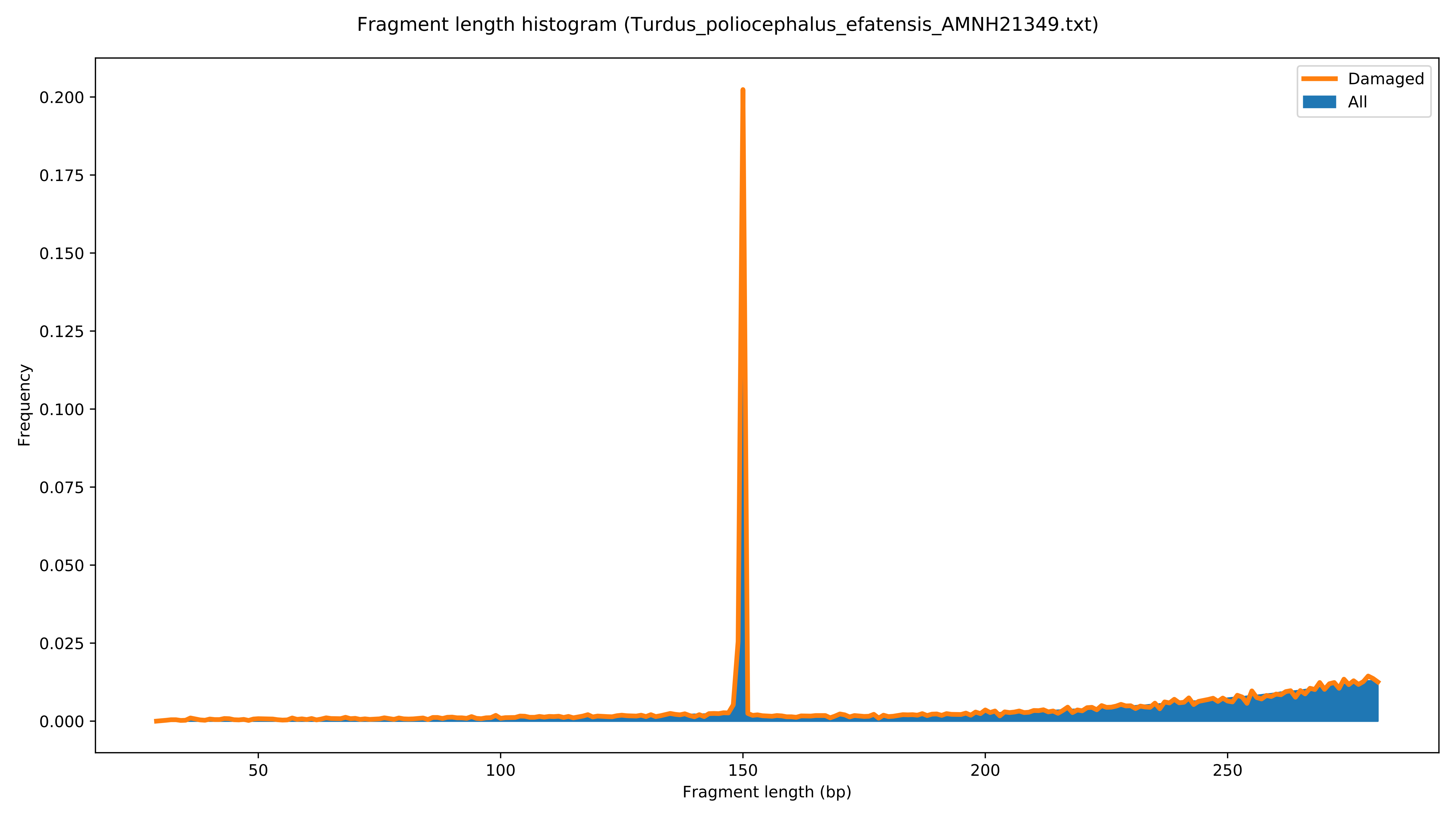

Fragment length histogram (Turdus_poliocephalus_efatensis_AMNH21349.txt)
Damaged
All
Frequency
0.000
0.025
0.050
0.075
0.100
0.125
0.150
0.175
0.200
50
100
150
200
250
Fragment length (bp)

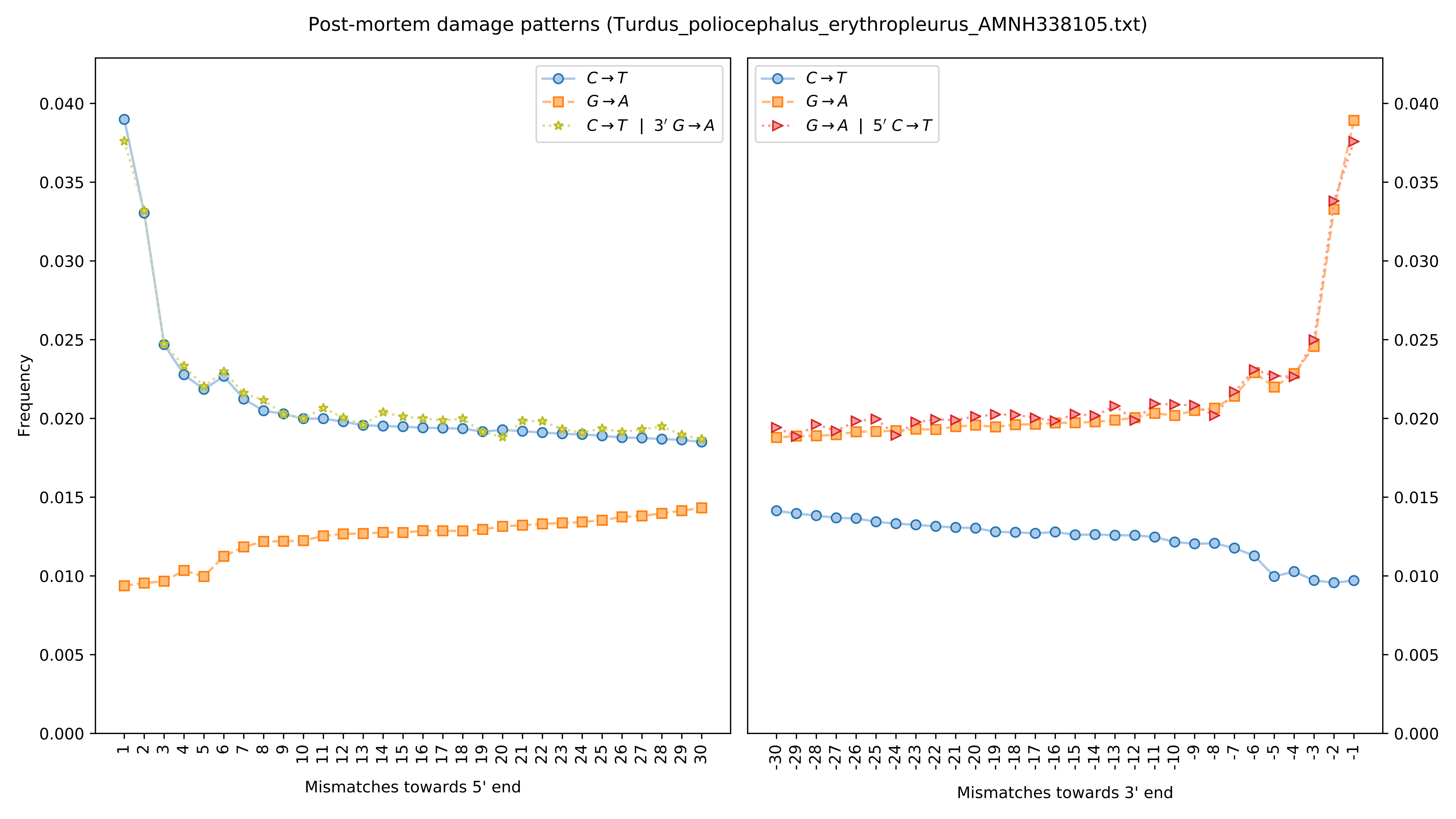

Post-mortem damage patterns (Turdus_poliocephalus_erythropleurus_AMNH338105.txt)
$C \rightarrow T$
$G \rightarrow A$
$C \rightarrow T$ | $3' G \rightarrow A$
$G \rightarrow A$ | $5' C \rightarrow T$
Frequency
Mismatches towards 5' end
Mismatches towards 3' end

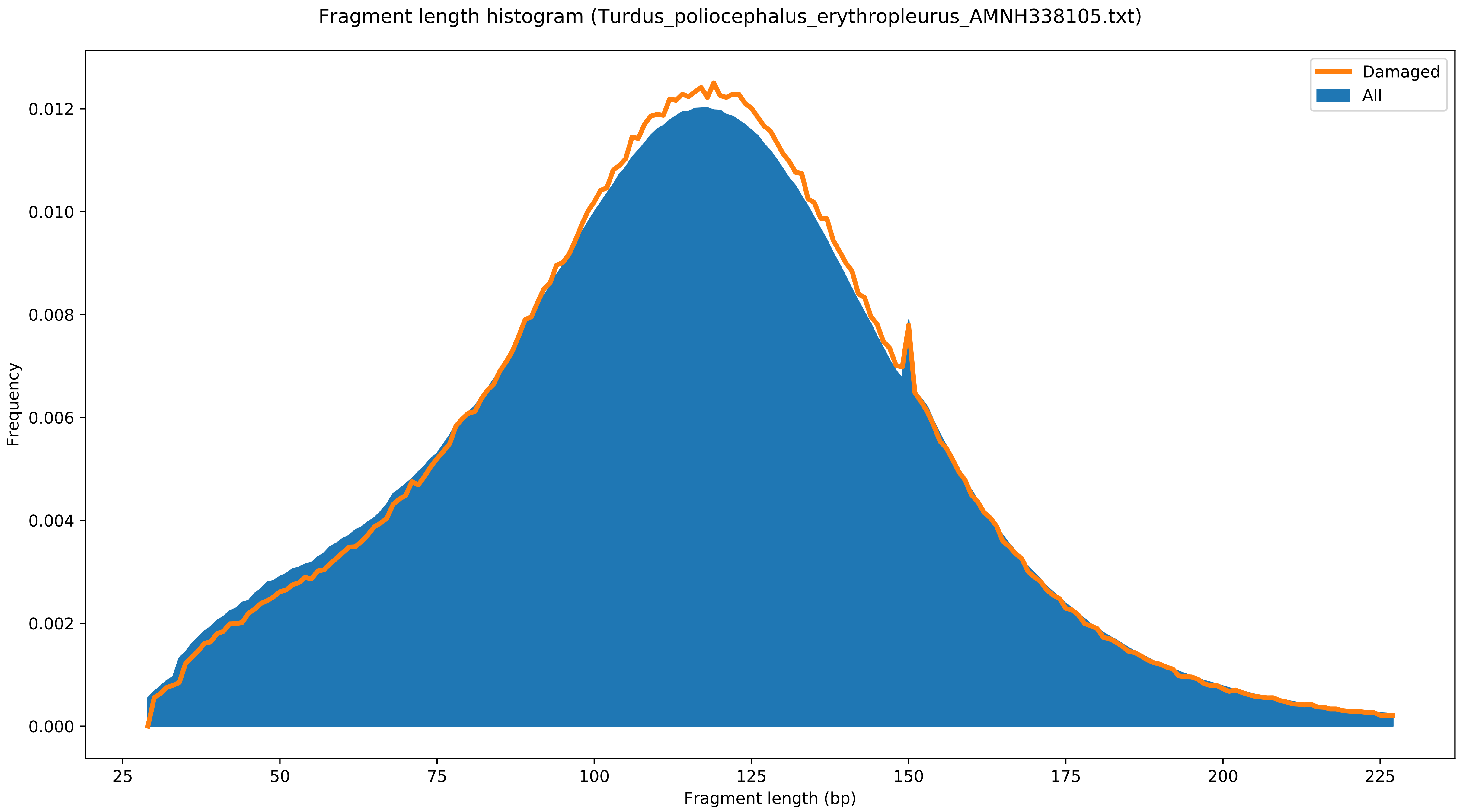

Fragment length histogram (Turdus_poliocephalus_erythropleurus_AMNH338105.txt)
Damaged
All
Frequency
0.000
0.002
0.004
0.006
0.008
0.010
0.012
25
50
75
100
125
150
175
200
225
Fragment length (bp)

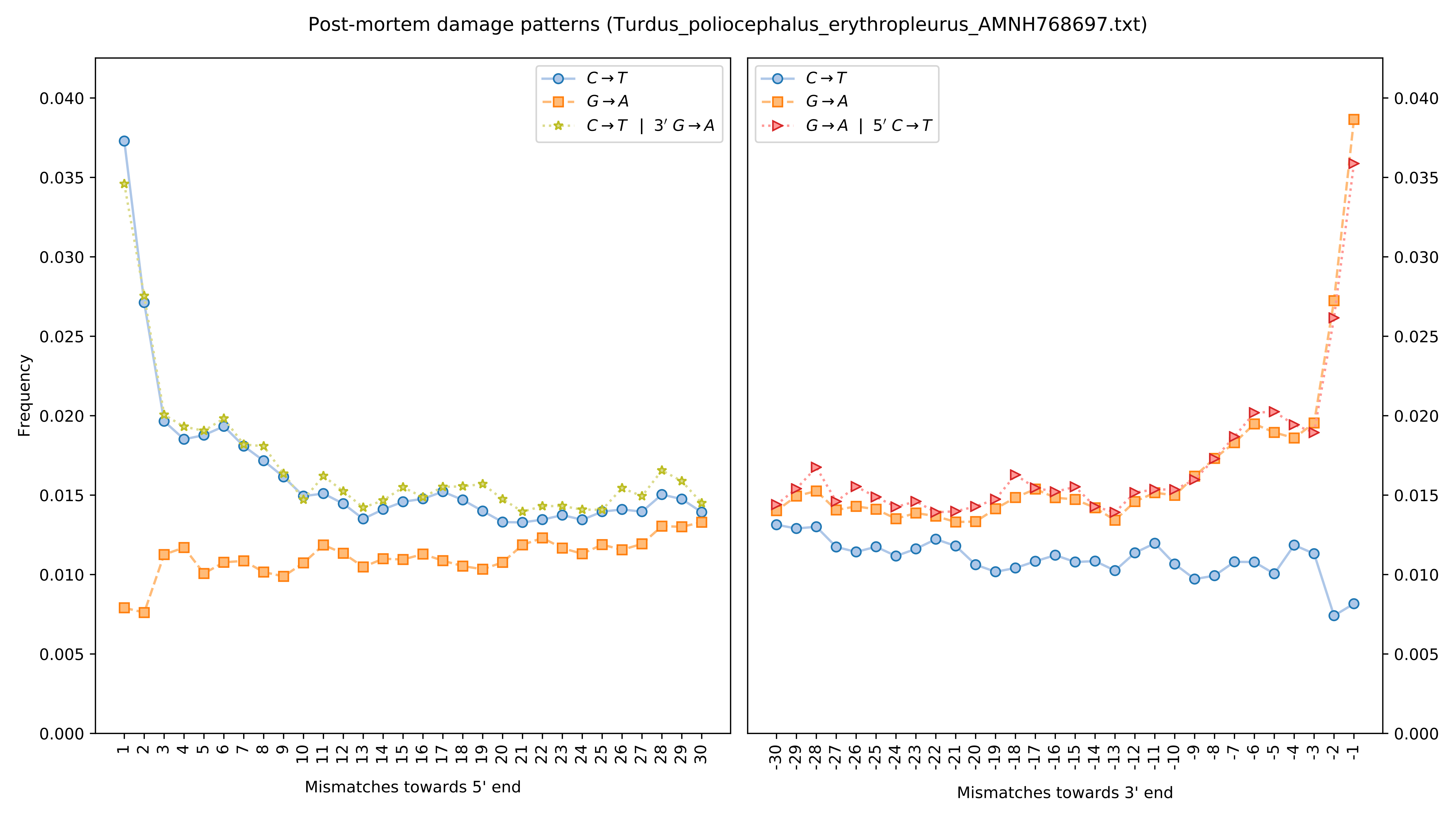
Post-mortem damage patterns (Turdus_poliocephalus_erythropleurus_AMNH768697.txt)
Frequency
C→T
G→A
C→T | 3′ G→A
G→A | 5′ C→T
0.000
0.005
0.010
0.015
0.020
0.025
0.030
0.035
0.040
1 2 3 4 5 6 7 8 9 10 11 12 13 14 15 16 17 18 19 20 21 22 23 24 25 26 27 28 29 30
-30 -29 -28 -27 -26 -25 -24 -23 -22 -21 -20 -19 -18 -17 -16 -15 -14 -13 -12 -11 -10 -9 -8 -7 -6 -5 -4 -3 -2 -1
Mismatches towards 5' end
Mismatches towards 3' end

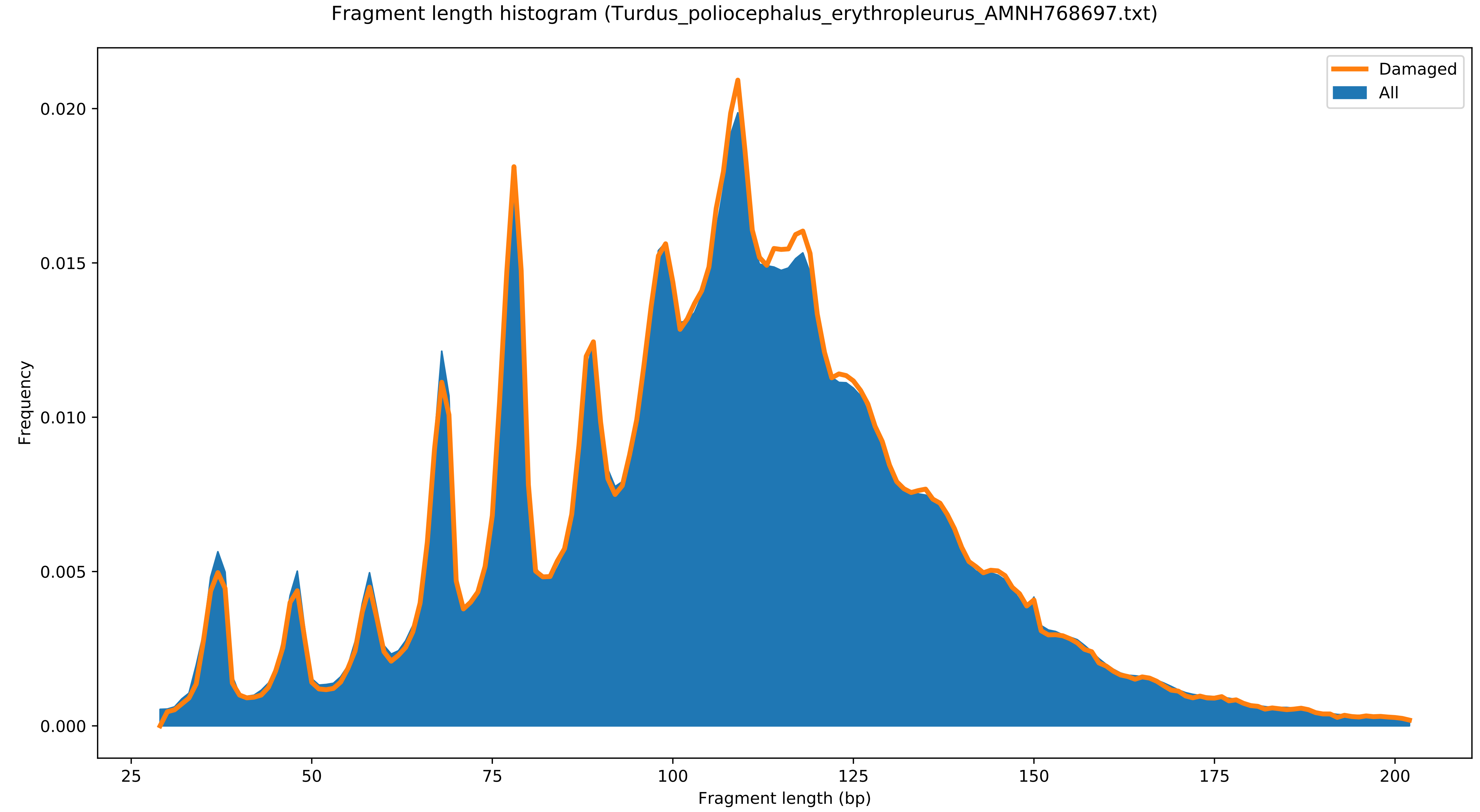

Fragment length histogram (Turdus_poliocephalus_erythropleurus_AMNH768697.txt)
Damaged
All
Frequency
0.000
0.005
0.010
0.015
0.020
25
50
75
100
125
150
175
200
Fragment length (bp)

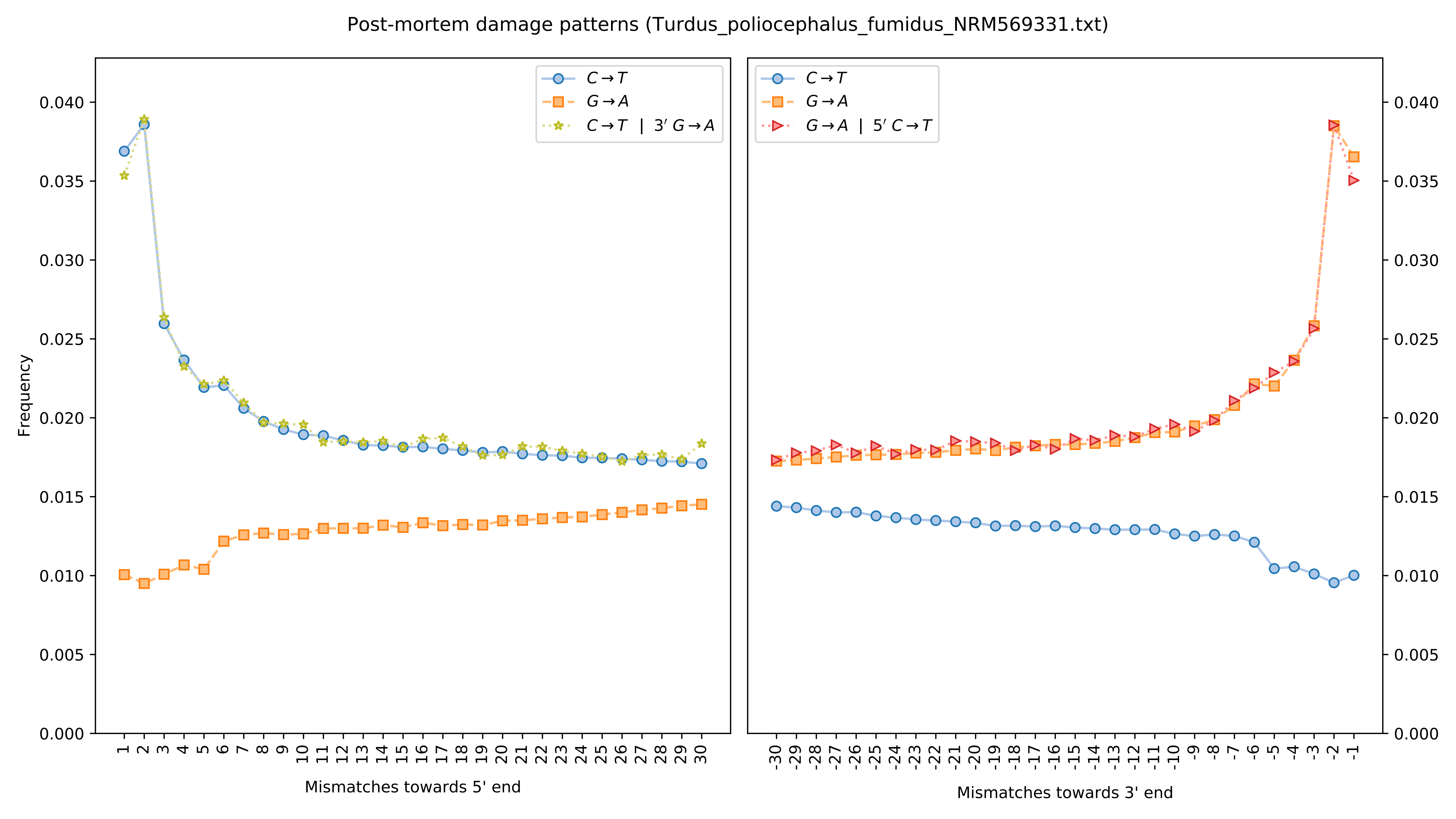
Post-mortem damage patterns (Turdus_poliocephalus_fumidus_NRM569331.txt)
Frequency
Mismatches towards 5' end
Mismatches towards 3' end
$C \rightarrow T$
$G \rightarrow A$
$C \rightarrow T$ | $3' G \rightarrow A$
$G \rightarrow A$ | $5' C \rightarrow T$

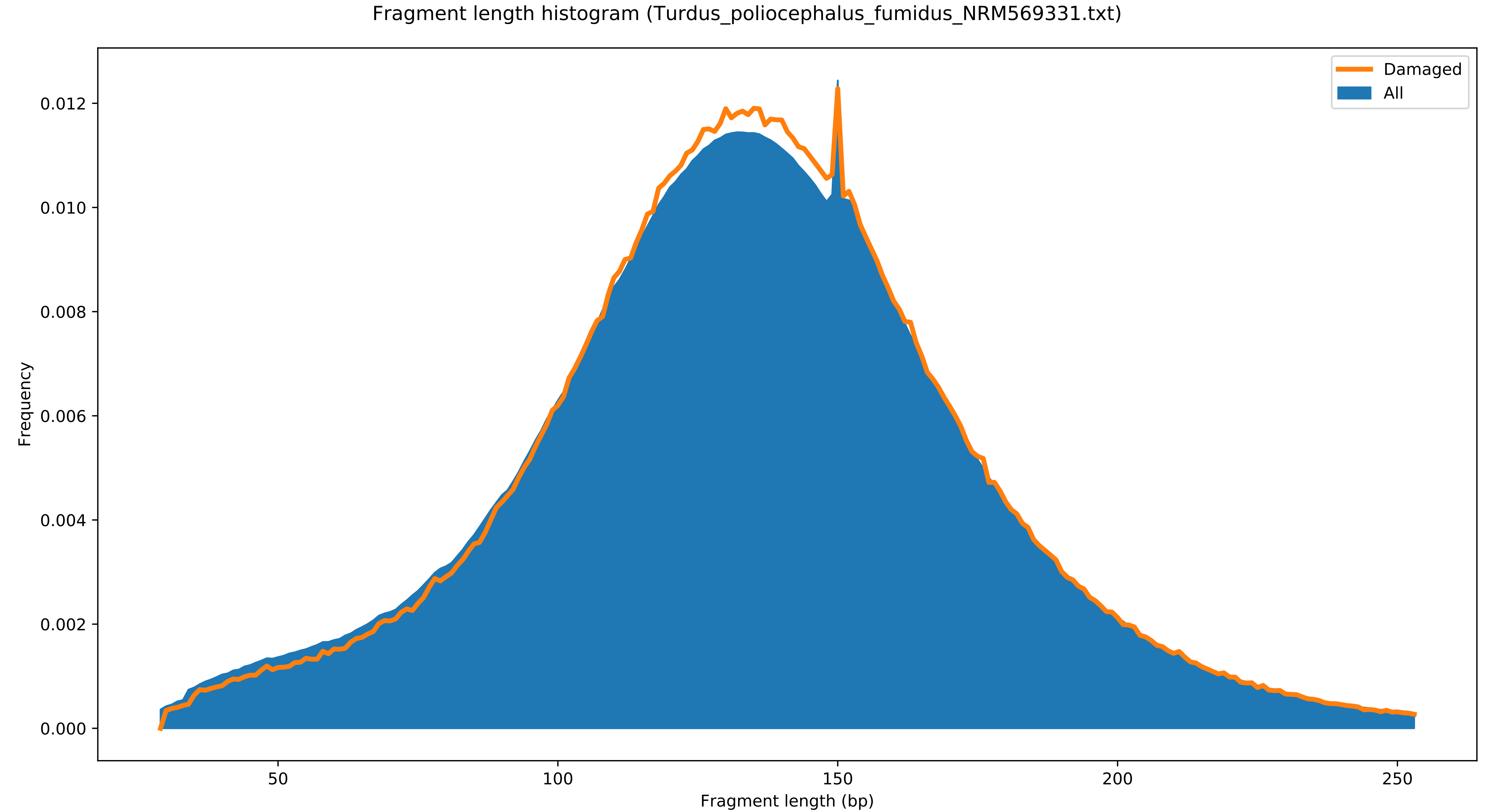

Fragment length histogram (Turdus_poliocephalus_fumidus_NRM569331.txt)
Damaged
All
Frequency
0.000
0.002
0.004
0.006
0.008
0.010
0.012
50
100
150
200
250
Fragment length (bp)

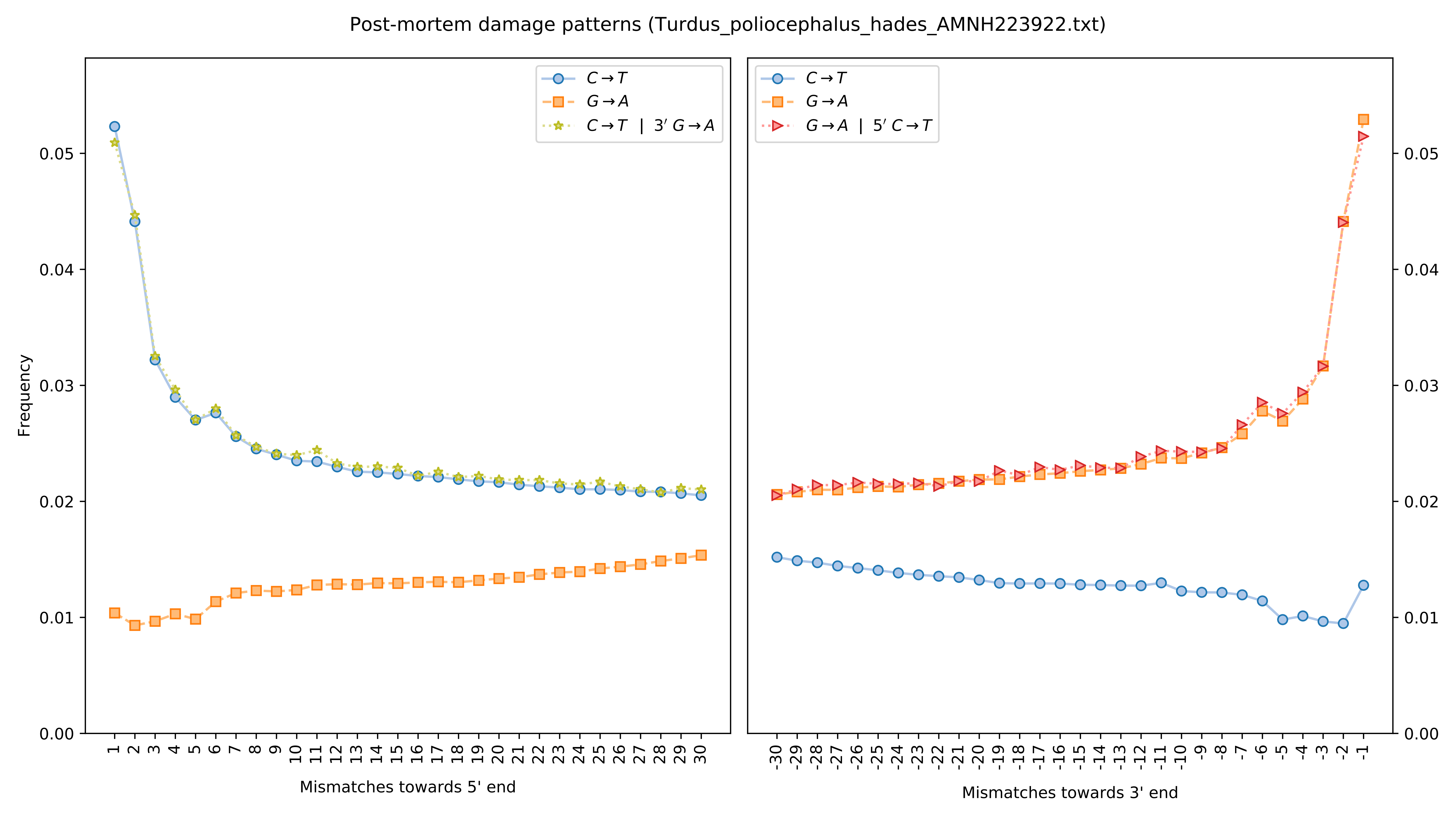

Post-mortem damage patterns (Turdus_poliocephalus_hades_AMNH223922.txt)
C→T
G→A
C→T | 3′ G→A
G→A | 5′ C→T
Frequency
Mismatches towards 5' end
Mismatches towards 3' end

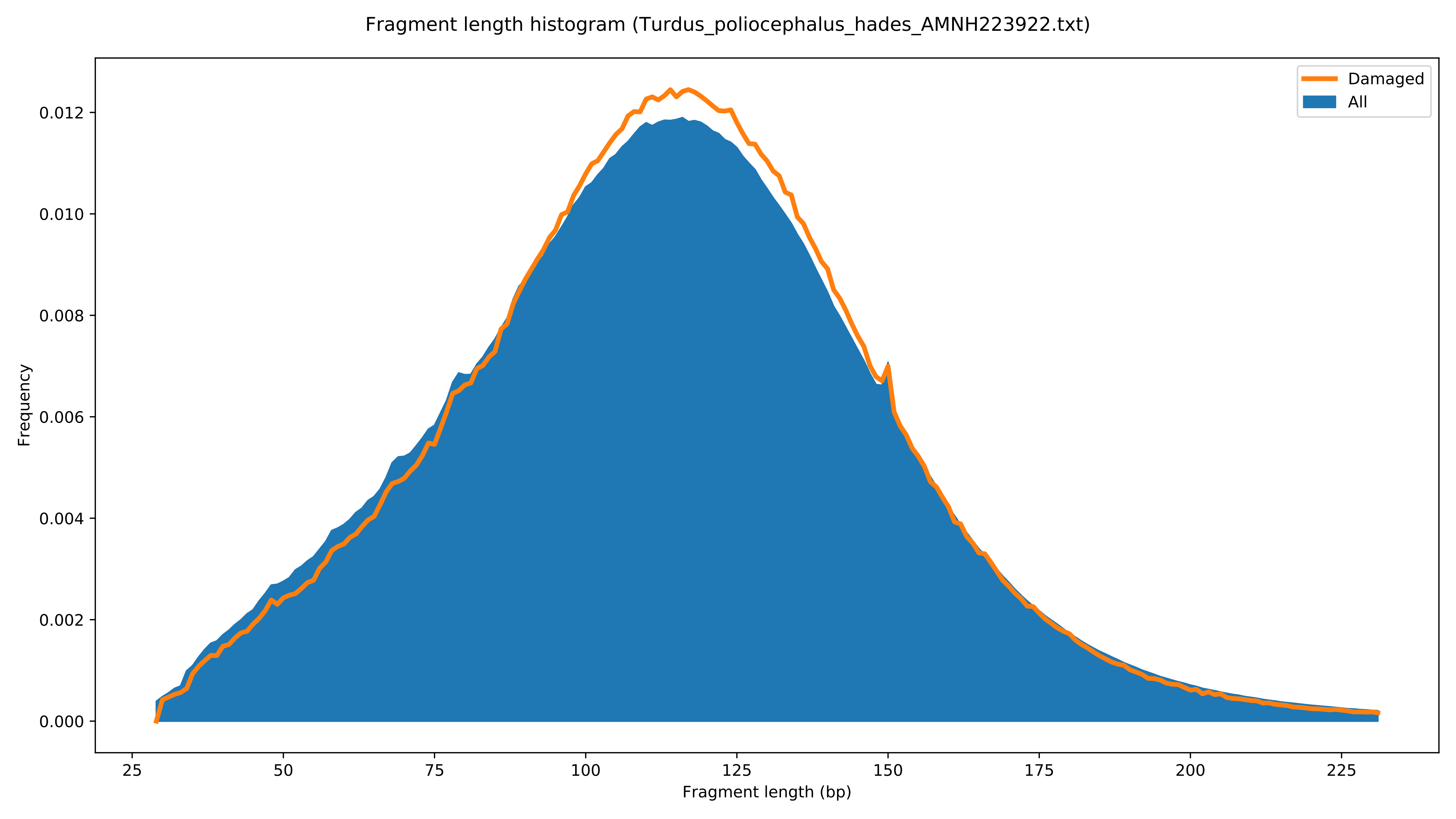
Fragment length histogram (Turdus_poliocephalus_hades_AMNH223922.txt)
Damaged
All
Frequency
Fragment length (bp)
0.000
0.002
0.004
0.006
0.008
0.010
0.012
25
50
75
100
125
150
175
200
225

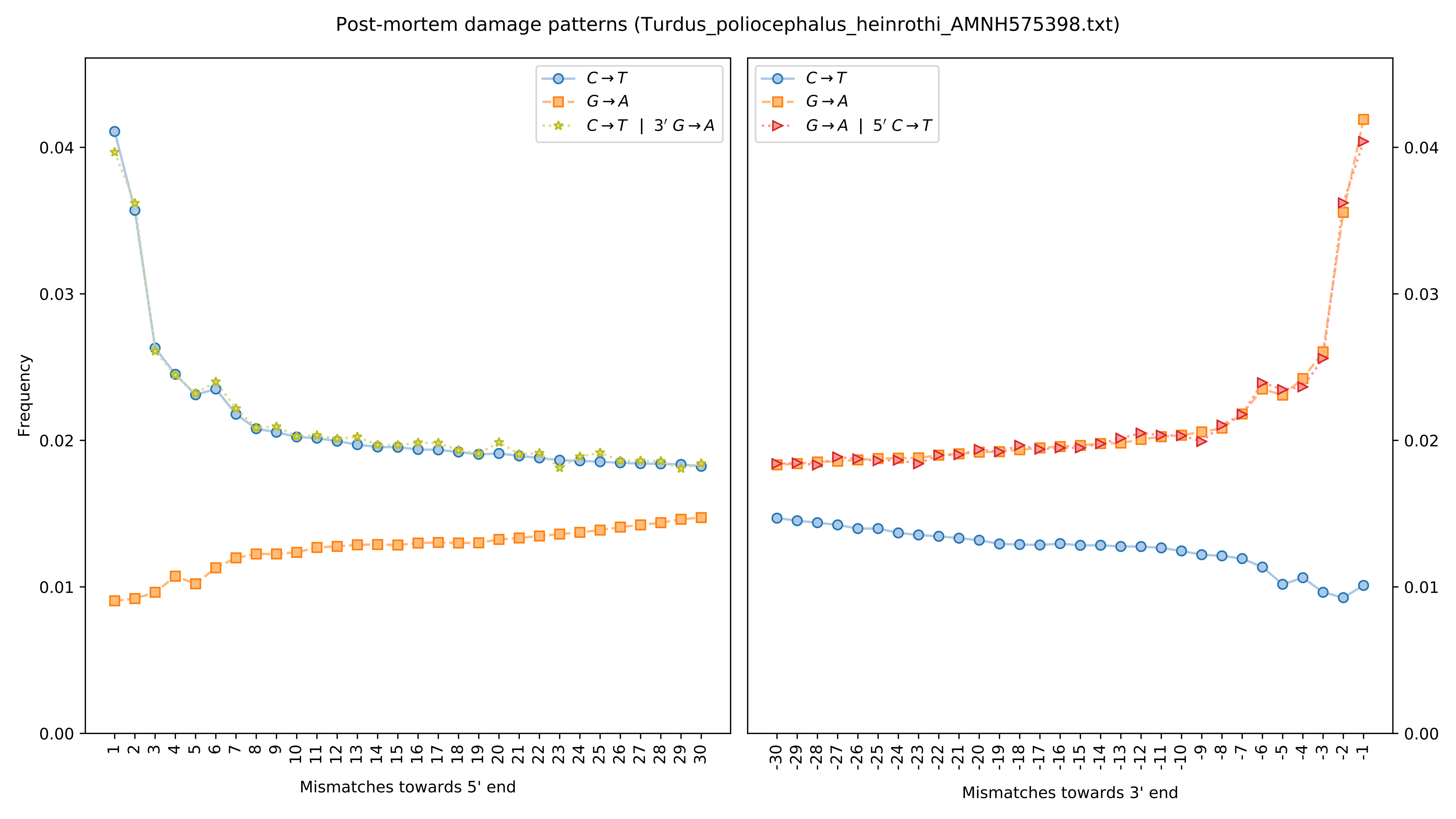

Post-mortem damage patterns (Turdus_poliocephalus_heinrothi_AMNH575398.txt)
Frequency
Mismatches towards 5' end
Mismatches towards 3' end
C→T
G→A
C→T | 3' G→A
G→A | 5' C→T

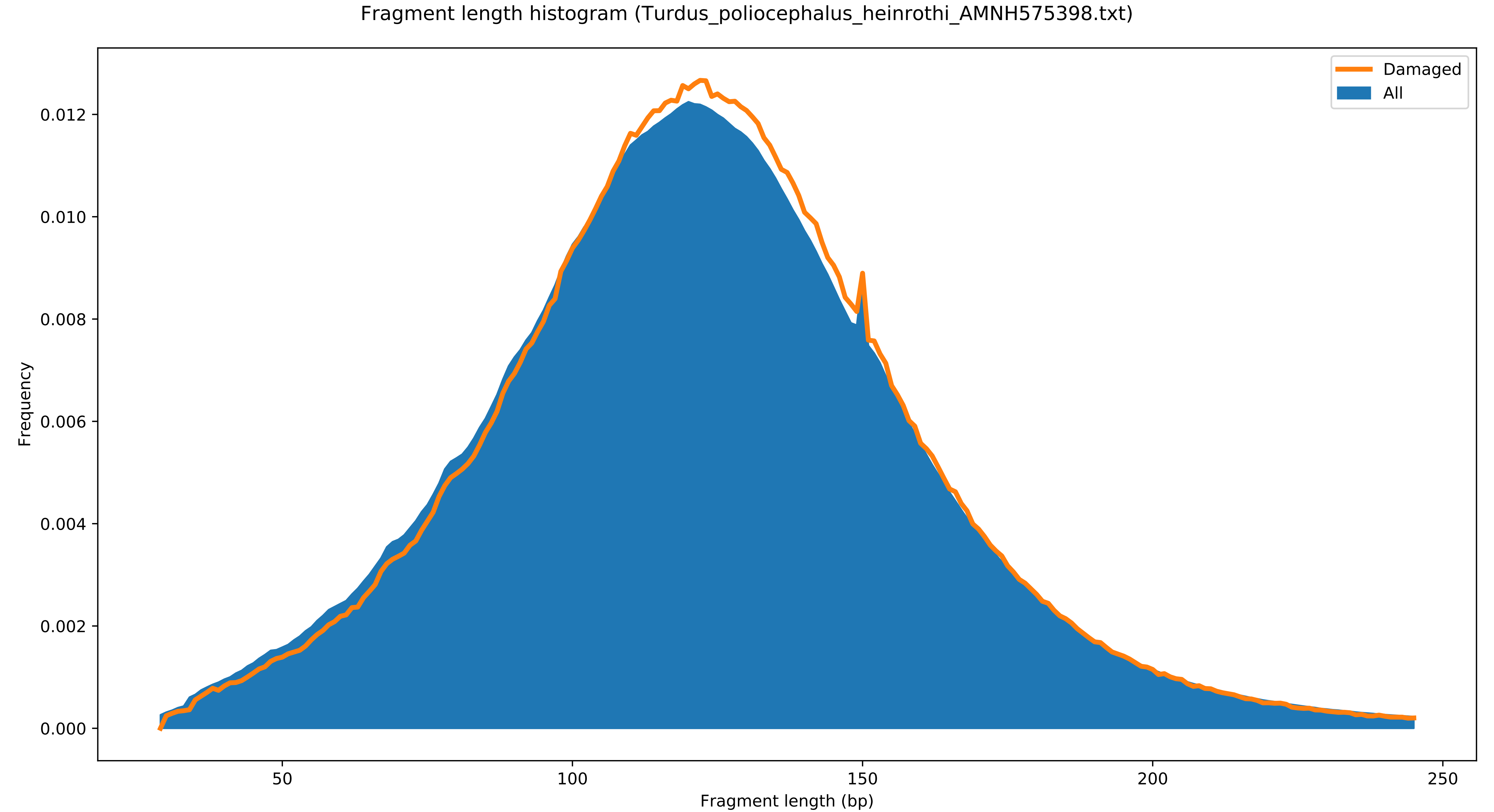

Fragment length histogram (Turdus_poliocephalus_heinrothi_AMNH575398.txt)
Damaged
All
Frequency
0.000
0.002
0.004
0.006
0.008
0.010
0.012
50
100
150
200
250
Fragment length (bp)

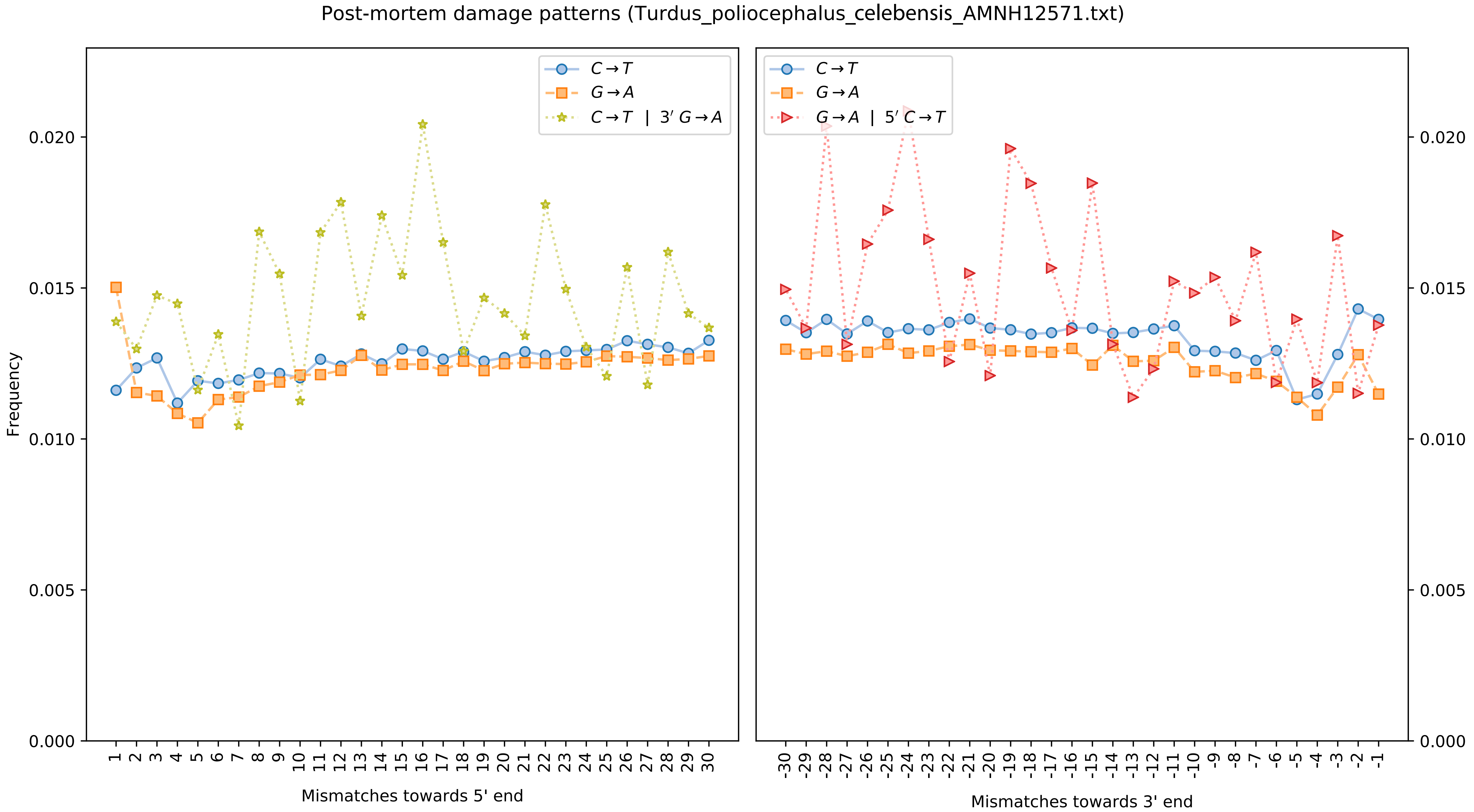

Post-mortem damage patterns (Turdus_poliocephalus_celebensis_AMNH12571.txt)
C→T
G→A
C→T | 3′ G→A
G→A | 5′ C→T
Frequency
Mismatches towards 5' end
Mismatches towards 3' end

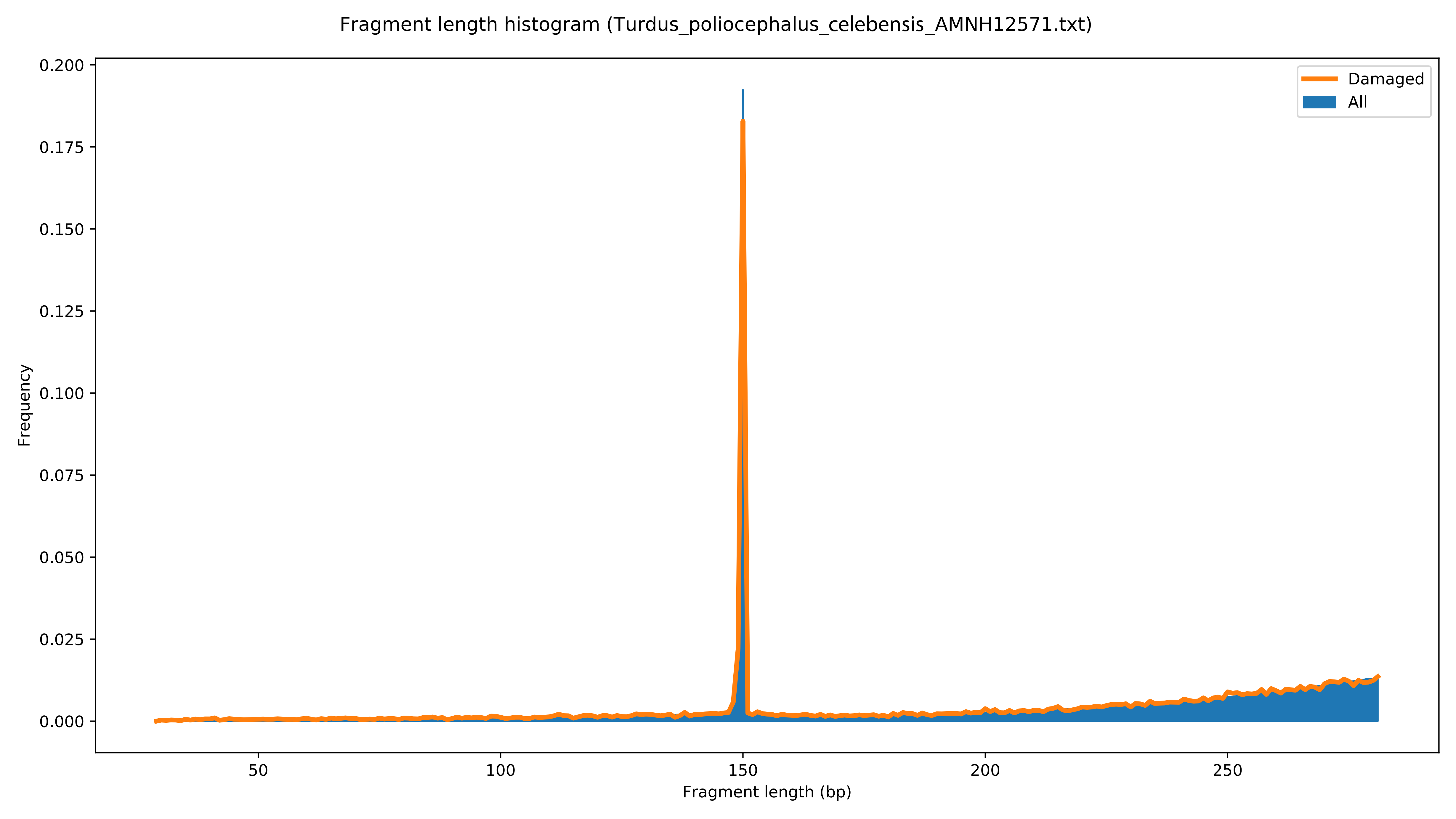

Fragment length histogram (Turdus_poliocephalus_celebensis_AMNH12571.txt)
Damaged
All
Frequency
Fragment length (bp)
0.000
0.025
0.050
0.075
0.100
0.125
0.150
0.175
0.200
50
100
150
200
250

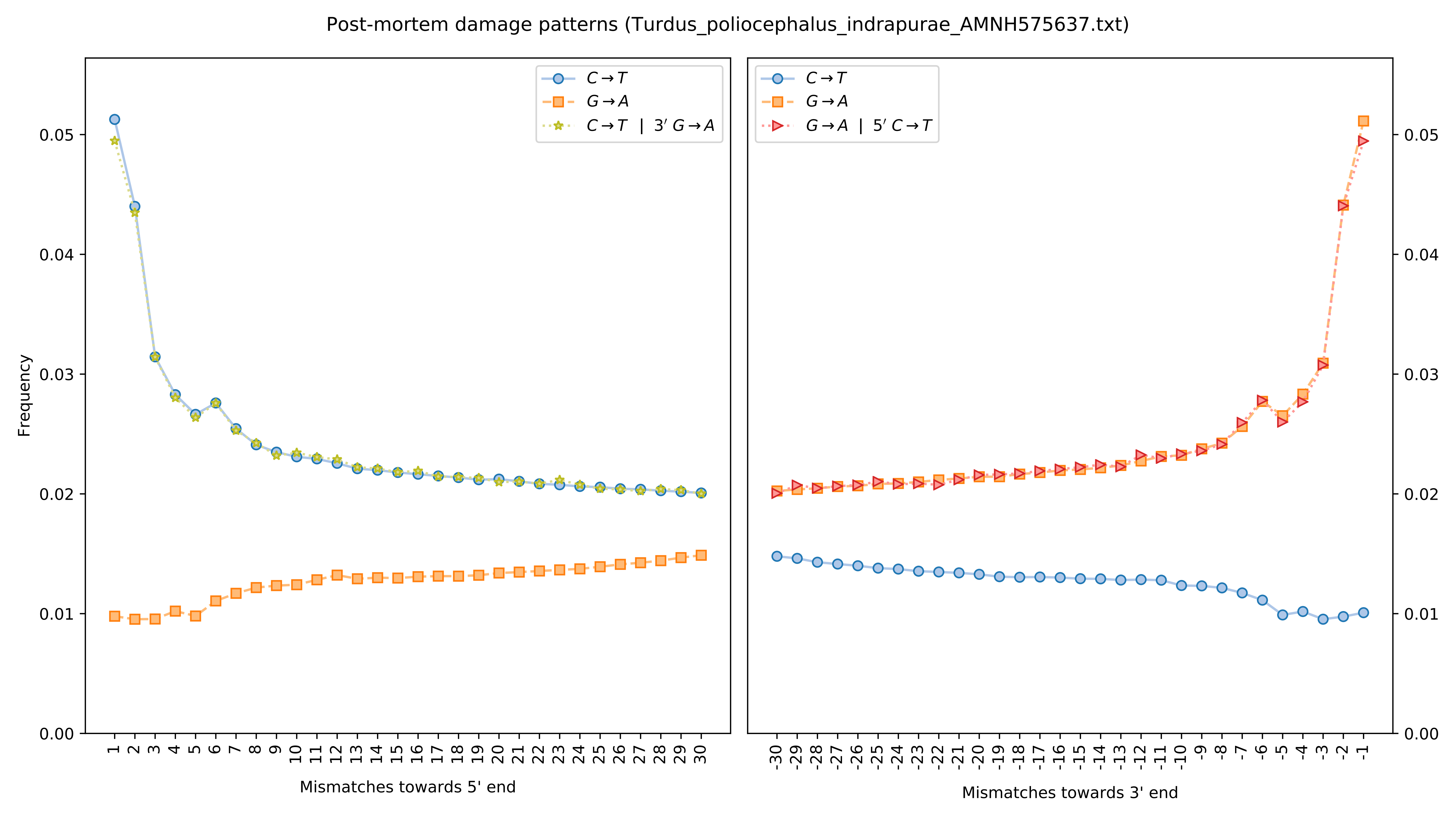
Post-mortem damage patterns (Turdus_poliocephalus_indrapurae_AMNH575637.txt)
Frequency
Mismatches towards 5' end
Mismatches towards 3' end
C→T
G→A
C→T | 3′ G→A
G→A | 5′ C→T
0.00
0.01
0.02
0.03
0.04
0.05

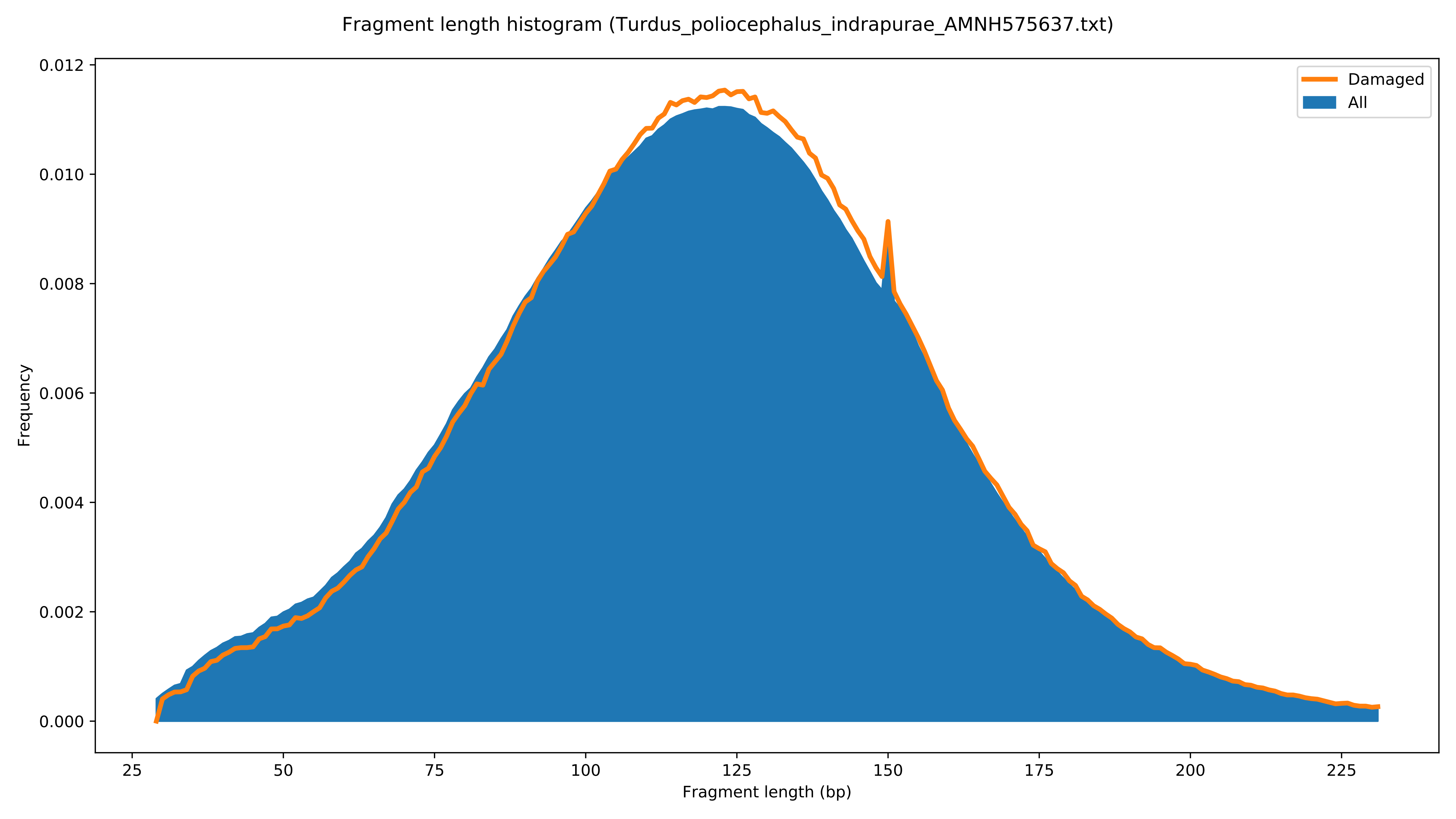

Fragment length histogram (Turdus_poliocephalus_indrapurae_AMNH575637.txt)
Damaged
All
Frequency
0.000
0.002
0.004
0.006
0.008
0.010
0.012
25
50
75
100
125
150
175
200
225
Fragment length (bp)

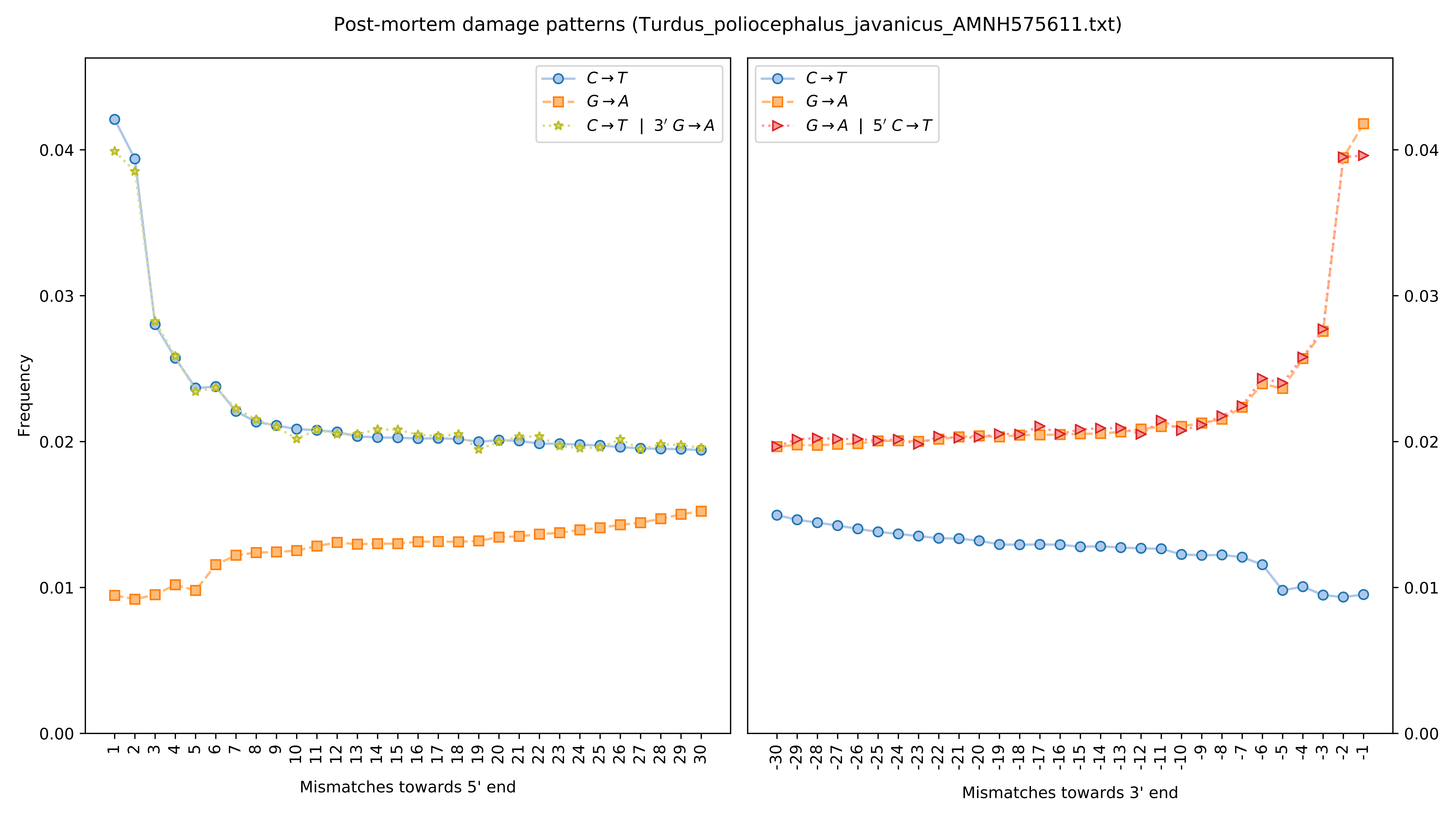
Post-mortem damage patterns (Turdus_poliocephalus_javanicus_AMNH575611.txt)
C→T
G→A
C→T | 3′ G→A
G→A | 5′ C→T
Frequency
Mismatches towards 5' end
Mismatches towards 3' end

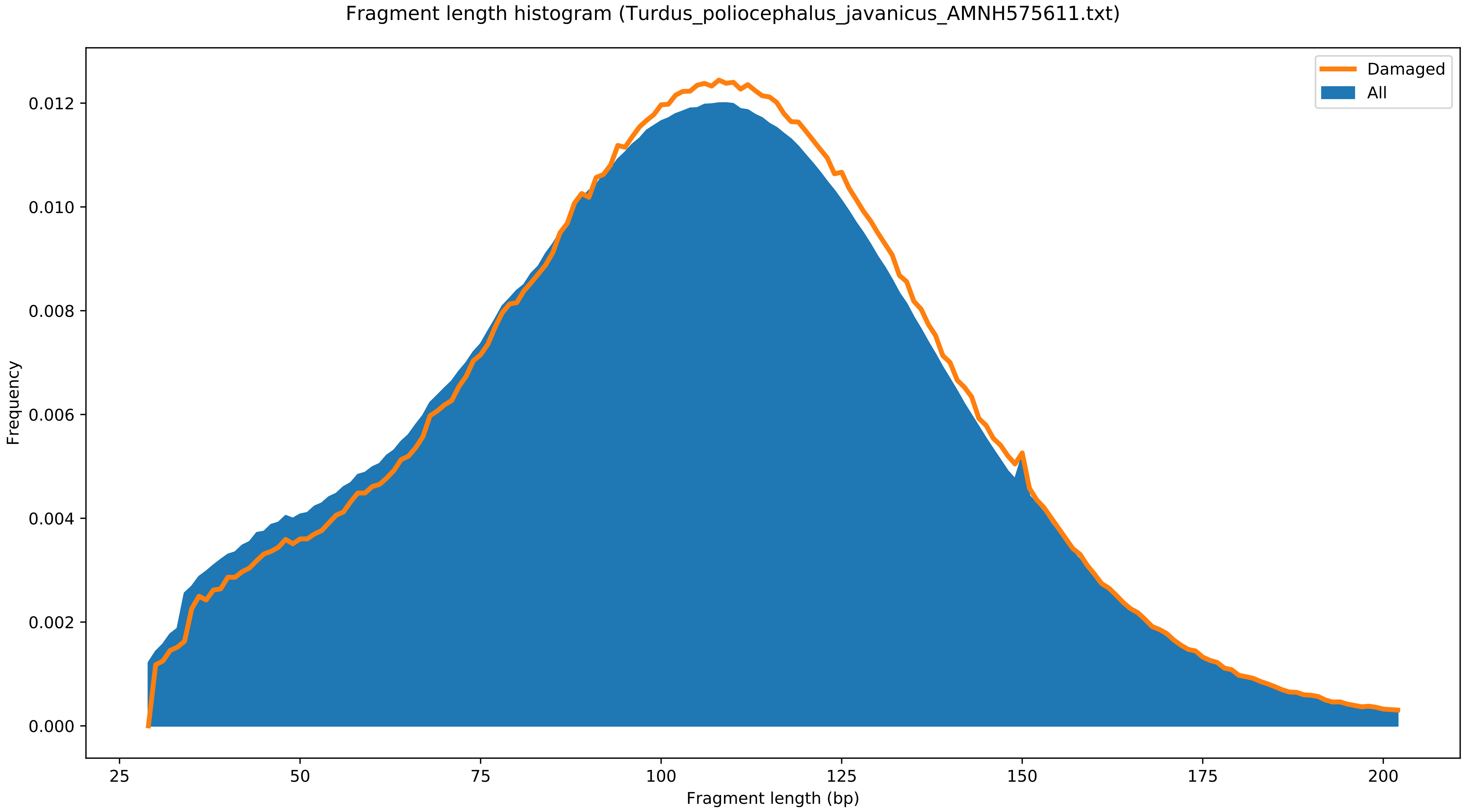

Fragment length histogram (Turdus_poliocephalus_javanicus_AMNH575611.txt)
Damaged
All
Frequency
0.000
0.002
0.004
0.006
0.008
0.010
0.012
25
50
75
100
125
150
175
200
Fragment length (bp)

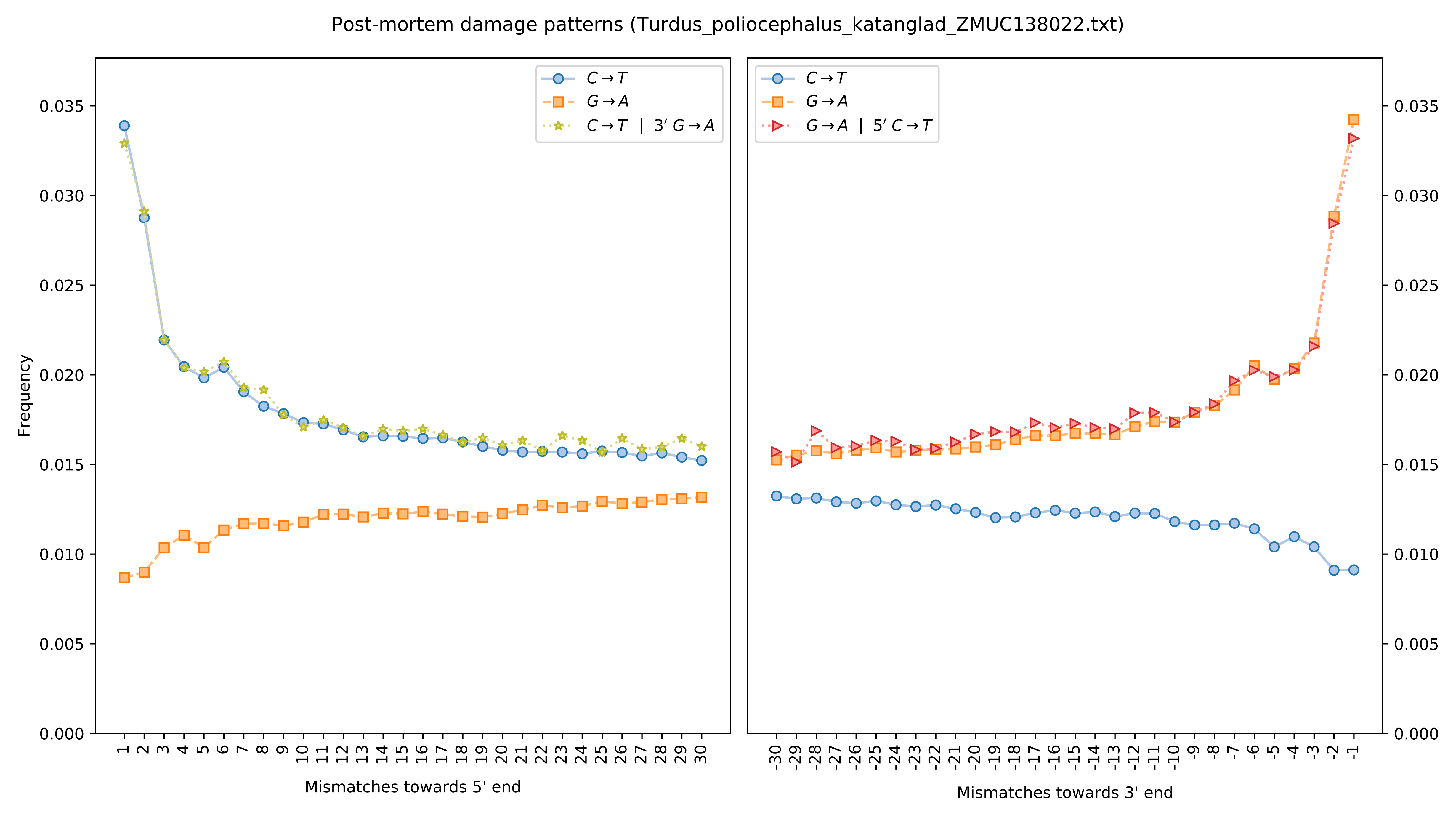
Post-mortem damage patterns (Turdus_poliocephalus_katanglad_ZMUC138022.txt)
Frequency
Mismatches towards 5' end
Mismatches towards 3' end
C→T
G→A
C→T | 3′ G→A
G→A | 5′ C→T
0.000
0.005
0.010
0.015
0.020
0.025
0.030
0.035

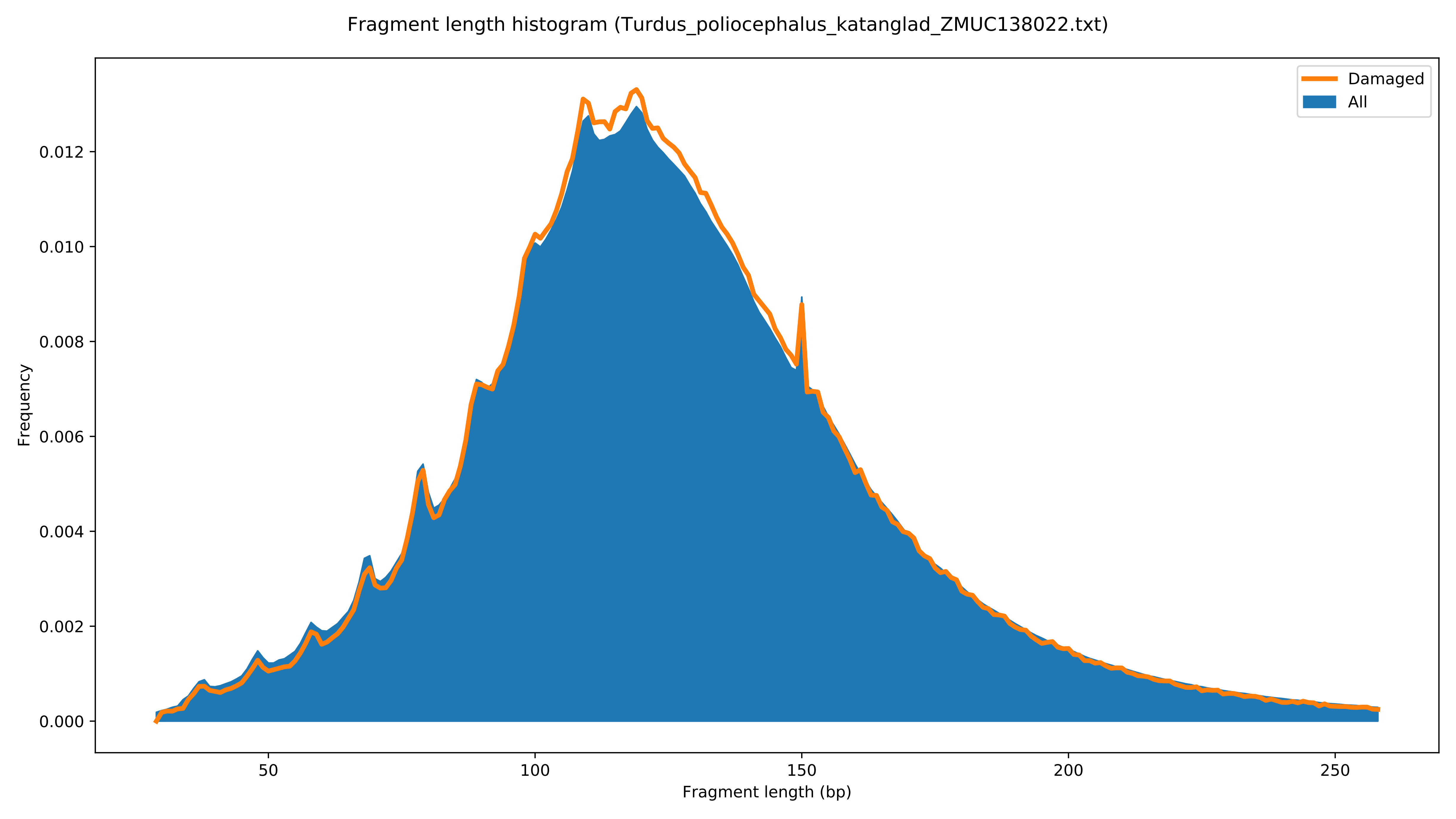

Fragment length histogram (Turdus_poliocephalus_katanglad_ZMUC138022.txt)
Damaged
All
Frequency
0.000
0.002
0.004
0.006
0.008
0.010
0.012
50
100
150
200
250
Fragment length (bp)

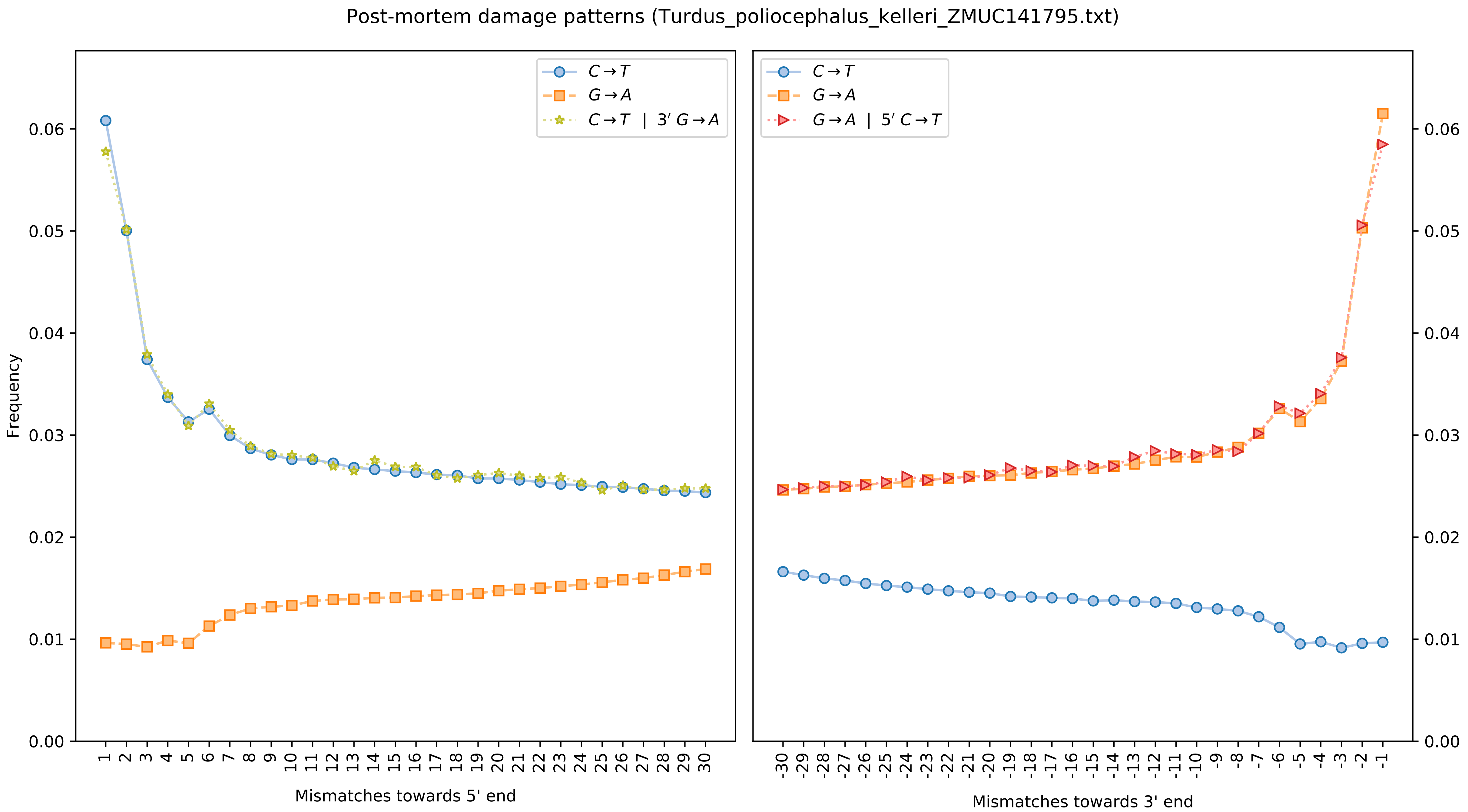
Post-mortem damage patterns (Turdus_poliocephalus_kelleri_ZMUC141795.txt)
C→T
G→A
C→T | 3′ G→A
G→A | 5′ C→T
Frequency
0.00
0.01
0.02
0.03
0.04
0.05
0.06
Mismatches towards 5' end
Mismatches towards 3' end

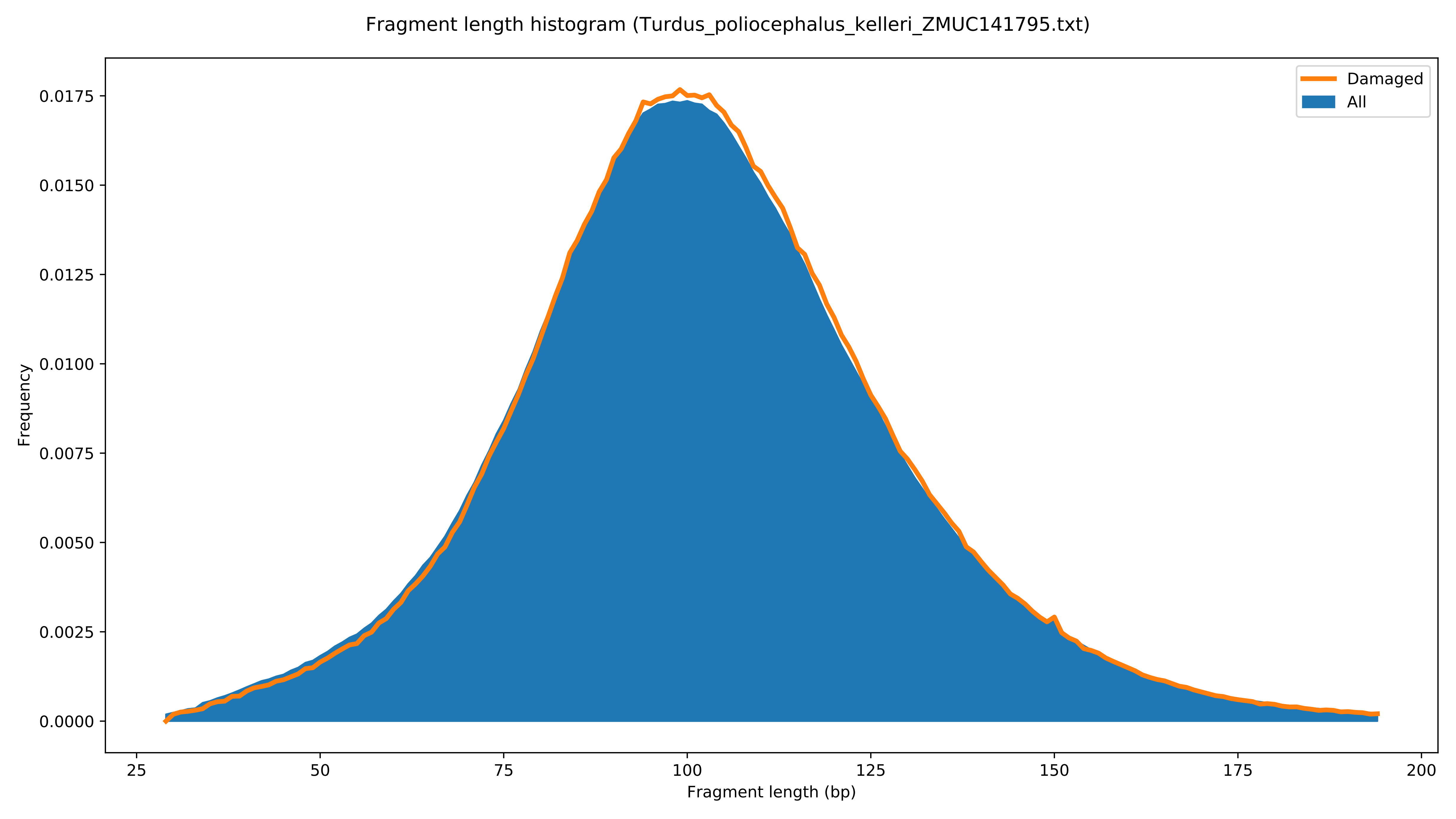

Fragment length histogram (Turdus_poliocephalus_kelleri_ZMUC141795.txt)
Damaged
All
Frequency
0.0000
0.0025
0.0050
0.0075
0.0100
0.0125
0.0150
0.0175
25
50
75
100
125
150
175
200
Fragment length (bp)

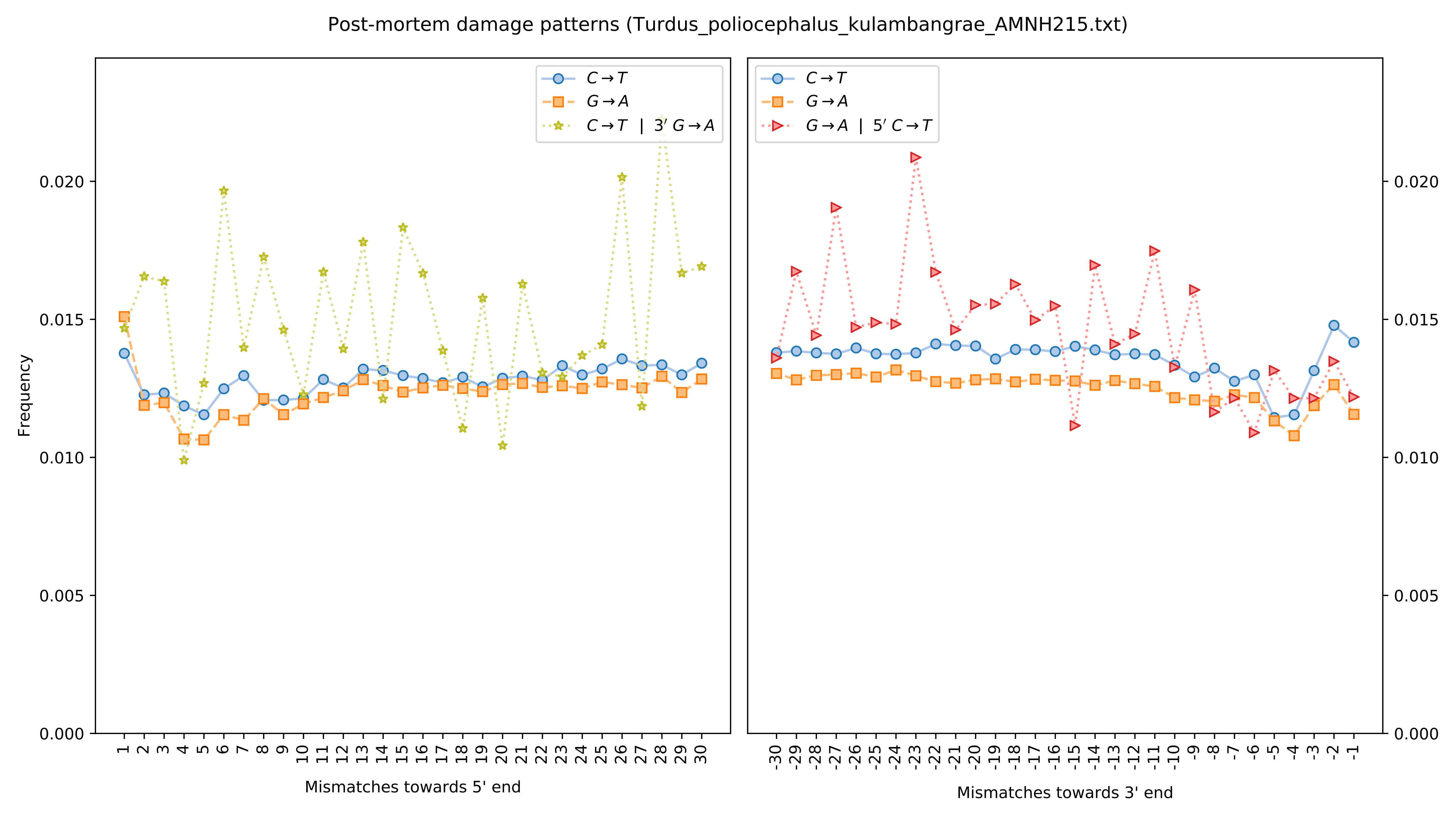
Post-mortem damage patterns (Turdus_poliocephalus_kulambangrae_AMNH215.txt)
C→T
G→A
C→T | 3′ G→A
G→A | 5′ C→T
Frequency
Mismatches towards 5' end
Mismatches towards 3' end
0.000
0.005
0.010
0.015
0.020

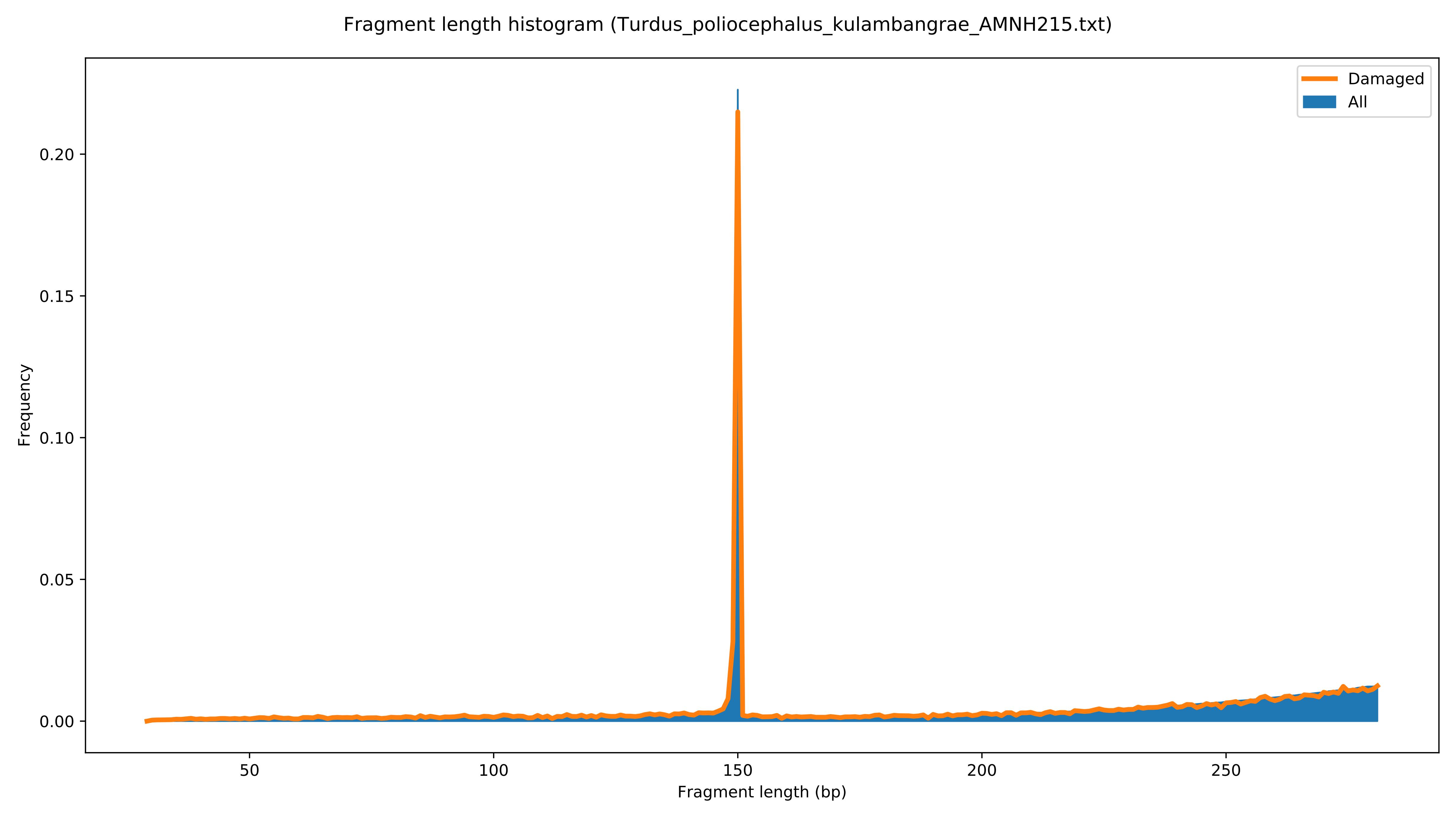
Fragment length histogram (Turdus_poliocephalus_kulambangrae_AMNH215.txt)
Damaged
All
Frequency
0.00
0.05
0.10
0.15
0.20
50
100
150
200
250
Fragment length (bp)

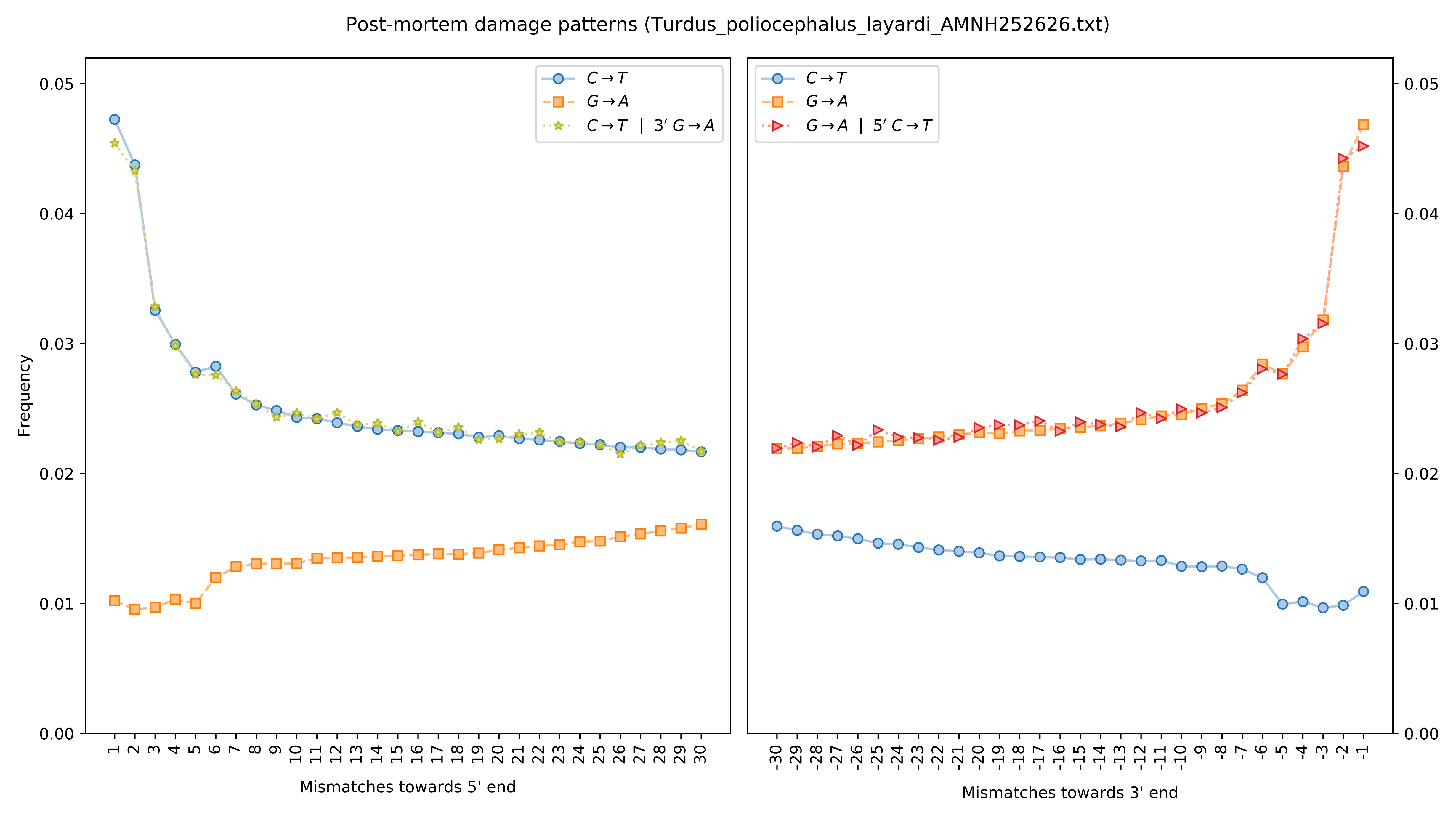
Post-mortem damage patterns (Turdus_poliocephalus_layardi_AMNH252626.txt)
Frequency
C→T
G→A
C→T | 3′ G→A
G→A | 5′ C→T
Mismatches towards 5' end
Mismatches towards 3' end

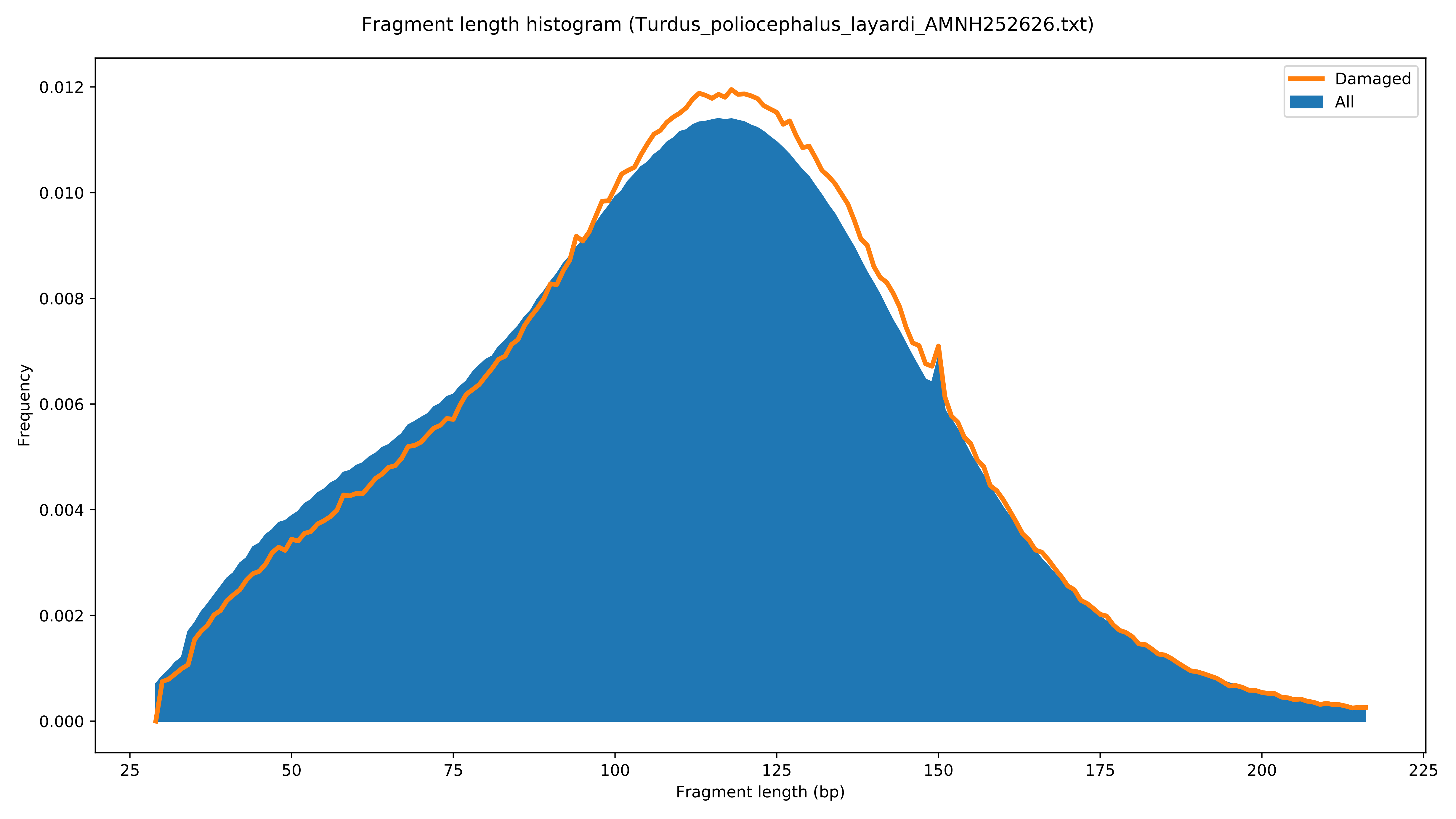

Fragment length histogram (Turdus_poliocephalus_layardi_AMNH252626.txt)
Frequency
Fragment length (bp)
Damaged
All
0.000
0.002
0.004
0.006
0.008
0.010
0.012
25
50
75
100
125
150
175
200
225

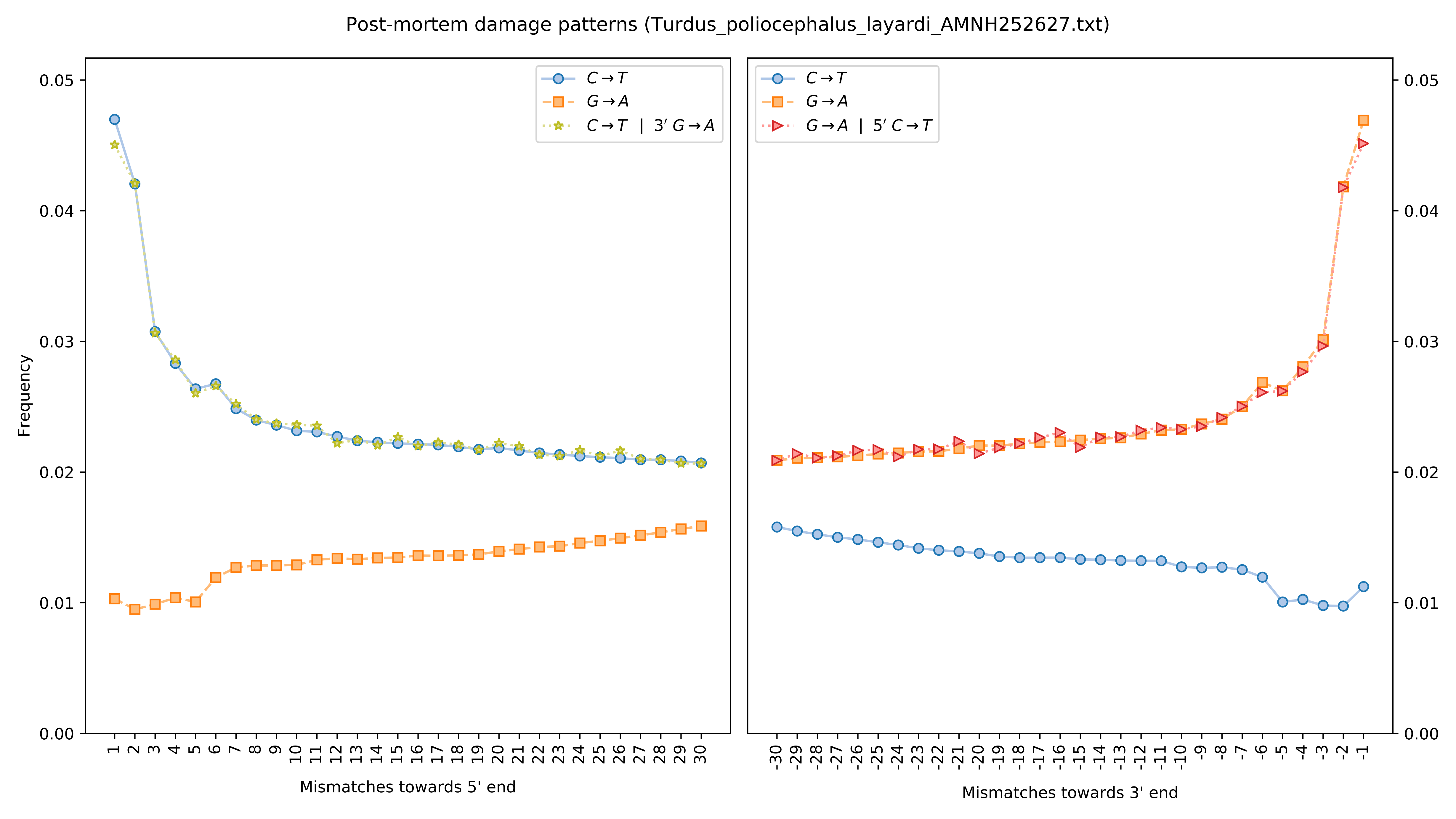

Post-mortem damage patterns (Turdus_poliocephalus_layardi_AMNH252627.txt)
Frequency
Mismatches towards 5' end
Mismatches towards 3' end
C→T
G→A
C→T | 3′ G→A
G→A | 5′ C→T

Fragment length histogram (Turdus_poliocephalus_layardi_AMNH252627.txt)

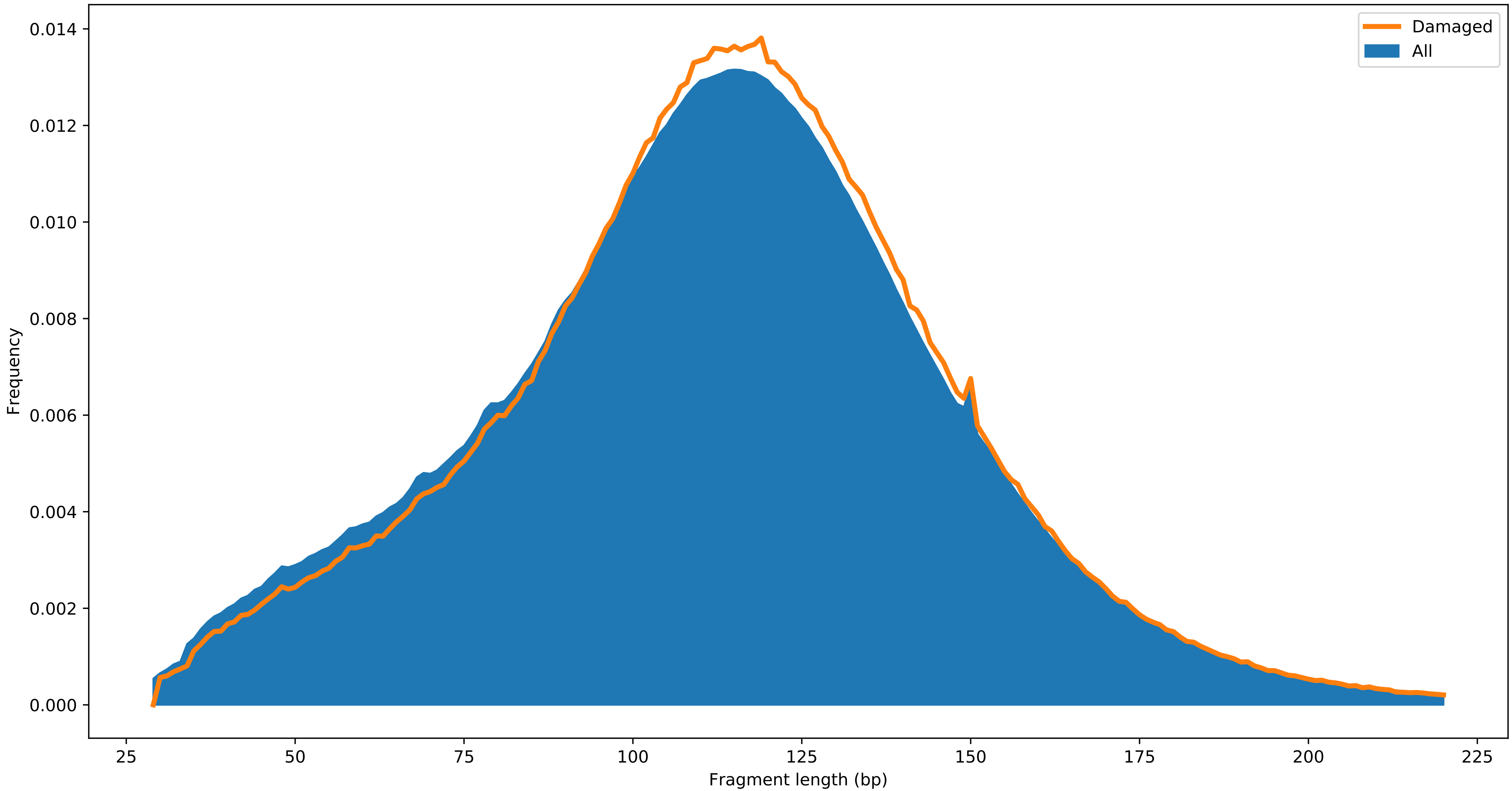

Post-mortem damage patterns (Turdus_poliocephalus_layardi_AMNH252636.txt)

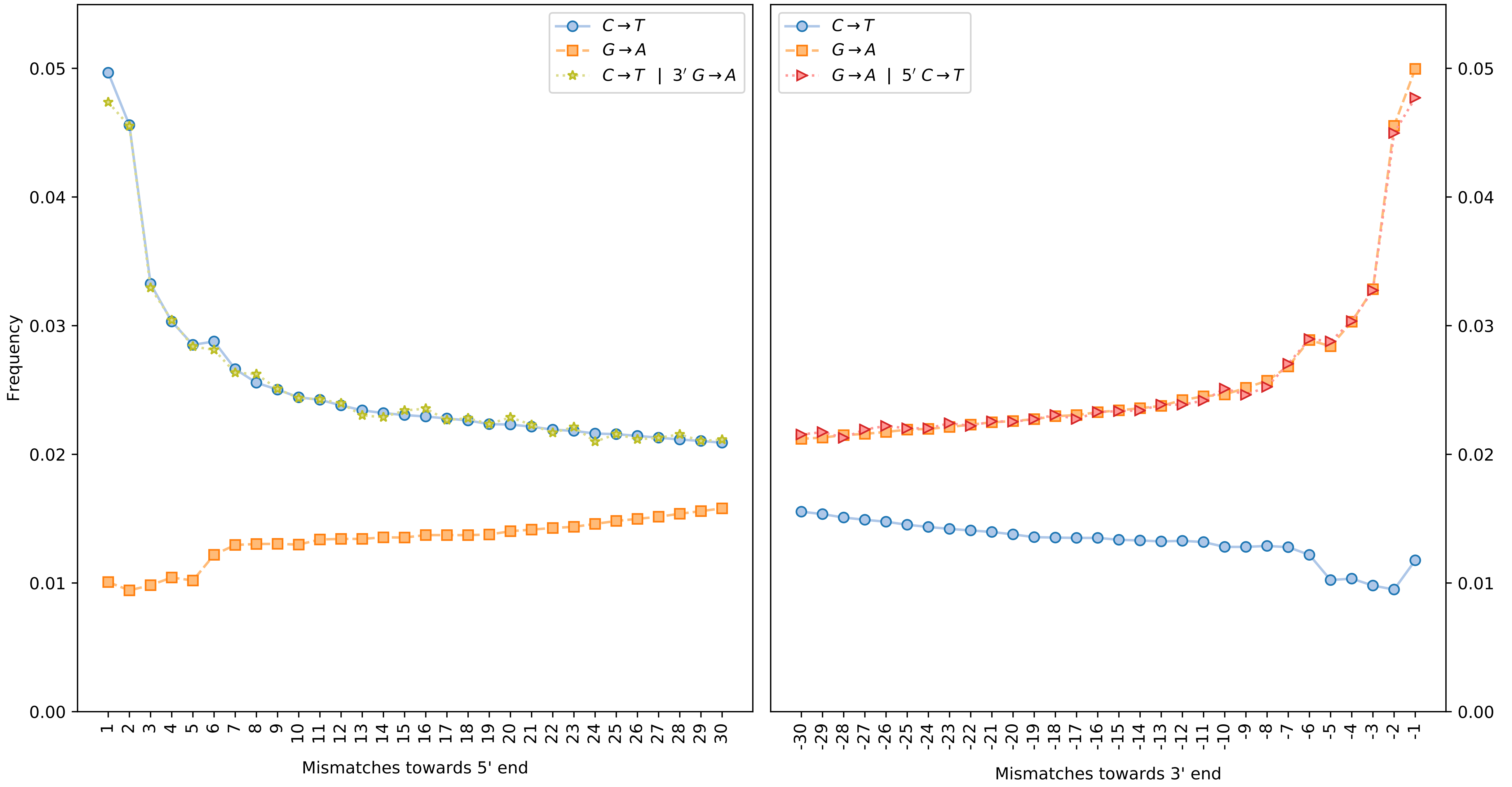

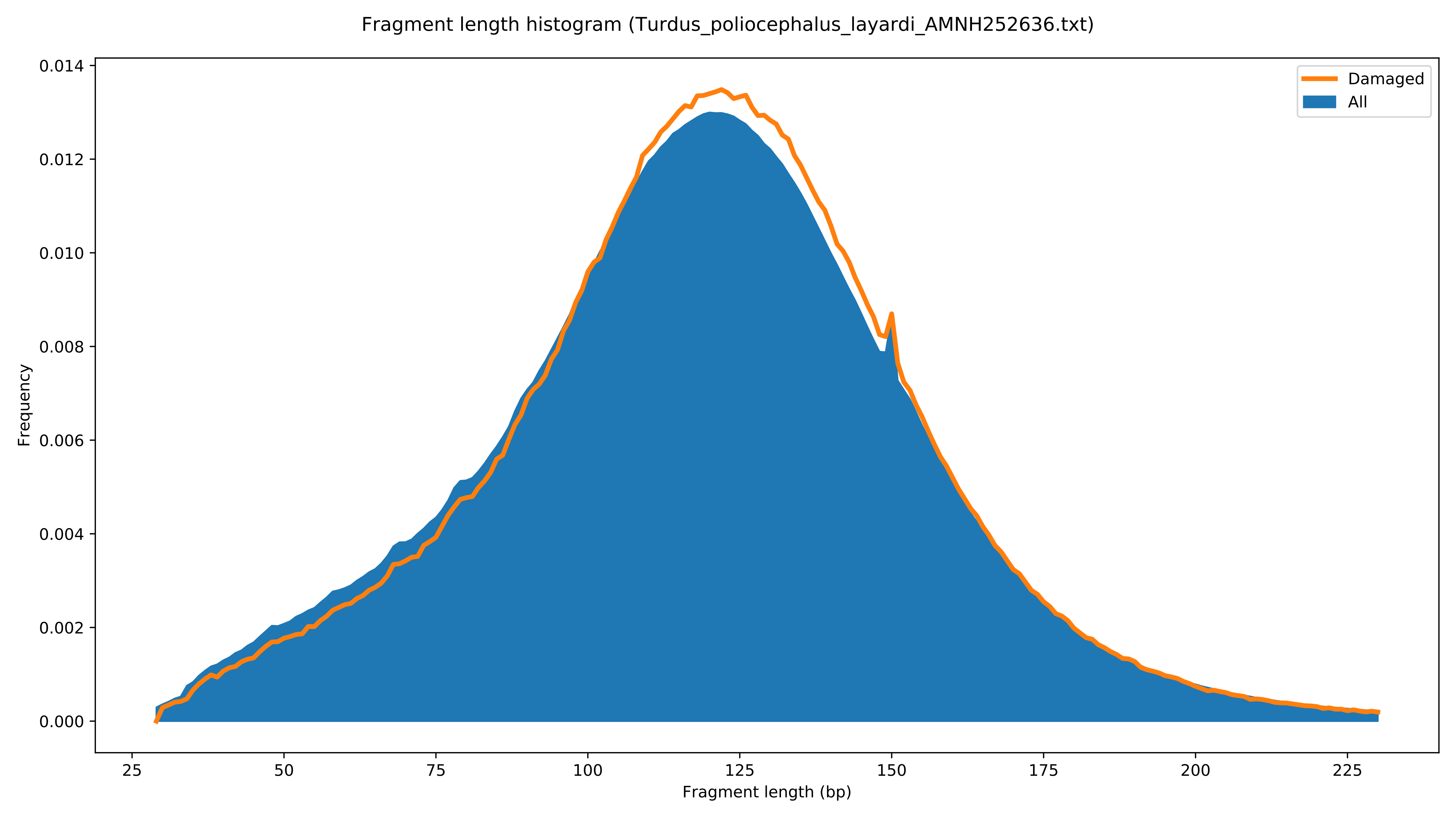

Fragment length histogram (Turdus_poliocephalus_layardi_AMNH252636.txt)
Damaged
All
Frequency
0.000
0.002
0.004
0.006
0.008
0.010
0.012
0.014
25
50
75
100
125
150
175
200
225
Fragment length (bp)

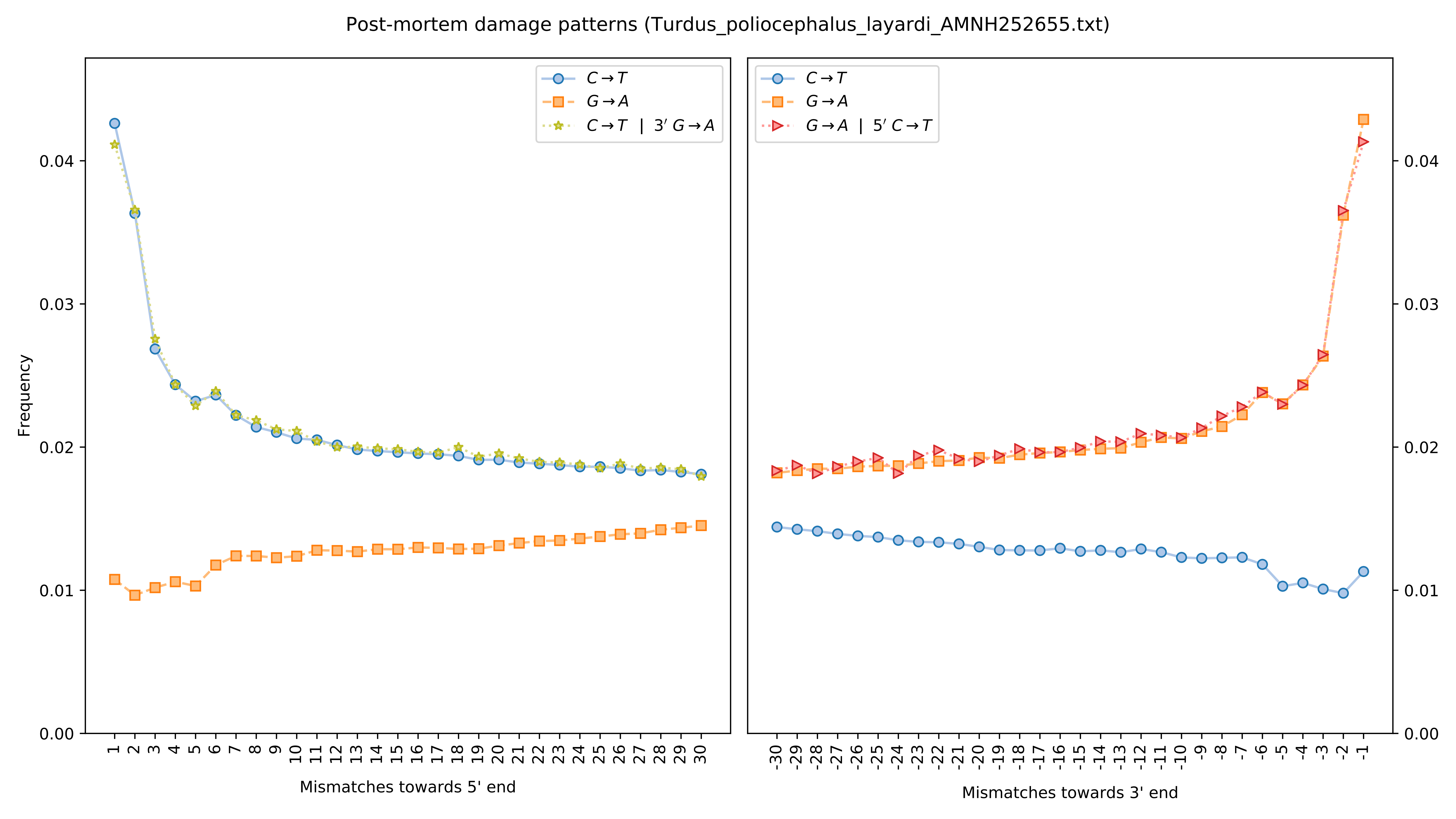
Post-mortem damage patterns (Turdus_poliocephalus_layardi_AMNH252655.txt)
$C \rightarrow T$
$G \rightarrow A$
$C \rightarrow T$ | $3' G \rightarrow A$
$G \rightarrow A$ | $5' C \rightarrow T$
Frequency
Mismatches towards 5' end
Mismatches towards 3' end

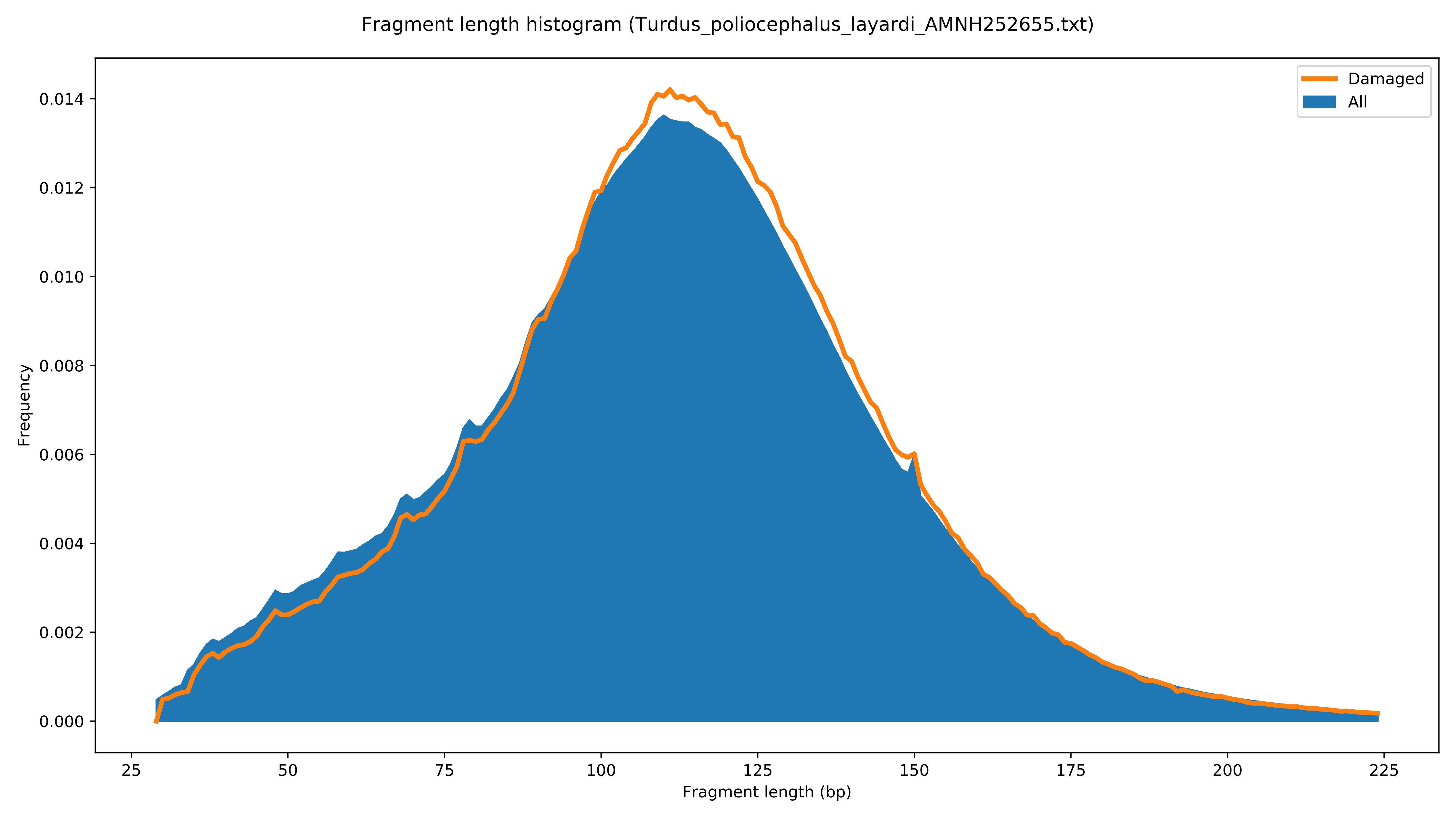

Fragment length histogram (Turdus_poliocephalus_layardi_AMNH252655.txt)
Damaged
All
Frequency
0.000
0.002
0.004
0.006
0.008
0.010
0.012
0.014
25
50
75
100
125
150
175
200
225
Fragment length (bp)

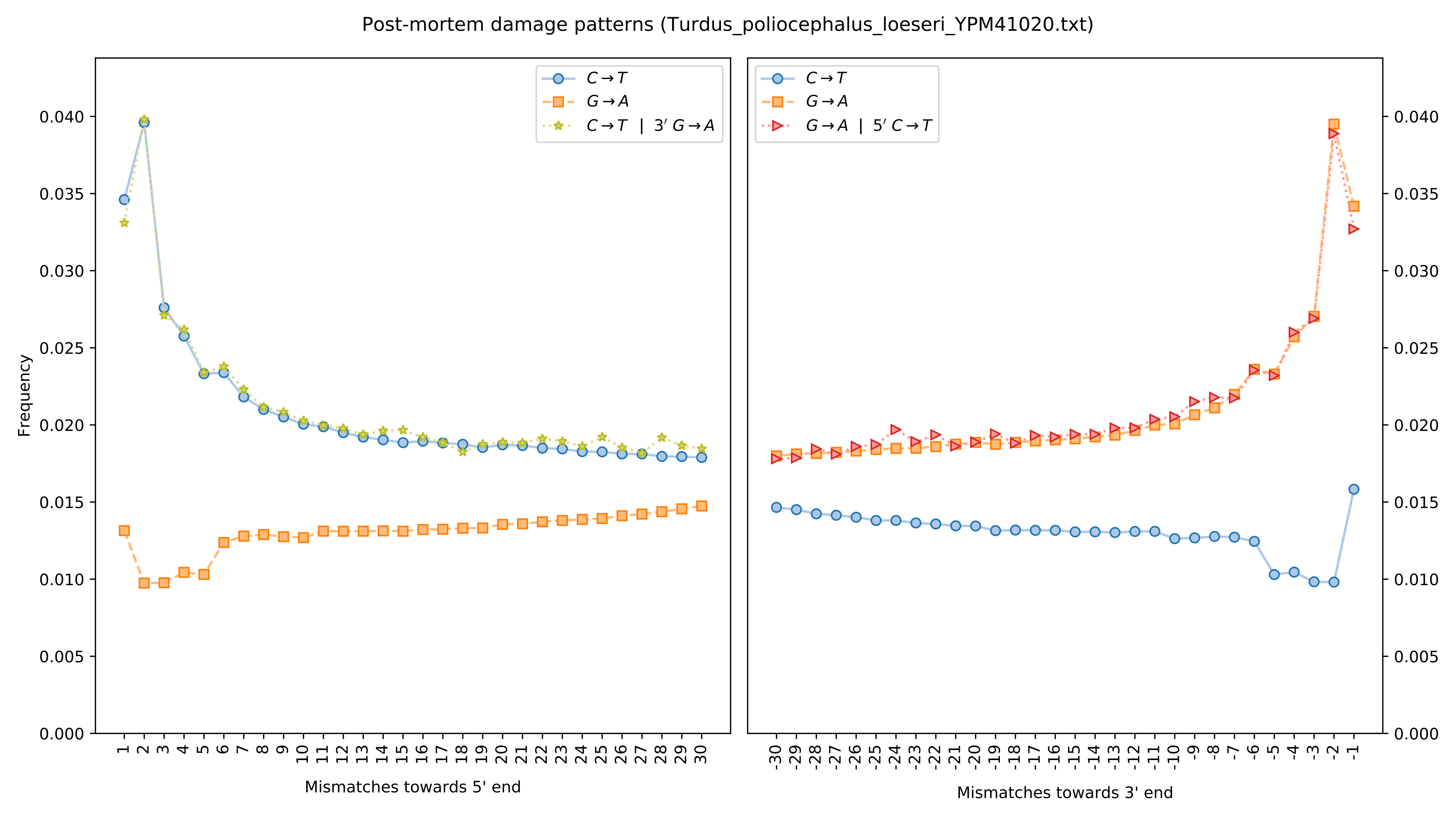

Post-mortem damage patterns (Turdus_poliocephalus_loeseri_YPM41020.txt)
Frequency
C→T
G→A
C→T | 3′ G→A
G→A | 5′ C→T
0.000
0.005
0.010
0.015
0.020
0.025
0.030
0.035
0.040
Mismatches towards 5' end
Mismatches towards 3' end

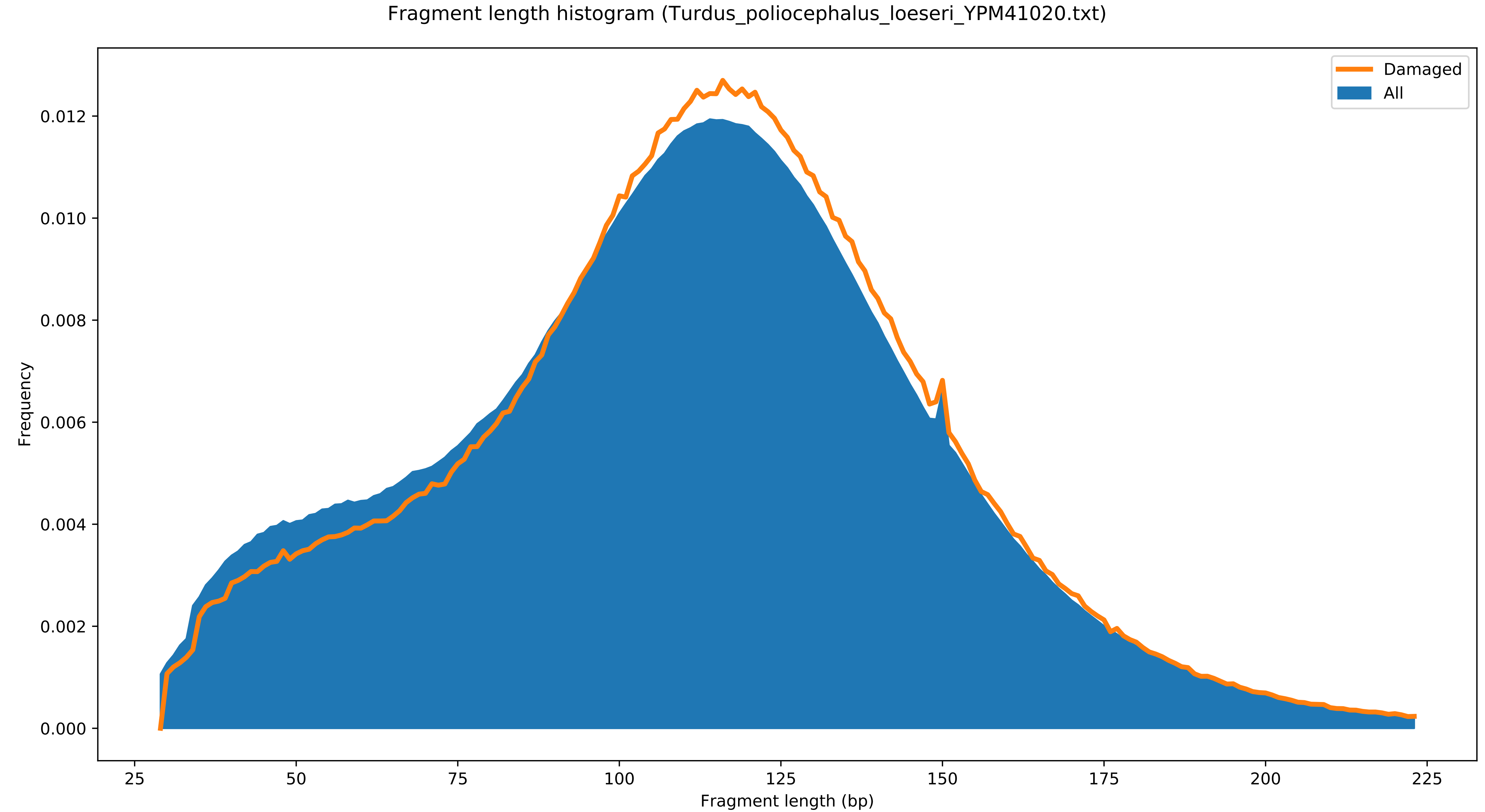
Fragment length histogram (Turdus_poliocephalus_loeseri_YPM41020.txt)
Damaged
All
Frequency
Fragment length (bp)
0.000
0.002
0.004
0.006
0.008
0.010
0.012
25
50
75
100
125
150
175
200
225

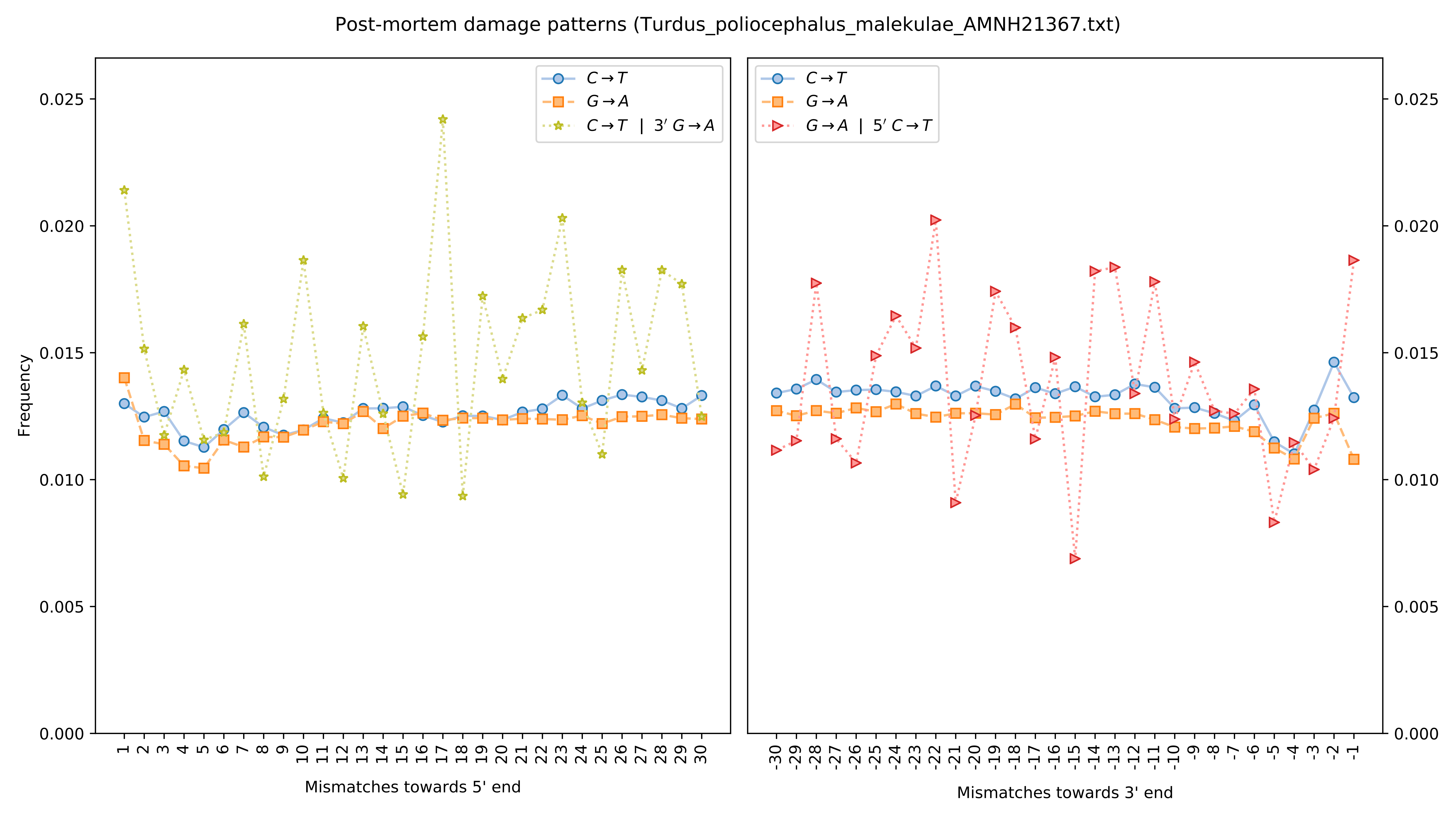
Post-mortem damage patterns (Turdus_poliocephalus_malekulae_AMNH21367.txt)
C→T
G→A
C→T | 3′ G→A
G→A | 5′ C→T
Frequency
0.000
0.005
0.010
0.015
0.020
0.025
Mismatches towards 5' end
Mismatches towards 3' end

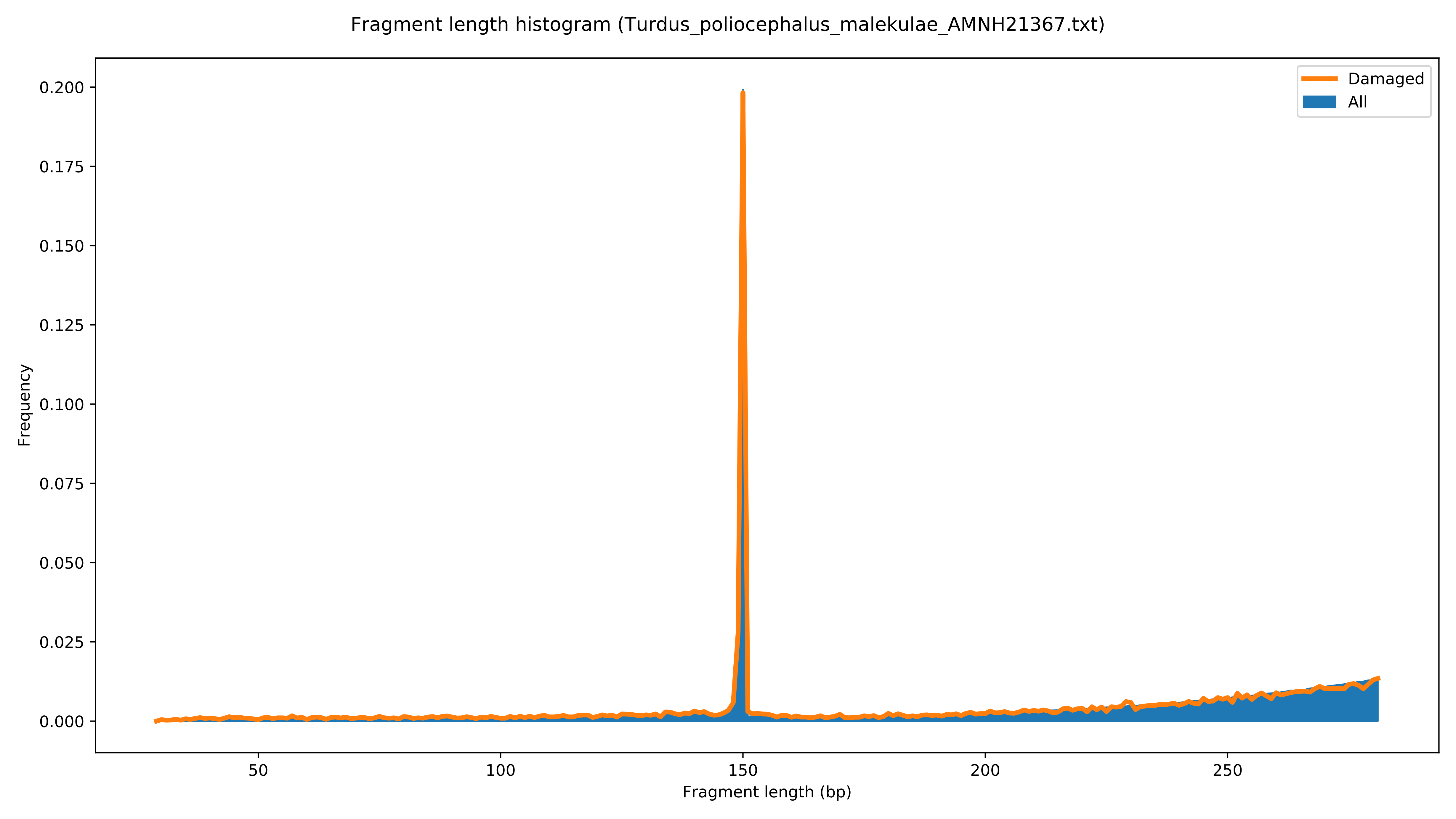

Fragment length histogram (Turdus_poliocephalus_malekulae_AMNH21367.txt)
Damaged
All
Frequency
0.000
0.025
0.050
0.075
0.100
0.125
0.150
0.175
0.200
50
100
150
200
250
Fragment length (bp)

Post-mortem damage patterns (Turdus_poliocephalus_malekulae_AMNH214424.txt)

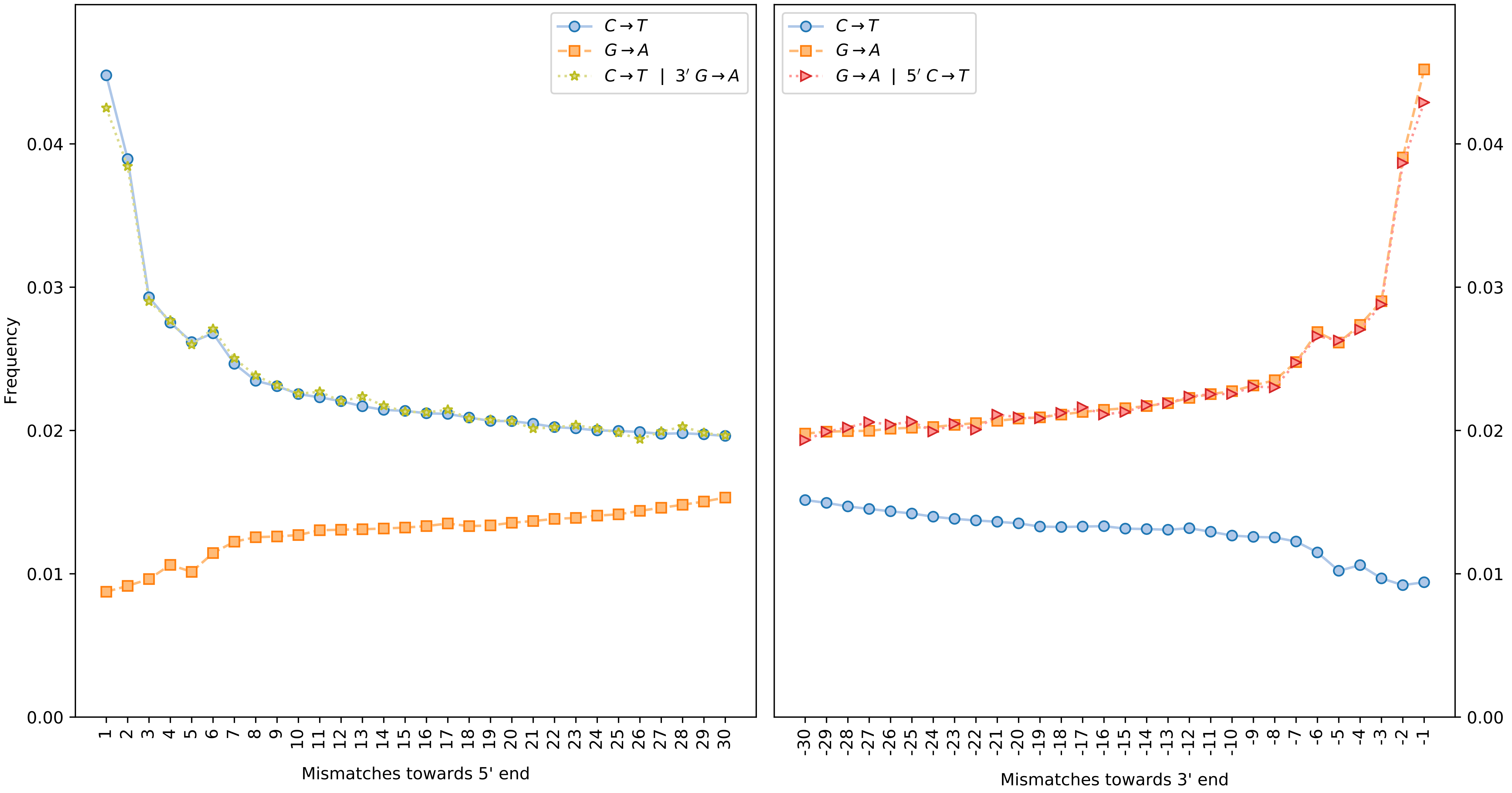

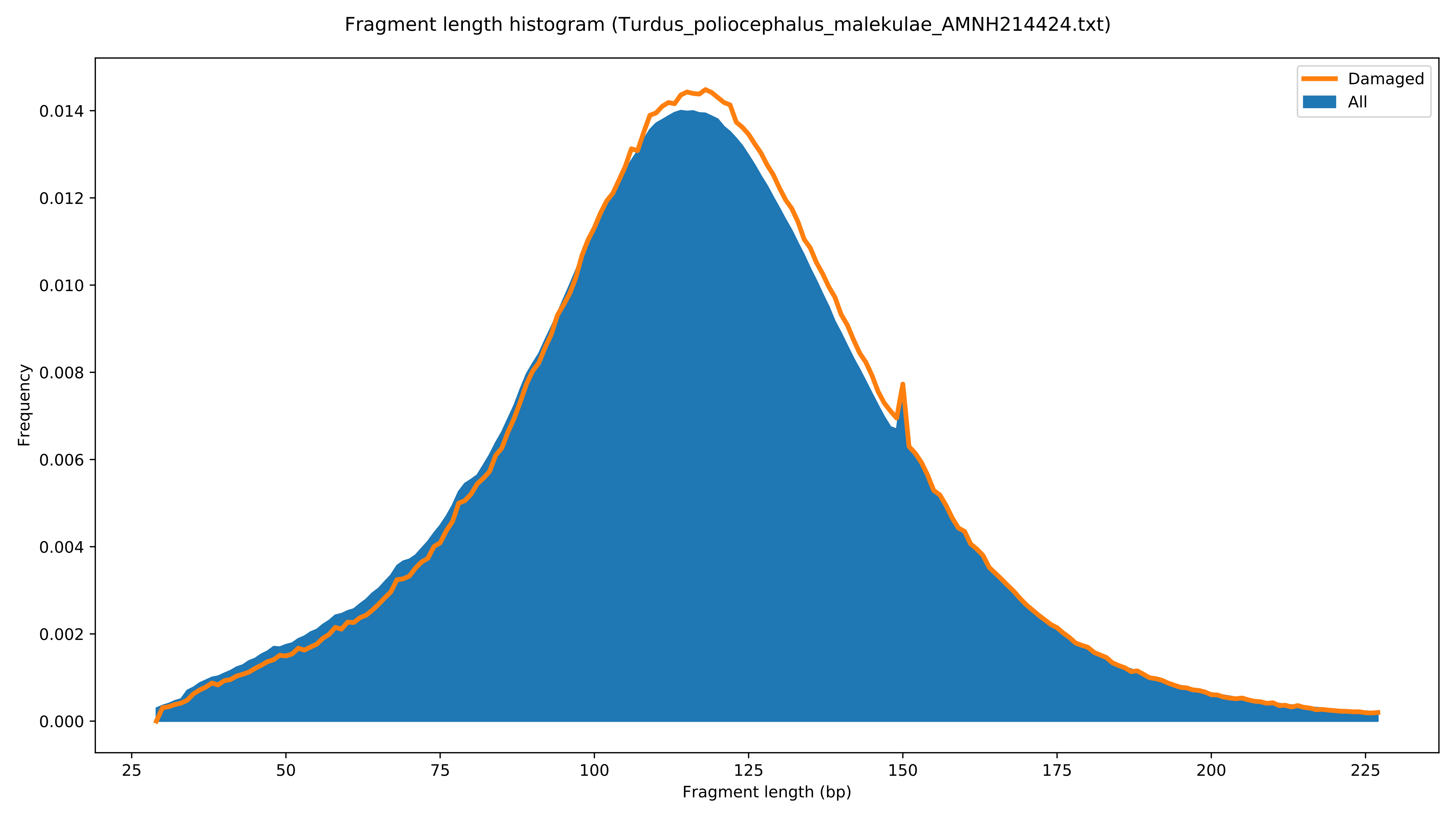
Fragment length histogram (Turdus_poliocephalus_malekulae_AMNH214424.txt)
Frequency
0.000
0.002
0.004
0.006
0.008
0.010
0.012
0.014
Fragment length (bp)
25
50
75
100
125
150
175
200
225
Damaged
All

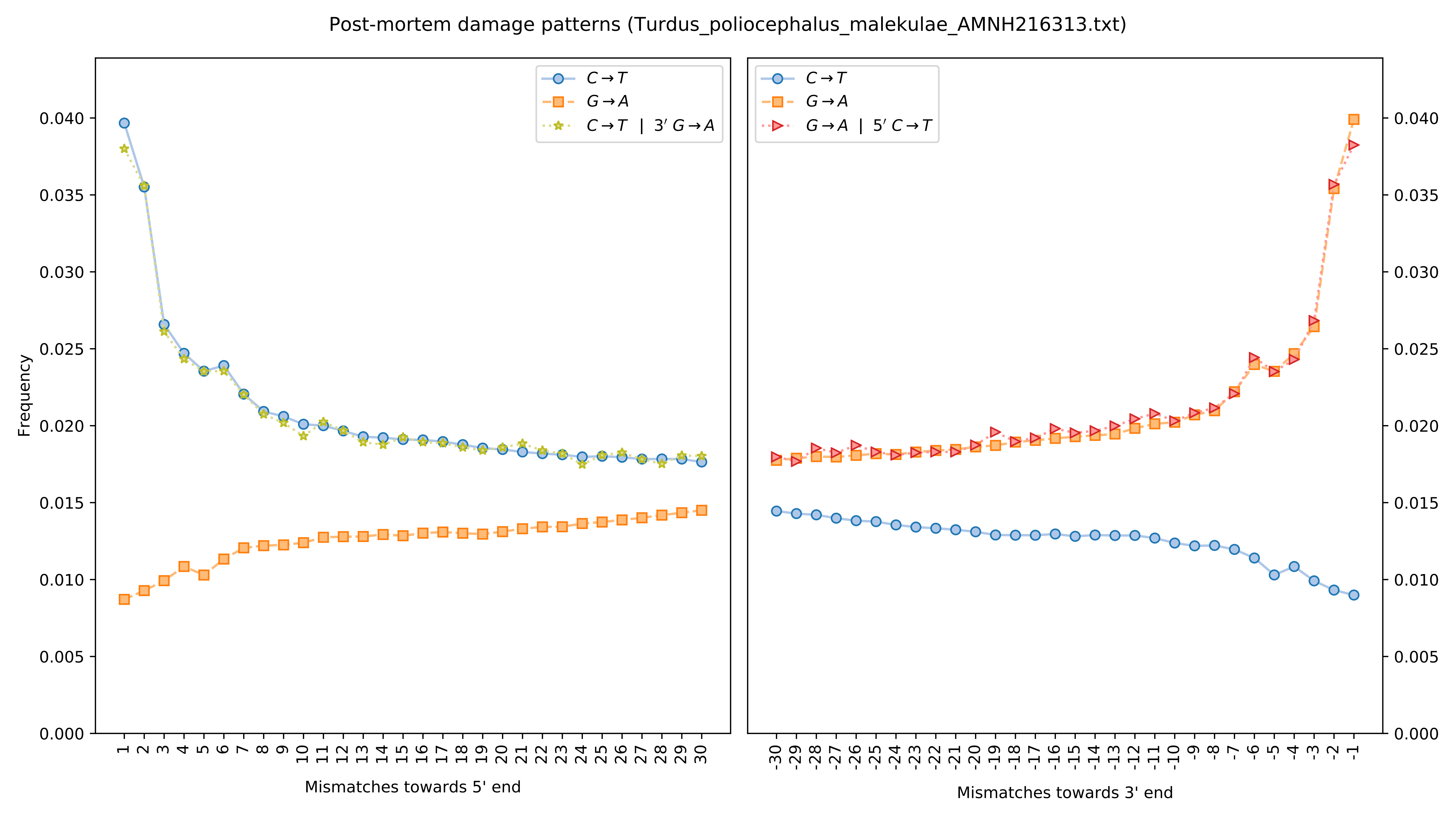

Post-mortem damage patterns (Turdus_poliocephalus_malekulae_AMNH216313.txt)
Frequency
Mismatches towards 5' end
Mismatches towards 3' end
C→T
G→A
C→T | 3′ G→A
G→A | 5′ C→T

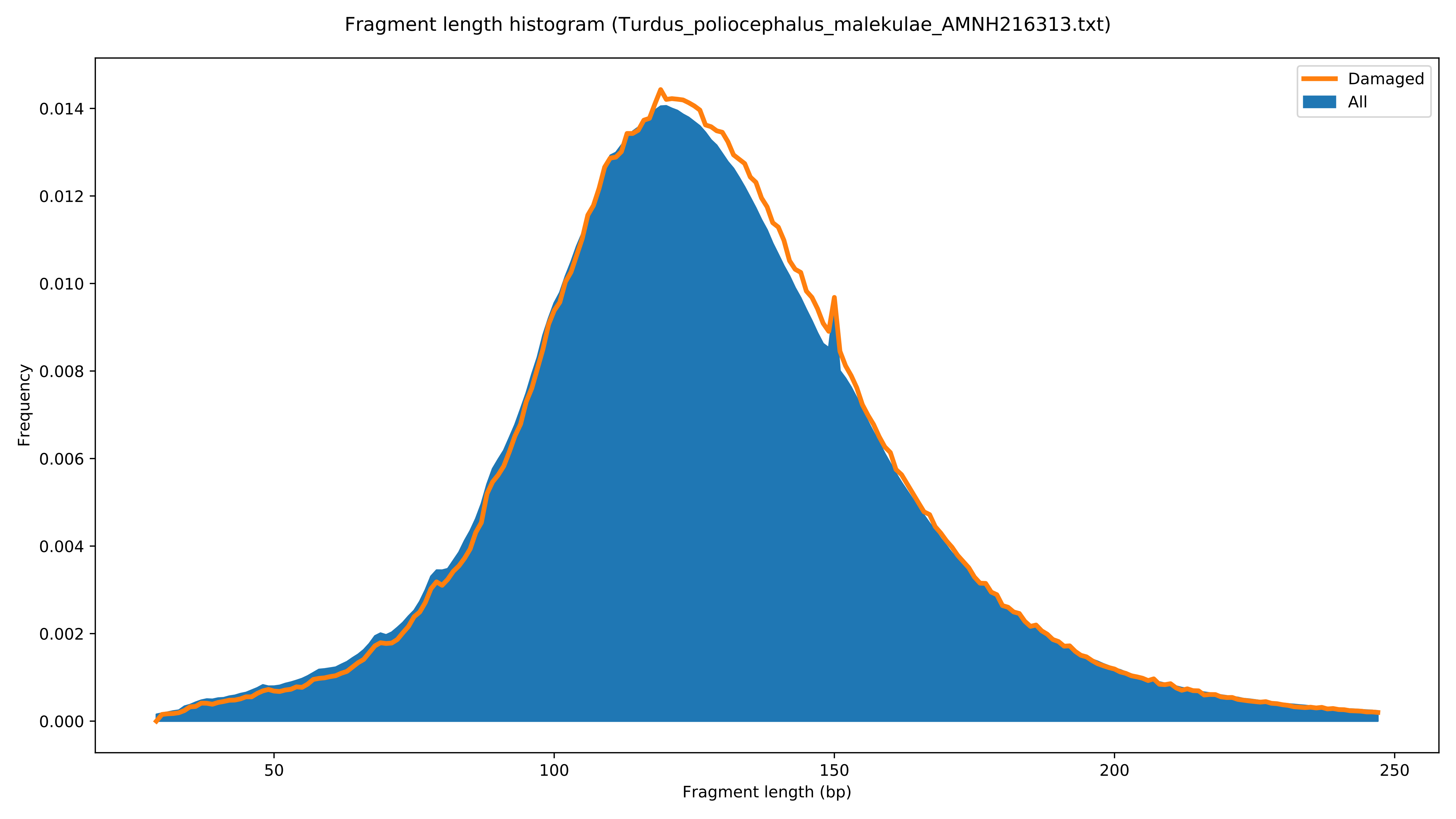

Fragment length histogram (Turdus_poliocephalus_malekulae_AMNH216313.txt)
Damaged
All
Frequency
0.000
0.002
0.004
0.006
0.008
0.010
0.012
0.014
50
100
150
200
250
Fragment length (bp)

Post-mortem damage patterns (Turdus_poliocephalus_malindangensis_ZMUC141794.txt)

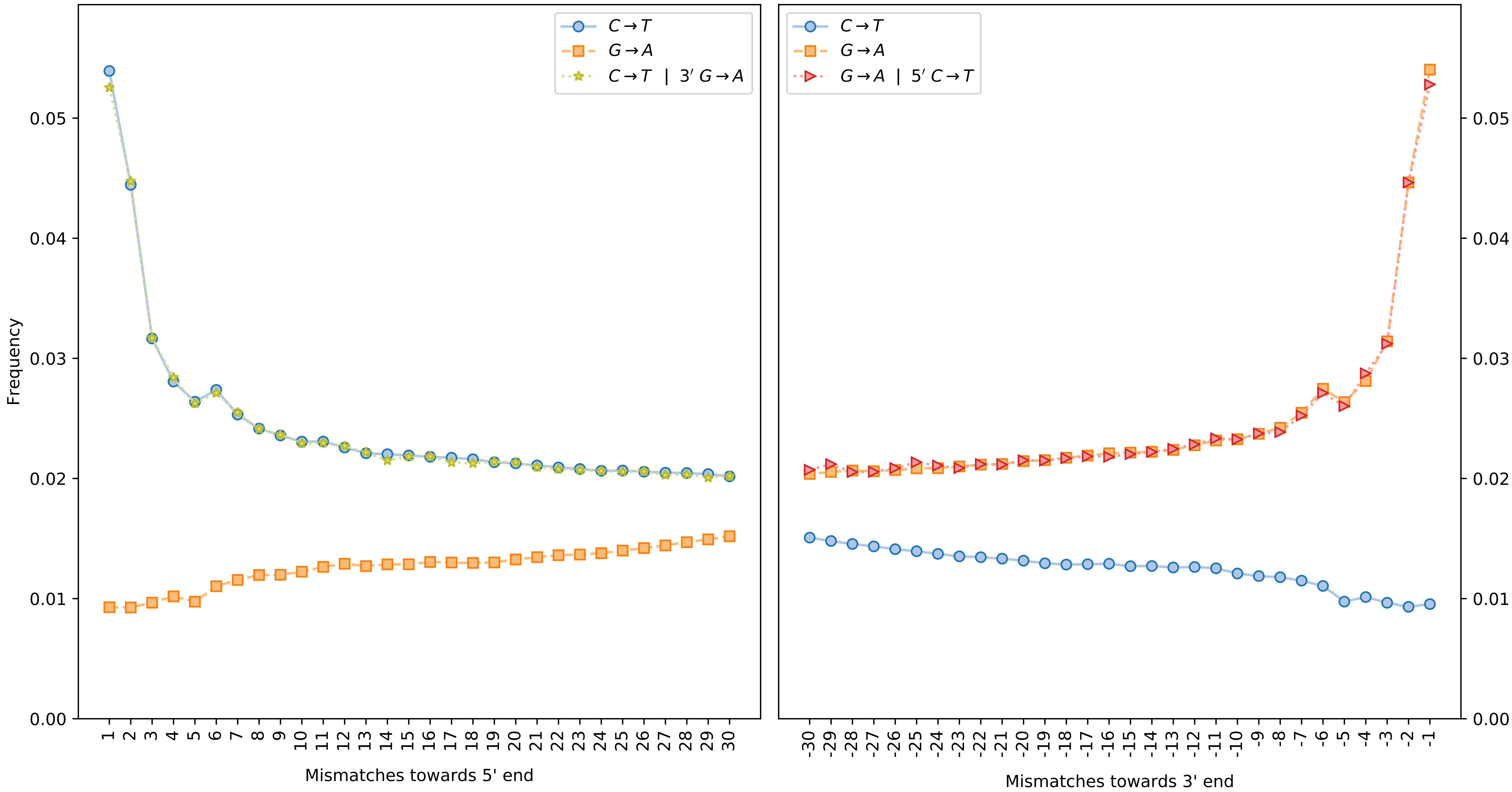

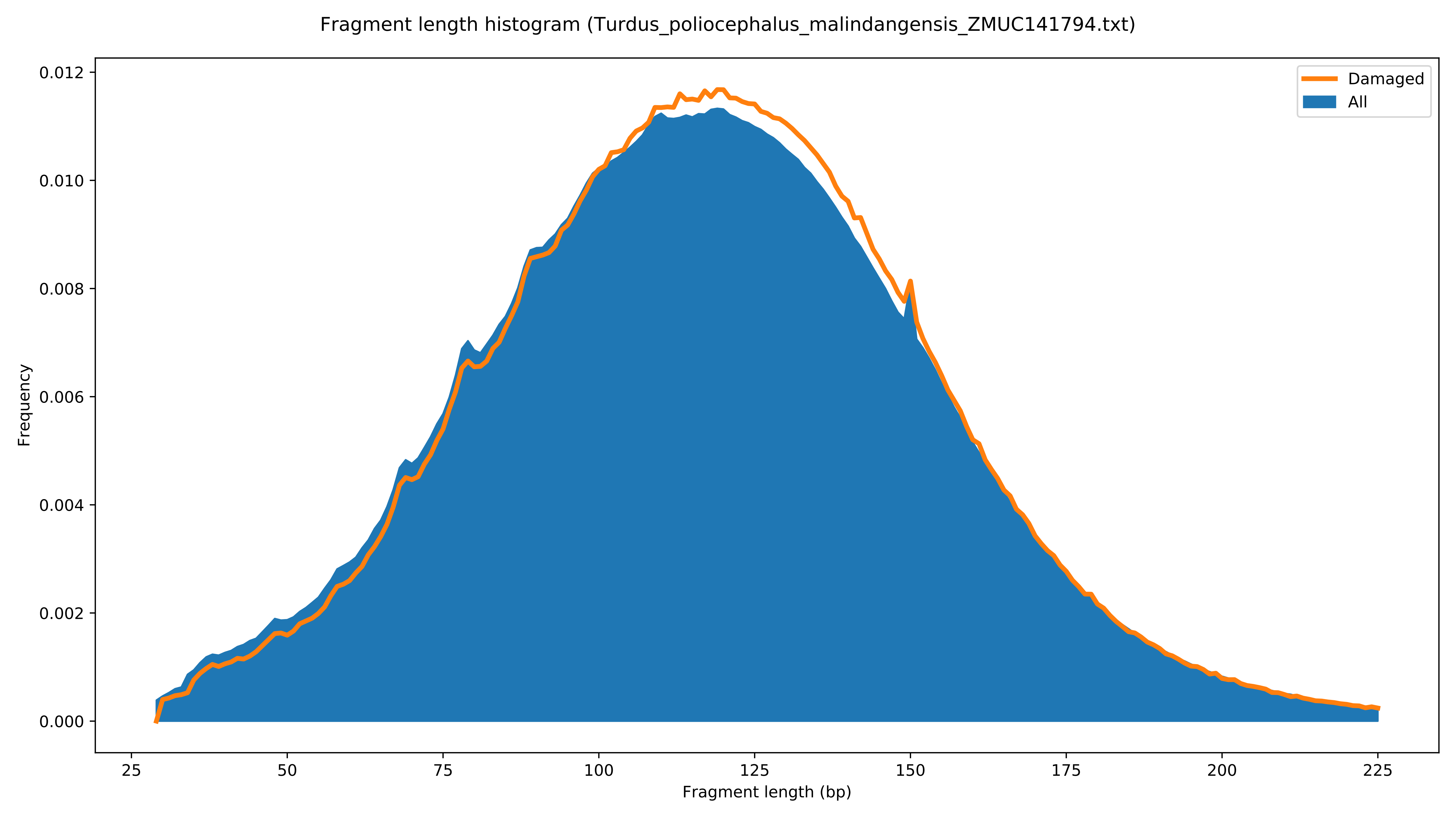
Fragment length histogram (Turdus_poliocephalus_malindangensis_ZMUC141794.txt)
Damaged
All
Frequency
0.000
0.002
0.004
0.006
0.008
0.010
0.012
25
50
75
100
125
150
175
200
225
Fragment length (bp)

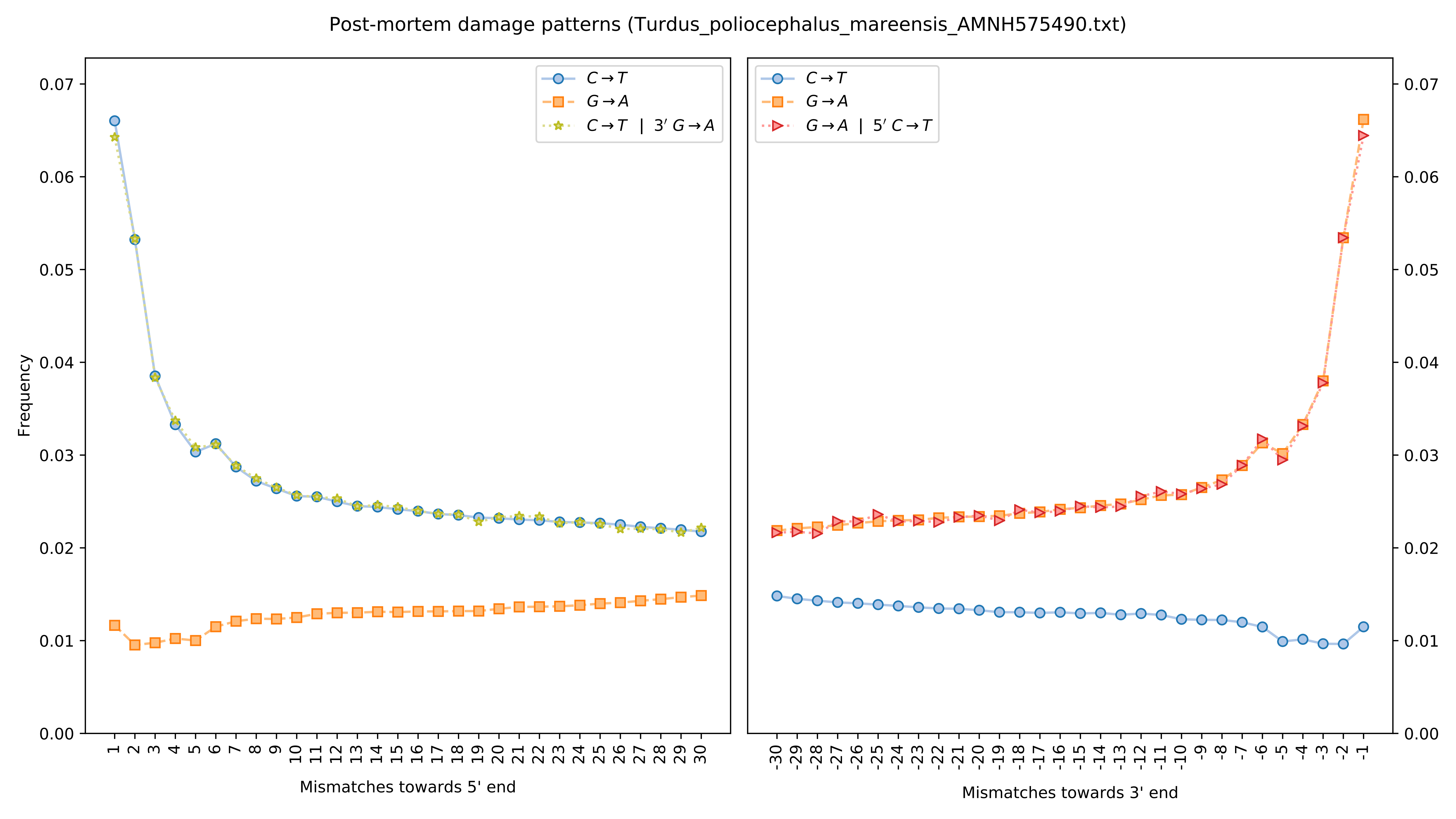

Post-mortem damage patterns (Turdus_poliocephalus_mareensis_AMNH575490.txt)
Frequency
Mismatches towards 5' end
Mismatches towards 3' end
C→T
G→A
C→T | 3′ G→A
G→A | 5′ C→T

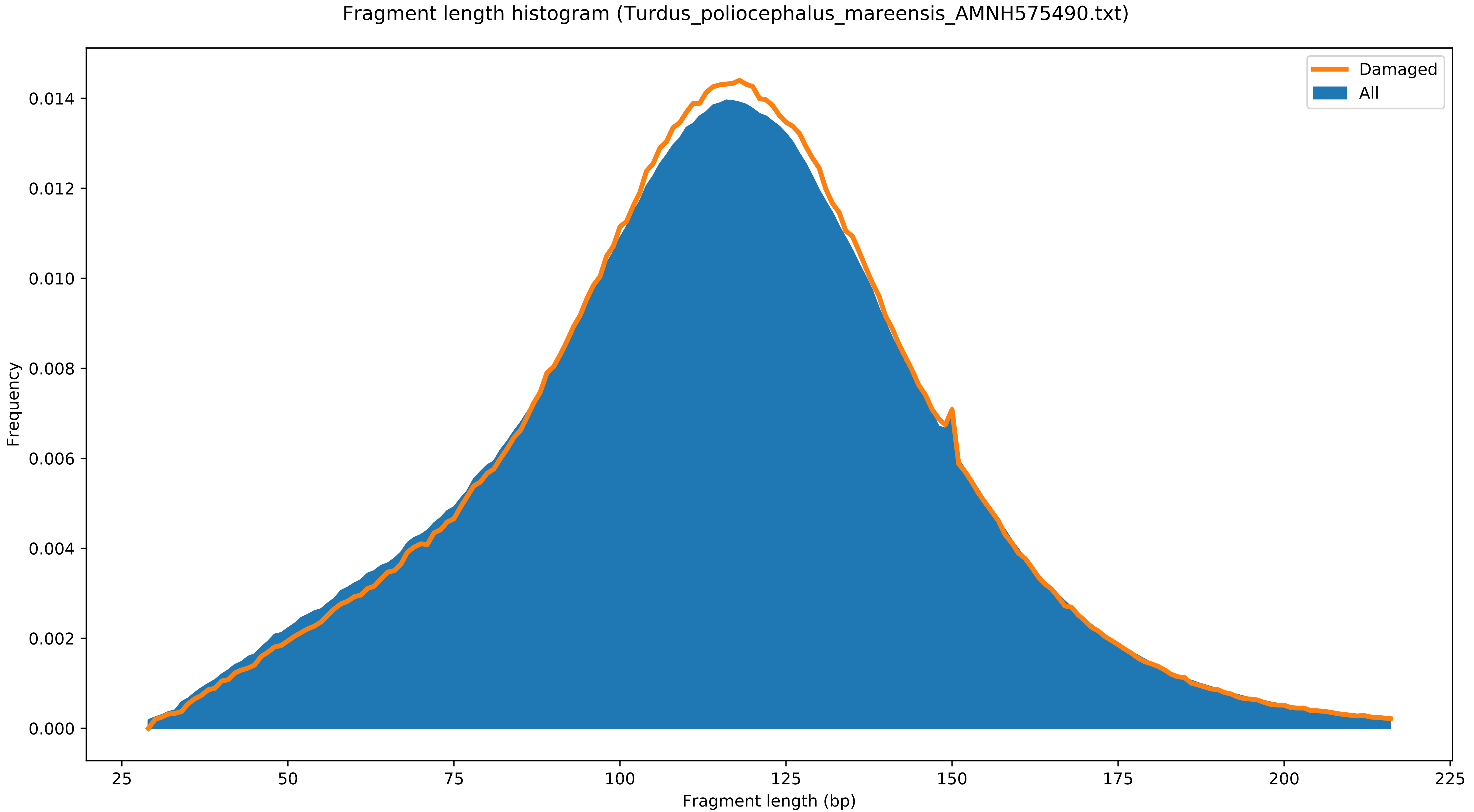

Fragment length histogram (Turdus_poliocephalus_mareensis_AMNH575490.txt)
Damaged
All
Frequency
0.000
0.002
0.004
0.006
0.008
0.010
0.012
0.014
Fragment length (bp)
25
50
75
100
125
150
175
200
225

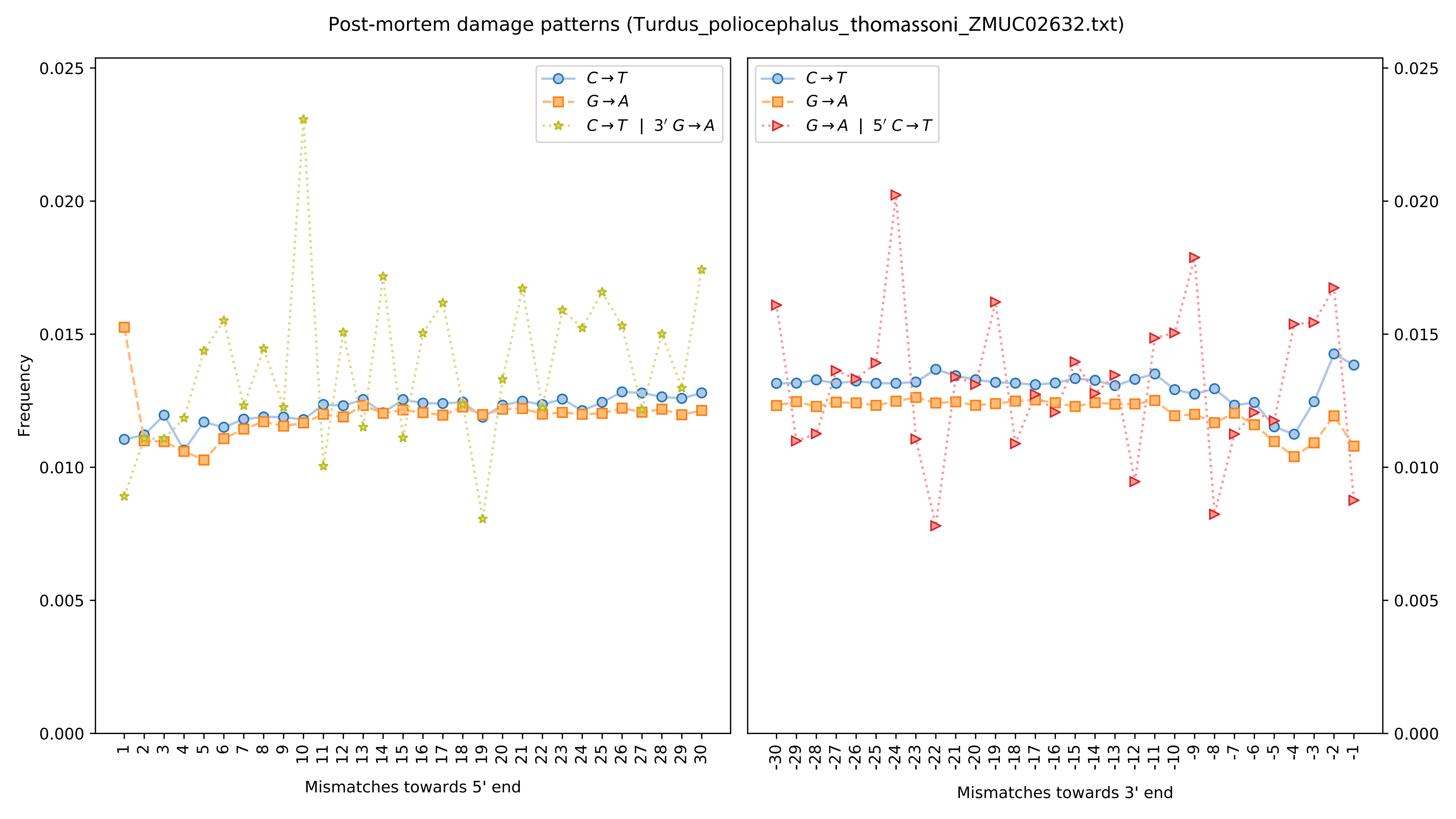

Post-mortem damage patterns (Turdus_poliocephalus_thomassoni_ZMUC02632.txt)
Frequency
Mismatches towards 5' end
Mismatches towards 3' end
C→T
G→A
C→T | 3′ G→A
G→A | 5′ C→T
0.000
0.005
0.010
0.015
0.020
0.025

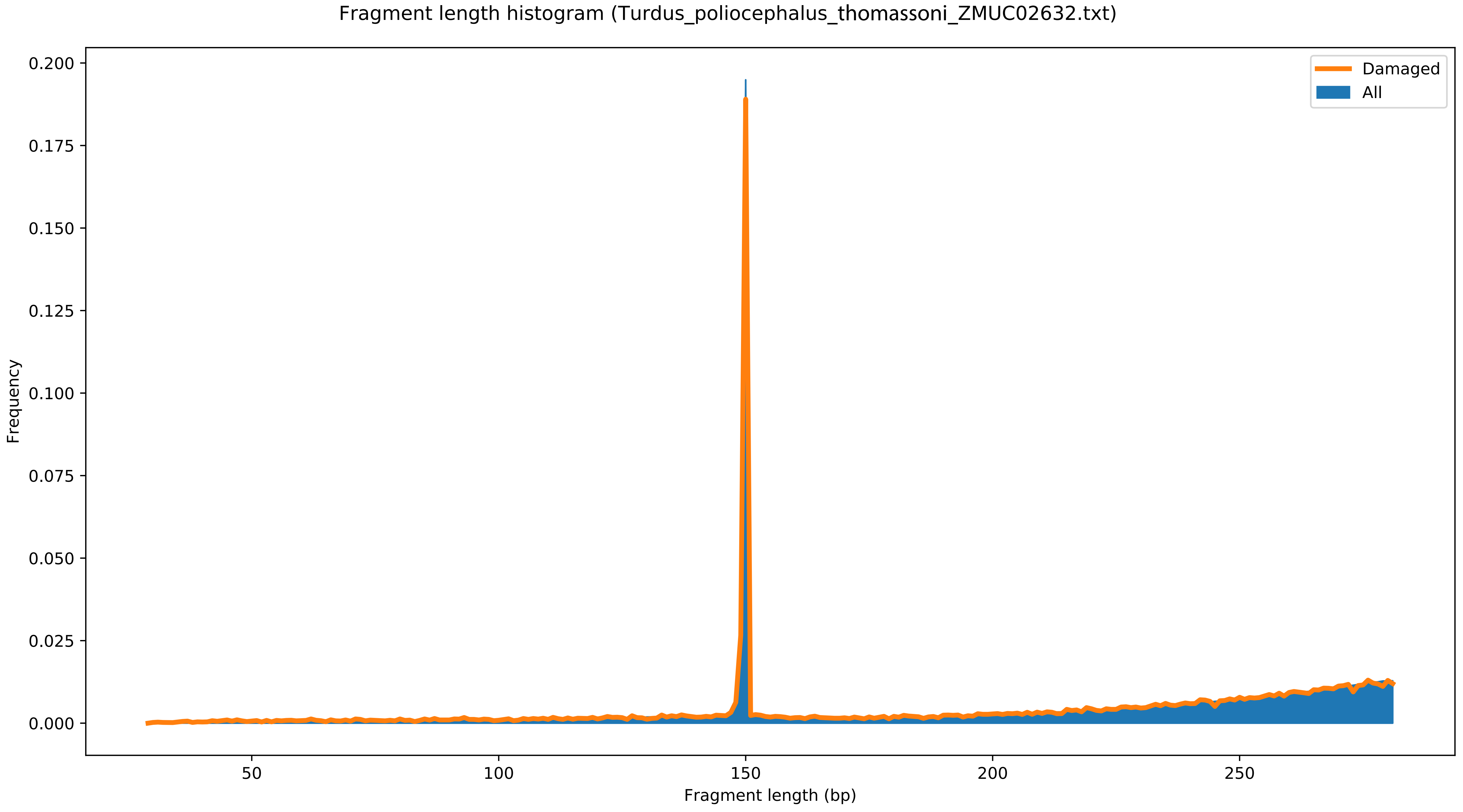

Fragment length histogram (Turdus_poliocephalus_thomassoni_ZMUC02632.txt)
Damaged
All
Frequency
0.200
0.175
0.150
0.125
0.100
0.075
0.050
0.025
0.000
50
100
150
200
250
Fragment length (bp)

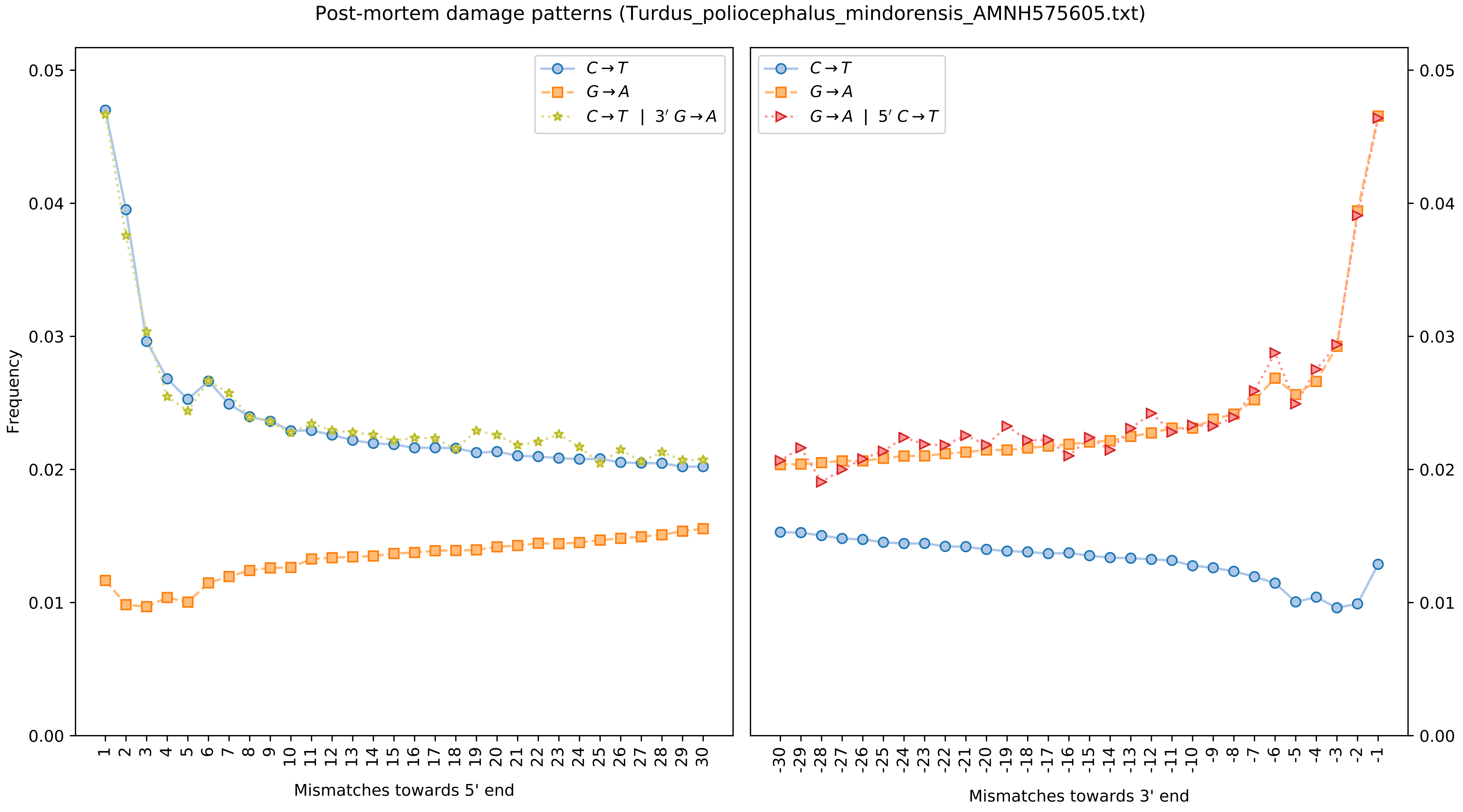

Post-mortem damage patterns (Turdus_poliocephalus_mindorensis_AMNH575605.txt)
Frequency
Mismatches towards 5' end
Mismatches towards 3' end
C→T
G→A
C→T | 3′ G→A
G→A | 5′ C→T
0.00
0.01
0.02
0.03
0.04
0.05

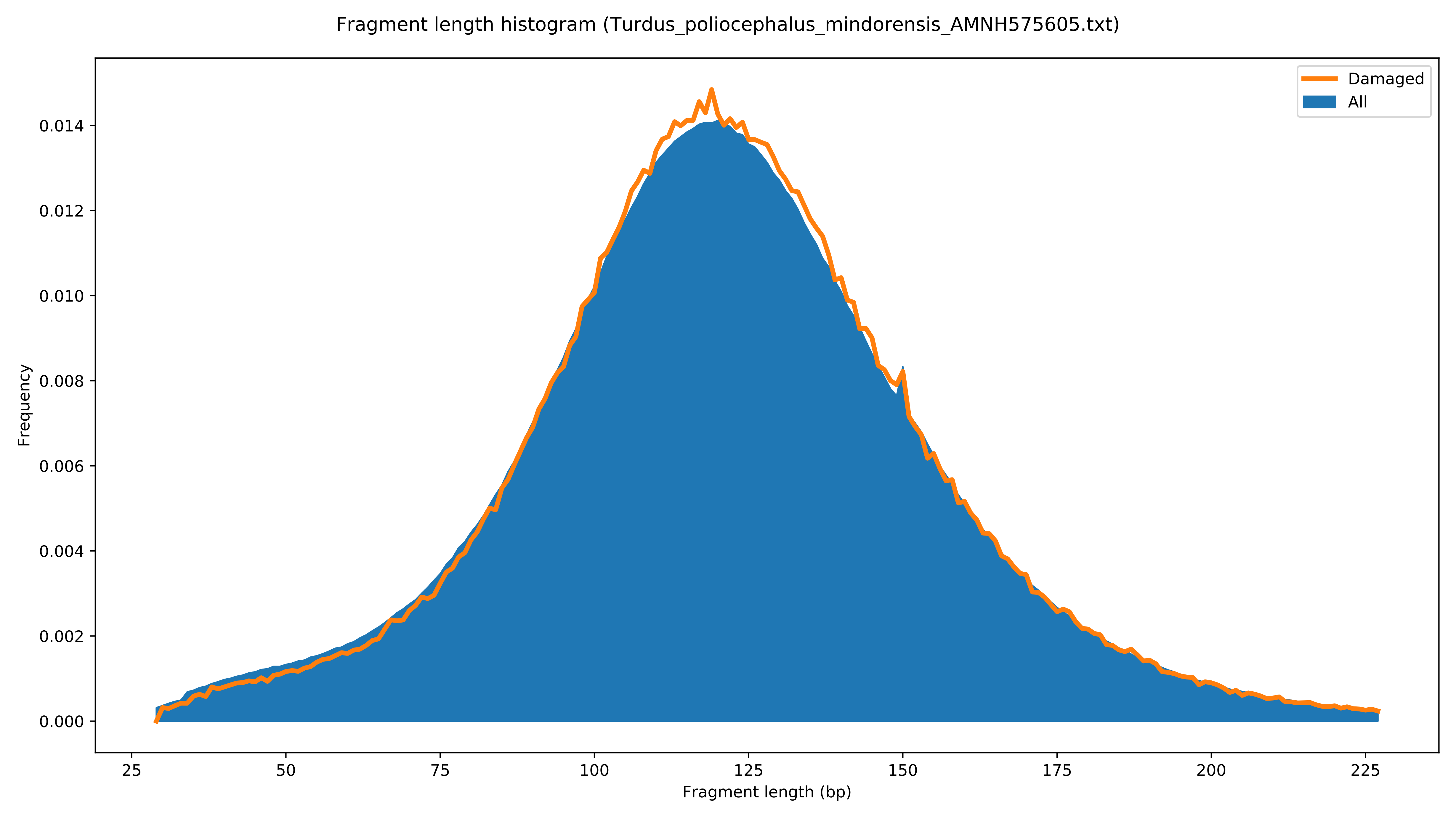
Fragment length histogram (Turdus_poliocephalus_mindorensis_AMNH575605.txt)
Damaged
All
Frequency
0.000
0.002
0.004
0.006
0.008
0.010
0.012
0.014
25
50
75
100
125
150
175
200
225
Fragment length (bp)

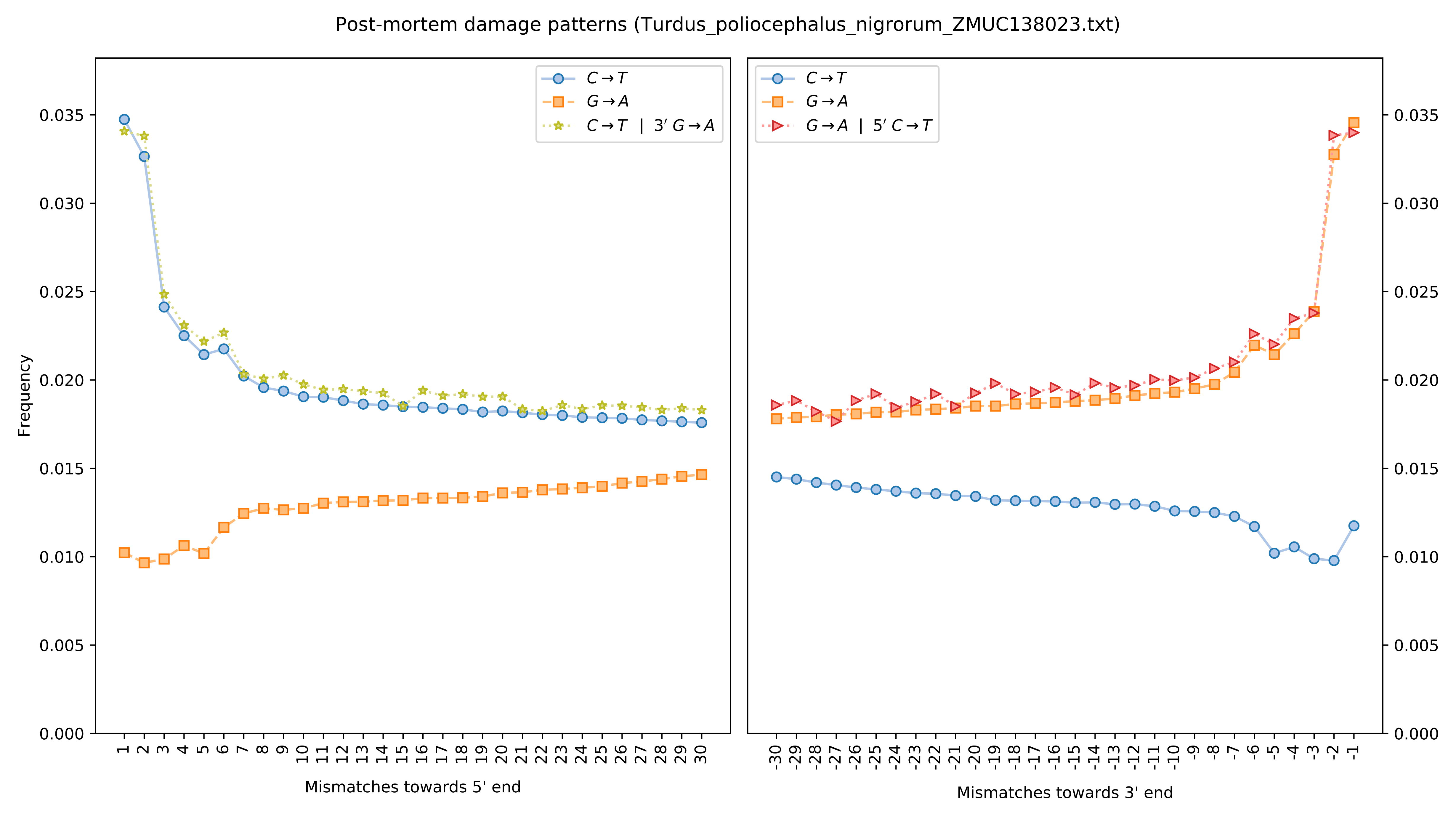

Post-mortem damage patterns (Turdus_poliocephalus_nigrorum_ZMUC138023.txt)
Frequency
Mismatches towards 5' end
Mismatches towards 3' end
$C \rightarrow T$
$G \rightarrow A$
$C \rightarrow T$ | $3' G \rightarrow A$
$G \rightarrow A$ | $5' C \rightarrow T$
0.000
0.005
0.010
0.015
0.020
0.025
0.030
0.035

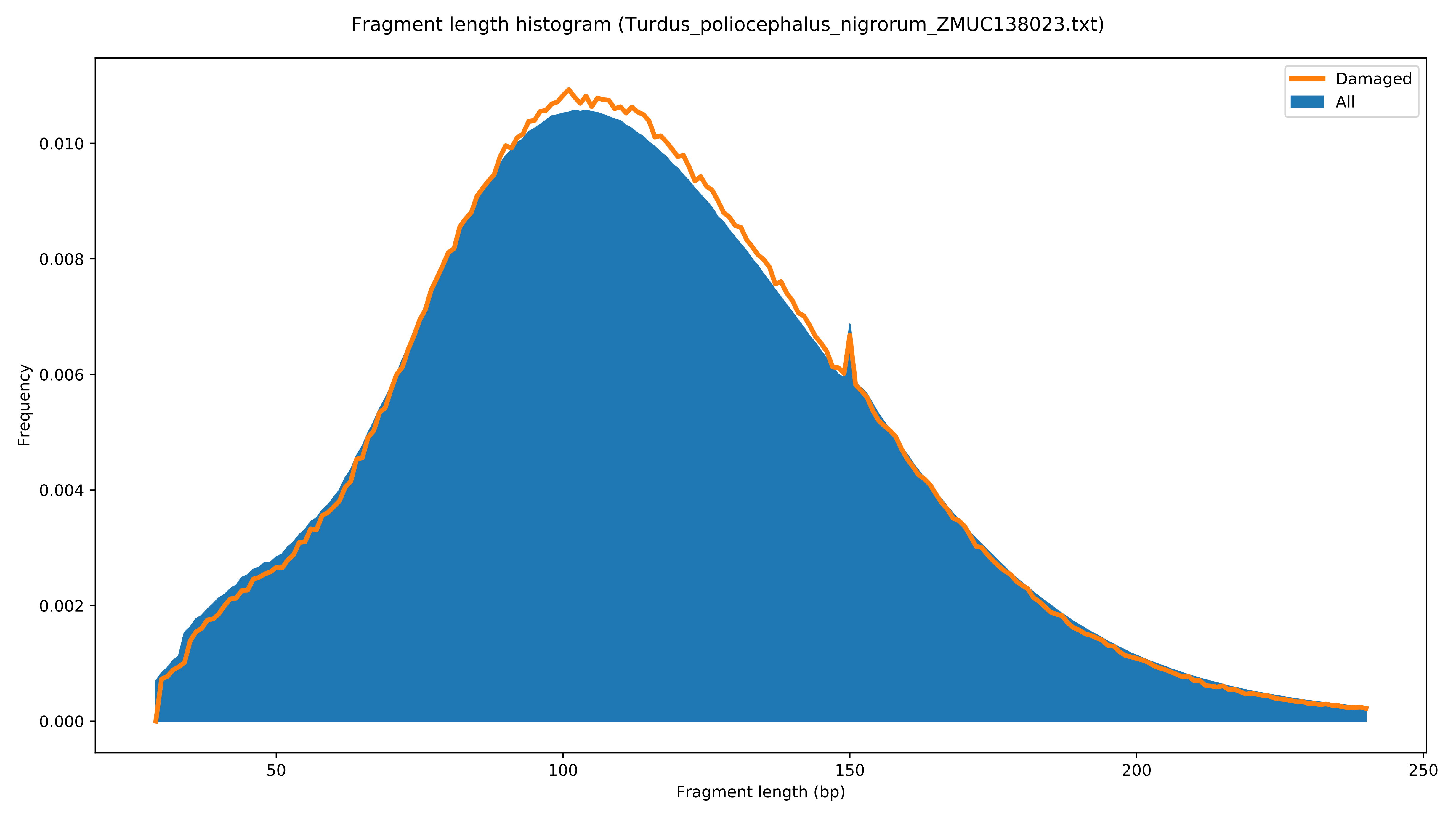

Fragment length histogram (Turdus_poliocephalus_nigrorum_ZMUC138023.txt)
Damaged
All
Frequency
0.000
0.002
0.004
0.006
0.008
0.010
50
100
150
200
250
Fragment length (bp)

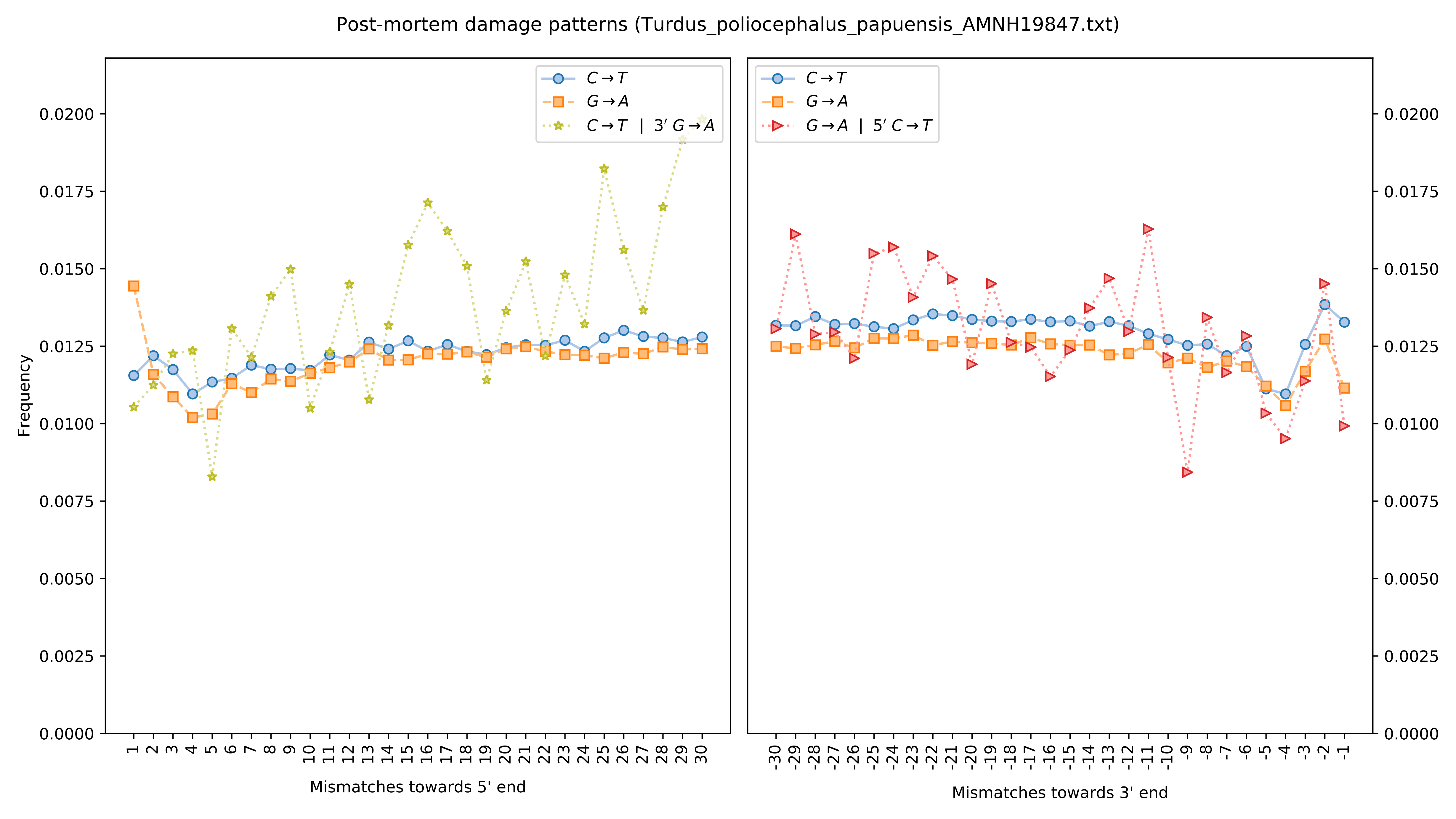

Post-mortem damage patterns (Turdus_poliocephalus_papuensis_AMNH19847.txt)
Frequency
Mismatches towards 5' end
Mismatches towards 3' end
C→T
G→A
C→T | 3′ G→A
G→A | 5′ C→T
0.0000
0.0025
0.0050
0.0075
0.0100
0.0125
0.0150
0.0175
0.0200

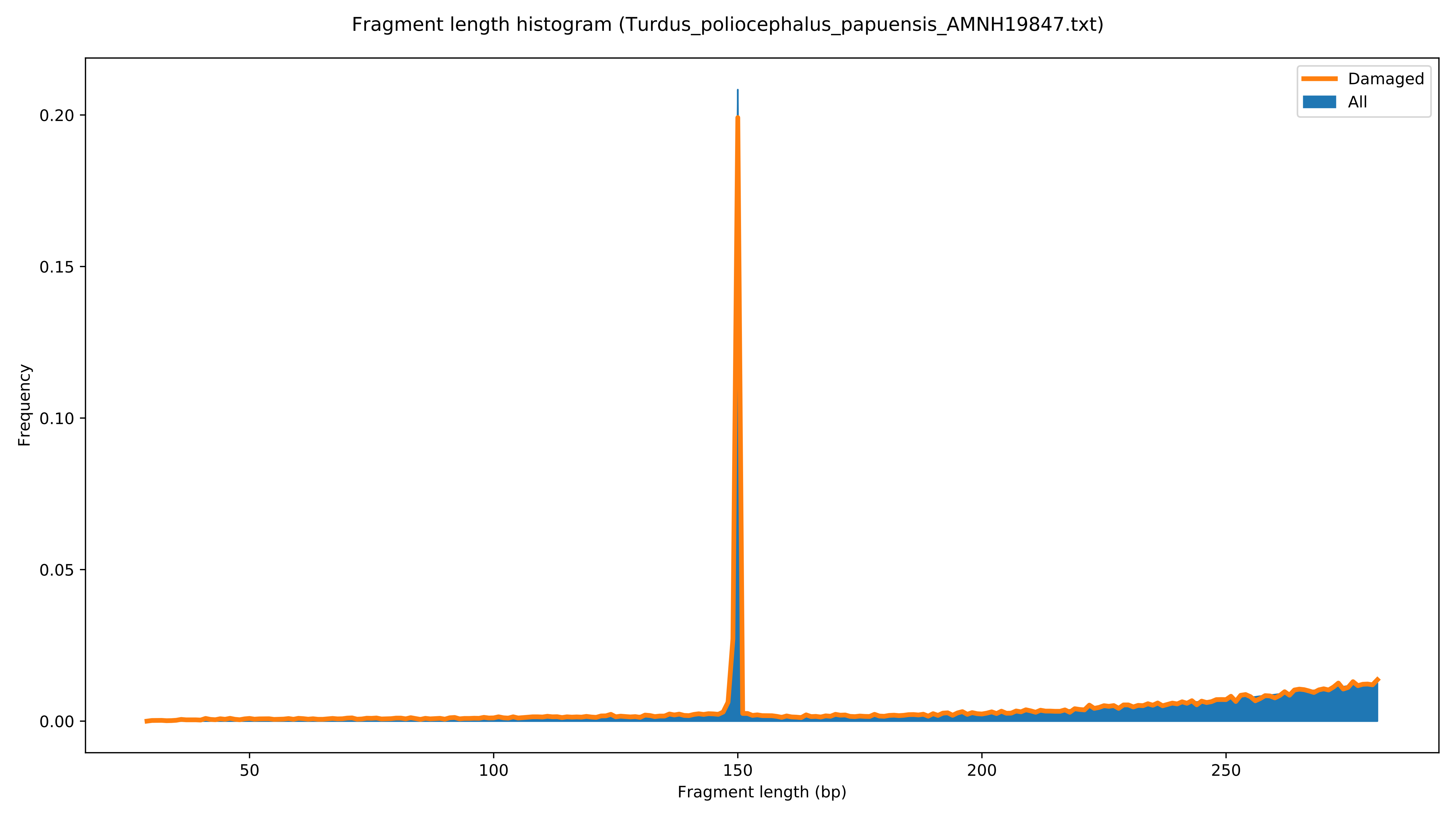
Fragment length histogram (Turdus_poliocephalus_papuensis_AMNH19847.txt)
Damaged
All
Frequency
0.00
0.05
0.10
0.15
0.20
50
100
150
200
250
Fragment length (bp)

Post-mortem damage patterns (Turdus_poliocephalus_papuensis_ZMUC150143.txt)

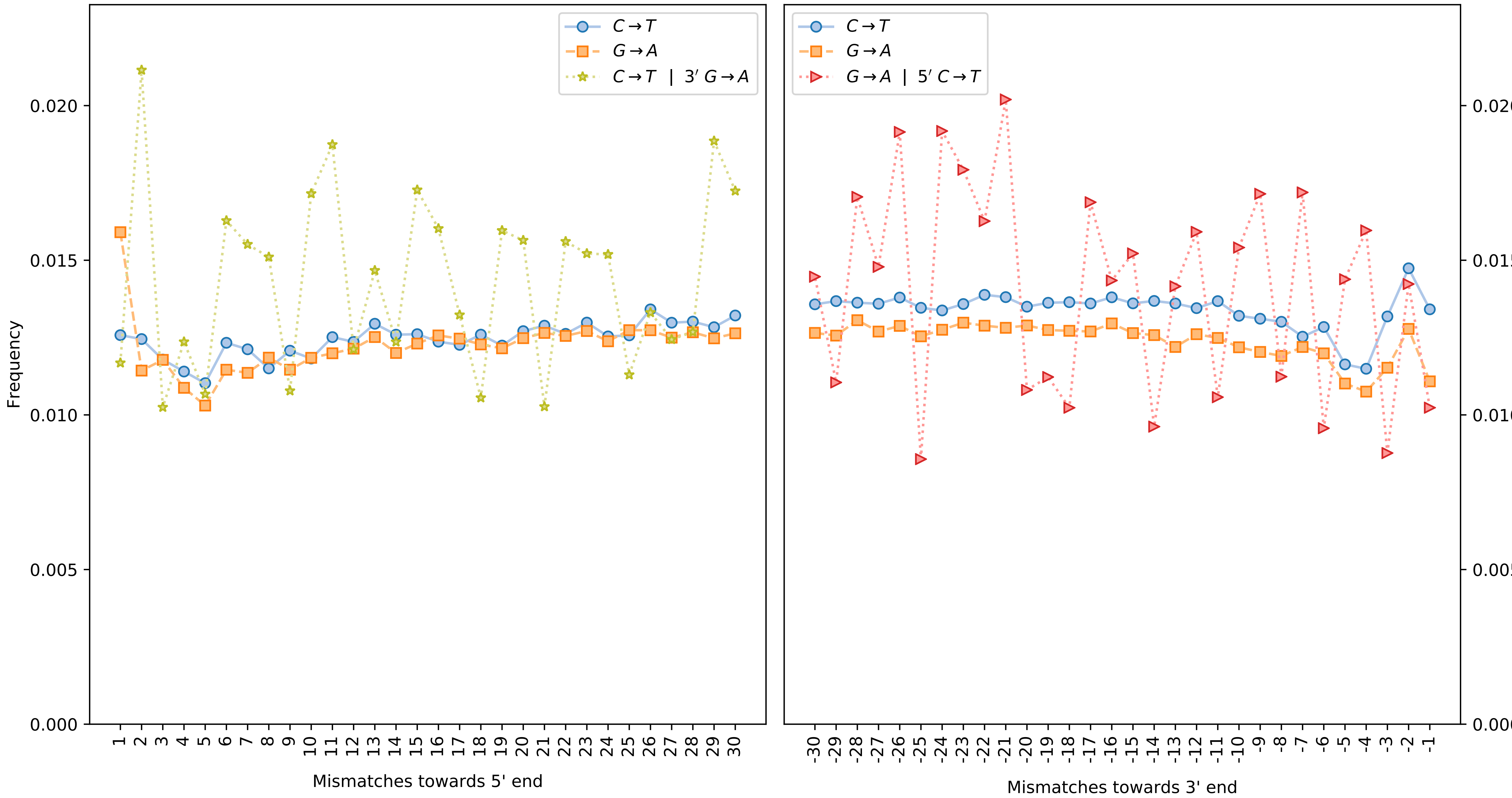

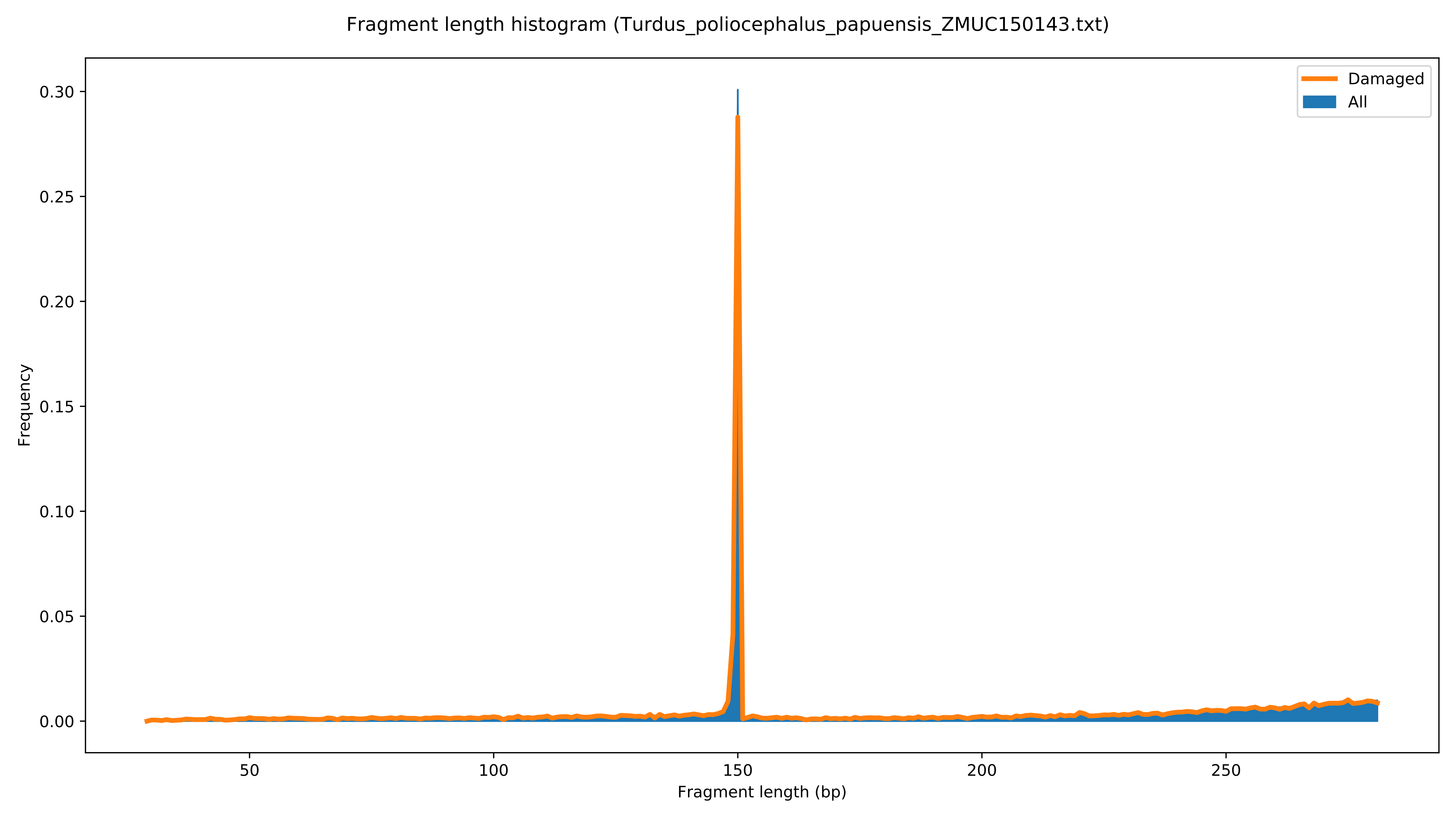
Fragment length histogram (Turdus_poliocephalus_papuensis_ZMUC150143.txt)
Damaged
All
Frequency
0.00
0.05
0.10
0.15
0.20
0.25
0.30
50
100
150
200
250
Fragment length (bp)

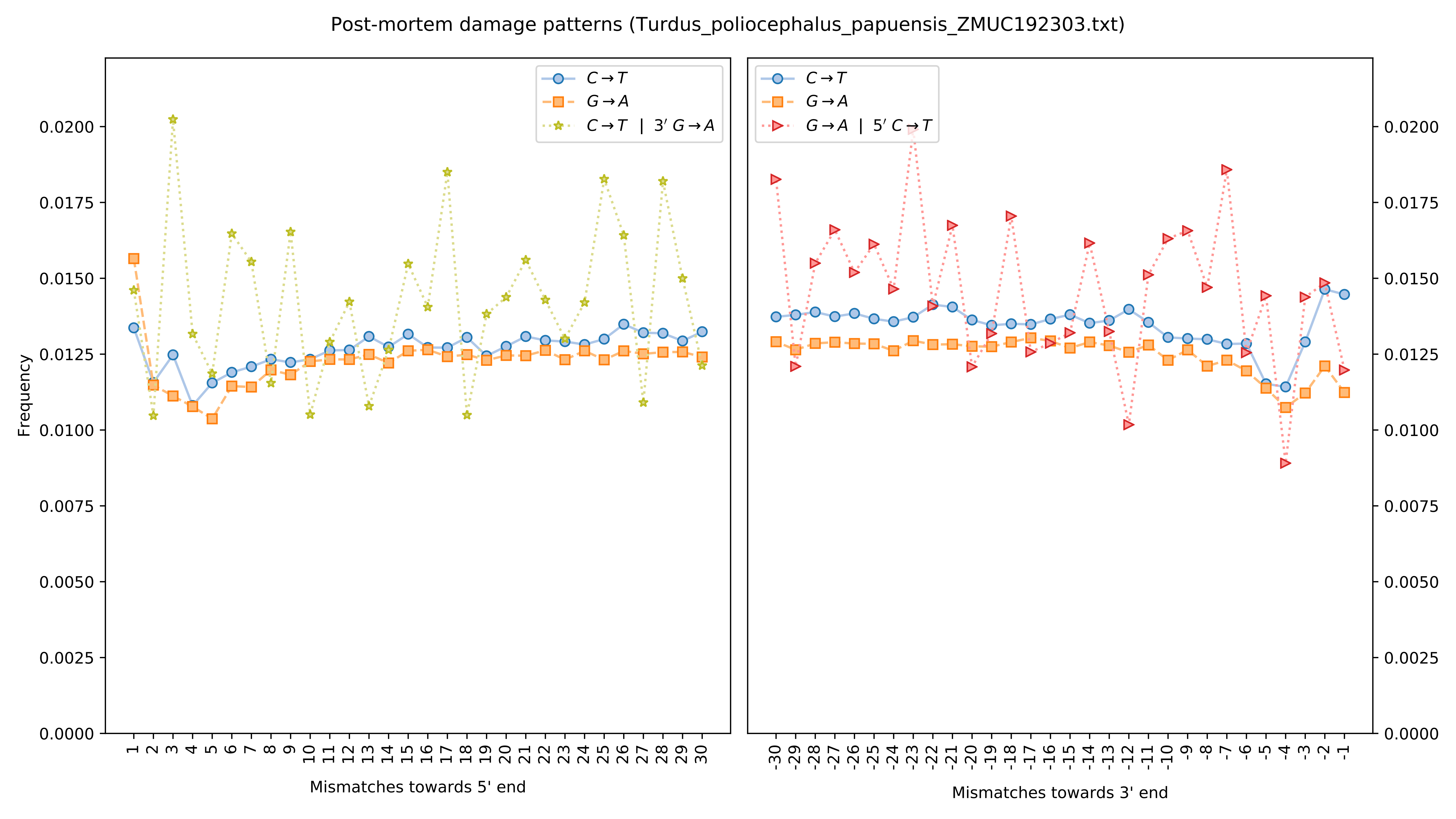

Post-mortem damage patterns (Turdus_poliocephalus_papuensis_ZMUC192303.txt)
C→T
G→A
C→T | 3′ G→A
G→A | 5′ C→T
Frequency
0.0000
0.0025
0.0050
0.0075
0.0100
0.0125
0.0150
0.0175
0.0200
Mismatches towards 5' end
Mismatches towards 3' end

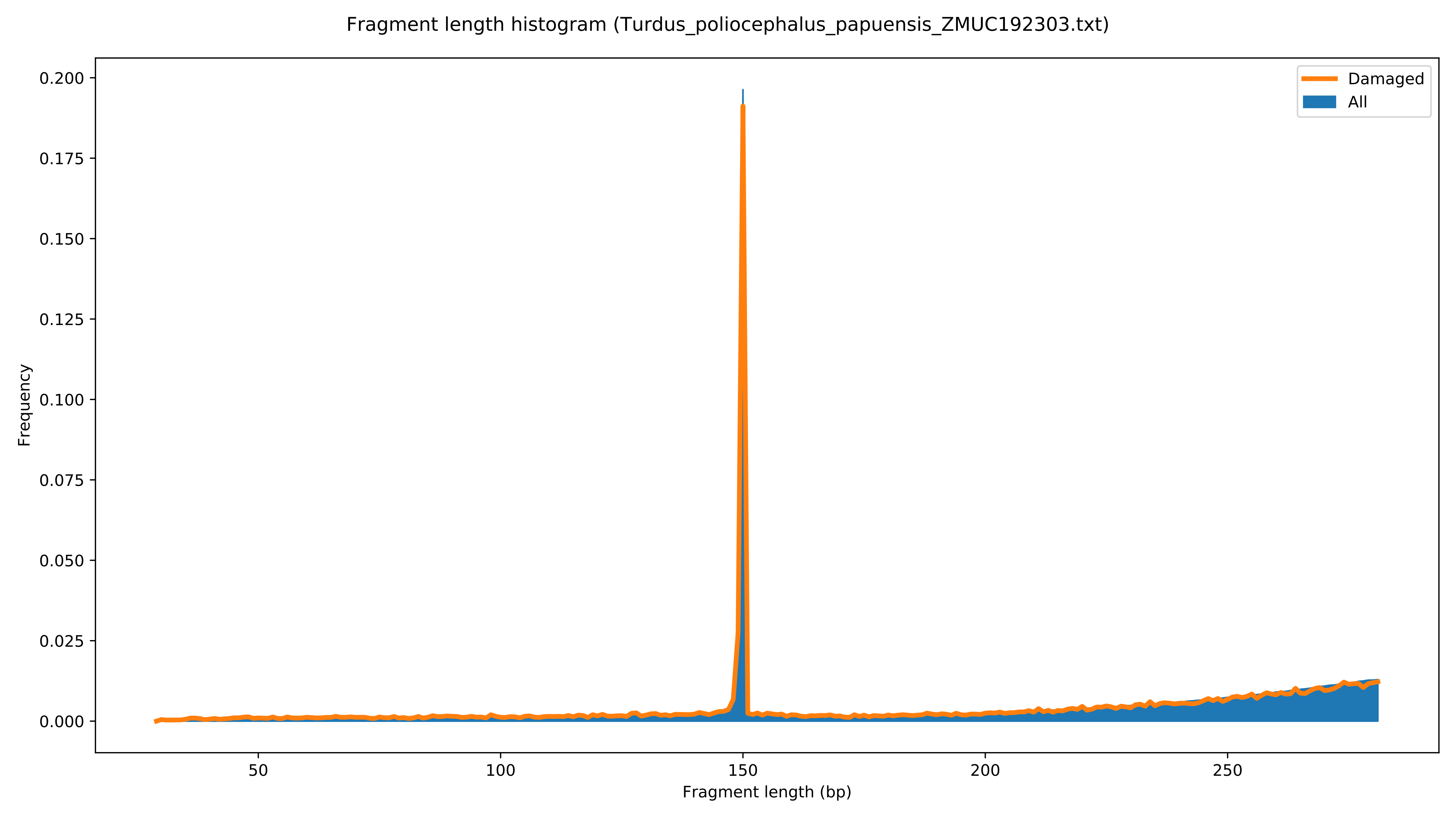

Fragment length histogram (Turdus_poliocephalus_papuensis_ZMUC192303.txt)
Damaged
All
Frequency
0.200
0.175
0.150
0.125
0.100
0.075
0.050
0.025
0.000
50
100
150
200
250
Fragment length (bp)

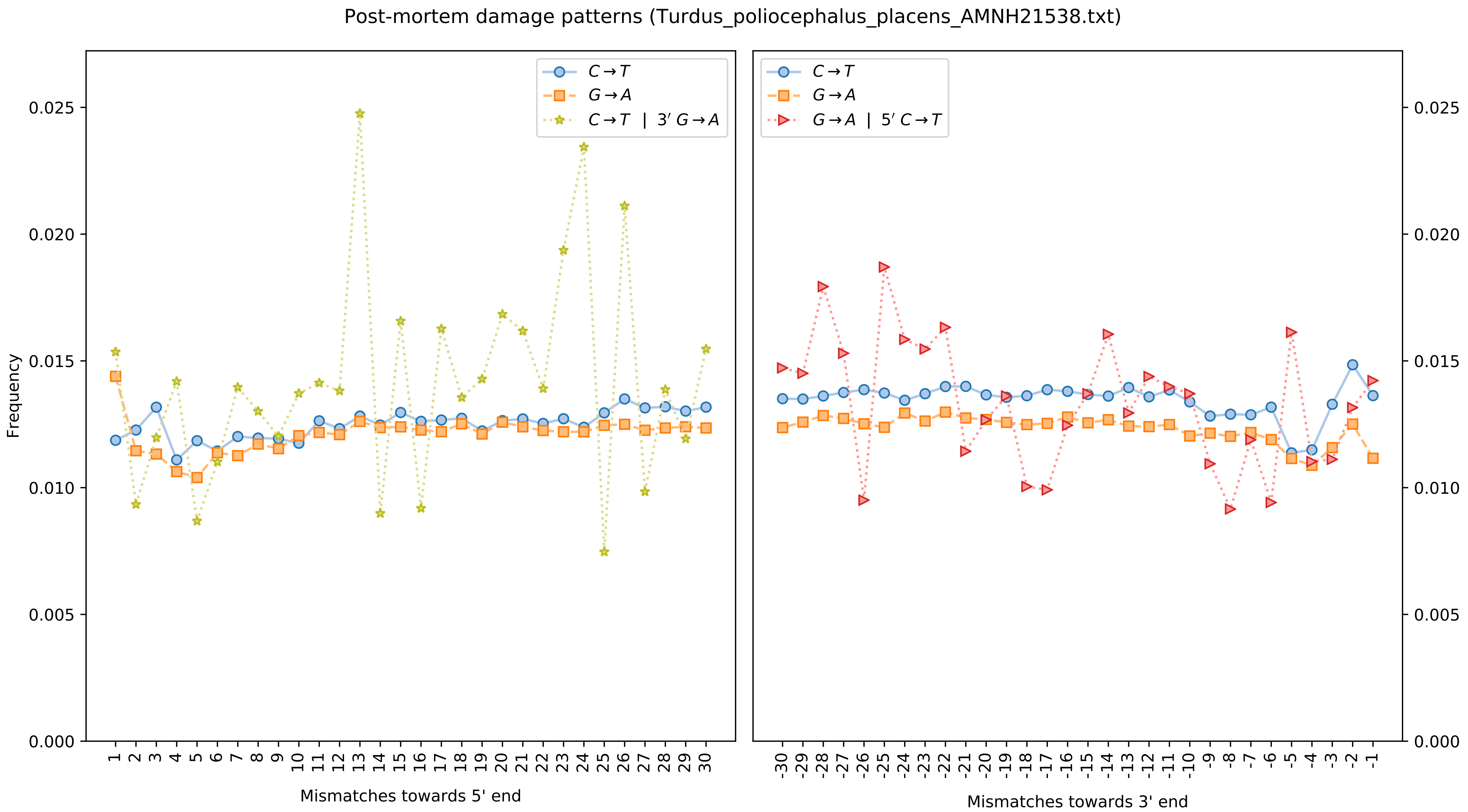

Post-mortem damage patterns (Turdus_poliocephalus_placens_AMNH21538.txt)
Frequency
$C \to T$
$G \to A$
$C \to T$ | $3' G \to A$
$G \to A$ | $5' C \to T$
0.000
0.005
0.010
0.015
0.020
0.025
Mismatches towards 5' end
Mismatches towards 3' end

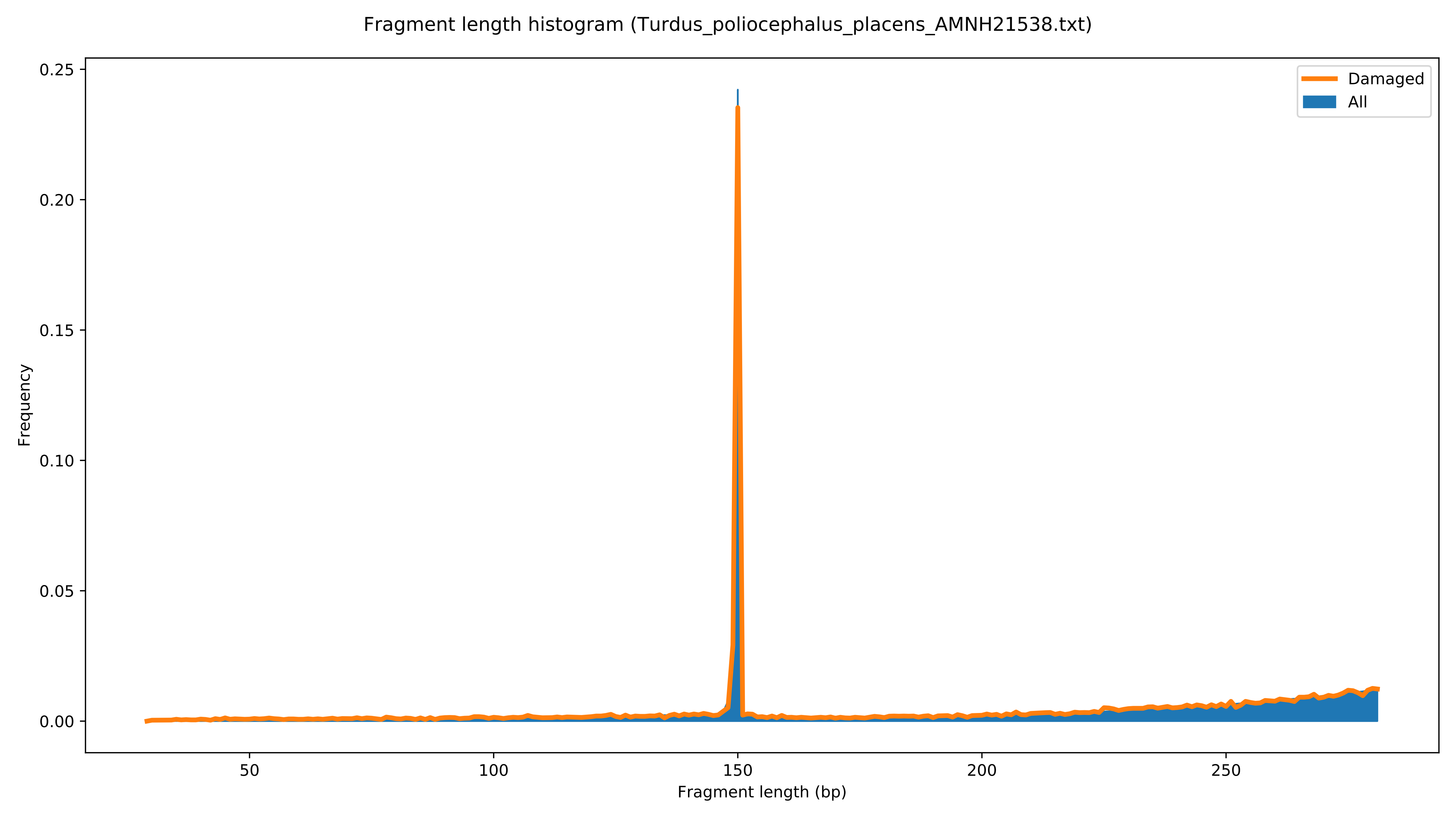
Fragment length histogram (Turdus_poliocephalus_placens_AMNH21538.txt)
Damaged
All
Frequency
0.25
0.20
0.15
0.10
0.05
0.00
50
100
150
200
250
Fragment length (bp)

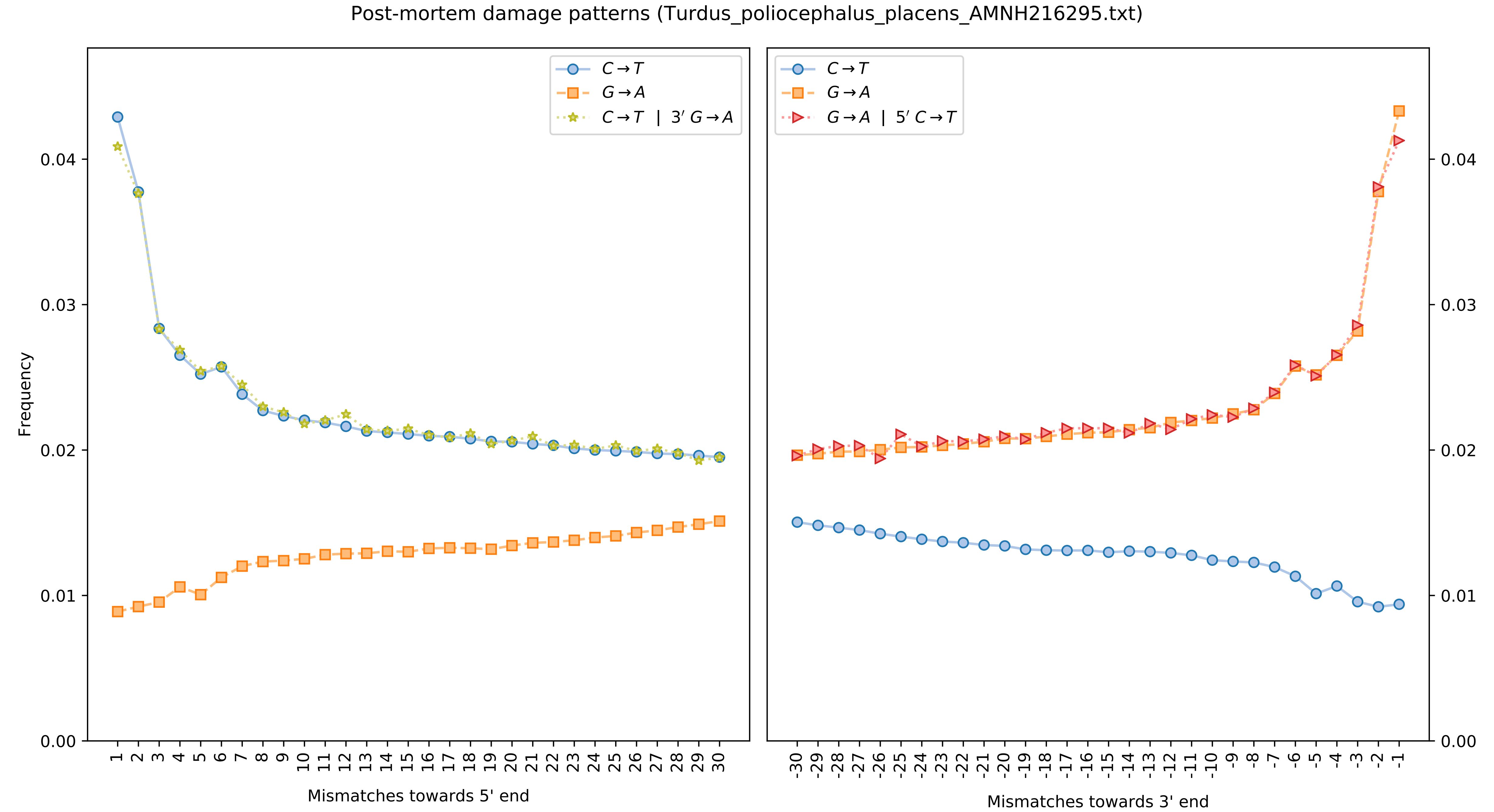
Post-mortem damage patterns (Turdus_poliocephalus_placens_AMNH216295.txt)
Frequency
Mismatches towards 5' end
Mismatches towards 3' end
C→T
G→A
C→T | 3′ G→A
G→A | 5′ C→T

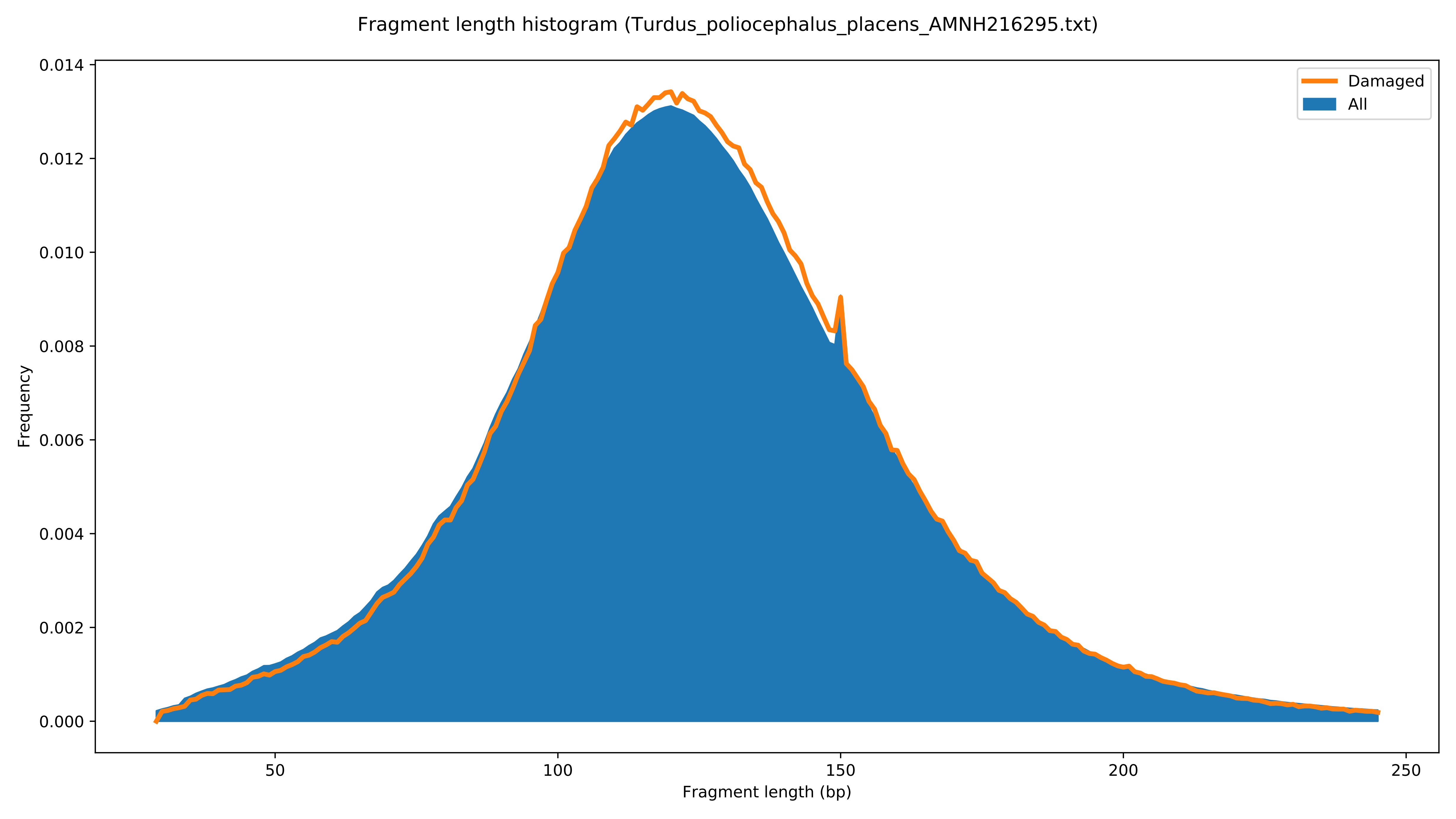

Fragment length histogram (Turdus_poliocephalus_placens_AMNH216295.txt)
Damaged
All
Frequency
0.000
0.002
0.004
0.006
0.008
0.010
0.012
0.014
50
100
150
200
250
Fragment length (bp)

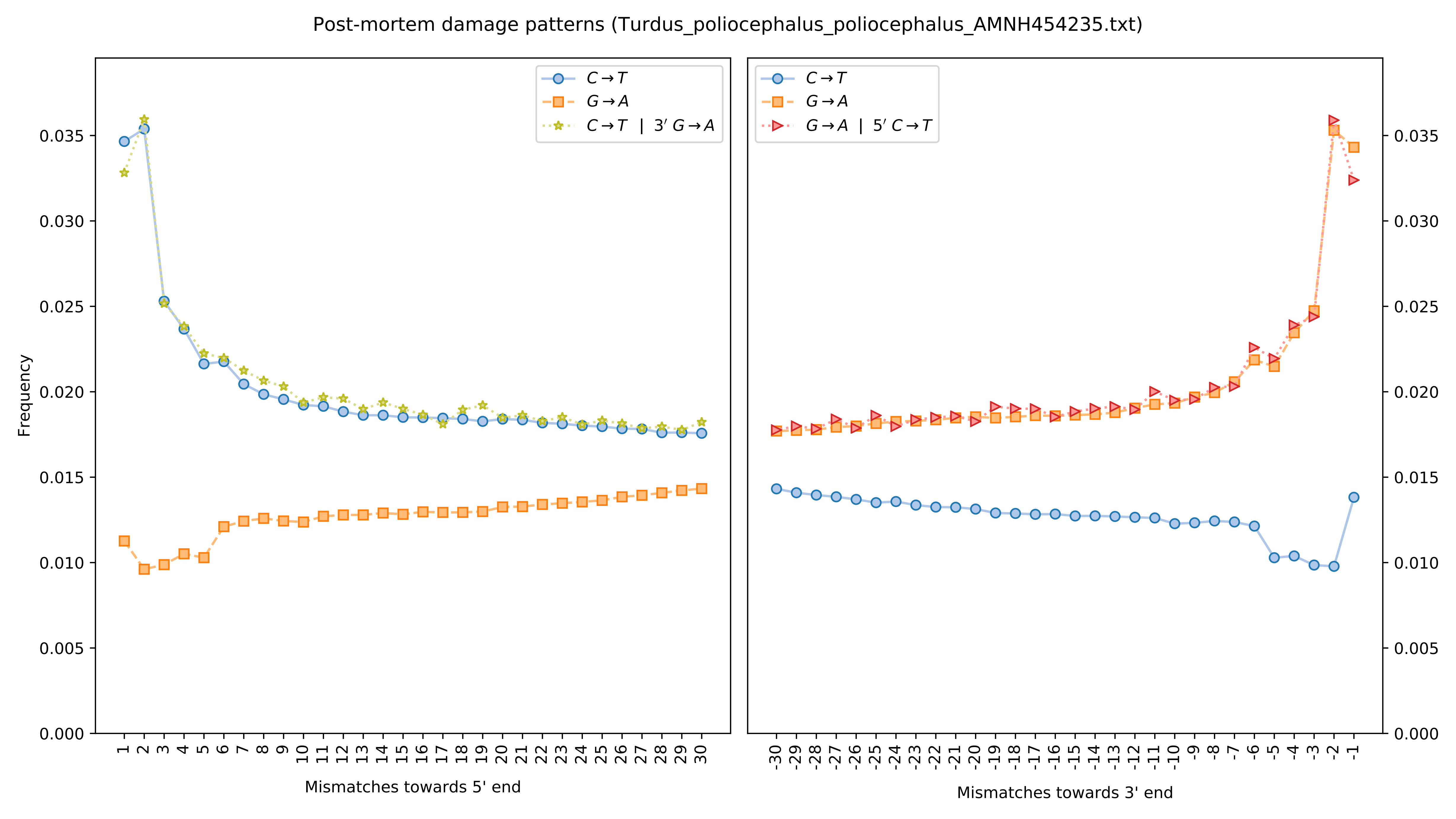

Post-mortem damage patterns (Turdus_poliocephalus_poliocephalus_AMNH454235.txt)
Frequency
Mismatches towards 5' end
Mismatches towards 3' end
C→T
G→A
C→T | 3′ G→A
G→A | 5′ C→T

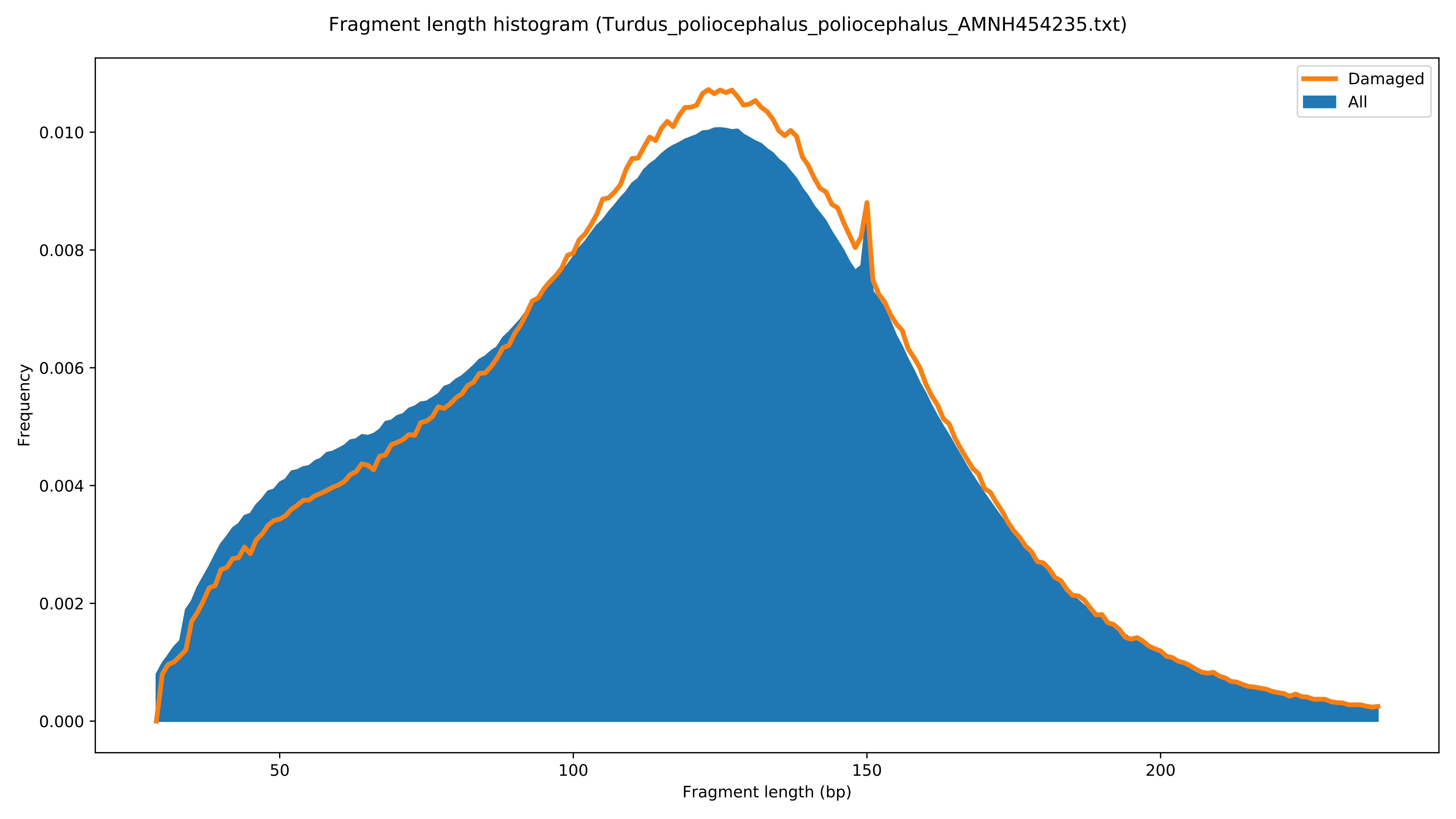

Fragment length histogram (Turdus_poliocephalus_poliocephalus_AMNH454235.txt)
Damaged
All
Frequency
0.000
0.002
0.004
0.006
0.008
0.010
50
100
150
200
Fragment length (bp)

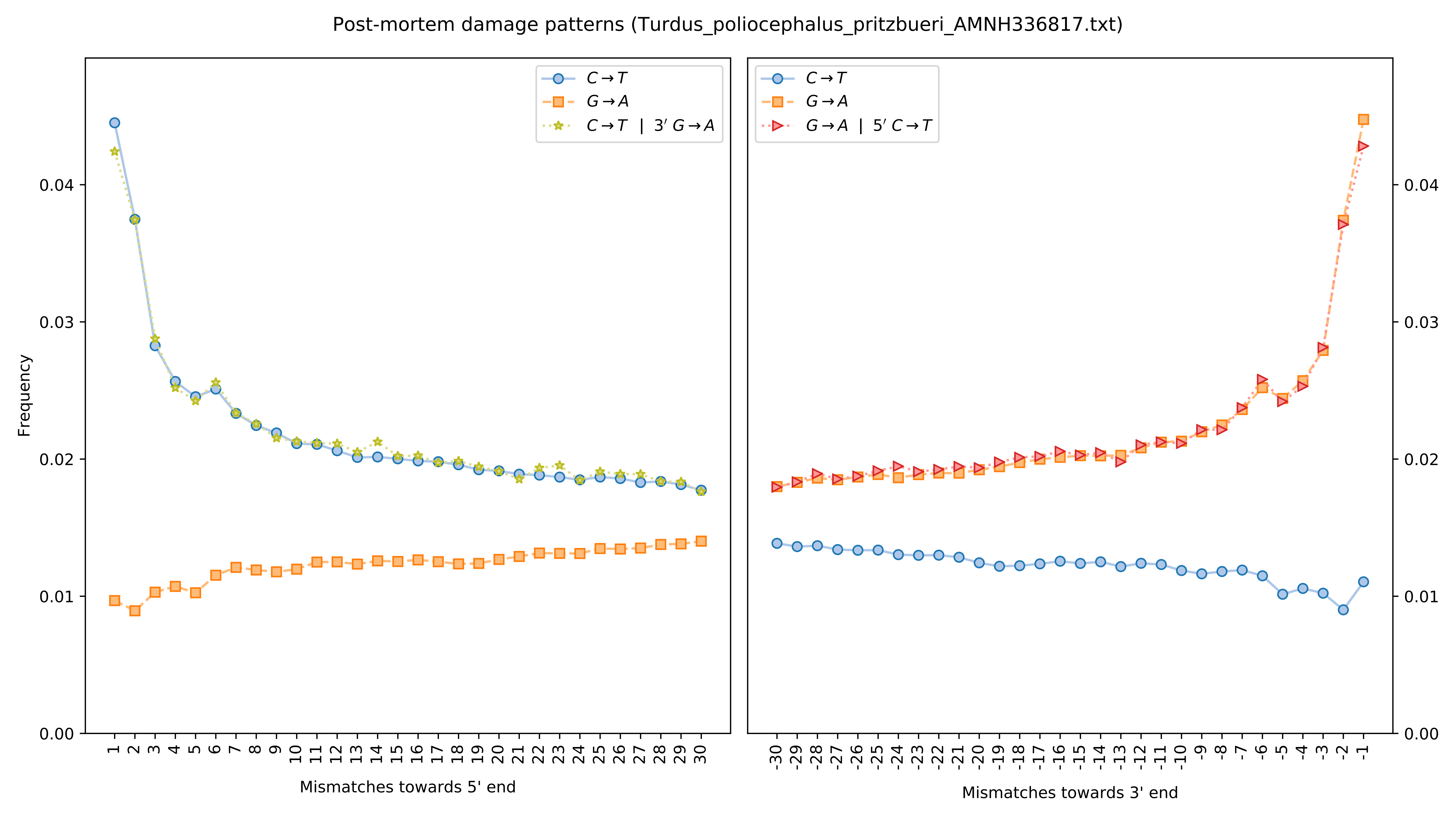

Post-mortem damage patterns (Turdus_poliocephalus_pritzbueri_AMNH336817.txt)
Frequency
Mismatches towards 5' end
Mismatches towards 3' end
C→T
G→A
C→T | 3′ G→A
G→A | 5′ C→T
0.00
0.01
0.02
0.03
0.04

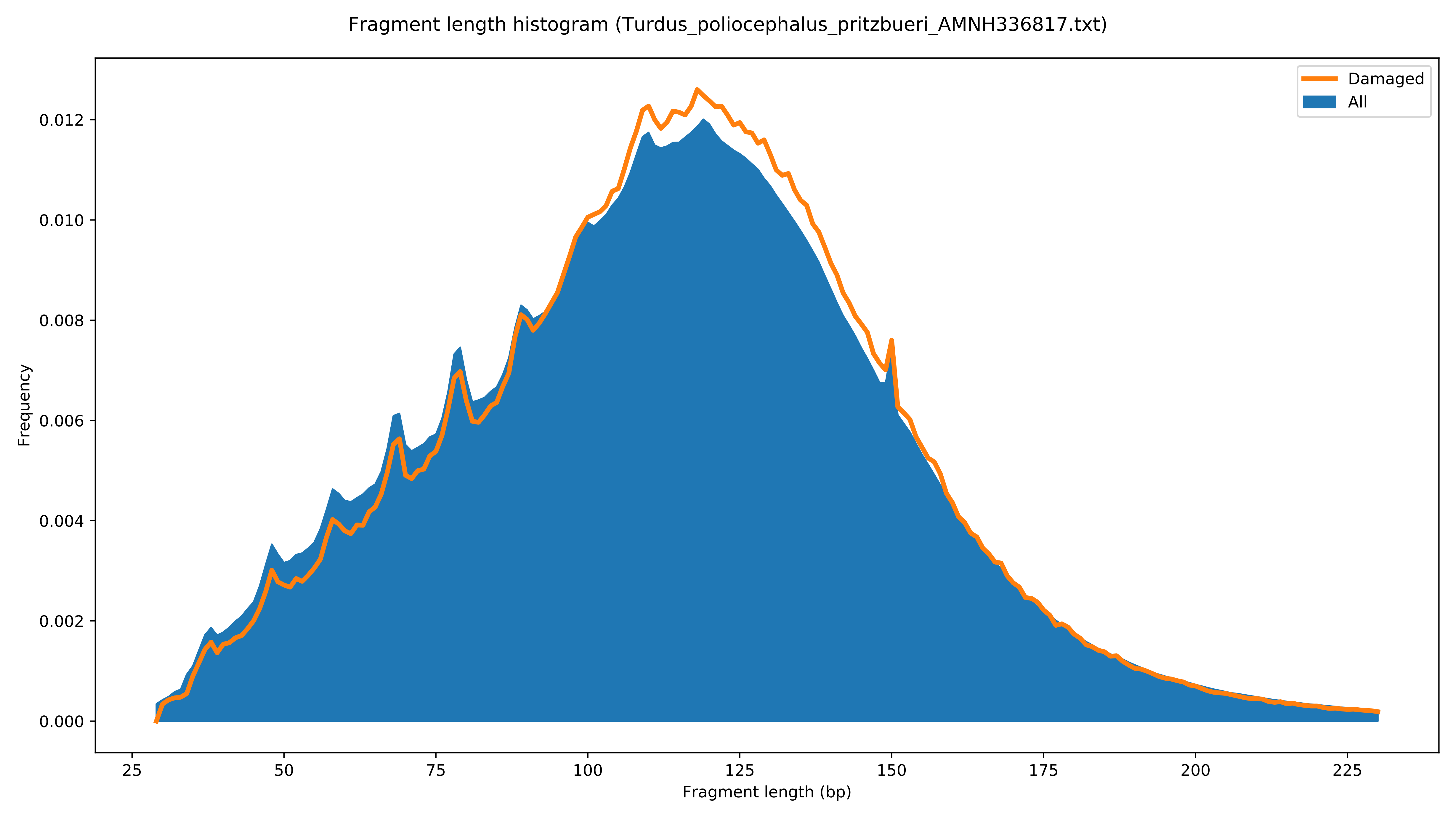

Fragment length histogram (Turdus_poliocephalus_pritzbueri_AMNH336817.txt)
Damaged
All
Frequency
0.000
0.002
0.004
0.006
0.008
0.010
0.012
25
50
75
100
125
150
175
200
225
Fragment length (bp)

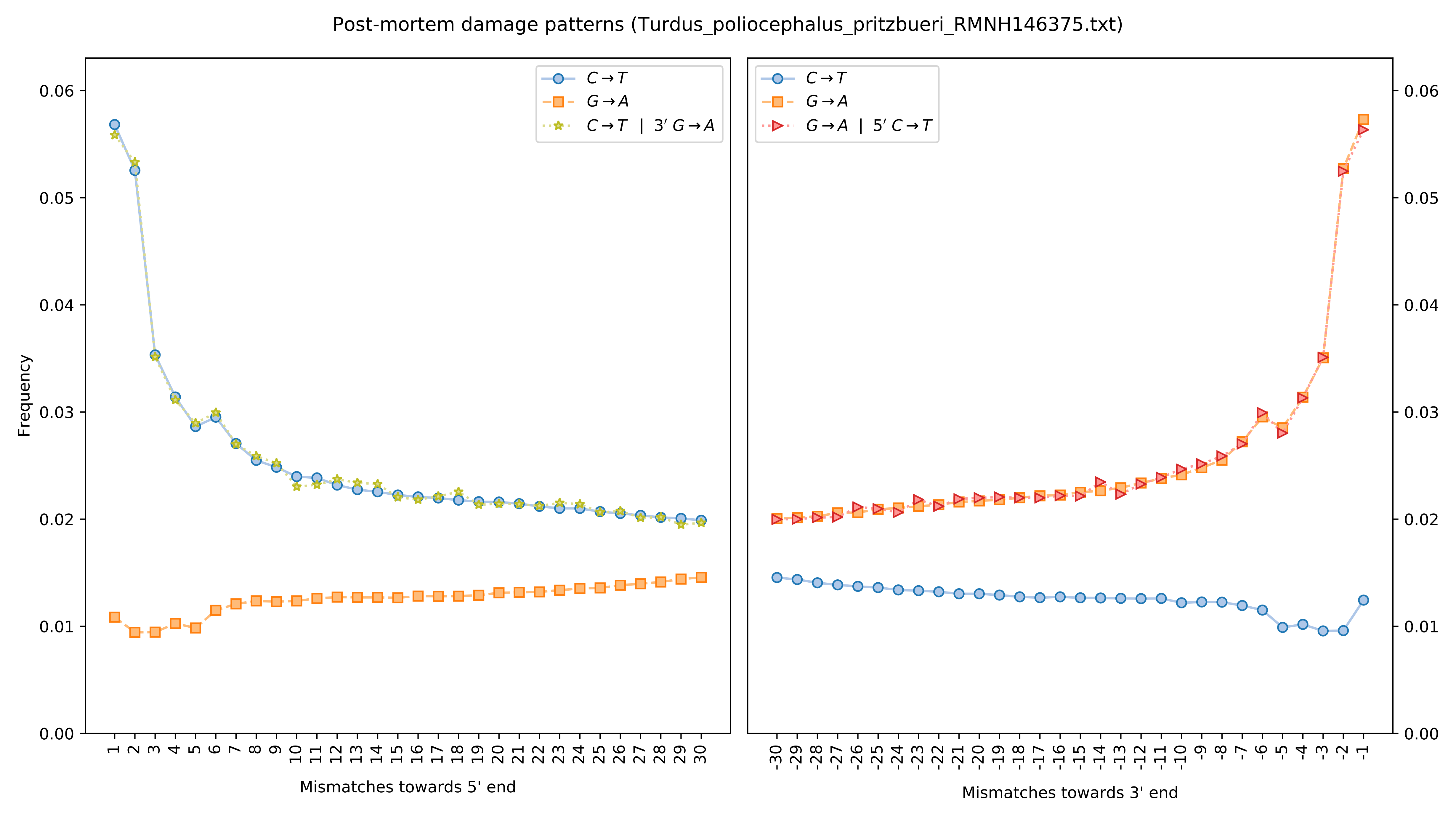

Post-mortem damage patterns (Turdus_poliocephalus_pritzbueri_RMNH146375.txt)
Frequency
0.00
0.01
0.02
0.03
0.04
0.05
0.06
C→T
G→A
C→T | 3′ G→A
G→A | 5′ C→T
1 2 3 4 5 6 7 8 9 10 11 12 13 14 15 16 17 18 19 20 21 22 23 24 25 26 27 28 29 30
Mismatches towards 5' end
-30 -29 -28 -27 -26 -25 -24 -23 -22 -21 -20 -19 -18 -17 -16 -15 -14 -13 -12 -11 -10 -9 -8 -7 -6 -5 -4 -3 -2 -1
Mismatches towards 3' end

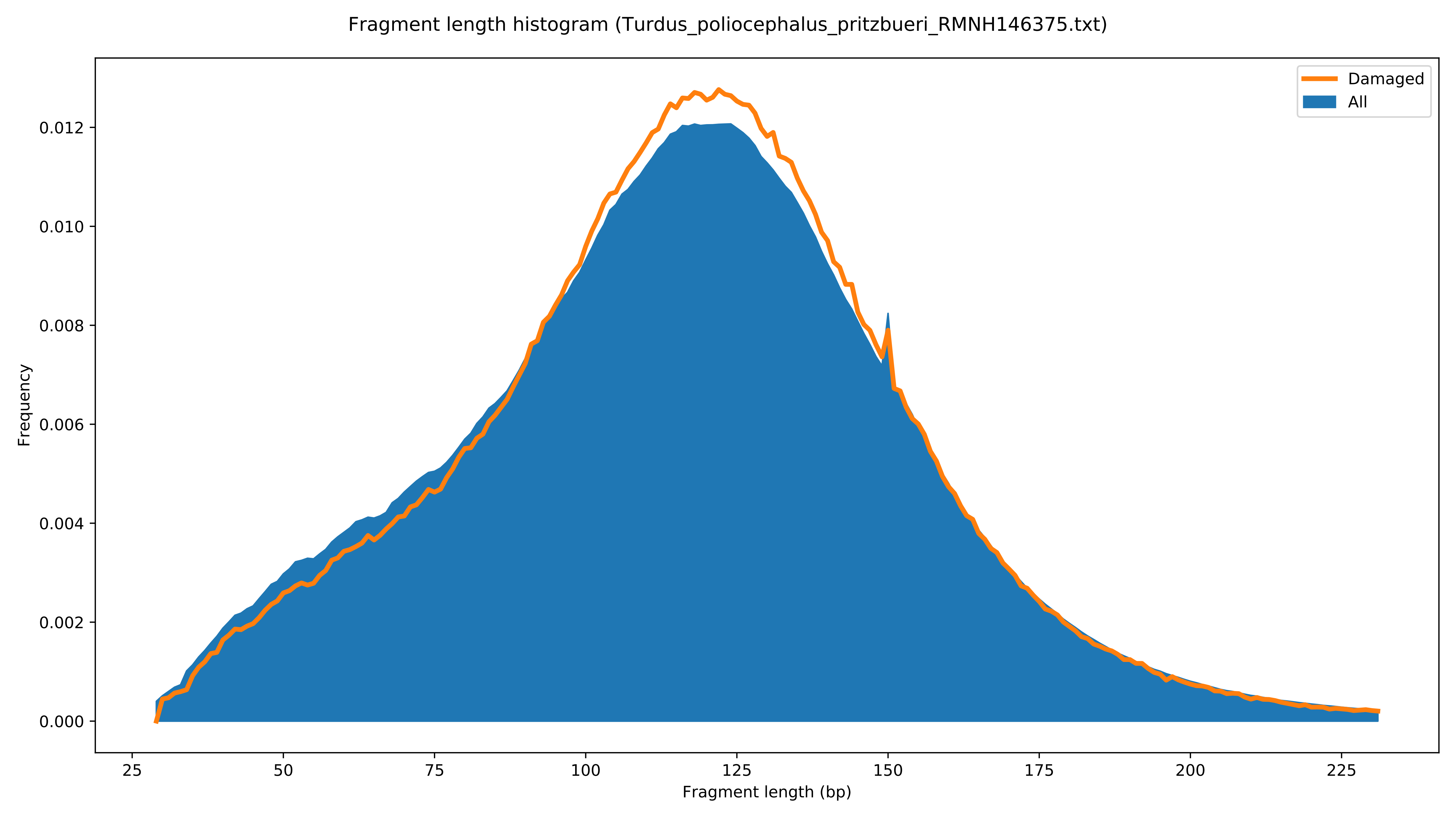

Fragment length histogram (Turdus_poliocephalus_pritzbueri_RMNH146375.txt)
Damaged
All
Frequency
0.000
0.002
0.004
0.006
0.008
0.010
0.012
25
50
75
100
125
150
175
200
225
Fragment length (bp)

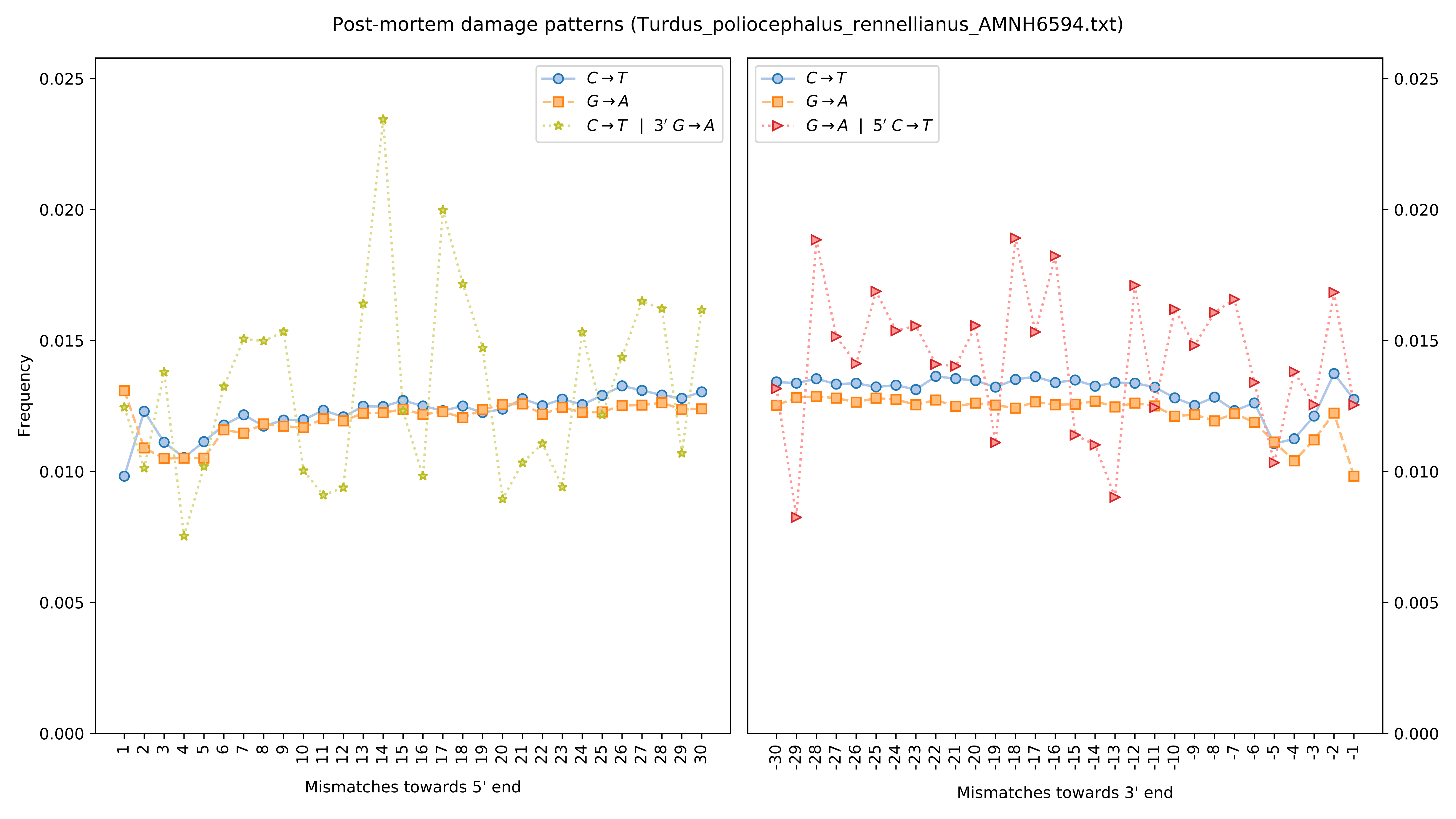

Post-mortem damage patterns (Turdus_poliocephalus_rennellianus_AMNH6594.txt)
Frequency
Mismatches towards 5' end
Mismatches towards 3' end
C→T
G→A
C→T | 3′ G→A
G→A | 5′ C→T
0.000
0.005
0.010
0.015
0.020
0.025

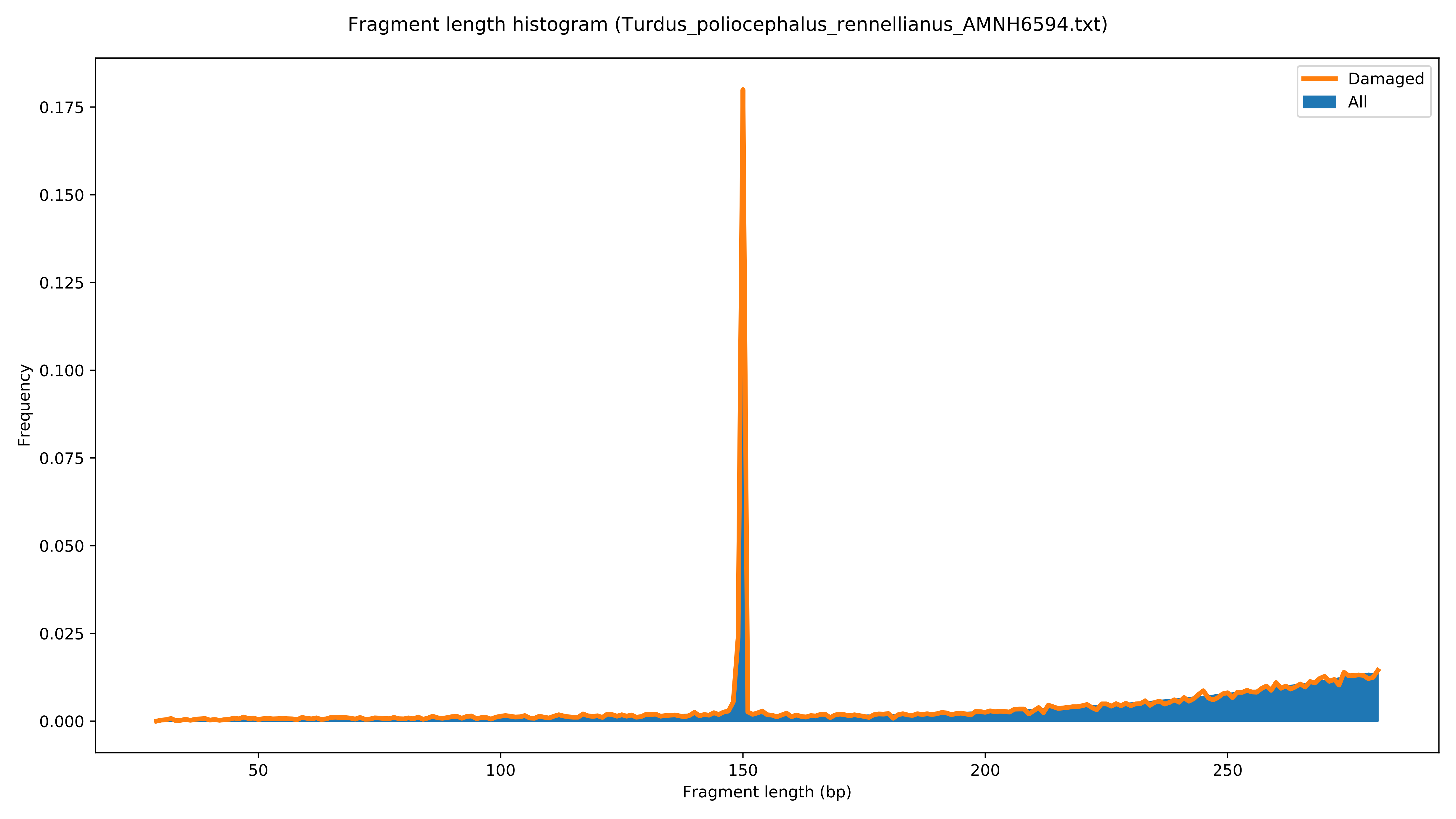

Fragment length histogram (Turdus_poliocephalus_rennellianus_AMNH6594.txt)
Damaged
All
Frequency
Fragment length (bp)
0.000
0.025
0.050
0.075
0.100
0.125
0.150
0.175
50
100
150
200
250

Post-mortem damage patterns (Turdus_poliocephalus_ruficeps_AMNH252598.txt)

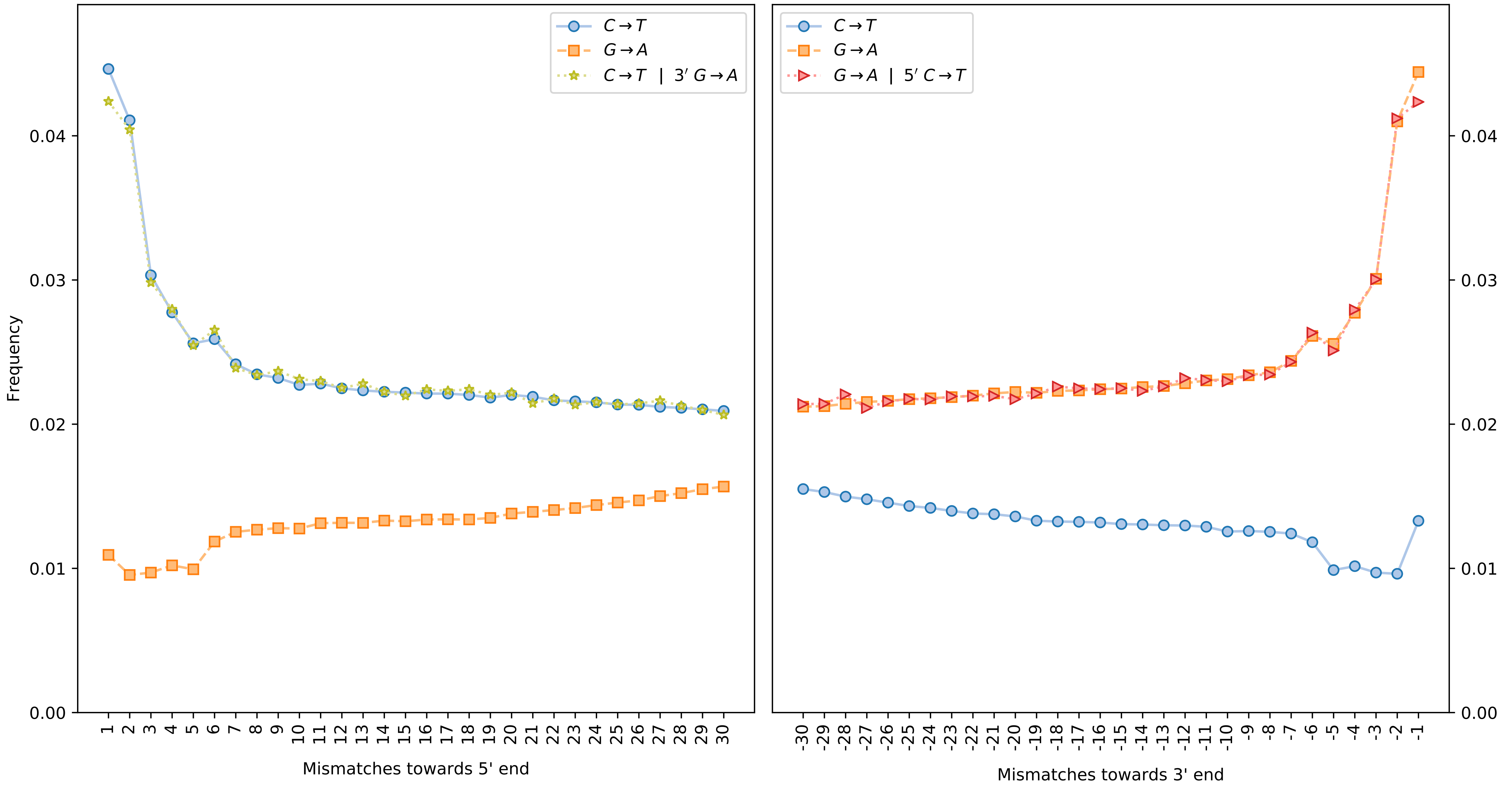

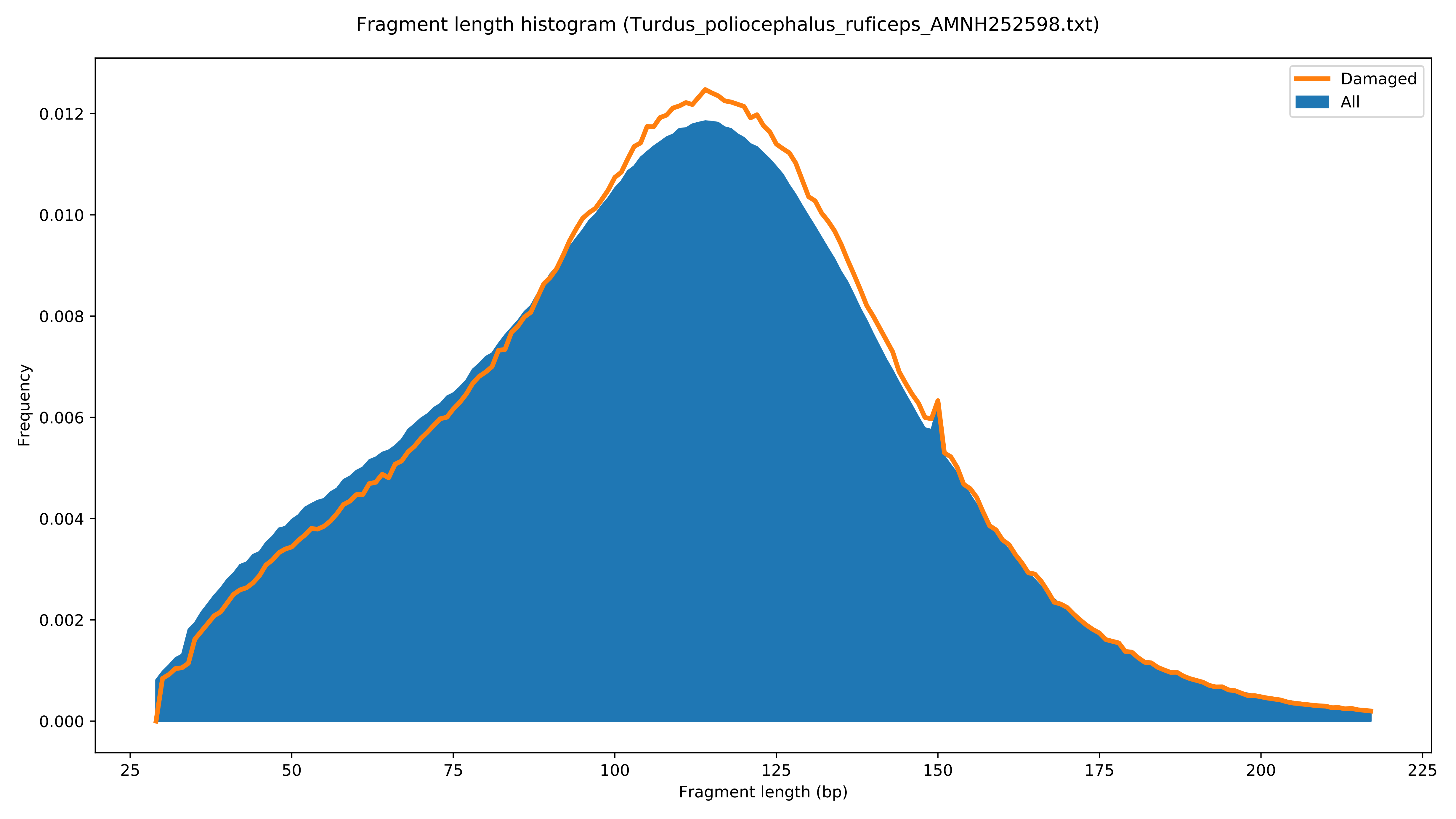
Fragment length histogram (Turdus_poliocephalus_ruficeps_AMNH252598.txt)
Damaged
All
Frequency
0.000
0.002
0.004
0.006
0.008
0.010
0.012
25
50
75
100
125
150
175
200
225
Fragment length (bp)

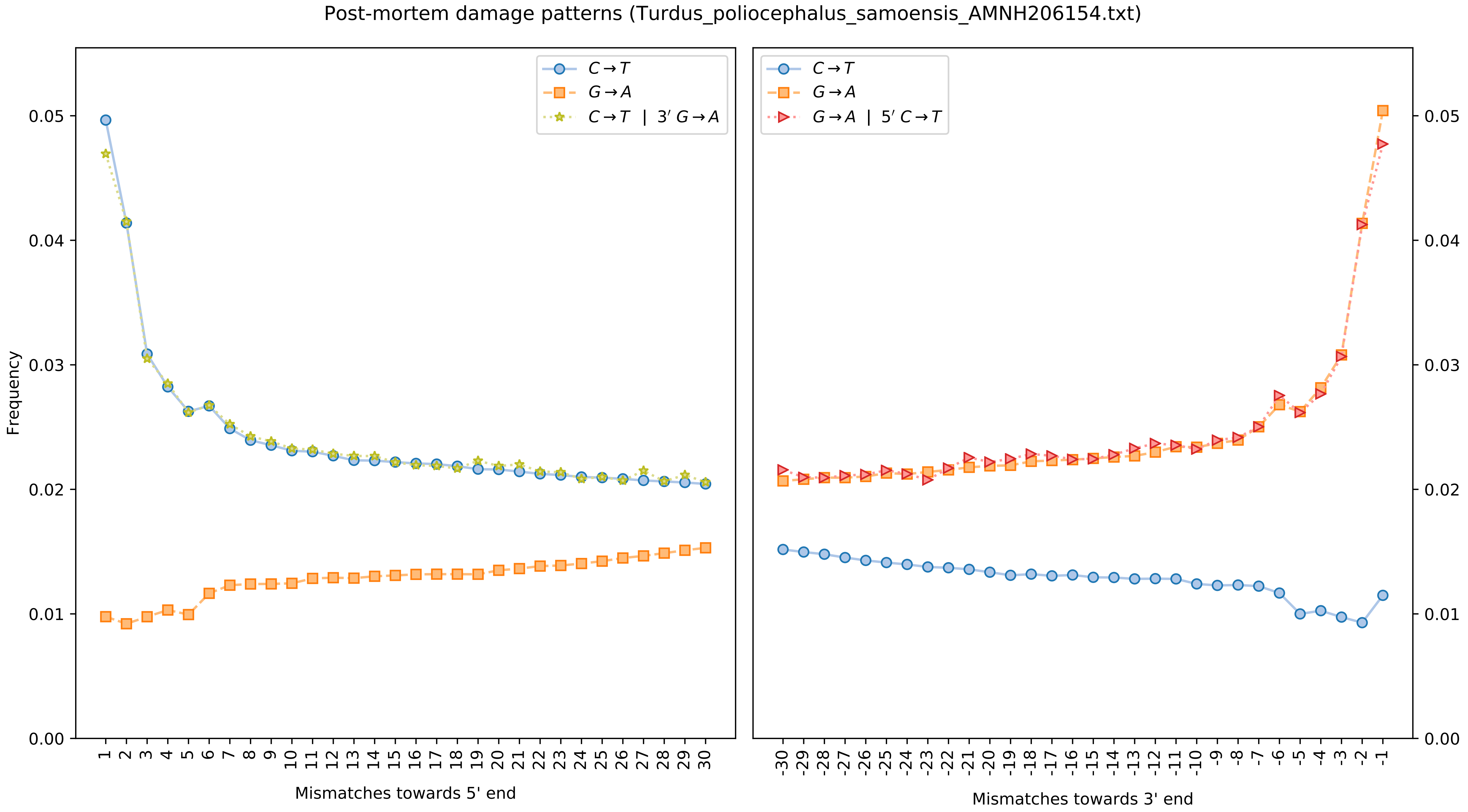

Post-mortem damage patterns (Turdus_poliocephalus_samoensis_AMNH206154.txt)
Frequency
Mismatches towards 5' end
Mismatches towards 3' end
C→T
G→A
C→T | 3′ G→A
G→A | 5′ C→T

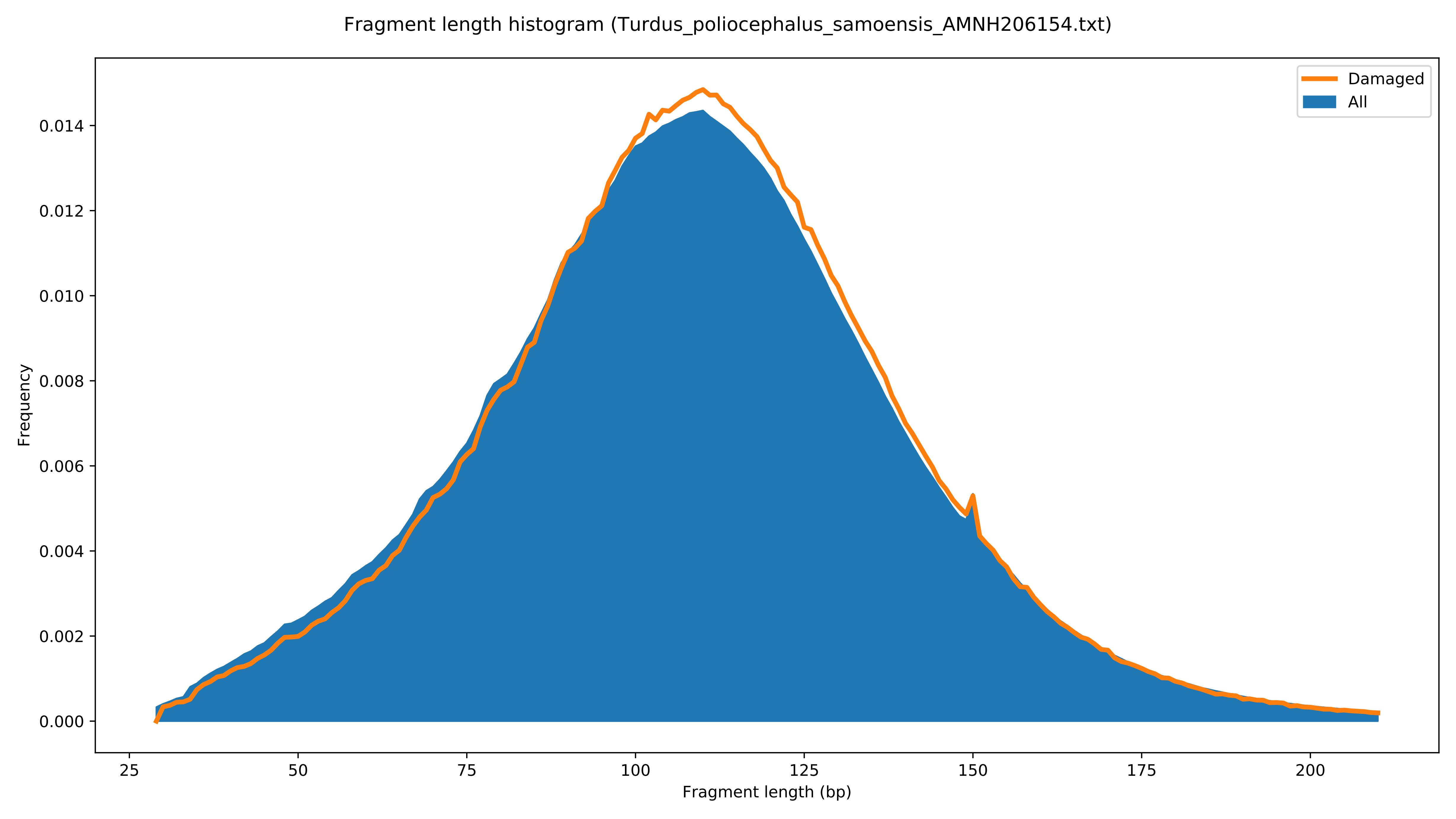

Fragment length histogram (Turdus_poliocephalus_samoensis_AMNH206154.txt)
Damaged
All
Frequency
0.000
0.002
0.004
0.006
0.008
0.010
0.012
0.014
25
50
75
100
125
150
175
200
Fragment length (bp)

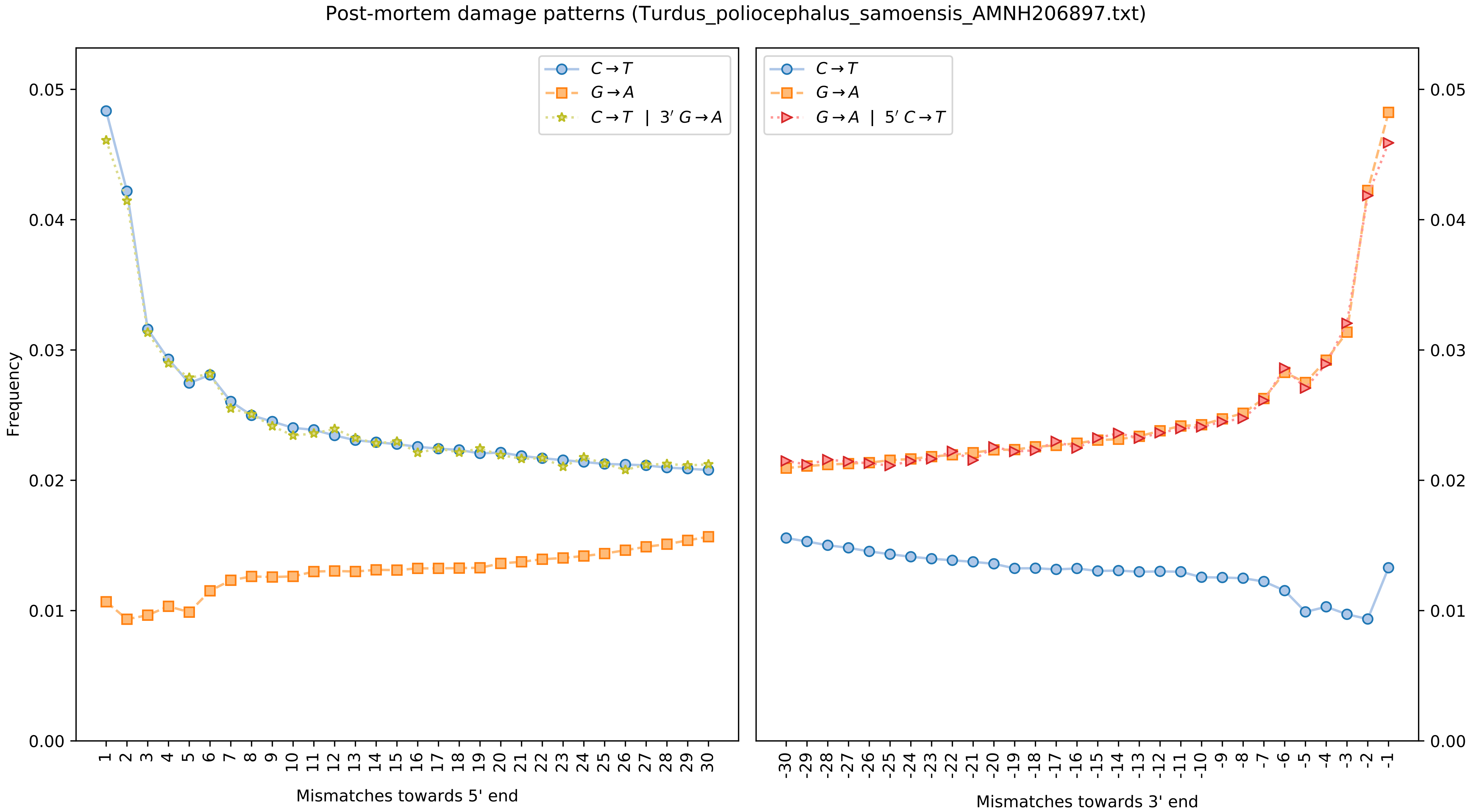

Post-mortem damage patterns (Turdus_poliocephalus_samoensis_AMNH206897.txt)
C→T
G→A
C→T | 3′ G→A
G→A | 5′ C→T
Frequency
Mismatches towards 5' end
Mismatches towards 3' end

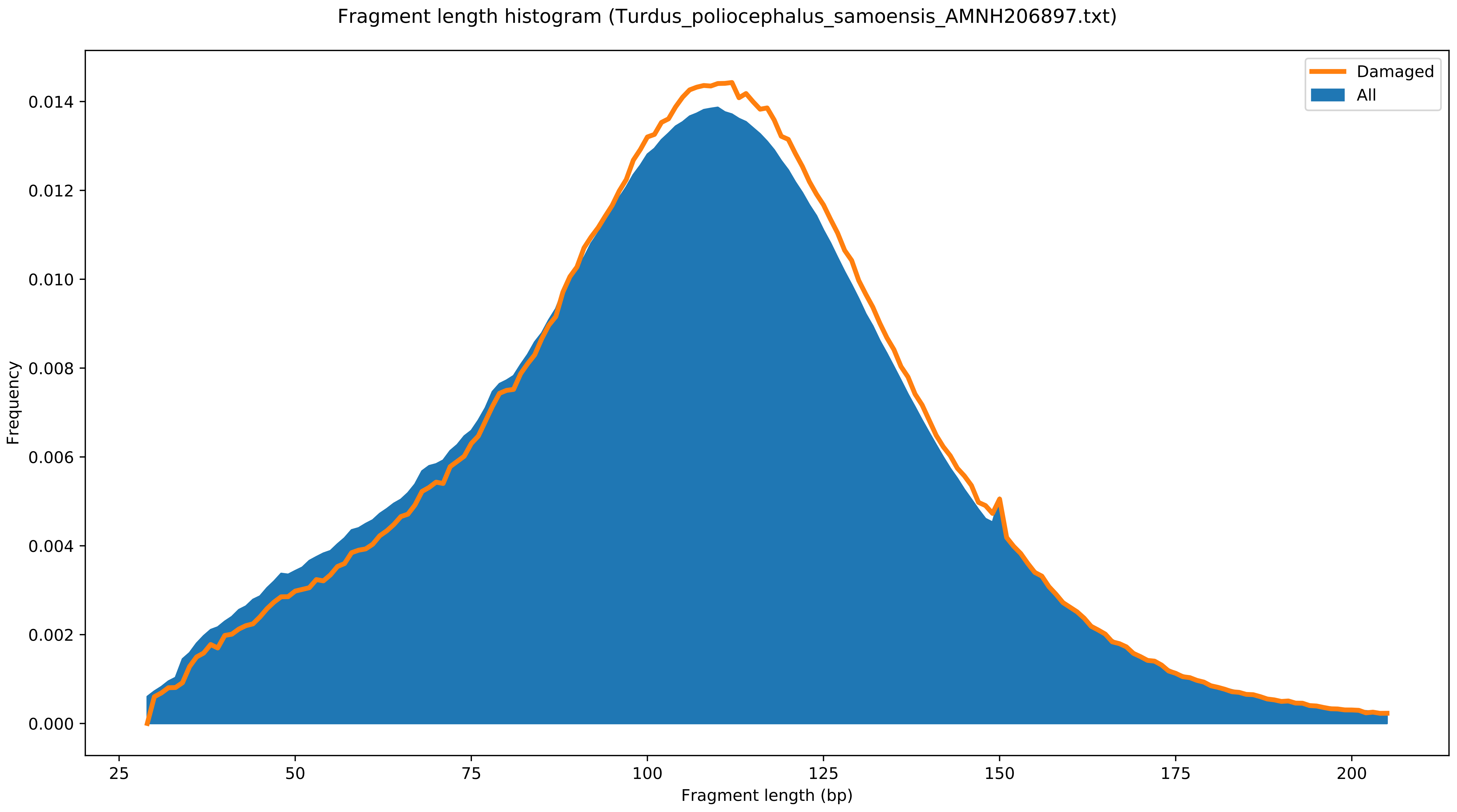
Fragment length histogram (Turdus_poliocephalus_samoensis_AMNH206897.txt)
Damaged
All
Frequency
0.000
0.002
0.004
0.006
0.008
0.010
0.012
0.014
25
50
75
100
125
150
175
200
Fragment length (bp)

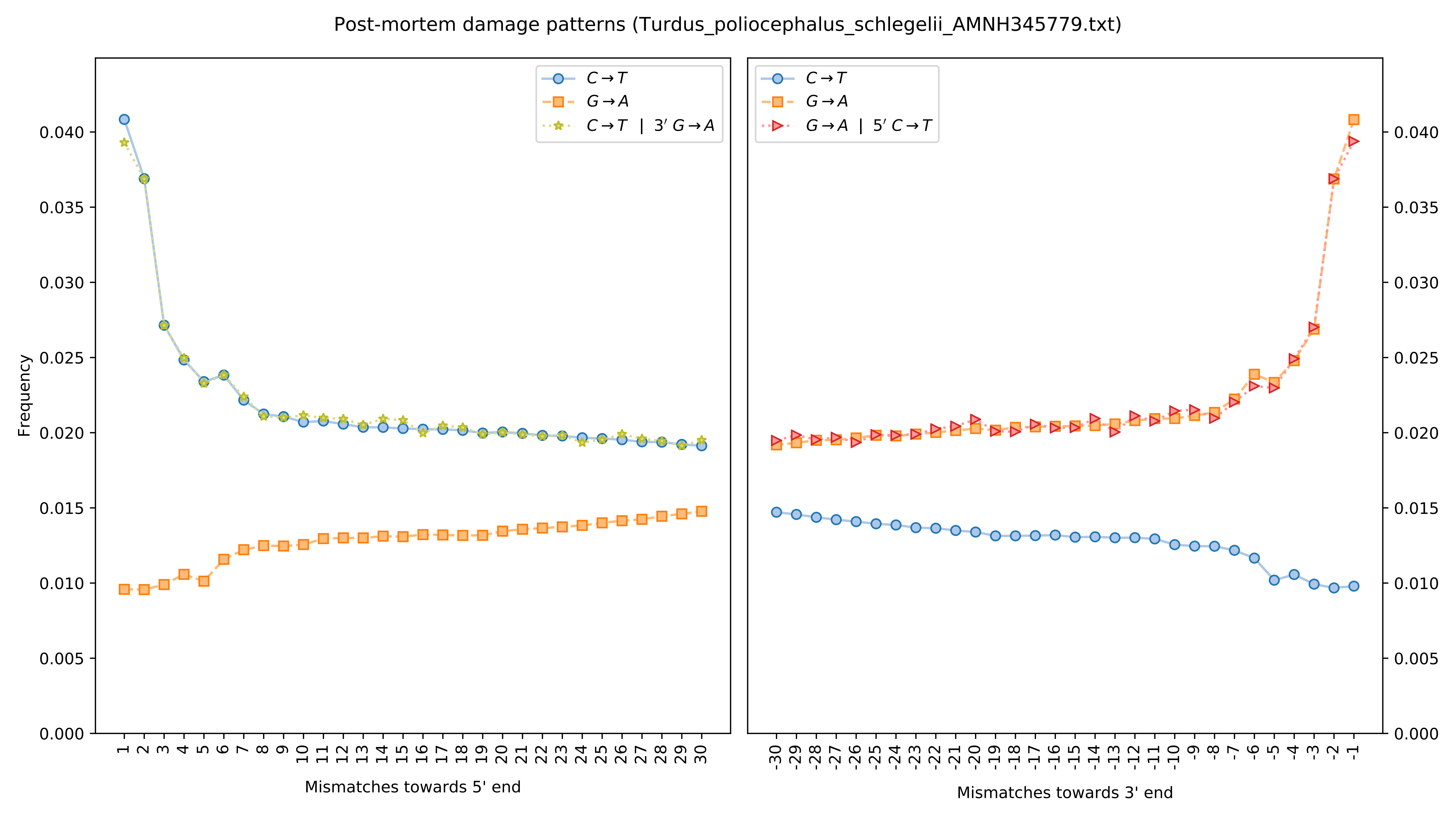

Post-mortem damage patterns (Turdus_poliocephalus_schlegelii_AMNH345779.txt)
$C \to T$
$G \to A$
$C \to T \ | \ 3' \ G \to A$
$G \to A \ | \ 5' \ C \to T$
Frequency
Mismatches towards 5' end
Mismatches towards 3' end

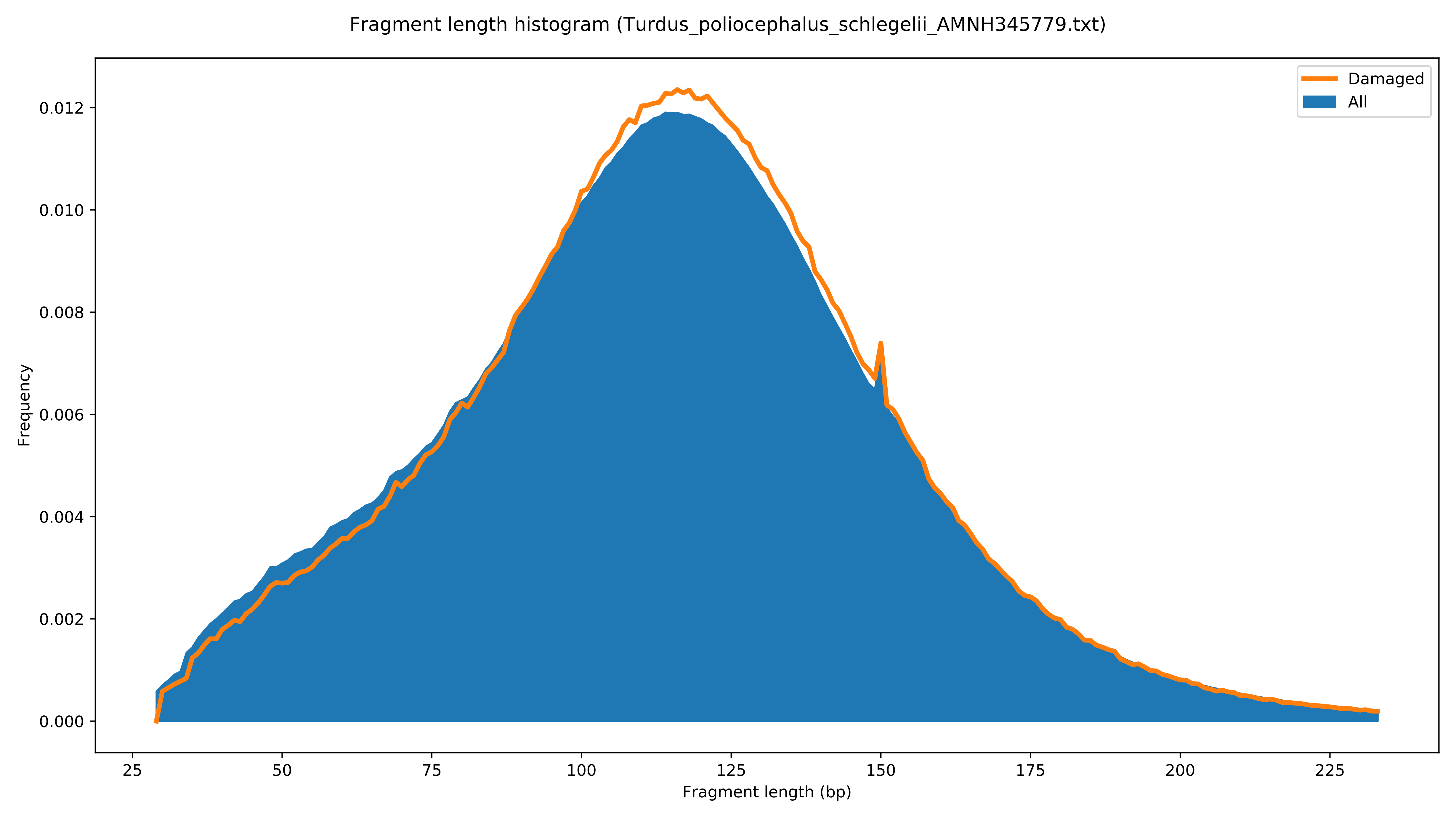

Fragment length histogram (Turdus_poliocephalus_schlegelii_AMNH345779.txt)
Damaged
All
Frequency
0.000
0.002
0.004
0.006
0.008
0.010
0.012
25
50
75
100
125
150
175
200
225
Fragment length (bp)

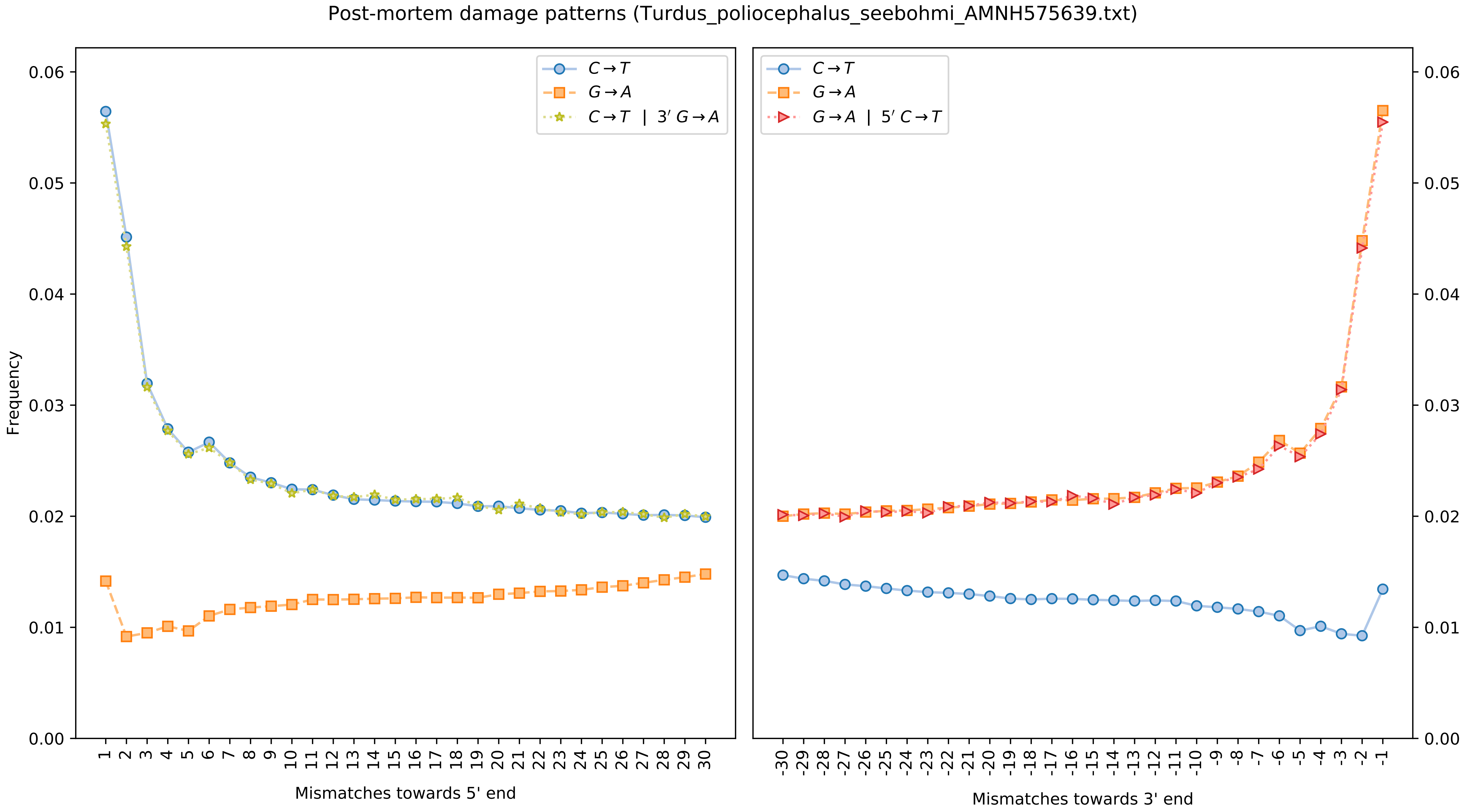

Post-mortem damage patterns (Turdus_poliocephalus_seebohmi_AMNH575639.txt)
Frequency
C→T
G→A
C→T | 3′ G→A
G→A | 5′ C→T
Mismatches towards 5' end
Mismatches towards 3' end

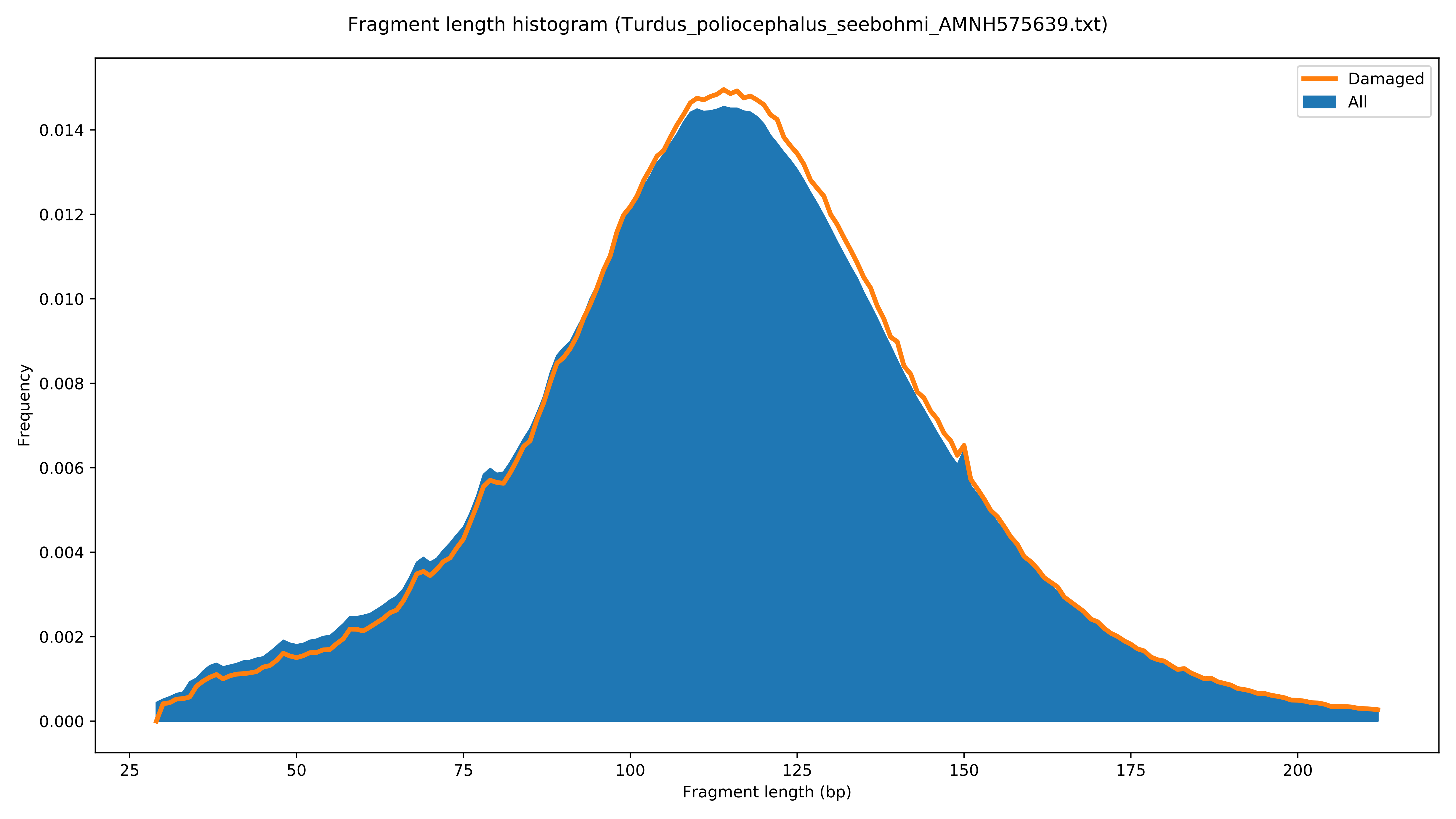

Fragment length histogram (Turdus_poliocephalus_seebohmi_AMNH575639.txt)
Damaged
All
Frequency
0.000
0.002
0.004
0.006
0.008
0.010
0.012
0.014
25
50
75
100
125
150
175
200
Fragment length (bp)

Post-mortem damage patterns (Turdus_poliocephalus_sladeni_AMNH22868.txt)

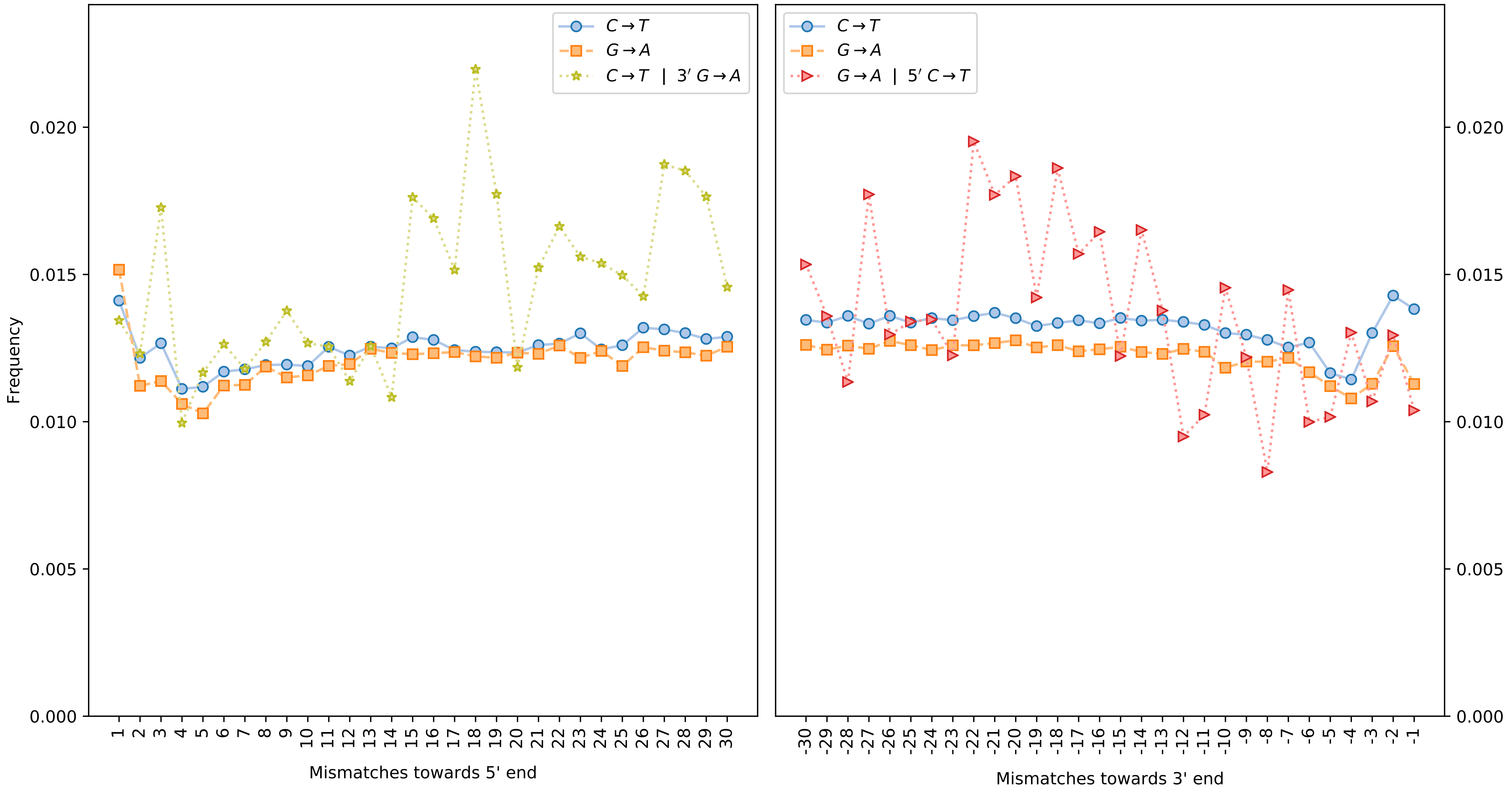

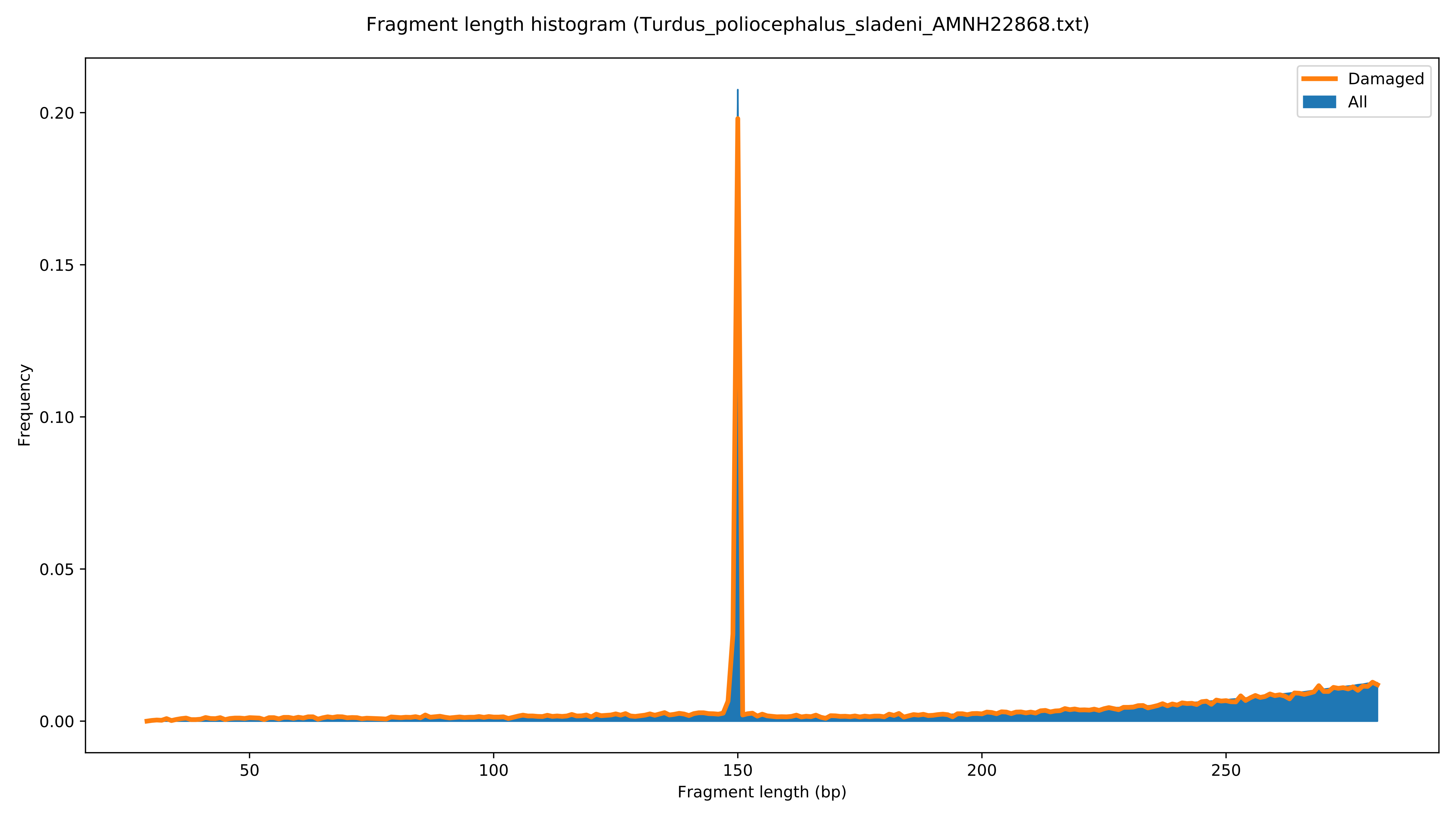

Fragment length histogram (Turdus_poliocephalus_sladeni_AMNH22868.txt)
Damaged
All
Frequency
0.00
0.05
0.10
0.15
0.20
50
100
150
200
250
Fragment length (bp)

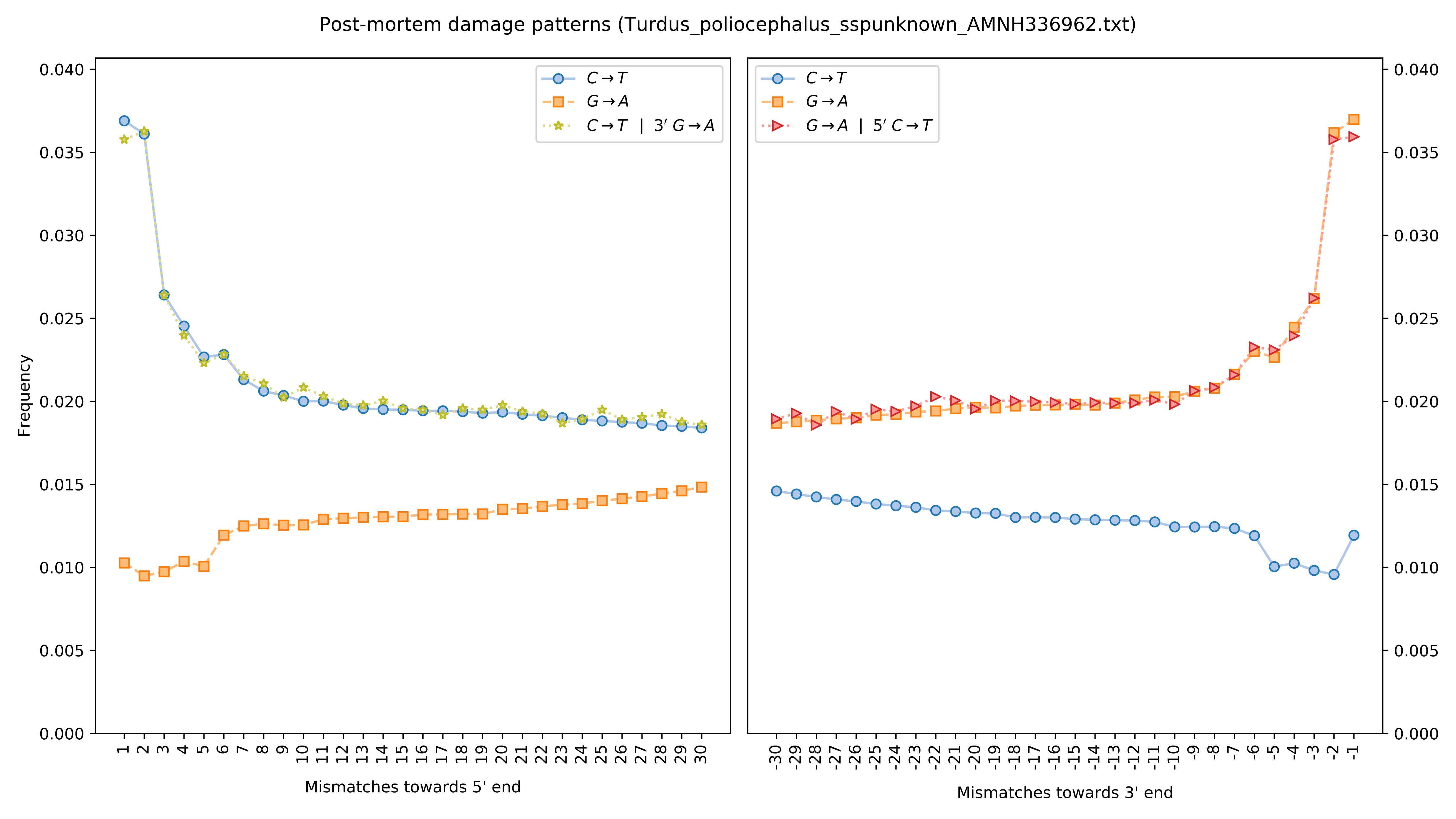
Post-mortem damage patterns (Turdus_poliocephalus_sspunknown_AMNH336962.txt)
C→T
G→A
C→T | 3′ G→A
G→A | 5′ C→T
Frequency
Mismatches towards 5' end
Mismatches towards 3' end

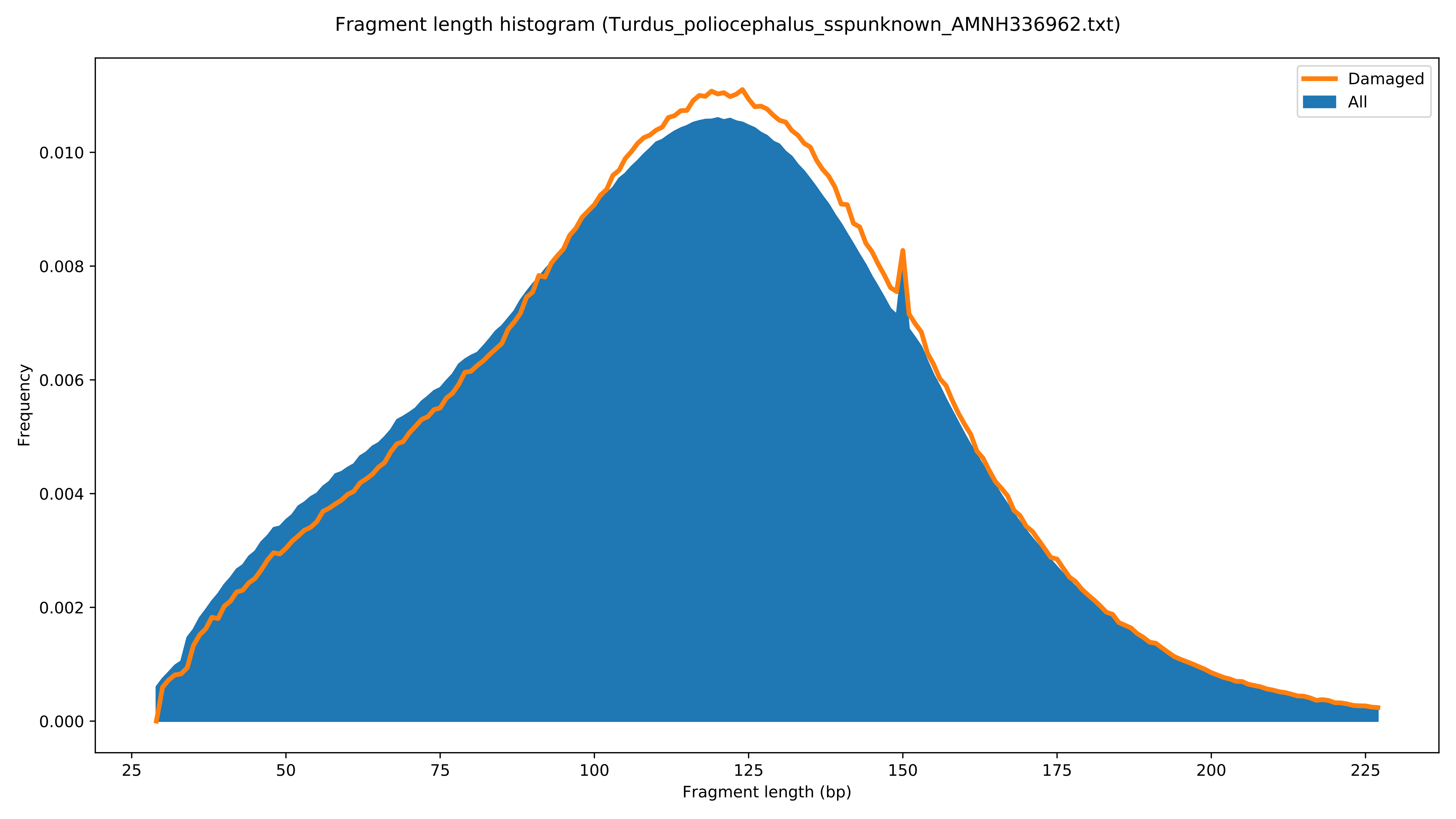

Fragment length histogram (Turdus_poliocephalus_sspunknown_AMNH336962.txt)
Damaged
All
Frequency
0.000
0.002
0.004
0.006
0.008
0.010
25
50
75
100
125
150
175
200
225
Fragment length (bp)

Post-mortem damage patterns (Turdus_poliocephalus_sspunknown_CMCB36870.txt)

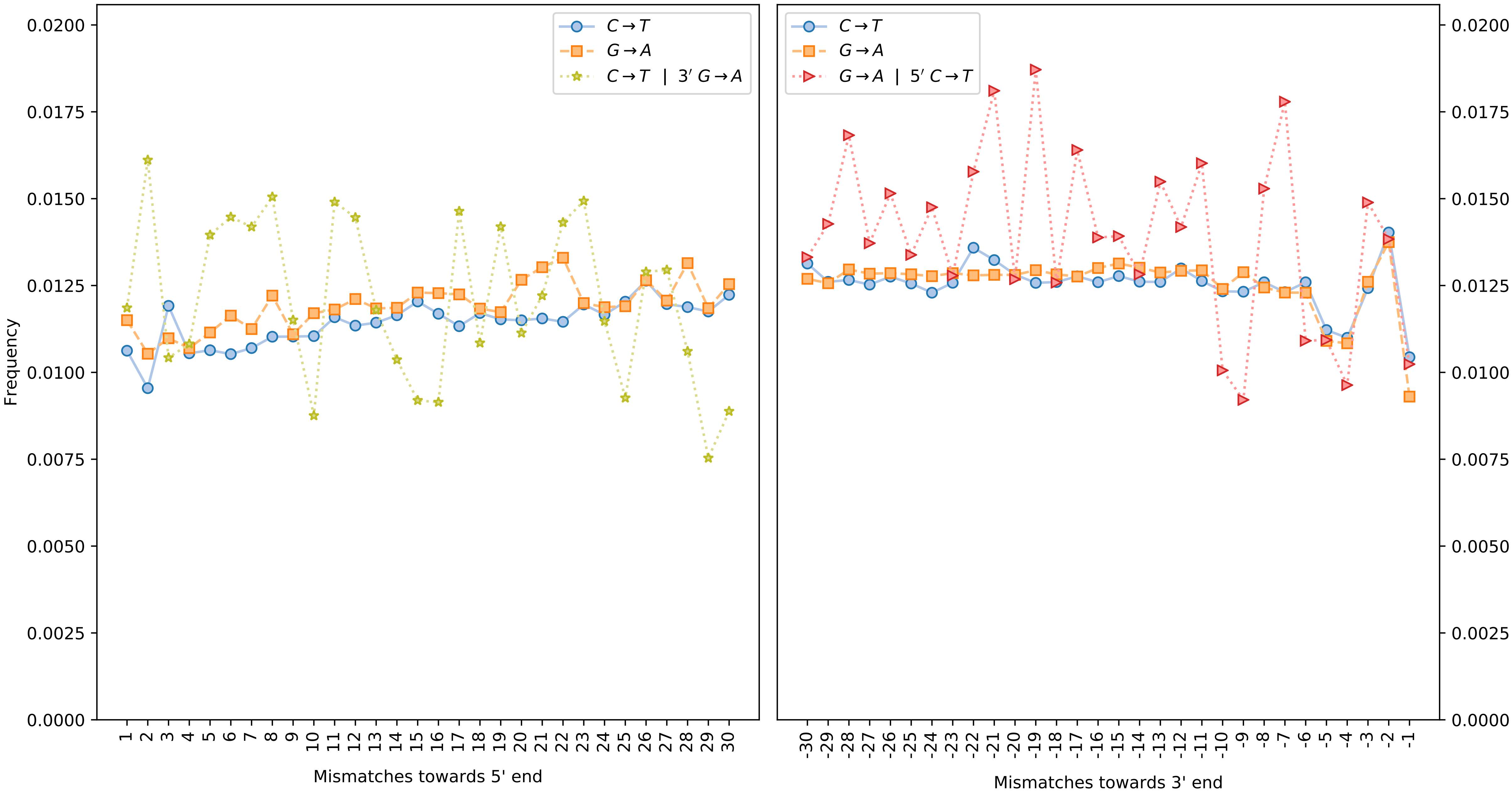

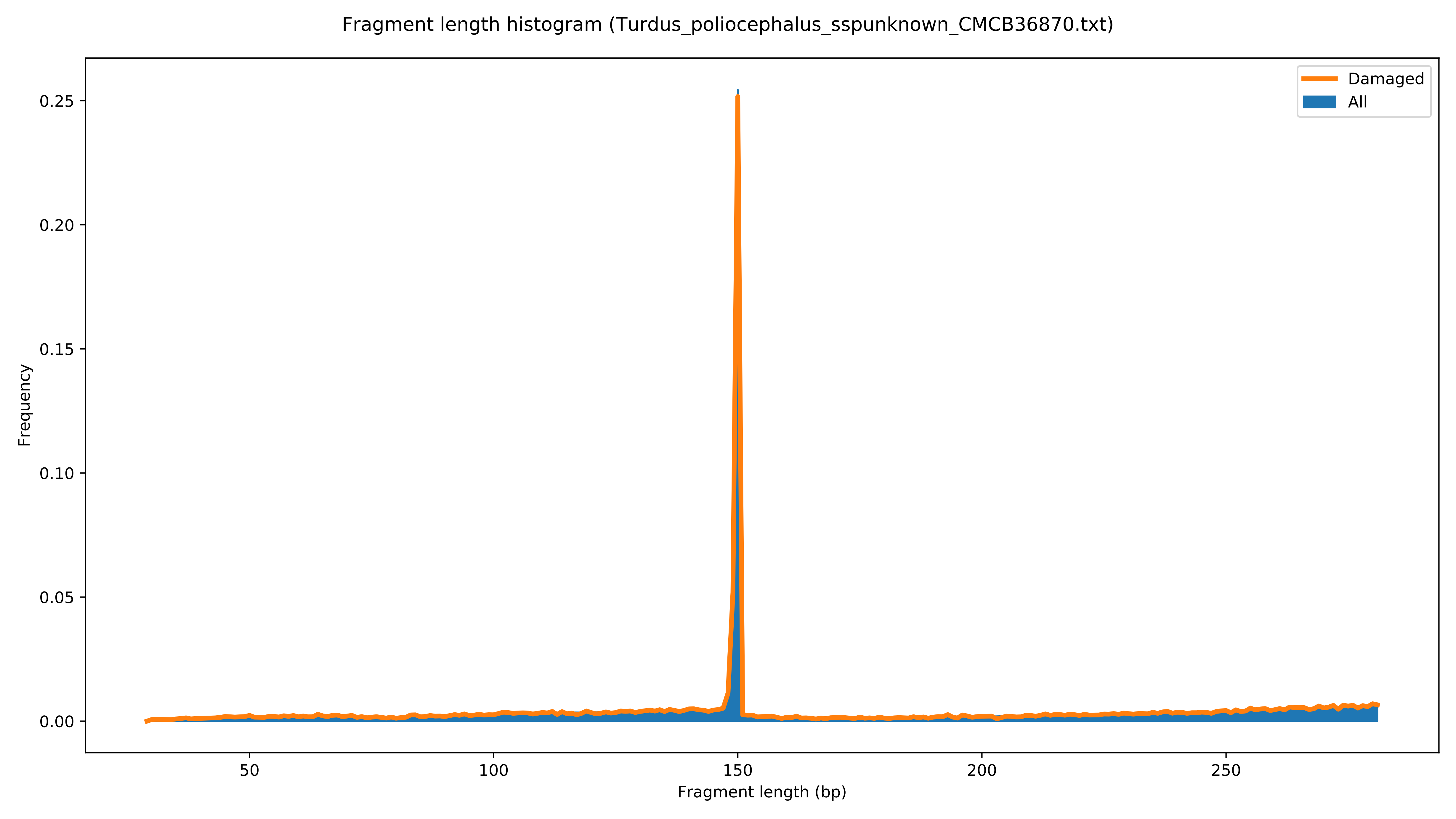

Fragment length histogram (Turdus_poliocephalus_sspunknown_CMCB36870.txt)
Damaged
All
Frequency
0.00
0.05
0.10
0.15
0.20
0.25
50
100
150
200
250
Fragment length (bp)

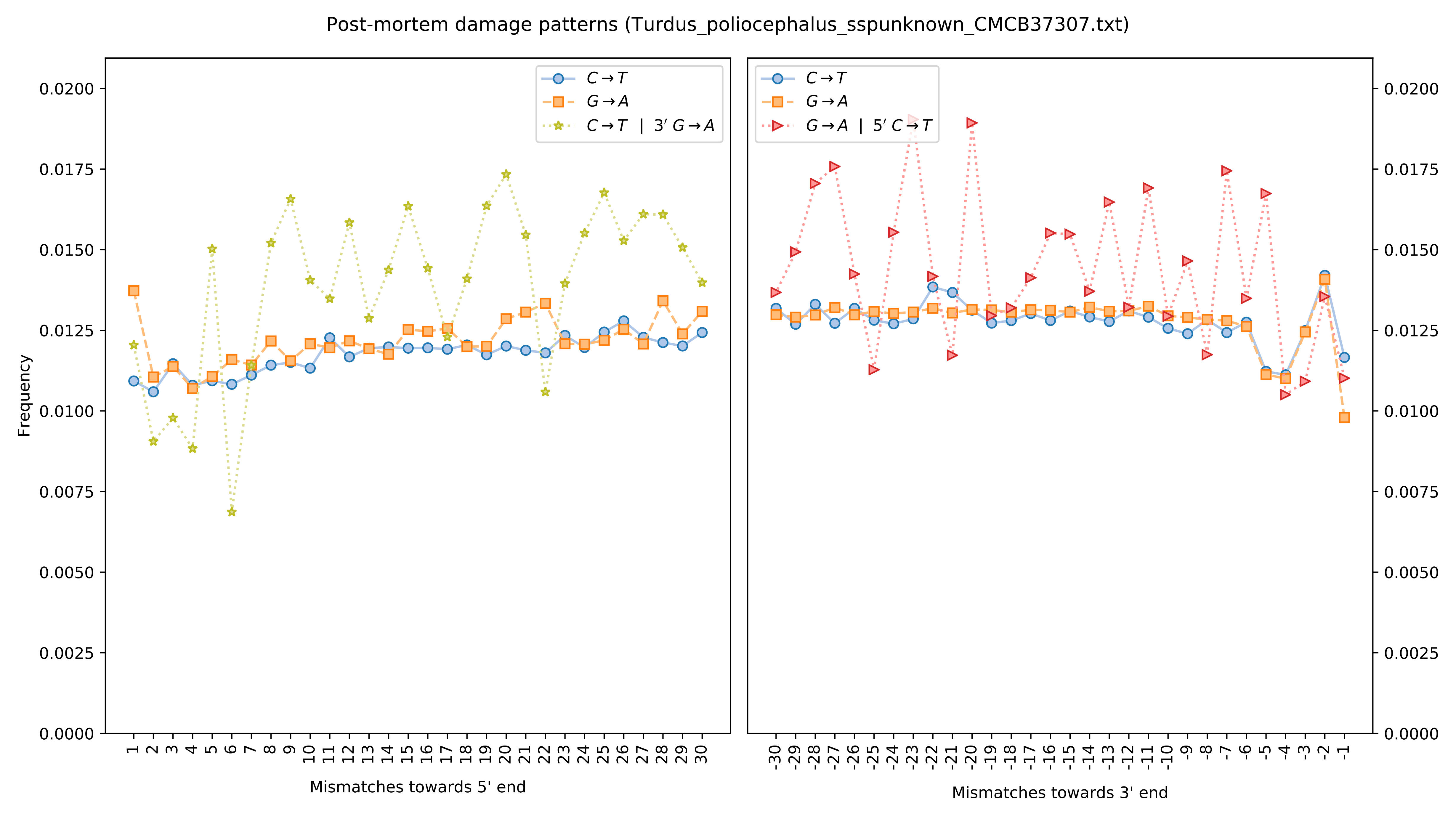

Post-mortem damage patterns (Turdus_poliocephalus_sspunknown_CMCB37307.txt)
Frequency
Mismatches towards 5' end
Mismatches towards 3' end
C→T
G→A
C→T | 3′ G→A
G→A | 5′ C→T

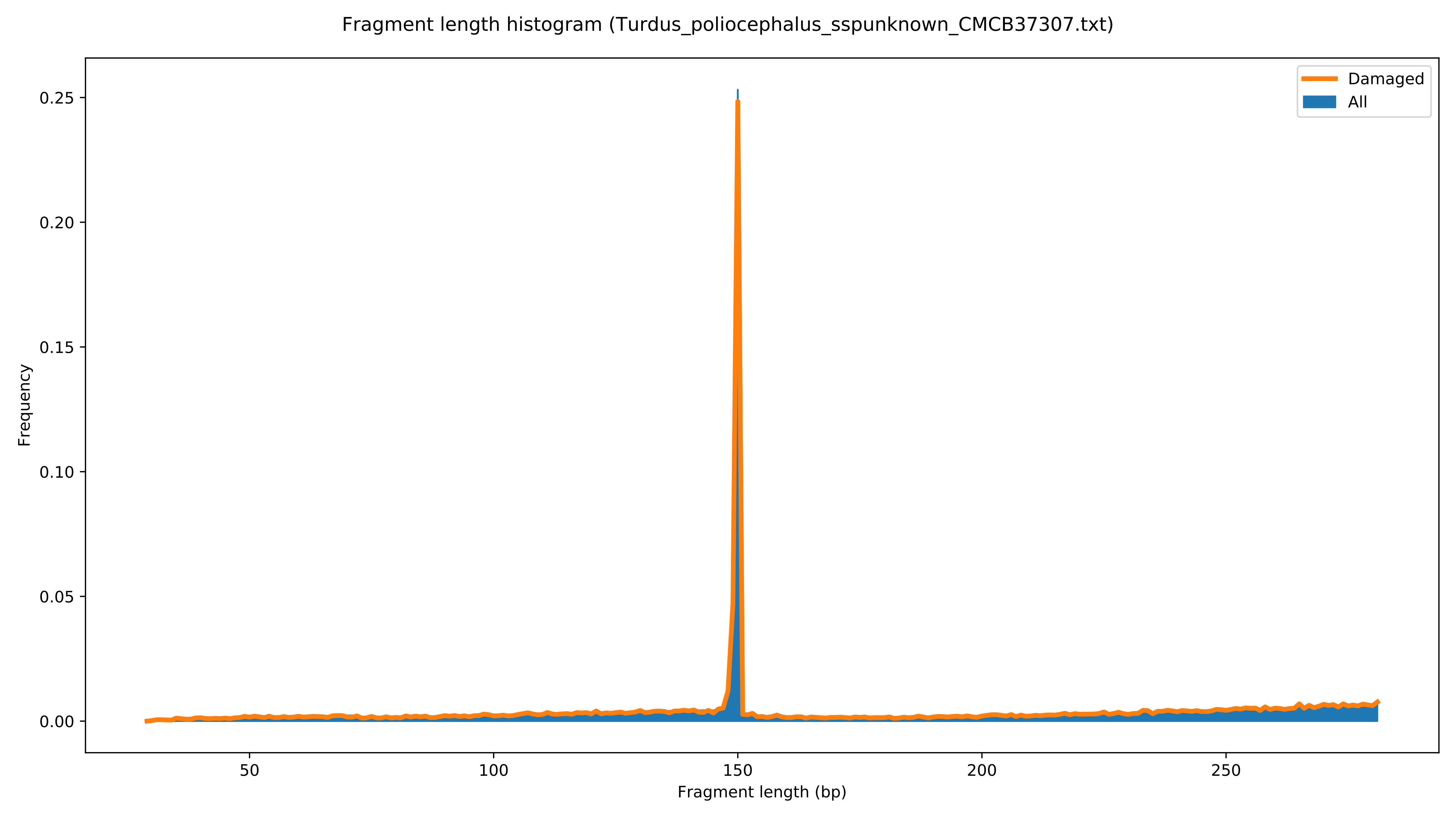

Fragment length histogram (Turdus_poliocephalus_sspunknown_CMCB37307.txt)
Damaged
All
Frequency
0.00
0.05
0.10
0.15
0.20
0.25
50
100
150
200
250
Fragment length (bp)

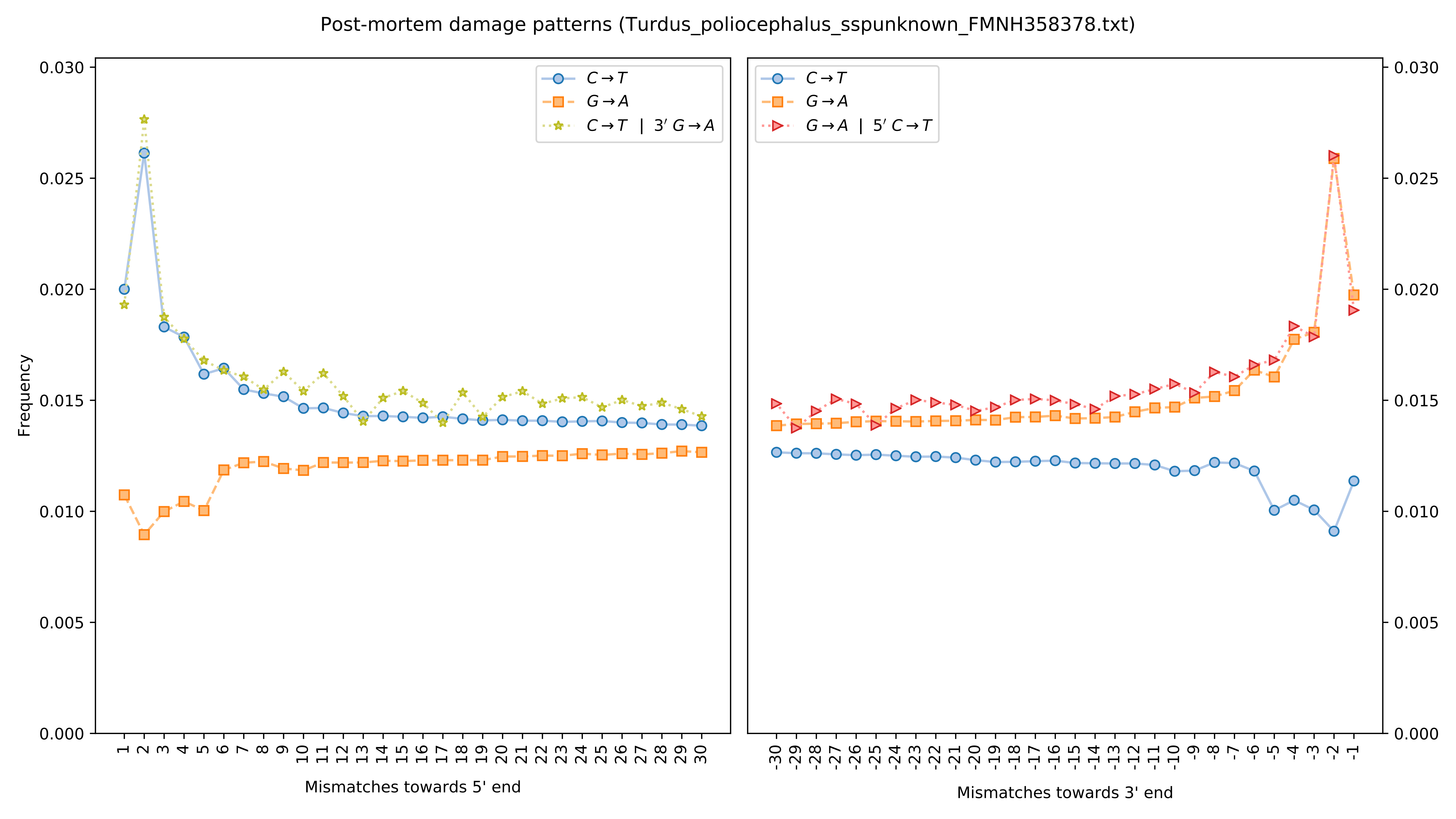

Post-mortem damage patterns (Turdus_poliocephalus_sspunknown_FMNH358378.txt)
Frequency
Mismatches towards 5' end
Mismatches towards 3' end
C→T
G→A
C→T | 3′ G→A
G→A | 5′ C→T
0.000
0.005
0.010
0.015
0.020
0.025
0.030

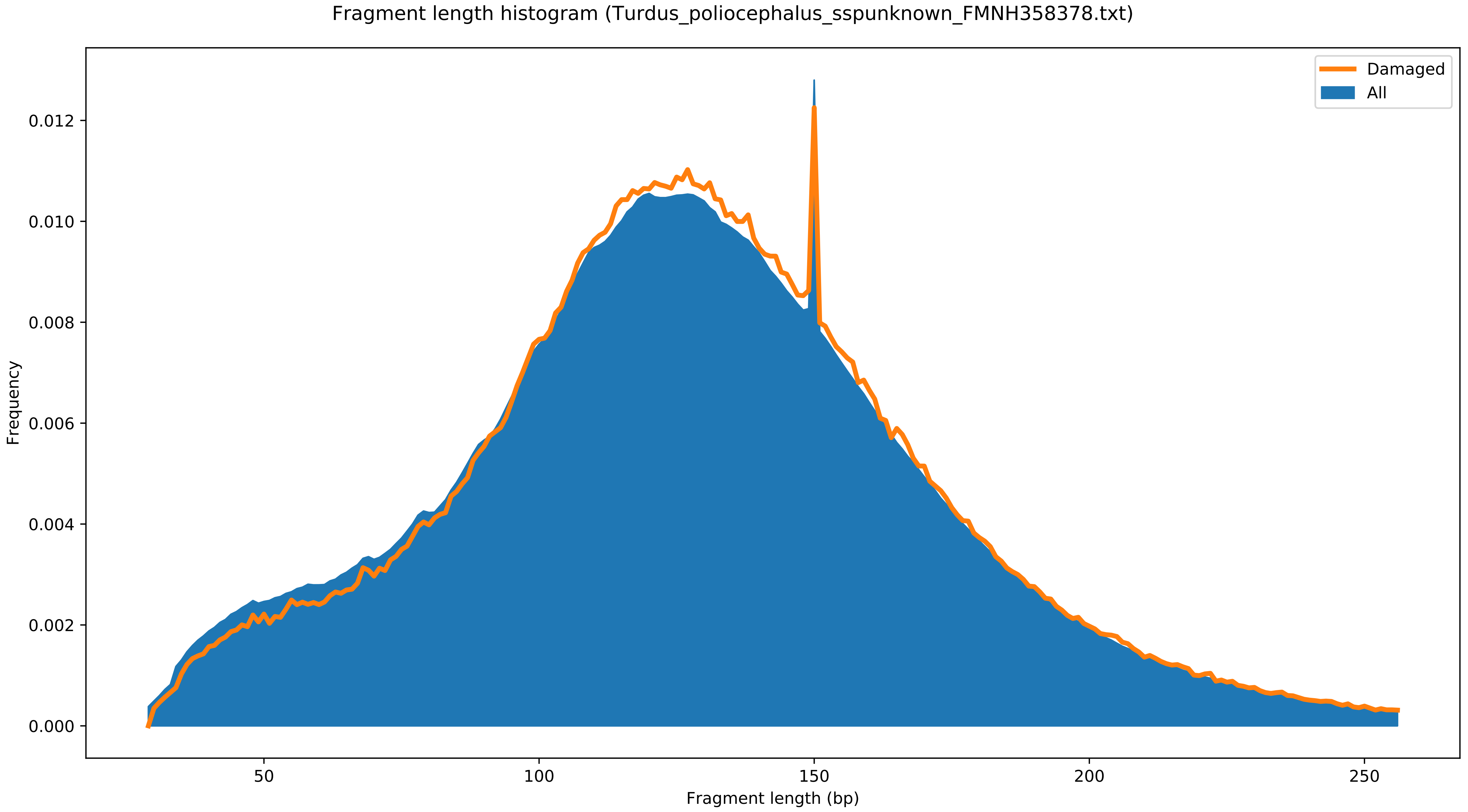

Fragment length histogram (Turdus_poliocephalus_sspunknown_FMNH358378.txt)
Damaged
All
Frequency
0.000
0.002
0.004
0.006
0.008
0.010
0.012
50
100
150
200
250
Fragment length (bp)

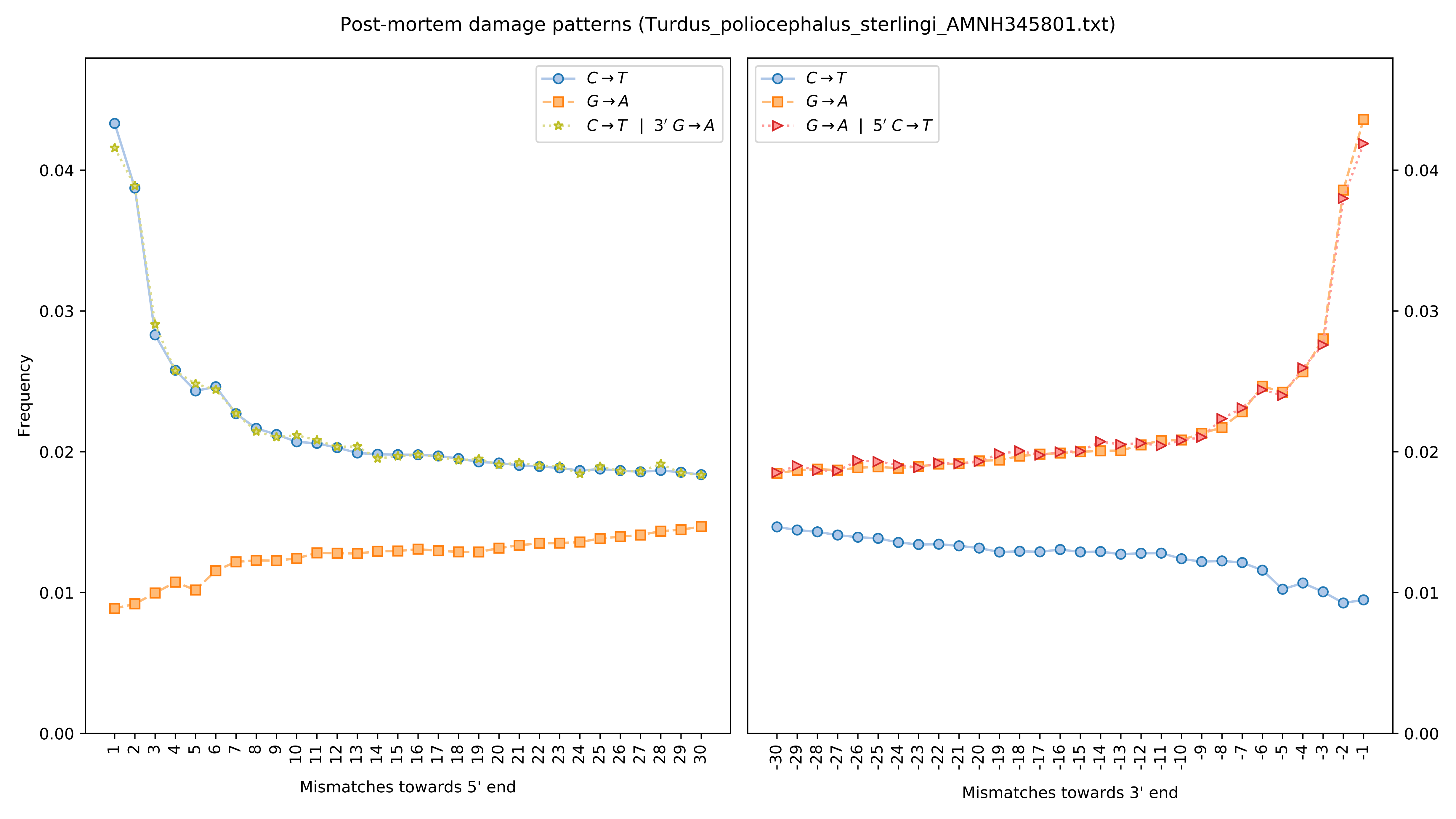

Post-mortem damage patterns (Turdus_poliocephalus_sterlingi_AMNH345801.txt)
Frequency
Mismatches towards 5' end
Mismatches towards 3' end
C→T
G→A
C→T | 3′ G→A
G→A | 5′ C→T

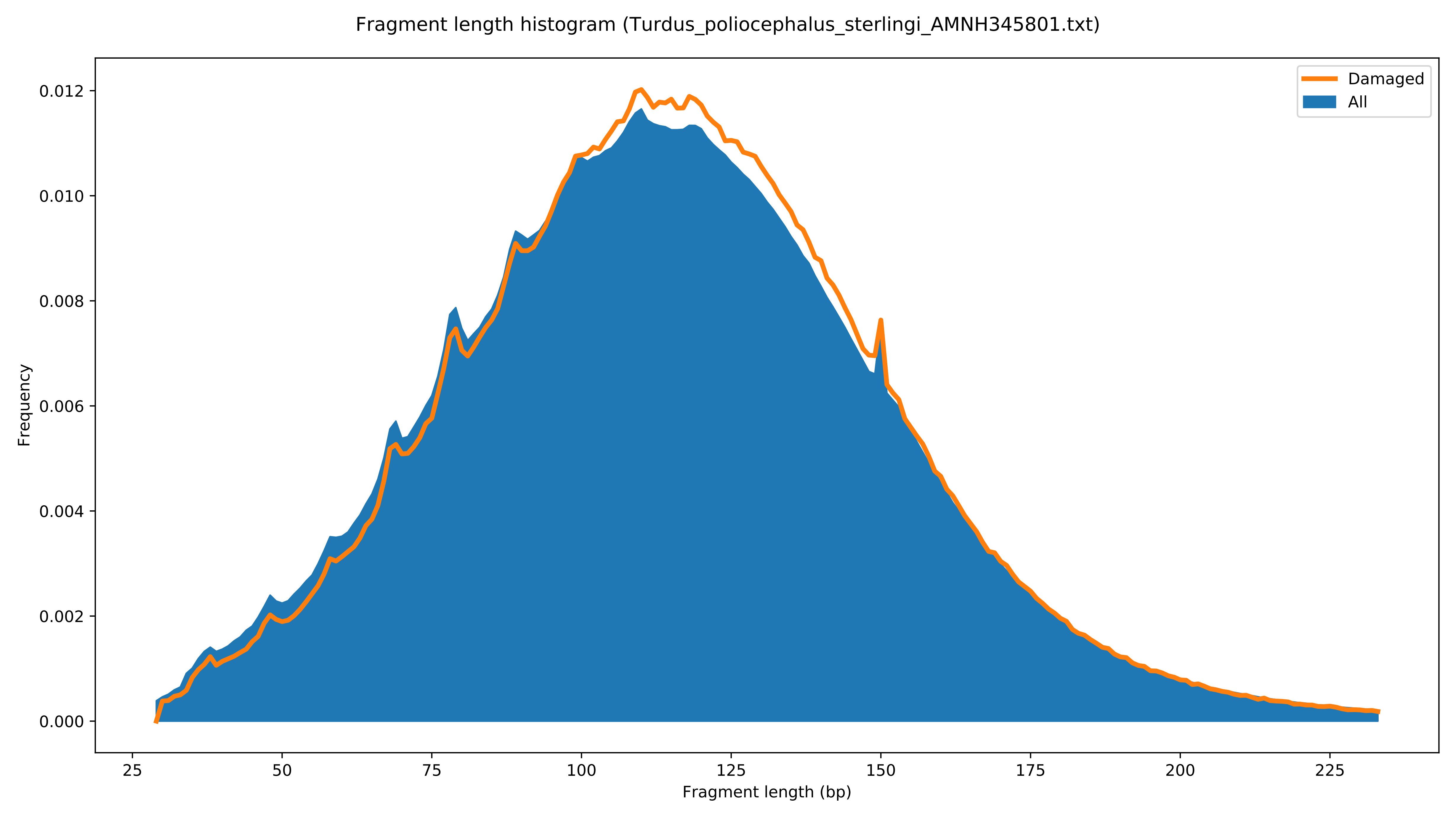

Fragment length histogram (Turdus_poliocephalus_sterlingi_AMNH345801.txt)
Damaged
All
Frequency
Fragment length (bp)
0.000
0.002
0.004
0.006
0.008
0.010
0.012
25
50
75
100
125
150
175
200
225

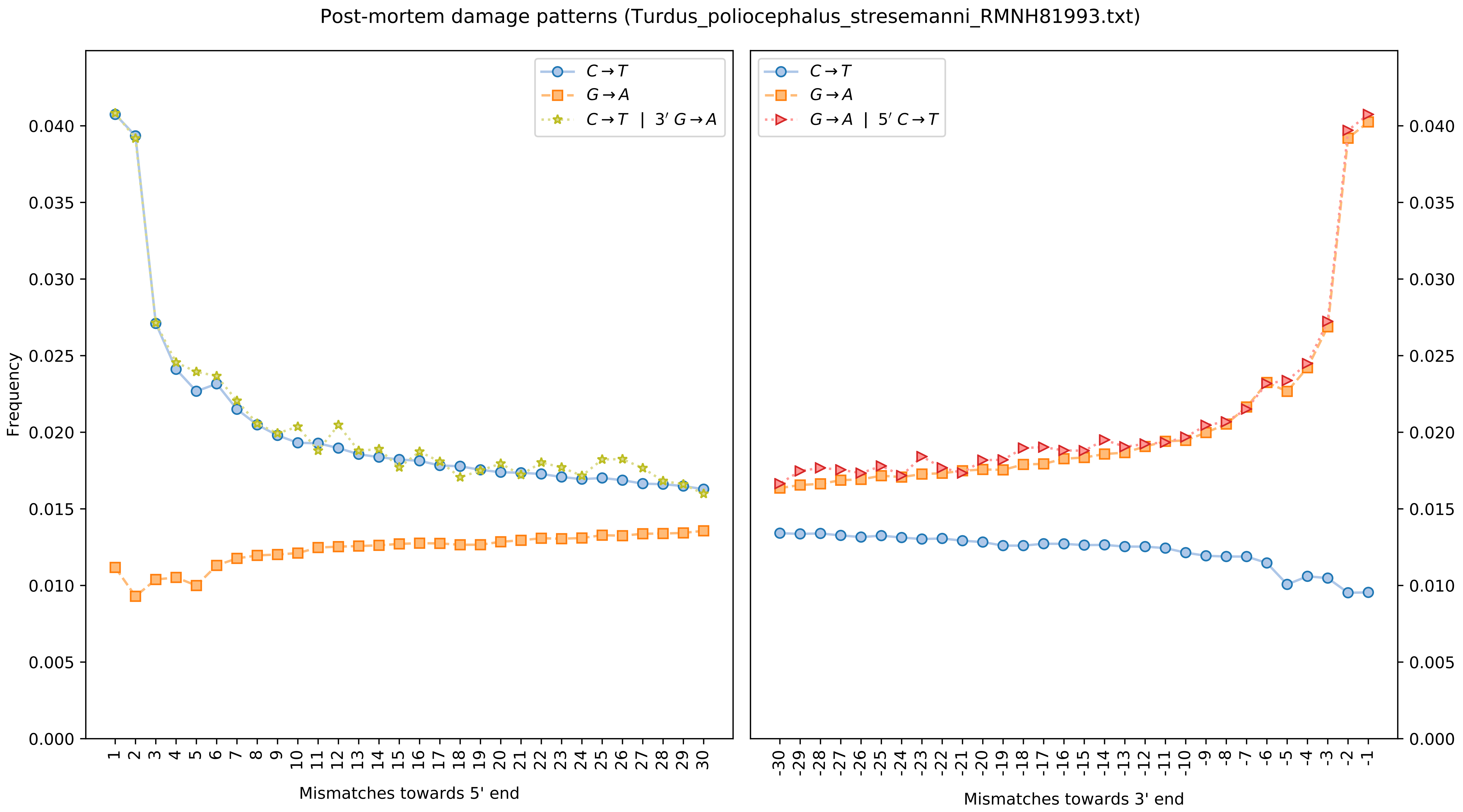

Post-mortem damage patterns (Turdus_poliocephalus_stresemanni_RMNH81993.txt)
Frequency
C→T
G→A
C→T | 3′ G→A
G→A | 5′ C→T
Mismatches towards 5' end
Mismatches towards 3' end

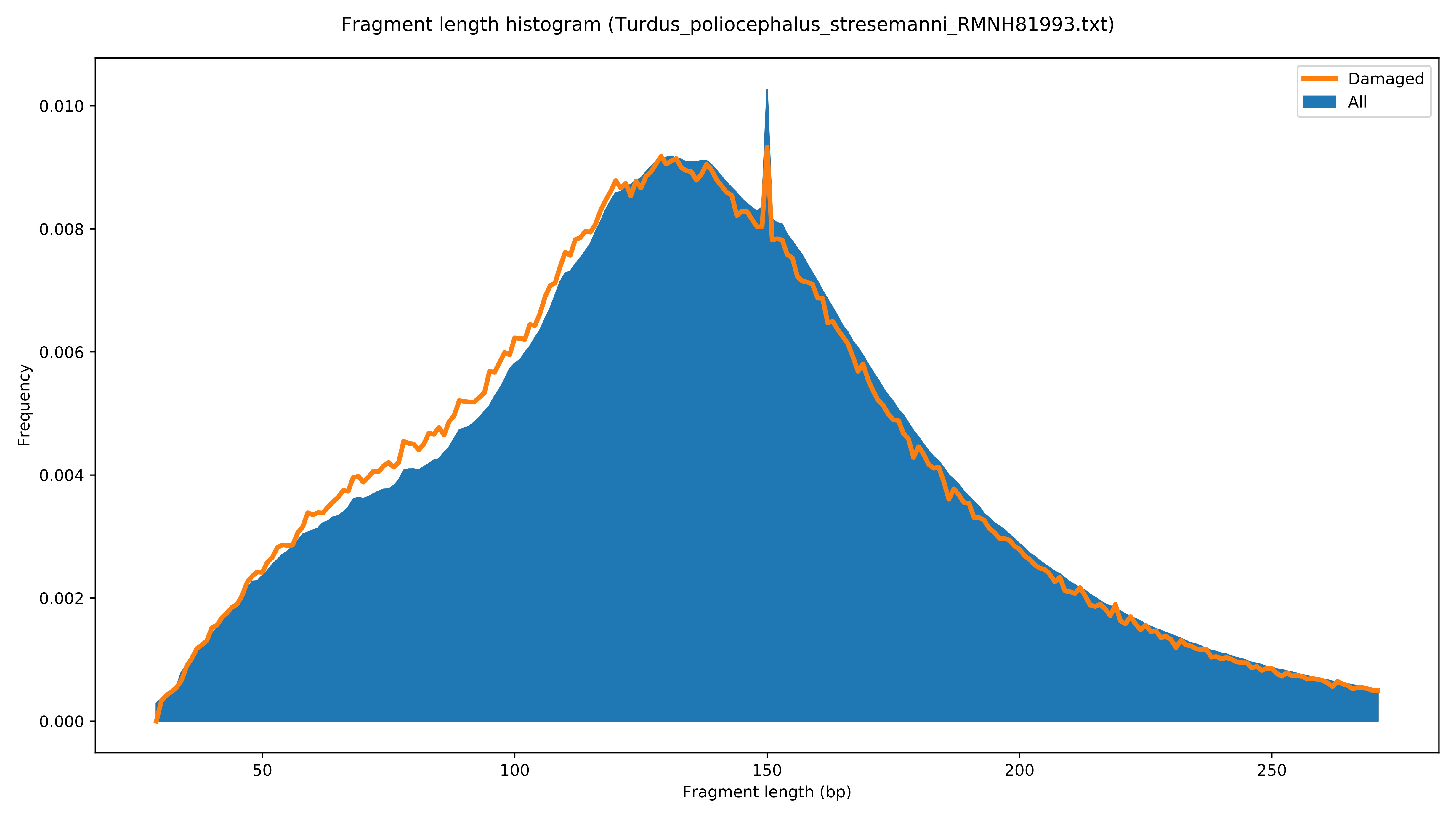

Fragment length histogram (Turdus_poliocephalus_stresemanni_RMNH81993.txt)
Damaged
All
Frequency
0.000
0.002
0.004
0.006
0.008
0.010
50
100
150
200
250
Fragment length (bp)

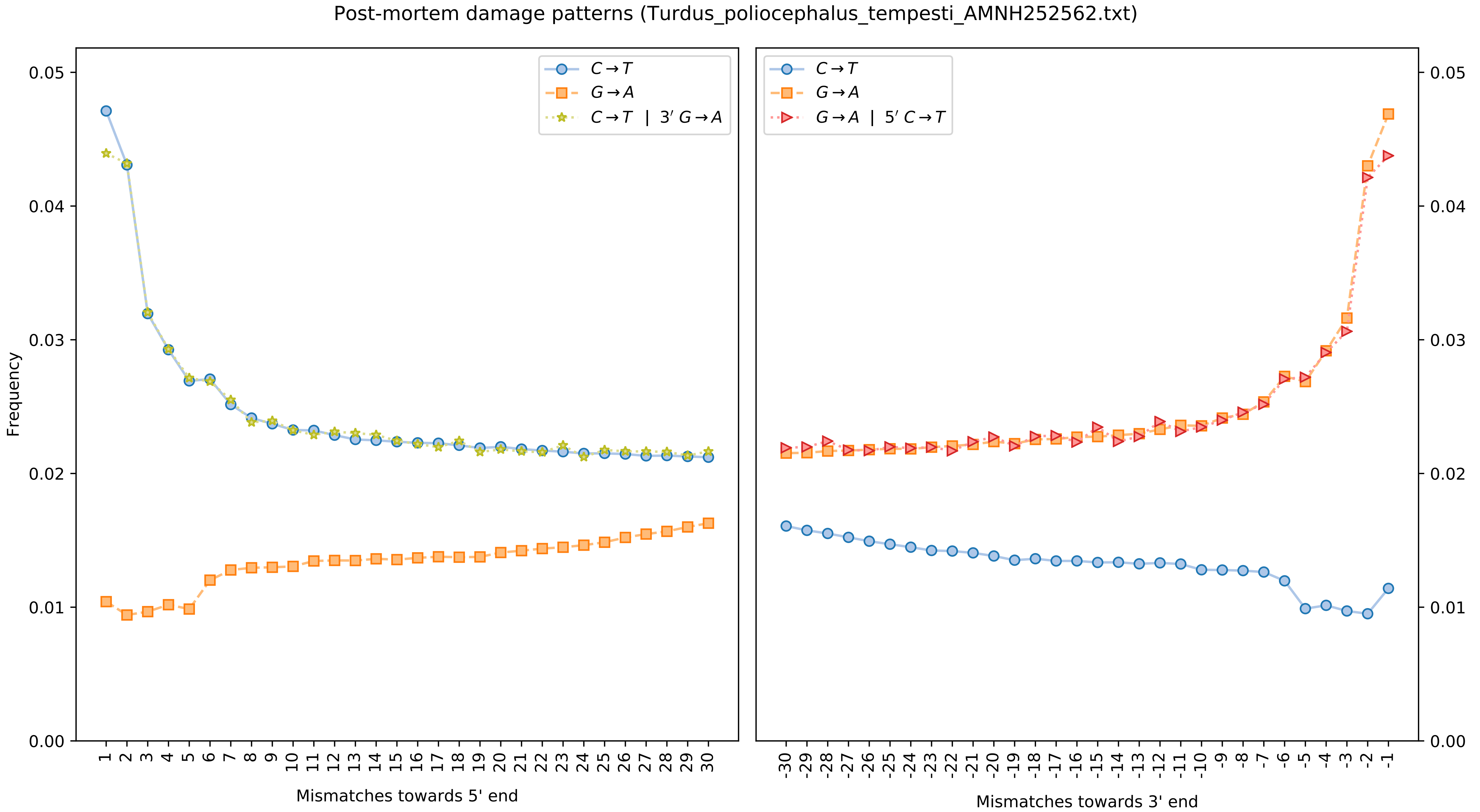

Post-mortem damage patterns (Turdus_poliocephalus_tempesti_AMNH252562.txt)
Frequency
0.00
0.01
0.02
0.03
0.04
0.05
$C \rightarrow T$
$G \rightarrow A$
$C \rightarrow T \mid 3' G \rightarrow A$
$G \rightarrow A \mid 5' C \rightarrow T$
1 2 3 4 5 6 7 8 9 10 11 12 13 14 15 16 17 18 19 20 21 22 23 24 25 26 27 28 29 30
Mismatches towards 5' end
-30 -29 -28 -27 -26 -25 -24 -23 -22 -21 -20 -19 -18 -17 -16 -15 -14 -13 -12 -11 -10 -9 -8 -7 -6 -5 -4 -3 -2 -1
Mismatches towards 3' end

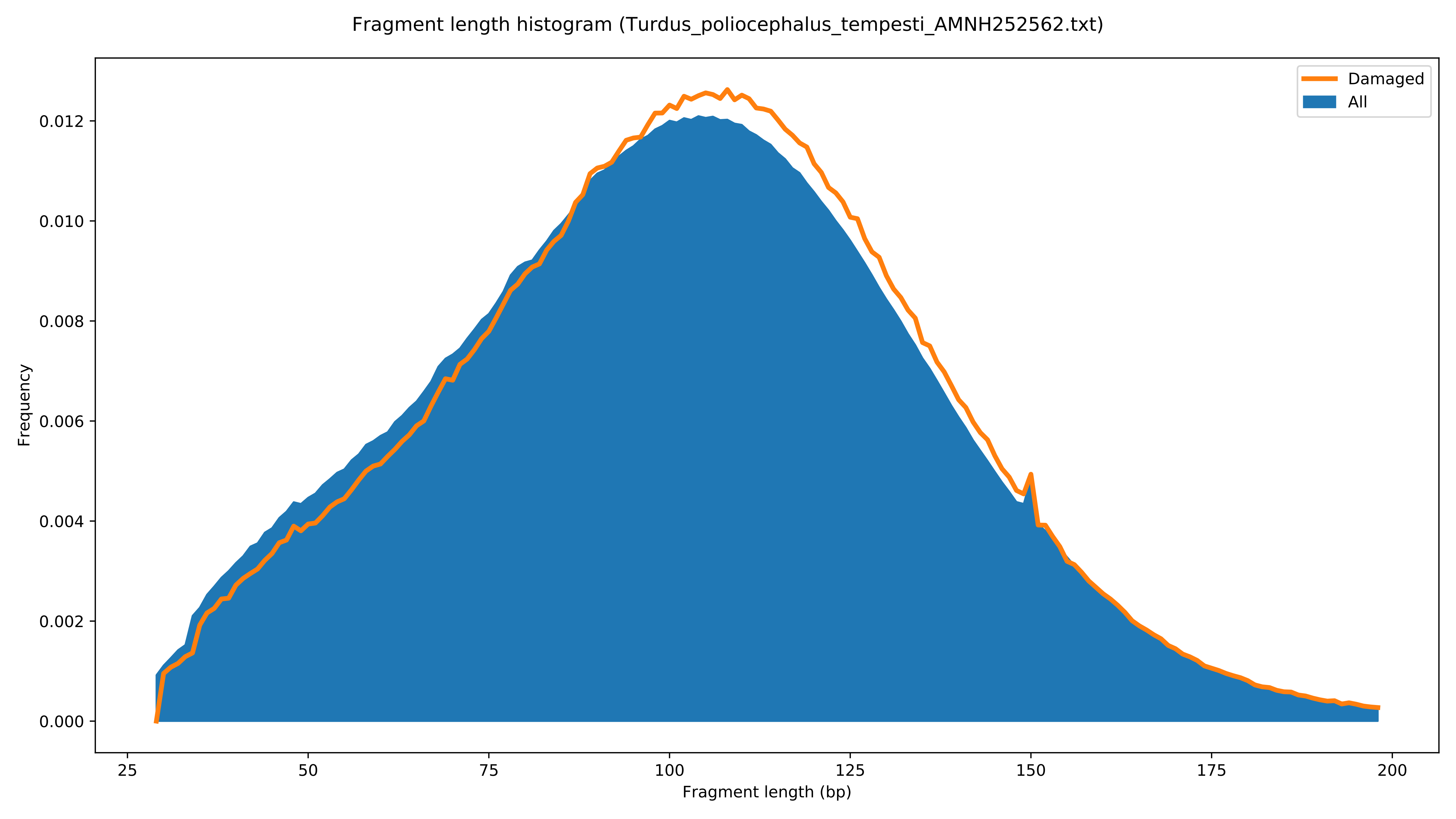

Fragment length histogram (Turdus_poliocephalus_tempesti_AMNH252562.txt)
Damaged
All
Frequency
0.000
0.002
0.004
0.006
0.008
0.010
0.012
25
50
75
100
125
150
175
200
Fragment length (bp)

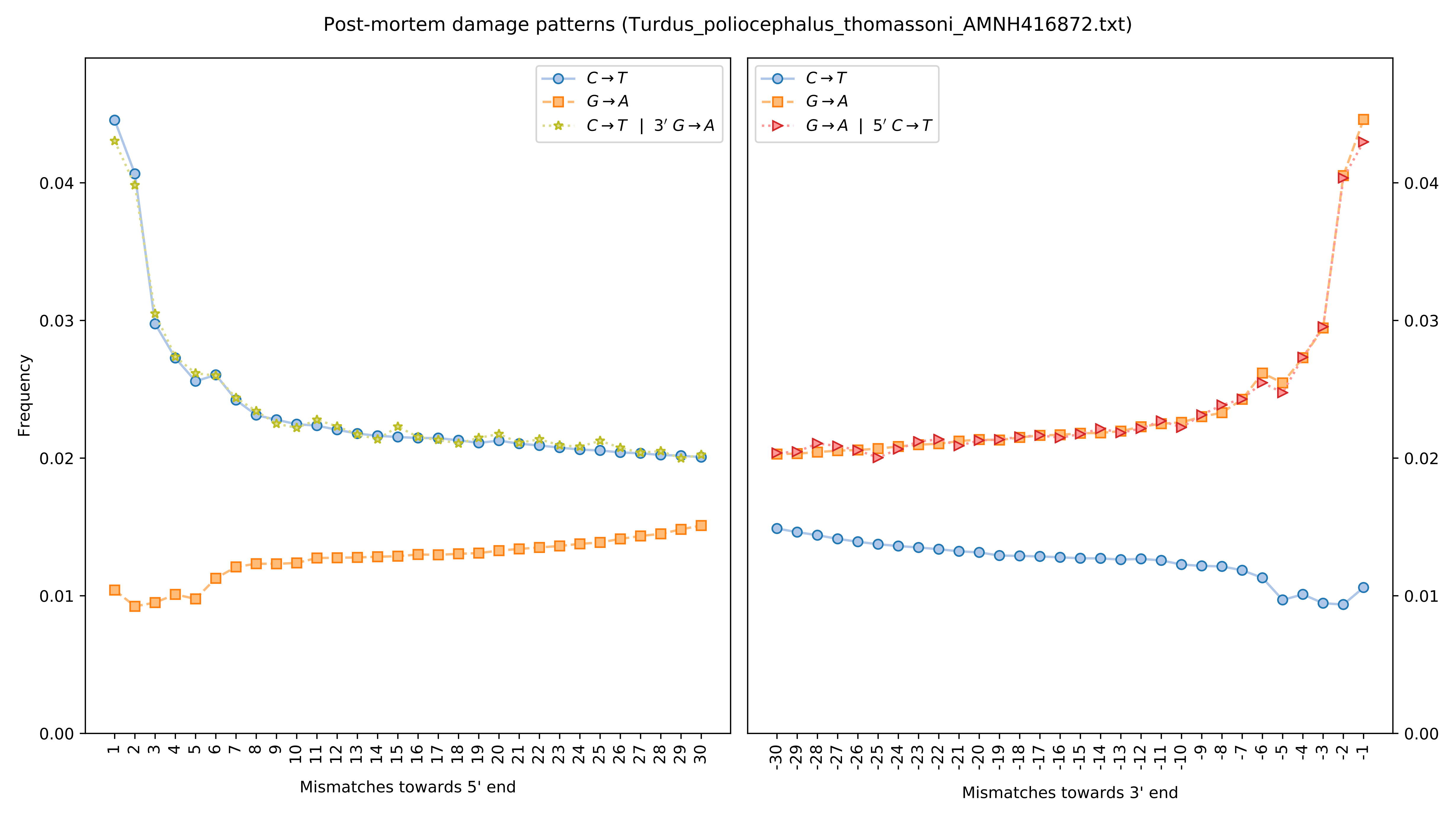

Post-mortem damage patterns (Turdus_poliocephalus_thomassoni_AMNH416872.txt)
Frequency
Mismatches towards 5' end
Mismatches towards 3' end
C→T
G→A
C→T | 3′ G→A
G→A | 5′ C→T
0.00
0.01
0.02
0.03
0.04

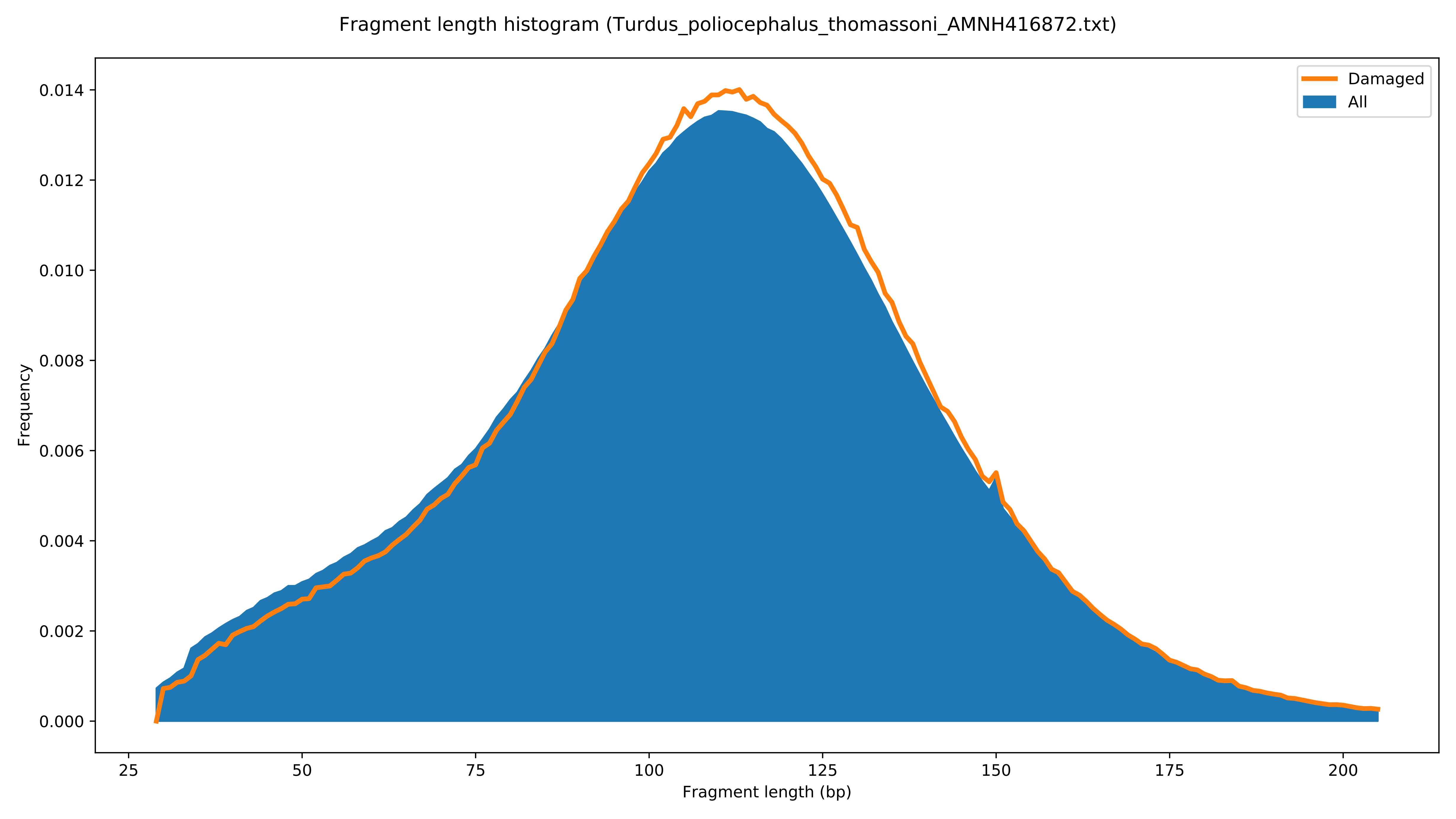

Fragment length histogram (Turdus_poliocephalus_thomassoni_AMNH416872.txt)
Frequency
0.000
0.002
0.004
0.006
0.008
0.010
0.012
0.014
25
50
75
100
125
150
175
200
Fragment length (bp)
Damaged
All

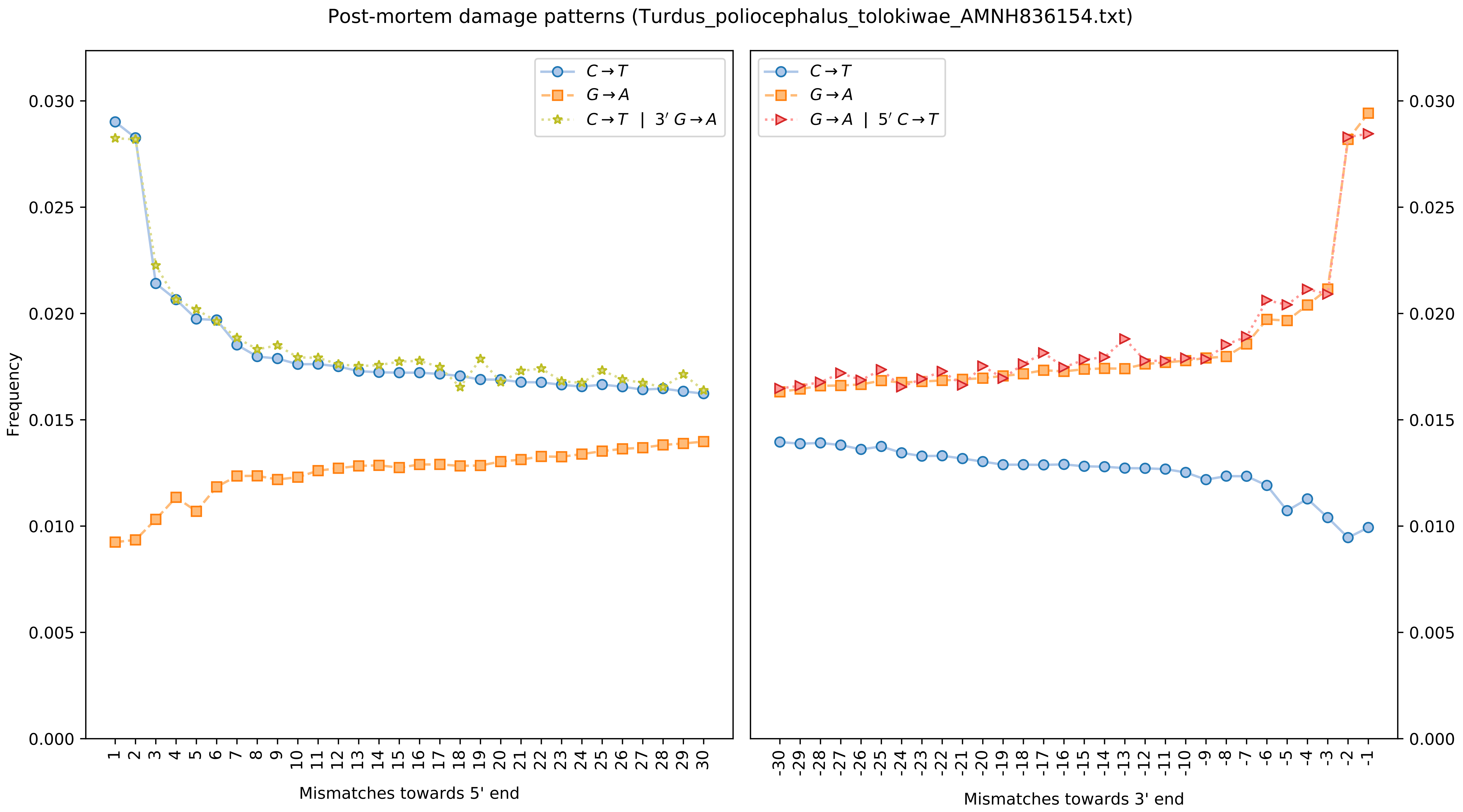

Post-mortem damage patterns (Turdus_poliocephalus_tolokiwae_AMNH836154.txt)
Frequency
Mismatches towards 5' end
Mismatches towards 3' end
C→T
G→A
C→T | 3′ G→A
G→A | 5′ C→T

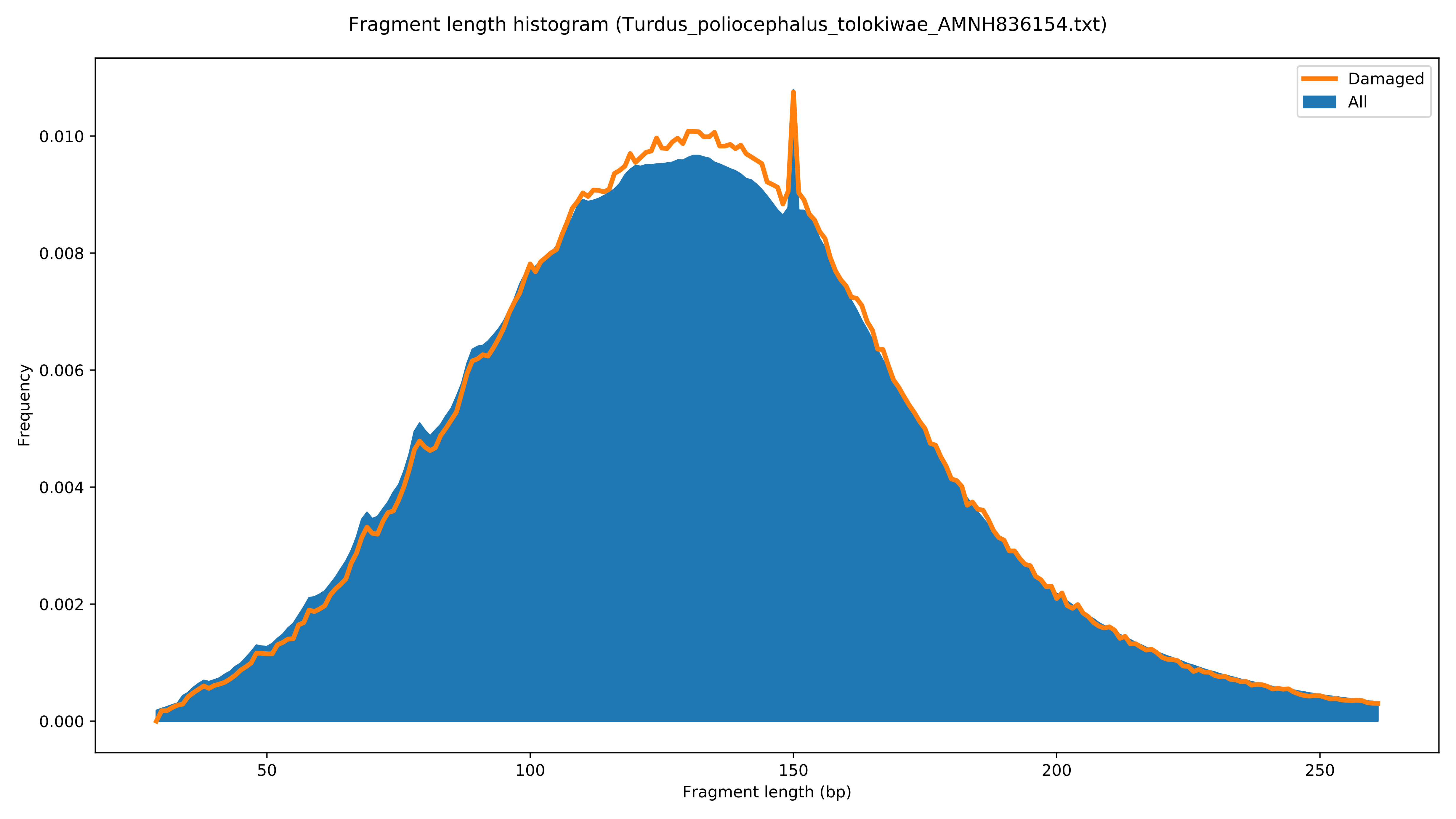
Fragment length histogram (Turdus_poliocephalus_tolokiwae_AMNH836154.txt)
Damaged
All
Frequency
0.000
0.002
0.004
0.006
0.008
0.010
50
100
150
200
250
Fragment length (bp)

Post-mortem damage patterns (Turdus_poliocephalus_vanikorensis_AMNH214400.txt)

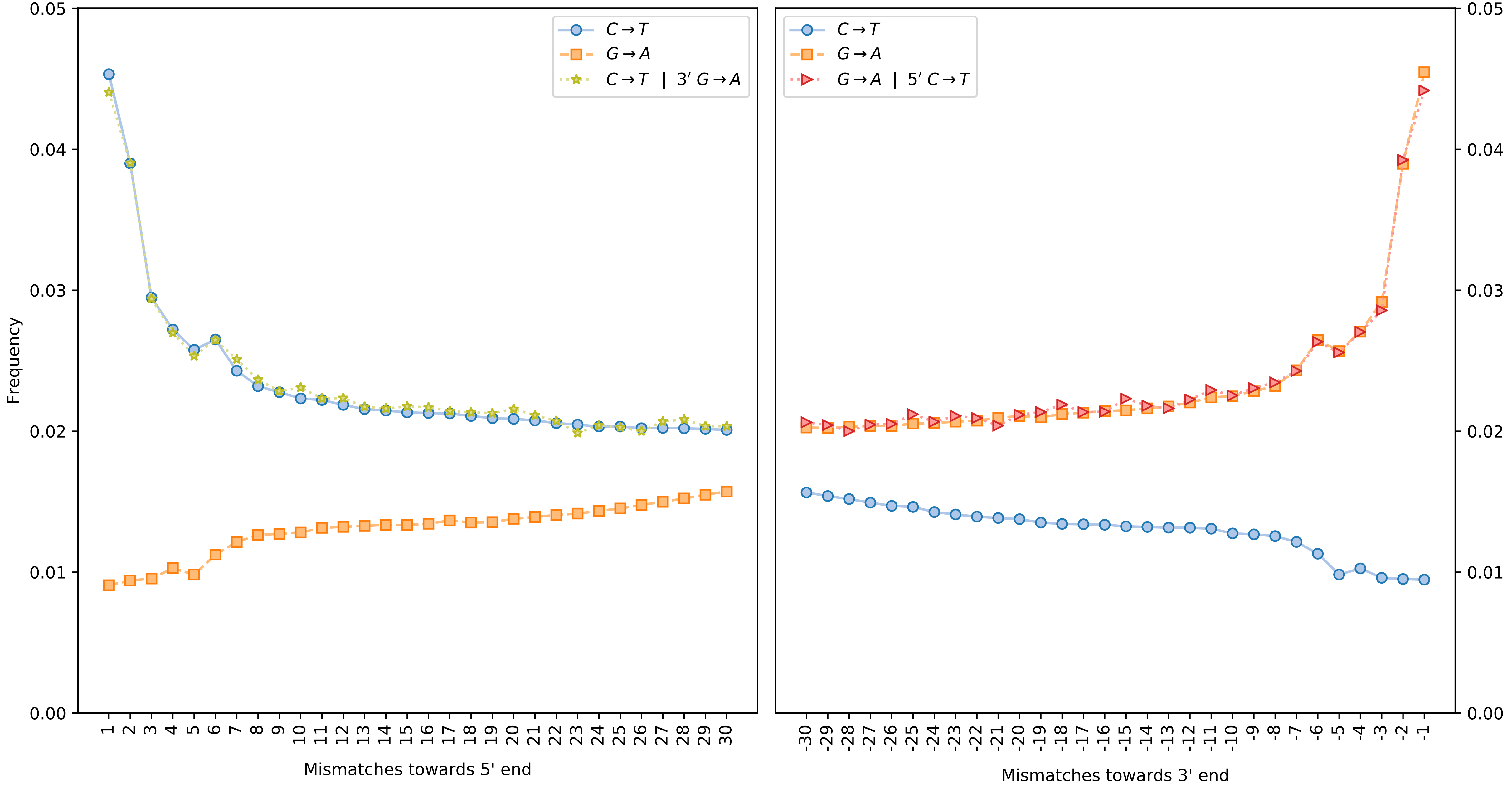

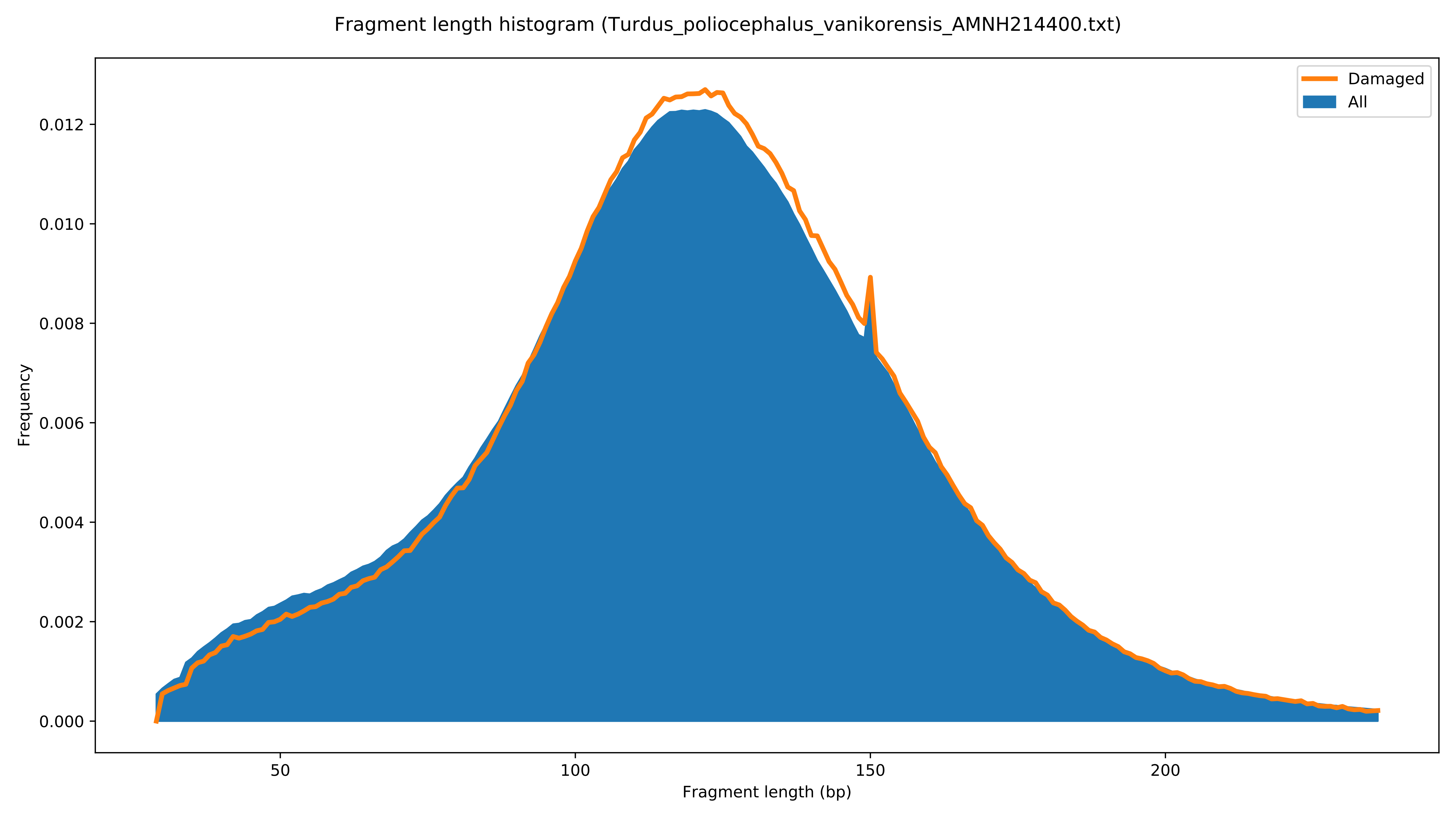

Fragment length histogram (Turdus_poliocephalus_vanikorensis_AMNH214400.txt)
Damaged
All
Frequency
0.000
0.002
0.004
0.006
0.008
0.010
0.012
50
100
150
200
Fragment length (bp)

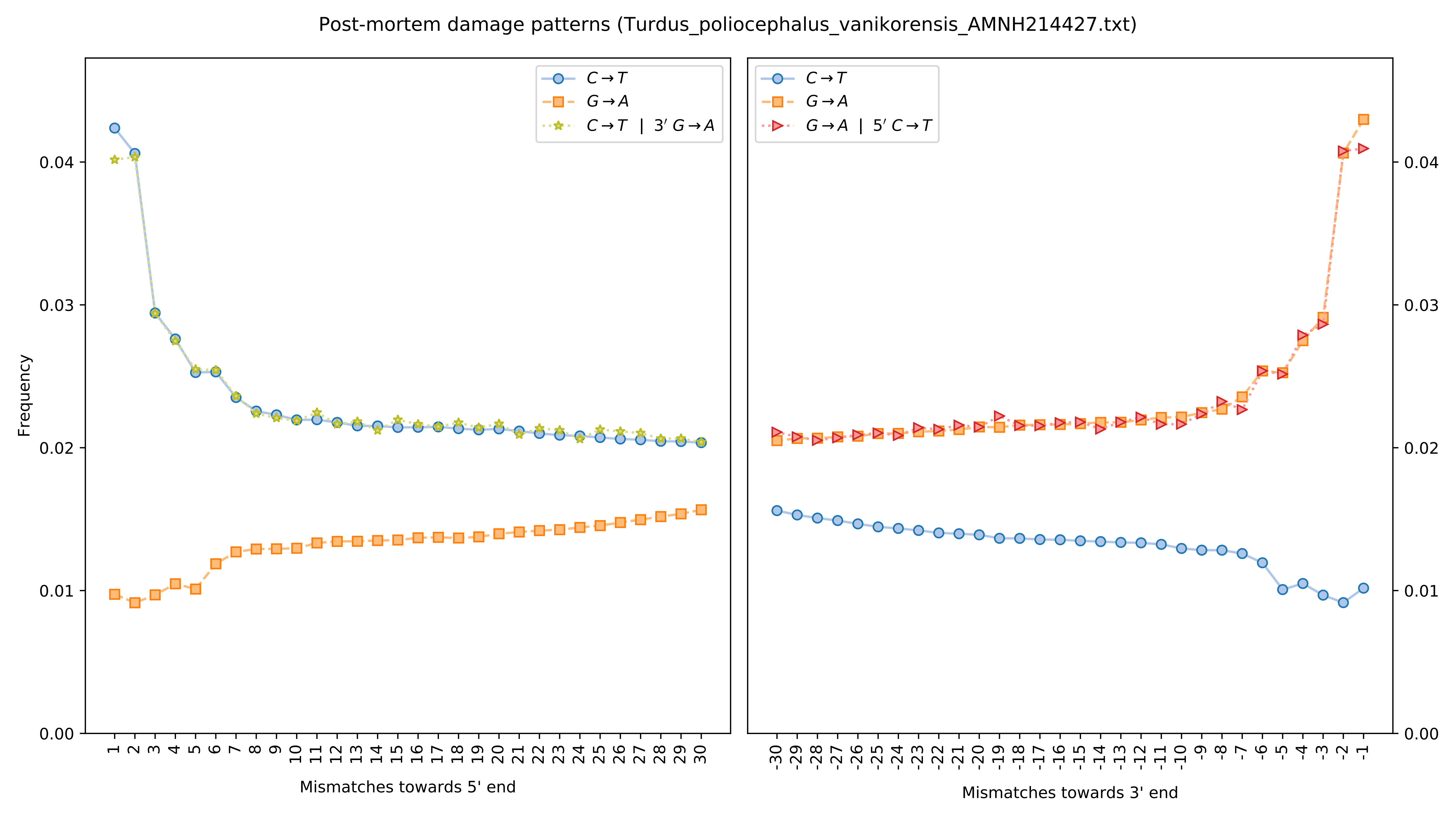

Post-mortem damage patterns (Turdus_poliocephalus_vanikorensis_AMNH214427.txt)
Frequency
Mismatches towards 5' end
Mismatches towards 3' end
C→T
G→A
C→T | 3′ G→A
G→A | 5′ C→T

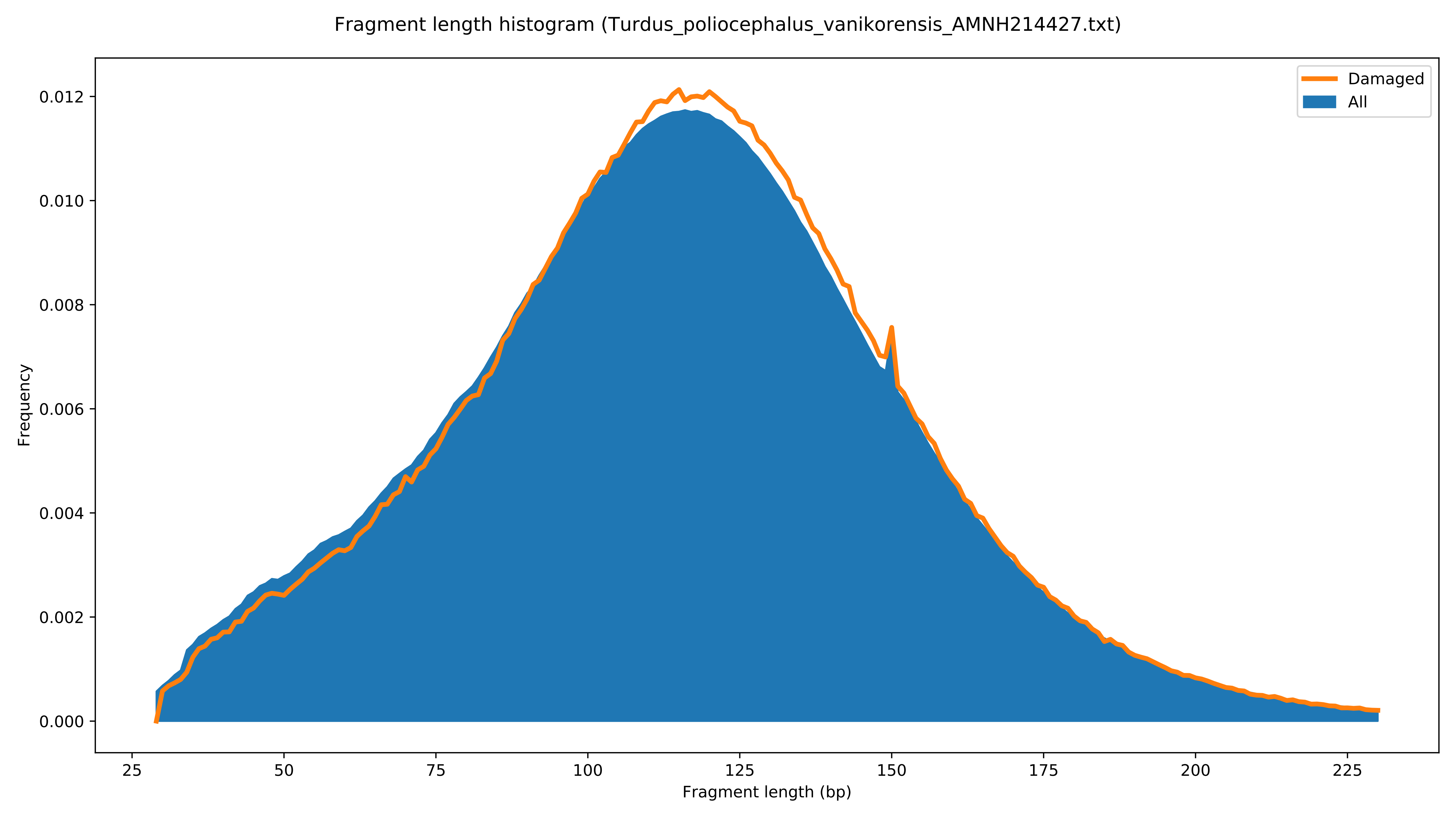

Fragment length histogram (Turdus_poliocephalus_vanikorensis_AMNH214427.txt)
Damaged
All
Frequency
0.000
0.002
0.004
0.006
0.008
0.010
0.012
25
50
75
100
125
150
175
200
225
Fragment length (bp)

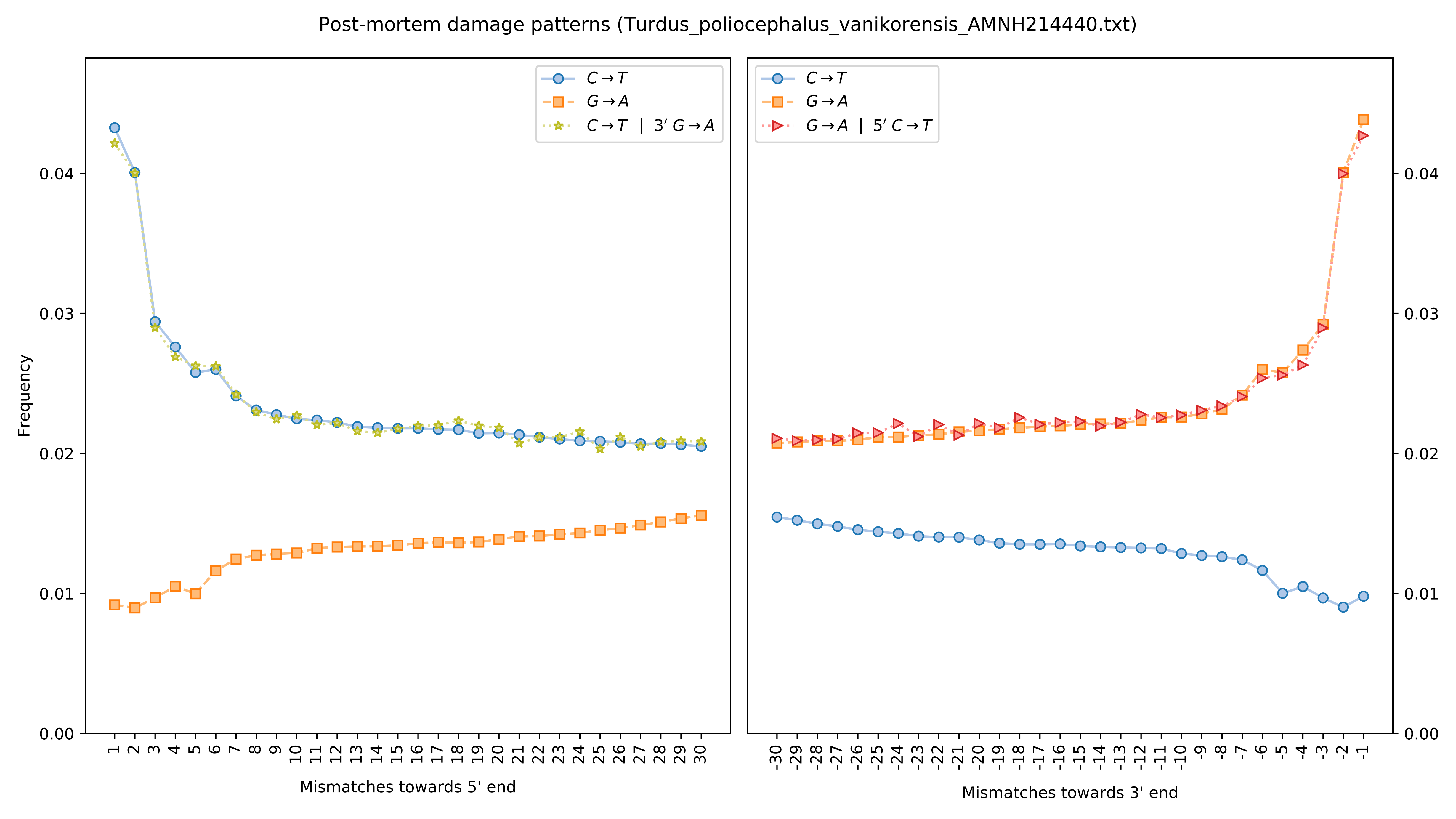
Post-mortem damage patterns (Turdus_poliocephalus_vanikorensis_AMNH214440.txt)
Frequency
Mismatches towards 5' end
Mismatches towards 3' end
C→T
G→A
C→T | 3′ G→A
G→A | 5′ C→T
0.00
0.01
0.02
0.03
0.04

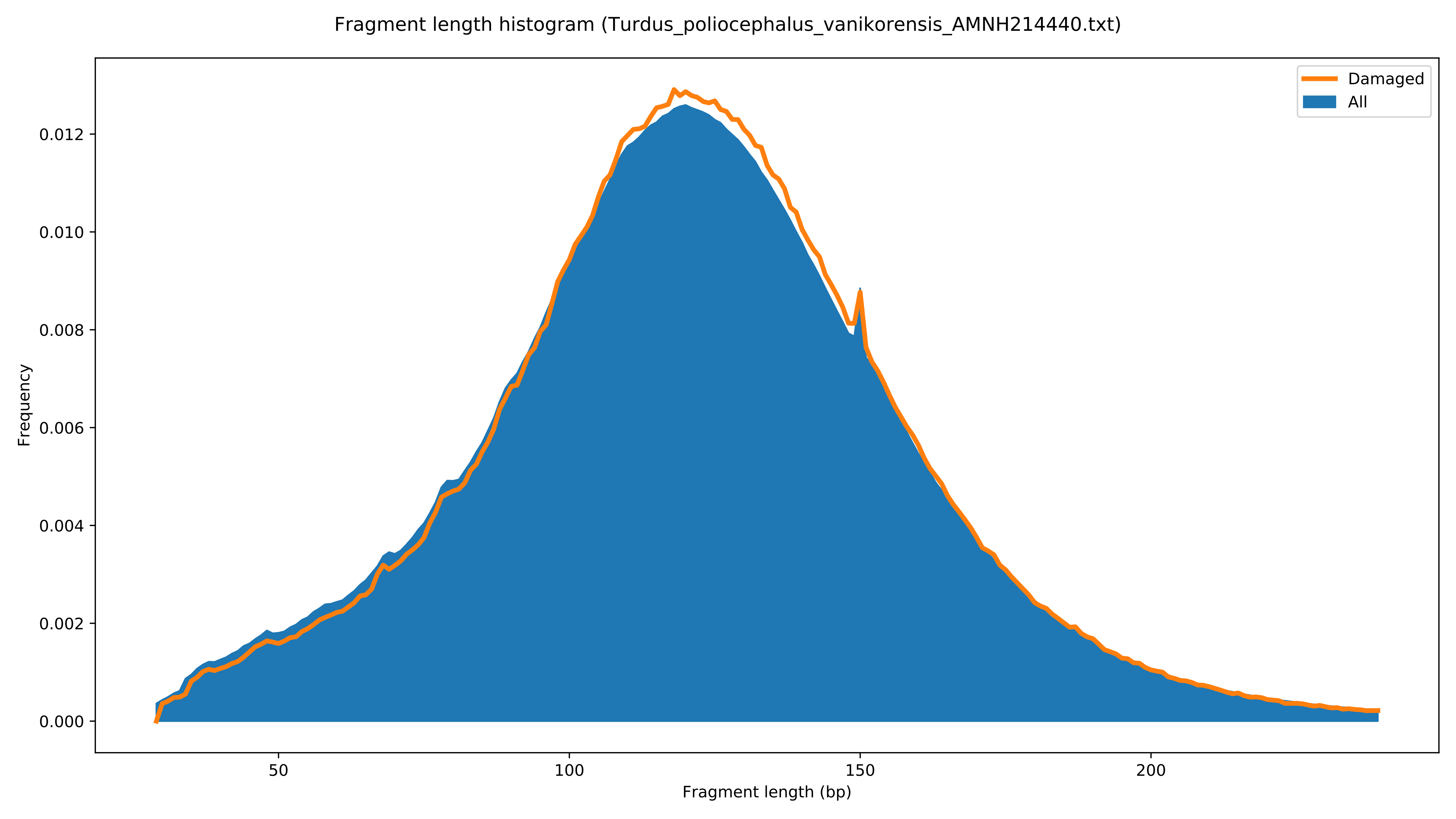

Fragment length histogram (Turdus_poliocephalus_vanikorensis_AMNH214440.txt)
Damaged
All
Frequency
0.000
0.002
0.004
0.006
0.008
0.010
0.012
50
100
150
200
Fragment length (bp)

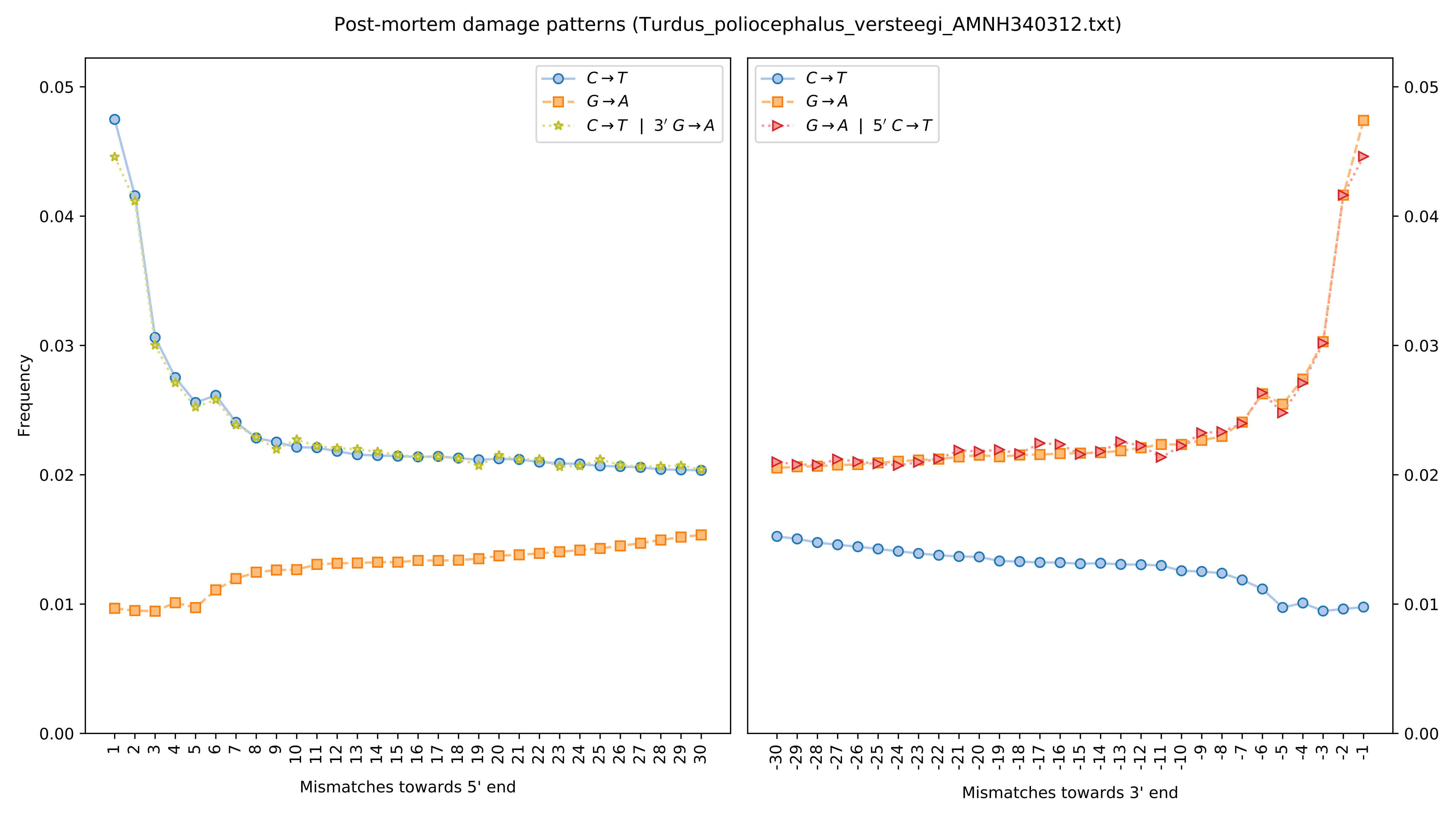

Post-mortem damage patterns (Turdus_poliocephalus_versteegi_AMNH340312.txt)
Frequency
Mismatches towards 5' end
Mismatches towards 3' end
C→T
G→A
C→T | 3′ G→A
G→A | 5′ C→T

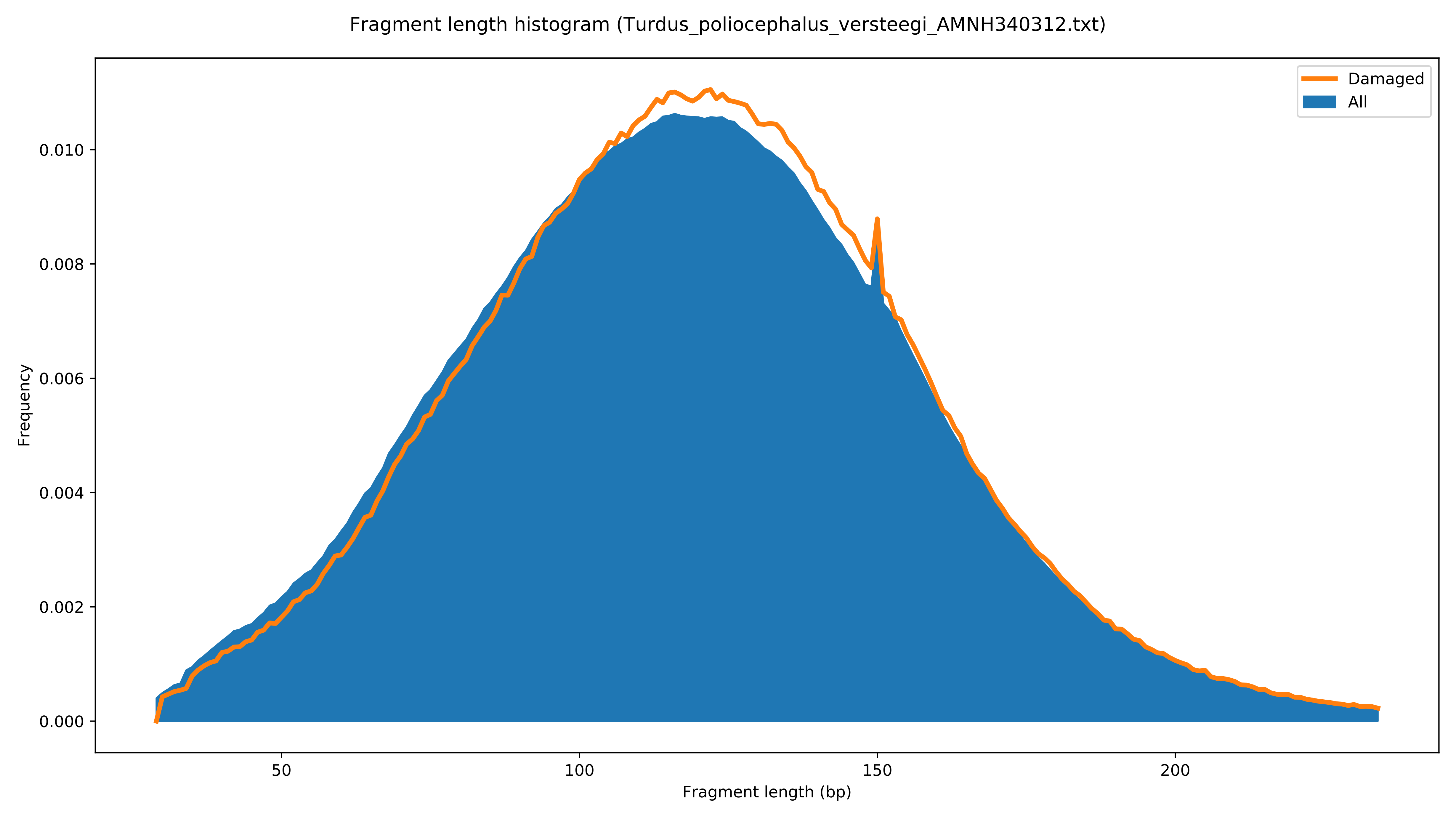

Fragment length histogram (Turdus_poliocephalus_versteegi_AMNH340312.txt)
Damaged
All
Frequency
0.000
0.002
0.004
0.006
0.008
0.010
50
100
150
200
Fragment length (bp)

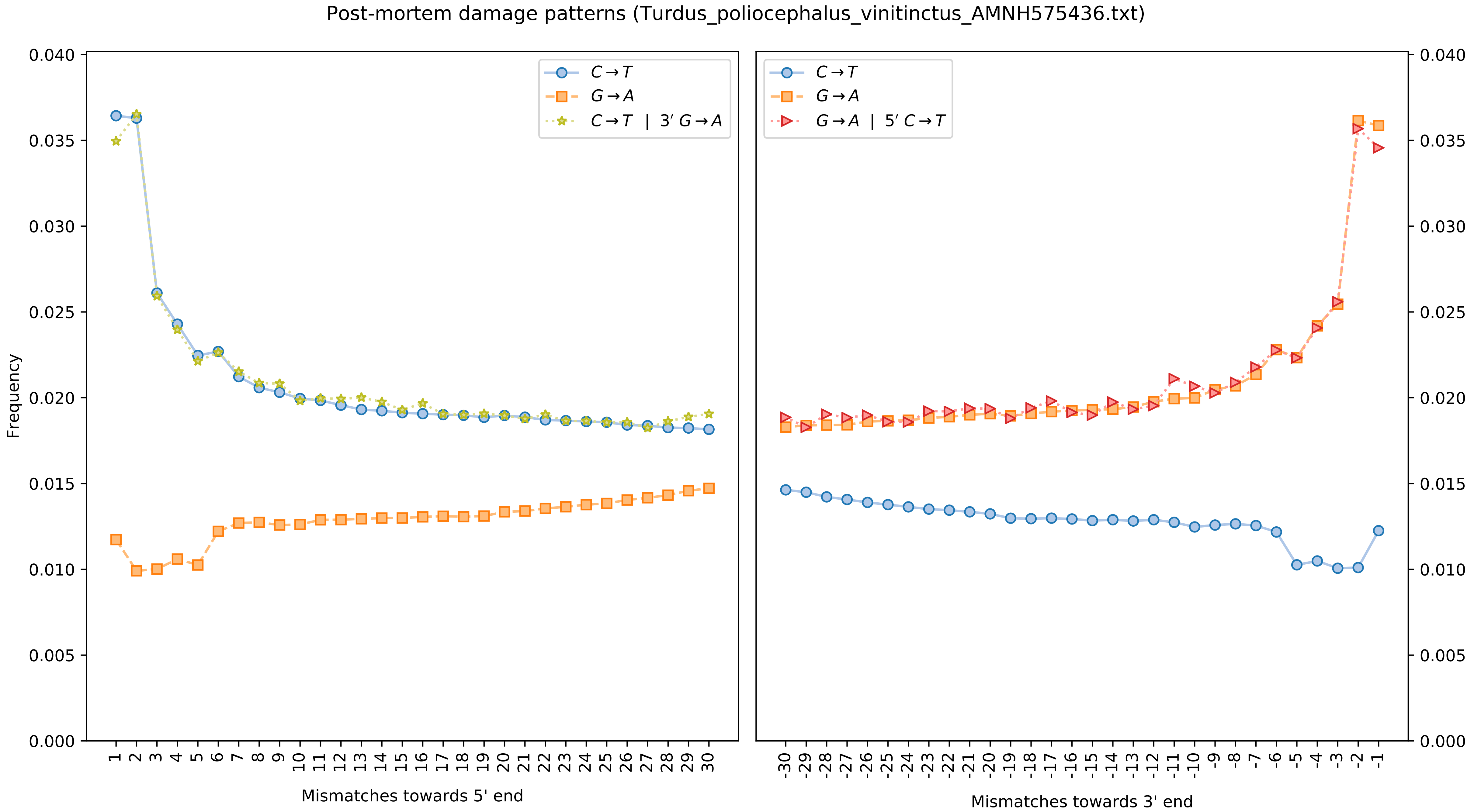
Post-mortem damage patterns (Turdus_poliocephalus_vinitinctus_AMNH575436.txt)
Frequency
Mismatches towards 5' end
Mismatches towards 3' end
C→T
G→A
C→T | 3′ G→A
G→A | 5′ C→T

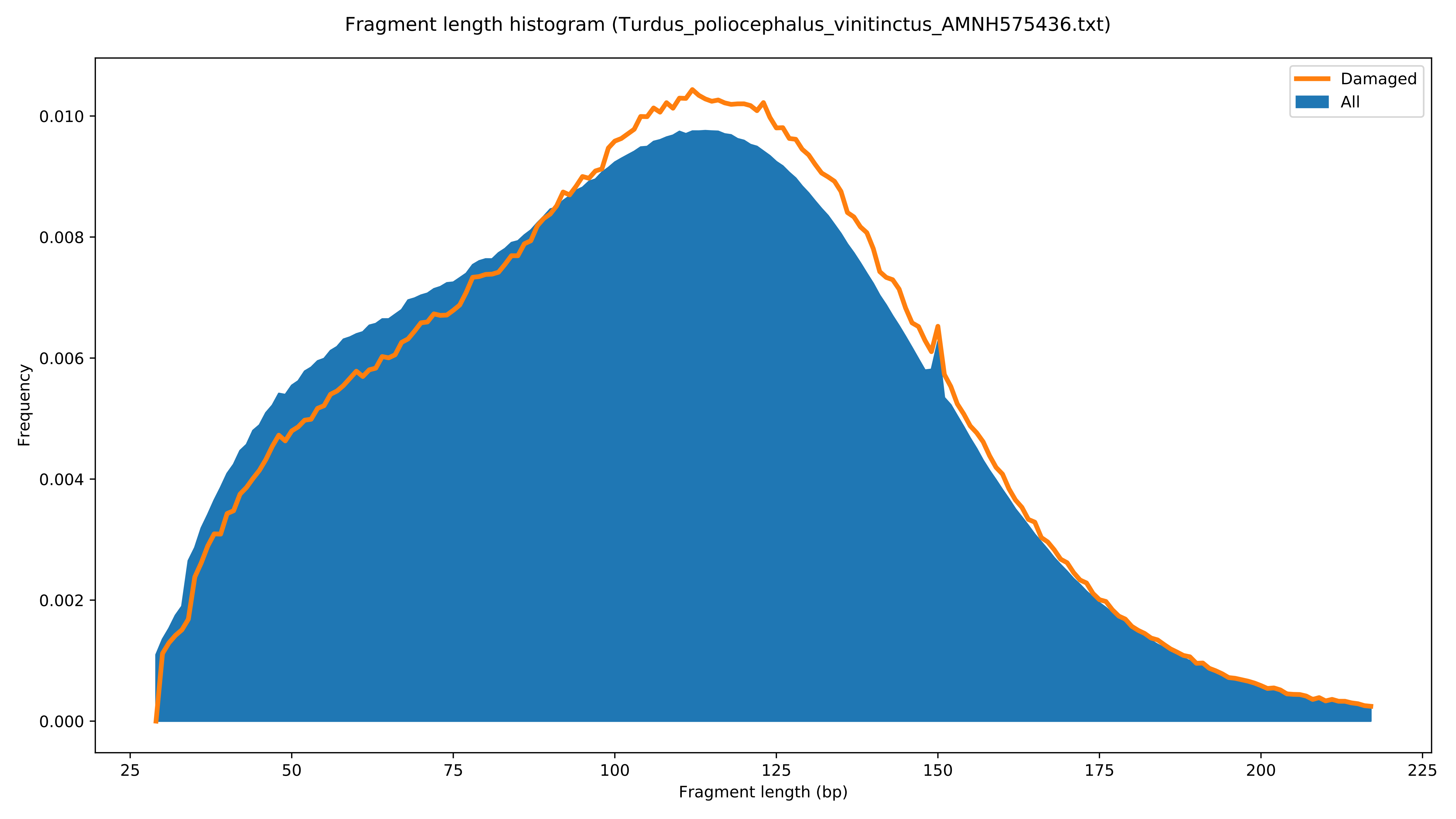

Fragment length histogram (Turdus_poliocephalus_vinitinctus_AMNH575436.txt)
Damaged
All
Frequency
0.000
0.002
0.004
0.006
0.008
0.010
25
50
75
100
125
150
175
200
225
Fragment length (bp)

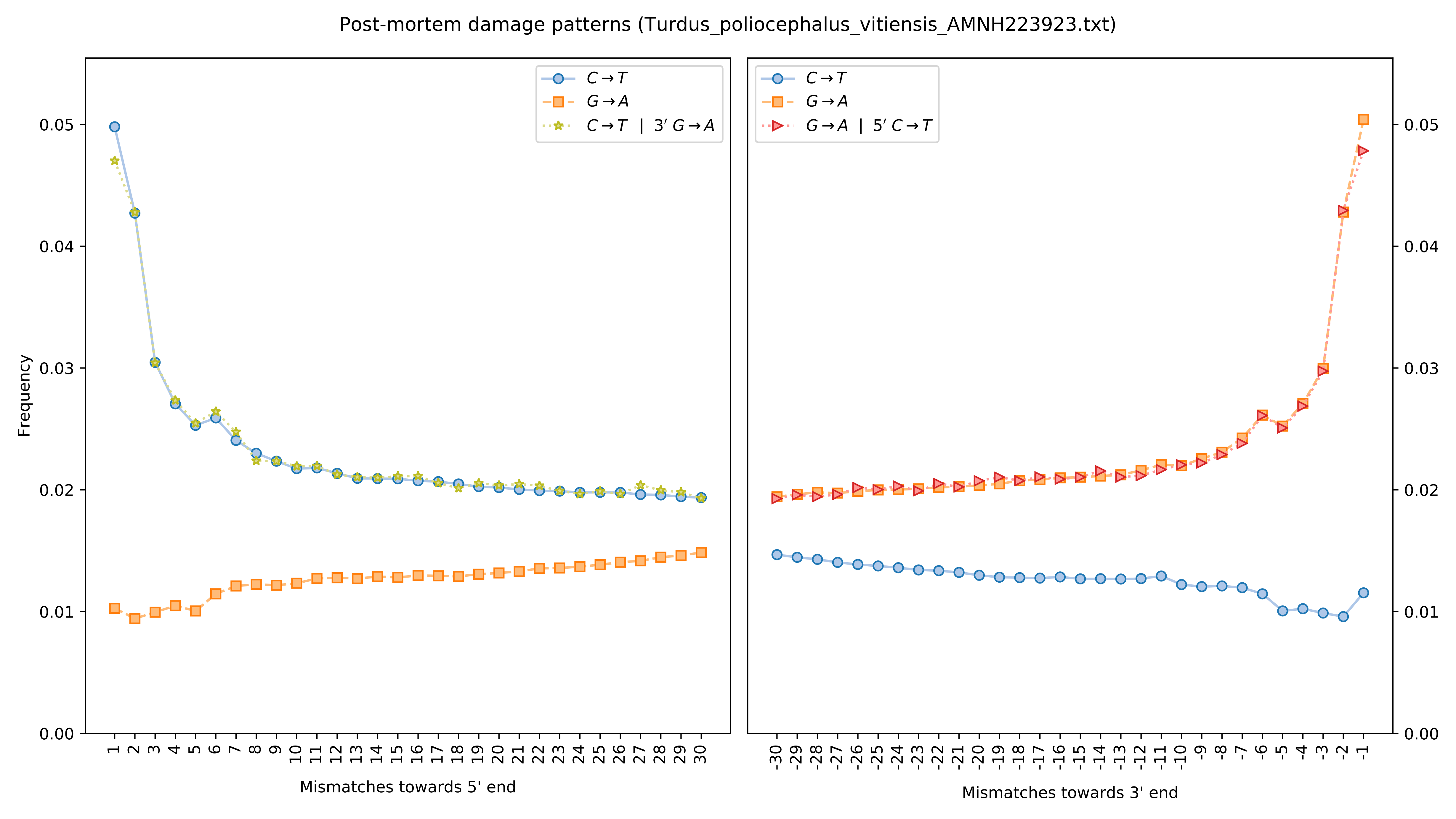

Post-mortem damage patterns (Turdus_poliocephalus_vitiensis_AMNH223923.txt)
Frequency
Mismatches towards 5' end
Mismatches towards 3' end
C→T
G→A
C→T | 3′ G→A
G→A | 5′ C→T
0.00
0.01
0.02
0.03
0.04
0.05

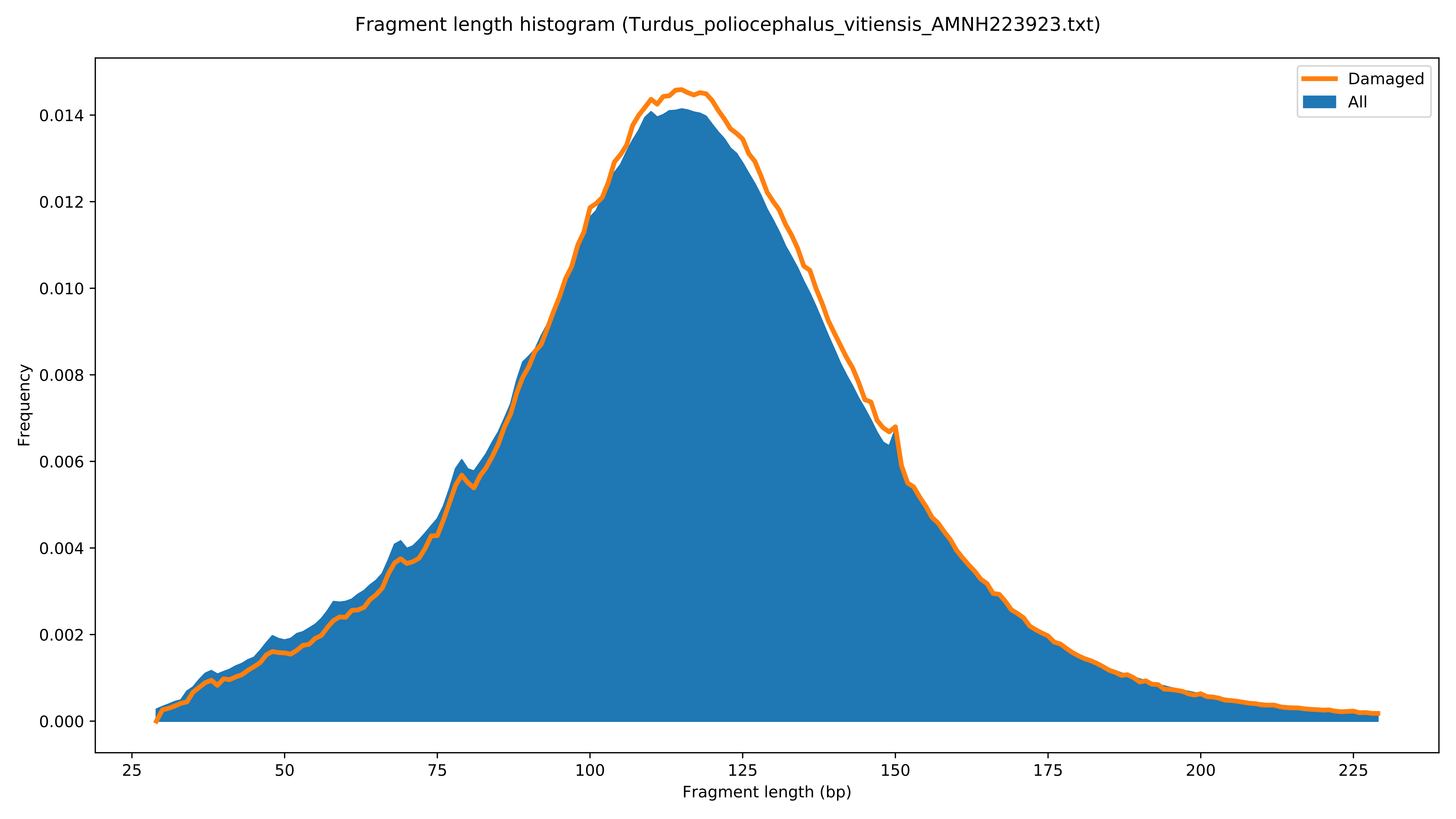

Fragment length histogram (Turdus_poliocephalus_vitiensis_AMNH223923.txt)
Damaged
All
Frequency
0.000
0.002
0.004
0.006
0.008
0.010
0.012
0.014
25
50
75
100
125
150
175
200
225
Fragment length (bp)

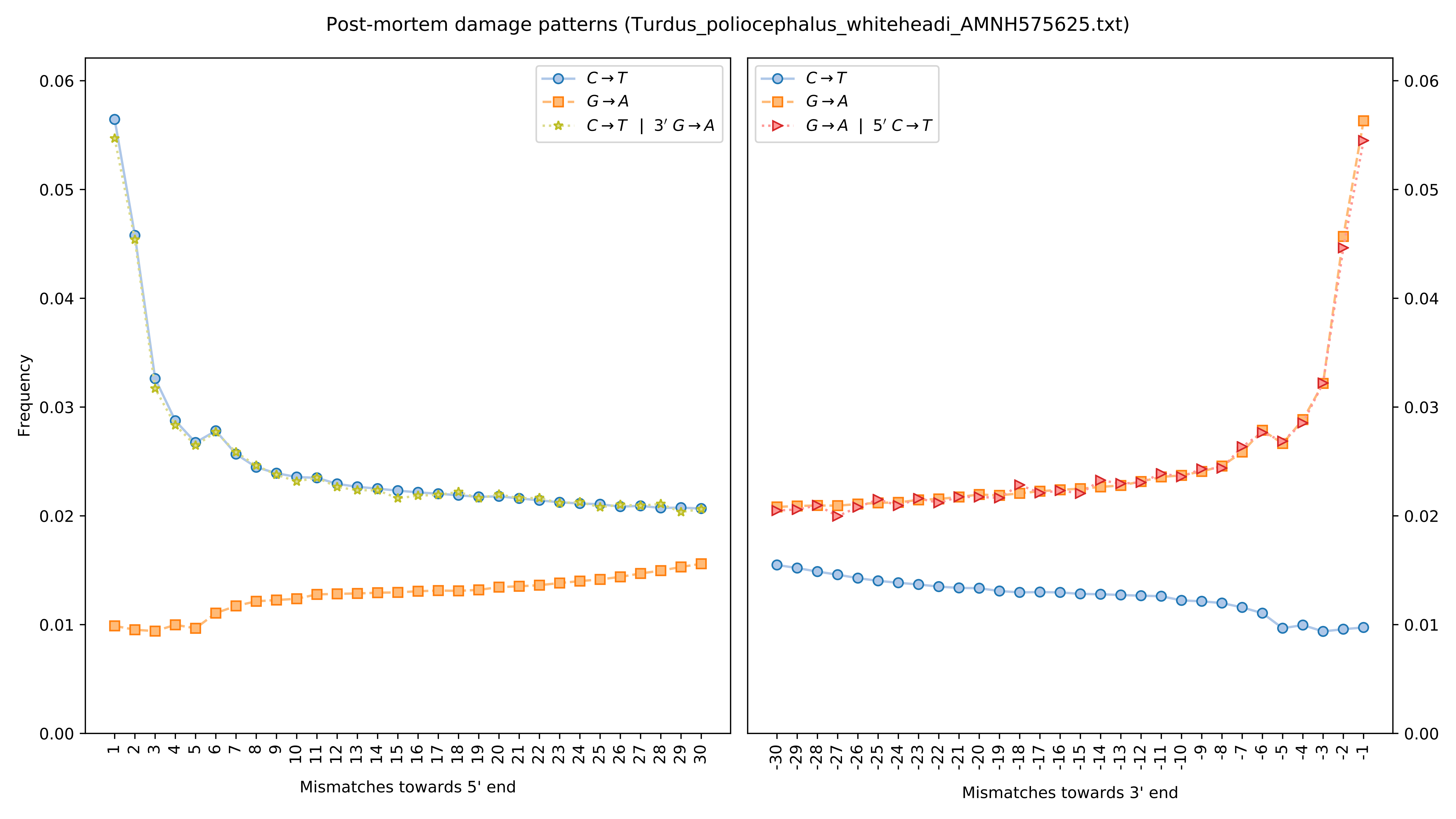

Post-mortem damage patterns (Turdus_poliocephalus_whiteheadi_AMNH575625.txt)
Frequency
Mismatches towards 5' end
Mismatches towards 3' end
C→T
G→A
C→T | 3′ G→A
G→A | 5′ C→T

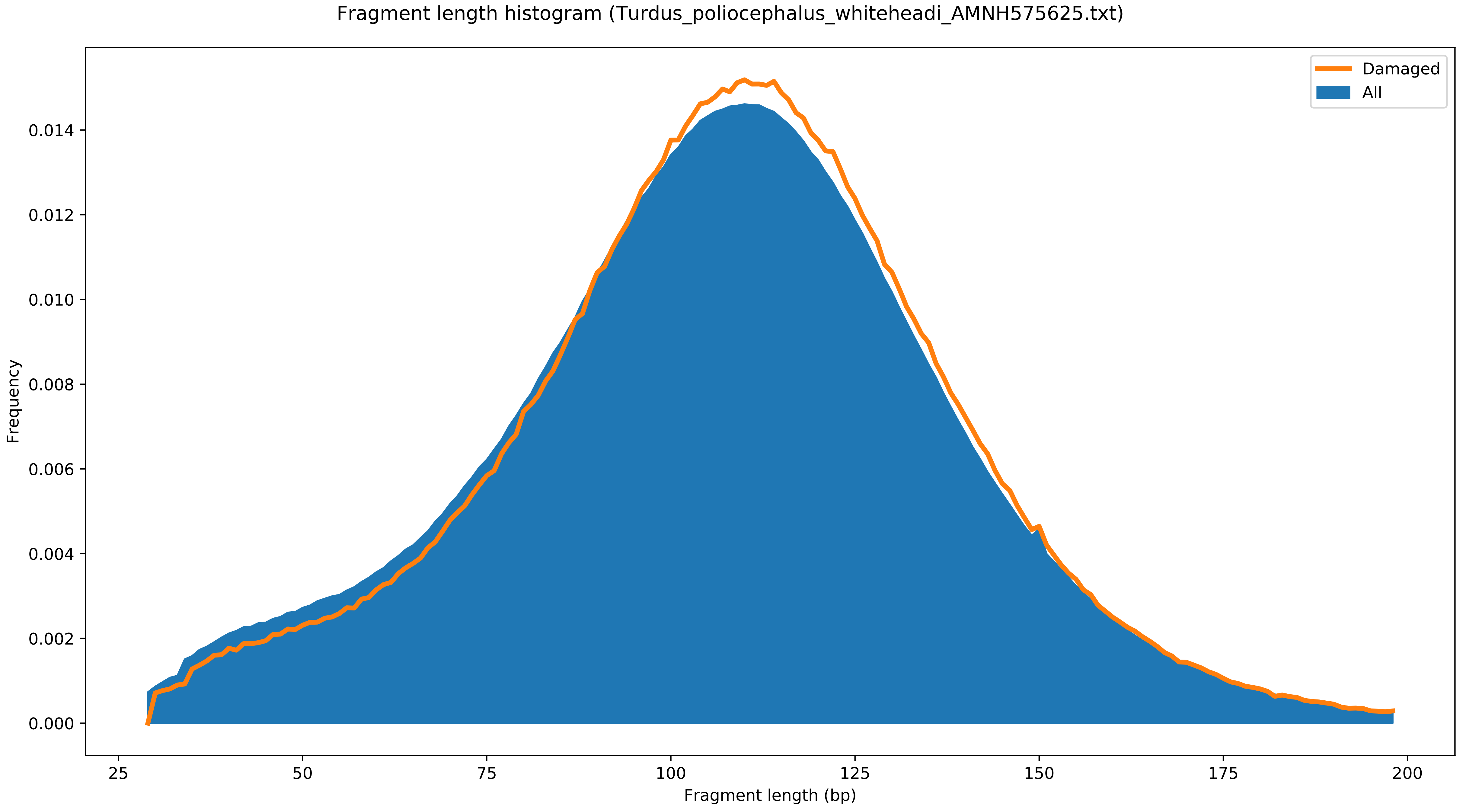

Fragment length histogram (Turdus_poliocephalus_whiteheadi_AMNH575625.txt)
Damaged
All
Frequency
0.014
0.012
0.010
0.008
0.006
0.004
0.002
0.000
25
50
75
100
125
150
175
200
Fragment length (bp)

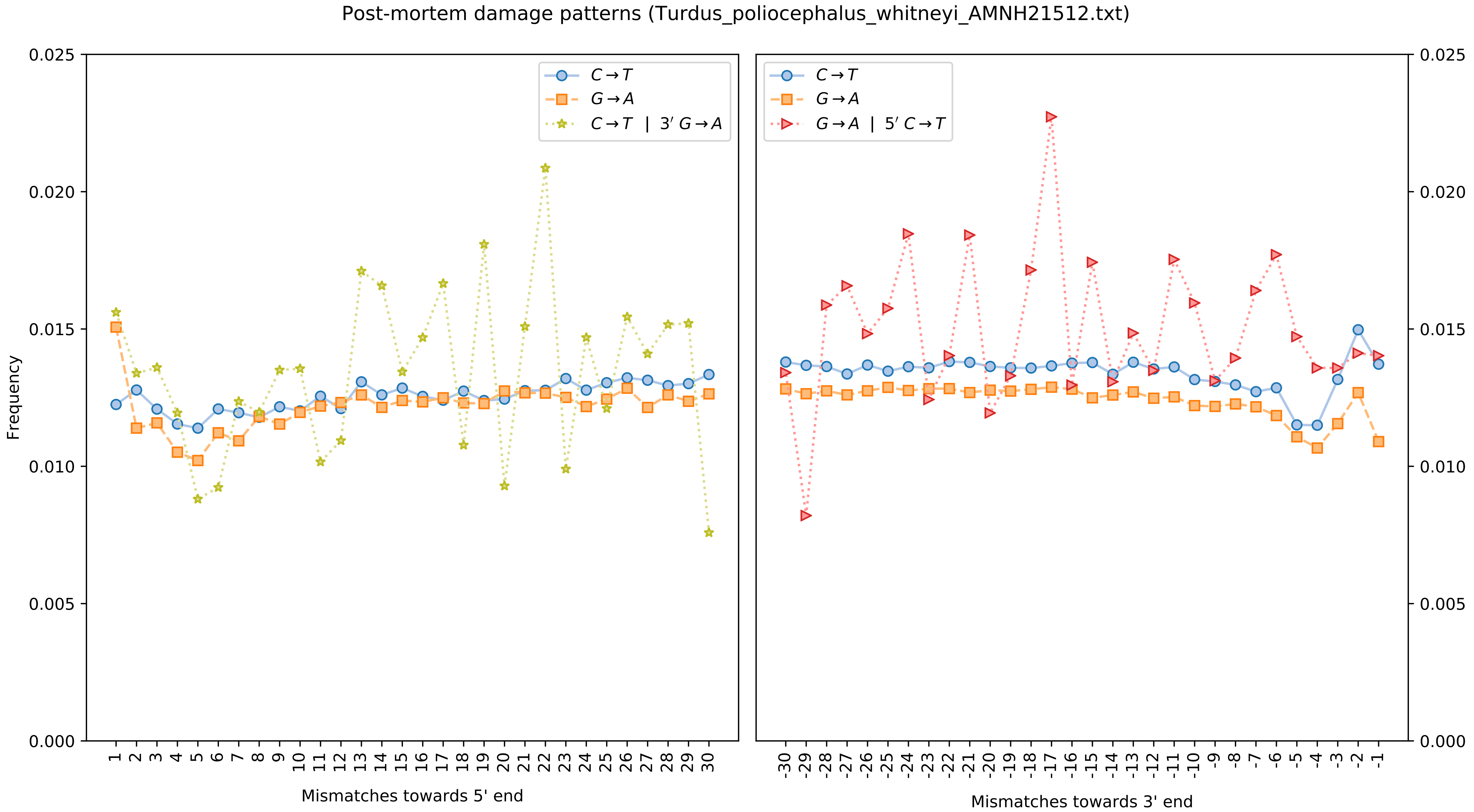

Post-mortem damage patterns (Turdus_poliocephalus_whitneyi_AMNH21512.txt)
Frequency
0.000
0.005
0.010
0.015
0.020
0.025
C→T
G→A
C→T | 3′ G→A
G→A | 5′ C→T
1 2 3 4 5 6 7 8 9 10 11 12 13 14 15 16 17 18 19 20 21 22 23 24 25 26 27 28 29 30
Mismatches towards 5' end
-30 -29 -28 -27 -26 -25 -24 -23 -22 -21 -20 -19 -18 -17 -16 -15 -14 -13 -12 -11 -10 -9 -8 -7 -6 -5 -4 -3 -2 -1
Mismatches towards 3' end

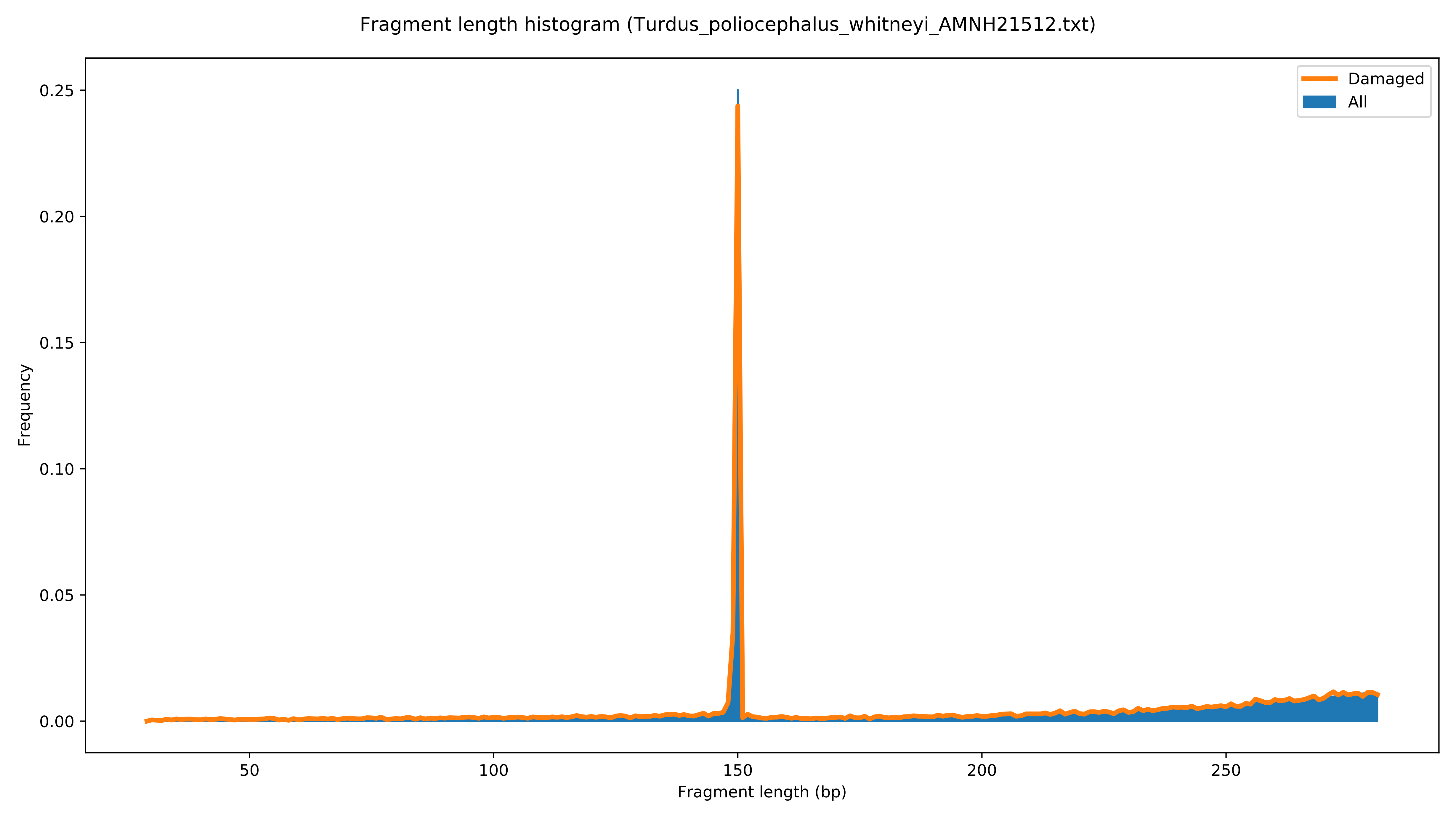
Fragment length histogram (Turdus_poliocephalus_whitneyi_AMNH21512.txt)
Damaged
All
Frequency
0.00
0.05
0.10
0.15
0.20
0.25
50
100
150
200
250
Fragment length (bp)

Post-mortem damage patterns (Turdus_poliocephalus_xanthopus_AMNH575554.txt)

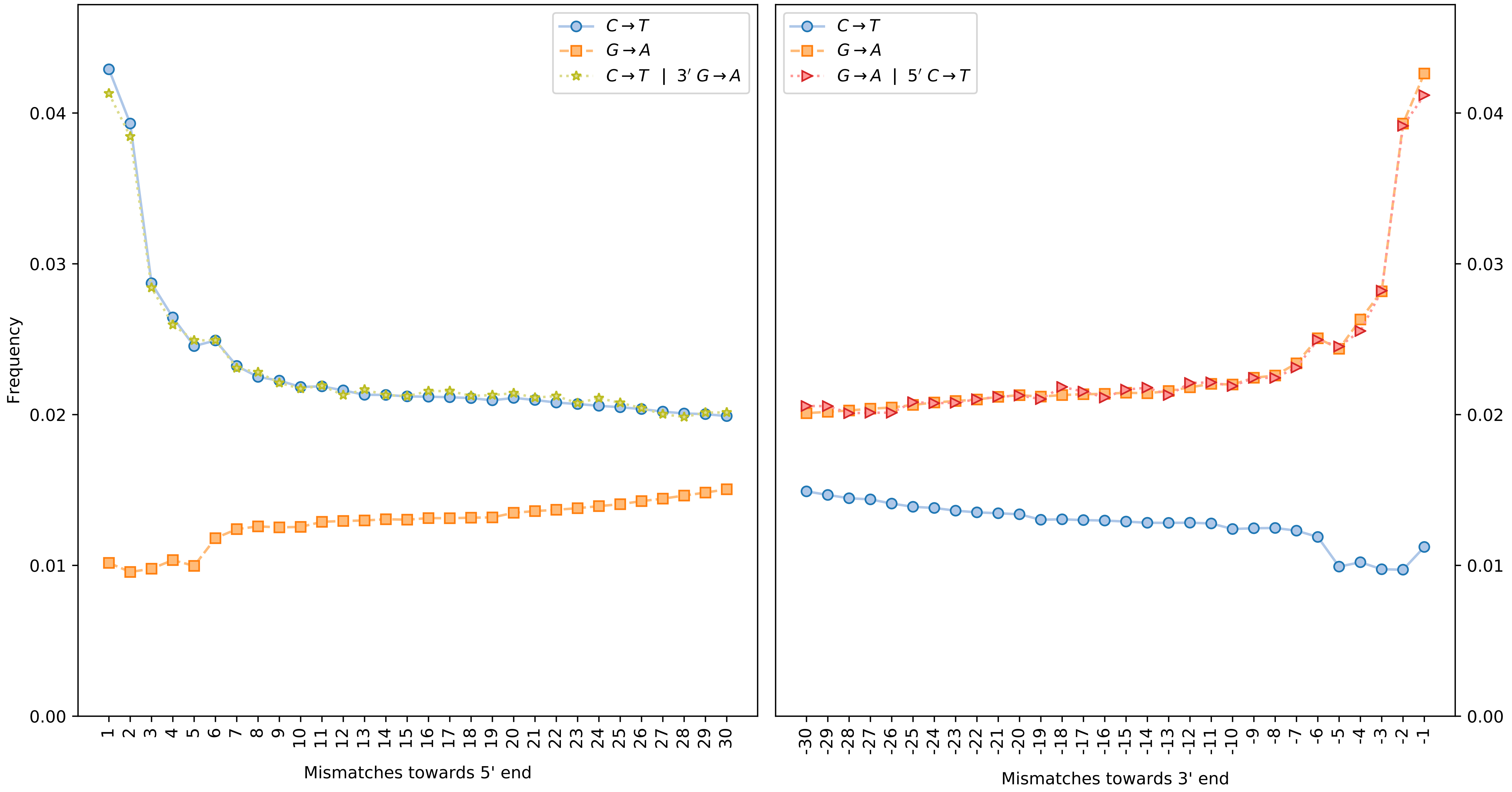

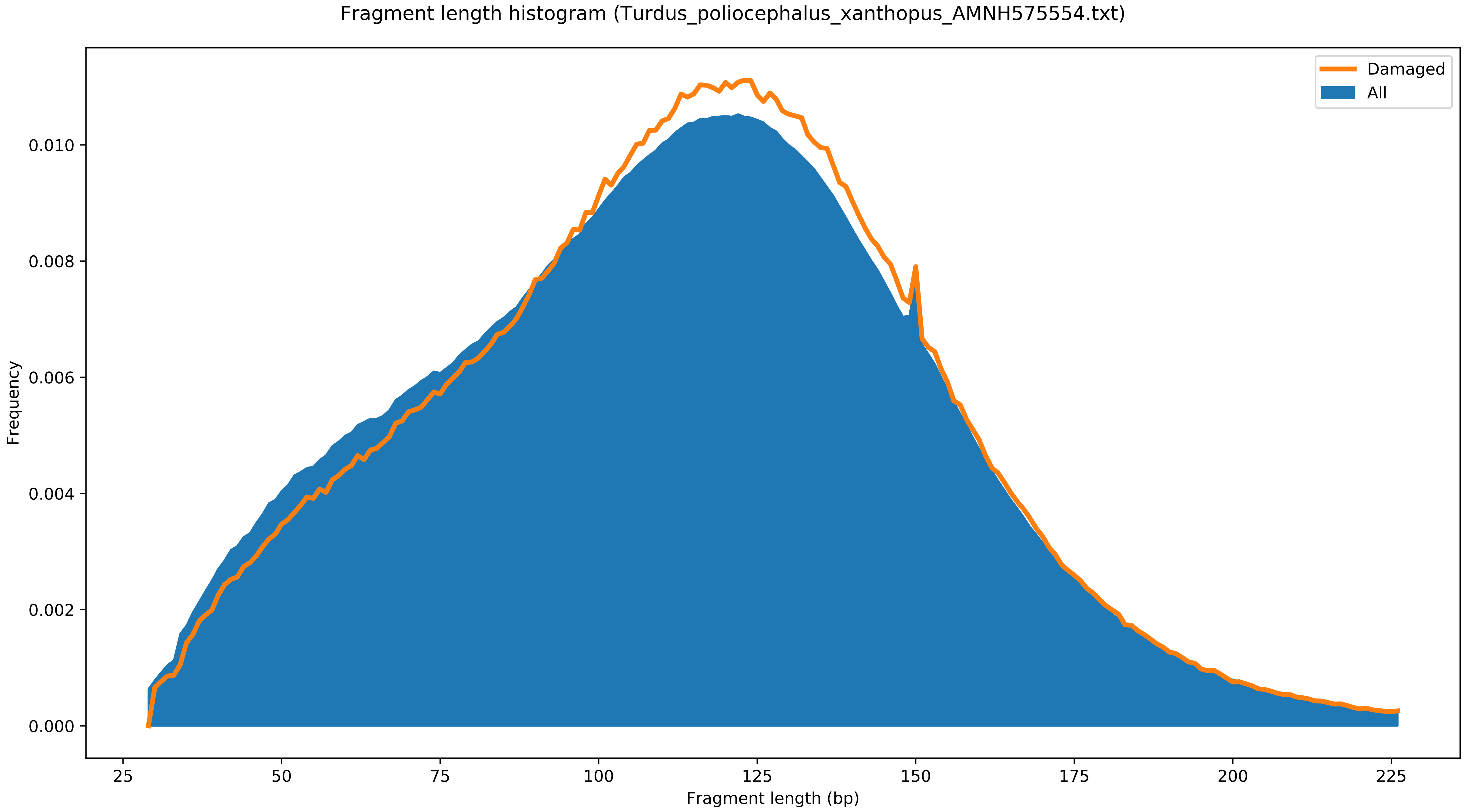

Fragment length histogram (Turdus_poliocephalus_xanthopus_AMNH575554.txt)
Damaged
All
Frequency
0.000
0.002
0.004
0.006
0.008
0.010
25
50
75
100
125
150
175
200
225
Fragment length (bp)
