## Supplementary File 4 for "Population genomics of the island thrush elucidates one of earth’s great archipelagic radiations"

*Turdus celaenops* NRM 90188019 (Japan)

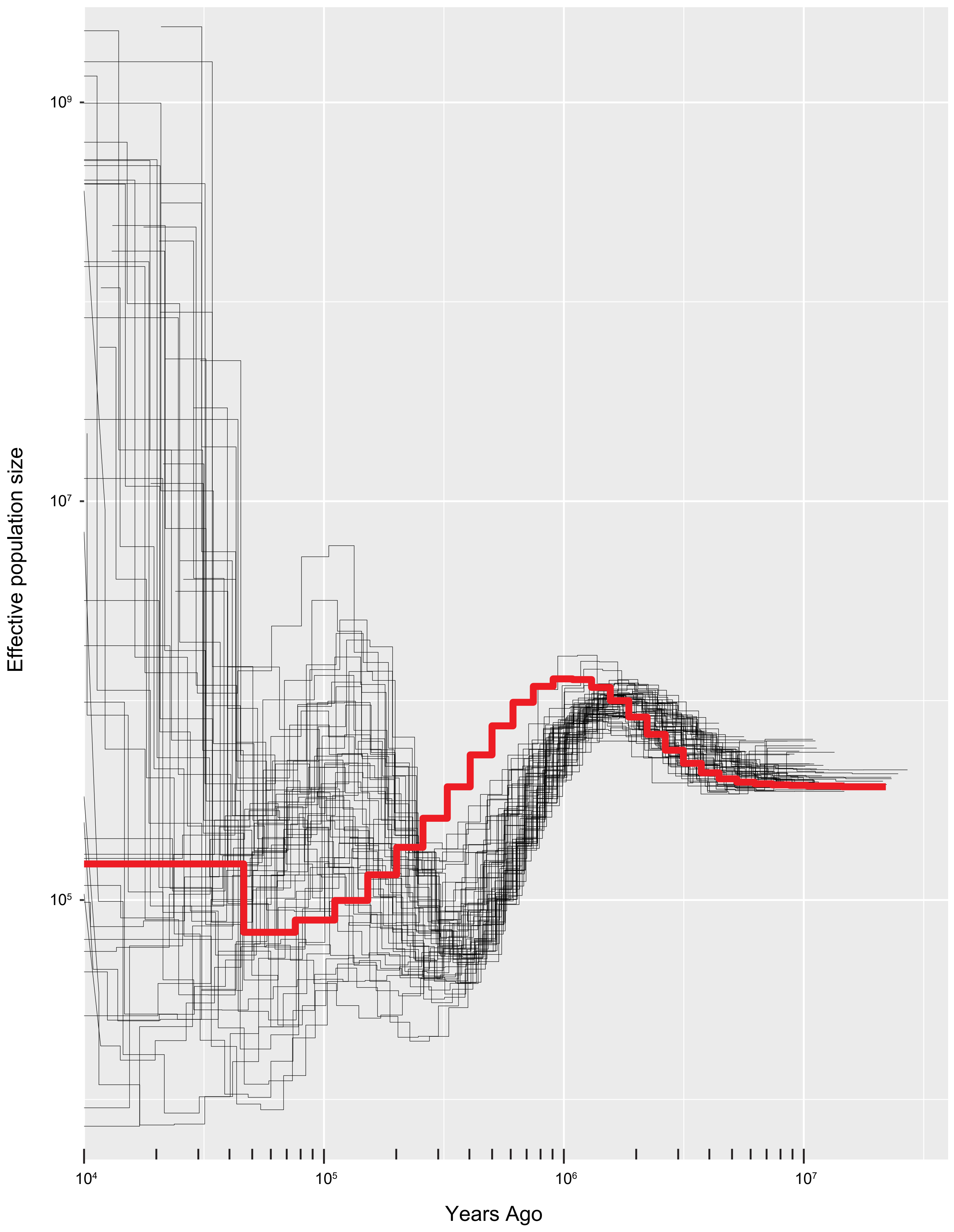

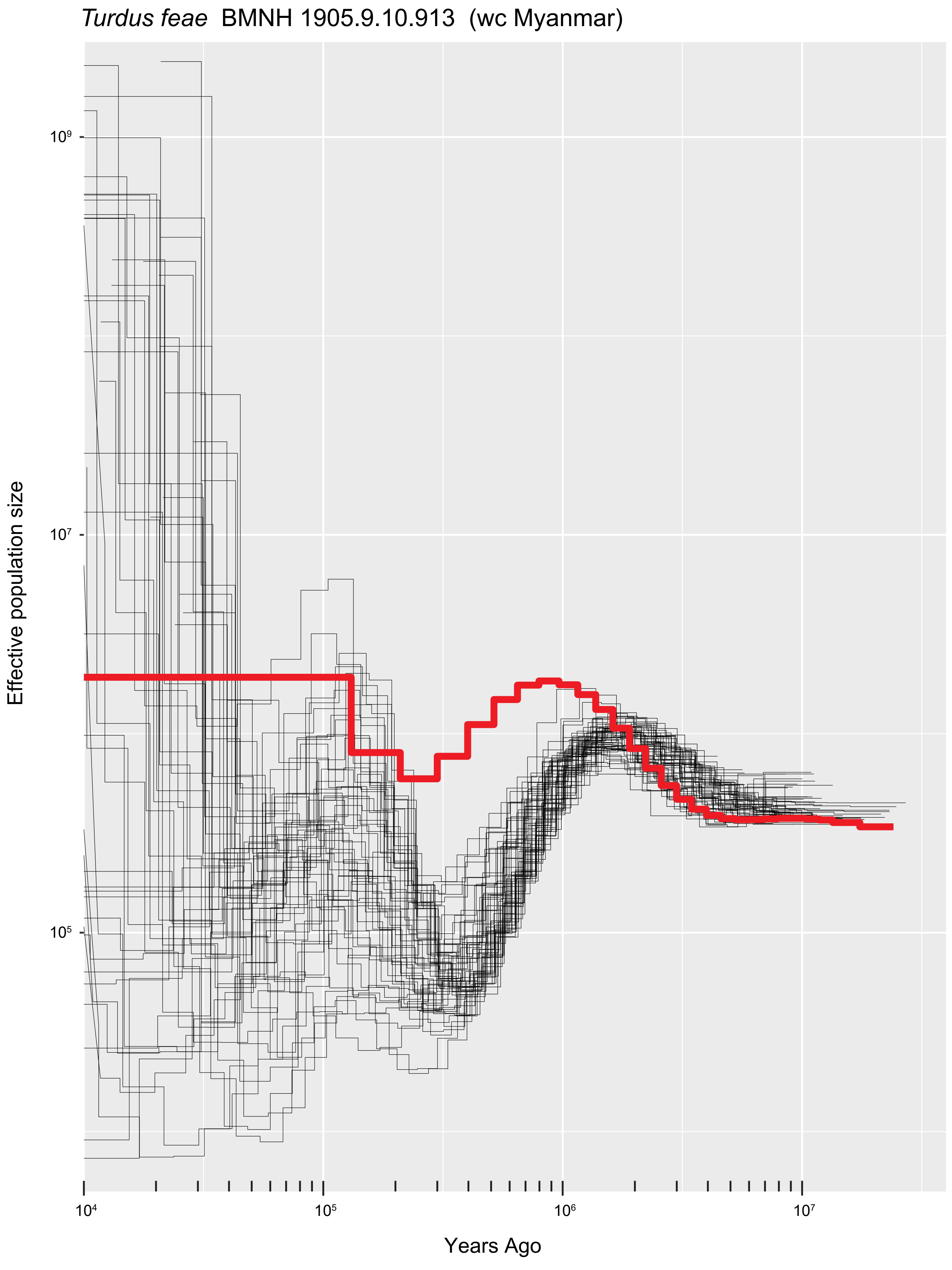
Turdus feae  BMNH 1905.9.10.913  (wc Myanmar)
Effective population size
$10^9$
$10^7$
$10^5$
$10^4$
$10^5$
$10^6$
$10^7$
Years Ago

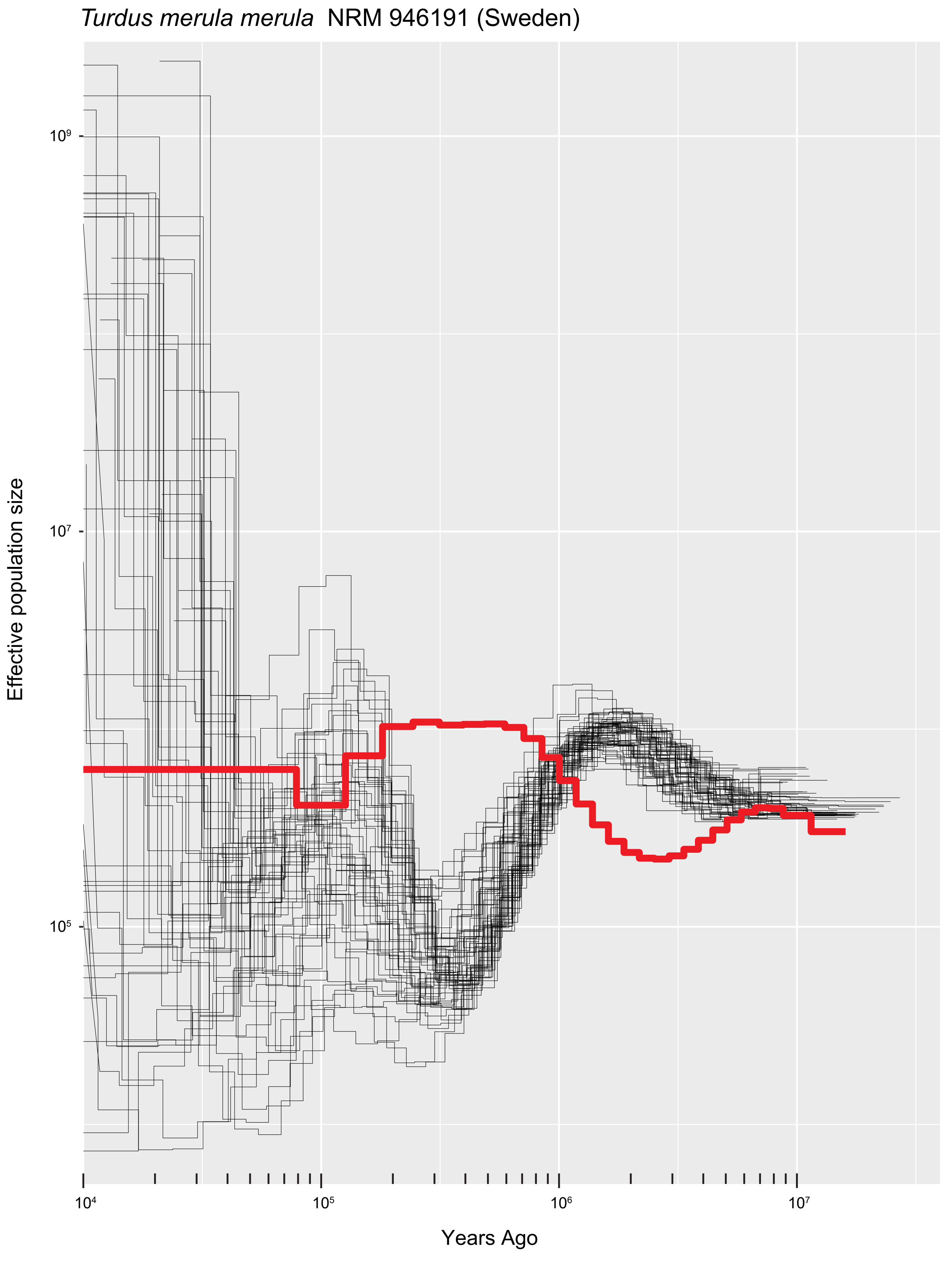

*Turdus merula merula* NRM 946191 (Sweden)
Effective population size
10⁹
10⁷
10⁵
10⁴
10⁵
10⁶
10⁷
Years Ago

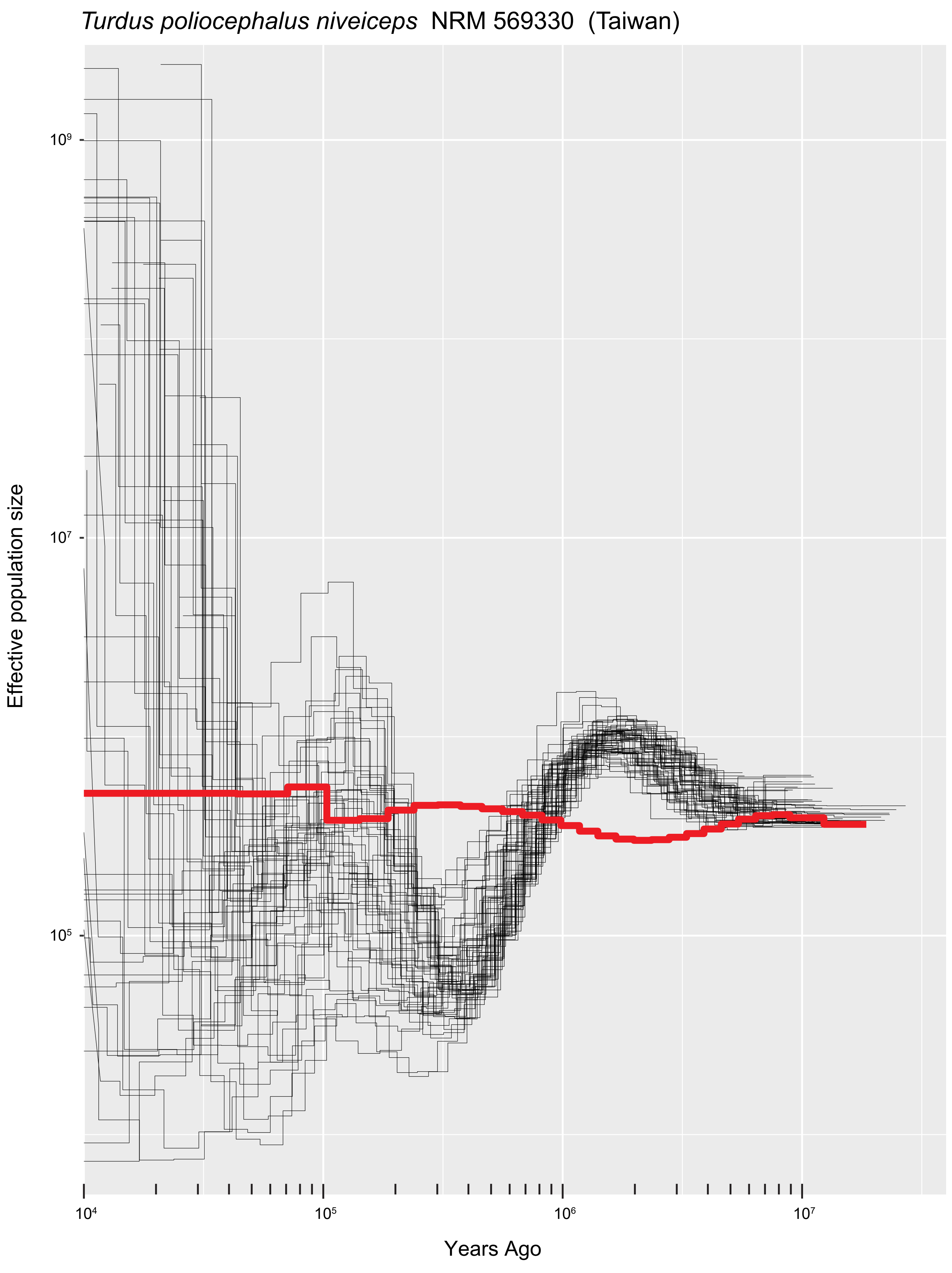

*Turdus poliocephalus niveiceps* NRM 569330 (Taiwan)
Effective population size
10⁹
10⁷
10⁵
Years Ago
10⁴
10⁵
10⁶
10⁷

### *Turdus obscurus* UWBM 78352 (e Russia)

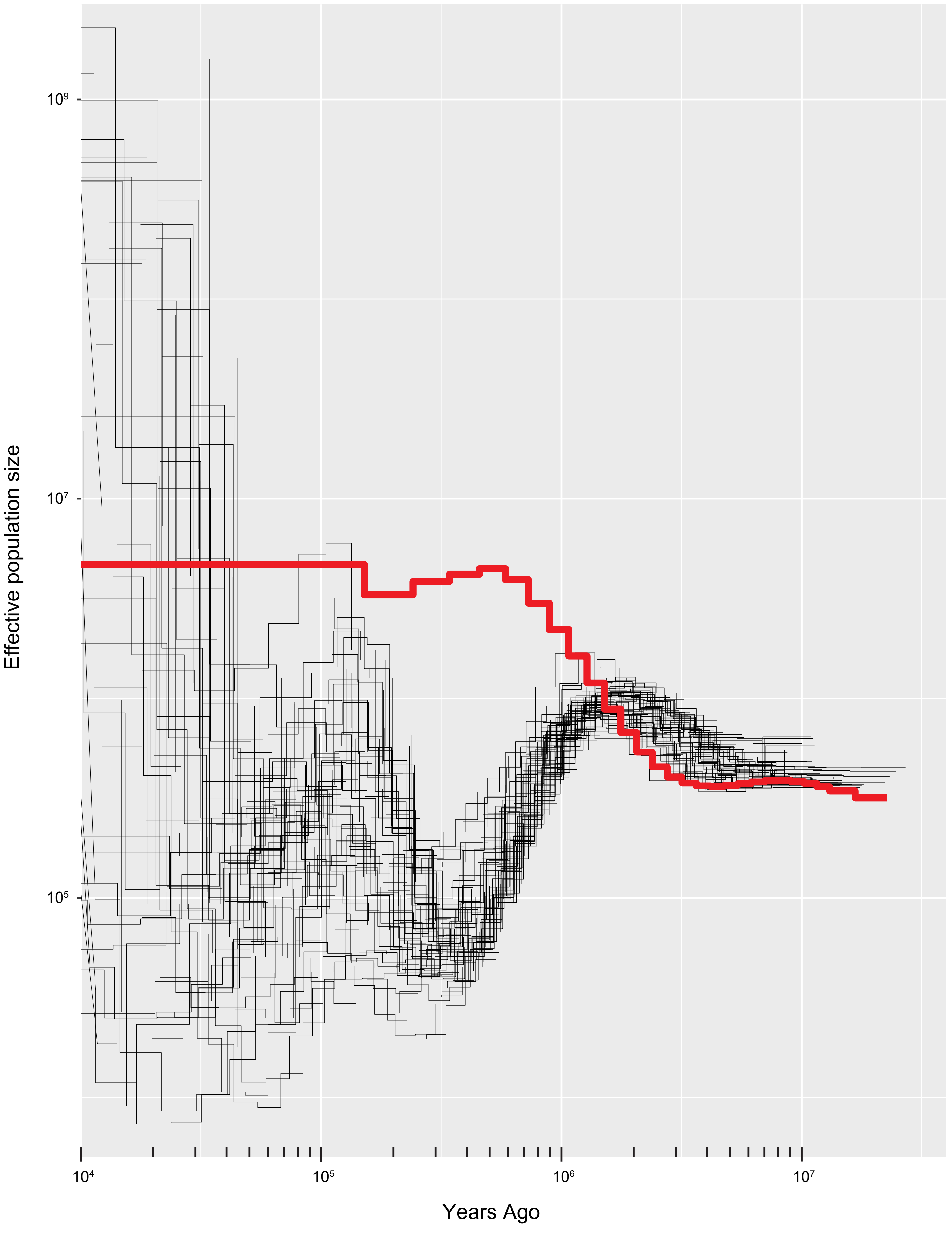

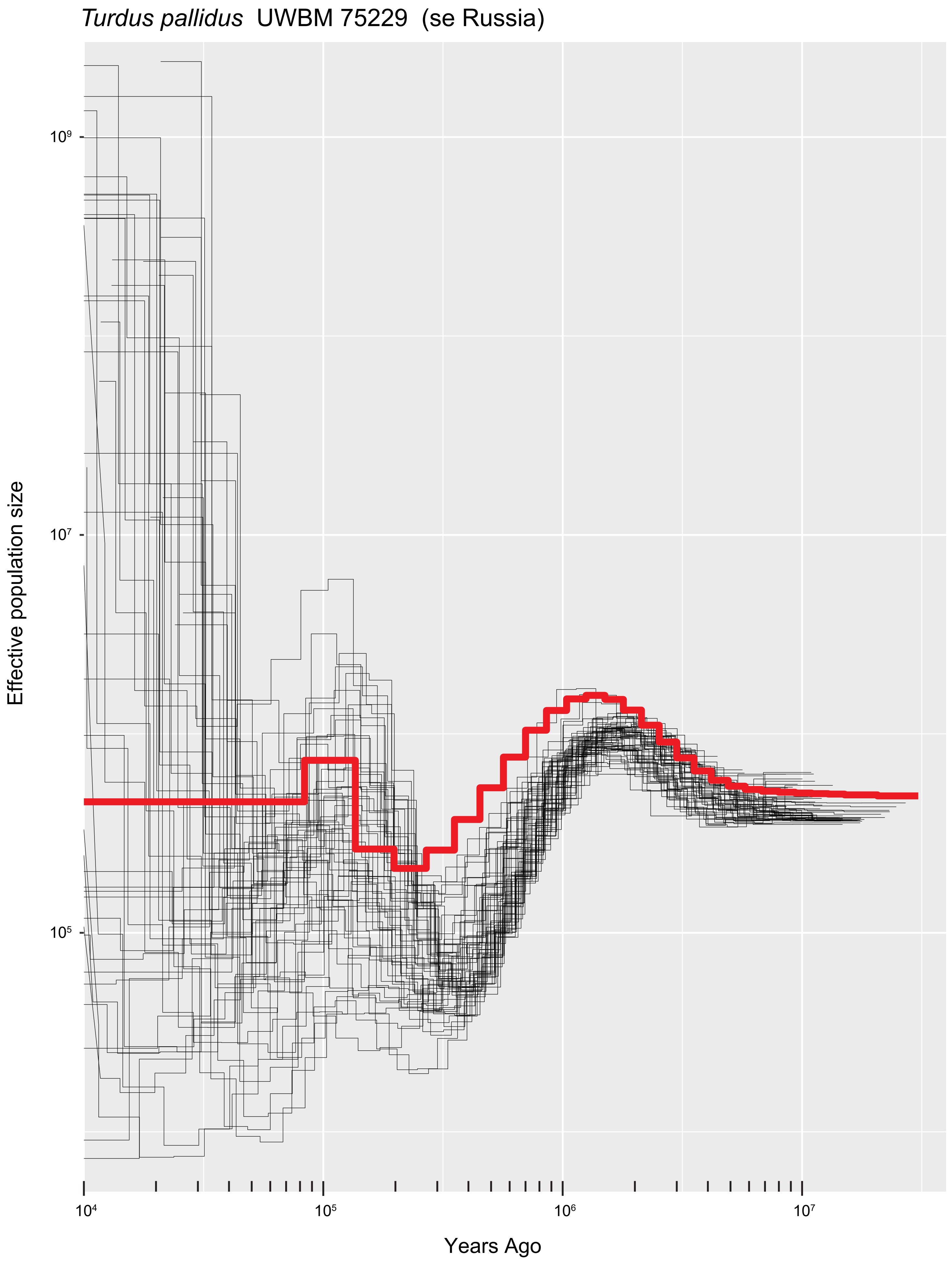

*Turdus pallidus* UWBM 75229 (se Russia)
Effective population size
10⁹
10⁷
10⁵
10⁴
10⁵
10⁶
10⁷
Years Ago

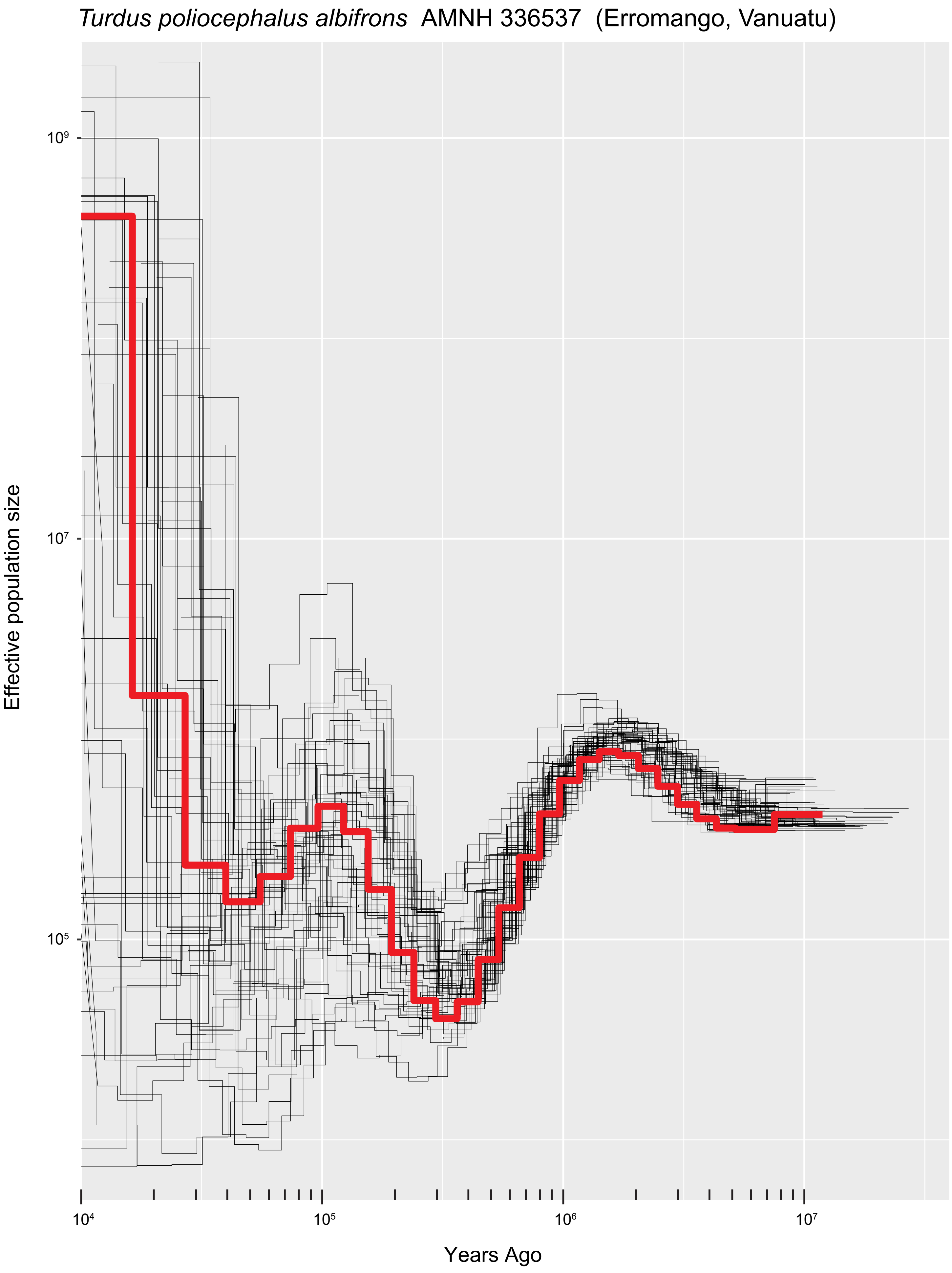

*Turdus poliocephalus albifrons* AMNH 336537 (Erromango, Vanuatu)
Effective population size
$10^9$
$10^7$
$10^5$
$10^4$
$10^5$
$10^6$
$10^7$
Years Ago

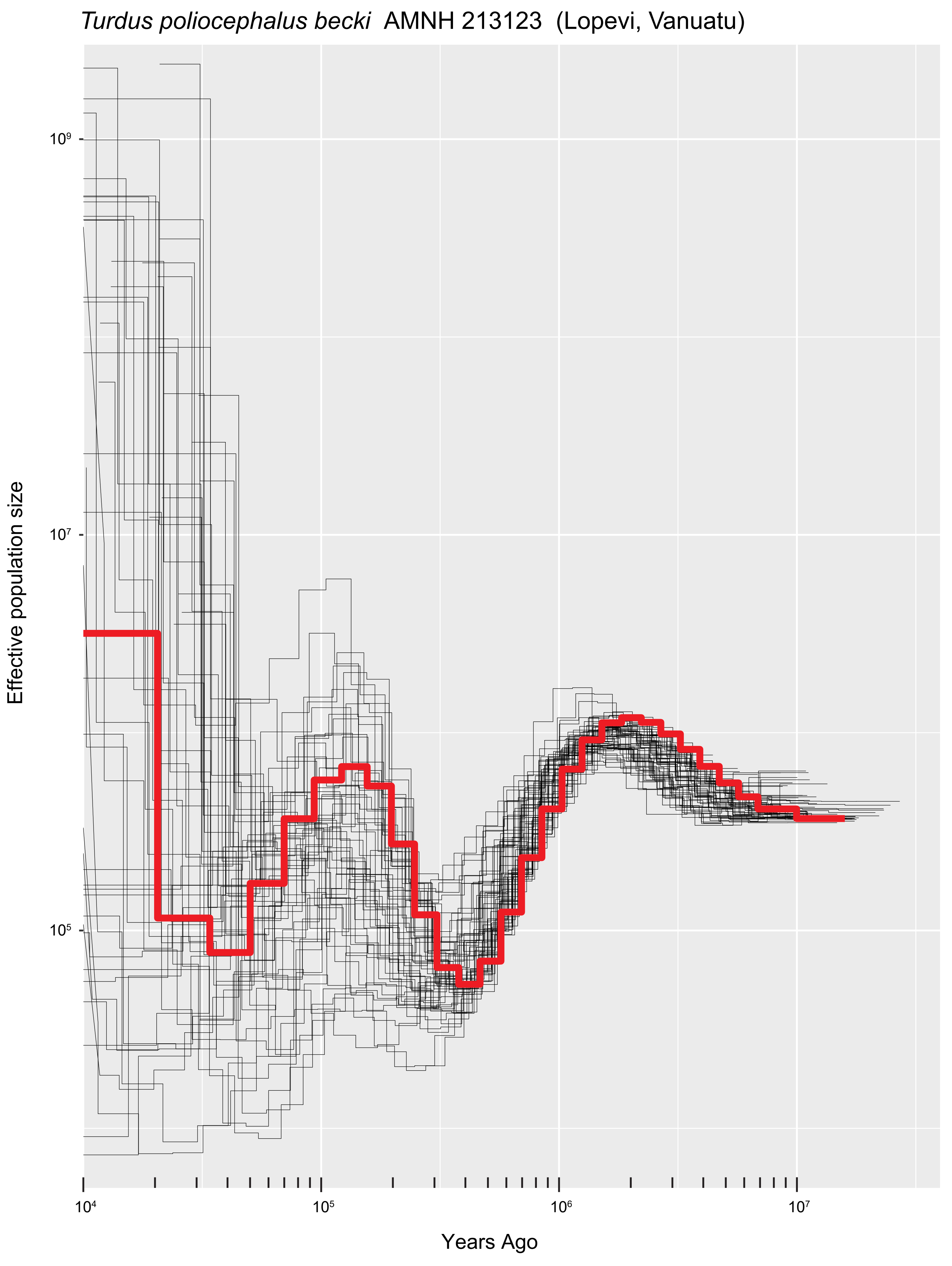

*Turdus poliocephalus becki* AMNH 213123 (Lopevi, Vanuatu)
Effective population size
$10^9$
$10^7$
$10^5$
$10^4$
$10^5$
$10^6$
$10^7$
Years Ago

*Turdus poliocephalus becki* AMNH 213130 (Epi, Vanuatu)

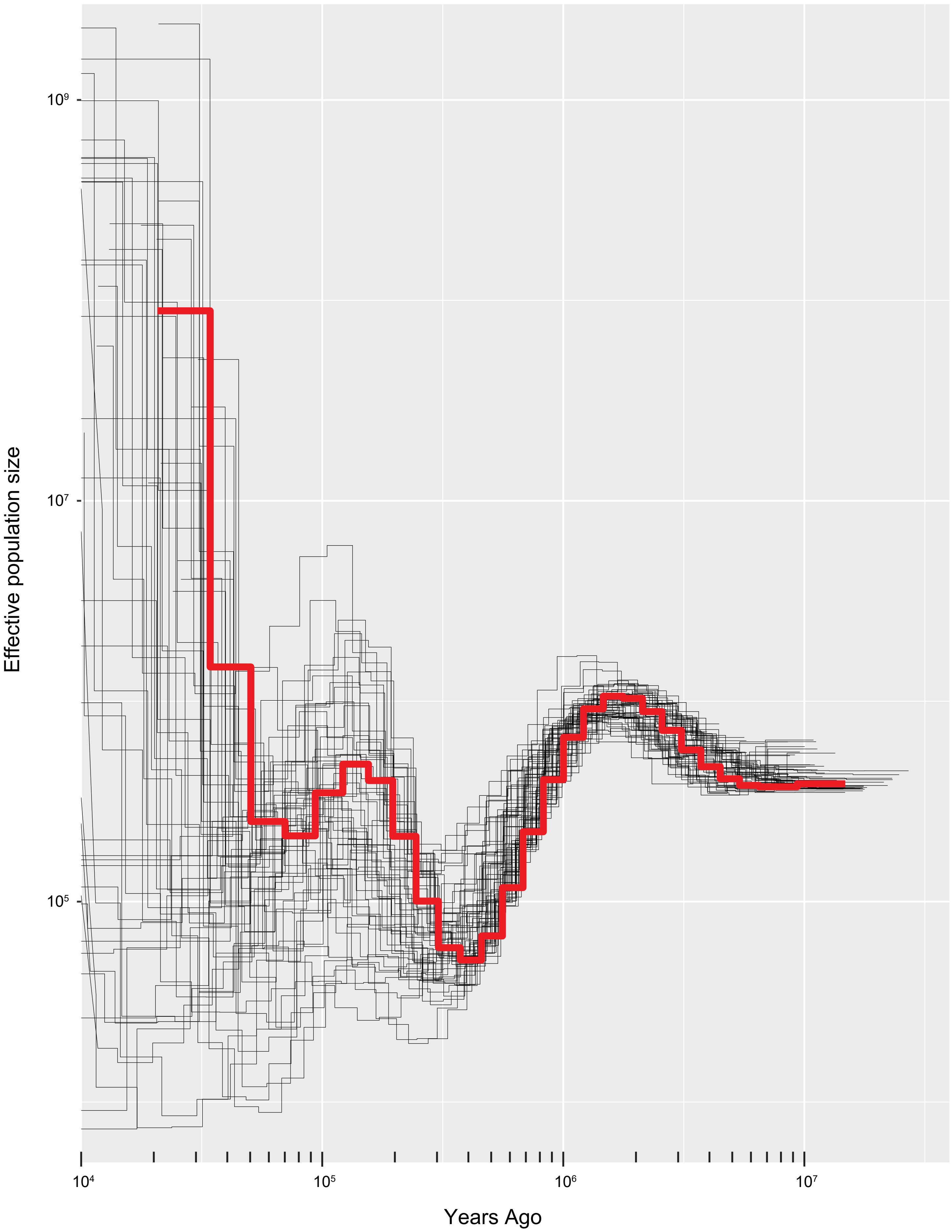

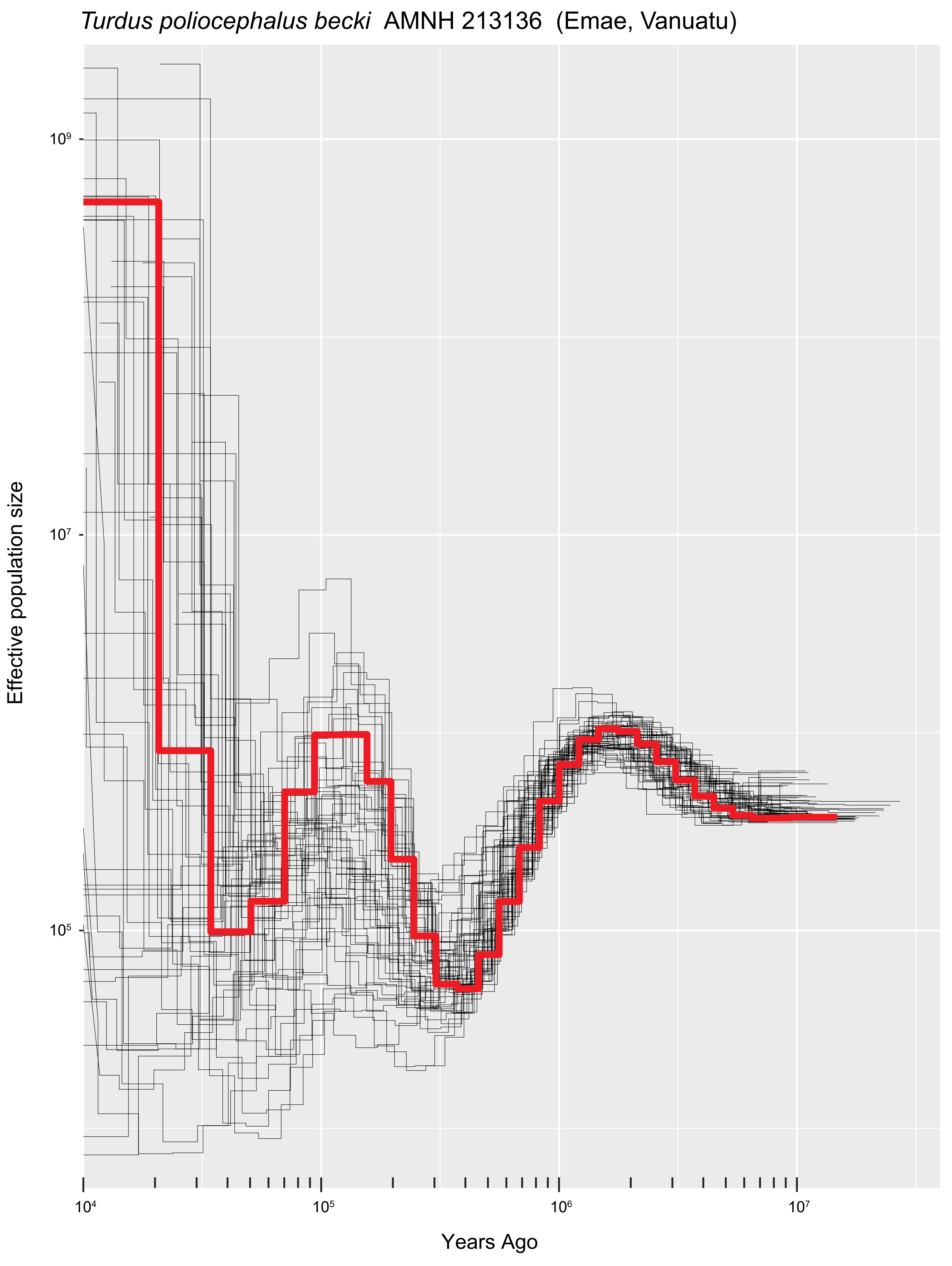

*Turdus poliocephalus becki* AMNH 213136 (Emae, Vanuatu)
Effective population size
$10^9$
$10^7$
$10^5$
$10^4$
$10^5$
$10^6$
$10^7$
Years Ago

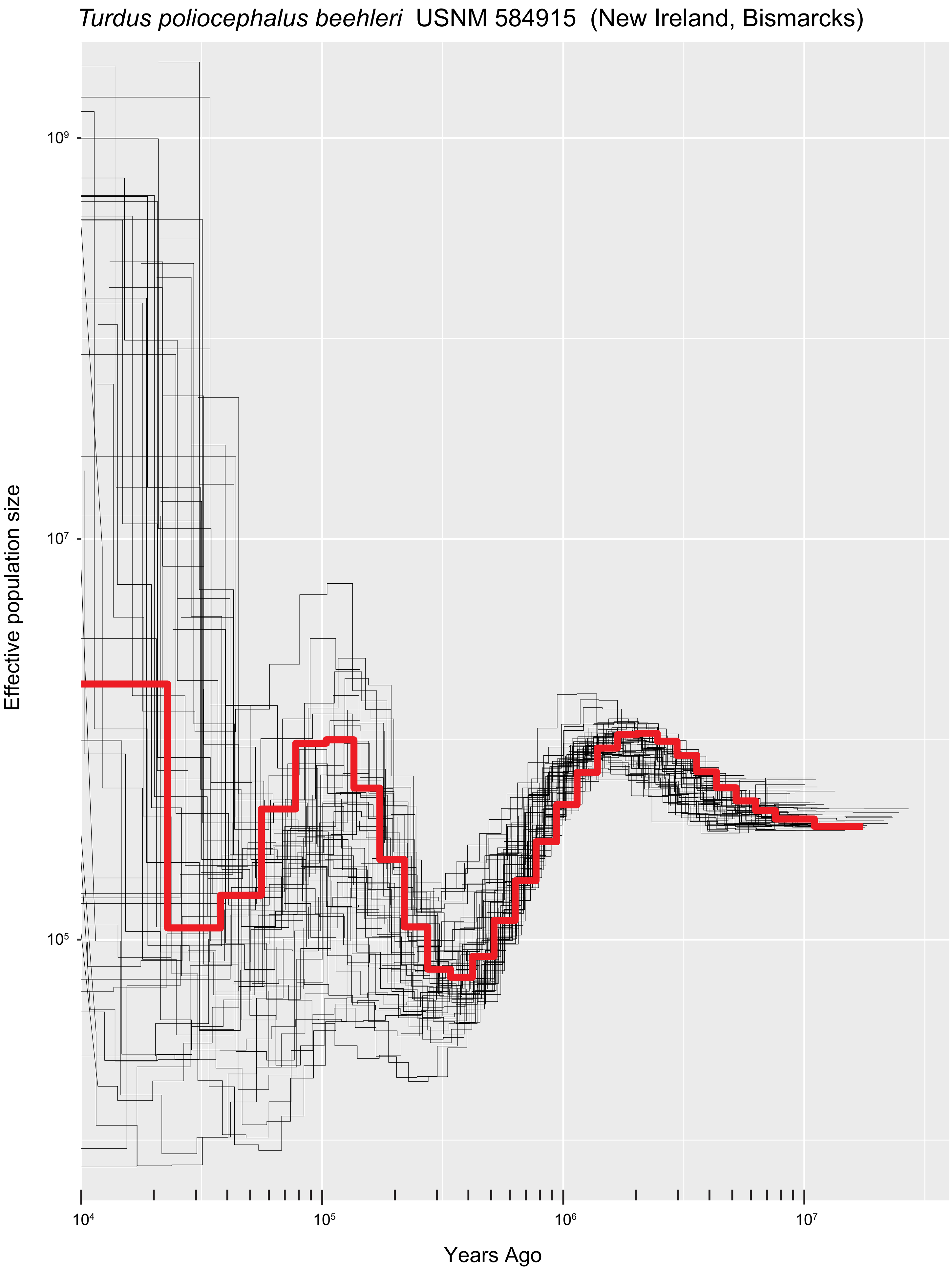

*Turdus poliocephalus beehleri* USNM 584915 (New Ireland, Bismarcks)
Effective population size
$10^9$
$10^7$
$10^5$
$10^4$
$10^5$
$10^6$
$10^7$
Years Ago

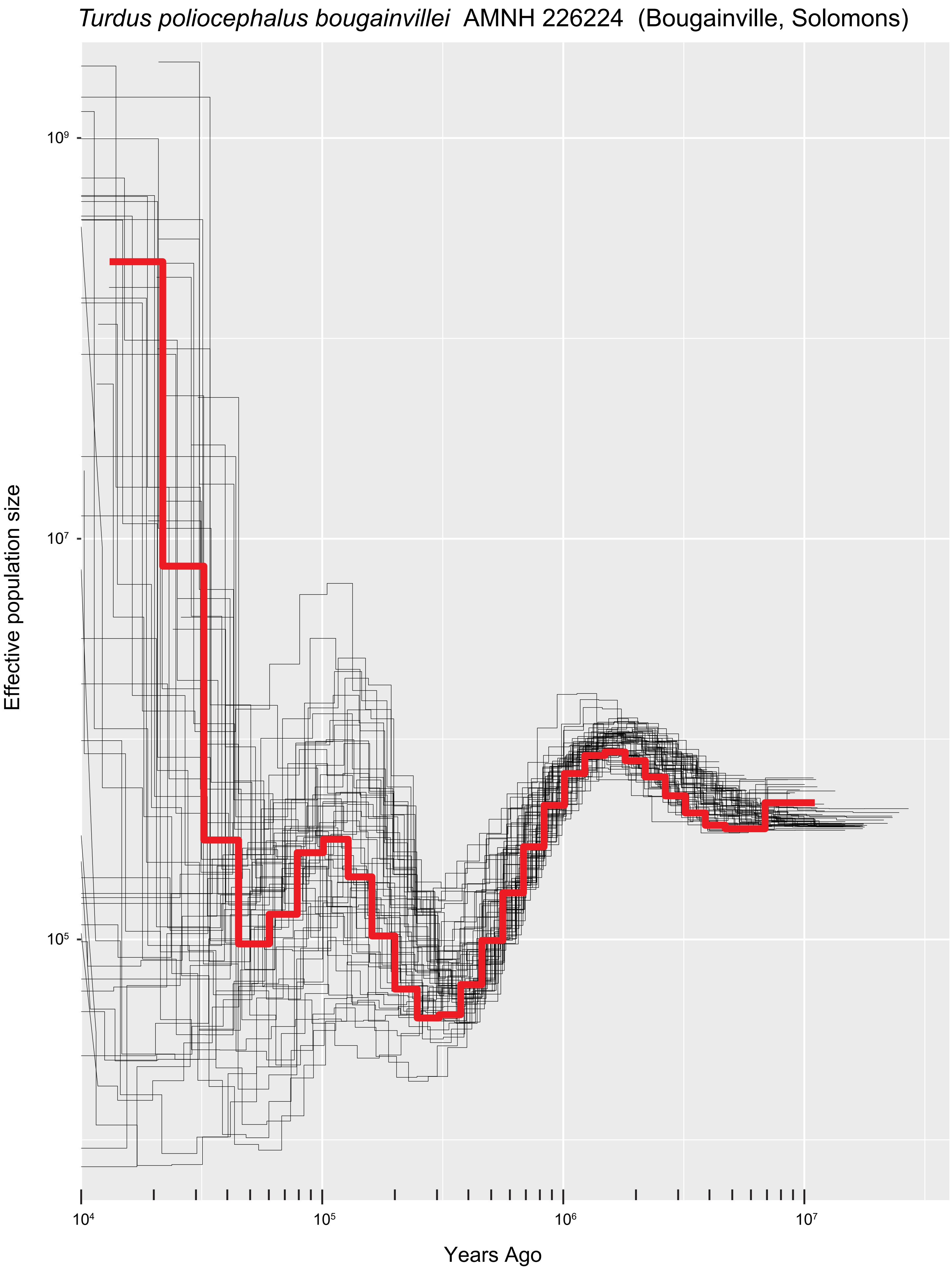
*Turdus poliocephalus bougainvillei* AMNH 226224 (Bougainville, Solomons)
Effective population size
$10^9$
$10^7$
$10^5$
$10^4$
$10^5$
$10^6$
$10^7$
Years Ago

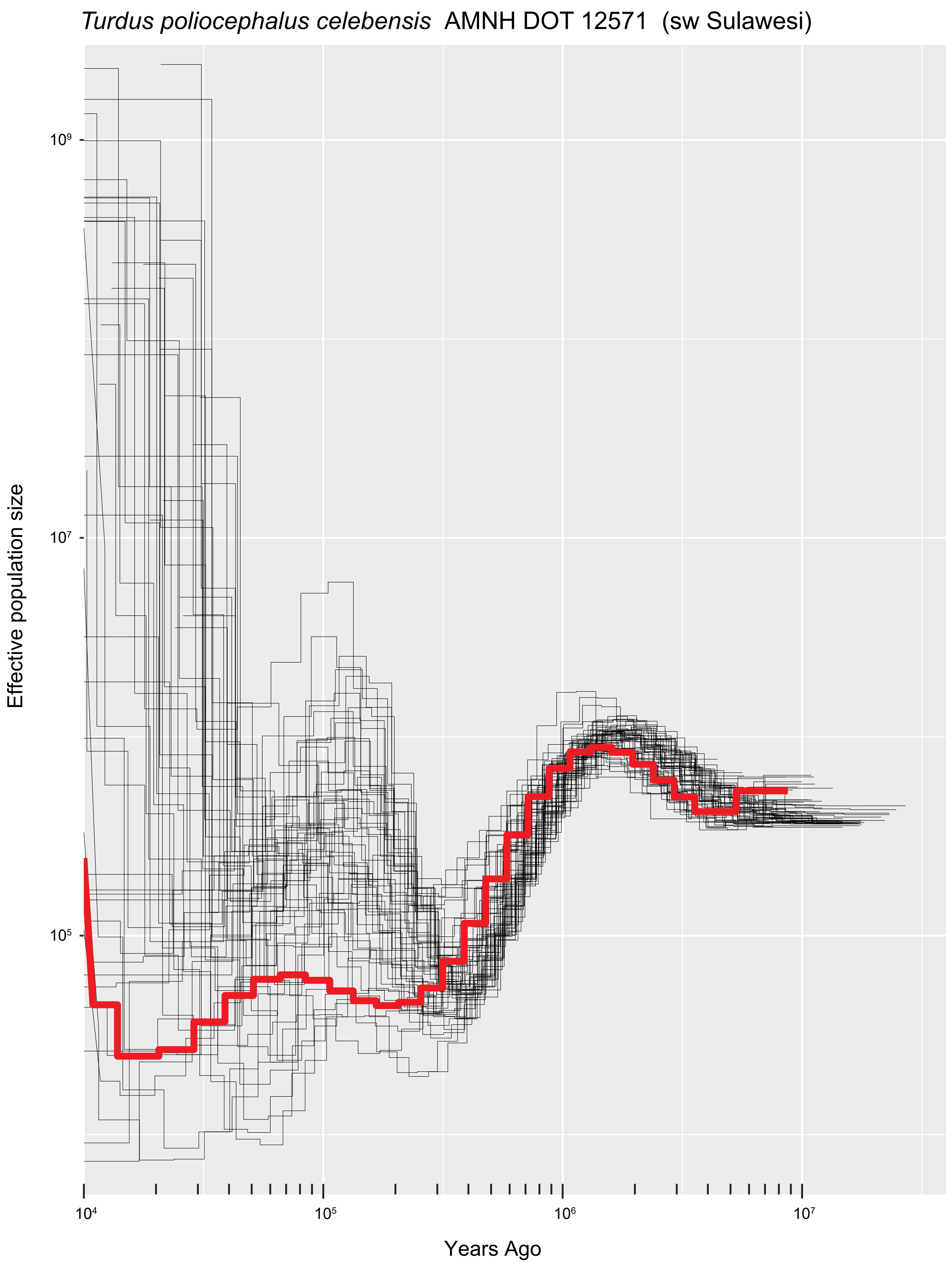
*Turdus poliocephalus celebensis* AMNH DOT 12571 (sw Sulawesi)
Effective population size
$10^9$
$10^7$
$10^5$
$10^4$
$10^5$
$10^6$
$10^7$
Years Ago

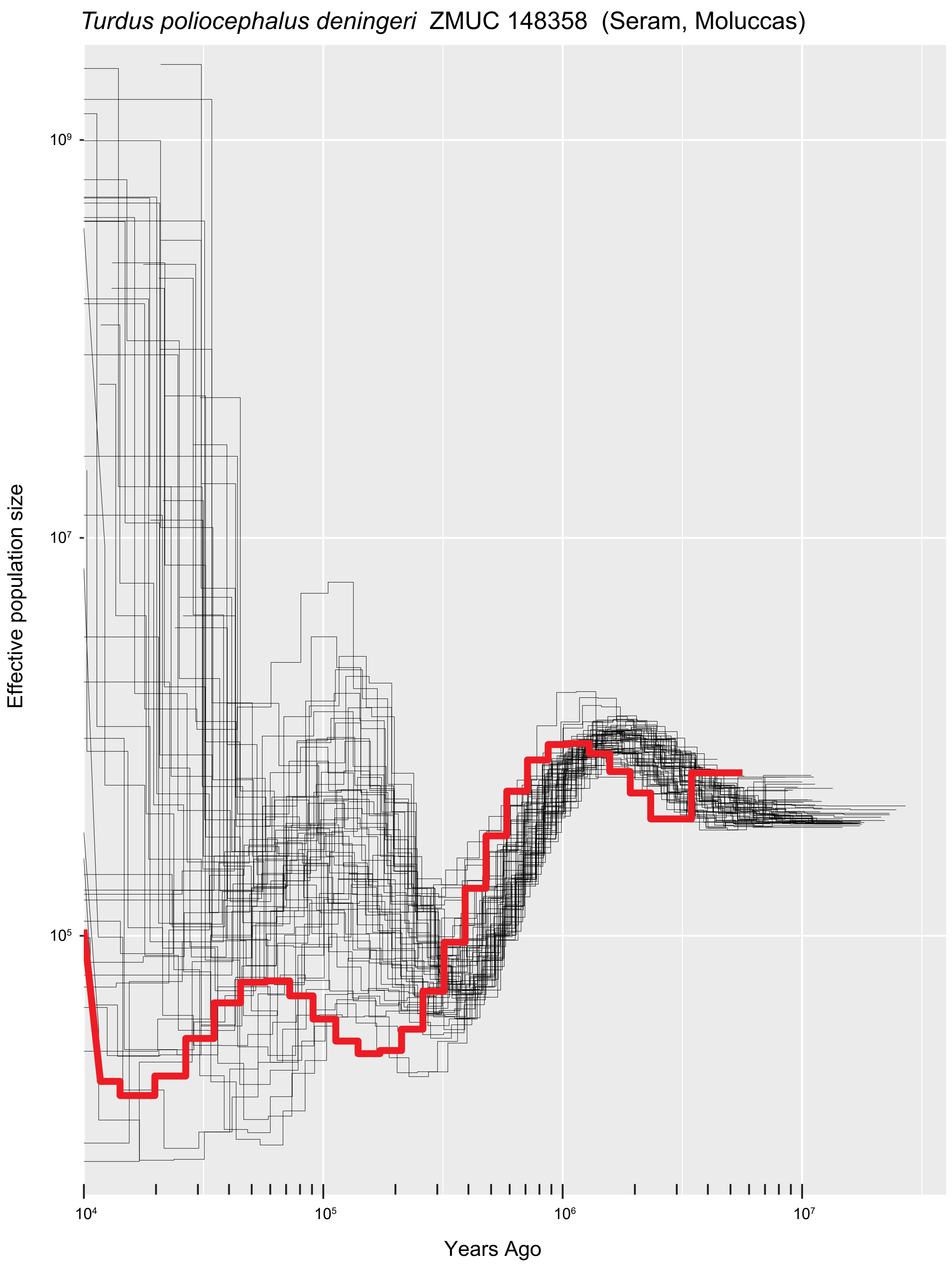
*Turdus poliocephalus deningeri* ZMUC 148358 (Seram, Moluccas)
Effective population size
$10^9$
$10^7$
$10^5$
$10^4$
$10^5$
$10^6$
$10^7$
Years Ago

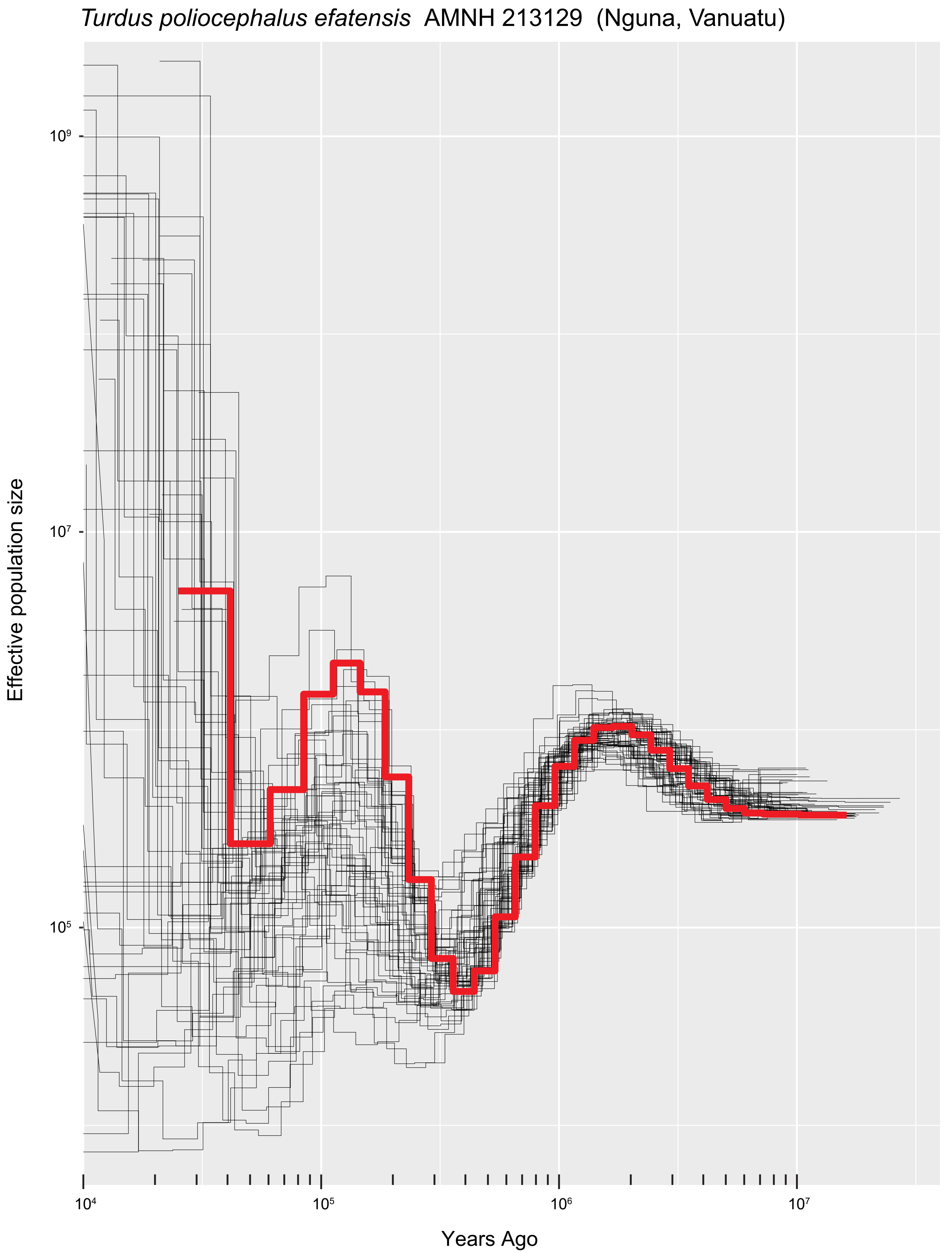
*Turdus poliocephalus efatensis* AMNH 213129 (Nguna, Vanuatu)
Effective population size
10⁹
10⁷
10⁵
10⁴
10⁵
10⁶
10⁷
Years Ago

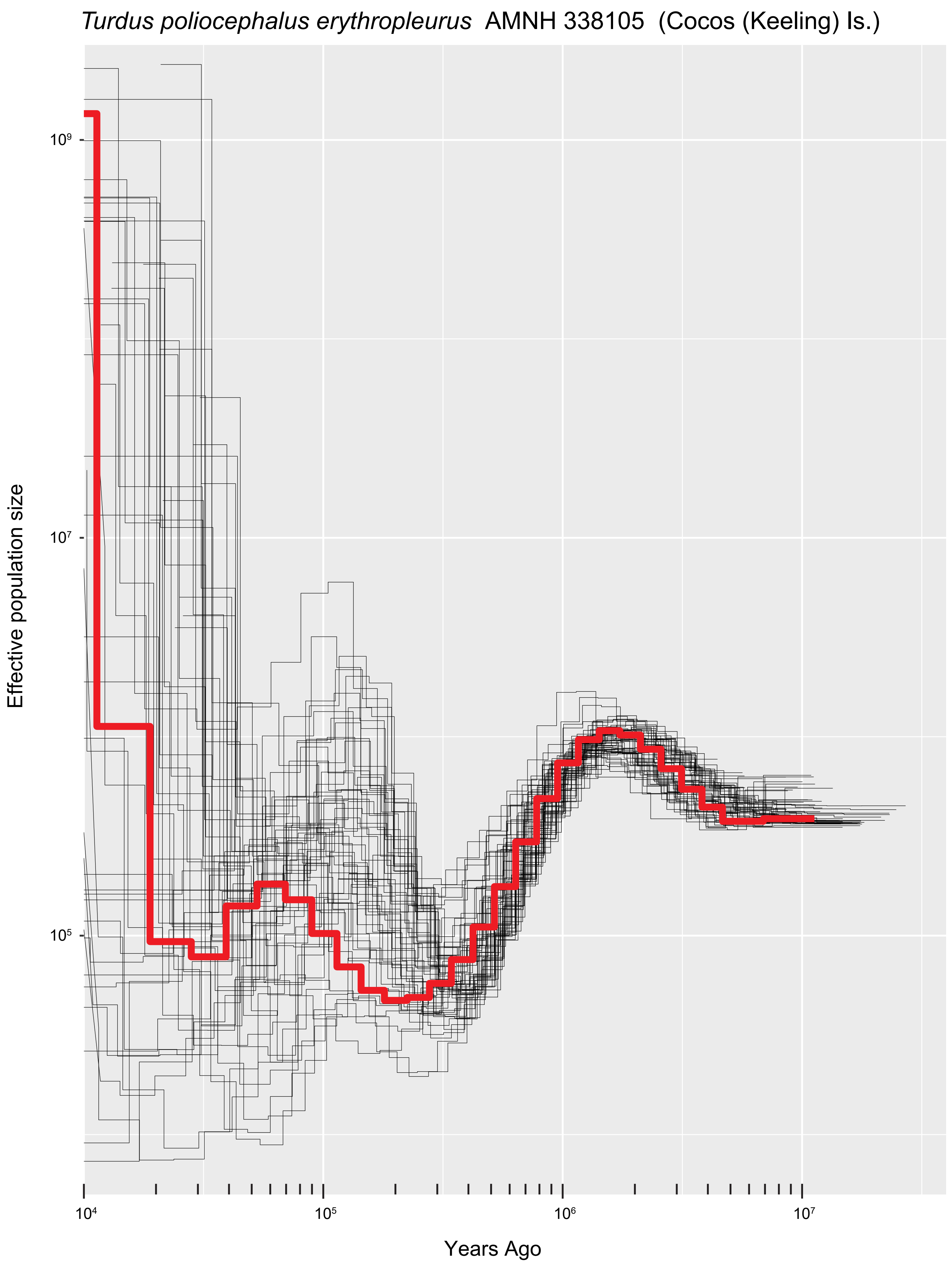
*Turdus poliocephalus erythropleurus* AMNH 338105 (Cocos (Keeling) Is.)
Effective population size
$10^9$
$10^7$
$10^5$
$10^4$
$10^5$
$10^6$
$10^7$
Years Ago

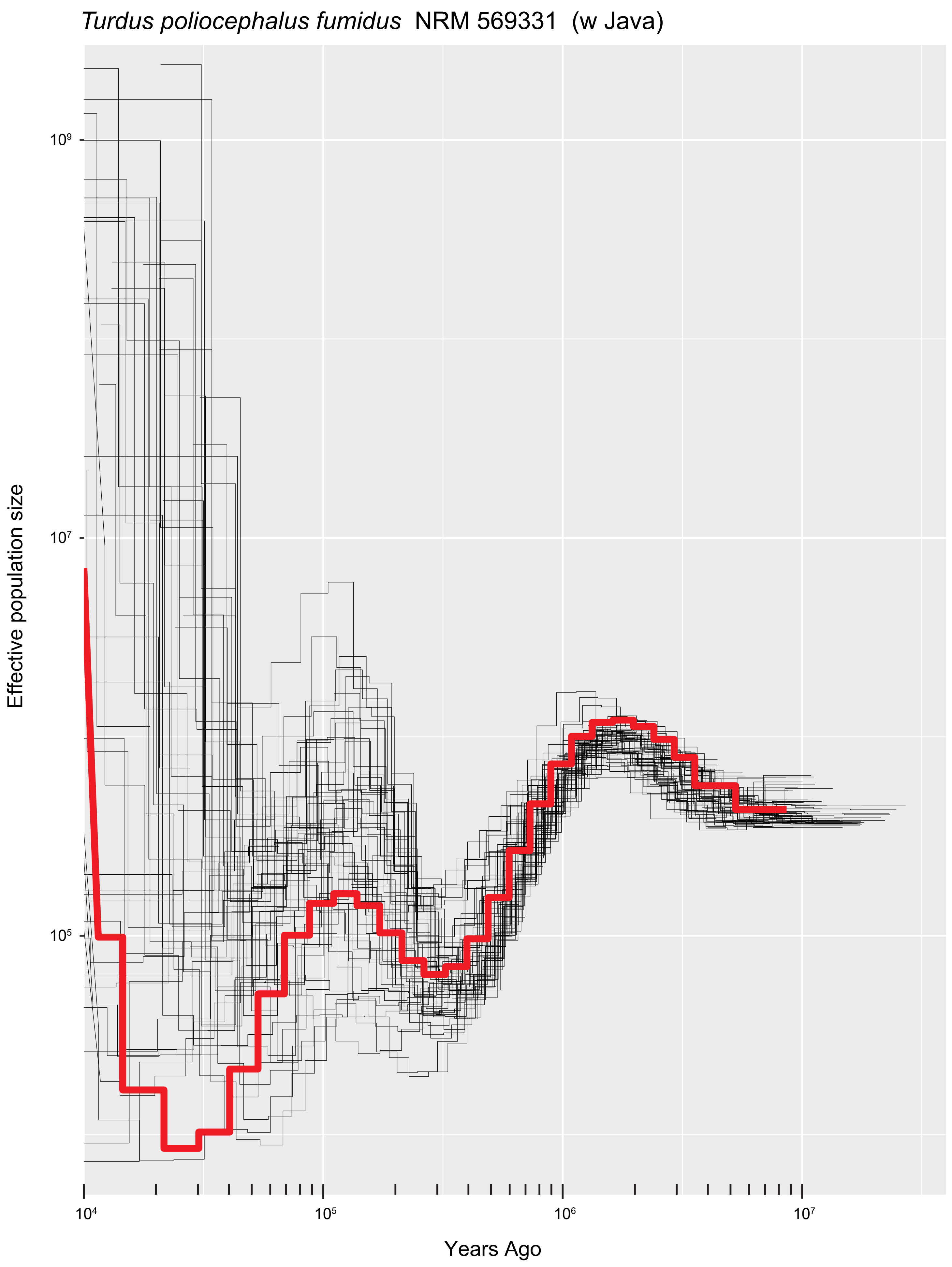

*Turdus poliocephalus fumidus* NRM 569331 (w Java)
Effective population size
10⁹
10⁷
10⁵
10⁴
10⁵
10⁶
10⁷
Years Ago

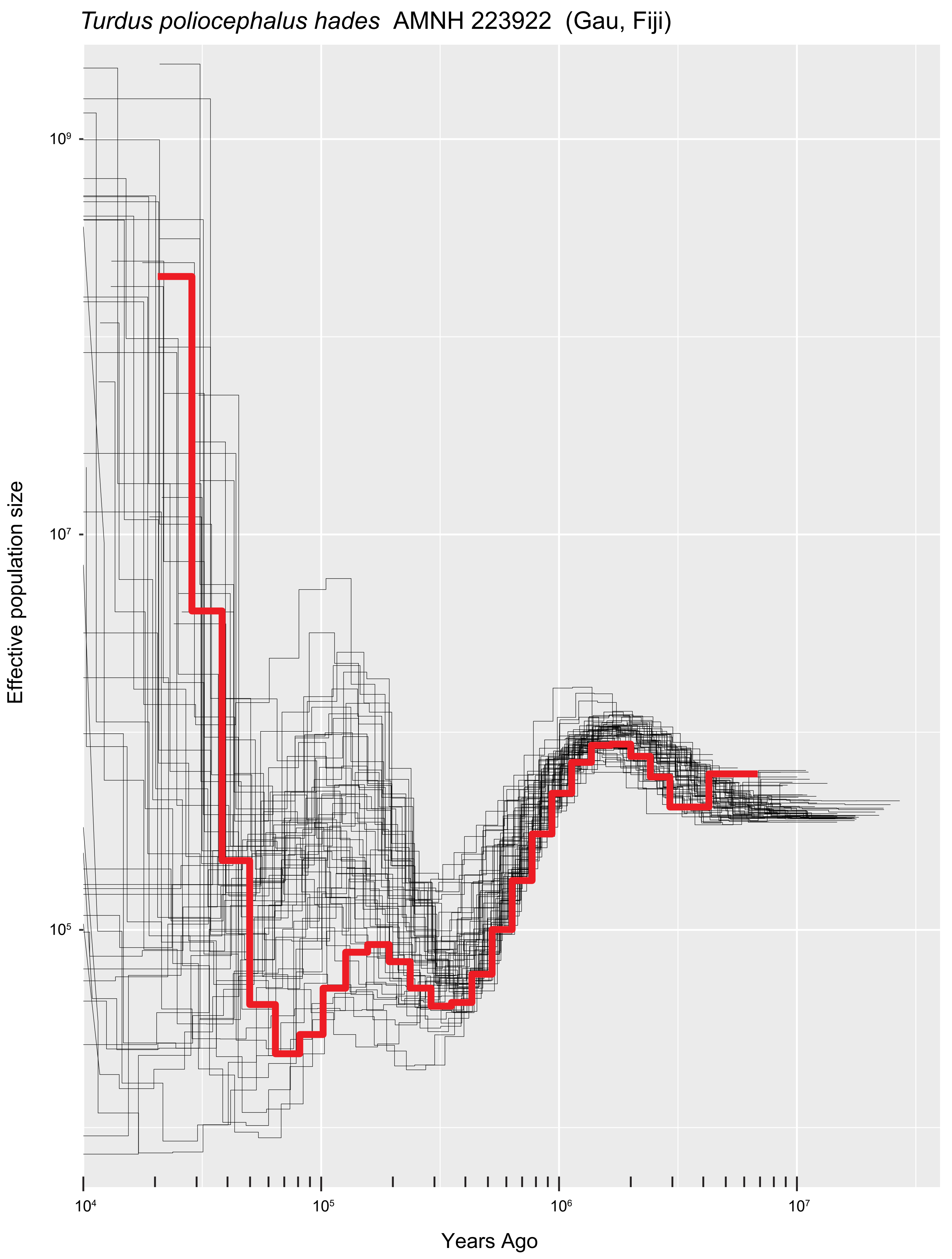
Turdus poliocephalus hades AMNH 223922 (Gau, Fiji)
Effective population size
10⁹
10⁷
10⁵
Years Ago
10⁴
10⁵
10⁶
10⁷

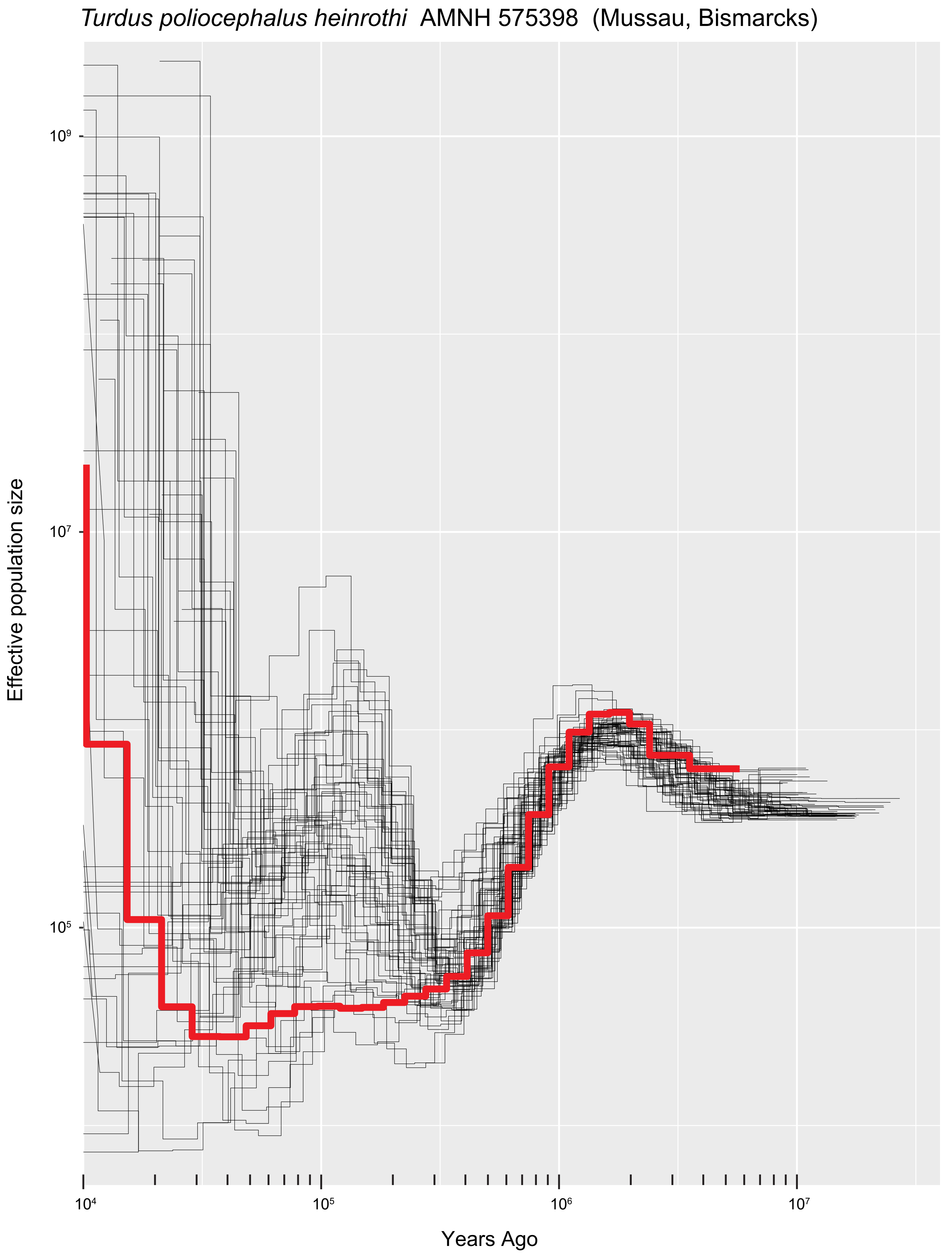
*Turdus poliocephalus heinrothi* AMNH 575398 (Mussau, Bismarcks)
Effective population size
10⁹
10⁷
10⁵
10⁴
10⁵
10⁶
10⁷
Years Ago

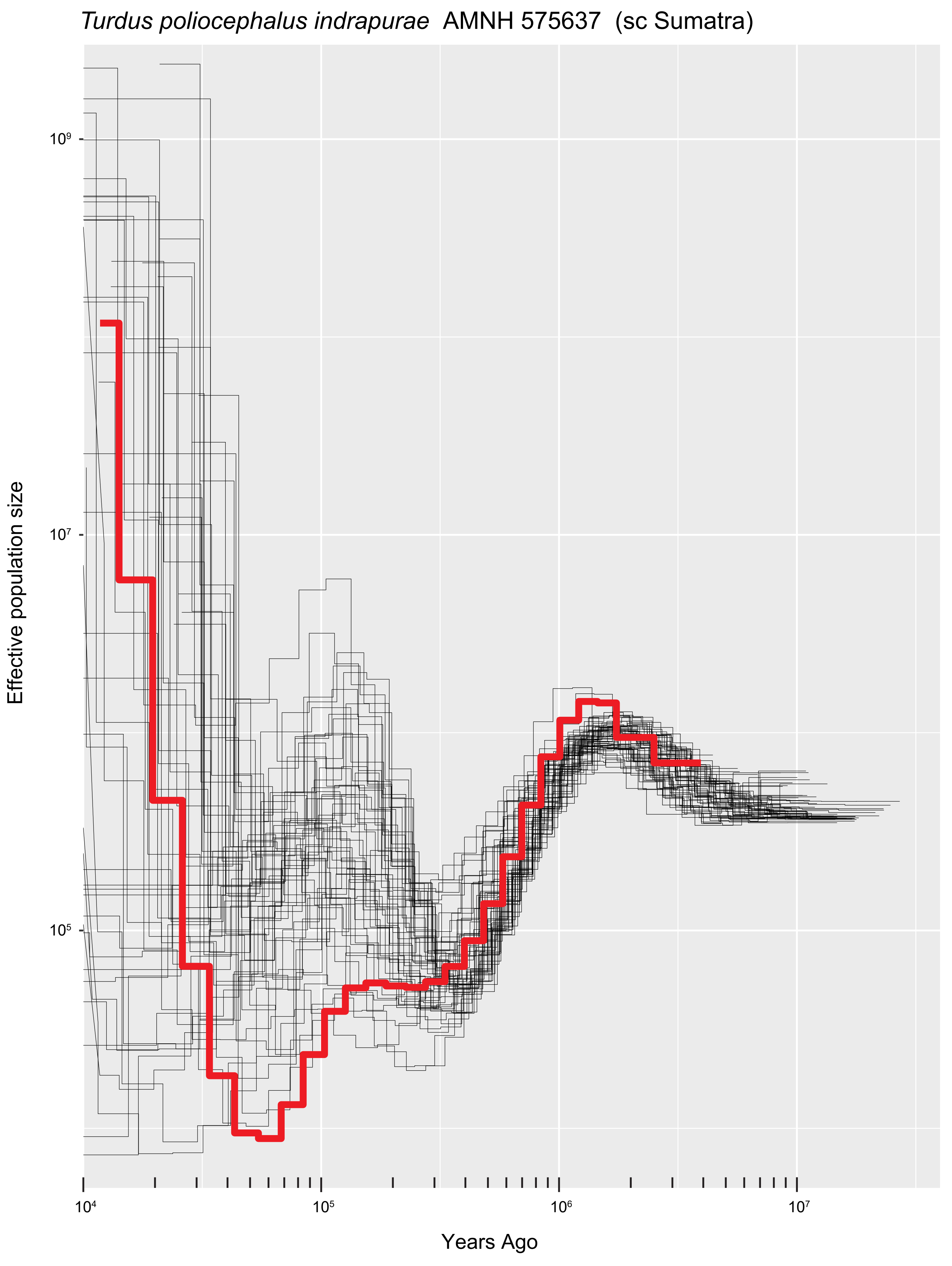
*Turdus poliocephalus indrapurae* AMNH 575637 (sc Sumatra)
Effective population size
$10^9$
$10^7$
$10^5$
$10^4$
$10^5$
$10^6$
$10^7$
Years Ago

*Turdus poliocephalus javanicus* AMNH 575611 (c Java)

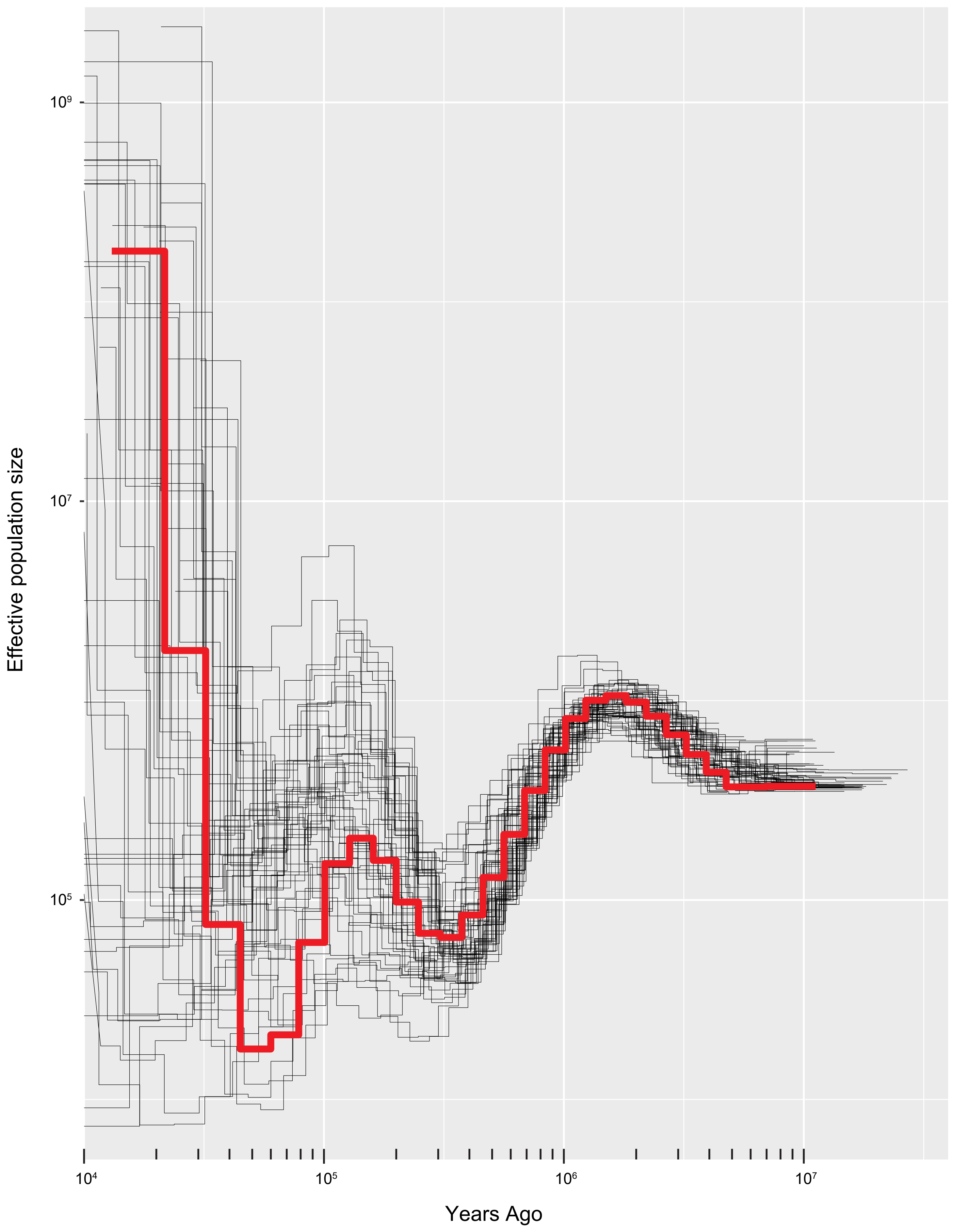

*Turdus poliocephalus katanglad* NHMD 129066 (c Mindanao, Philippines)

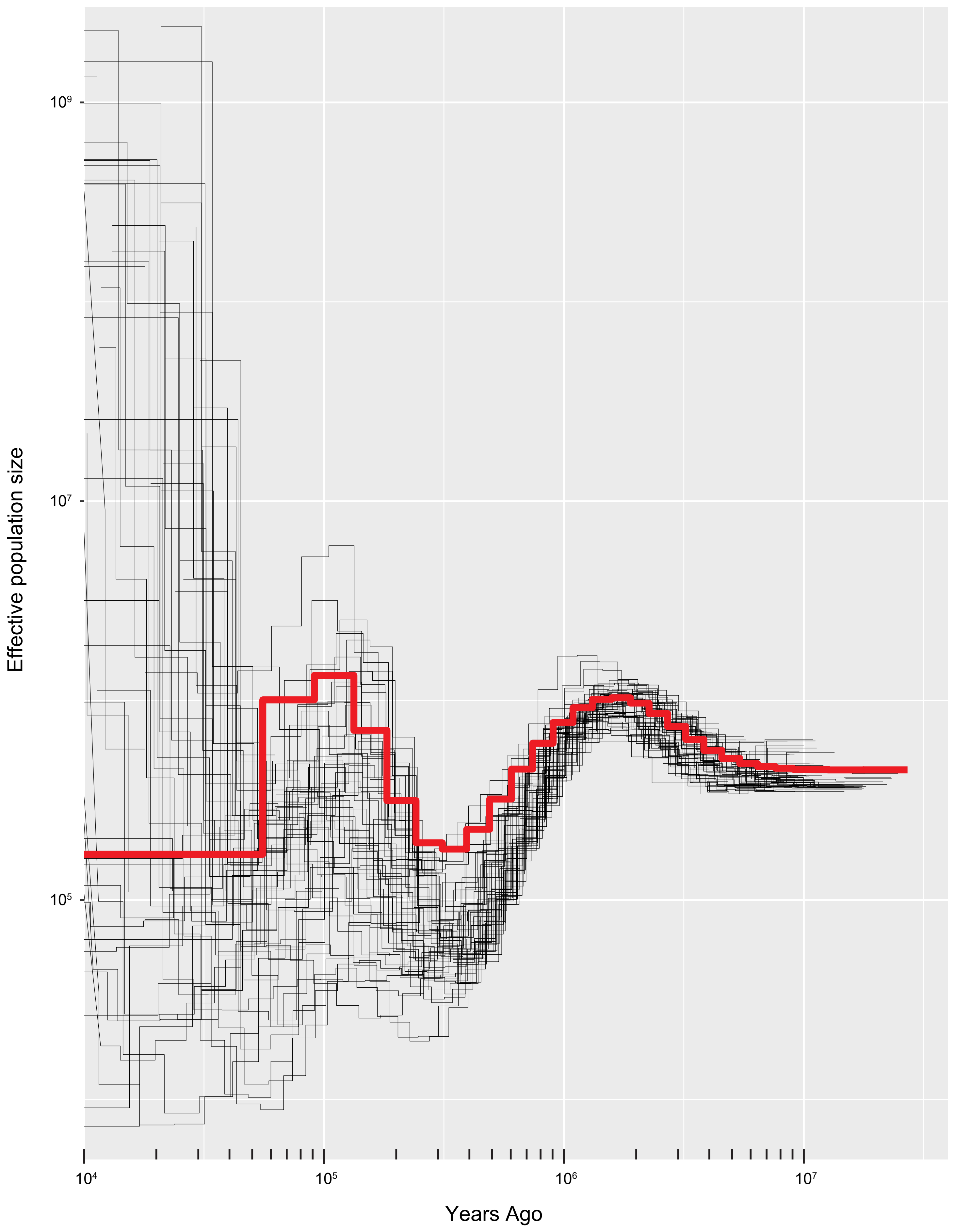

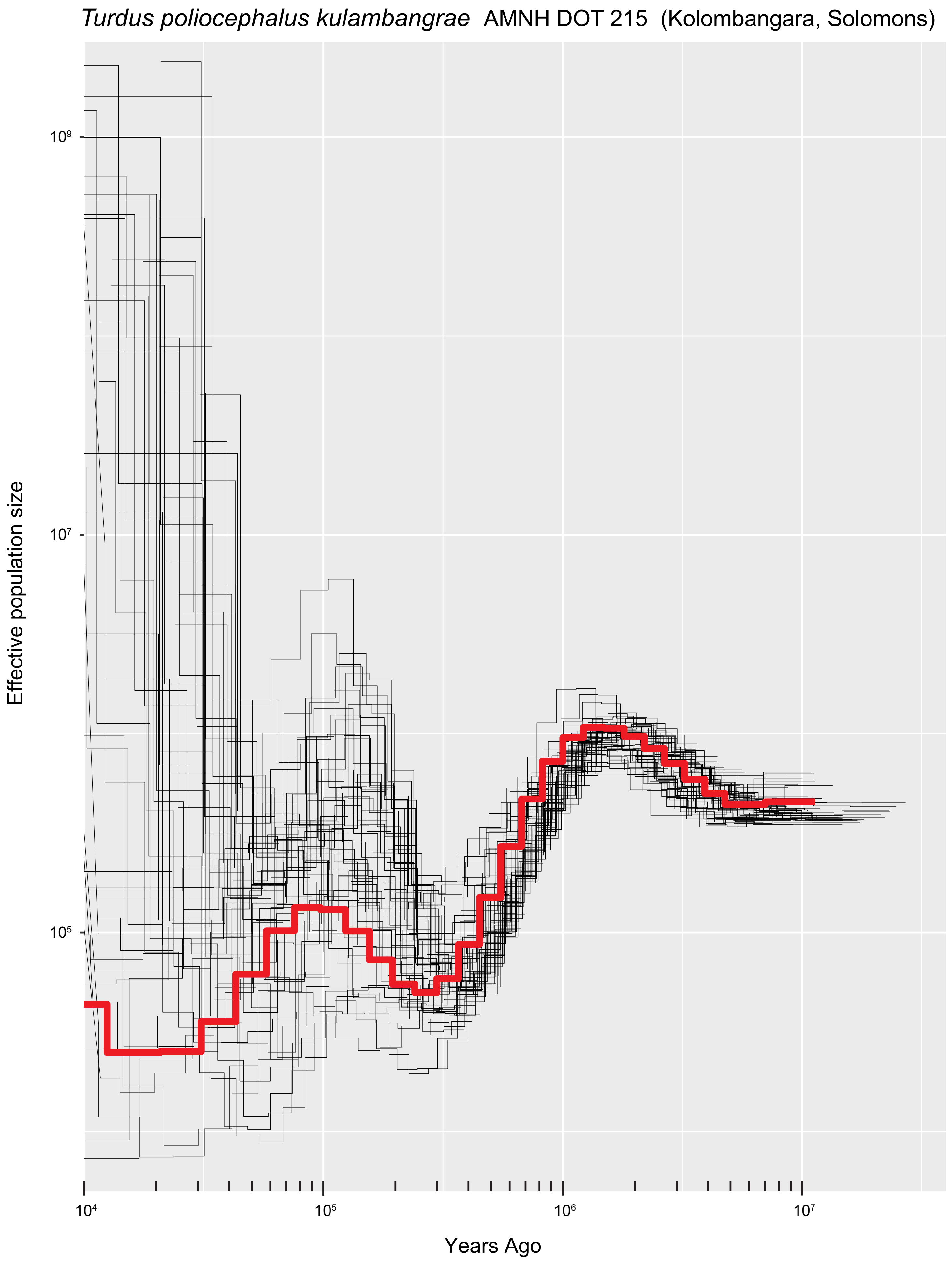

Turdus poliocephalus kulambangrae  AMNH DOT 215  (Kolombangara, Solomons)
Effective population size
$10^9$
$10^7$
$10^5$
$10^4$
$10^5$
$10^6$
$10^7$
Years Ago

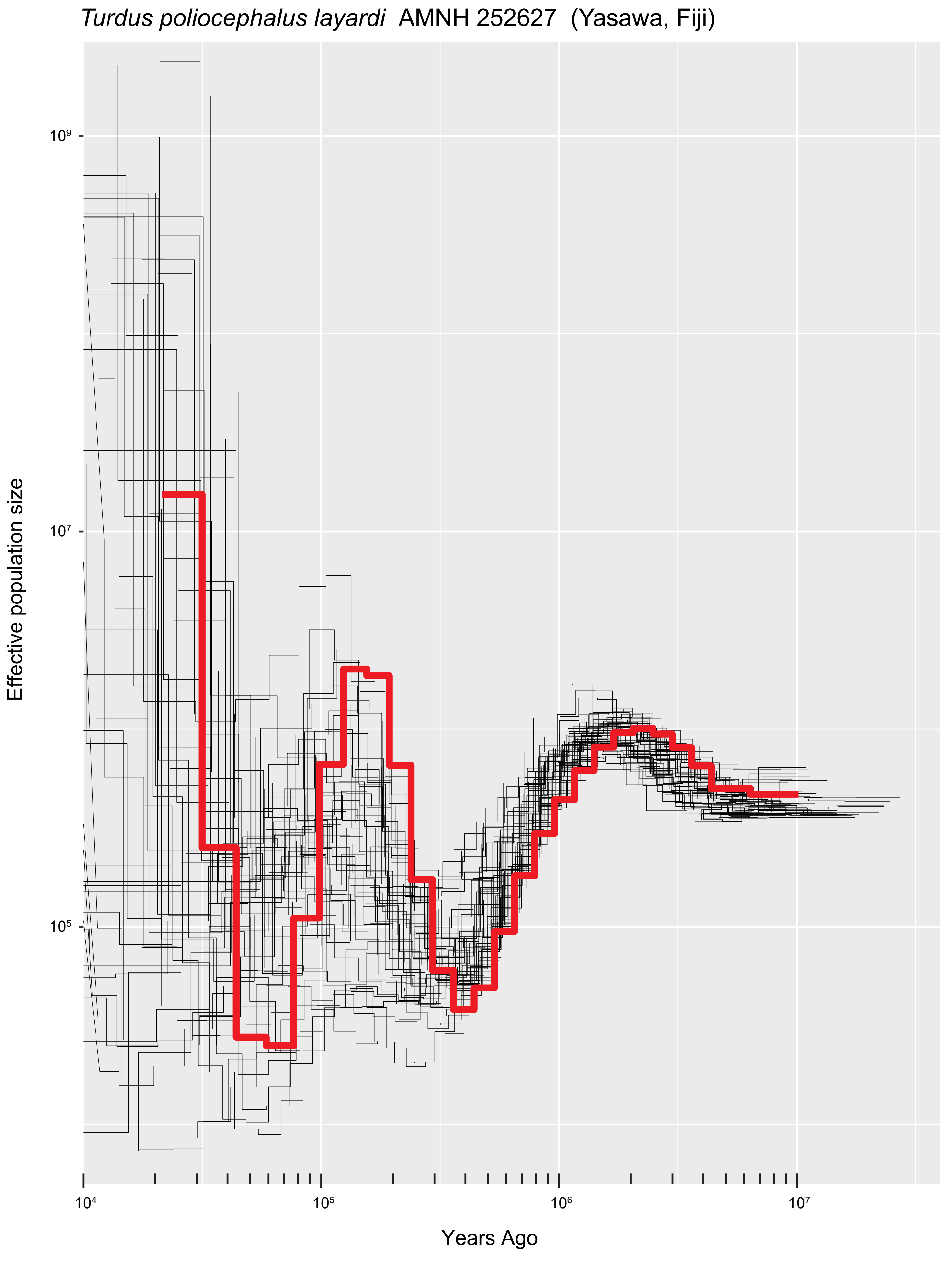

*Turdus poliocephalus layardi* AMNH 252627 (Yasawa, Fiji)
Effective population size
10^9
10^7
10^5
10^4
10^5
10^6
10^7
Years Ago

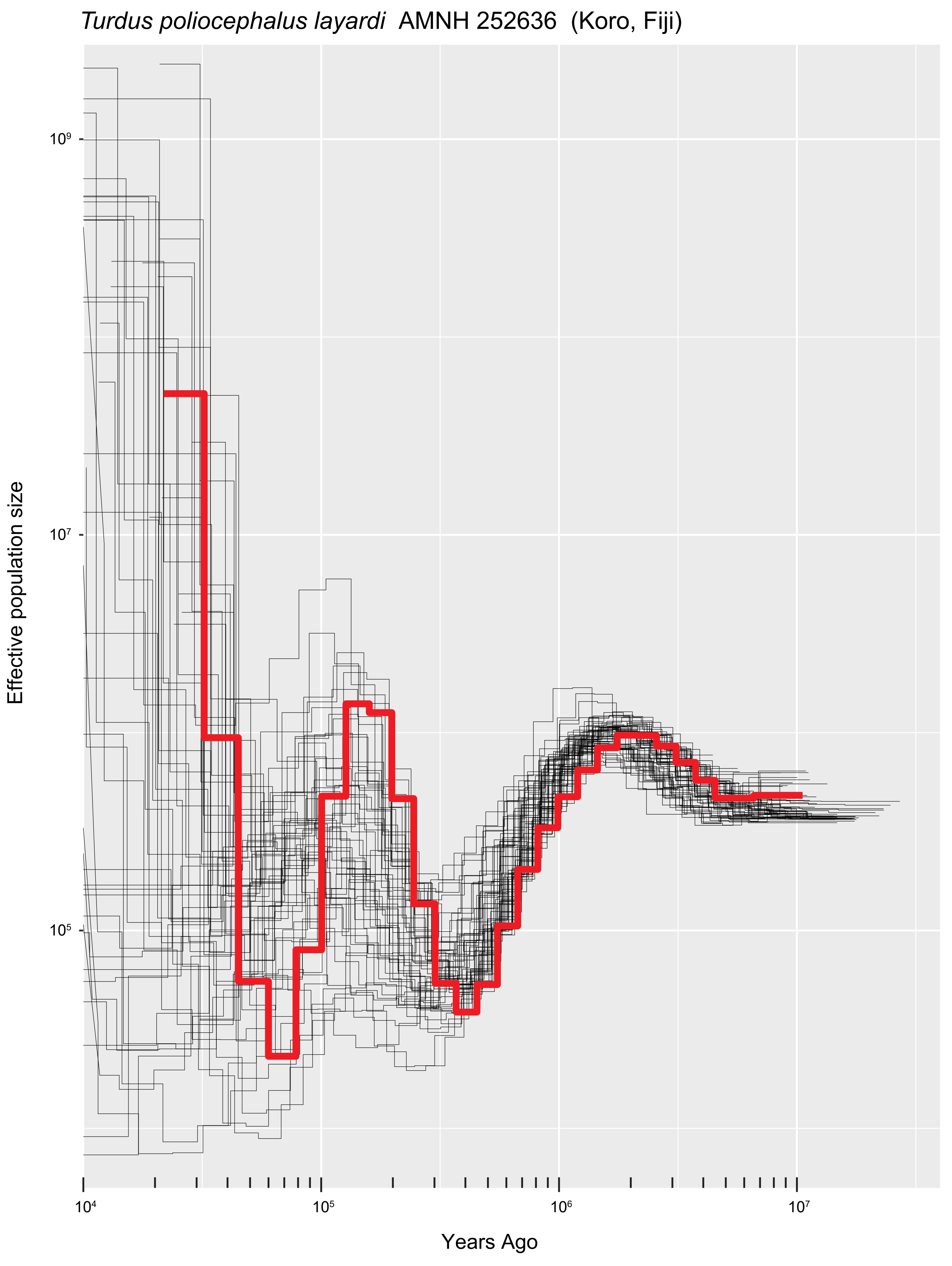
*Turdus poliocephalus layardi* AMNH 252636 (Koro, Fiji)
Effective population size
$10^9$
$10^7$
$10^5$
$10^4$
$10^5$
$10^6$
$10^7$
Years Ago

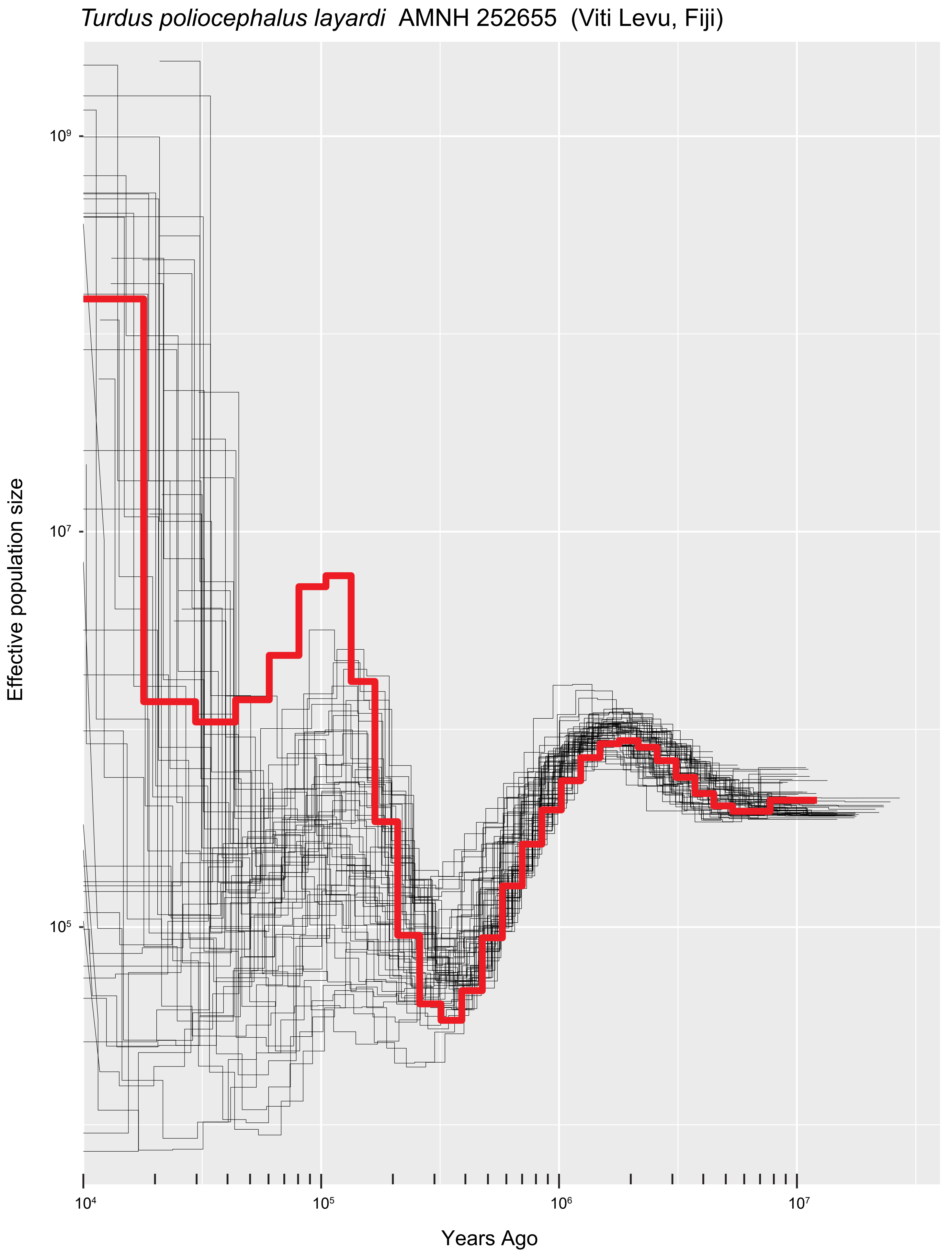

*Turdus poliocephalus layardi*  AMNH 252655  (Viti Levu, Fiji)
Effective population size
$10^9$
$10^7$
$10^5$
$10^4$
$10^5$
$10^6$
$10^7$
Years Ago

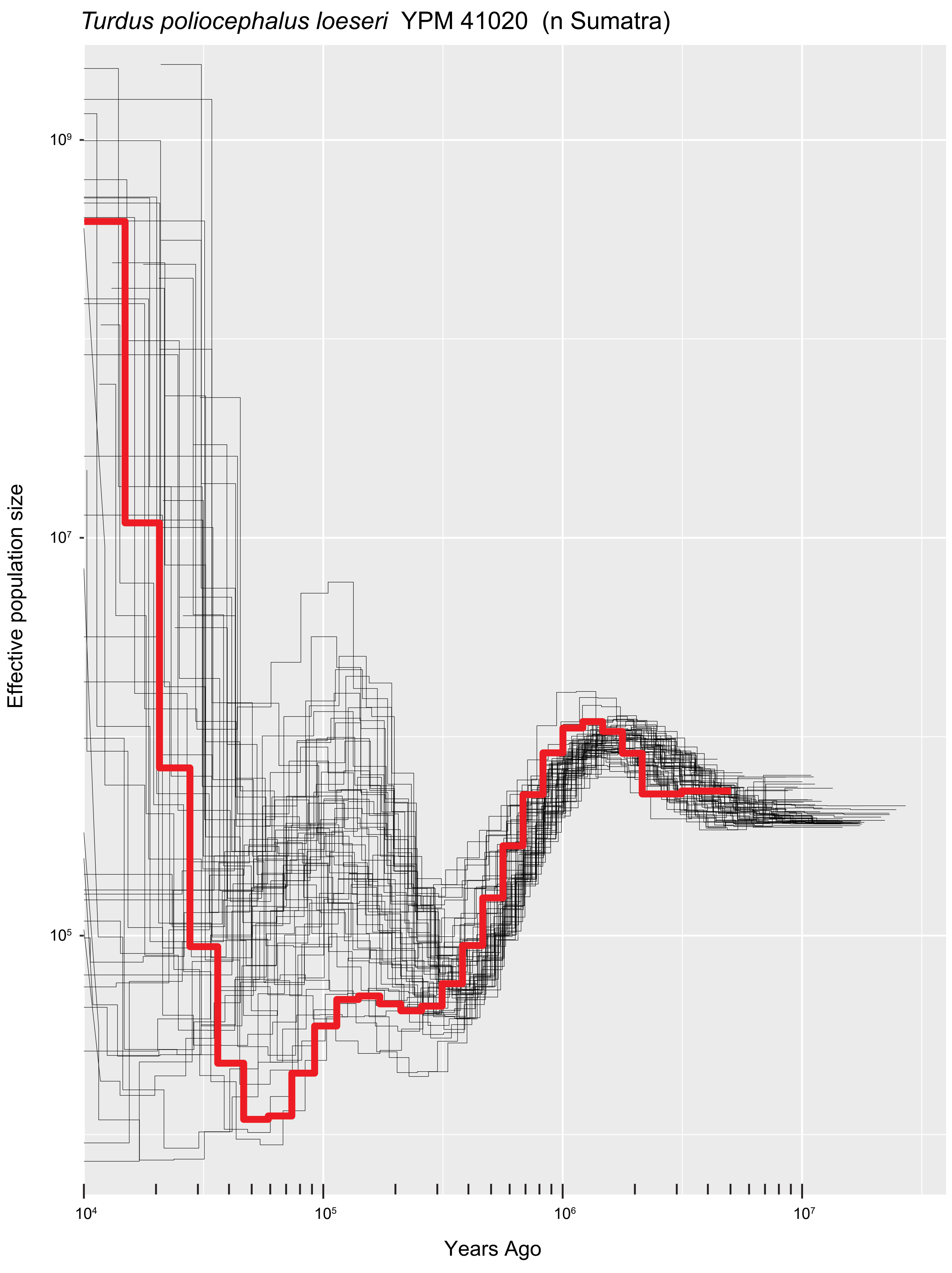

*Turdus poliocephalus loeseri* YPM 41020 (n Sumatra)
Effective population size
$10^9$
$10^7$
$10^5$
$10^4$
$10^5$
$10^6$
$10^7$
Years Ago

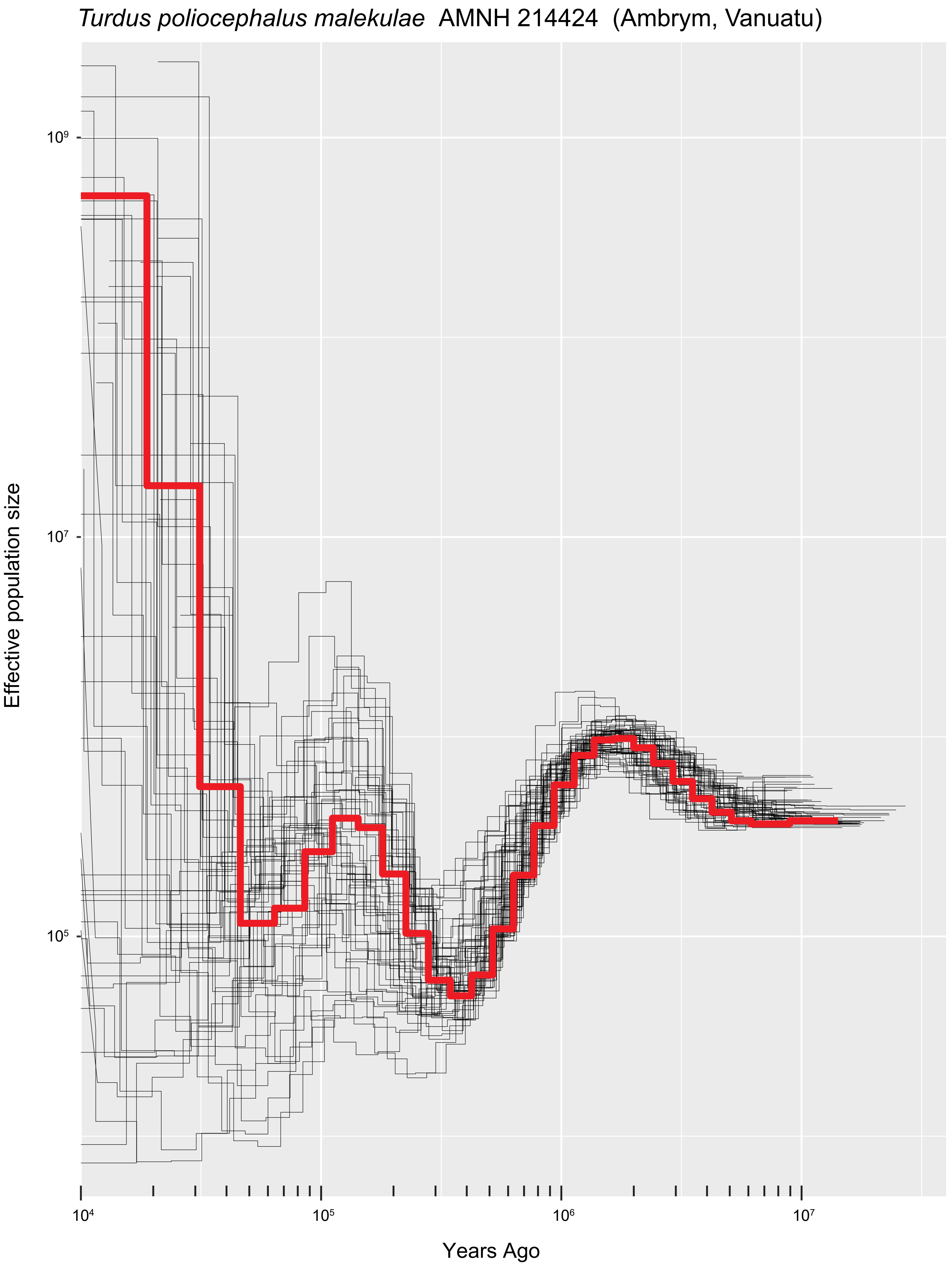
*Turdus poliocephalus malekulae* AMNH 214424 (Ambrym, Vanuatu)
Effective population size
$10^9$
$10^7$
$10^5$
$10^4$
$10^5$
$10^6$
$10^7$
Years Ago

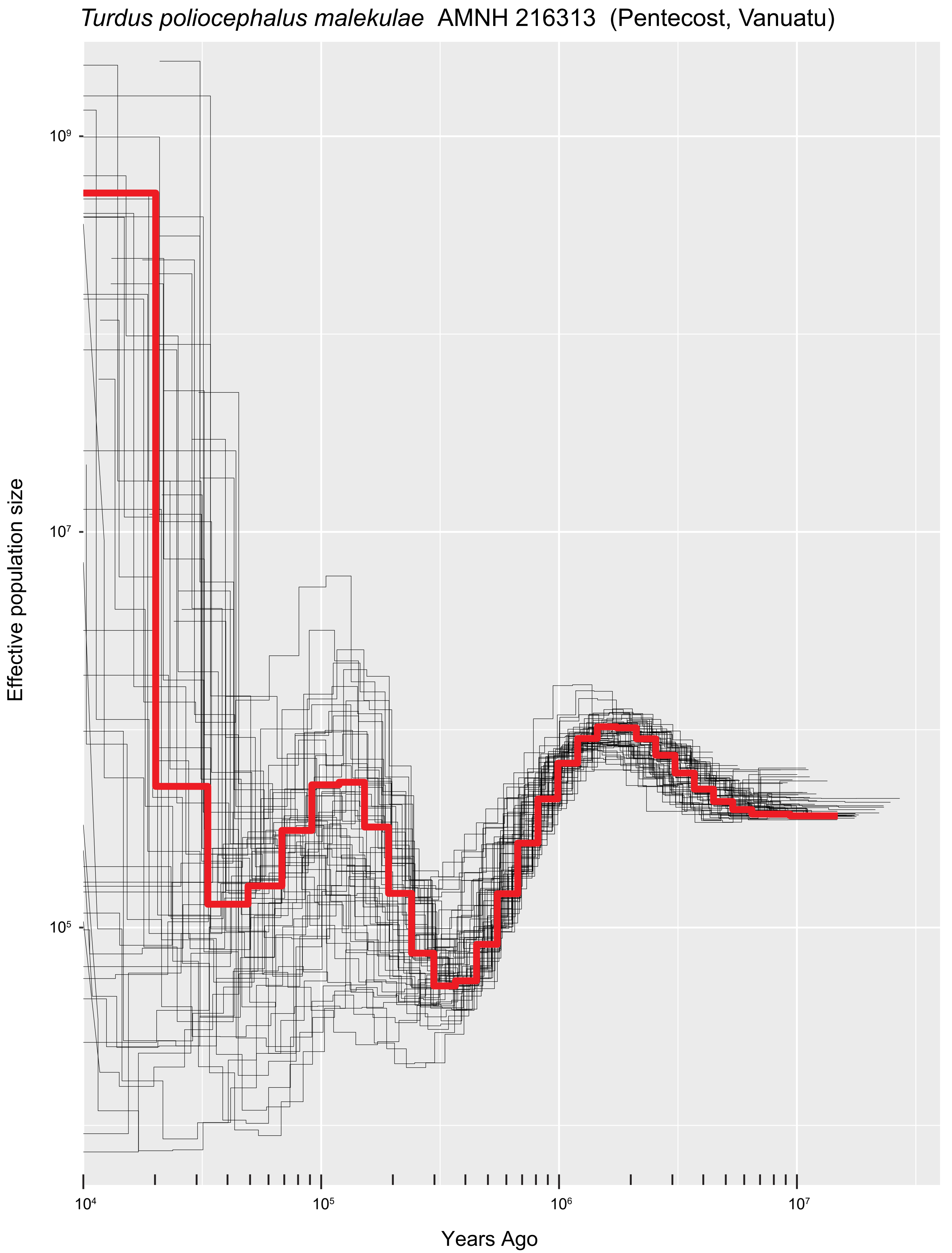

*Turdus poliocephalus malekulae* AMNH 216313 (Pentecost, Vanuatu)
Effective population size
$10^9$
$10^7$
$10^5$
$10^4$
$10^5$
$10^6$
$10^7$
Years Ago

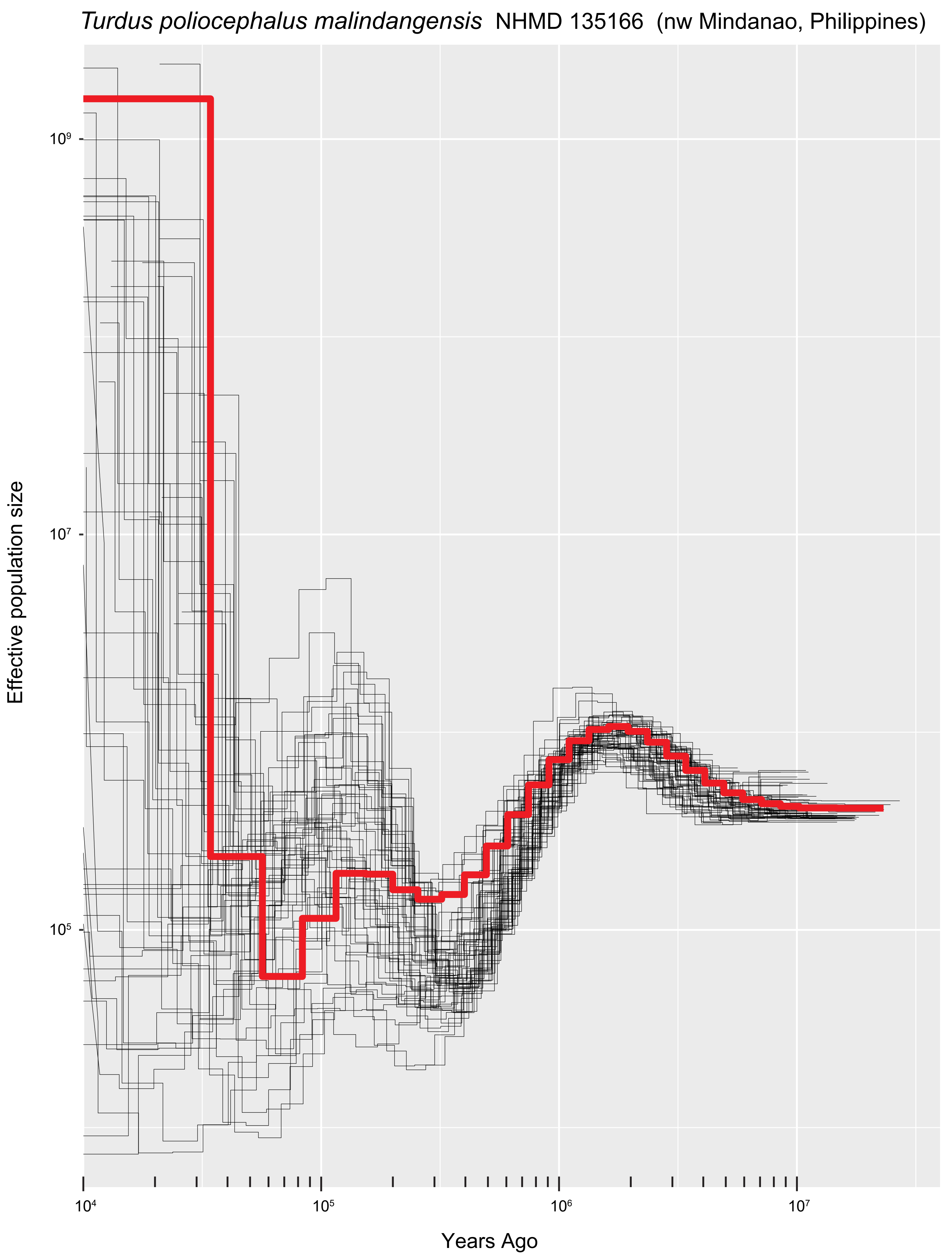
*Turdus poliocephalus malindangensis* NHMD 135166 (nw Mindanao, Philippines)
Effective population size
$10^9$
$10^7$
$10^5$
$10^4$
$10^5$
$10^6$
$10^7$
Years Ago

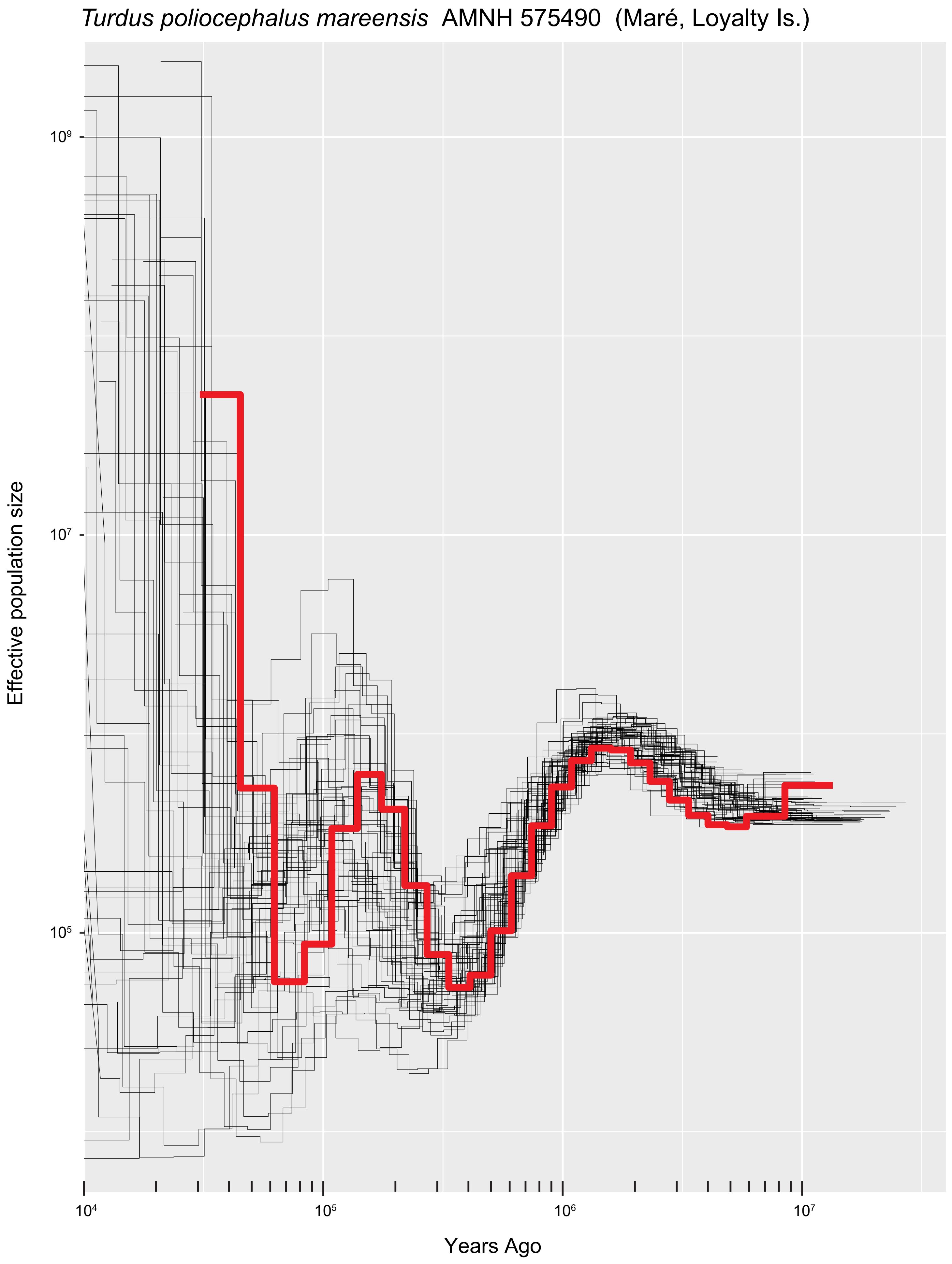
*Turdus poliocephalus mareensis* AMNH 575490 (Maré, Loyalty Is.)
Effective population size
$10^9$
$10^7$
$10^5$
$10^4$
$10^5$
$10^6$
$10^7$
Years Ago

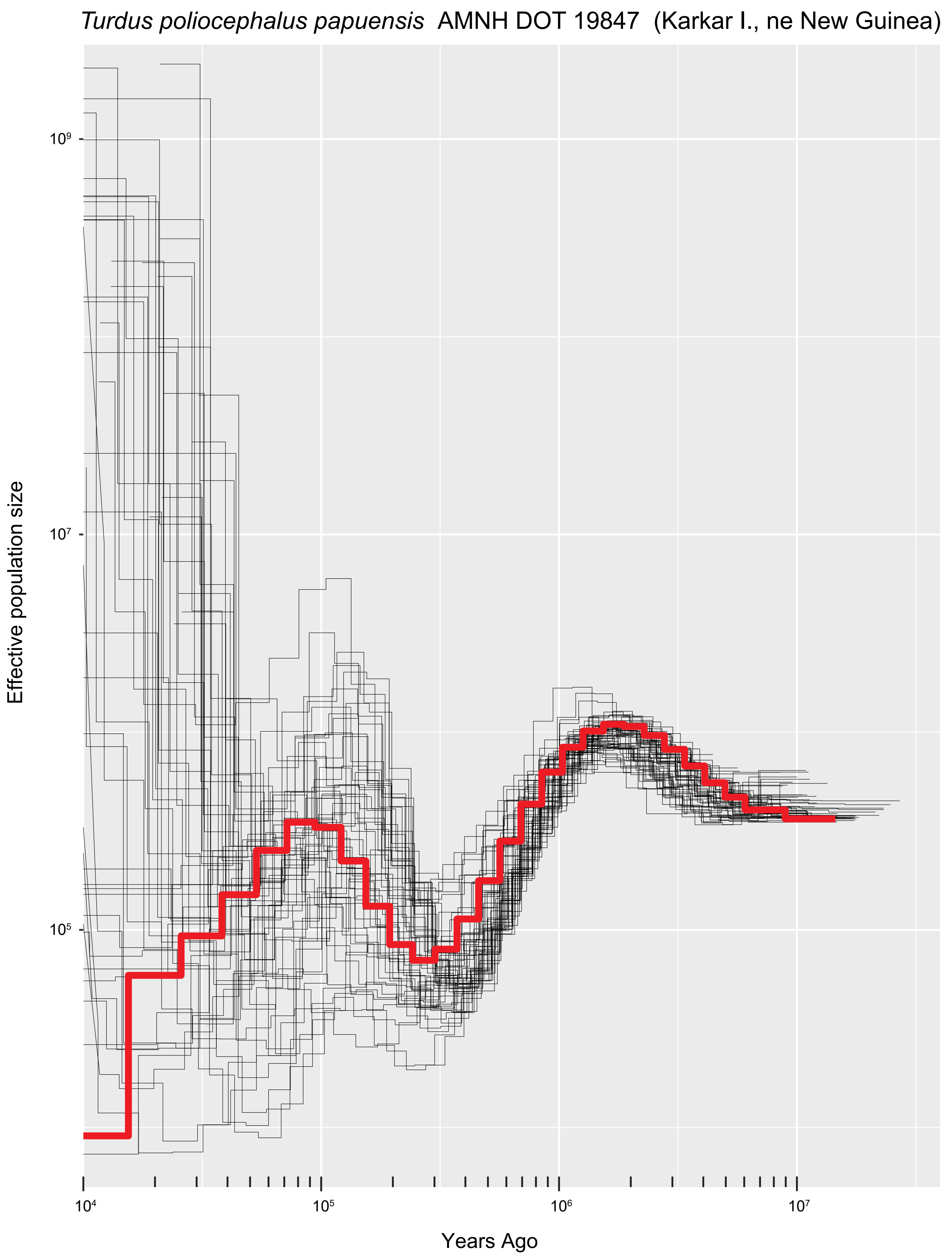
*Turdus poliocephalus papuensis* AMNH DOT 19847 (Karkar I., ne New Guinea)
Effective population size
$10^9$
$10^7$
$10^5$
$10^4$
$10^5$
$10^6$
$10^7$
Years Ago

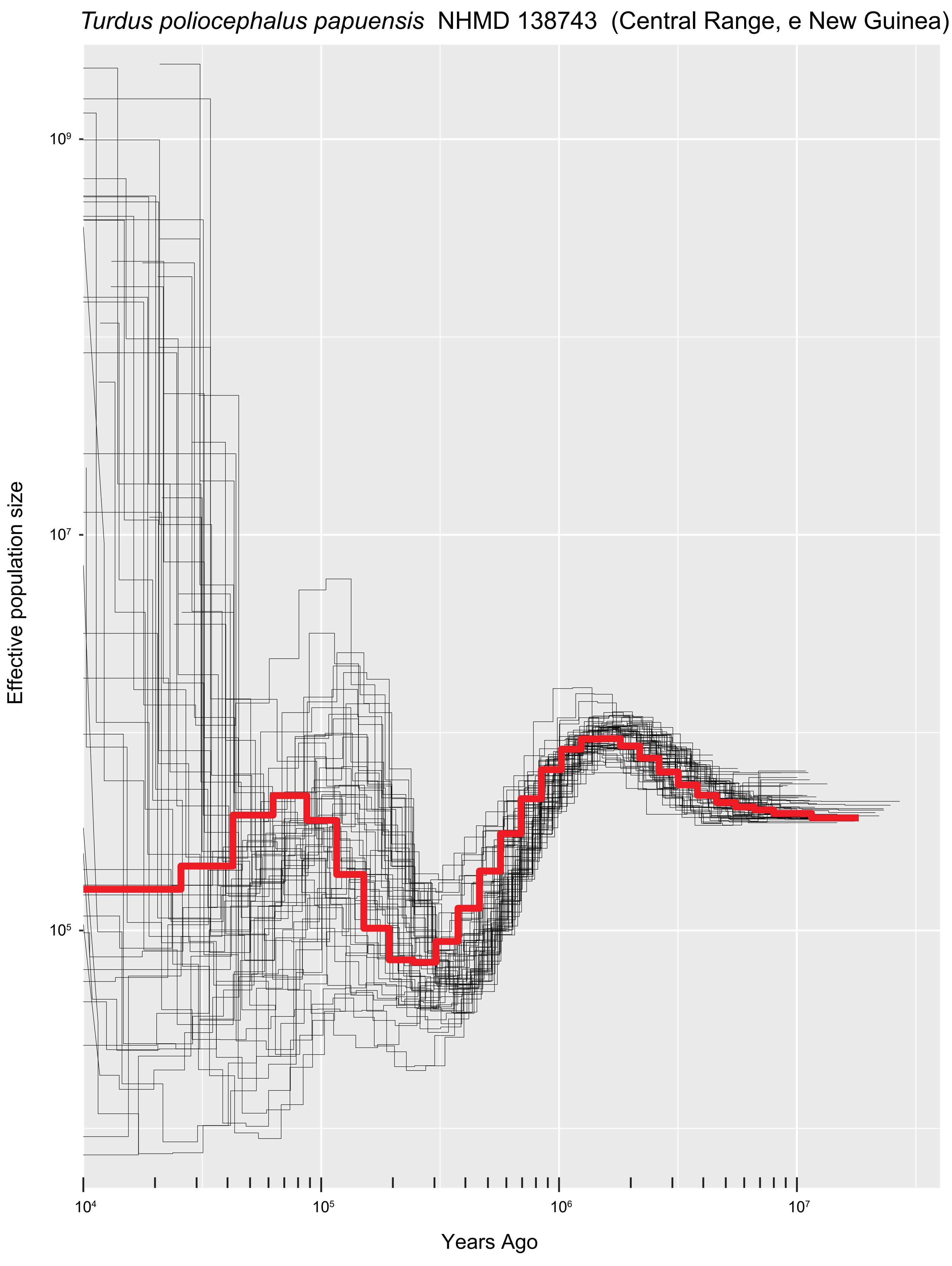

*Turdus poliocephalus papuensis* NHMD 138743 (Central Range, e New Guinea)
Effective population size
$10^9$
$10^7$
$10^5$
$10^4$
$10^5$
$10^6$
$10^7$
Years Ago

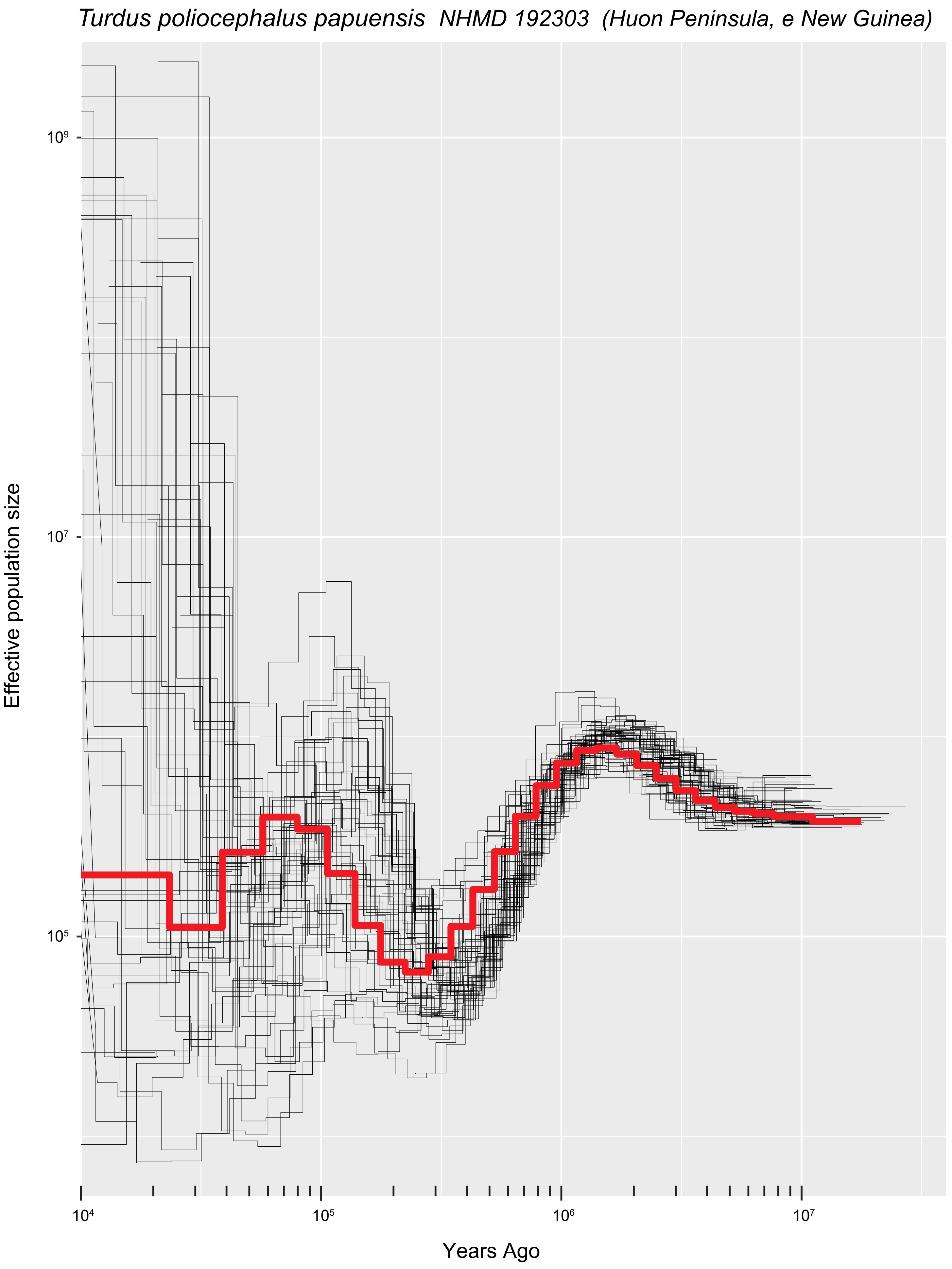

*Turdus poliocephalus papuensis* NHMD 192303 (Huon Peninsula, e New Guinea)
Effective population size
$10^9$
$10^7$
$10^5$
$10^4$
$10^5$
$10^6$
$10^7$
Years Ago

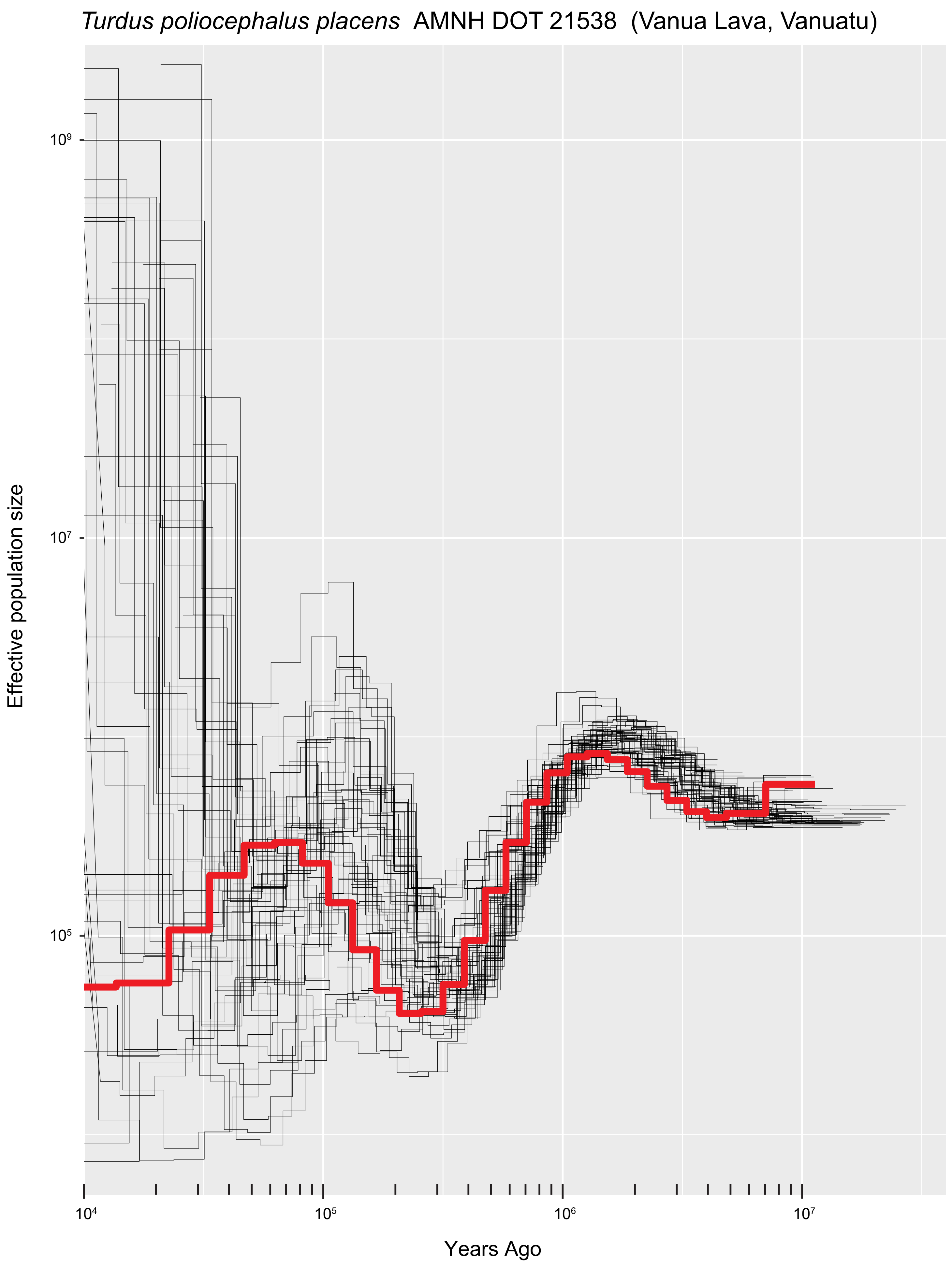
*Turdus poliocephalus placens* AMNH DOT 21538 (Vanua Lava, Vanuatu)
Effective population size
$10^9$
$10^7$
$10^5$
Years Ago
$10^4$
$10^5$
$10^6$
$10^7$

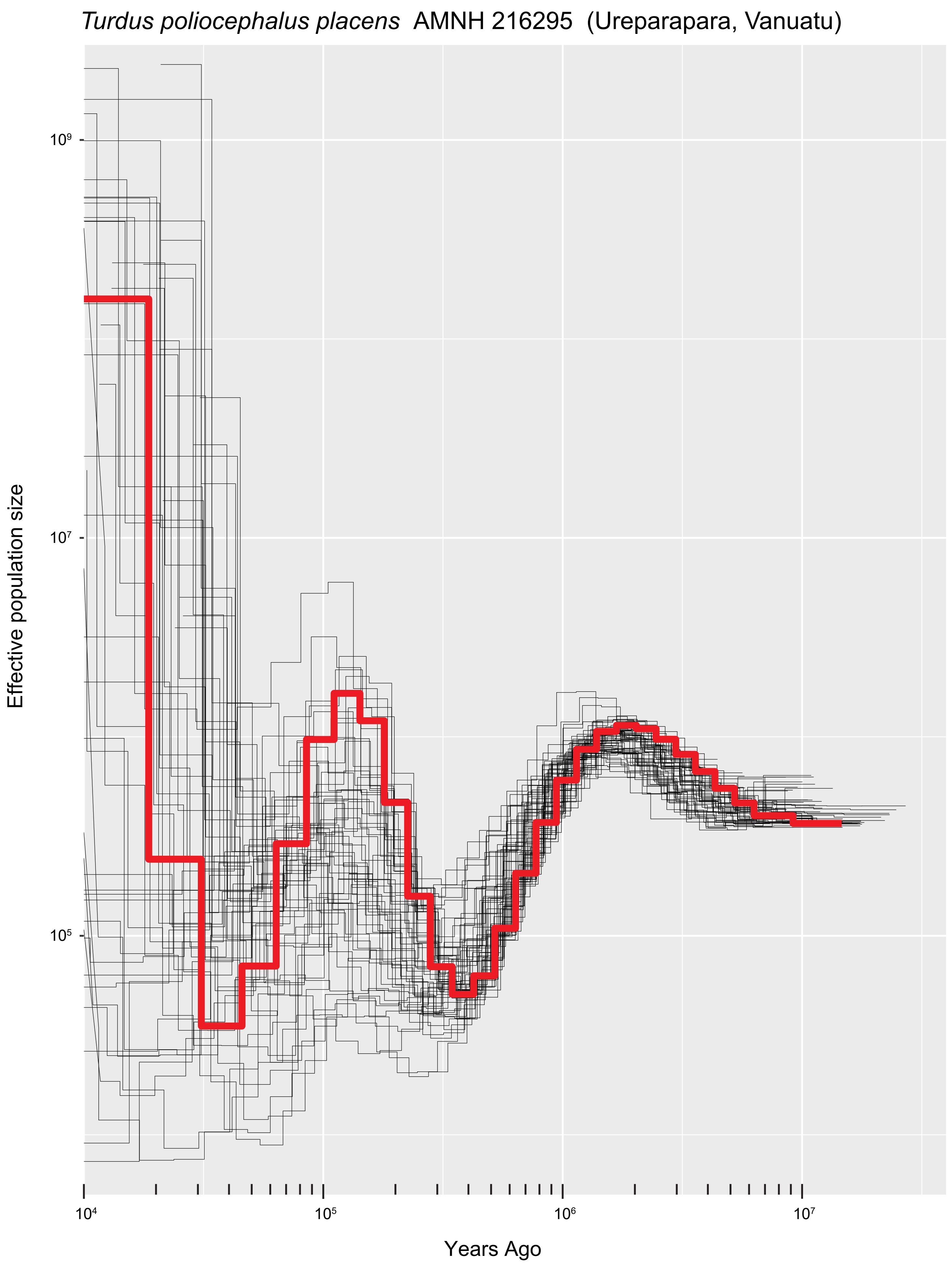

*Turdus poliocephalus placens* AMNH 216295 (Ureparapara, Vanuatu)
Effective population size
$10^9$
$10^7$
$10^5$
$10^4$
$10^5$
$10^6$
$10^7$
Years Ago

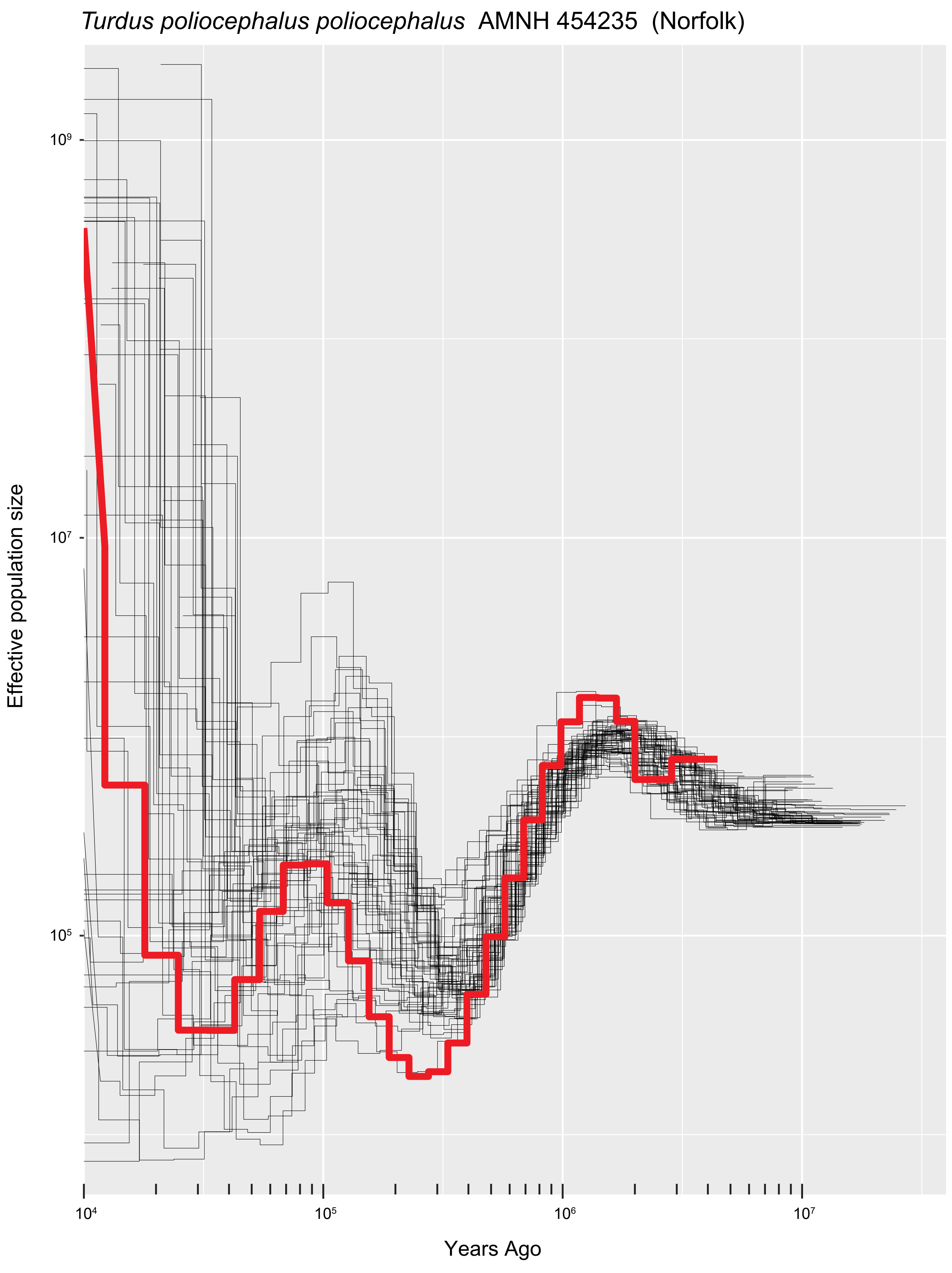
*Turdus poliocephalus poliocephalus* AMNH 454235 (Norfolk)
Effective population size
10^9
10^7
10^5
10^4
10^5
10^6
10^7
Years Ago

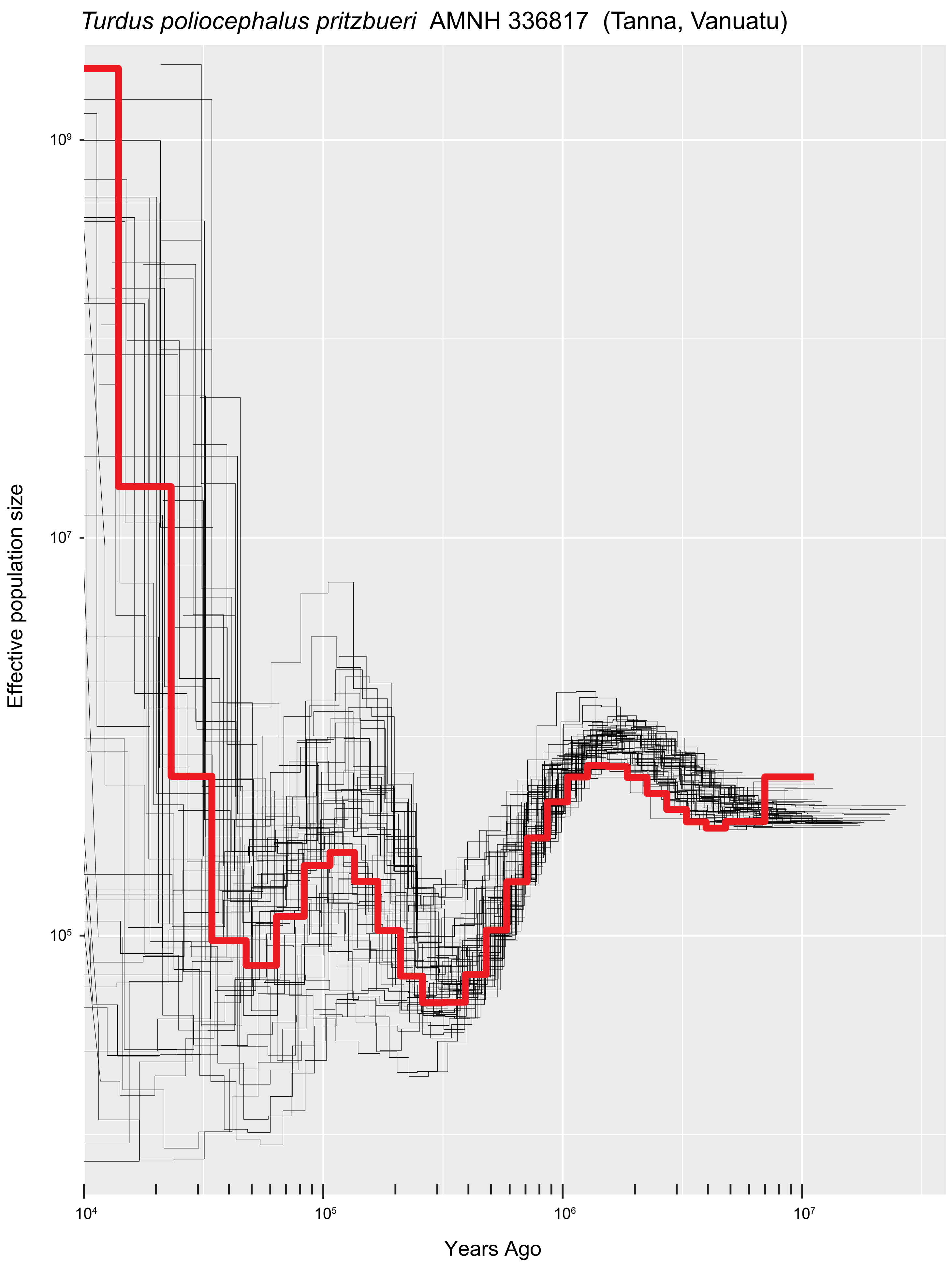

*Turdus poliocephalus pritzbueri* AMNH 336817 (Tanna, Vanuatu)
Effective population size
10⁹
10⁷
10⁵
10⁴
10⁵
10⁶
10⁷
Years Ago

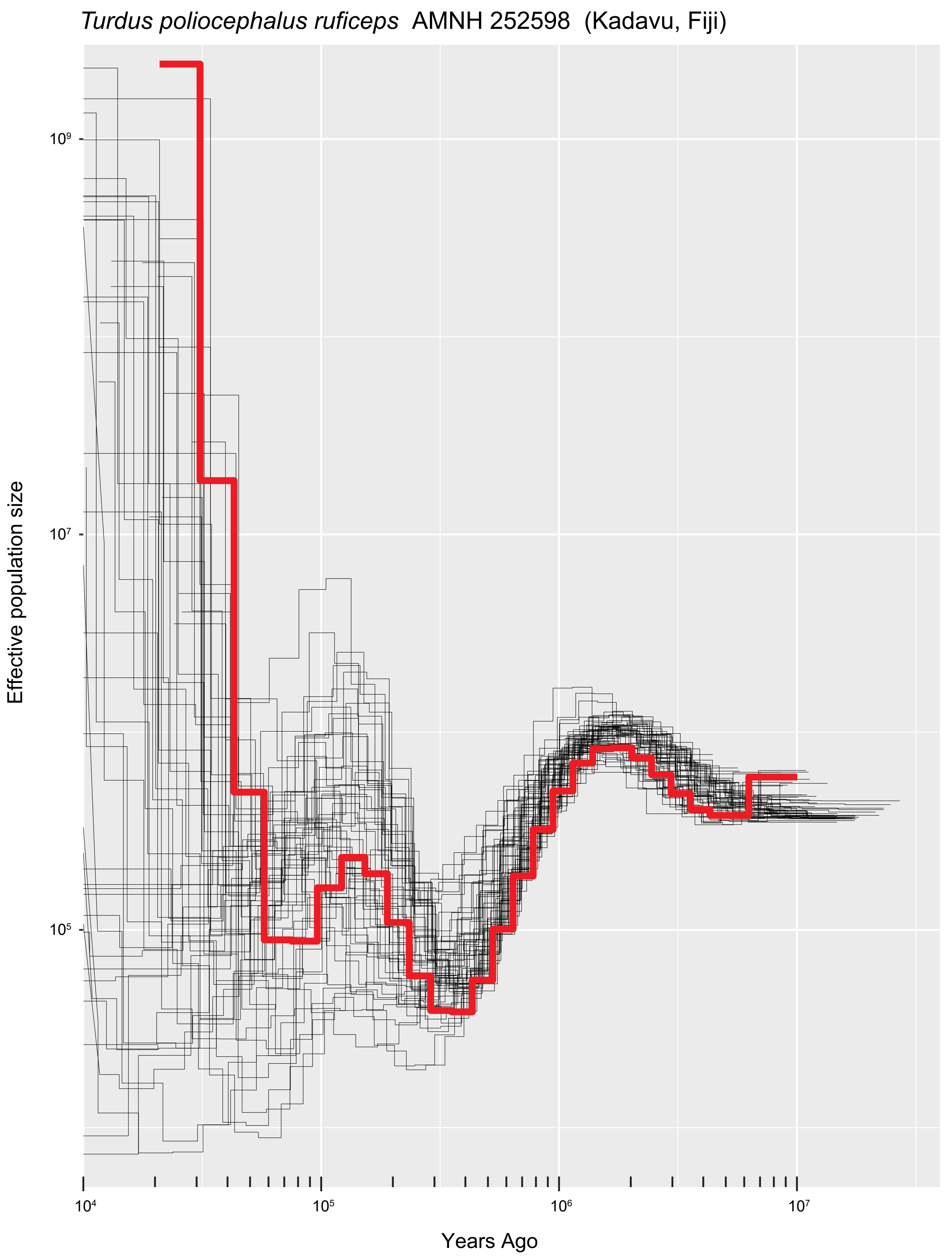

*Turdus poliocephalus ruficeps* AMNH 252598 (Kadavu, Fiji)
Effective population size
$10^9$
$10^7$
$10^5$
$10^4$
$10^5$
$10^6$
$10^7$
Years Ago

*Turdus poliocephalus samoensis* AMNH 206154 (Savai'i, Samoa)

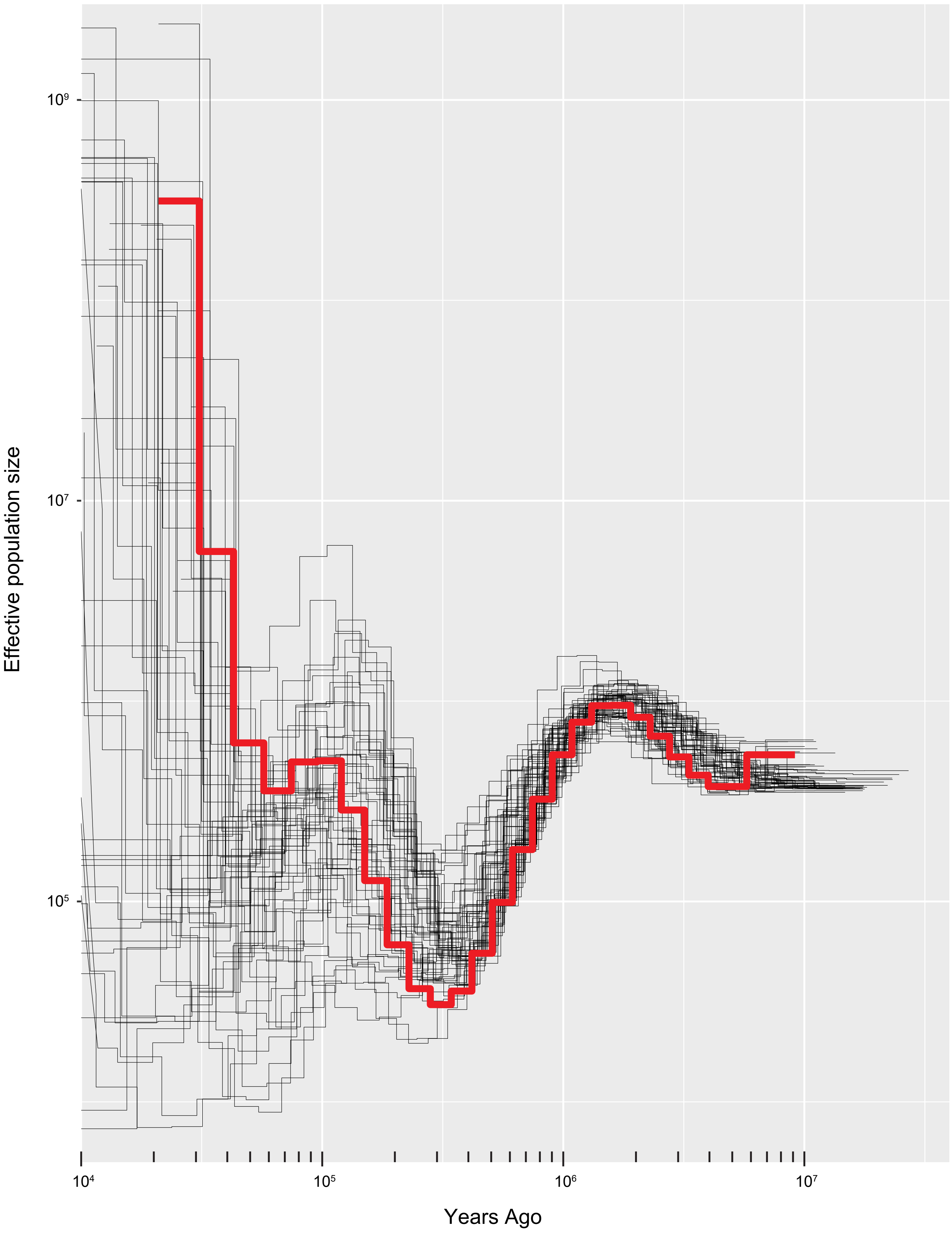

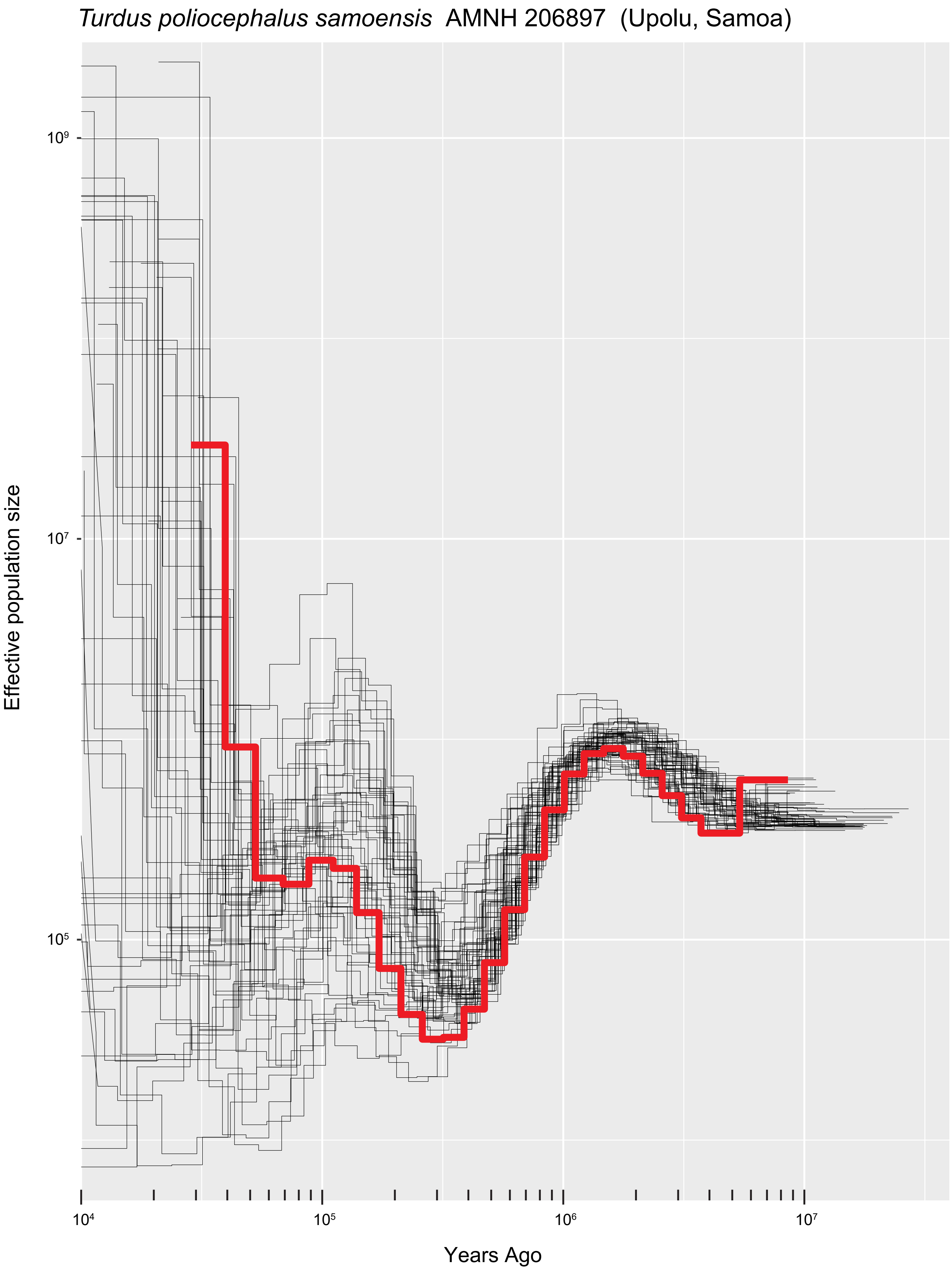
Turdus poliocephalus samoensis AMNH 206897 (Upolu, Samoa)
Effective population size
$10^9$
$10^7$
$10^5$
$10^4$
$10^5$
$10^6$
$10^7$
Years Ago

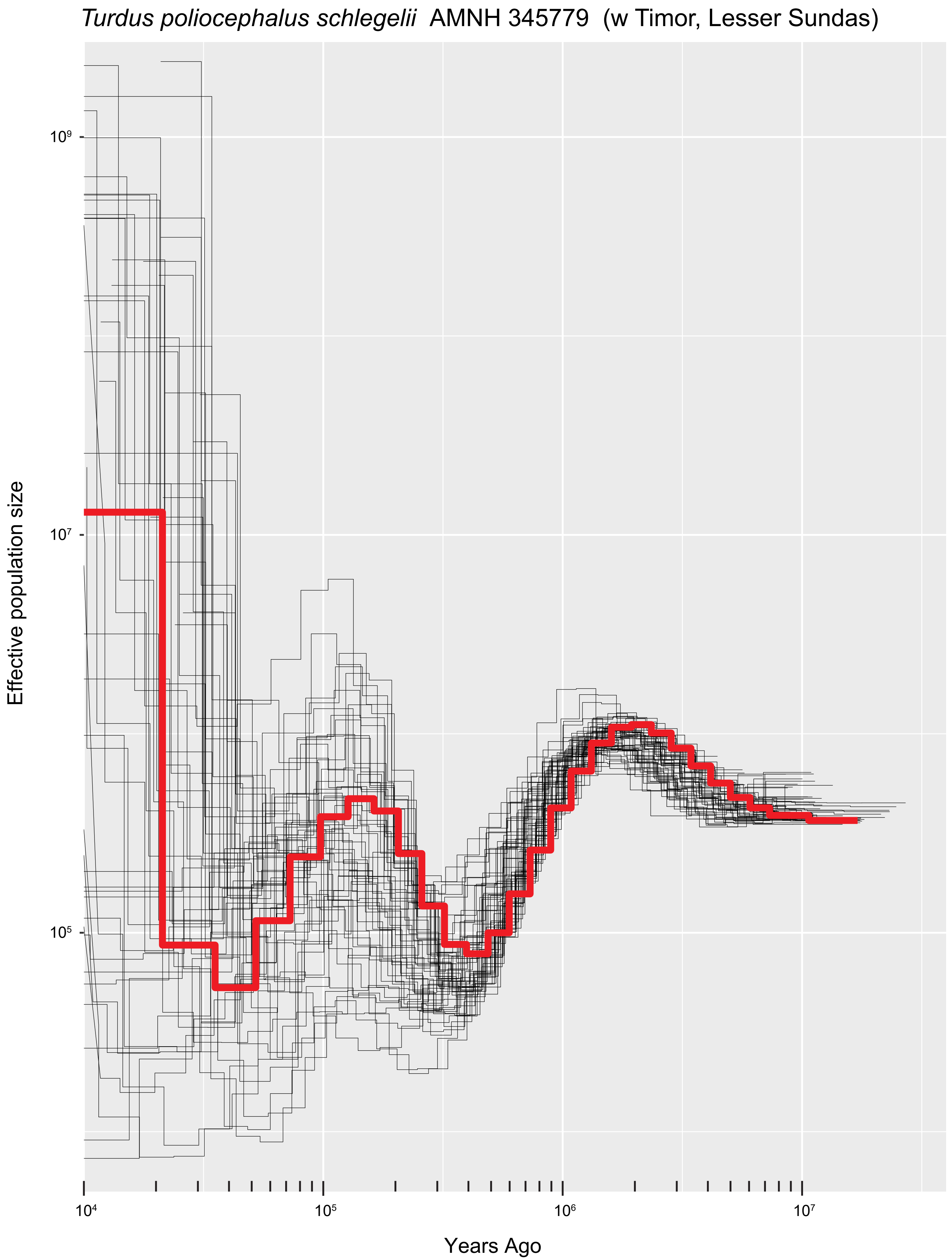

Turdus poliocephalus schlegelii AMNH 345779 (w Timor, Lesser Sundas)
Effective population size
10⁹
10⁷
10⁵
10⁴
10⁵
10⁶
10⁷
Years Ago

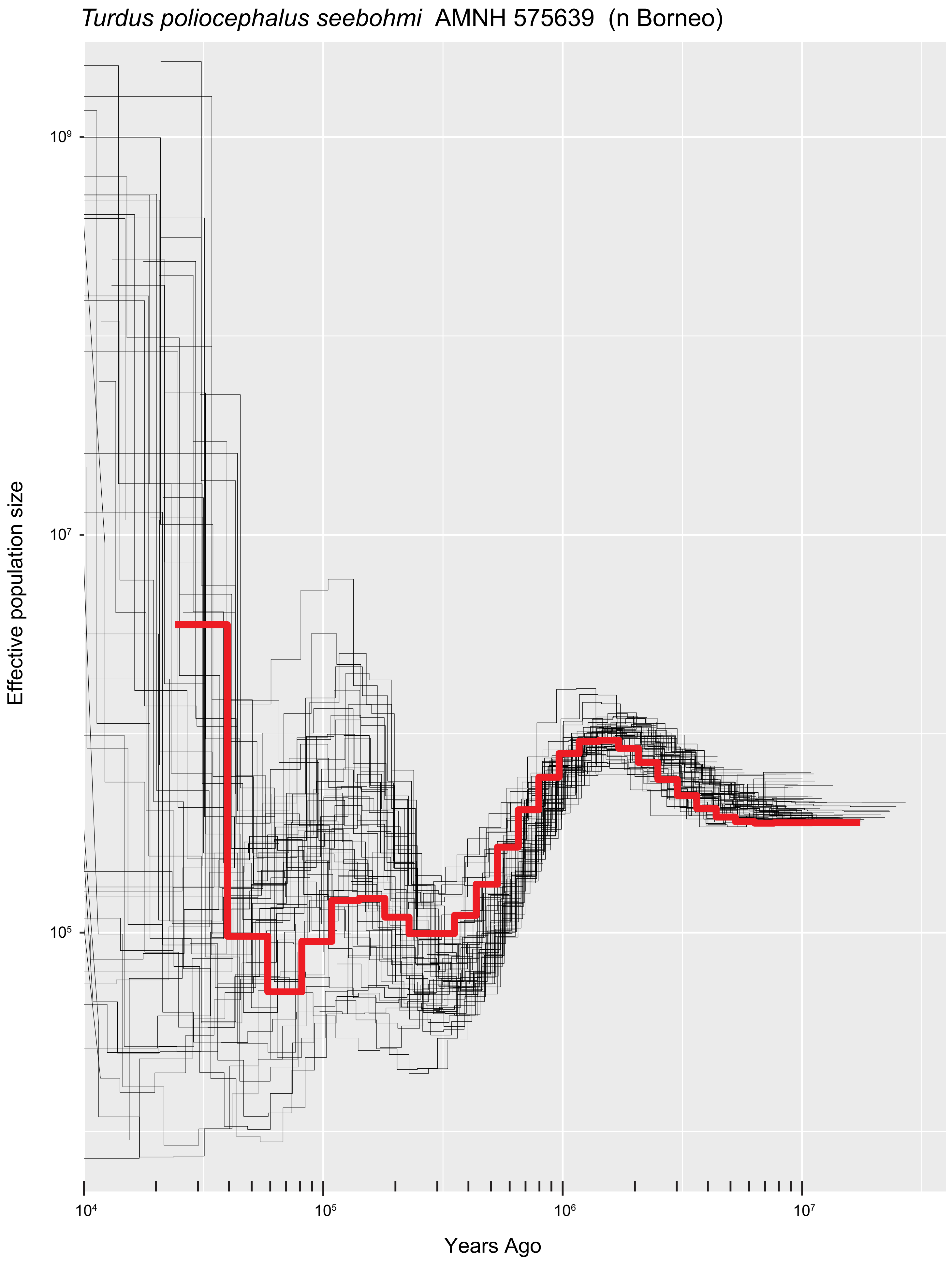
*Turdus poliocephalus seebohmi* AMNH 575639 (n Borneo)
Effective population size
$10^9$
$10^7$
$10^5$
$10^4$
$10^5$
$10^6$
$10^7$
Years Ago

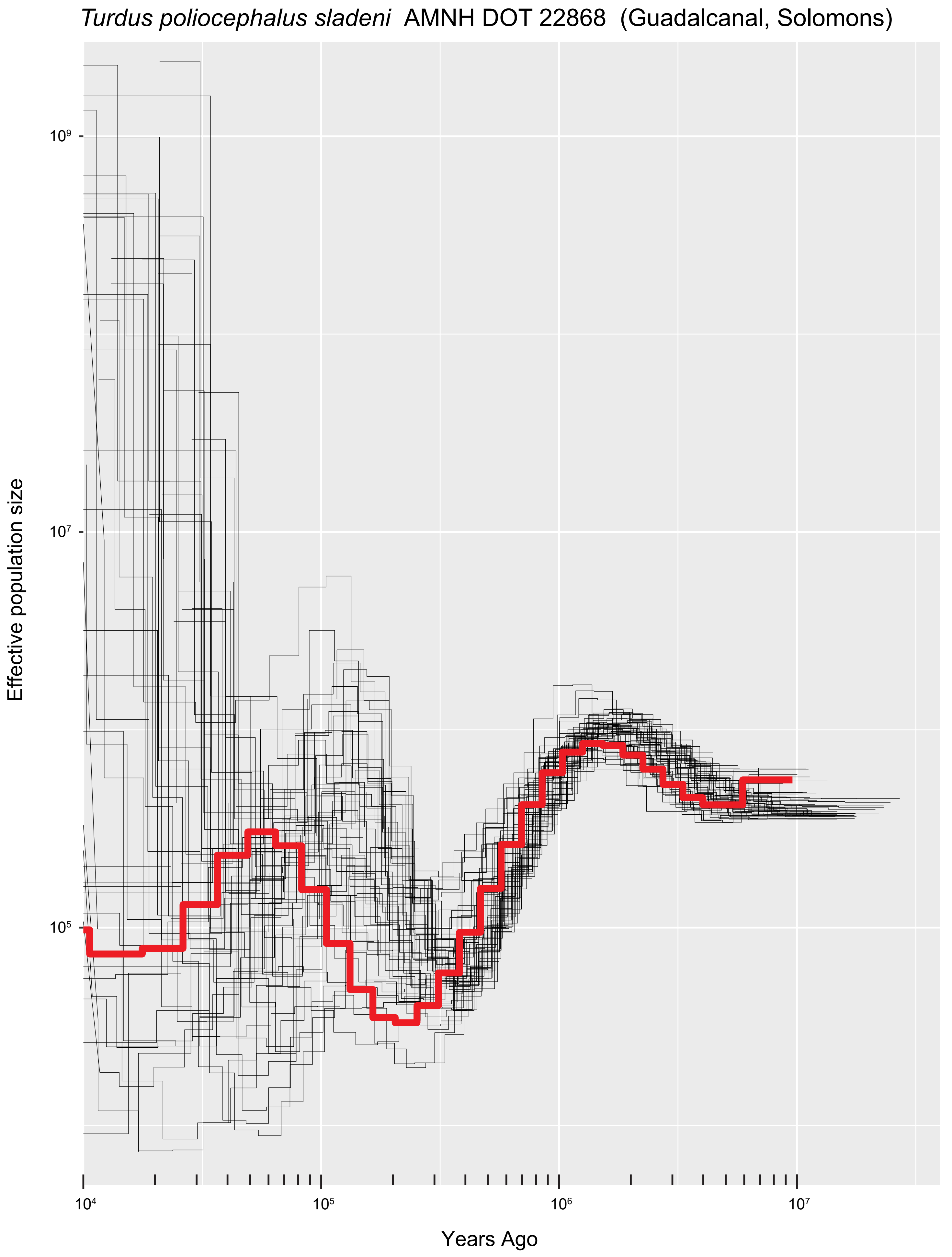
*Turdus poliocephalus sladeni* AMNH DOT 22868 (Guadalcanal, Solomons)
Effective population size
$10^9$
$10^7$
$10^5$
$10^4$
$10^5$
$10^6$
$10^7$
Years Ago

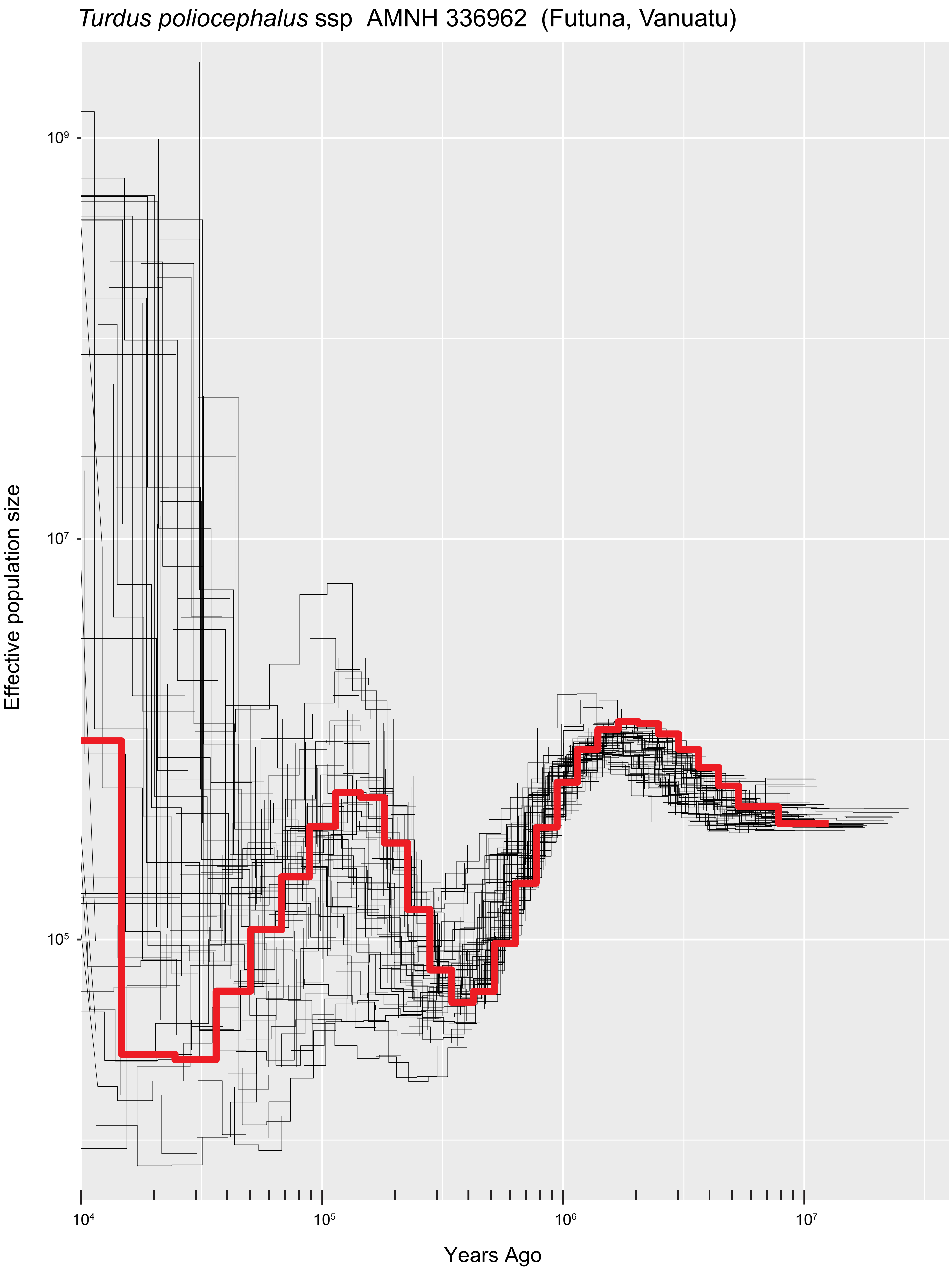
*Turdus poliocephalus* ssp AMNH 336962 (Futuna, Vanuatu)
Effective population size
$10^9$
$10^7$
$10^5$
$10^4$
$10^5$
$10^6$
$10^7$
Years Ago

*Turdus poliocephalus* ssp  B36870  (Panay, Philippines)

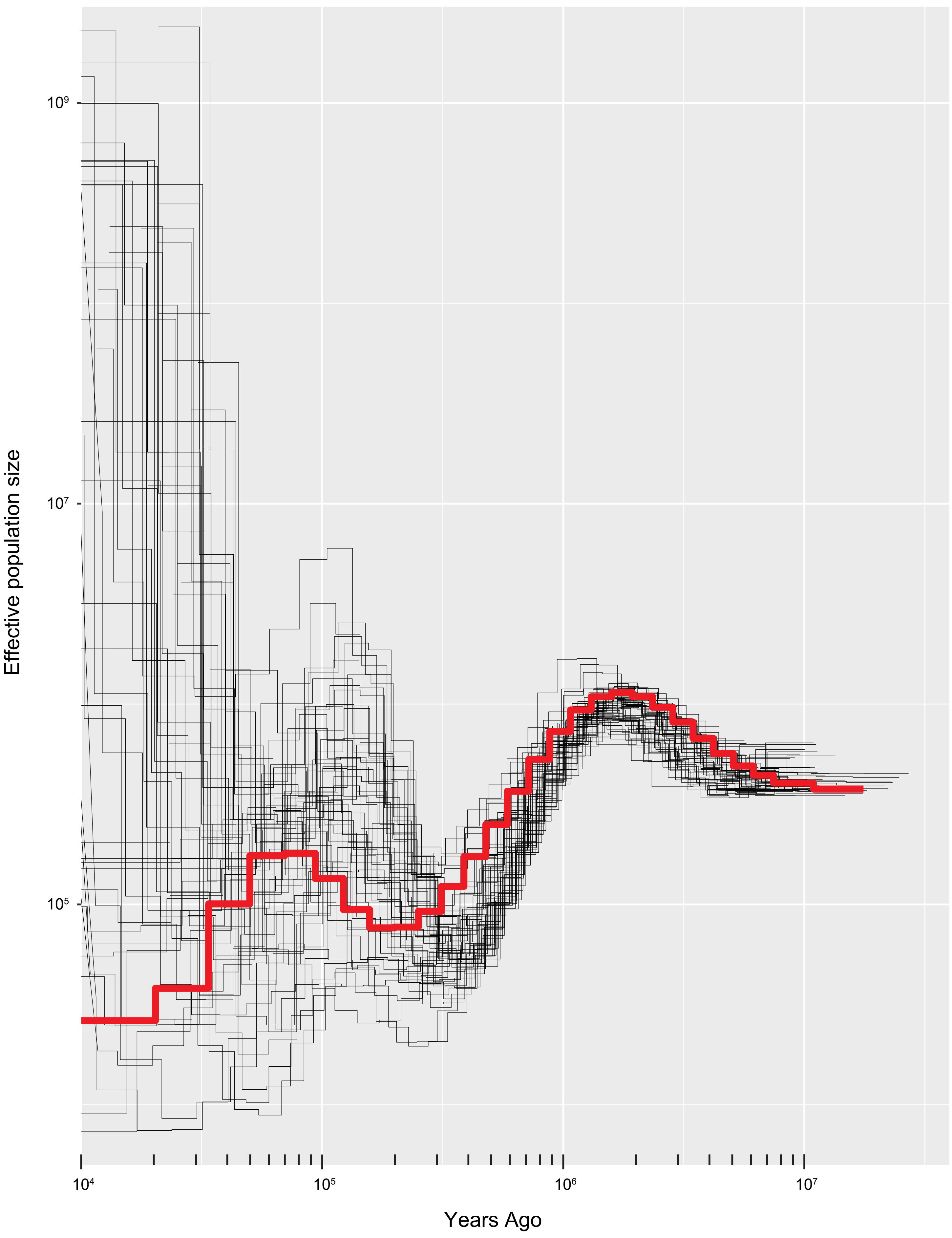

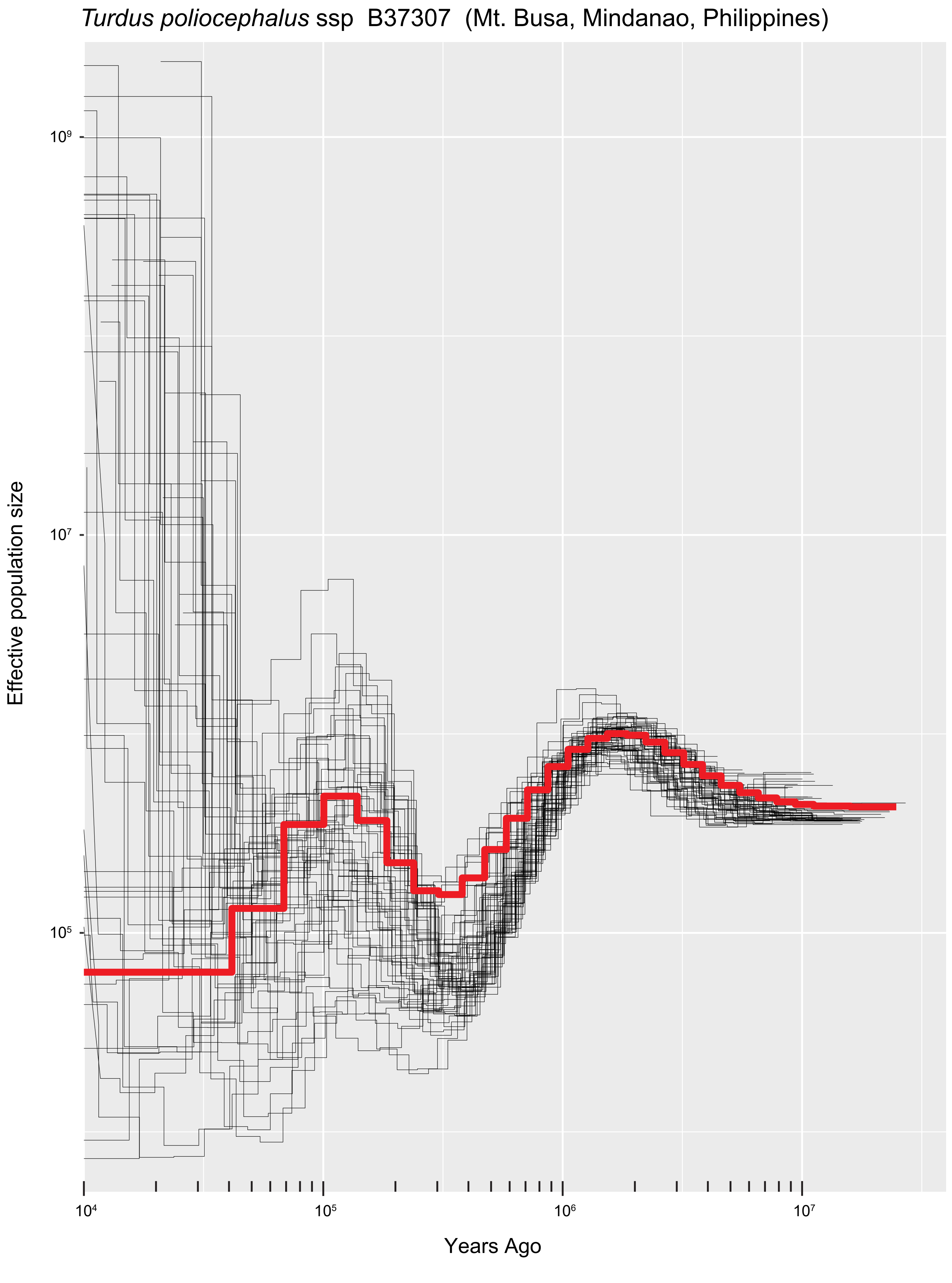
*Turdus poliocephalus* ssp B37307 (Mt. Busa, Mindanao, Philippines)
Effective population size
$10^9$
$10^7$
$10^5$
$10^4$
$10^5$
$10^6$
$10^7$
Years Ago

*Turdus poliocephalus* ssp  FMNH 358378  (Sibuyan, Philippines)

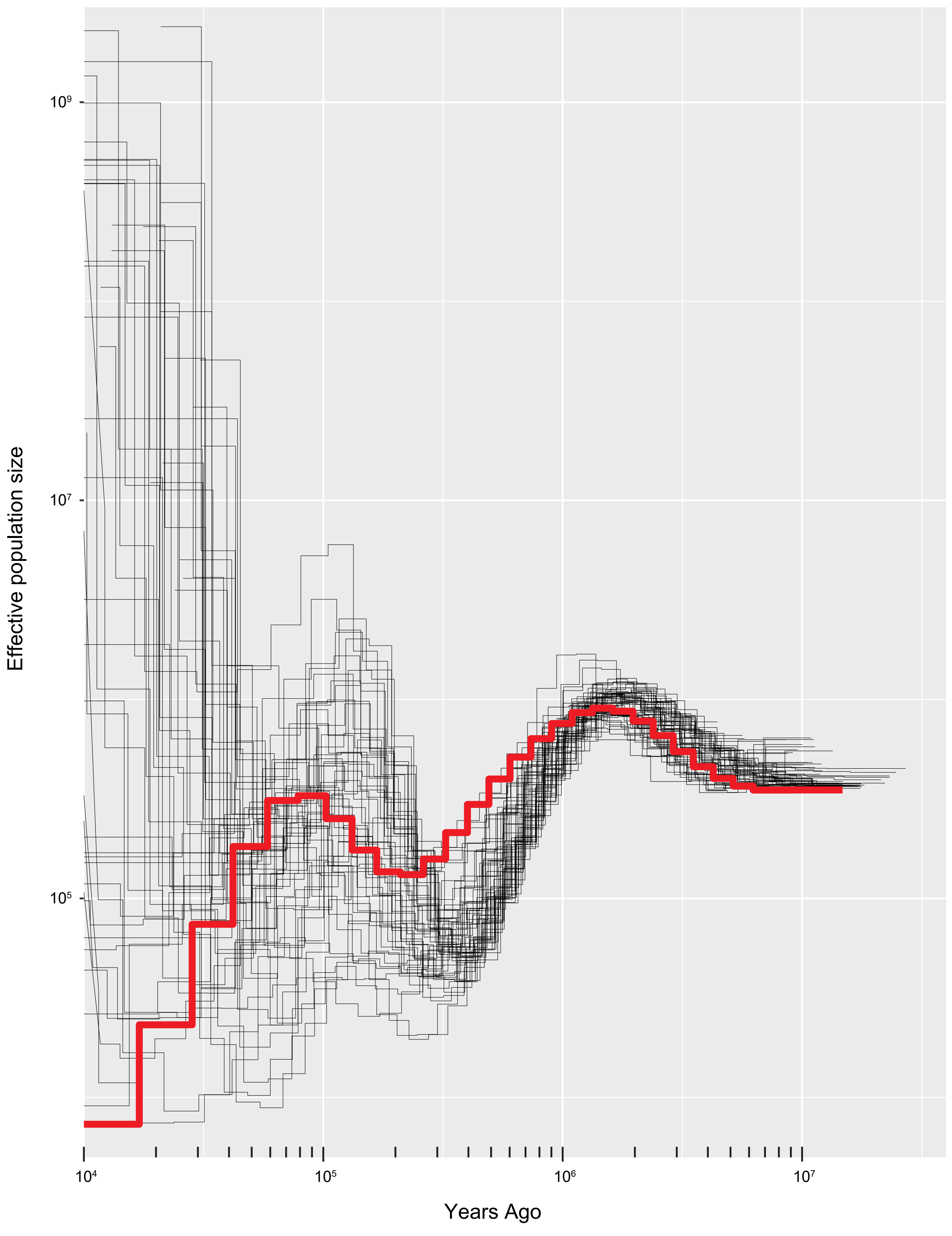

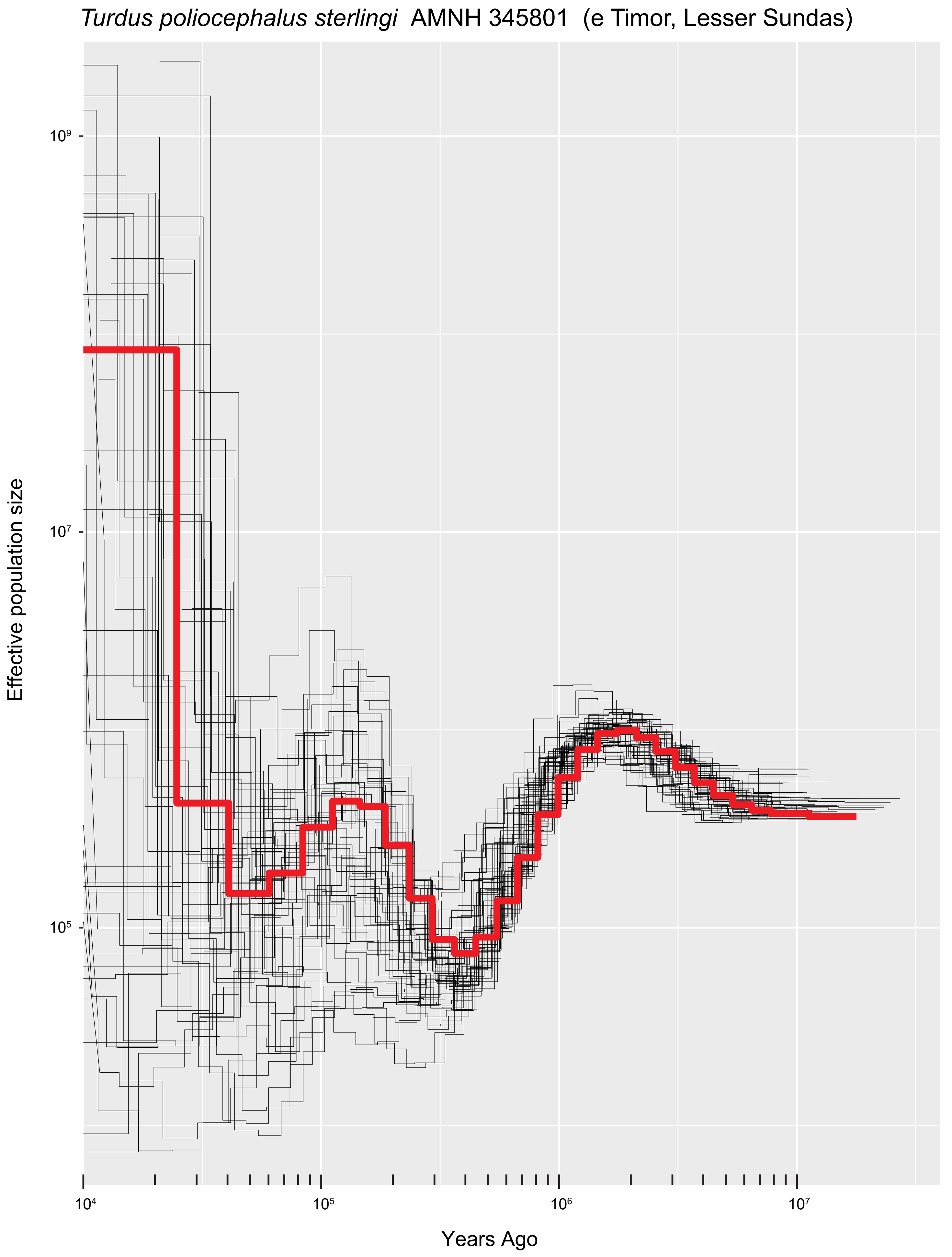

*Turdus poliocephalus sterlingi* AMNH 345801 (e Timor, Lesser Sundas)
Effective population size
$10^9$
$10^7$
$10^5$
$10^4$
$10^5$
$10^6$
$10^7$
Years Ago

*Turdus poliocephalus thomassoni* AMNH 416872 (n Luzon, Philippines)

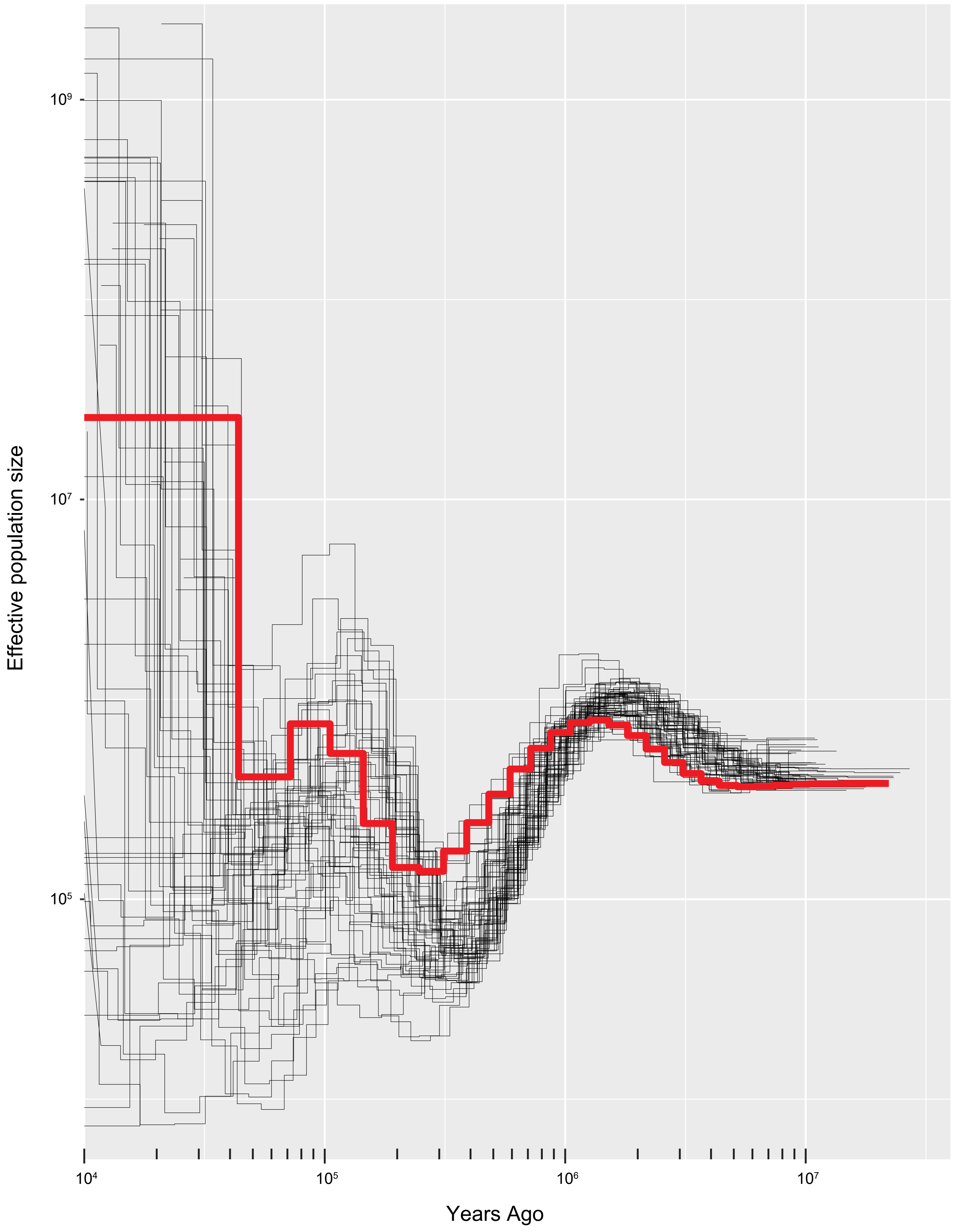

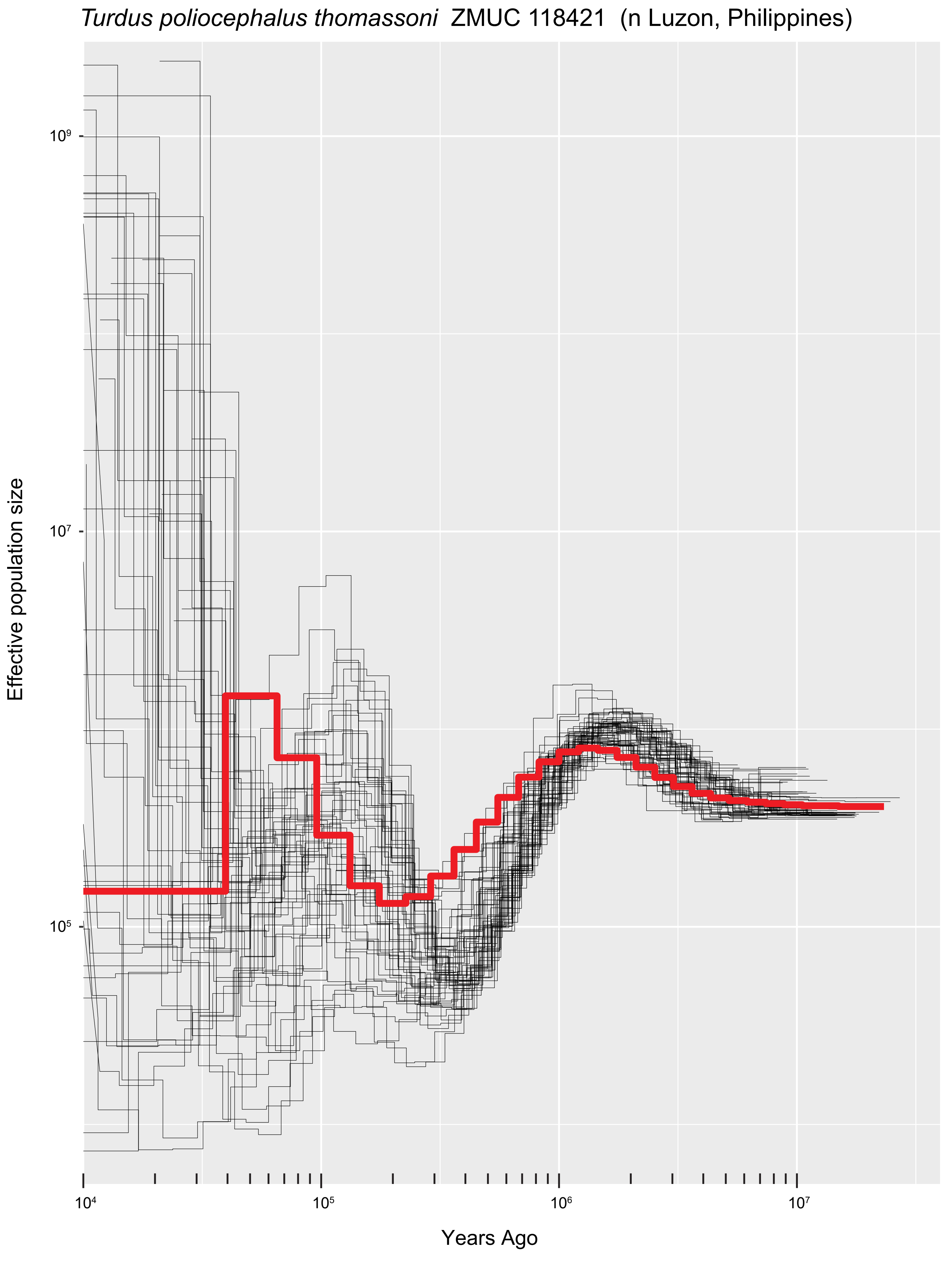

Turdus poliocephalus thomassoni  ZMUC 118421  (n Luzon, Philippines)
Effective population size
10⁹
10⁷
10⁵
10⁴
10⁵
10⁶
10⁷
Years Ago

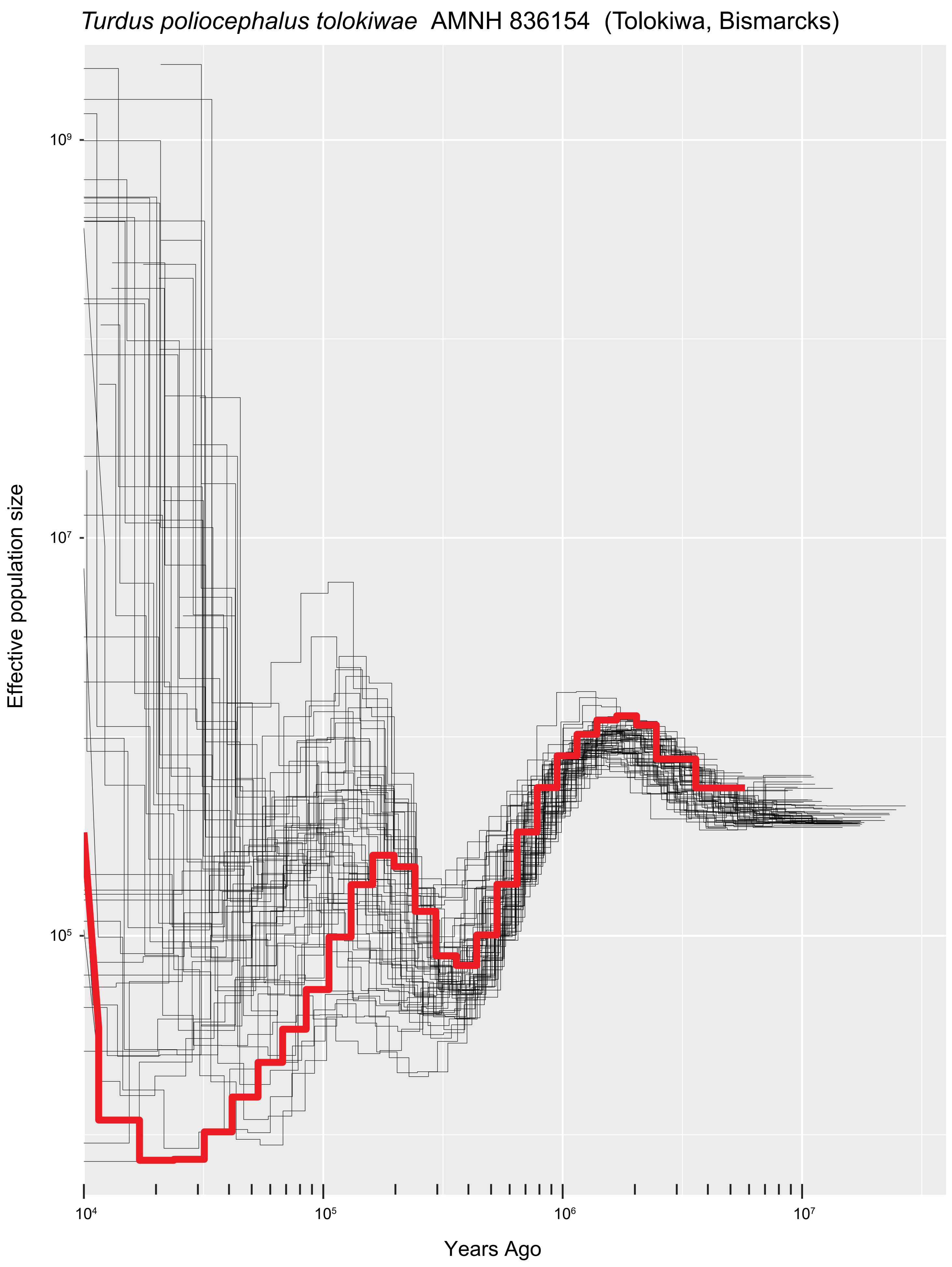

*Turdus poliocephalus tolokiwae* AMNH 836154 (Tolokiwa, Bismarcks)
Effective population size
$10^9$
$10^7$
$10^5$
$10^4$
$10^5$
$10^6$
$10^7$
Years Ago

### *Turdus poliocephalus vanikorensis* AMNH 214400 (Utupua, Santa Cruz Is.)

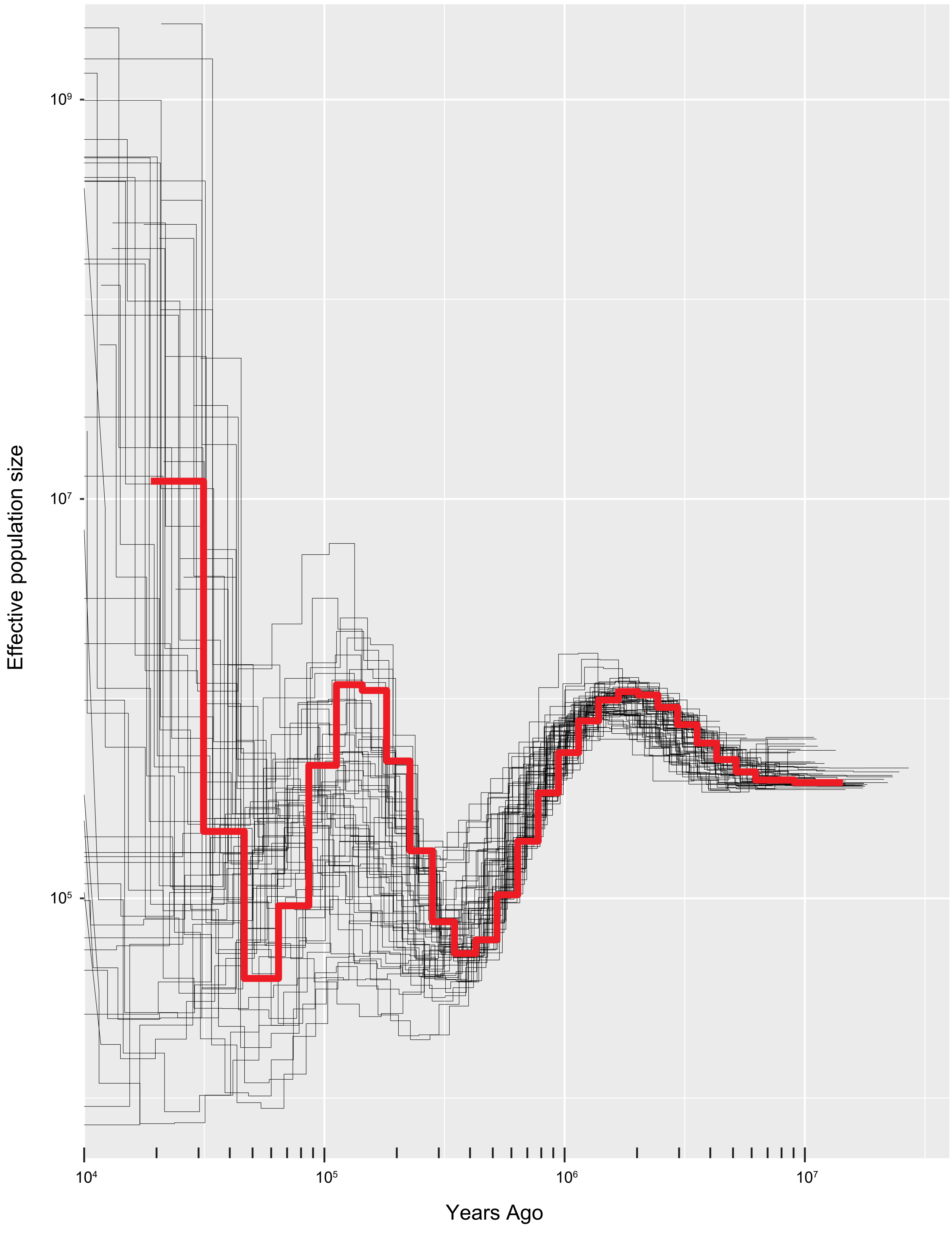

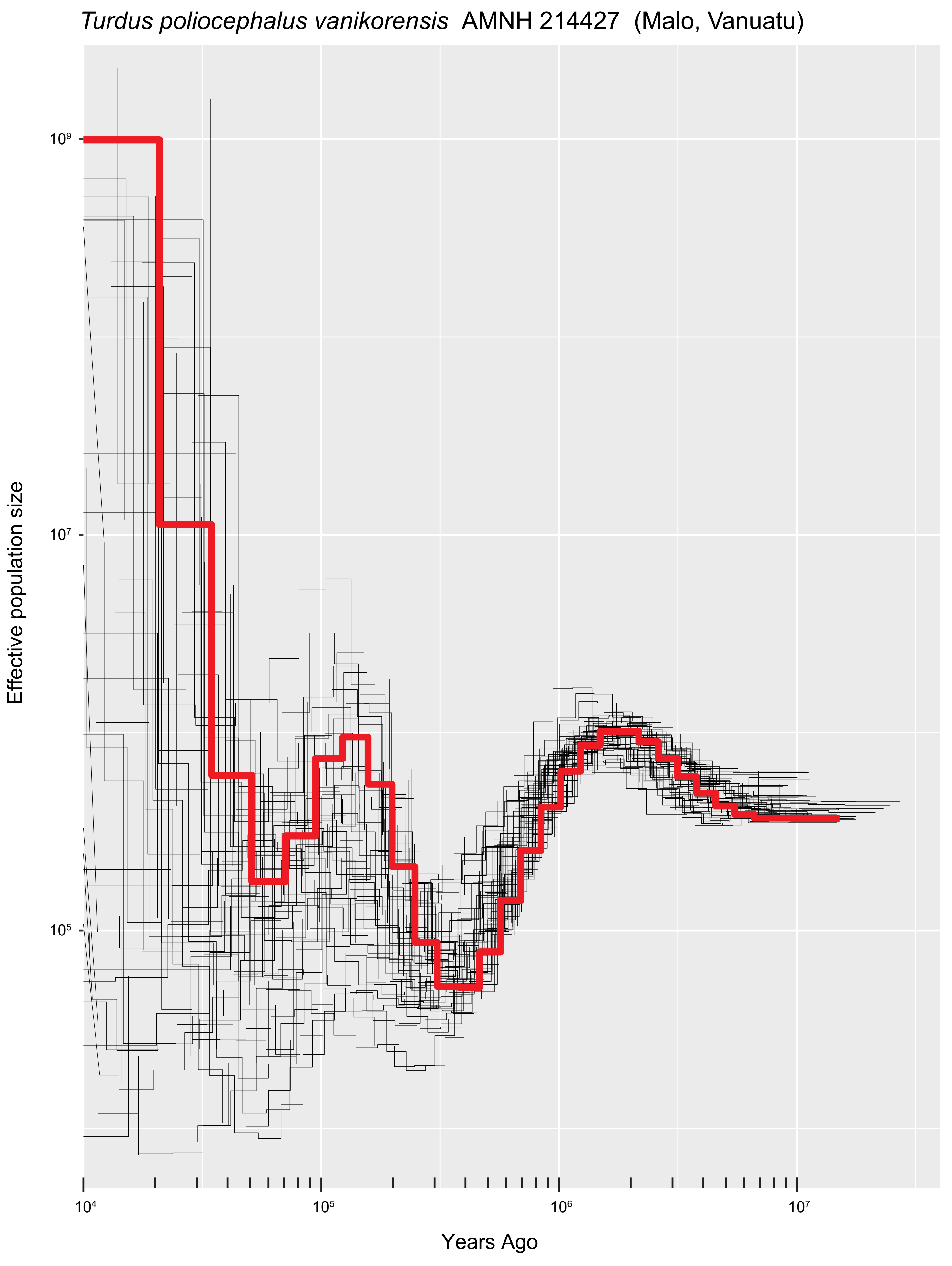

*Turdus poliocephalus vanikorensis* AMNH 214427 (Malo, Vanuatu)
Effective population size
$10^9$
$10^7$
$10^5$
Years Ago
$10^4$
$10^5$
$10^6$
$10^7$

*Turdus poliocephalus vanikorensis* AMNH 214440 (Espiritu Santo, Vanuatu)

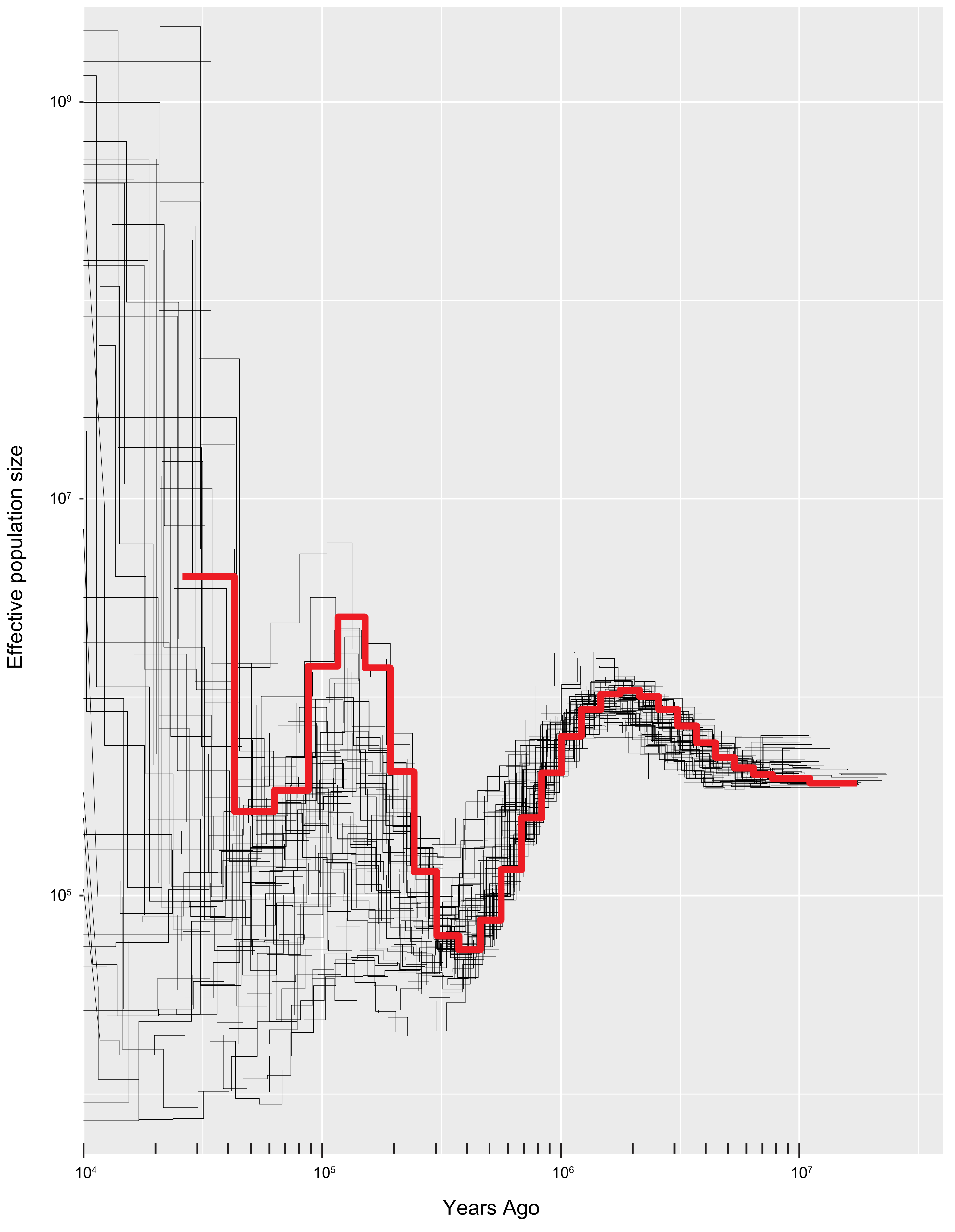

*Turdus poliocephalus versteegi* AMNH 340312 (w New Guinea)

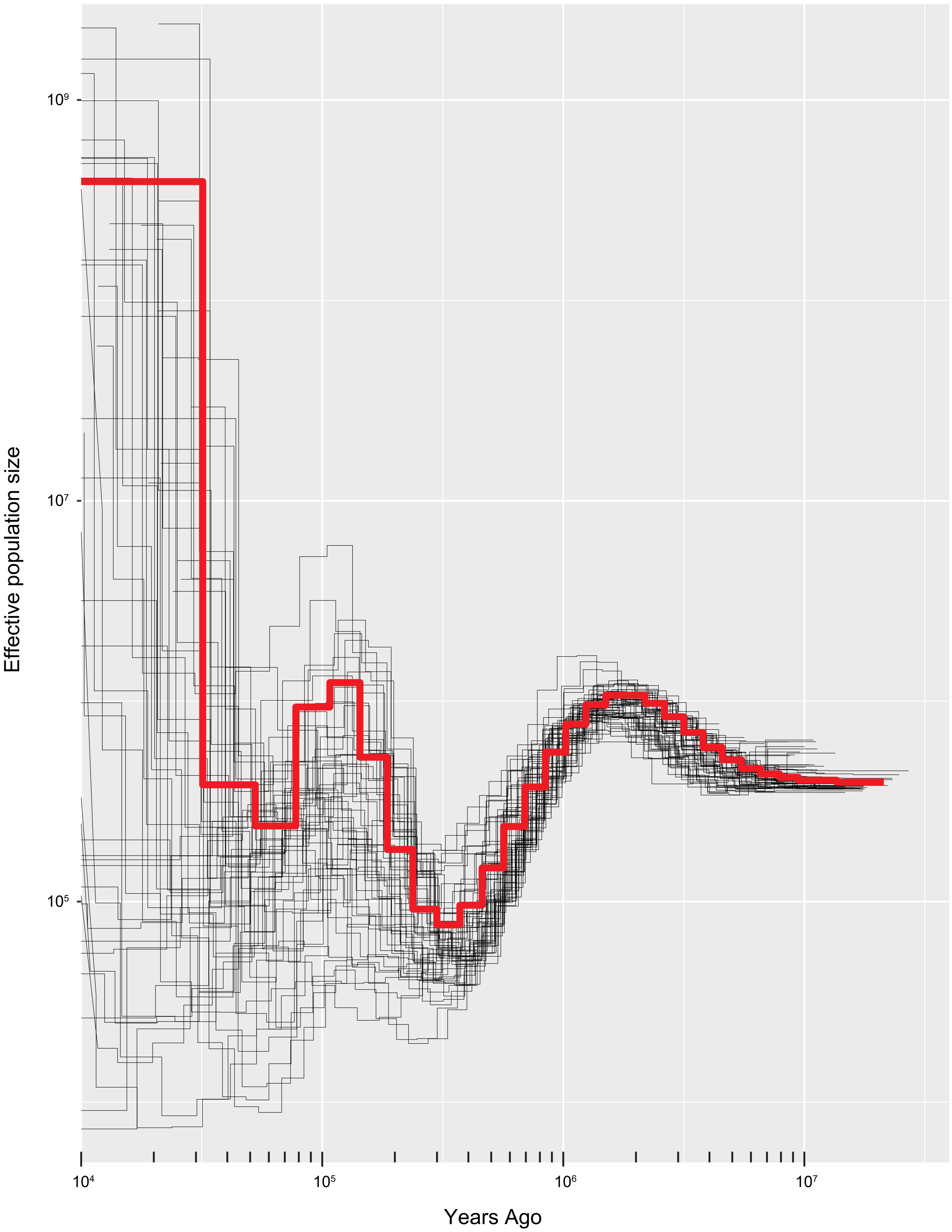

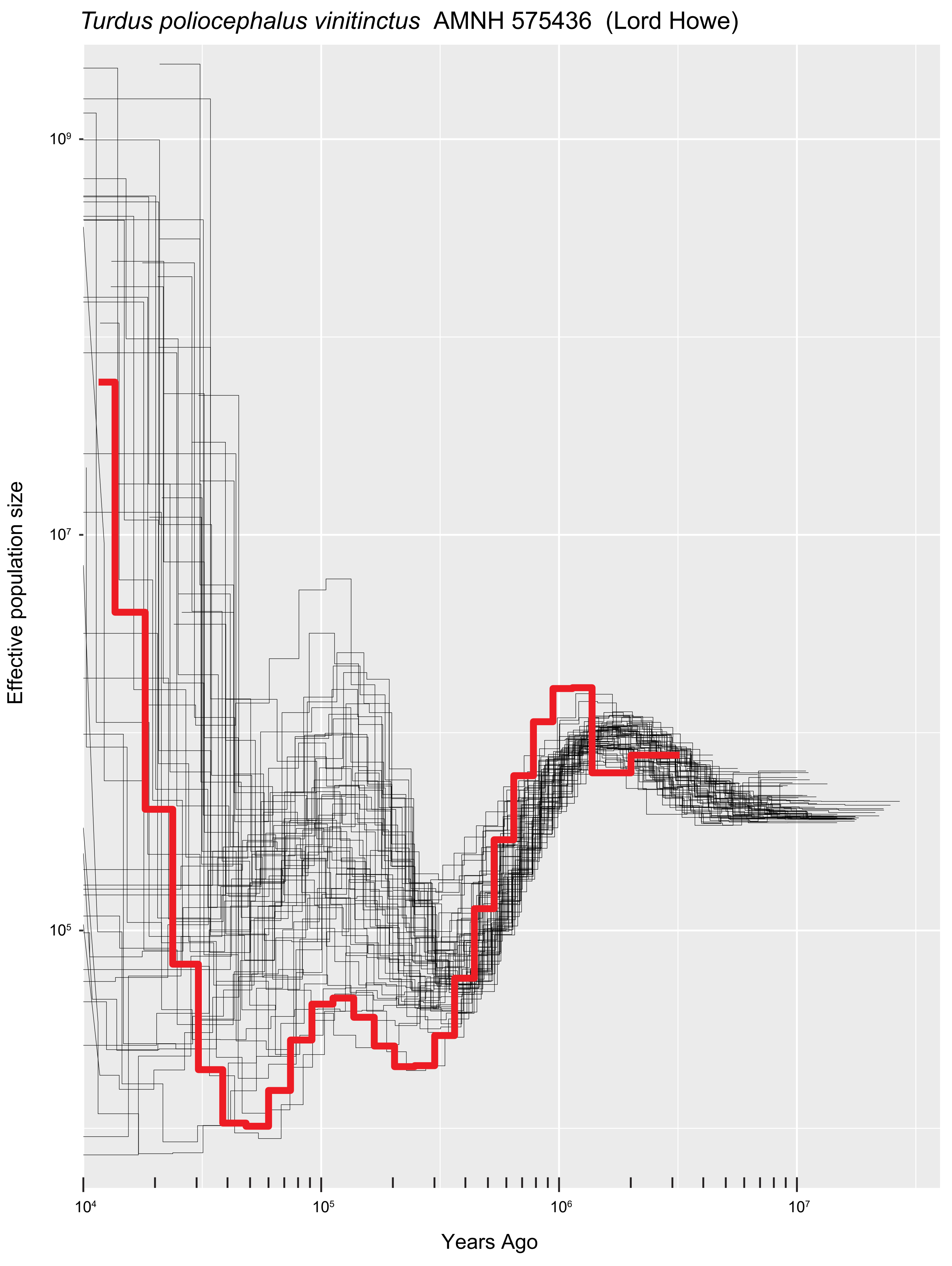

Turdus poliocephalus vinitinctus AMNH 575436 (Lord Howe)
Effective population size
10^9
10^7
10^5
10^4
10^5
10^6
10^7
Years Ago

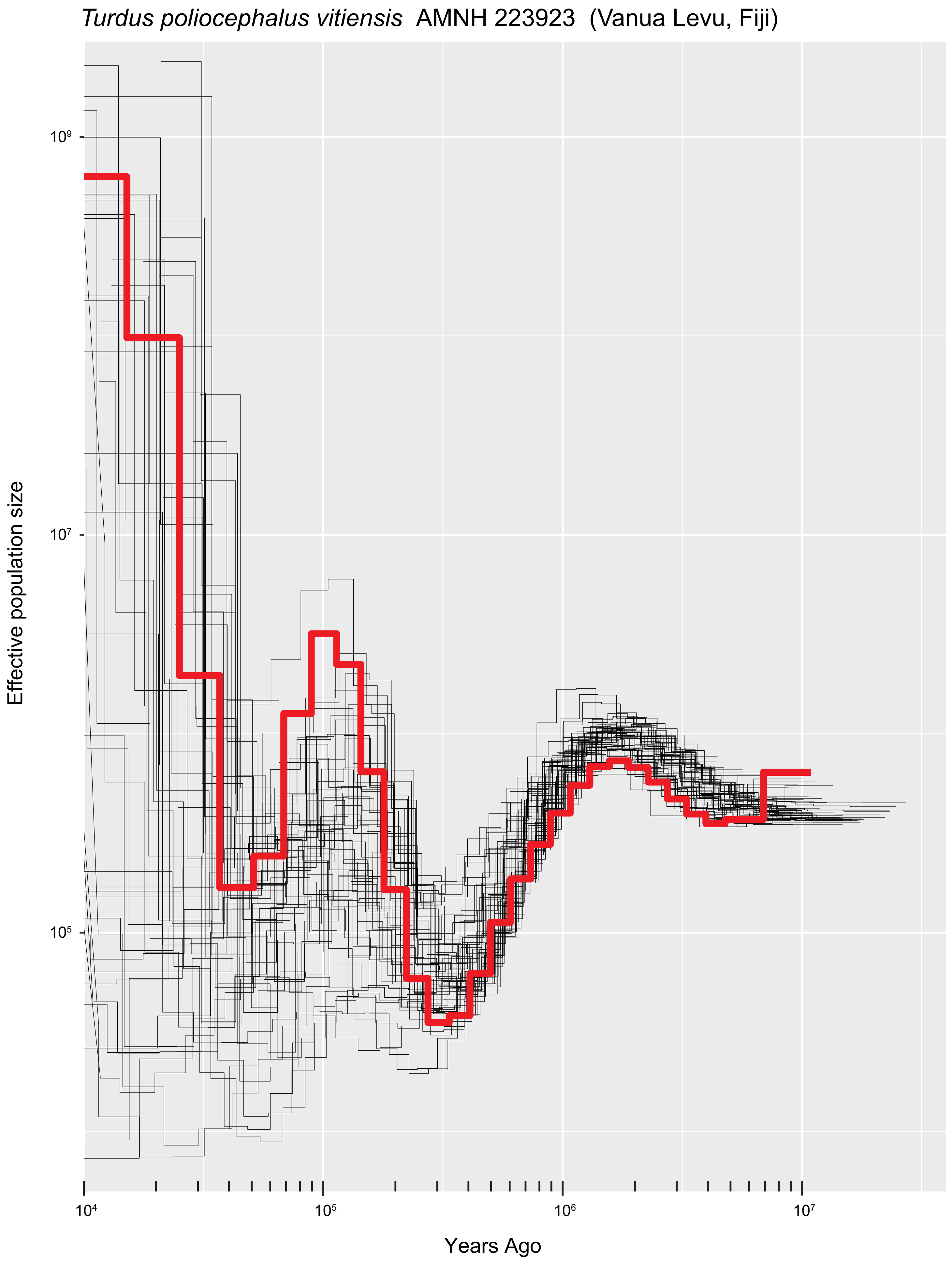

*Turdus poliocephalus vitiensis* AMNH 223923 (Vanua Levu, Fiji)
Effective population size
10^9
10^7
10^5
10^4
10^5
10^6
10^7
Years Ago

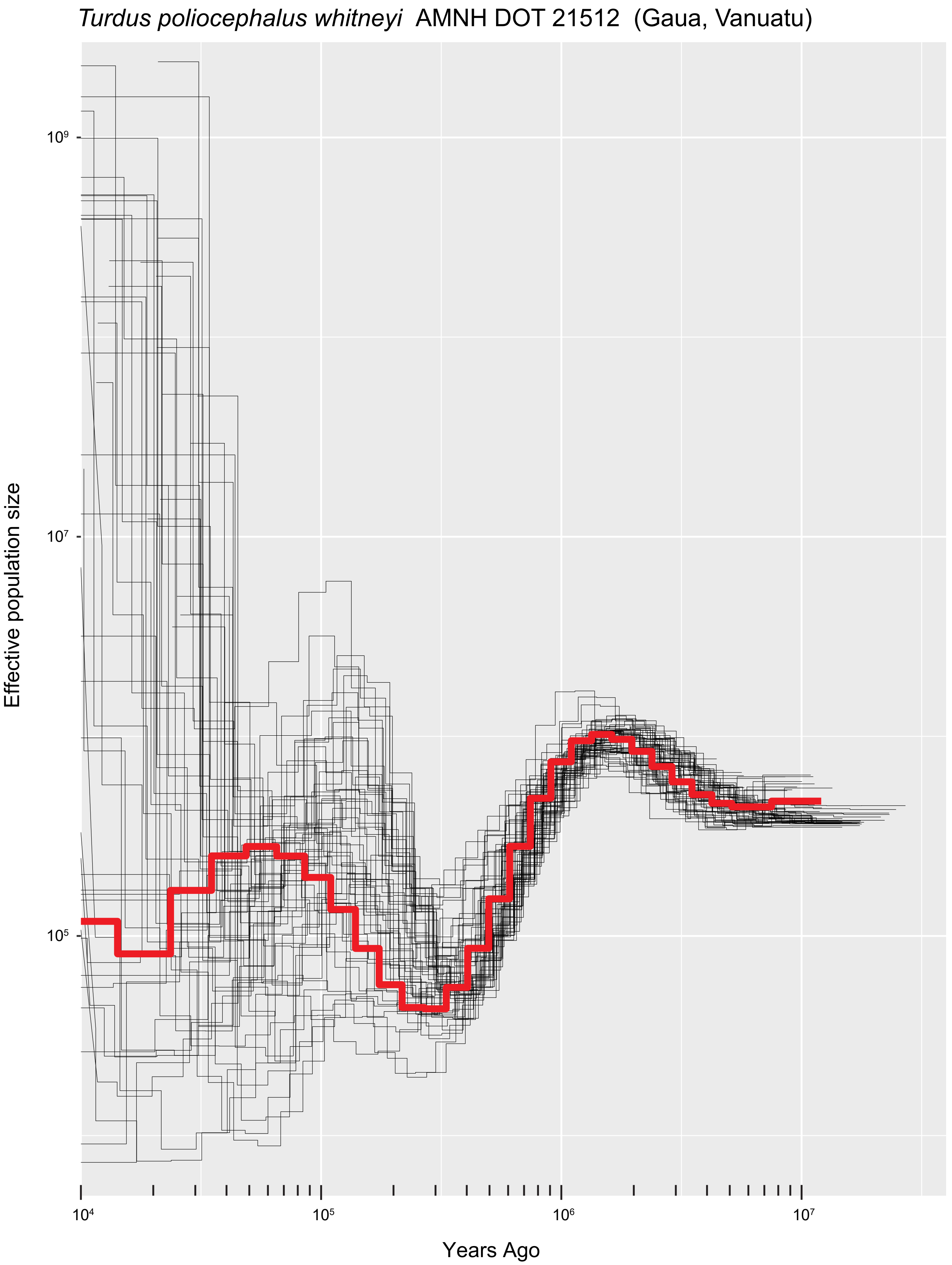

*Turdus poliocephalus whitneyi* AMNH DOT 21512 (Gaua, Vanuatu)
Effective population size
10⁹
10⁷
10⁵
10⁴
10⁵
10⁶
10⁷
Years Ago

*Turdus poliocephalus xanthopus* AMNH 575554 (Grande Terre, New Caledonia)

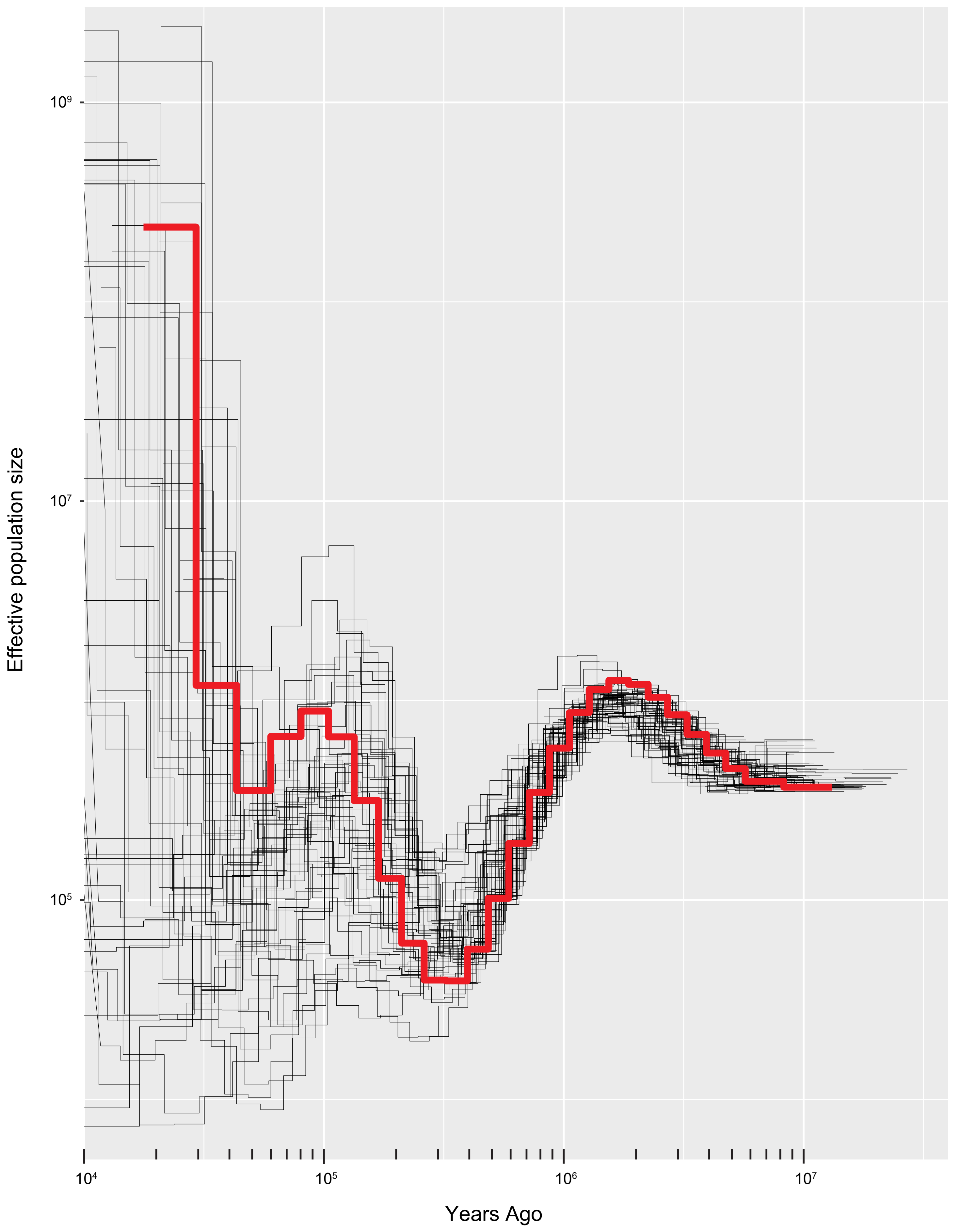
